# Integrin-dependent trafficking of CD98hc reduces endocytic noise to maintain metabolic homeostasis

**DOI:** 10.64898/2026.07.24.740607

**Authors:** F. Sila Rizalar, Hans Peter Grimm, Angelique Augustin, Jose L Garcia-Cordero, Roberto Villaseñor

## Abstract

A stable supply of amino acids is essential for cellular homeostasis. CD98hc is an indispensable component of the LAT1 amino acid transporter. Here, we found an unexpected role for β1-integrin in amino acid homeostasis via the regulation of CD98hc trafficking. We identified Fast Endophilin-Mediated Endocytosis (FEME) as the primary internalization route for CD98hc in both osteosarcoma and brain endothelial cells. Endocytosis was followed by β1-integrin-dependent recycling, which diverted CD98hc from early endosomes. By combining mathematical modelling with high-content imaging, we demonstrate that this trafficking pathway exhibited emergent properties that filtered high-frequency endocytic events to stabilize CD98hc at the plasma membrane. Mutating CD98hc to disrupt its interaction with β1-integrin led to redistribution of the transporter into early endosomes, reduced intracellular amino acid levels and impaired growth under metabolic stress. We propose that regulation of transporter trafficking dynamics is a general mechanism to buffer endocytic noise and maintain metabolic homeostasis.

## Introduction

Cellular homeostasis requires mechanisms to maintain a stable supply of amino acids during periods of high metabolic demand. CD98hc is the chaperone for the amino acid transporter LAT1^1,2^ which is essential for the cellular supply of branched-chain amino acids (BCAA)^3^ and is frequently upregulated in cancer cells to support continuous proliferation^4,5^. While other nutrient receptors (e.g. transferrin receptor (TfR) for iron^6^) supply their cargoes via clathrin-mediated endocytosis (CME) ^7^, BCAA transport requires a stable pool of LAT1 at the plasma membrane. However, CD98hc is internalized by clathrin-independent endocytosis (CIE) and subsequently recycled in an Arf6-dependent manner without transiting through early endosomes^8–10^. The specific CIE mechanisms^11^ responsible for CD98hc internalization are unknown. How the cell coordinates high-capacity^8^ CIE endocytosis with recycling to ensure prolonged LAT1 residence at the plasma membrane and thus a stable supply of BCAA remains an outstanding question.

In addition to its crucial role in amino acid transport, CD98hc has recently emerged as an effective target for receptor-mediated transcytosis across the BBB^12–15^. Similar to the clinically validated transferrin receptor (TfR)^16^, engaging CD98hc increases the brain exposure of peripherally administered antibodies by 6-8-fold *in vivo*^13^. Therefore, understanding the mechanisms of CD98hc trafficking is of high translational relevance for brain delivery.

To resolve this, we mapped the molecular mechanisms regulating CD98hc endocytosis in both osteosarcoma and human brain endothelial cells. We identified fast endophilin-mediated endocytosis (FEME) as the primary driver of CD98hc internalization. By analyzing the interactome of CD98hc, we identified that β1-integrin remains associated with CD98hc within endosomes and is required for fast recycling back to the plasma membrane. The dynamics of this trafficking pathway led to unexpected emergent properties that filter transient endocytosis bursts to stabilize CD98hc at the plasma membrane. We investigated the physiological consequences of perturbing this “biological filter” by mutating CD98hc to prevent its interaction with β1-integrin and measuring its impact on trafficking, intracellular localization, amino acid transport and cell growth under metabolic stress.

## Results

To identify the trafficking pathway for CD98hc, we first compared the localization of an anti-CD98hc antibody to that of an anti-TfR antibody, a canonical cargo for CME^7^, after binding the respective receptors at the plasma membrane. In stem-cell-derived brain endothelial cells^17^, CD98hc-positive vesicles did not overlap with TfR-positive vesicles after 2 hours of incubation with the respective antibodies (Figure 1A). Furthermore, CD98hc exhibited minimal colocalization with the early endosome marker EEA1 and late endo-lysosomal marker LAMP1 (Figure 1B, Figure S1A) and was not enriched in intracellular sorting tubules (Figure 1C, Figure S1B, Suppl. Video 1, 2). Endogenously tagged mGreenLantern-CD98hc (mGL-CD98hc knock in, Figure S1C) fusion protein did not colocalize with TfR nor with EEA1 (Figure 1D). Kinetic transport analysis showed that TfR antibodies were equally distributed between endosomes and plasma membrane, reflecting the saturation of the limited receptor pool at the plasma membrane^18^ (Figure 1E). In contrast, the majority of CD98hc antibodies remained at the plasma membrane (Figure 1F). Strikingly, the extent of antibody internalization was comparable for both receptors (Figure 1F). These findings show that after internalization, CD98hc follows a distinct trafficking pathway in brain endothelial cells that diverges from TfR sorting, maintaining a stable surface pool of CD98hc despite high internalization activity.

**Figure 1.**
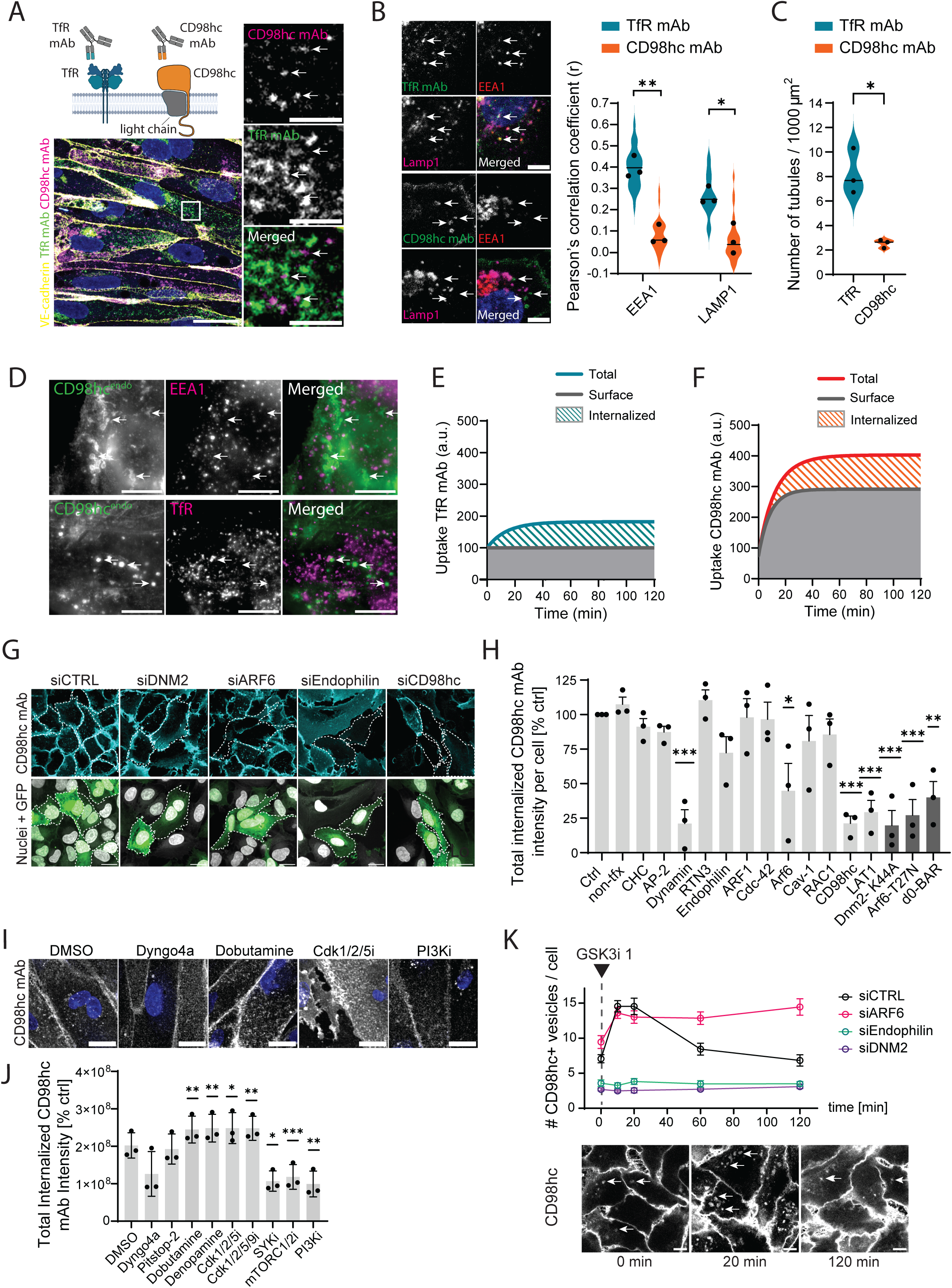
FEME regulates CD98hc endocytosis in brain endothelial cells. **A**, Top: scheme of mAbs used to target TfR and CD98hc. Bottom: Representative confocal images of iCE-BECs (day 14) treated with AF647-conjugated CD98hc mAb and AF488-conjugated TfR mAb, immunostained for VE-Cadherin and DAPI. Scale bar, 25 μm. Zoom-ins, Scale bar, 5 μm. Arrows point to CD98hc mAb-positive vesicles. **B,** Left: Representative confocal images of iCE-BECs (day 14) treated with TfR mAb (top) or CD98hc mAb (bottom), and immunostained for EEA1 and LAMP1. Scale bar, 10 μm. Right: Pearson’s correlation coefficient of the respective mAbs with EEA1 or LAMP1. Violin plots show single-cell distributions (minimum 50 cells per condition), with horizontal lines indicating the overall mean, and data points representing the mean values from three independent biological experiments. Statistical significance was determined by a two-tailed Student’s *t*-test; *, *P* < 0.05; **, *P* < 0.01. **C,** Quantification of endosomal tubule density in hiCE-BECs (day 14) following a 3-hour treatment with TfR or CD98hc mAbs. Violin plots display single-cell distributions normalized per 1,000 μm² (minimum 50 cells per condition), with horizontal lines indicating the overall mean, and data points representing the mean values from three independent biological experiments. Statistical significance was determined by a paired two-tailed Student’s t-test; *, P < 0.05. **D,** Representative confocal images of mGreenLantern-CD98hc KI iCE-BECs (day 14) immunostained for EEA1 (top) or TfR (bottom). Scale bar, 10 μm. **E, F**. Graphs show the simulation of uptake based on experimental data on TfR (**E**) or CD98hc (**F**) mAbs in iCE-BECs (day 14) over a 120-minute time course. Gray areas show the antibody retained on the surface, striped areas show the internalized fraction. **G,** Representative images of intracellular CD98hc mAb localization in resting human U2OS cells upon knockdown of selected endocytosis-related proteins. Nucleofected cells were highlighted with dashed lines. Scale bar, 10 μm. **H,** Targeted CD98hc mAb internalization screen using siRNAs or dominant-negative mutants of selected proteins across diverse endocytic pathways. WT U2OS cells were nucleofected with selected siRNAs together alongside a GFP reporter plasmid (light-gray bars), or with dominant-negative mutation-expressing plasmids fused to a GFP or HA reporter (dark-gray bars). 48 h post-nucleofection cells were treated with CD98hc mAb for 1 h. Bars show the mean ± SEM from a minimum of 100 cells per condition, from three independent biological experiments. Statistical analysis was performed by one-way ANOVA followed by Dunnet’s test. *P < 0.05, **P < 0.01, ***P < 0.001. **I,** Representative confocal images showing the intracellular distribution of CD98hc mAb in iCE-BECs (day 14) upon treatment with DMSO, Dyngo4a, dobutamine, Cdk1/2/5i, SYKi, or GDC-0941 (PI3Ki) for 15 minutes before incubation with CD98hc mAb for 1 hour. Scale bar, 20 μm. **J,** Bar graph shows the normalized total CD98hc mAb vesicle intensity after incubation with DMSO, (vehicle); Dyngo4a, 80 μg/ml; Pitstop-2, 30 μM; dobutamine, 10 μM; denopamine, 10 μM; Roscovitine (Cdk1/2/5i), 1 μM; Dinaciclib (Cdk1/2/5/9i), 1 μM; P505-15 (SYKi), 1 μM; Torin 1 (mTORC1/2i), 10 μM; and GDC-0941 (PI3Ki), 1 μM. Bars show the mean ± SEM from minimum 100 cells per condition, from three independent biological experiments. Statistical analysis was performed by one-way ANOVA followed by Dunnet’s test. *P < 0.05, **P < 0.01, ***P < 0.001. **K,** Top: Graph shows the number of intracellular CD98hc vesicles upon treatment with with 10 μM CHIR-99021 (GSK3i 1) over a 120-minute time course. U2OS cells were nucleofected with either a non-targeting siRNA (siCTRL) or with siRNAs against Arf6 (siARF6), Endophilin A1 + A2 (siEndophilin), or Dynamin-2 (siDNM). Points show the mean ± SEM from minimum 100 cells per condition, from three independent biological experiments. Bottom: Representative confocal images showing the localization of CD98hc in U2OS cells upon GSK3i 1 incubation for different timepoints (t=0, 20 or 120 minutes). Scale bar, 10 μm. In image panels, arrows point to CD98hc-positive vesicles.

To dissect the molecular mechanisms of CD98hc endocytosis, we used U2OS cells as a more amenable cell model to perform a targeted siRNA screen against key machinery across diverse endocytic pathways (Figure 1G, H). As expected, knockdown of either CD98hc or its heterodimeric partner, the LAT1 light chain, blocked internalization of an anti-CD98hc antibody (Figure 1G, H). Consistent with prior reports^9,19^, CD98hc uptake was unaffected by down-regulation of clathrin-mediated endocytosis (Clathrin heavy chain (CHC) or AP-2 knock-down) but was modestly reduced upon downregulation of Arf6 or overexpression of its dominant-negative mutant, T27N. Unexpectedly, depletion (siDNM2) or dominant-negative inhibition (Dnm2-K44A) of Dynamin-2 robustly suppressed CD98hc internalization by approximately 75% (Figure 1G, H). Simultaneous knock-down of Endophilin A1 and A2 reduced CD98hc endocytosis (Figure 1G, H) while expressing an Endophilin mutant lacking membrane-binding activity (d0-BAR)^20^ reduced CD98hc internalization by nearly 75% (Figure 1H). Importantly, these effects were specific to CD98hc as internalization of an anti-TfR antibody relied strictly on CHC and the AP-2 adaptor complex (Figure S1E, F)., The need for both dynamin and endophilin shows that CD98hc endocytosis does not occur via CLIC/GEEC, as previously reported^8^. Instead, our data point towards an endophilin-dependent pathway.

To clarify the role of Arf6 on CD98hc endocytosis, we performed live imaging in mGL-CD98hc knock-in iPSC lines (Figure S1C, D). Knockdown of Dynamin-2 or Endophilin A1 and A2 decreased intracellular CD98hc-positive vesicles, an observation consistent with their role in endocytosis. In contrast, knockdown of Arf6 led to accumulation of enlarged, intracellular CD98hc-positive structures (Figure S1D), indicating that Arf6 does not drive CD98hc endocytosis, but rather acts downstream to regulate sorting as previously reported^9^.

Since CD98hc uptake required endophilin but not CHC nor AP2, we asked whether this pathway corresponded to fast endophilin-mediated endocytosis (FEME)^20^. To this end, we modulated known upstream FEME regulators in U2OS cells^21^. Pharmacological activation via the beta-adrenergic agonist Dobutamine, or inhibition of negative regulators GSK3 (GSK3i 1/2) or Cdk1/2/5 increased by 50% the number of CD98hc-positive intracellular vesicles (Figure S1I, J). Conversely, inhibition of kinases essential for FEME carrier maturation, such as SYK (SYKi) or PI3K (PI3Ki), decreased vesicle formation by half (Figure S1I, J). We validated the role of FEME in CD98hc internalization in stem-cell-derived brain endothelial cells with pharmacological interventions. Pharmacologic inhibition of dynamin (Dyngo4A) reduced internalization of an anti-CD98hc antibody, whereas inhibiting clathrin-mediated endocytosis (Pitstop-2) had no effect (Figure 1I, J). Furthermore, multiple FEME activators (Dobutamine, Denopamine, and Cdk inhibitors) increased antibody uptake, while inhibitors of FEME carrier maturation (Syk, mTORC and PI3K inhibitors) suppressed it by roughly 50% (Figure 1J). We reproduced the same findings in both primary human brain endothelial cells (Figure S1K) and the immortalized hCMEC/D3 cerebral microvascular endothelial cell line (Figure S1L). Furthermoe, expression of the membrane-binding deficient endophilin mutant (ΔH0-BAR) in hCMEC/D3 cells further confirmed the role of FEME in CD98hc internalization (Figure S1G, H), whereas disrupting the CLIC/GEEC pathway with Cdc42-T17N had no impact (Figure S1H). Altogether, these data show that FEME drives CD98hc internalization in both U2OS and brain endothelial cells.

Next, we assessed how Endophilin and Arf6 specifically regulate CD98hc intracellular trafficking. To do so, we knocked down different proteins in U2OS cells and measured the dynamics of CD98hc trafficking after incubation with the GSK3 inhibitor CHIR-99021 (GSK3i 1) to promote the formation of endophilin carrier assembly. In control cells (siCTRL), GSK3i 1 triggered a transient burst of CD98hc internalization that reached a maximum within 10 to 20 minutes, doubling the number of intracellular CD98hc-positive vesicles. This initial uptake was progressively resolved with a half-life of 20 minutes, returning to baseline two hours after the initial stimulus (Figure 1K). Such timescale, together with our previous observation of low CD98hc colocalization with early and late endosomes suggest that the decay of the curve reflects recycling. Vesicle influx was abolished upon siRNA-mediated depletion of Dynamin-2 (siDNM2) or Endophilin-A1/A2 (siEndophilin) (Figure 1K), confirming the role of FEME for CD98hc endocytosis. In contrast, depletion of Arf6 (siARF6) did not alter the initial endocytic burst, but prevented the decline of vesicles, which remained elevated for the duration of the 120-minute time course (Figure 1K). This experiment confirms that Arf6 regulates CD98hc recycling.

FEME drives the rapid, high-volume internalization of its cargoes^20^, yet its activation in U2OS cells led only to a modest 50% increase in intracellular CD98hc vesicles. This discrepancy suggested the presence of an equally fast sorting mechanism that maintains steady-state transporter levels at the plasma membrane. To identify this machinery, we mapped the endogenous CD98hc interactome in mGL-CD98hc knock-in iPSCs using quantitative mass spectrometry (Figure 2A). By employing mild detergent lysis conditions optimized to preserve endosome-associated macromolecular complexes^22^ (Figure S2A), we identified 51 high-confidence interacting proteins (Figure 2A, Table S3). Notably, the CD98hc interactome was heavily enriched for vesicular trafficking machinery, particularly a distinct signature of Rab-family small GTPases, including Rab9, Rab11a/b and Rab14, as well as β1-integrin, a known interactor of CD98hc^23,24^. To spatially map these pathways, we systematically quantified the steady-state CD98hc colocalization across a comprehensive panel of Rab GTPases using high-resolution confocal microscopy (Figure 2B). In agreement with the interactome data, endogenous CD98hc preferentially localized to recycling (Rab14, Rab11a/b) and late endosomes/retrograde carriers (Rab9)^25^ (Figure 2B, Figure S2B). Importantly, CD98hc remained colocalized with β1-integrin within intracellular vesicles (Figure 2C). Since β1-integrin relies on Arf6 and Rab11 for endosomal recycling^26^, we hypothesized that the CD98hc-integrin complex is required for endosomal sorting of CD98hc. To test this, we generated CD98hc-knockout U2OS cells and stably re-expressed either wild-type mGL-CD98hc (CD98hc WT) or an integrin-binding site mutant (Integrin signaling mutant; CD98hc ISM) that disrupts the physical interaction between the two proteins ^27^. The expression levels of CD98hc were comparable between cell lines (Figure S2C). Strikingly, CD98hc ISM cells showed a two-fold increase in intracellular anti-CD98hc-positive vesicles (Figure 2D, E; Figure S2D). Upon incubation with GSK3i, CD98hc WT cells showed a transient increase followed by rapid decay of intracellular CD98hc-positive vesicles. In contrast, the number of CD98hc-positive vesicles did not decay in CD98hc ISM cells (Figure 2F). Together, these data show that CD98hc is sorted to recycling endosomes after FEME internalization and suggest that interaction with β1-integrin is required for recycling back to the plasma membrane.

**Figure 2.**
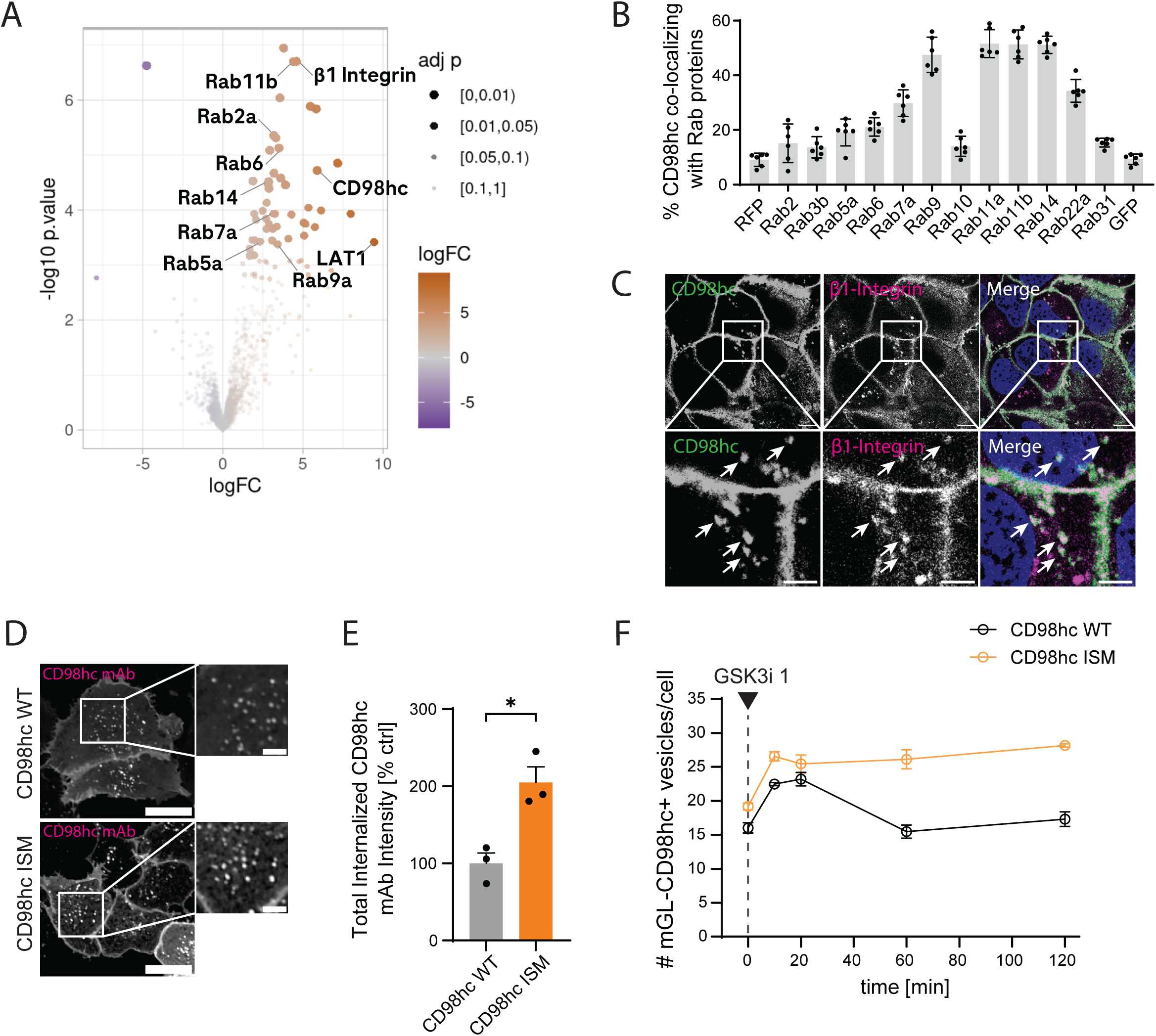
β1-integrin regulates intracellular transport of CD98hc. A, Endogenous CD98hc interactome in human iPSCs. Volcano plot from quantitative label-free quantification (LFQ) affinity purification-mass spectrometry (MS/MS) of endogenously tagged mGL-CD98hc iPSCs (n = 3 biological replicates) relative to parental untagged wild-type cells. The axes display log2 fold change (logFC) against-log(10) P-value calculated via a linear model fit per protein using the limma framework (contrast tests of the given conditions are extracted and corrected for multiple testing). Dot color indicates enrichment magnitude; dot size corresponds to Benjamini-Hochberg adjusted P-value (adj p) tiers. Labeled nodes highlight the bait (CD98hc), its heterodimeric light chain partner (LAT1), β1-Integrin, and an enriched network of vesicle-trafficking Rab GTPases. **B,** Localization screen with Rab small GTPases. Graph shows the fraction of CD98hc positive vesicles co-localizing with selected Rab proteins expressed as mRFP-or eGFP-fusion proteins in U2OS cells. Bars show the mean ± SD from minimum 50 cells per condition, from six independent biological experiments. **C,** Representative images of endogenous CD98hc with endogenous β1-Integrin in resting U2OS cells. Arrows show CD98hc-positive vesicles colocalized with β1-Integrin. Scale bar, 20 μm; Zoom-ins, Scale bar, 5 μm. **D,** Representative confocal images showing CD98hc WT (top) and CD98hc ISM (bottom) U2OS cells after incubation with CD98hc mAb for 30 minutes. Scale bar, 20 μm. Zoom-ins, Scale bar, 5 μm. **E,** CD98hc mAb internalization in CD98hc WT or CD98hc ISM U2OS lines. Bars show the mean ± SEM from a minimum of 100 cells per condition, from three independent biological experiments. Statistical analysis was performed by a paired two-tailed t-test. *P < 0.05. **F,** Graph shows the number of intracellular CD98hc vesicles in mGL-WT-CD98hc or mGL-ISM-CD98hc expressing U2OS cells upon incubation with 10 μM CHIR-99021 (GSK3i 1) over a 120-minute time course. Data show the mean ± SEM from minimum 100 cells per condition, from n=3 independent biological experiments.

To precisely quantify the impact of β1-integrin on CD98hc trafficking, we combined kinetic analysis with mathematical modelling. We first evaluated trafficking during sustained endocytosis by stimulating cells with the β-adrenergic agonist Dobutamine (Figure S3A). Continuous Dobutamine exposure led to a monotonic increase in CD98hc vesicles that reached a steady-state within 20 minutes in cells expressing mGL-CD98hc-WT (Figure 3A). Conversely, vesicles continuously accumulated in cells expressing mGL-CD98hc ISM during the experimental 2-hour window (Figure 3D). To model these dynamics, we adapted a deterministic, three-compartment ordinary differential equation (ODE) system^28^ tracking receptor distribution across the plasma membrane (PM), early endosomes (EE) and recycling endosomes (RE) (Figure 3C, F). Fitting this model to an experimental kinetic dataset across a 1,000-fold agonist concentration range revealed striking differences in transport kinetics between cells expressing either mGL-CD98hc WT or ISM (Figure 3A-D). Even at the highest concentration of dobutamine (100 μM), the levels of CD98hc at the plasma membrane of cells expressing mGL-CD98hc-WT were highly resilient, showing only a minor, transient drop (<5%) before rapidly establishing a compensated steady-state (Figure 3A). In contrast, cells expressing mGL-CD98hc-ISM showed depletion of CD98hc from the plasma membrane by 15%, i.e. 3-fold lower compared to CD98hc-WT cells (Figure 3D). The depletion from the plasma membrane was accompanied by a sustained increase in the CD98hc vesicular intensity (Figure 3E, G), likely reflecting accumulation into endosomes. The model revealed that in CD98hc WT cells internalization is balanced by fast (1 minute timescale, Table S4) recycling from early endosomes, explaining both CD98hc plasma membrane resilience and reduced localization in early endosomes. In cells expressing mGL-CD98hc-ISM, however, both processes are altered: basal internalization (k_basal_) dropped 2.5-fold while recycling from both early (k_rec_) and recycling endosomes (k_rec-2_) were both reduced by nearly 4-fold (Table S4). We need to stress that in ISM cells recycling back to the plasma membrane is not fully impaired but occurs at a substantially slower rate. These altered transport rates predict that CD98hc should accumulate in early endosomes in cells expressing mGL-CD98hc-ISM. High-resolution colocalization analysis with compartment-specific Rab GTPases confirmed this prediction. While in CD98hc WT cells the majority of CD98hc was localized to recycling endosomes (40% and 42% with Rab14 and Rab11b, respectively) (Figure 3I, Figure S3B, S3C), in CD98hc ISM cells intracellular CD98hc shifted away from these compartments and accumulated in Rab5-positive early endosomes (Figure 3H, I). Together, our combined experimental and computational modelling approach shows that the interaction between CD98hc and integrin is required for fast recycling, which stabilizes transporter levels at the plasma membrane to counteract sustained FEME activation.

**Figure 3.**
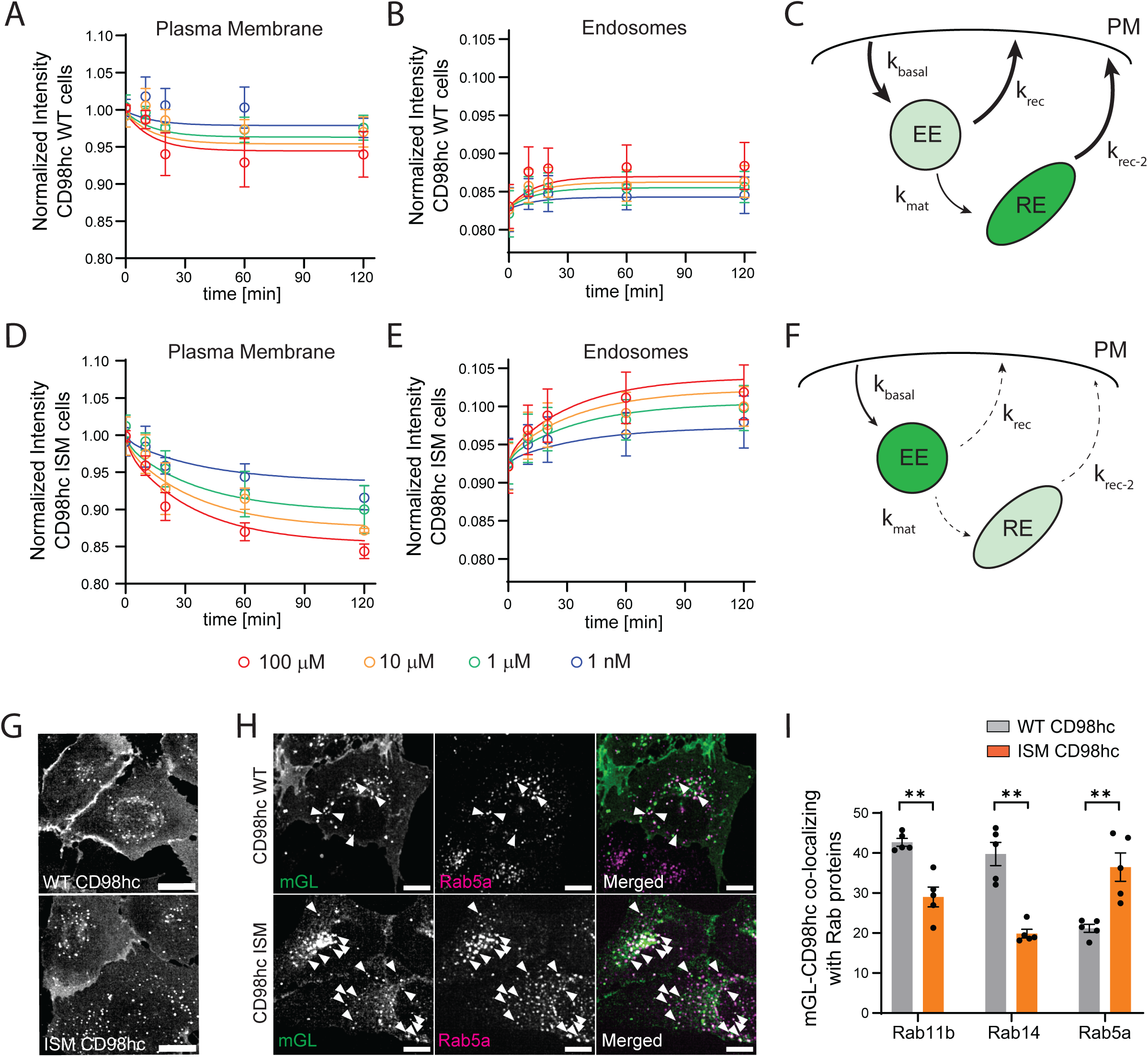
β1-integrin-mediated recycling stabilizes surface CD98hc levels during sustained endocytosis. A, B, Graphs show normalized CD98hc intensity at the plasma membrane (**A**) or in endosomes (**B**) in CD98hc WT cells. **D, E,** Graphs show normalized CD98hc intensity at the plasma membrane (**D**) or in endosomes (**E**) in CD98hc ISM cells. In all graphs, points show the experimental data upon treatment with different concentrations of Dobutamine (10 nM, 1 μM, 10 μM or 100 μM) over 120 minutes with mean ± SEM from minimum 100 cells per condition, from three independent biological experiments. Solid lines show the simulation of the ODE using the best fit parameters to the experimental dataset. **C, F,** Schematics of intracellular pathway with three compartments for CD98hc WT (**C**) and ISM (**F**) cells. A deterministic three-compartment ordinary differential equation system was used to model receptor flux across the plasma membrane (PM), early endosomal (EE), and deep/recycling endosomal (RE) pools governed by transition rate constants (kbasal, krec, kmat and krec-2). **G,** Representative confocal image of CD98hc WT (top) ISM (bottom) cells following Dobutamine (10 μM) treatment for 20 minutes. Scale bars, 20 µm. **H,** Representative confocal images of CD98hc WT or CD98hc ISM cell lines co-stained for the early endosomal marker Rab5a (magenta). Arrowheads highlight mGL-CD98-positive vesicles colocalized with Rab5. Scale bars, 20 µm. **I,** Colocalization of mGL-CD8hc-WT or mGL-CD98hc-ISM with Rab11b, Rab14 and Rab5a. Bars show the percentage of total mGL-CD98hc intensity colocalized with Rab11b, Rab14 or Rab5a in WT and ISM mutant lines. Data are presented as mean ± SEM; **P < 0.01 by paired two-tailed Student’s t-test.

Having established the molecular mechanisms of CD98hc trafficking, we investigated the emergent dynamic properties resulting from coupling a stochastic burst process like FEME^20^ to a very high recycling rate (i.e. with a timescale of one minute). It is well known from the regulation of signaling networks that the timescale between two biochemical reactions can filter high-frequency (i.e. noisy) inputs^29^. Therefore, we asked whether the dynamics of CD98hc trafficking could filter fluctuating endocytosis resulting from transient FEME activation. To this end, we used periodic pulses of dobutamine of different durations (reflecting the frequency of stimulation) and measured the amount of internalized CD98hc (reflecting the response amplitude) at each pulse duration (Figure S4). Experimental data showed that while both WT and ISM cells reduced the number of intracellular vesicles during the washout period, the maximum internalization amplitude was significantly higher in the ISM genotype compared to WT cells (Figure 4A). Analyzing these dynamics via a frequency-response plot (Figure 4B) revealed that the CD98hc WT trafficking circuit acts as a low-pass filter as it reduces the internalization of CD98hc at high stimulation frequency, i.e. shorter FEME bursts. In contrast, CD98hc ISM cells exhibited a highly sensitive response profile with higher amplitude at all frequencies tested. These data show that fast integrin-dependent recycling filters transient FEME bursts, thus limiting internalization of CD98hc and maintaining a stable plasma membrane pool of transporters.

**Figure 4.**
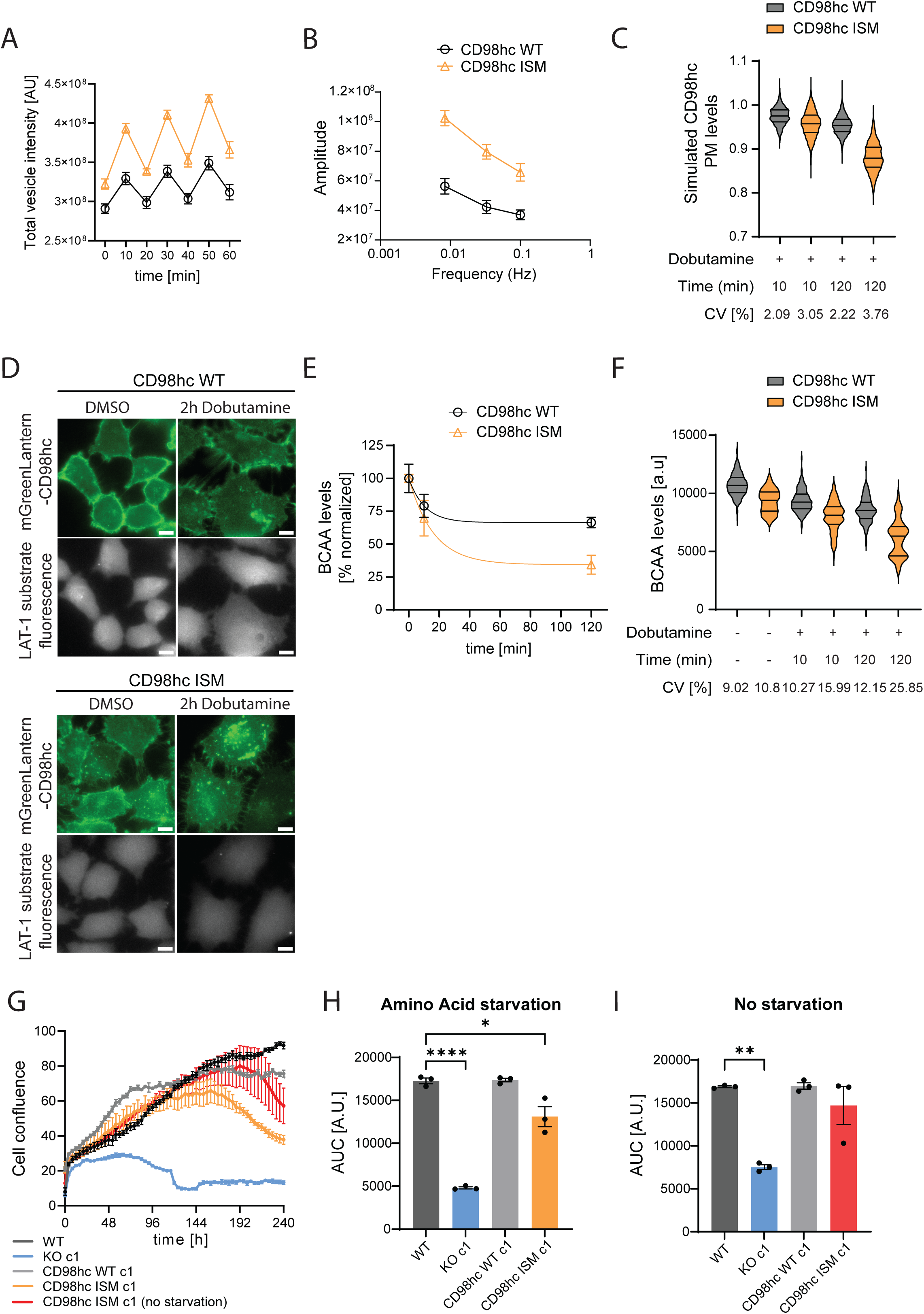
CD98hc trafficking operates as an RC circuit that buffers noise to maintain amino acid homeostasis. A, Experimental data of total CD98hc vesicle intensity after three oscillatory 10-minute pulses of 10 μM dobutamine comparing CD98hc WT and CD98hc ISM cells. Points show mean ± SEM from n=3 independent biological experiments. **B,** Frequency-amplitude plot comparing the amplitude-to-frequency response of CD98hc WT and CD98hc cells for pulse durations of 10, 30 and 120 minutes. Points show mean ± SEM from n=3 independent biological experiments. **C,** Simulation of CD98hc levels at the plasma membrane using the Ornstein-Uhlenbeck stochastic differentiation equation with the best fit parameters from Table S4 for CD98hc WT and ISM cells after incubation with 10 μM dobutamine for 10 or 120 minutes. Violin plots show single-cell distributions from 1000 independent simulated cells, with horizontal lines indicating the median and quartiles. **D,** Representative images of CD98hc WT or ISM lines treated with DMSO or 10 μM dobutamine for 2 h showing mGL-CD98hc expression (top) and the fluorescence reporter measuring LAT1 substrate (bottom). Scale bars, 20 µm. **E,** Branch chain amino acid (BCAA) levels over time for CD98hc WT or ISM cells upon dobutamine treatment for 10 min and 2 h. Data show the mean ± SEM from minimum 50 cells per condition, from three independent biological experiments. **F,** Experimental data showing the BCAA levels of both CD98hc WT and ISM lines change upon incubation with 10 μM dobutamine for different time points. Violin plots show single-cell distributions from three independent biological experiments with minimum 50 cells per condition, with horizontal lines indicating the median and quartiles. **G,** Cell growth measured as increase in confluency of WT, CD98hc KO, and CD98hc WT or ISM U2OS lines in media containing low amino acid levels (10%) for 10 days. For comparison, the red line shows the growth trajectory of CD98hc ISM cells in non-starved amino acid media. Points show mean ± SEM from three independent experiments with minimum 100 cells from each experiment. **H, I,** Comparison of cell growth measured as area under the curve (AUC) of trajectories shown in **G** for cells grown under amino acid starvation (**H**) or full media (**I**). Bars show mean ± SEM from n=3 independent experiments. Statistical analysis was performed by one-way Anova followed by Dunnett’s test. *P < 0.05, **P < 0.01.

We next asked whether the filtering function of CD98hc trafficking dynamics could buffer the intrinsic cellular noise^30^ arising from stochastic endocytosis events and thus reduce phenotypic heterogeneity. To address this, we used stochastic modeling to simulate the levels of CD98hc at the plasma membrane across a large virtual cell population (see Methods for details). Our model predicted that the lower recycling rate in the ISM genotype increases cell-to-cell heterogeneity of CD98hc levels at the plasma membrane compared to WT cells. Specifically, the coefficient of variation (CV) of the ISM single-cell distribution is predicted to increase in a time-dependent manner up to a 70% higher value after two hours of FEME activation compared to WT cells (Figure 4C). This simulation demonstrates that fast recycling is required to reduce the phenotypic heterogeneity of CD98hc levels at the plasma membrane.

Given the essential role of CD98hc in amino acid transport^1^, the consequences of higher heterogeneity in CD98hc levels at the plasma membrane should be reflected in altered intracellular BCAA levels. To test this prediction, we developed a live-cell imaging assay to measure real-time BCAA levels in single cells following sustained FEME internalization in both WT and ISM CD98hc lines. At steady-state, both genotypes showed a similar degree of CD98hc localization at the plasma membrane and similar levels of intracellular BCAAs (Figure 4D). The average BCAA levels across the cell population showed that sustained FEME activation reduced intracellular amino acids by 25% in CD98hc WT cells, compared to a 70% depletion in CD98hc ISM cells (Figure 4E). Prior to FEME activation, the CV of single-cell measurements were comparable between both cell lines (Figure 4F). This shows that overexpression of the ISM mutant is not driving higher phenotypic heterogeneity. Strikingly, the heterogeneity of BCAA levels increased in a time-dependent manner, reaching a 2-fold increase in the CV of single-cell measurements in ISM compared to WT CD98hc cells (Figure 4F). The higher fold-change between the measured CVs compared to the simulation is expected, as the model excluded any extrinsic source of variability (e.g. differences in CD98hc expression across cells). This result confirms the model’s prediction and shows that the fast recycling of CD98hc filters stochastic endocytic events to maintain stable and homogeneous intracellular BCAA levels.

To determine the long-term physiological consequence of altered CD98hc trafficking dynamics, we tracked U2OS cell growth under amino acid depletion and sustained FEME activation. Over a 10-day period, CD98hc-KO cells exhibited severe growth impairment, a defect that was rescued by the re-expression of CD98hc WT (Figure 4G, H). Strikingly, re-expression of CD98hc ISM led to an initial proliferation phase but showed reduced growth after 96 hours of culture. This phenotype in CD98hc ISM cells was partially rescued in amino acid-supplemented media (Figure 4J), where cells continued to grow normally for more than 192 hours in culture. This strongly suggests that reduced growth is driven by amino acid stress rather than impaired cell adhesion. We reproduced these data in an additional independent clone for each genotype (Figure S5). Together, these experiments demonstrate that fast β1-integrin-mediated recycling of CD98hc is essential to sustain long-term proliferation under metabolic stress.

## Discussion

Prior foundational work established that CD98hc is internalized via CIE and is subsequently sorted in an Arf6-dependent manner^9^. Cell fractionation in fibroblasts detected CD98hc in clathrin-independent carriers after pharmacological inhibition of dynamin^8^. This led to the assumption that CD98hc internalization occurs via the CLIC/GEEC pathway. Our data challenge this assumption and identify an alternative uptake mechanism for CD98hc. Through orthogonal methods (siRNA, expression of dominant negative mutants and pharmacological perturbations) we show that FEME is the primary internalization route for CD98hc in both the U2OS osteosarcoma cell line and human brain endothelial cells. While we don’t exclude that CD98hc could still be internalized via CLIC/GEEC, it is also plausible that in earlier studies the protein was mislocalized to a different compartment upon dynamin inhibition.

The interaction between β1-integrin and CD98hc is currently described as a mechanism to regulate integrin signaling and cell adhesion^31,32^. Here we expand on the physiological relevance of this interaction by showing that β1-integrin regulates recycling of CD98hc to the plasma membrane. The recycling of a CD98hc mutant that cannot interact with β1-integrin was 4 times slower compared to the wild-type protein. Consequently, CD98hc was sequestered within Rab5-positive early endosomes, ultimately leading to reduced BCAA transport. These findings provide a novel link between integrins and cell metabolism through the regulation of trafficking dynamics and suggest that mechanotransduction could modulate amino acid metabolism.

Our findings on CD98hc endocytosis shed new light on the molecular mechanisms for therapeutic delivery across the BBB. Compared to TfR, CD98hc delivers a lower concentration of payload after peripheral administration but has prolonged residence time in the brain^13,33^. The established paradigm assumed that this pharmacokinetic profile resulted from slow, rate-limiting target internalization of CD98hc. Here, we challenge this notion by demonstrating that CD98hc undergoes rapid internalization via FEME at rates comparable to clathrin-mediated endocytosis of TfR. The prolonged brain exposure observed *in vivo* is likely governed by distinct intracellular routing and fast recycling rather than slow entry. While we established the molecular regulation of CD98hc internalization, a key open question is how its intracellular sorting is regulated to achieve transcytosis. Recent *in vivo* mapping indicates that β1-integrin is predominantly enriched at the abluminal membrane of brain endothelial cells^34^. Therefore, we do not anticipate that the β1-integrin mechanism we described in non-polarized cells will affect transcytosis across the BBB.

Beyond transport, endocytosis is increasingly recognized as a signal processing system^35–37^. Endosomal recycling is a well-established mechanism to sustain receptor signaling and resensitize cells for repeated inputs^38^. Beyond these canonical functions, our data shows that recycling can also act as a low-pass filter. High-frequency, transient endocytic events (i.e. extrinsic or intrinsic noise) are suppressed while low-frequency, sustained endocytosis is allowed. This emergent property stabilizes CD98hc at the plasma membrane while still allowing for the redistribution of the transporter in response to prolonged stimuli. The structure of this trafficking pathway is analogous to a parallel resistor-capacitor (RC) circuit (Figure S6) ^39^. The recycling loop acts as a resistor that rapidly redirects charge (CD98hc) back to the capacitor (plasma membrane), thus maintaining a stable voltage (plasma membrane CD98hc levels) despite being drained by transient current sinks (FEME bursts). Importantly, similarly to an electronic RC-circuit, the timescale of recycling determines the strength of the filter. Unlike constitutive recycling loops (e.g. TfR) that operate on slower timescales^18^, the fast recycling rate of CD98hc matches the burst dynamics of FEME and prevents accumulation within endosomes. Since multiple transmembrane receptors follow the same transport architecture (e.g. GLUT4^40^, EGFR^41^), we propose that coupling stimulus-driven internalization to fast recycling represents a general mechanism to filter extrinsic and intrinsic noise. Further work is required to fully investigate the physiological consequences of increased noise on amino acid metabolism.

## Materials and Methods

### Cell culture and Maintenance

Human osteosarcoma (U2OS), HeLa, and HEK293 cell lines (originally obtained from ATCC) were maintained in Dulbecco’s Modified Eagle Medium (DMEM) supplemented with 4.5 g/L glucose, 292 μg/mL L-glutamine (11965092, Life Technologies), 10% (v/v) fetal bovine serum (FBS; F4135, Sigma-Aldrich), 100 U/mL penicillin, and 100 μg/mL streptomycin (10378016, Gibco).

Immortalized human cerebral microvascular endothelial cells (hCMEC/D3) were obtained from Merck Millipore (SCC066) and cultured in Endothelial Growth Medium-2 (EGM-2) BulletKit (CC-3162, Lonza). To ensure experimental reproducibility, cells from three distinct passages were utilized.

Primary human brain microvascular endothelial cells (HBMVECs) were purchased from AngioProteomie (cAP-0002) from three independent biological batches. HBMVECs were cultured in Endothelial Growth Medium (cAP-02, AngioProteomie) on tissue culture vessels pre-treated with Quick Coating Solution (cAP-01, AngioProteomie). Each cell batch was cultured for two subsequent passages before seeding for internalization experiments.

Human induced pluripotent stem cell (hiPSC) experiments were performed using the BIONi010-C line, obtained from the European Bank for Induced Pluripotent Stem Cells (EBiSC). This line was originally derived from dermal fibroblasts of an 18-year-old male via electroporation of episomal plasmids encoding OCT4, SOX2, KLF4, L-MYC, and LIN28. hiPSCs were maintained under feeder-free conditions on culture plates coated with Geltrex LVT (A1413301, Thermo Fisher Scientific) and fed daily with mTeSR Plus medium (100-0276, STEMCELL Technologies). Routine passaging and single-cell dissociation were performed using Gentle Cell Dissociation Reagent (100-0485, STEMCELL Technologies) or StemPro Accutase Cell Dissociation Reagent (A1110501, Thermo Fisher Scientific).

All cell lines and primary cultures were systematically screened and confirmed negative for mycoplasma contamination. All cultures were maintained at 37°C in a humidified atmosphere containing 5% CO₂.

### Generation of Stable CD98hc Expression Lines

To generate stable U2OS lines expressing mGreenLantern (mGL)-tagged variants, cDNA sequences encoding mGL, a flexible GSGS linker, and human SLC3A2 (encoding CD98hc) were synthesized de novo and cloned in-frame downstream of the cytomegalovirus (CMV) promoter within the expression vector pcDNA3.1(+) (GenScript). Constructs included both wild-type CD98hc (mGL-CD98hc-WT) and the integrin-binding interface mutant (mGL-CD98hc-ISM).

Parental wild-type or CRISPR-mediated CD98hc-knockout (KO) U2OS cells were transfected using jetPRIME transfection reagent (101000046, Polyplus/Sartorius) at 60–80% confluency according to the manufacturer’s protocol. Stable transformants were selected under continuous pressure of 1 mg/mL Geneticin (G418 Sulfate; 10131027, Thermo Fisher Scientific). Following selection, uniform, low-to-moderate expression pools of mGL-positive cells were isolated via fluorescence-activated cell sorting (FACS).

Cell lines generated from CD89hc KO lines and stably express mGL-WT-or mGL-ISM-CD98hc constructs are called CD98hc WT and CD98hc ISM in this study.

### hiPSC differentiation into brain endothelial cells

Differentiation of hiPSCs into brain endothelial cells was previously described^17^. Briefly, 2 million hiPSCs were seeded in a 10 cm dish coated with Geltrex in 10 mL of mTeSR Plus Media supplemented with ROCK inhibitor, Y-27632 10 µM (SCM075, EMD Millipore). 24 h later, media was replaced with mesodermal induction media composed of DMEM/F12 (31331-028, Gibco) and Neurobasal medium (21103-049, LifeTechnologies) 1:1, 2-Mercaptoethanol (31350-10, ThermoFisher), B27 (17504044, Gibco), N2 (17502048, Gibco) supplemented with fresh CHIR-99021 8 μM (13122, Cayman) and BMP4 25ng/mL (120-05ET, Peprotech). On day 4 and 5, media was replaced with Endothelial differentiation medium consisting of StemPro-34 SFM Media (10639011, Life Technologies) with StemPro-34 supplement, Glutamax (35050061, Gibco), Penicillin-Streptomycin (15070063, Gibco), and freshly supplemented with VEGF165 50 ng/mL (293-VE-010, R&D) and Forskolin 2 µM (ab120058, abcam). On day 6, cells were replated in 10 cm dishes (1.2 million per dish) coated with Vitronectin 2.5 µg/ml (SRP3186, Sigma) and Fibronectin 7.5 µg/ml (F0895, Sigma) in BBB Identity Acquisition media consisting of Vasculife VEGF Endothelial Medium (LL-0003, Lifeline Cell technology) supplemented with iCell Endothelial cells medium supplement (M1019, Fuji cell dynamics) instead of the FBS included in the LL-0003 kit and 10 mL of L-Glutamine LifeFactor instead of the 25 mL. included in the kit. The media was freshly supplemented with CHIR-99021 4 µM (13122, Cayman), SB-431542 5 μM (72234, Stemcell), cAMP 50 nM (ab120424, abcam). On day 11, PECAM1-positive cells were MACS sorted using CD31 MicroBead Kit from MACS Miltenyi Biotec (130-091-935, Miltenyi Biotec), according to the manufacturer’s protocol. Sorted cells were either frozen in liquid nitrogen or replated in the same conditions with BBB Identity Maintenance media (same composition of BBB Identity Acquisition media).

Experiments were carried out on day 14 (three days after MACS sorting) and with iCE-BECs differentiated from the hiPS_ BIONi010-C13 parental line, if not indicated differently.

### Generation of genome-edited isogenic iPSC lines using CRISPR/Cas9

To generate an mGreenLantern-CD98hc KI hiPSC line via insertion of an mGreenLantern tag after the start codon of SLC3A2, the following synthetic gRNA targeting sequence was used: UGAGCCAGGACACCGAGGUG (Synthego). A donor vector was generated containing the original genomic sequence of human SLC3A2 approximately 1000 bp up-and downstream of the start-codon including the sequence of mGreenLantern and s GSGG linker after the start-codon of SLC3A2. The homology regions and the mGreenLantern plus the GSGG linker sequence were joined together via gene synthesis. Gene synthesis and plasmid purification were performed by Genscript. WT hiPSCs were nucleofected with 1 mg Donor vector and synthetic gRNA – Cas9 protein complex containing 300 pmol of sgRNA and 40 pmol of recombinant SpCas9 Nuclease (R20SPCAS9-SM, Synthego) using the P3 Primary Cell 4D-Nucleofector® X Kit S (F-16768, Lonza) according to manufacturer’s instructions. Post-nucleofection, hiPSCs were cultured in the presence of Y-27632 10 µM (SCM075, EMD Millipore) and Alt-R™ HDR Enhancer V2 0.69mM (10007910, IDT) overnight. Next day the medium was changed to mTeSR Plus Media (100-0276, Stemcell) and cells were maintained until confluency was reached.

Manually picked single knock-in (KI) iPS clones were screened by Western Blotting and by PCR amplification of the genomic DNA up-and downstream of homology arms by using these primers: forward, 5’-ctggtcgagctggacggcgacg-3’; reverse, 5’-agacgccgcgttcatcggctgctt-3’.

Clones with integration of the homology cassette in both alleles (absence of WT band) were sequenced to ensure integrity of both homology arms and correct introduction of the mGreenLantern tag. Correct clones were expanded and cryopreserved in liquid nitrogen.

### Generation of genome-edited isogenic U2OS lines using CRISPR/Cas9

To generate an mGreenLantern-CD98hc KI U2OS line via insertion of an mGreenLantern tag after the start codon of SLC3A2, the same synthetic gRNA targeting sequence previously used for human iPSCs was used: UGAGCCAGGACACCGAGGUG (Synthego). WT U2-O-S cells were nucleofected with 1 mg Donor vector and synthetic gRNA – Cas9 protein complex containing 300 pmol of sgRNA and 40 pmol of recombinant SpCas9 Nuclease (R20SPCAS9-SM, Synthego) using the SG Cell Line 4D-Nucleofector® X Kit S (V4XC-3032, Lonza) according to manufacturer’s instructions. 24h post-nucleofection medium was changed to fresh DMEM (11965092, Life Technologies) and cells were maintained until confluency was reached. 4-days after nucleofection GFP-positive cells were single-sorted into 96-well plates using fluorescence-assisted cell sorting. Clones were screened by Western Blotting and by PCR amplification of the genomic DNA up-and downstream of homology arms by using these primers: forward, 5’-ctggtcgagctggacggcgacg-3’; reverse, 5’-agacgccgcgttcatcggctgctt-3’.

To generate CD98hc KO U2OS lines, three synthetic gRNA targeting sequences were used: SLC3A2+62885185: CCUCUGCAGGCUCUUGAUUG; SLC3A2-62885206: CAGGAUCUGCUGAAGGUCGG; SLC3A2+62885340: UGAAUGCCACUGGCAAUCGC (Synthego). WT U2-O-S cells were nucleofected with 1 mg Donor synthetic gRNA – Cas9 protein complex containing 300 pmol of sgRNA (a mix of all three sgRNAs) and 40 pmol of recombinant SpCas9 Nuclease (R20SPCAS9-SM, Synthego), nucleofected and screened as explained above.

### Immunofluorescence detection of mAb localization *in vitro*

hiCE-BECs at day 11 were grown in glass-bottom 96-well plates (PBK96G-1.5-5-F, MatTek) for 72 hours before the experiment in full BBB Acquisition medium. Cells were incubated with TfR mAb or CD98hc mAb at final concentration of 100 nM, or 25 μg/mL of Transferrin-AlexaFluor647 (Thermo Scientific, Cat no. T23366) as control, for different amounts of time at 37 °C and 5% CO2.

For measurements of clearance, cells were first incubated with mAbs or Transferrin for 30 minutes at 37 °C and 5% CO2 and were afterwards incubated with starvation medium for different time points at 37 °C and 5% CO2. For measurements of internalization, cells were incubated with mAbs or Transferrin for different time periods (up to 120 minutes) before washing with 1X PBS and fixation with 2% PFA for 15 minutes at room temperature. For detection of mAb clusters at the plasma membrane, cells were instead incubated on ice for 30 minutes and medium. Afterwards, cells were washed once with PBS and fixed with 4% PFA/PBS for 30 minutes at room temperature.

For immunofluorescence, cells were first permeabilized with 0.1% Saponin (84510, BioChemika) for 10 minutes in the presence of 4% cold water fish gelatin (G7041, Sigma) for blocking. Afterwards, cells were incubated with the corresponding antibodies in PBS with 4% cold water fish gelatin and 0.01% saponin (SG-PBS) for 1 hour, washed three times for 5 minutes with 1X PBS and incubated for 30 minutes with appropriate secondary antibodies in SG-PBS. To detect Brain Shuttle constructs exclusively at the plasma membrane, all the incubations were carried out in the absence of saponin to prevent membrane permeabilization. Finally, cells were washed three times for 5 minutes with PBS and nuclei were stained with 1 µg/ml DAPI and/or 1 µg/ml HCS CellMask Blue Stain (H32720, ThermoFisher).

### Modeling CD98hc internalization kinetics in hiCE-BECs

The internalization kinetics was modeled using a deterministic, two-compartment ordinary differential equation (ODE) system. The model captures the distribution of antibodies associated with the plasma membrane *A_PM_* or with the endosomal compartments *A_TE_*.

The amount of antibodies associated with the plasma membrane *A_PM_* is modeled by a rapid initial binding of amount *A_PM_*_,0_, followed by a slower accumulation up to to the maximal capacity *A_PM_*_,*max*_ characterized by the dissociation rate *k_off_*.

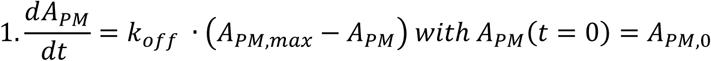

The initially bound amount is furthermore expressed as a fraction of the maximal capacity *A_PM_*_,0_ = *f_0_ · A_PM,max_.*

Subsequent internalization of antibodies associated with the plasma membrane leads to accumulation of endosomal antibodies *A_TE_*. This process is characterized by the relative strength of endocytosis *z_end_* and by the endosomal recycling/elimination rate *k_out_*:

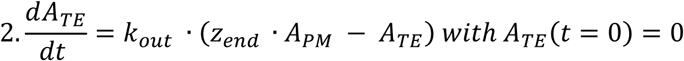

The signal from non-permeabilized cells is identified with *A_PM_* and the signal from permeabilized cells is constituted by the sum *A_PM_* + *A_TE_*.

Parameters were estimated using Monolix, the corrected Bayes Information Criterion was used for model optimization. For the final model, *f*_0_, *A_PM_*_,*max*_, and *z_end_* were allowed to vary between antibodies, *A_PM_*_,*max*_ between experiments.

### Sorting Tubule Formation Assay

iPSC-derived brain endothelial cells were seeded in a 96-well glass bottom imaging plate (15.000 cells per well) (PBK96G-1.5-F, MatTek USA) in BBB Identity Maintenance media 48 h before the experiment. After washing with PBS, cells were incubated with either BrS-1026 (TfR mAb) or CD98hc mAb at 100 nM final concentration for at least three hours at 37°C 5% CO2. After washing with BBB Identity maintenance media, cells were imaged with a DMI-8 fluorescence microscope (Leica Microsystems) equipped with a stage incubator controlling at 37 °C and 5% CO2. Full frame 1024 x 1024 images were acquired with HCX PL APO 100X/1.4 NA oil objective with a resolution of 0.288 µm, two z-stacks with 1 µm step size. Single color images were acquired for a final rate of 4 frames per second for 30 seconds each. The number of tubules per cell occurring within 30 second acquisition was quantified manually in the maximum intensity projection of the z-sections for each movie.

## Targeted siRNA Screen for Endocytosis

U2OS cells were co-transfected with target-specific siRNAs or a non-targeting control siRNA (sequences listed in **Table S1**) alongside a GFP reporter plasmid using the SG Cell 4D-Nucleofector X Kit S (VX4G-3032, Lonza) according to the manufacturer’s protocols. Transfected cells were seeded into black 96-well imaging plates (12,000 cells/well; 6055302, Revvity) and cultured for 48 hours. To quantify cargo internalization, cells were subjected to an antibody-feeding assay by incubation with 100 nM of either anti-TfR mAb (BrS-1026) or anti-CD98hc mAb for 1 hour at 37°C. Cells were then rinsed once with PBS to remove unbound surface antibodies, immediately fixed with 4% (w/v) paraformaldehyde (PFA) in PBS, and processed for quantitative immunofluorescence microscopy.

### Pharmacological Treatments and Small-Molecule Modulations

iPSC-derived brain endothelial cells (hiCE-BECs; treated at day 14 post-differentiation), hCMEC/D3 cells, and primary HBMVECs were cultured on glass-bottom 96-well plates (PBK96G-1.5-5-F, MatTek). Osteosarcoma U2OS lines were seeded into black-walled 96-well imaging plates (6055302, Revvity) at a baseline density of 12,000 cells per well.

Unless specified otherwise, cells were pre-incubated with targeted small-molecule inhibitors or pathway agonists in their respective complete growth media for 20 min at 37°C under 5% CO₂. Chemical modulators were maintained at their final working concentrations throughout the experimental time courses. A comprehensive directory of all chemical reagents, functional definitions, assigned study nomenclature and working concentrations is compiled in **Table S2**.

For antibody internalization experiments, BrS-1026 (TfR mAb) or CD98hc mAb’s (unconjugated or Alexa fluor-488 or-647 conjugated) were added at 100 nM final concentration and cells were incubated for 30 more minutes before live-or fixed imaging.

For pulse-chase experiments, cells were stimulated with Dobutamine for FEME upregulation at final concentrations ranging from 10 nM to 0.1 mM and cells were fixed with 4% (w/v) paraformaldehyde (PFA) in PBS immediately after the treatment or washes.

### Confocal or Epi-fluorescence Image Acquisition and Quantitative Internalization Analysis

For hiCE-BEC antibody internalization assays, z-series images consisting of 4 optical sections spaced at 0.75 μm intervals were acquired on a Leica SP8 confocal microscope utilizing an oil-immersion objective (HC PL APO CS2 63×/ NA 1.40 oil, 100 nm pixel size at 1024 x 1024 frame size). Image processing and quantification were performed in Fiji/ImageJ using a custom automated macro. Total cellular regions were segmented using the CellMask Blue channel to define individual regions of interest, and total vesicular fluorescence intensity was measured per cell. To strictly isolate and quantify the internalized cargo pool, total fluorescence intensity on the cell surface measured from parallel, non-permeabilized control conditions was subtracted from the intensity of permeabilized samples.

The Perkin Elmer’s Harmony high-content analysis software 5.1 (HH17000001) was used to set up the plate dimensions enabling the fast and efficient imaging with the Opera Phenix High Content Imaging System (PerkinElmer). For U2OS kinetic pulse-chase and steady-state internalization assays stimulated with GSK3i or Dobutamine, the 40× long WD confocal objective was used. Per well, 20 images were acquired, 4 planes per image with a section thickness of 0.8 µm per plane. Appropriate channels were selected (DAPI, Alexa-488, Alexa-647) and exposure time as well as focus height was set accordingly and kept the same between wells. Perkin Elmer’s Harmony high-content analysis software 5.1 was used to analyze the images. In brief, basic flat field correction and maximum projection was applied. Cell area was determined by a binary threshold mask using the CellMask Blue signal, followed by spot-detection profiling of the antibody channel to calculate sum vesicular intensity per cell. Mirroring the confocal processing pipeline, internalized antibody levels were quantified by subtracting the plasma membrane signal derived from parallel, non-permeabilized control wells.

Live cell imaging samples were acquired with a DMI-8 fluorescence microscope (Leica Microsystems) equipped with a stage incubator controlling at 37 °C and 5% CO2. Full frame 1024 x 1024 images were acquired with HCX PL APO 100X/1.4 NA oil objective with a resolution of 0.288 µm. Single or dual color images were acquired for a final rate of 2 frames per second.

### Colocalization analysis

To analyze cargo colocalization with endogenous organelle markers (EEA1, LAMP1) in brain endothelial cells, samples were imaged on a Leica SP8 confocal microscope using an oil-immersion objective (HC PL APO CS2 63x/ NA 1.40 oil, 100 nm pixel size at 1024 x 1024 frame size). Intensity-based colocalization was quantified using the Coloc2 plugin within Fiji/ImageJ to calculate Pearson’s correlation coefficients.

For the overexpressed Rab GTPase screening panel, cells were imaged on an Opera Phenix High-Content Screening System using the Opera Phenix High Content Imaging System (PerkinElmer) as described above. Object-based spatial overlap was analyzed via Harmony 5.1 software by segmenting individual CD98hc-positive vesicular carriers and calculating the percentage of vesicles scoring positive for the corresponding Rab protein signal.

### Immunoprecipitation using magnetic nanobeads

HEK293 cells were seeded on 10 cm tissue culture plates and transfected with mGL-CD98hc and mRFP-LAT1 expressing plasmids; or mGL-CD98hc iPSCs with endogenous CD98hc expression were used without overexpression. In the case of transfection, cells were incubated for 24 hours before processing. Plates were washed twice with ice-cold 1X PBS. 1 mL of IP lysis buffer (0.01% saponin, 20 mM HEPES pH7,4, 130 mM NaCl, 10 mM NaF, Protease Inhibitor (11836170001, Roche)) was added per plate, lysates were scraped into a tube and were centrifuged for 10 minutes at 10 000 x g at 4°C. Soluble fraction of the lysate was kept and pellet was discarded. 75 μL of lysate were transferred into a clean tube and mixed with 25 uL of 4x SDS-buffer to be used as the input on WB. Rest of the supernatant was mixed with 15 uL of previously washed GFP-or RFP-trap Magnetic Agarose beads (ChromoTek) and the mix was incubated for 3 h at 4°C on a rotator. Beads were washed three times with the IP buffer, and bound proteins were eluted in SDS–PAGE sample buffer, resolved by SDS–PAGE and analysed via immunoblot.

### Protein preparation and immunoblot

Confluent cells were briefly washed with PBS on ice and harvested in 100 μL 1X RIPA buffer (89900, LifeTech) for one well of a 6-well dish. After the lysates were collected with a cell scraper, they were centrifuged at 14,000 x g for 20 minutes at 4°C to remove cell debris, and the supernatant were retained for protein quantification using the Pierce bicinchoninic acid assay method (23225, ThermoFisher). Soluble proteins were diluted in NuPAGE LDS Sample Buffer (4x) (NP0007, Thermo Fisher) containing NuPAGE Sample Reducing Agent (NP0004, Thermo Fisher) as per manufacturer’s instructions, denatured for 5 minutes at 95°C, spun-down and stored at-20°C.

5-20 µg of protein was typically loaded on and resolved using 4-15% Mini-Protean TGX Stain-Free gels (4568085, Bio-Rad) and transferred on 0.2 µm nitrocellulose membranes (1704159, Bio-Rad) using a Trans-Blot Turbo Transfer System, (BioRad). Membranes were subsequently blocked with 5% Bovine Serum Albumin (BSA) (A561, Sigma-Aldrich) in Tris-buffered saline with 0.1% Tween® 20 Detergent (TBS-T) for 1h followed by primary antibody incubation in 5% BSA in TBS-T overnight. After three washes in TBS-T, appropriate HRP-conjugated secondary antibodies in 5% BSA in TBS-T were applied (donkey-anti-rabbit-HRP, A16035, ThermoFisher, 1:5000; donkey-anti-goat-HRP, A16005, ThermoFisher, 1:5000; donkey-anti-mouse-HRP, A32788, ThermoFisher, 1:2000) for 1h at RT. Immunoblots were washed three times in TBS-T for five minutes each, and protein bands were then visualised using SuperSignal West Pico Plus Chemiluminescent Substrate (34580, ThermoFisher). Image acquisition was performed using the ChemiDoc MP (BioRad).

### LC-MS/MS analysis CD98hc interactors

After enrichment of the proteins on magnetic beads, the beads were resuspended in denaturing buffer (1% (w/v) sodium deoxycholate, 10 mM tris(2-carboxyethyl)phosphine, 40 mM chloroacetamide, 100 mM Tris, pH 8.5) and incubated at 95°C for 10 minutes (min). Samples were subjected to trypsin digestion at 37°C for 1.5 hours and transferred to new tubes. The resulting peptides were purified using stage-tip solid-phase extraction and subsequently dried via vacuum centrifugation. Peptides were reconstituted in a solution of 2% Acetonitrile (ACN) and 0.5% Formic acid (FA). A total of 400 ng of peptides was analyzed by liquid chromatography (LC) using a nano capillary system (Vanquish Neo, Thermo Scientific). Chromatographic separation was achieved on a C18 reverse-phase nano-high-performance liquid chromatography (nano-HPLC) column connected to a mass spectrometer (Orbitrap Ascend™ Tribrid, Thermo Scientific) via electrospray ionization. The Data-Independent Acquisition (DIA) method consisted of one full-range MS1 scan from 340 to 1210 m/z at 120,000 resolution, utilizing a custom Automatic Gain Control (AGC) target and a maximum injection time of 20 ms. Subsequently, 28 DIA segments were acquired at 15,000 resolution, with a standard AGC target and a maximum injection time of 22 ms. High-Energy Collisional Dissociation (HCD) fragmentation was set with normalized collision energy optimized for each segment. All spectra were recorded in profile mode. Raw files were processed using Spectronaut 19 software. The experimental settings were based on the BGS Default SNE settings for a library-free DIA search, without imputation or normalization. Default settings included a 1% false discovery rate (FDR) control at both the peptide and protein levels. The mass spectrometric data were specifically analyzed using the Pulsar search engine, as implemented in Spectronaut software, maintaining a 1% FDR on peptide and protein levels. The search engine utilized a human UniProt FASTA database (Homo Sapiens, 2022 07 01). Search parameters allowed for a maximum of two missed cleavages and included variable modifications for N-terminal acetylation and methionine oxidation.

### Plasmids

All expression vectors were synthesized and sequence-verified by GenScript using a pcDNA3.1(+) backbone and human cDNA inserts. Fusion constructs utilized a flexible GSGG linker between the fluorescent tag and the protein boundary.Cargo constructs were N-terminally tagged with mGreenLantern (mGL) and comprised wild-type CD98hc (mGL-WT) and an integrin-binding site mutant harboring W179G and R181A mutations (mGL-ISM)^27^. Empty pcDNA3.1(+), eGFP, and mRFP vectors were used as controls.

Dominant-negative (DN) constructs for endocytic and recycling regulators included C-terminally tagged Arf6-T27N-HA/mRFP, and N-terminally tagged versions of Dynamin-2 (eGFP/mRFP-DNM2-K44A), Cdc42 (HA-CDC42-N17), and Endophilin A1 lacking the amphipathic H0 helix (eGFP/mRFP-Δ0-BAR).

The intracellular mapping panel consisted of the following N-terminally tagged human Rab GTPases: mRFP-Rab2a, mRFP-Rab9a, eGFP-Rab10, mRFP-Rab11a, mRFP-Rab11b, mRFP-Rab14, mRFP-Rab31; and eGFP-Rab3b, eGFP-Rab5a, eGFP-Rab6, eGFP-Rab7a, eGFP-Rab22a.

### Amino Acid Uptake Assay

To evaluate intracellular branched-chain amino acid (BCAA) uptake, an Amino Acid Uptake Assay Kit (UP04; Dojindo Laboratories, Kumamoto, Japan) was utilized according to the manufacturer’s protocol. Briefly, wild-type (WT) or integrin-mutant (ISM) mGreenLantern-CD98hc expressing U2-O-S cells were seeded in a 96-well microplate and cultured overnight at 37°C in a 5% CO2 incubator. Prior to the assay, cells were pre-incubated with either 10 µM dobutamine (for 10 min or 2 h), 100 µM BCH (for 10 min, a specific LAT-1 inhibitor used as a negative control), or DMSO (for 2h). To maintain consistent treatment conditions, all subsequent assay solutions and wash buffers were supplemented with the respective concentrations of dobutamine or BCH.

The culture medium was removed, and cells were washed three times with pre-warmed Hank’s Balanced Salt Solution (HBSS; 37°C). Cells were then incubated in 150 µL of pre-warmed HBSS at 37°C for 5 min. The supernatant was replaced with 150 µL of pre-warmed BPA uptake solution (prepared by mixing equal volumes of BPA Solution and BPA Dilution Buffer followed by a 50-fold dilution in HBSS) or pre-warmed HBSS as a blank control, and incubated at 37°C for 5 min. After washing three times with pre-warmed HBSS, cells were incubated with 150 µL of probe working solution (250-fold dilution in HBSS, centrifuged at 300 × g for 3 min to remove undissolved residue) at 37°C for 5 min. Live-cell imaging was immediately performed using a DMI-8 fluorescence microscope (Leica Microsystems) equipped with a stage incubator controlling at 37 °C and 5% CO2. Full frame 1024 x 1024 images were acquired with HCX PL APO 100X/1.4 NA oil objective with a resolution of 0.288 µm, five z-stacks with 0.5 µm step size, using GFP (for mGreenLantern) and DAPI (for the BPA probe; Ex/Em = 360/460 nm) channels. The working solution was maintained in the wells throughout imaging to prevent signal dissipation. Total intensity in cytoplasm or on the plasma membrane was quantified on Fiji using the maximum intensity projection of the z-sections for each image.

### Cell Proliferation Assay

For proliferation assays under low amino acid conditions, wild-type (WT) U2OS cells, two independent CD98hc knockout (KO) clones, and WT or integrin-mutant (ISM) mGreenLantern-CD98hc expressing lines (generated from the respective KO clones) were utilized. Cells were seeded into 96-well black PhenoPlates (6055302, Revvity) at a density of 6,000 cells per well. The following day, cells were washed twice with phosphate-buffered saline containing calcium and magnesium (PBS+/+) and incubated in an amino acid-free starvation medium (Dulbecco’s MEM (DMEM) w/o Amino Acids) for 2 h. Following starvation, the medium was replaced with a low amino acid treatment medium, consisting of an amino acid-free starvation medium supplemented with 10% complete DMEM (containing 10% dialyzed fetal bovine serum [FBS]) and 2 µM dobutamine. Following media change, the plates were immediately placed into an Incucyte live-cell imaging system (Sartorius). Phase-contrast images were automatically acquired at 10x magnification every 4 h for 10 days. Cell proliferation was evaluated by quantifying phase-contrast cell confluency over time using the Incucyte integrated analysis software (Incucyte 2022B Rev2).

### Mathematical model of CD98hc intracellular transport

CD98hc trafficking dynamics were modeled using a deterministic, three-compartment ordinary differential equation (ODE) system based on first-order mass-action kinetics.

The model captures the distribution of receptors across three functional compartments: plasma membrane, early endosomes and recycling endosomes.

The temporal dynamics of the mobile receptor pools are governed by the following system of ODEs:

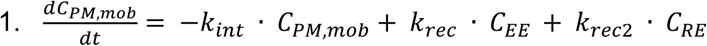

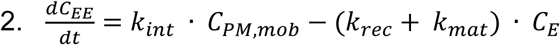

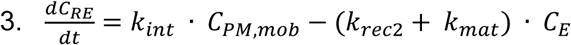

where *k_int_* is the total internalization rate, *k_rec_* is the recycling rate from early endosomes, *k_mat_* is the maturation rate from early endosomes to recycling endosomes and *k_rec_*_2_ is the recycling rate from recycling endosomes. kintrepresents the sum of a constant, constitutive basal endocytosis rate *k_basal_* and an agonist-driven rate. The dose-dependent effect of the agonist (dobutamine) was modeled using a Hill equation to account for receptor occupancy and cooperative signaling dynamics:

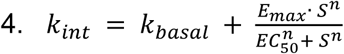

where S is the concentration of dobutamine, *E_max_* is the maximal induced internalization rate, *EC*_50_ is the half-maximal effective concentration and *n* is the Hill coefficient.

Because the experimental timeframe (120 minutes) was shorter than the expected half-life of CD98hc, receptor mass was assumed to be strictly conserved during the assay with the rate of degradation and rate of synthesis equal to 0.

To reconcile mass conservation with the experimentally measured fluorescence intensities, an optical scaling factor *scale_endo_* was introduced into the model. This parameter corrects for different fluorescence intensities between plasma membrane and endosomes arising from quenching and imaged volumes.

To account for the experimental observation of high receptor density at the plasma membrane, the model incorporates an immobile fraction pool *f_imm_*. At *t* = 0, the total baseline plasma membrane pool *C_P_*_0_ is thus partitioned into a mobile fraction *C_PM_*_,*mob*_ = *C_P_*_0_ · (1 − *f_imm_*) that can undergo endocytosis and a static, immobile fraction *C_PM_*_,*imm*_ = *C_P_*_0_ · *f_imm_*.

The simulated CD98hc concentrations were mapped to the experimental readout using:

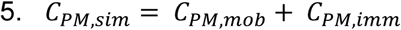

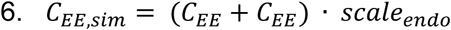

Model parameters were estimated by fitting the ODE system to the experimental time-course data spanning four dobutamine concentrations (10, 1000, 10000 and 100000 nM) simultaneously for both the plasma membrane and endosomal compartments.

To ensure structural identifiability of the model, initial steady-state conditions (i.e. prior to dobutamine stimulation) at *t* = 0 were not fitted arbitrarily but were defined from the experimental baseline data for each genotype. The initial basal internalization *k_basal_* was calculated analytically for each iteration to satisfy the steady-state assumption at *t* = 0:

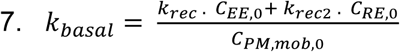

To prevent the optimization algorithm from artificially neglecting the lower-magnitude endosomal measurements in favor of the higher magnitude plasma membrane measurements, the loss function was formulated using relative (fractional residuals). The objective function minimized the sum of squared relative errors:

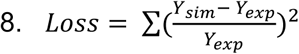

Custom Python scripts (v3.x) for ODE integration and global optimization were developed with coding assistance from Gemini (3.1 Pro). Parameter estimation was executed using the scipy.optimize.least_squares module employing the Trust Region Reflective algorithm with parameter bounds to ensure biological plausibility (all rates > 0). The deterministic ODE system was numerically integrated at each optimization using the scipy.integrate.odeint function in Python (v3.x). This function relies on the LSODA algorithm from the FORTRAN ODEPACK library. During the integration, the agonist concentration *S* was treated as a step function, transitioning instantly from 0 to the target dose at *t* = 0.

To identify the genotype-specific changes in parameters, the global pharmacological constants (*EC*_50_, *n*) and the optical scaling parameter (*scale_endo_*) were shared constraints across both CD98hc WT and ISM datasets. Conversely, to evaluate the different trafficking dynamics, the maximal induced response (*E_max_*), immobile fraction (*f_imm_*) and all intracellular kinetic rates were fitted independently for CD98hc WT and ISM genotypes. The goodness of fit was evaluated using the unweighted coefficient of determination (*R*^2^). The global fitting procedure was constrained by simultaneously fitting the numerical integration output (*Y_sim_*) against a matrix of 96 independent experimental data points. This dataset comprised paired readouts for both the plasma membrane and endosomal compartments, tracked across the two genotypes (CD98hc WT and ISM), stimulated at four dobutamine concentrations and sampled across six discrete time points.

### Mathematical model of CD98hc plasma membrane variability

To evaluate the buffering capacity of the CD98hc trafficking circuit, the deterministic ODE system was translated into a system of stochastic differential equations (SDEs). This framework allowed to simulate cell-to-cell variability driven strictly by intrinsic kinetic noise, i.e. the stochastic nature of endocytosis and vesicle fission, while intentionally controlling for extrinsic noise (e.g. baseline receptor expression and cell cycle stage).

By treating the total mobile receptor pool as conserved (*C_tot_* = *C_PM_* + *C_EE_* + *C_RE_*) and lumping together the endosomal recycling rates, the deterministic dynamics of the mobile plasma membrane pool can be analytically mapped to an Ornstein-Uhlenbeck (OU) process:

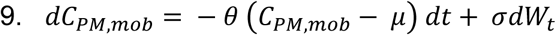

where *dW_t_* represents a standard Wiener process accounting for the continuous random nature of intracellular vesicular exchange at the macroscale level of a single cell, and *σ* represents the absolute amplitude of that intrinsic biological noise. In this OU formulation, the equilibrium set-point *μ* and the homeostatic restoring force *θ* are explicitly defined by the trafficking rates:

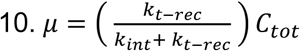

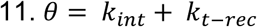

To simulate population-level distributions of CD98hc without violating physical mass conservation, the stochastic noise was modeled as a dynamic exchange flux between the mobile plasma membrane and the early endosome. The full three-compartment SDE system was formulated as:

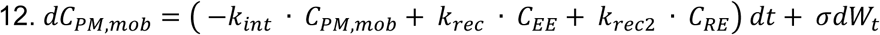

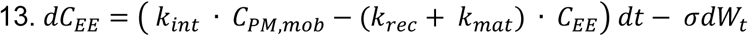

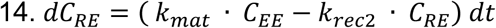

The SDE system was integrated numerically using the Euler-Maruyama method in Python (v3.x). To ensure computational stability and precision during the stochastic integration of stiff parameters, a high-resolution fixed time step (dt = 0.05 minutes) was used.

Population variability was simulated by initializing a computational cohort of N = 1000 independent cells for each genotype. All cells within a given cohort were strictly anchored to the experimental baseline measurements at t = 0 prior to dobutamine stimulation. This allowed to isolate the variance induced exclusively by the kinetic trafficking network differences rather than arbitrarily distributed starting conditions. An identical basal noise amplitude *σ* = 0.02 was applied to both genotype cohorts.

Each of the 1000 single-cell trajectories was simulated over a continuous 120 minute window corresponding to continuous dobutamine exposure at 10 μM. To generate the population distribution maps, the total surface CD98hc level was extracted at specific time points t = 0, 10 and 120 minutes.

To fairly assess the spread of populations possessing distinct baseline macroscopic averages, the relative inter-cellular variability was quantified using the coefficient of variation (CV), defined as the standard deviation of the computational cohort normalized by its time-matched arithmetic mean 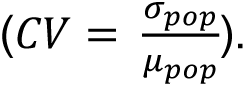

### Antibodies

The following primary antibodies were used for immunostaining: anti-eGFP (ab13970, Abcam; used at 1:500 dilution); anti-β1-Integrin (ab30394 (clone 12G10), Abcam; used at 1:500 dilution); anti-EEA1 (3288S, Cell Signaling; used at 1:300 dilution); anti-EEA1 (610457, BD Biosciences; used at 1:300 dilution); anti-LAMP1 (9091, Cell Signaling; used at 1:500 dilution); VE-Cadherin (2500S, Cell Signaling; used at 1:200 dilution); TfR (13-6800, ThermoFisher; used at 1:300 dilution).

The following primary antibodies were used for immunobloting: anti-CD98hc (ab307587 [EPR27110-42], Abcam, used at 1:1000 dilution); anti-LAT1 (#32683, Cell Signaling, used at 1:1000 dilution); anti-β-Actin (#12262, Cell Signaling, used at 1:100 dilution); anti-eGFP (ab290, Abcam; or ab13970, Abcam; both used at 1:1000 dilution)

The following primary antibodies were used for microscopy: anti-eGFP (ab290, Abcam; or ab13970, Abcam; both used at 1:1000 dilution); anti-Rab5 (C8B1) (#3547, Cell Signaling, used at 1:200 dilution); B1-Integrin (ab30394, Abcam, used at 1:500 dilution), anti-CD98hc mAb’s conjugated to AF488 or AF647 (315602, BioLegend, used at final 100nM concentration for live cell imaging)

We used a monovalent antibodies against the extracellular domain of the human transferrin receptor (TfR)^42^ and against the extracellular domain of the human CD98hc.

The following secondary antibodies were used for microscopy: Alexa Fluor 488 Donkey Anti-Chicken IgY (IgG) (703-545-155; Jackson ImmunoResearch), Alexa Fluor 488 donkey anti-mouse IgG (715-545-150, Jackson ImmunoResearch), Alexa Fluor 488 donkey anti-rabbit IgG, (711-545-152, Jackson ImmunoResearch), Alexa Fluor 488 donkey Anti-human IgG (709-545-149, Jackson ImmunoResearch), Alexa Fluor 594 donkey Anti-mouse IgG (715-585-150, Jackson ImmunoResearch), Alexa Fluor 647 donkey Anti-rabbit IgG (711-605-152, Jackson ImmunoResearch), Alexa Fluor 647 donkey Anti-human IgG (709-605-149, Jackson ImmunoResearch), and for immunoblot; donkey anti-mouse IgG-HRP conjugated (A16011, Invitrogen) and donkey anti-rabbit IgG-HRP conjugated (A16035, Invitrogen). DNA was stained using DAPI (D9542, Sigma), cytoplasm was stained with CellMask Blue Stain (H32720, ThermoFisher), and transferrin was labeled with Transferrin-488 (T2871, ThermoFisher) or Transferrin-AF647 (T23366, ThermoFisher) at 25 μg/mL final concentration.

## Author contributions

F.S.R contributed to project administration, conceptualization, data curation, methodology, investigation, visualization, writing—original draft, and writing—review & editing. H.P.G. contributed to methodology, simulation and data analysis. A.A. contributed to study design, data curation, data generation and analysis. JL.G.C contributed to conceptualization, data analysis and simulation. R.V. contributed to project administration, supervision, conceptualization, data curation, methodology, visualization, writing—original draft, and writing—review & editing and decision to submit. All authors reviewed and edited the manuscript.

## Acknowledgements

We thank Marcus Bantscheff and Balazs Banfai from 360 labs, pRED, for support in data generation and analysis. We thank Telma Lopes and Antoine Rizkallah for excellent technical support. We thank Jens Niewoehner, Thomas Kraft and Anna-Lena Bolender for reagent generation. We thank Martina Pigoni for critically reviewing the manuscript. Finally, we thank Gilbert di Paolo for his feedback and guidance. Sila Rizalar was supported by a fellowship from the Roche Postdoctoral program. The manuscript was reviewed for consistency and grammar using a large language model (Gemini 3.1 Pro).

---

**Table S1.**
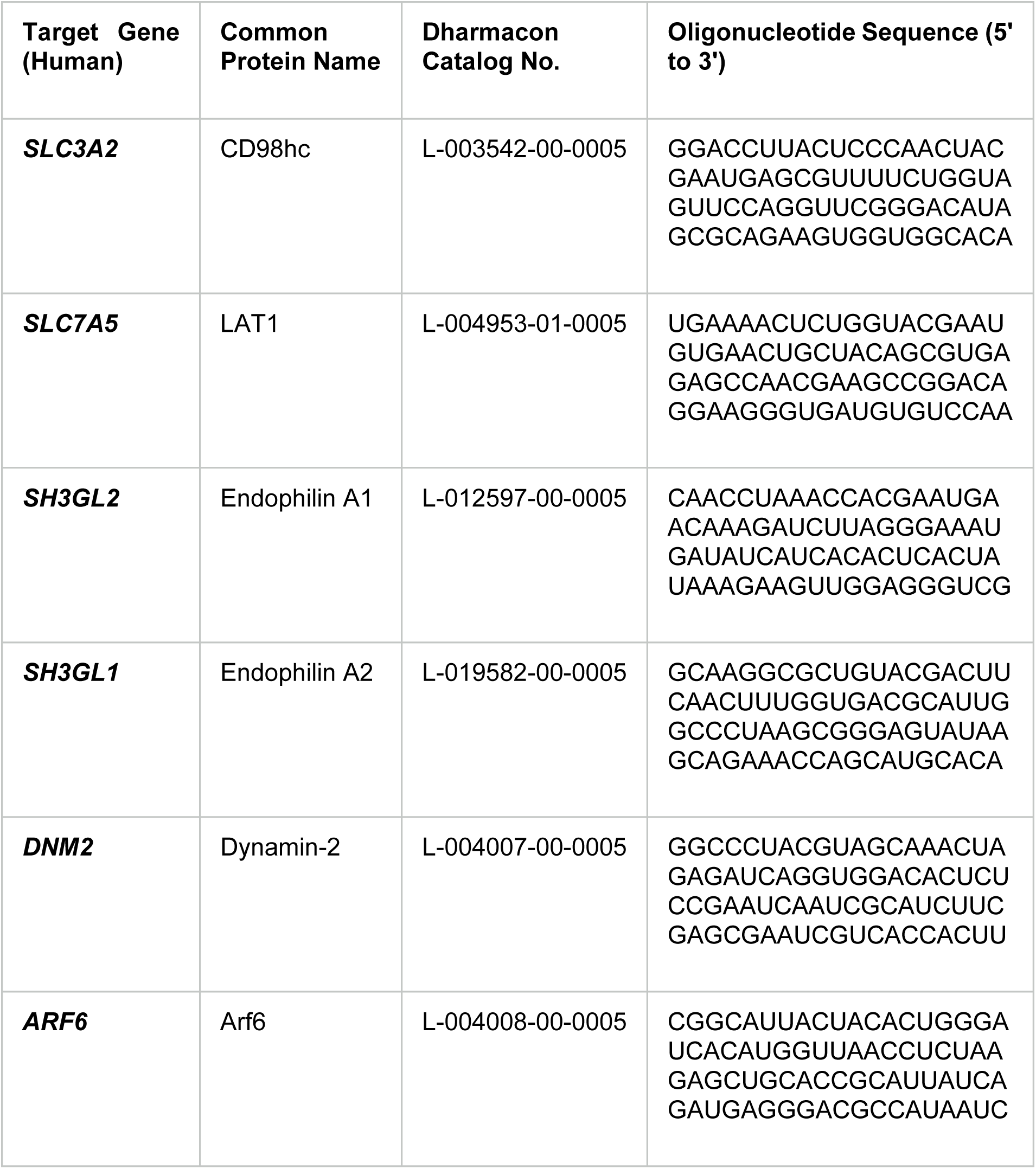
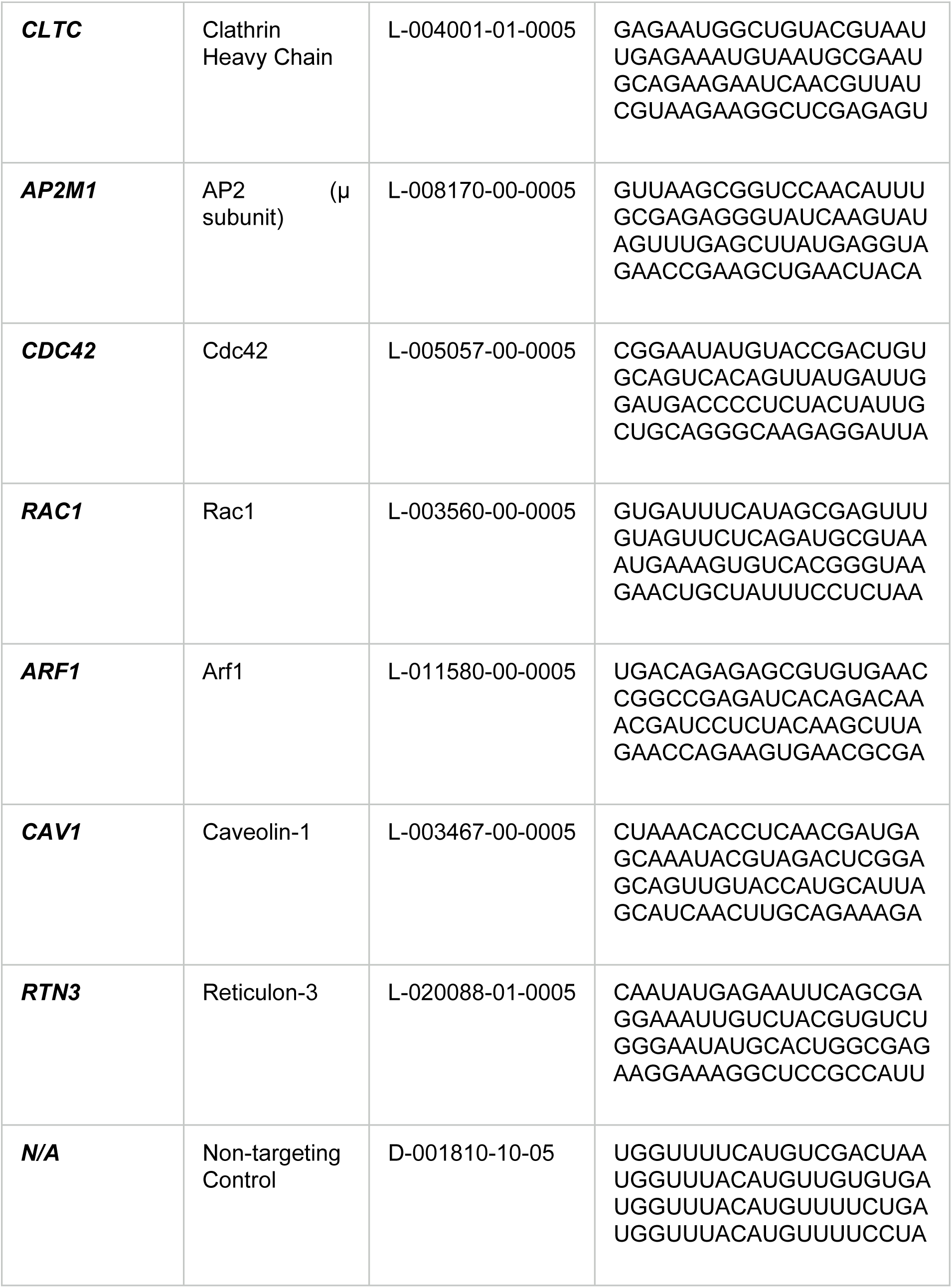
List of siRNAs.

**Table S2.** List of Small-Molecule Inhibitors and Pathway Agonists.

| Compound Name | Functional Target / Pathway | Source / Catalog No. | Final Concentration |
| --- | --- | --- | --- |
| **Dyngo 4a** | Dynamin inhibitor | Abcam (ab120689) | 80 µg/mL |
| **Pitstop 2** | Clathrin-mediated endocytosis inhibitor | Abcam (ab120687) | 30 µM |
| **GDC-0941** | Broad and specific PI(3)K inhibitor (called PI3Ki in this study) | MedChemExpress (HY-50094) | 1 µM |
| **P505-15** | Spleen tyrosine kinase (SYK) inhibitor (called SYKi in this study) | MedChemExpress (HY-15322) | 1 µM |
| **CHIR-99021** | Glycogen synthase kinase 3 (GSK3) inhibitor (called GSK3i 1 in this study) | Cayman Chemical (13122) | 1 µM |
| **BIO-6-bromoindirubin-3'-oxime** | GSK3α/β inhibitor (called GSK3i 2 in this study) | Sigma-Aldrich (B1686) | 1 µM |
| **Dinaciclib** | CDK1, CDK2, CDK5, and CDK9 inhibitor (called Cdk1/2/5/9i in this study) | MedChemExpress (HY-10492) | 1 µM |
| **Roscovitine** | CDK1, CDK2, and CDK5 selective inhibitor (called Cdk1/2/5i in this study) | MedChemExpress (HY-30237) | 1 µM |
| **Dobutamine** | Partial β<sub>1</sub>-adrenergic receptor agonist | MedChemExpress (HY-15746A) | 10 µM |
| **Denopamine** | Selective $\beta_1$-adrenergic receptor agonist | MedChemExpress (HY-119515) | 10 μM |
| **EIPA (5- (N-ethyl-N-isopropyl)amiloride)** | Na<sup>+</sup>/H<sup>+</sup> exchanger (NHE) inhibitor | MedChemExpress (HY-101840) | 25 μM |
| **Imipramine** | Fluid-phase endocytosis/macropinocytosis inhibitor | MedChemExpress (HY-B1490A) | 5 μM |
| **Latrunculin A** | Actin polymerization inhibitor (G-actin sequestering) | MedChemExpress (HY-16929) | 1 μM |
| **Cytochalasin D** | Actin filament depolymerizer | MedChemExpress (HY-N6682) | 1 μM |
| **Torin 1** | mTORC1/mTORC2 inhibitor (called mTORCi in this study) | Tocris Bioscience (4247) | 10 μM |

