## Supplementary Figures for "Integrin-dependent trafficking of CD98hc reduces endocytic noise to maintain metabolic homeostasis"

#### An integrin-dependent trafficking circuit buffers noise to maintain metabolic homeostasis

**Table S4**. Kinetic rate constants and best-fit parameters for CD98hc intracellular transport

Parameter fits $R^2 = 0.99$

| **Parameter** | **WT** | **Mutant** |
|---|---|---|
| $E_{max}$ | 0.0259 min-1 | 0.0392 min-1 |
| $EC_{50}$ | 38.209 nM | |
| n | 0.1693 | |
| $scale_{endo}$ | 0.0789 | |
| $f_{imm}$ | 20.91% | 40.81% |
| $k_{basal}$ | 0.1061 min-1 | 0.043 min-1 |
| $k_{rec}$ | 0.97 min-1 | 0.2546 min-1 |
| $k_{mat}$ | 0.56267 min-1 | 0.1384 min-1 |
| $k_{rec2}$ | 0.0310 min-1 | 0.0081 min-1 |

Rates expressed as normalized fluorescence intensity units/minute. Goodness of fit ($R^2 = 0.99$) represents the agreement between the experimental time course data and the model-predicted mean trajectory.

Figure S1

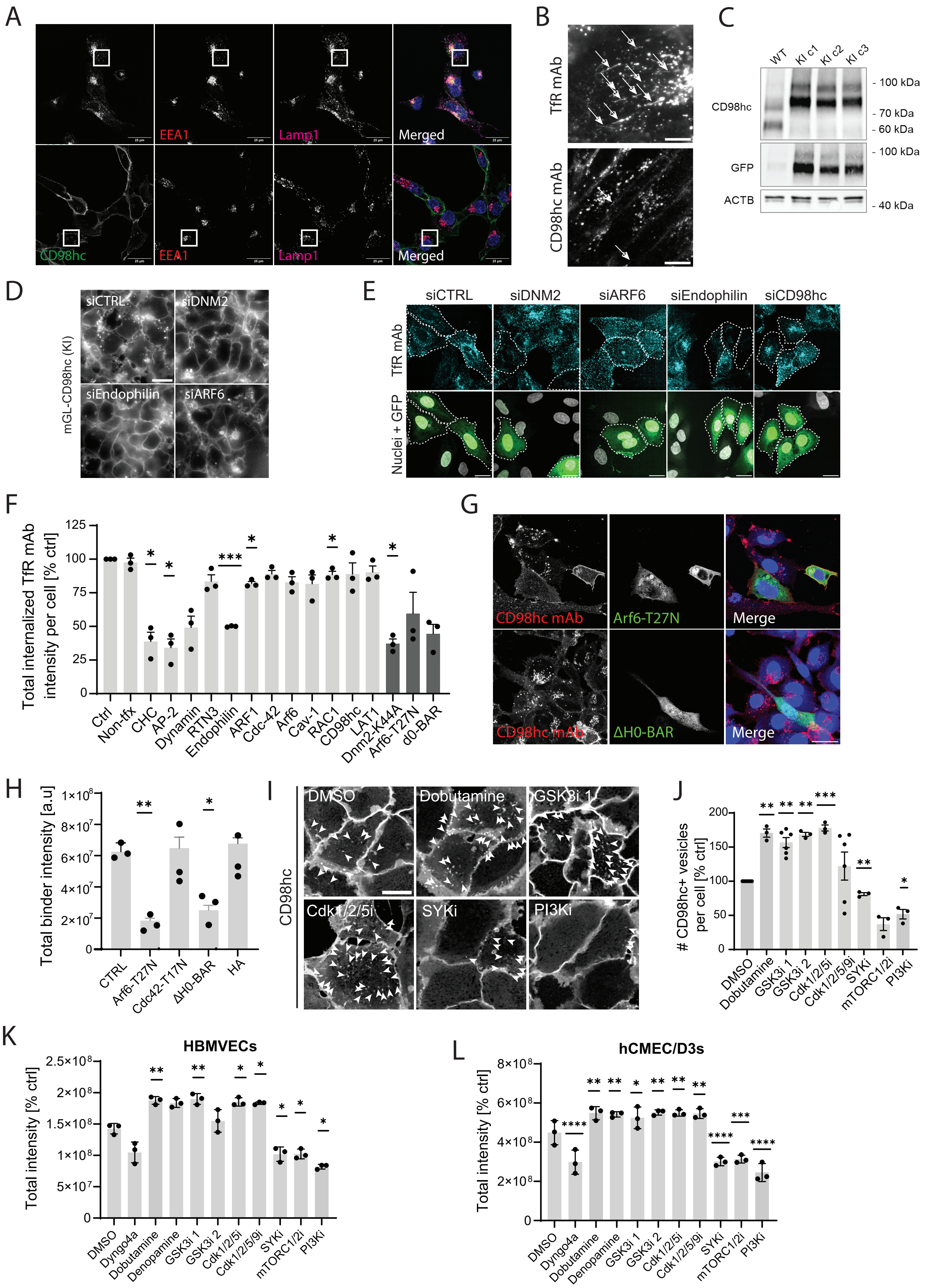
A
EEA1
Lamp1
Merged
CD98hc
EEA1
Lamp1
Merged
B
TfR mAb
CD98hc mAb
C
WT
KI c1
KI c2
KI c3
CD98hc
GFP
ACTB
- 100 kDa
- 70 kDa
- 60 kDa
- 100 kDa
- 40 kDa
D
mGL-CD98hc (KI)
siCTRL
siDNM2
siEndophilin
siARF6
E
siCTRL
siDNM2
siARF6
siEndophilin
siCD98hc
TfR mAb
Nuclei + GFP
F
Total internalized TfR mAb intensity per cell [% ctrl]
Ctrl
Non-tfx
CHC
AP-2
Dynamin
RTN3
Endophilin
ARF1
Cdc-42
Arf6
Cav-1
RAC1
CD98hc
LAT1
Dnm2-K44A
Arf6-T27N
d0-BAR
G
CD98hc mAb
Arf6-T27N
Merge
CD98hc mAb
ΔH0-BAR
Merge
H
Total binder intensity [a.u]
CTRL
Arf6-T27N
Cdc42-T17N
ΔH0-BAR
HA
I
CD98hc
DMSO
Dobutamine
GSK3i 1
Cdk1/2/5i
SYKi
PI3Ki
J
### CD98hc+ vesicles per cell [% ctrl]
DMSO
Dobutamine
GSK3i 1
GSK3i 2
Cdk1/2/5i
Cdk1/2/5/9i
SYKi
mTORC1/2i
PI3Ki
K
HBMVECs
Total intensity [% ctrl]
DMSO
Dyngo4a
Dobutamine
Denopamine
GSK3i 1
GSK3i 2
Cdk1/2/5i
Cdk1/2/5/9i
SYKi
mTORC1/2i
PI3Ki
L
hCMEC/D3s
Total intensity [% ctrl]
DMSO
Dyngo4a
Dobutamine
Denopamine
GSK3i 1
GSK3i 2
Cdk1/2/5i
Cdk1/2/5/9i
SYKi
mTORC1/2i
PI3Ki

**Figure S1. A,** Representative confocal images of iCE-BECs (day 14) treated with TfR mAb (top) or CD98hc mAb (bottom), and stained for EEA1 and LAMP1, showing the full panel of the image in Figure 1B. Scale bar, 10 µm. **B,** Representative images of iCE-BECs (day 14) treated with fluorescence-conjugated TfR mAb (top) or CD98hc mAb (bottom) for 3 hours. Arrows highlight tubule structures. Scale bar, 10 µm. **C,** Western blot showing three independent iPS clones (c1, c2 and c3) with successful endogenous eGFP-CD98hc expression alongside the parental control clone (WT). β-Actin was used as a loading control. **D,** Representative images showing mGL-CD98hc knockin hiPSCs nucleofected with siRNAs against a non-targeting control (siCTRL), Dynamin-2 (siDNM2), Arf6 (siARF6) or Endophilin A1 and A2 (siEndophilin). Scale bar, 20 µm. **E,** Representative images of TfR mAb localization in resting U2OS cells upon knockdown of selected endocytosis-related proteins. Nucleofected cells were highlighted with dashed lines. Scale bar, 10 µm. **F,** Targeted TfR mAb internalization screen using siRNAs or dominant-negative mutants of selected proteins across diverse endocytic pathways. WT U2OS cells were nucleofected with selected siRNAs together alongside a GFP reporter plasmid (shown in light-gray bars), or with dominant-negative mutation-expressing plasmids fused to a GFP or HA reporter (shown in dark-gray bars). 48 h post-nucleofection cells were treated with TfR mAb for 1 h. Bars show the mean ± SEM from a minimum of 100 cells per condition, from three independent biological experiments. Statistical analysis was performed by one-way ANOVA followed by Dunnet's test. *P < 0.05, ***P < 0.001 **G,** Representative images of hMEC/D3 cells nucleofected with dominant-negative Arf6 (Arf6-T27N-HA) (top) or dominant negative Endophilin BAR domain (eGFP-ΔH0-BAR) (bottom) expressing constructs and 48 h after incubated with CD98hc mAb for 1 h. Scale bar, 10 µm. **H,** Quantification of internalized CD98hc mAb intensity in hMEC/D3 cells which are non-transfected (CTRL) or expressing Arf6-T27N-HA, Cdc42-T17N-HA, eGFP-ΔH0-BAR or HA constructs. Bars show the mean ± SEM from minimum 30 cells per condition, from three independent biological experiments. Statistical analysis was performed by one-way ANOVA followed by Dunnet's test. *P < 0.05, **P < 0.01. **I,** Representative images of intracellular distribution of CD98hc in resting U2OS cells treated with 10 µM DMSO, 10 µM dobutamine, 10 µM GSK3i1, 1 µM Cdk1/2/5i, 1 µM SYKi, and 10 nM GDC-0941 (PI3Ki). Arrows highlight selected CD98hc-positive vesicles. Scale bar, 20 µm. **J,** Bar graph shows the normalized number of CD98hc-positive vesicles after incubation with DMSO, (vehicle); dobutamine, 10 µM ; CHIR-99041 (GSK3i 1), 1 µM; BIO-6-bromoindirubin- 3′-oxime (GSK3i 2), 1µM; Roscovitine (Cdk1/2/5i), 1 µM; Dinaciclib (Cdk1/2/5/9i), 1 µM; P505-15 (SYKi), 1 µM; Torin 1 (mTORC1/2i), 10 µM; and GDC-0941 (PI3Ki), 1 µM. Bars show the mean ± SEM from minimum 100 cells per condition, from three independent biological experiments. Statistical analysis was performed by one-way ANOVA followed by Dunnet's test. *P < 0.05, **P < 0.01, ***P < 0.001. **K,L,** Graphs show the normalized total CD98hc mAb vesicle intensity in primary human brain microvascular endothelial cells (HBMVECs, **K**) and an immortalized brain endothelial cell line (hCMEC/D3, **L**) after incubation with FEME activators and inhibitors as in Figure 1; DMSO, (vehicle); Dyngo4a, 80 µg/ml; dobutamine, 10 µM ; denopamine, 10 µM; CHIR-99041 (GSK3i 1), 1 µM; BIO-6-bromoindirubin- 3′-oxime (GSK3i 2), 1µM; Roscovitine (Cdk1/2/5i), 1 µM; Dinaciclib (Cdk1/2/5/9i), 1 µM; P505-15 (SYKi), 1 µM; Torin 1 (mTORC1/2i), 10 µM; and GDC-0941 (PI3Ki), 1 µM. Bars show the mean ± SEM from minimum 100 cells per condition, from three independent biological experiments. Statistical analysis was performed by one-way ANOVA followed by Dunnet's test. *P < 0.05, **P < 0.01, ***P < 0.001.

Figure S2

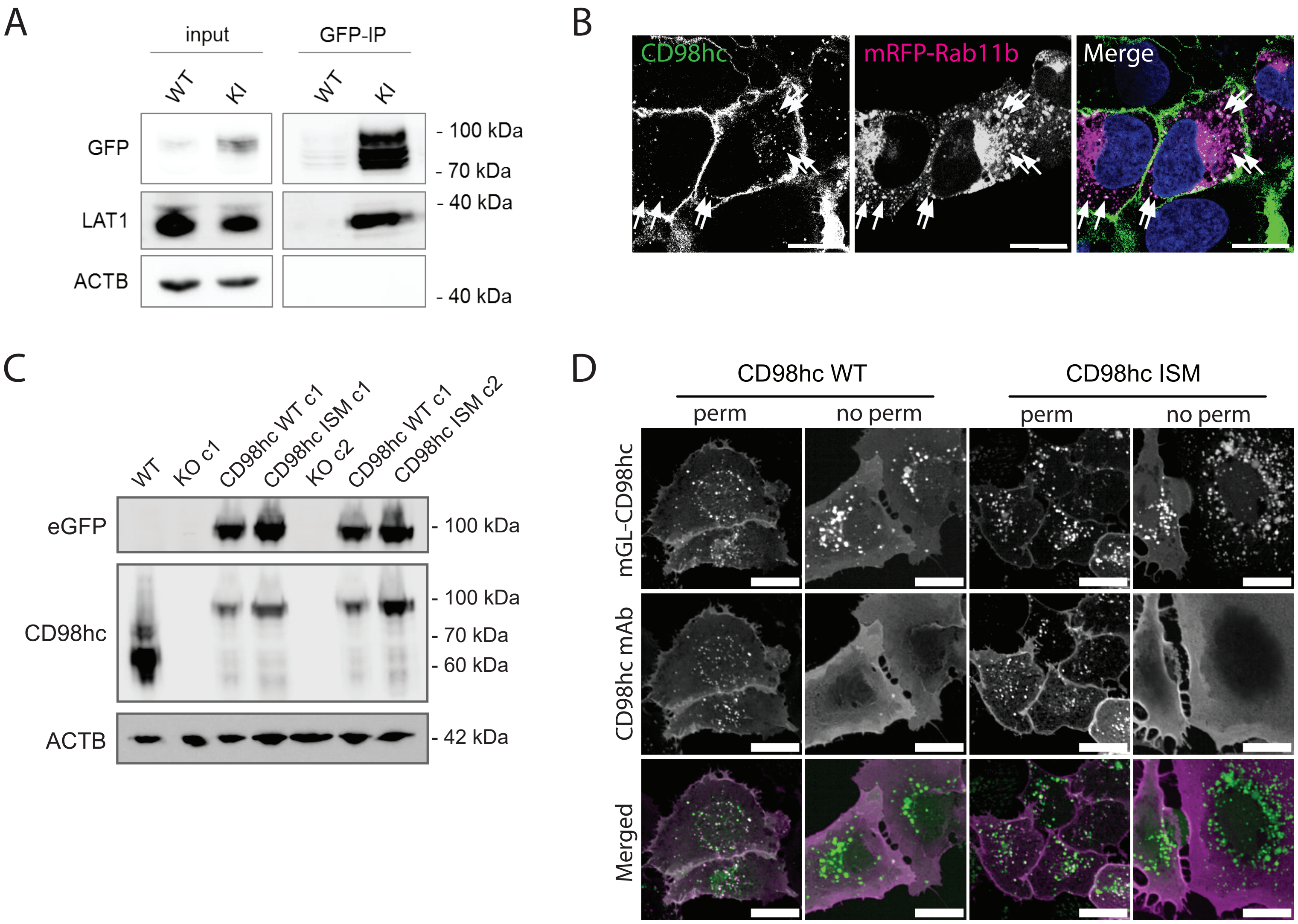

A
input
GFP-IP
WT
KI
WT
KI
GFP
LAT1
ACTB
- 100 kDa
- 70 kDa
- 40 kDa
- 40 kDa
B
CD98hc
mRFP-Rab11b
Merge
C
WT
KO c1
CD98hc WT c1
CD98hc ISM c1
KO c2
CD98hc WT c1
CD98hc ISM c2
eGFP
CD98hc
ACTB
- 100 kDa
- 100 kDa
- 70 kDa
- 60 kDa
- 42 kDa
D
CD98hc WT
CD98hc ISM
perm
no perm
perm
no perm
mGL-CD98hc
CD98hc mAb
Merged

**Figure S2. A,** Immunoprecipitation of endogenous mGreenLantern-CD98hc in WT or CD98hc-KI iPSCs. Bound proteins were detected by immunoblotting. Total lysate blots represent 5% of material used as input for coimmunoprecipitation. IP, immunoprecipitate. β-Actin was used as the loading control. **B,** Representative images of endogenous CD98hc with mRFP-Rab11b in U2OS cells. Arrows highlight selected co-localized puncta. Scale bar, 20 µm. **C,** Western blot showing successful generation of CD98hc knockout U2OS clones (KO c1 and c2) as well as the stable KO lines expressing WT CD98hc (CD98hc WT c1 and CD98hc WT c2) or ISM CD98hc (CD98hc ISM c1 and CD98hc ISM c2). β-Actin was used as the loading control. **D,** Full panel of representative confocal images shown in Figure 2 H, showing CD98hc WT or CD98hc ISM cells after incubation with CD98hc mAb for 30 minutes. CD98hc mAb was stained with- or without-permeabilization (perm; no perm). Scale bar, 20 µm.

Figure S3

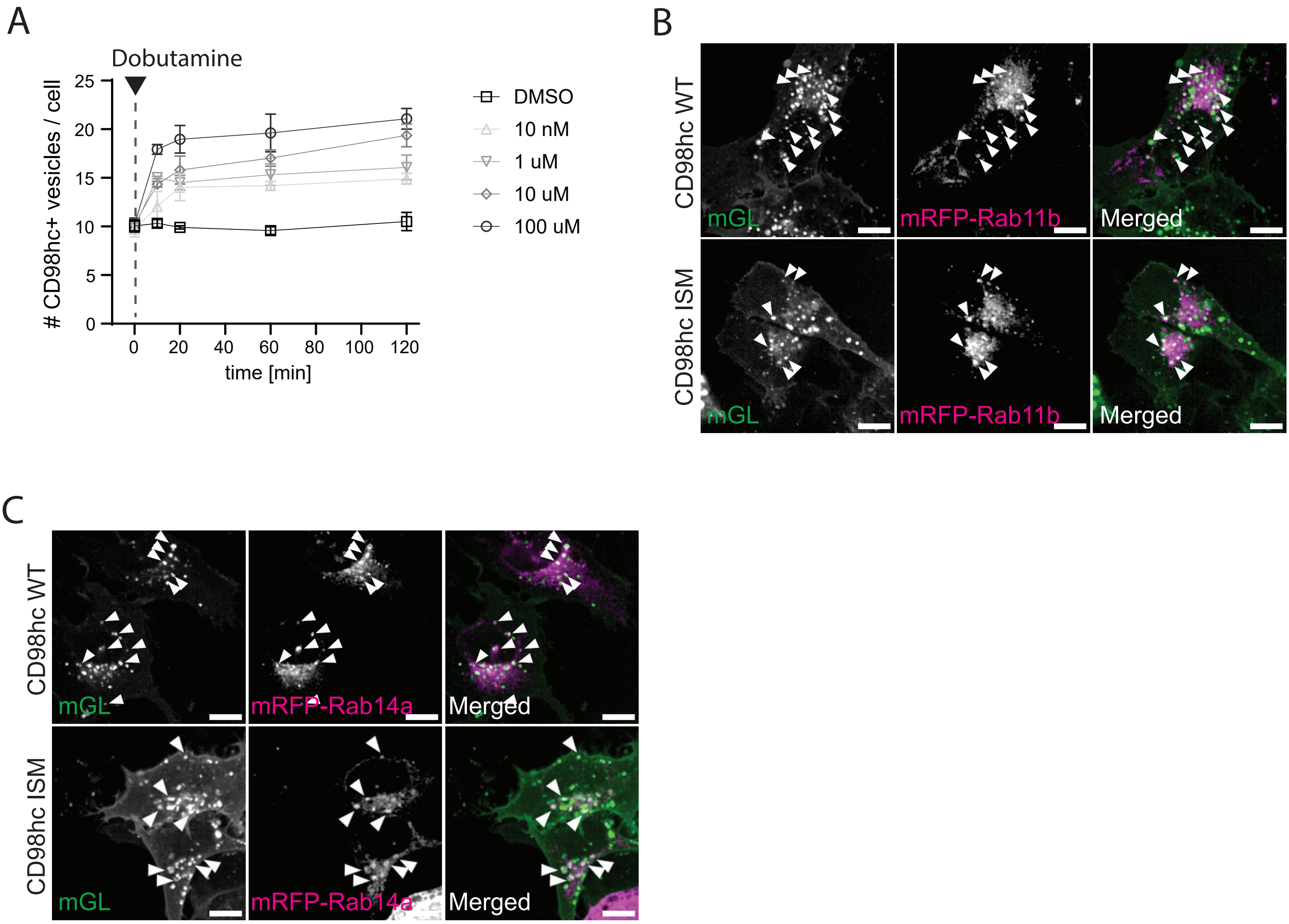
A
Dobutamine
### CD98hc+ vesicles / cell
25
20
15
10
5
0
0
20
40
60
80
100
120
time [min]
DMSO
10 nM
1 uM
10 uM
100 uM
B
CD98hc WT
CD98hc ISM
mGL
mRFP-Rab11b
Merged
C
CD98hc WT
CD98hc ISM
mGL
mRFP-Rab14a
Merged

**Figure S3. A,** Dose-response of CD98hc vesicle accumulation in wild-type U2OS cells across indicated concentrations of Dobutamine (10 nM to 100 µM) over a 120-minute time course. Data are presented as mean ± SEM. **B,** Representative confocal images of CD98hc WT or CD98hc ISM U2OS lines co-expressing the endosomal marker mRFP-Rab11b (magenta). Arrowheads highlight mGL-CD98hc positive vesicles colocalizing with Rab11. Scale bars, 20 µm. **C,** Representative confocal images of CD98hc WT or CD98hc ISM U2OS lines co-expressing the endosomal marker mRFP-Rab14 (magenta). Arrowheads highlight mGL-CD98hc positive vesicles colocalizing with Rab14. Scale bars, 20 µm.

Figure S4

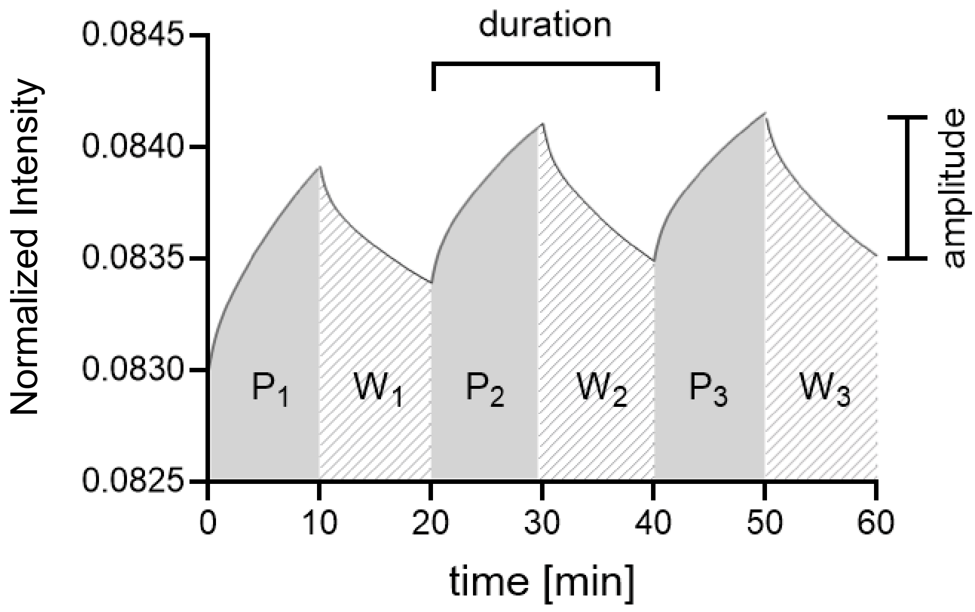

duration
amplitude
Normalized Intensity
0.0845
0.0840
0.0835
0.0830
0.0825
P1
W1
P2
W2
P3
W3
0
10
20
30
40
50
60
time [min]

**Figure S4.** Simulation of intracellular CD98hc intensity using best fit parameters of ODE from Figure 3 after oscillatory 10 minute pulses of 10 μM dobutamine. The amplitude is defined as the mean difference between the value at the end of the pulse minus the value at the end of the washout period. The duration is defined as the length of each pulse.

Figure S5

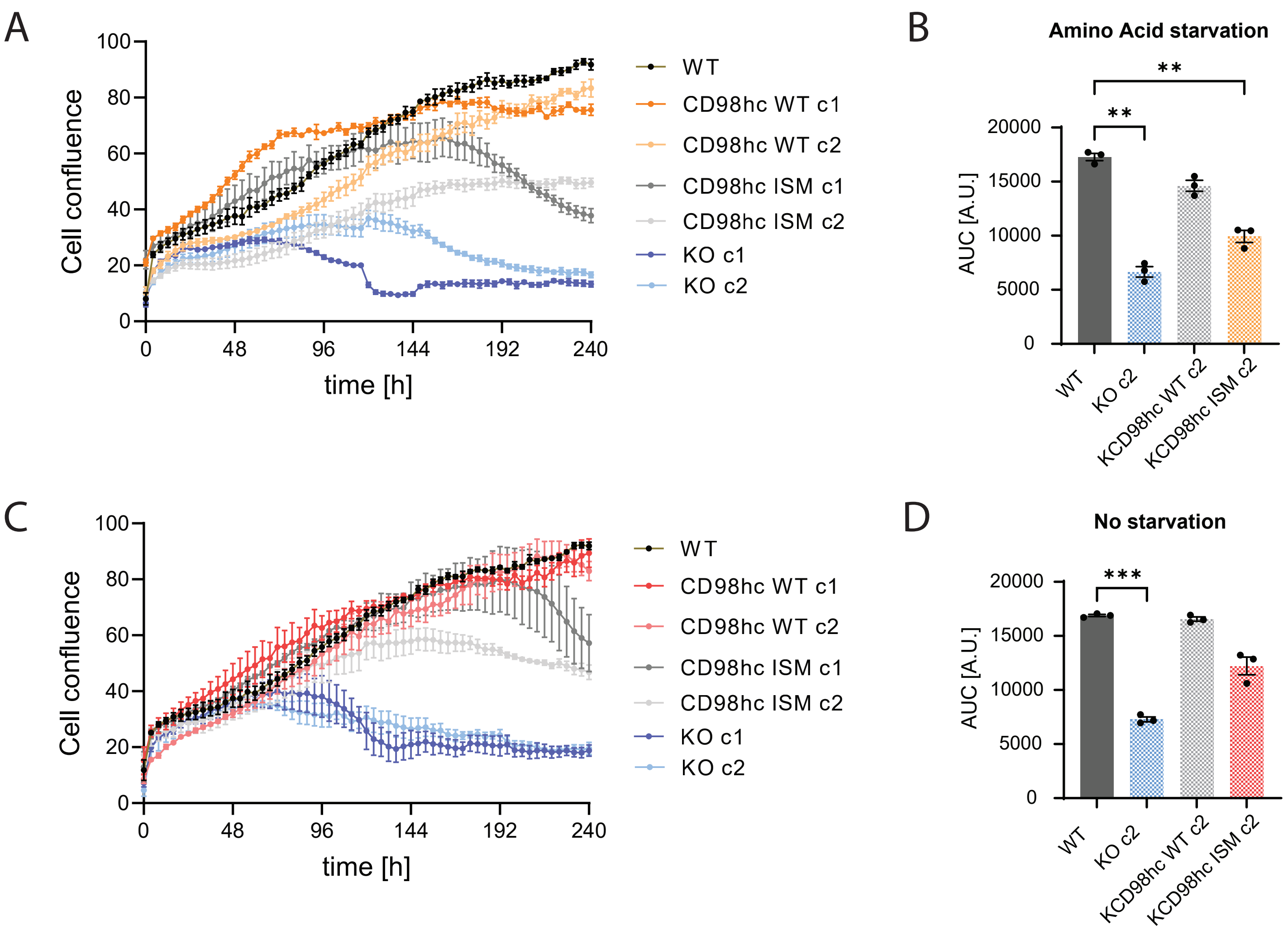
A
Cell confluence
time [h]
0
48
96
144
192
240
100
80
60
40
20
0
WT
CD98hc WT c1
CD98hc WT c2
CD98hc ISM c1
CD98hc ISM c2
KO c1
KO c2
B
Amino Acid starvation
AUC [A.U.]
20000
15000
10000
5000
0
**
**
WT
KO c2
KCD98hc WT c2
KCD98hc ISM c2
C
Cell confluence
time [h]
0
48
96
144
192
240
100
80
60
40
20
0
WT
CD98hc WT c1
CD98hc WT c2
CD98hc ISM c1
CD98hc ISM c2
KO c1
KO c2
D
No starvation
AUC [A.U.]
20000
15000
10000
5000
0
***
WT
KO c2
KCD98hc WT c2
KCD98hc ISM c2

**Figure S5. A, C,** Cell growth measured as increase in confluency of WT, CD98hc KO, and CD98hc WT or ISM U2OS lines for two independent clones in media containing low amino acid levels (10%) (**A**) or full growth media (**C**) for 10 days. The trajectories for clone 1 are also shown in Figure 4I. Data represent mean ± SEM from n=3 independent experiments. **B and D,** Comparison of cell growth for clone 2 measured as area under the curve (AUC) of trajectories shown in **A** and **C** for cells grown under amino acid starvation (**B**) or full media (**D**). Bars show mean ± SEM from n=3 independent experiments. Statistical analysis was performed by one-way Anova followed by Dunnett's test. $^{**}P < 0.01$, $^{***}P < 0.001$.

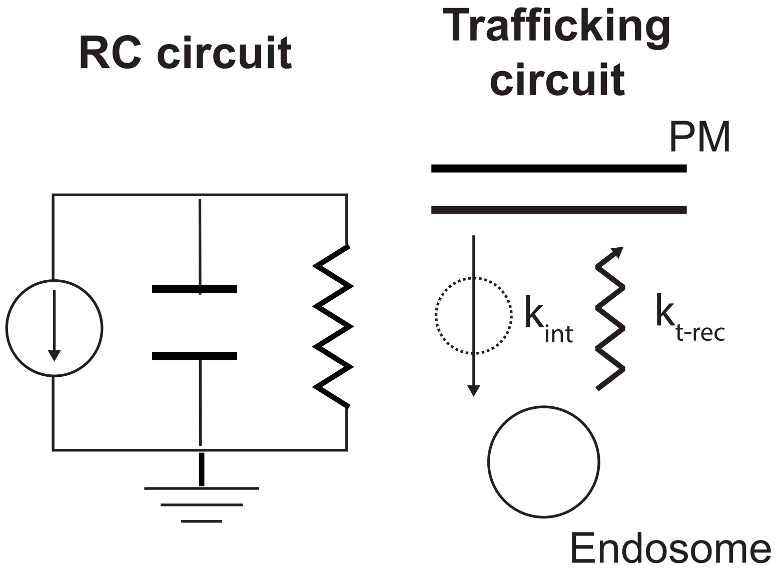
RC circuit
Trafficking circuit
PM
$k_{int}$
$k_{t\text{-}rec}$
Endosome

**Figure S6.** Diagram of a parallel RC circuit with a current sink, capacitor and resistor (left) compared to the trafficking circuit of CD98hc (right). The plasma membrane (PM) is analogous to the capacitor, endocytosis to the current sink, and recycling to the resistor.

**Supplementary Video 1,** Fluorescence-conjugated TfR mAb in iCE-BECs (day 14) after treatment for 3 hours. Scale bar, 10 µm. Related to Figure 1C and Figure S1 B.

**Supplementary Video 2,** Fluorescence-conjugated CD98hc mAb in iCE-BECs (day 14) after treatment for 3 hours. Scale bar, 10 µm. Related to Figure 1C and Figure S1 B.
