## Supplementary Table 3 for "Integrin-dependent trafficking of CD98hc reduces endocytic noise to maintain metabolic homeostasis"

| Contrast | FeatureID | GeneName | UniquePeptic | Organism | logFC | CI.L | CI.R | AveExpr | TStat | PValue | FDR | B | direction |
|---|---|---|---|---|---|---|---|---|---|---|---|---|---|
| KI - WT, cell_origin: iPSC | P35222 | CTNNB1 | 28 | Homo sapien | 3.7916955 | 2.92887566 | 4.65451534 | 0.23400181 | 9.36255631 | 1.13E-07 | 0.00020595 | 7.83522859 | up |
| KI - WT, cell_origin: iPSC | P05556 | ITGB1 | 8 | Homo sapien | 4.58700048 | 3.49913834 | 5.67486262 | -1.1314084 | 8.9858112 | 1.9709E-07 | 0.00020595 | 7.34090691 | up |
| KI - WT, cell_origin: iPSC | Q15907 | RAB11B | 13 | Homo sapien | 4.42028563 | 3.37096412 | 5.46960714 | 1.79721041 | 8.97726666 | 1.9951E-07 | 0.00020595 | 7.33003213 | up |
| KI - WT, cell_origin: iPSC | Q13636 | RAB31 | 5 | Homo sapien | -4.7584333 | -5.9026048 | -3.6142617 | 0.02440346 | -8.8628866 | 2.3503E-07 | 0.00020595 | 7.18351306 | down |
| KI - WT, cell_origin: iPSC | O43681 | GET3 | 13 | Homo sapien | 3.56556916 | 2.61071418 | 4.52042414 | 0.46990772 | 7.95781454 | 9.0716E-07 | 0.00063592 | 5.9599811 | up |
| KI - WT, cell_origin: iPSC | P02786 | TFRC | 34 | Homo sapien | 5.46377011 | 3.95715642 | 6.9703838 | 1.25129513 | 7.72845845 | 1.2979E-06 | 0.00072043 | 5.63115008 | up |
| KI - WT, cell_origin: iPSC | Q16563 | SYPL1 | 4 | Homo sapien | 5.82374313 | 4.20419686 | 7.44328941 | 0.50115622 | 7.66321866 | 1.4388E-06 | 0.00072043 | 5.53618308 | up |
| KI - WT, cell_origin: iPSC | P61019 | RAB2A | 6 | Homo sapien | 3.17925559 | 2.20840427 | 4.15010692 | 0.3591883 | 6.97870892 | 4.3886E-06 | 0.00188487 | 4.50073238 | up |
| KI - WT, cell_origin: iPSC | Q15836 | VAMP3 | 5 | Homo sapien | 3.31224005 | 2.29225665 | 4.33222346 | 1.02875405 | 6.92039791 | 4.8399E-06 | 0.00188487 | 4.40918551 | up |
| KI - WT, cell_origin: iPSC | P20340 | RAB6A | 8 | Homo sapien | 3.5011003 | 2.3824422 | 4.6197584 | 3.66184275 | 6.66974987 | 7.4104E-06 | 0.00259734 | 4.00963117 | up |
| KI - WT, cell_origin: iPSC | O14828 | SCAMP3 | 8 | Homo sapien | 2.92813679 | 1.98430106 | 3.87197251 | 0.32127141 | 6.6114591 | 8.1922E-06 | 0.00261033 | 3.91530245 | up |
| KI - WT, cell_origin: iPSC | O43752 | STX6 | 9 | Homo sapien | 7.16541782 | 4.74322561 | 9.58761003 | -2.4890599 | 6.30427696 | 1.4005E-05 | 0.00409074 | 3.40942276 | up |
| KI - WT, cell_origin: iPSC | P08195 | SLC3A2 | 29 | Homo sapien | 5.88890997 | 3.84234606 | 7.93547388 | 2.57323339 | 6.13213873 | 1.9023E-05 | 0.00512882 | 3.11950331 | up |
| KI - WT, cell_origin: iPSC | P26232 | CTNNA2 | 3 | Homo sapien | 3.19248493 | 2.07113301 | 4.31383686 | -1.0869018 | 6.06721371 | 2.1374E-05 | 0.00535116 | 3.00896221 | up |
| KI - WT, cell_origin: iPSC | P02787 | TF | 29 | Homo sapien | 3.61281554 | 2.31964561 | 4.90598547 | 0.3548421 | 5.95377695 | 2.6238E-05 | 0.00613088 | 2.81426773 | up |
| KI - WT, cell_origin: iPSC | P61106 | RAB14 | 12 | Homo sapien | 2.85814049 | 1.8208374 | 3.89544358 | 1.14416094 | 5.87192187 | 3.0455E-05 | 0.00627901 | 2.67255514 | up |
| KI - WT, cell_origin: iPSC | P54920 | NAPA | 14 | Homo sapien | 1.87685198 | 1.19602582 | 2.55767814 | 1.26243619 | 5.87128914 | 2.9638E-05 | 0.00627901 | 2.69515391 | up |
| KI - WT, cell_origin: iPSC | P04899 | GNAI2 | 14 | Homo sapien | 2.83344708 | 1.79693667 | 3.86995748 | 0.85131189 | 5.82564218 | 3.3145E-05 | 0.00642603 | 2.59198251 | up |
| KI - WT, cell_origin: iPSC | O15400 | STX7 | 9 | Homo sapien | 3.90752403 | 2.47142376 | 5.34362429 | 1.08262923 | 5.79854807 | 3.4834E-05 | 0.00642603 | 2.54466166 | up |
| KI - WT, cell_origin: iPSC | P05023 | ATP1A1 | 44 | Homo sapien | 2.8794852 | 1.80621123 | 3.95275917 | 2.33848488 | 5.71750589 | 4.0442E-05 | 0.00708748 | 2.40246022 | up |
| KI - WT, cell_origin: iPSC | P35221 | CTNNA1 | 39 | Homo sapien | 2.73694262 | 1.65649977 | 3.81738547 | 0.71461388 | 5.3984149 | 7.3417E-05 | 0.01225362 | 1.83315902 | up |
| KI - WT, cell_origin: iPSC | Q6RW13 | AGTRAP | 3 | Homo sapien | 5.34563959 | 3.19223485 | 7.49904432 | -1.7222554 | 5.29025223 | 9.0136E-05 | 0.01436032 | 1.63687589 | up |
| KI - WT, cell_origin: iPSC | P11166 | SLC2A1 | 6 | Homo sapien | 6.13176692 | 3.63198554 | 8.63154829 | -1.5456665 | 5.22740294 | 0.00010162 | 0.01459518 | 1.52208063 | up |
| KI - WT, cell_origin: iPSC | Q99747 | NAPG | 4 | Homo sapien | 2.48740016 | 1.47187052 | 3.5029298 | -1.0312734 | 5.21943576 | 0.0001029 | 0.01459518 | 1.50970354 | up |
| KI - WT, cell_origin: iPSC | P11717 | IGF2R | 66 | Homo sapien | 3.16489182 | 1.85714832 | 4.47263533 | -0.6230574 | 5.15749322 | 0.00011619 | 0.01459518 | 1.39376168 | up |
| KI - WT, cell_origin: iPSC | P30825 | SLC7A1 | 5 | Homo sapien | 7.98091302 | 4.68080795 | 11.2810181 | -4.1124982 | 5.15379546 | 0.00011701 | 0.01459518 | 1.38695629 | up |
| KI - WT, cell_origin: iPSC | P51149 | RAB7A | 14 | Homo sapien | 3.25585407 | 1.90943608 | 4.60227207 | 1.79797331 | 5.15332332 | 0.00011712 | 0.01459518 | 1.38608723 | up |
| KI - WT, cell_origin: iPSC | Q00610 | CLTC | 51 | Homo sapien | 1.9630126 | 1.15023474 | 2.77579045 | 1.50240529 | 5.14480745 | 0.00011723 | 0.01459518 | 1.38282705 | up |
| KI - WT, cell_origin: iPSC | Q92542 | NCSTN | 7 | Homo sapien | 4.27765885 | 2.50320602 | 6.05211168 | -0.7962641 | 5.13740797 | 0.00012076 | 0.01459518 | 1.35677483 | up |
| KI - WT, cell_origin: iPSC | P62820 | RAB1A | 5 | Homo sapien | 2.71052395 | 1.5654205 | 3.85562739 | 0.33548321 | 5.044416 | 0.00014448 | 0.01688062 | 1.18484332 | up |
| KI - WT, cell_origin: iPSC | Q92581 | SLC9A6 | 2 | Homo sapien | 2.07480225 | 1.19339475 | 2.95620975 | -1.9347574 | 5.01348418 | 0.00015024 | 0.01698692 | 1.14406805 | up |
| KI - WT, cell_origin: iPSC | Q9Y6M5 | SLC30A1 | 9 | Homo sapien | 5.04510444 | 2.87813466 | 7.21207422 | -2.5475839 | 4.96157629 | 0.00016967 | 0.01849121 | 1.03075135 | up |
| KI - WT, cell_origin: iPSC | Q9Y624 | F11R | 11 | Homo sapien | 5.15400269 | 2.92863163 | 7.37937375 | -1.238819 | 4.93565248 | 0.00017845 | 0.01849121 | 0.98235423 | up |
| KI - WT, cell_origin: iPSC | Q8TB96 | ITFG1 | 2 | Homo sapien | 2.87573854 | 1.63176619 | 4.11971088 | -2.7284816 | 4.92389289 | 0.00017937 | 0.01849121 | 0.97448705 | up |
| KI - WT, cell_origin: iPSC | Q9BV40 | VAMP8 | 5 | Homo sapien | 3.36308142 | 1.89362916 | 4.83253368 | -0.9515742 | 4.8773539 | 0.00019995 | 0.01974212 | 0.87321782 | up |
| KI - WT, cell_origin: iPSC | Q7Z3C6 | ATG9A | 6 | Homo sapien | 5.74205695 | 3.22821011 | 8.25590379 | -3.9202197 | 4.8677812 | 0.00020373 | 0.01974212 | 0.85525848 | up |
| KI - WT, cell_origin: iPSC | Q99805 | TM9SF2 | 5 | Homo sapien | 3.06923733 | 1.72031592 | 4.41815875 | -0.8935382 | 4.84893307 | 0.00021138 | 0.01974212 | 0.81986578 | up |
| KI - WT, cell_origin: iPSC | P14923 | JUP | 24 | Homo sapien | 3.11792848 | 1.74210747 | 4.49374948 | -0.7732093 | 4.82954904 | 0.00021956 | 0.01974212 | 0.78342319 | up |
| KI - WT, cell_origin: iPSC | Q9H0U4 | RAB1B | 10 | Homo sapien | 2.72301928 | 1.52139297 | 3.92464559 | 3.13811383 | 4.82929411 | 0.00021967 | 0.01974212 | 0.78294364 | up |
| KI - WT, cell_origin: iPSC | O15127 | SCAMP2 | 3 | Homo sapien | 5.07757176 | 2.76795873 | 7.38718479 | -1.0361046 | 4.68510379 | 0.00029176 | 0.02556519 | 0.51051358 | up |
| KI - WT, cell_origin: iPSC | P08962 | CD63 | 2 | Homo sapien | 4.08715347 | 2.20159 | 5.97271694 | -0.1367122 | 4.6193637 | 0.00033231 | 0.02835294 | 0.38555307 | up |
| KI - WT, cell_origin: iPSC | P61020 | RAB5B | 6 | Homo sapien | 1.85249226 | 0.99242057 | 2.71256394 | 1.92469856 | 4.58885666 | 0.00035029 | 0.02835294 | 0.33350262 | up |
| KI - WT, cell_origin: iPSC | P20339 | RAB5A | 3 | Homo sapien | 1.98520857 | 1.06319026 | 2.90722688 | 0.53751458 | 4.58848128 | 0.00035331 | 0.02835294 | 0.32669633 | up |
| KI - WT, cell_origin: iPSC | P51151 | RAB9A | 6 | Homo sapien | 3.07946257 | 1.64806676 | 4.51085839 | -2.3235647 | 4.58477001 | 0.00035593 | 0.02835294 | 0.31961675 | up |
| KI - WT, cell_origin: iPSC | P61026 | RAB10 | 9 | Homo sapien | 2.32698625 | 1.23947214 | 3.41450036 | 1.63853118 | 4.55996264 | 0.00037392 | 0.02912436 | 0.27225931 | up |

| | | | | | | | | | | | | | |
|---|---|---|---|---|---|---|---|---|---|---|---|---|---|
| KI - WT, cell_origin: iPSC | Q01650 | SLC7A5 | 4 | Homo sapien | 9.43843711 | 5.01512184 | 13.8617524 | -4.6167098 | 4.54730998 | 0.00038346 | 0.02921767 | 0.24808189 | up |
| KI - WT, cell_origin: iPSC | Q01628 | IFITM3 | 2 | Homo sapien | 3.41942066 | 1.80072982 | 5.03811151 | -0.1923599 | 4.50184969 | 0.00041982 | 0.03130764 | 0.1610862 | up |
| KI - WT, cell_origin: iPSC | Q15904 | ATP6AP1 | 5 | Homo sapien | 1.67627494 | 0.86482361 | 2.48772628 | -0.8563719 | 4.40013974 | 0.00050782 | 0.03708117 | -0.024247 | up |
| KI - WT, cell_origin: iPSC | P61764 | STXBP1 | 7 | Homo sapien | 1.95630496 | 0.98015567 | 2.93245424 | -1.5174616 | 4.26900942 | 0.00066244 | 0.04738497 | -0.2791174 | up |
| KI - WT, cell_origin: iPSC | P21281 | ATP6V1B2 | 13 | Homo sapien | 1.67915732 | 0.8383058 | 2.52000883 | 0.5001085 | 4.25428817 | 0.00068429 | 0.04796902 | -0.3097023 | up |
| KI - WT, cell_origin: iPSC | Q13277 | STX3 | 4 | Homo sapien | 4.89074 | 2.38551292 | 7.39596708 | -1.9239704 | 4.16034979 | 0.0008342 | 0.05509908 | -0.4980674 | up |
| KI - WT, cell_origin: iPSC | P61421 | ATP6V0D1 | 7 | Homo sapien | 4.17576304 | 2.03261735 | 6.31890874 | 0.11669315 | 4.15227892 | 0.00084794 | 0.05509908 | -0.5137486 | up |
| KI - WT, cell_origin: iPSC | Q92520 | FAM3C | 2 | Homo sapien | 2.65120051 | 1.2898051 | 4.01259593 | -1.9426539 | 4.15012005 | 0.00085166 | 0.05509908 | -0.5179439 | up |
| KI - WT, cell_origin: iPSC | Q13740 | ALCAM | 3 | Homo sapien | 2.29631872 | 1.11691649 | 3.47572096 | -1.0038621 | 4.14927851 | 0.00085311 | 0.05509908 | -0.5195793 | up |
| KI - WT, cell_origin: iPSC | Q9UHN6 | CEMIP2 | 23 | Homo sapien | 4.74189048 | 2.30254558 | 7.18123538 | -3.3413795 | 4.14267323 | 0.00086461 | 0.05509908 | -0.5324172 | up |
| KI - WT, cell_origin: iPSC | Q14344 | GNA13 | 10 | Homo sapien | 2.73544764 | 1.32192683 | 4.14896845 | 1.2445164 | 4.12409411 | 0.00089779 | 0.05619198 | -0.568541 | up |
| KI - WT, cell_origin: iPSC | P36543 | ATP6V1E1 | 2 | Homo sapien | 1.24306333 | 0.59571764 | 1.89040903 | -0.102722 | 4.08974916 | 0.0009498 | 0.05840459 | -0.6255187 | up |
| KI - WT, cell_origin: iPSC | Q9NRW7 | VPS45 | 15 | Homo sapien | 5.26717664 | 2.48844986 | 8.04590343 | -2.4415211 | 4.03956317 | 0.00106591 | 0.06410654 | -0.7331381 | up |
| KI - WT, cell_origin: iPSC | Q9UBG0 | MRC2 | 5 | Homo sapien | 3.25984613 | 1.53751673 | 4.98217553 | -2.4681862 | 4.03351109 | 0.00107911 | 0.06410654 | -0.7449366 | up |
| KI - WT, cell_origin: iPSC | O14662 | STX16 | 8 | Homo sapien | 6.77145325 | 3.13047046 | 10.412436 | -4.1494531 | 3.9633792 | 0.00124475 | 0.07271392 | -0.8817784 | up |
| KI - WT, cell_origin: iPSC | Q96FQ6 | S100A16 | 2 | Homo sapien | 1.99676339 | 0.90253701 | 3.09098977 | -2.8886821 | 3.88650388 | 0.00143896 | 0.08260524 | -1.0234734 | up |
| KI - WT, cell_origin: iPSC | P04216 | THY1 | 3 | Homo sapien | 5.32435613 | 2.39492817 | 8.25378409 | -1.603264 | 3.87334911 | 0.00149579 | 0.08260524 | -1.0577182 | up |
| KI - WT, cell_origin: iPSC | Q13443 | ADAM9 | 5 | Homo sapien | 1.86111583 | 0.83703008 | 2.88520157 | -1.5544053 | 3.87128358 | 0.00148978 | 0.08260524 | -1.0558466 | up |
| KI - WT, cell_origin: iPSC | P09496 | CLTA | 6 | Homo sapien | 1.35749548 | 0.60922036 | 2.10577059 | -0.1815021 | 3.86401448 | 0.00150834 | 0.08260524 | -1.0682984 | up |
| KI - WT, cell_origin: iPSC | P27824 | CANX | 15 | Homo sapien | 2.02167613 | 0.89815529 | 3.14519696 | 0.62112254 | 3.8347126 | 0.00161871 | 0.08728568 | -1.1332953 | up |
| KI - WT, cell_origin: iPSC | P13987 | CD59 | 2 | Homo sapien | 2.42390242 | 1.06855802 | 3.77924682 | -1.0973158 | 3.8112535 | 0.00169827 | 0.08954509 | -1.1792 | up |
| KI - WT, cell_origin: iPSC | Q9HCE6 | ARHGEF10L | 2 | Homo sapien | -7.8855915 | -12.299342 | -3.4718413 | -4.9517934 | -3.8074034 | 0.0017117 | 0.08954509 | -1.1867349 | down |
| KI - WT, cell_origin: iPSC | P35613 | BSG | 9 | Homo sapien | 5.35247804 | 2.34393333 | 8.36102276 | 0.8057828 | 3.79141035 | 0.00176865 | 0.09076973 | -1.2180368 | up |
| KI - WT, cell_origin: iPSC | Q86Y82 | STX12 | 7 | Homo sapien | 2.91712985 | 1.27528403 | 4.55897568 | 0.21369819 | 3.7863936 | 0.00178691 | 0.09076973 | -1.2278565 | up |
| KI - WT, cell_origin: iPSC | O43747 | AP1G1 | 9 | Homo sapien | 3.44177316 | 1.47147463 | 5.41207168 | -2.8002127 | 3.72265319 | 0.00203611 | 0.10195075 | -1.3526467 | up |
| KI - WT, cell_origin: iPSC | P08754 | GNAI3 | 3 | Homo sapien | 3.67193942 | 1.53393208 | 5.80994675 | -0.052453 | 3.66006404 | 0.0023149 | 0.11427777 | -1.4752027 | up |
| KI - WT, cell_origin: iPSC | P38606 | ATP6V1A | 20 | Homo sapien | 1.2684811 | 0.52256847 | 2.01439374 | 1.18212431 | 3.62236087 | 0.00248146 | 0.12041928 | -1.5435906 | up |
| KI - WT, cell_origin: iPSC | P52306 | RAP1GDS1 | 3 | Homo sapien | 1.90329499 | 0.78231138 | 3.02427861 | -2.6651747 | 3.61615072 | 0.00250802 | 0.12041928 | -1.5543091 | up |
| KI - WT, cell_origin: iPSC | P42212 | GFP | 8 | Aequorea vic | 6.58164008 | 2.69481655 | 10.4684636 | 2.31279295 | 3.60862433 | 0.00257258 | 0.12185011 | -1.5759144 | up |
| KI - WT, cell_origin: iPSC | P11279 | LAMP1 | 6 | Homo sapien | 2.12422281 | 0.85619321 | 3.39225241 | -0.2775563 | 3.57003969 | 0.00278463 | 0.13013502 | -1.6514357 | up |
| KI - WT, cell_origin: iPSC | Q12846 | STX4 | 8 | Homo sapien | 2.54486322 | 1.01336518 | 4.07636125 | -0.6243427 | 3.54119929 | 0.00295451 | 0.13601414 | -1.707866 | up |
| KI - WT, cell_origin: iPSC | Q5VW38 | GPR107 | 2 | Homo sapien | 3.14346419 | 1.24851622 | 5.03841217 | -3.4995628 | 3.5347543 | 0.00298804 | 0.13601414 | -1.7191514 | up |
| KI - WT, cell_origin: iPSC | O15427 | SLC16A3 | 4 | Homo sapien | 5.09050981 | 1.9208152 | 8.26020443 | -3.1905698 | 3.42252181 | 0.00376998 | 0.16940729 | -1.9398108 | up |
| KI - WT, cell_origin: iPSC | Q5TC84 | OGFRL1 | 7 | Homo sapien | 3.37100312 | 1.24985932 | 5.49214691 | -2.3687517 | 3.38681451 | 0.00405682 | 0.17982206 | -2.0094836 | up |
| KI - WT, cell_origin: iPSC | P80303 | NUCB2 | 7 | Homo sapien | -3.0363509 | -4.9501259 | -1.1225759 | -1.7900725 | -3.3811423 | 0.00410436 | 0.17982206 | -2.0205453 | down |
| KI - WT, cell_origin: iPSC | Q15131 | CDK10 | 2 | Homo sapien | -1.7124479 | -2.8004591 | -0.6244367 | -2.3396605 | -3.3521519 | 0.00432026 | 0.1869448 | -2.0716451 | down |
| KI - WT, cell_origin: iPSC | Q9UKS6 | PACSIN3 | 6 | Homo sapien | 3.64920269 | 1.27762979 | 6.02077558 | -2.6357333 | 3.27916958 | 0.00505999 | 0.2162837 | -2.2190735 | up |
| KI - WT, cell_origin: iPSC | Q10589 | BST2 | 4 | Homo sapien | 5.3351238 | 1.83293922 | 8.83730838 | -2.4825523 | 3.24644342 | 0.0054113 | 0.22700942 | -2.282632 | up |
| KI - WT, cell_origin: iPSC | Q9GZT9 | EGLN1 | 5 | Homo sapien | 1.82579081 | 0.62455421 | 3.0270274 | 0.39295667 | 3.23910354 | 0.00549336 | 0.22700942 | -2.2968752 | up |
| KI - WT, cell_origin: iPSC | P20020 | ATP2B1 | 18 | Homo sapien | 3.041785 | 1.03986324 | 5.04370677 | 1.18256161 | 3.23805172 | 0.00550522 | 0.22700942 | -2.2989159 | up |
| KI - WT, cell_origin: iPSC | P20338 | RAB4A | 5 | Homo sapien | 2.83908298 | 0.95562148 | 4.72254448 | -1.5187815 | 3.21235628 | 0.00580297 | 0.23650495 | -2.3487401 | up |
| KI - WT, cell_origin: iPSC | Q53GQ0 | HSD17B12 | 3 | Homo sapien | 1.31213374 | 0.43593369 | 2.18833379 | -1.1927198 | 3.18985179 | 0.00604215 | 0.2434223 | -2.388721 | up |
| KI - WT, cell_origin: iPSC | P42892 | ECE1 | 5 | Homo sapien | 0.96858903 | 0.31562818 | 1.62154989 | 0.40548677 | 3.15930884 | 0.00642332 | 0.25583777 | -2.4469696 | up |
| KI - WT, cell_origin: iPSC | Q9UHW9 | SLC12A6 | 4 | Homo sapien | 3.12122781 | 1.00788578 | 5.23456984 | -3.4776094 | 3.14744424 | 0.00662811 | 0.26102835 | -2.4743347 | up |
| KI - WT, cell_origin: iPSC | O75167 | PHACTR2 | 9 | Homo sapien | 1.85659709 | 0.59394099 | 3.11925319 | -1.4003175 | 3.1335379 | 0.00681944 | 0.26539365 | -2.5011863 | up |
| KI - WT, cell_origin: iPSC | Q8IVL0 | NAV3 | 3 | Homo sapien | 2.75155287 | 0.87721858 | 4.62588716 | -3.2568478 | 3.12847853 | 0.00689039 | 0.26539365 | -2.5109503 | up |

| | | | | | | | | | | | | |
|---|---|---|---|---|---|---|---|---|---|---|---|---|
| KI - WT, cell_origin: iPSC | Q8N3R9 | PALS1 | 6 | Homo sapien | 2.17677552 | 0.68632567 | 3.66722536 | -2.7888068 | 3.11156398 | 0.00711078 | 0.27090526 | -2.541652 up |
| KI - WT, cell_origin: iPSC | Q9UBI6 | GNG12 | 2 | Homo sapien | 1.21591812 | 0.37274293 | 2.05909332 | 0.42757736 | 3.07209527 | 0.00770228 | 0.28598016 | -2.6172046 up |
| KI - WT, cell_origin: iPSC | O15126 | SCAMP1 | 8 | Homo sapien | 4.12862081 | 1.26377977 | 6.99346184 | -1.4086552 | 3.07119107 | 0.00774645 | 0.28598016 | -2.6213078 up |
| KI - WT, cell_origin: iPSC | P04439 | HLA-A | 7 | Homo sapien | 2.70377773 | 0.82410308 | 4.58345237 | -1.6499017 | 3.06542483 | 0.00783822 | 0.28598016 | -2.6323944 up |
| KI - WT, cell_origin: iPSC | O75923 | DYSF | 35 | Homo sapien | 1.99954716 | 0.60771082 | 3.3913835 | -1.9080385 | 3.06158166 | 0.00789997 | 0.28598016 | -2.6397813 up |
| KI - WT, cell_origin: iPSC | P41218 | MNDA | 7 | Homo sapien | -2.5596927 | -4.3442141 | -0.7751713 | -0.1469445 | -3.0568099 | 0.00797732 | 0.28598016 | -2.6489504 down |
| KI - WT, cell_origin: iPSC | Q92544 | TM9SF4 | 2 | Homo sapien | 2.34771209 | 0.71036115 | 3.98506303 | -1.2048652 | 3.05566282 | 0.00799602 | 0.28598016 | -2.6511542 up |
| KI - WT, cell_origin: iPSC | O00592 | PODXL | 10 | Homo sapien | 3.85869046 | 1.15922104 | 6.55815988 | -0.1771496 | 3.04623846 | 0.00815133 | 0.28859017 | -2.6692537 up |
| KI - WT, cell_origin: iPSC | P54709 | ATP1B3 | 12 | Homo sapien | 7.96211875 | 2.36241172 | 13.5618258 | -3.0428692 | 3.03016106 | 0.00842317 | 0.29292064 | -2.7001042 up |
| KI - WT, cell_origin: iPSC | O43760 | SYNGR2 | 2 | Homo sapien | 4.8103043 | 1.42610368 | 8.19450492 | -1.1721511 | 3.02913631 | 0.00844079 | 0.29292064 | -2.7020694 up |
| KI - WT, cell_origin: iPSC | P18084 | ITGB5 | 7 | Homo sapien | 3.03490018 | 0.88952655 | 5.18027382 | -1.8445177 | 3.0146982 | 0.00869302 | 0.29871589 | -2.7297433 up |
| KI - WT, cell_origin: iPSC | O75165 | DNAJC13 | 68 | Homo sapien | 3.88820314 | 1.12802961 | 6.64837667 | -1.0402106 | 3.00202931 | 0.00892042 | 0.30355414 | -2.7540028 up |
| KI - WT, cell_origin: iPSC | Q9ULF5 | SLC39A10 | 6 | Homo sapien | 2.67283671 | 0.76907173 | 4.5766017 | -2.648585 | 2.99199912 | 0.0091046 | 0.30553715 | -2.7731938 up |
| KI - WT, cell_origin: iPSC | Q8WY07 | SLC7A3 | 6 | Homo sapien | 4.73816686 | 1.36040098 | 8.11593274 | -1.8440357 | 2.98939411 | 0.00915304 | 0.30553715 | -2.7781757 up |
| KI - WT, cell_origin: iPSC | P17612 | PRKACA | 2 | Homo sapien | 1.38176154 | 0.39262773 | 2.37089534 | -0.6861429 | 2.97701088 | 0.00938679 | 0.3097246 | -2.8018445 up |
| KI - WT, cell_origin: iPSC | O00461 | GOLIM4 | 3 | Homo sapien | 2.5197426 | 0.71049544 | 4.32898975 | -4.4843289 | 2.96618314 | 0.00953836 | 0.3097246 | -2.8189602 up |
| KI - WT, cell_origin: iPSC | O43617 | TRAPPC3 | 4 | Homo sapien | 0.84204786 | 0.23737879 | 1.44671693 | 0.38170953 | 2.96591501 | 0.00954358 | 0.3097246 | -2.8194732 up |
| KI - WT, cell_origin: iPSC | Q9Y6M7 | SLC4A7 | 10 | Homo sapien | 1.33858512 | 0.37292296 | 2.30424728 | -1.2425347 | 2.95283163 | 0.00981912 | 0.31574327 | -2.8455359 up |
| KI - WT, cell_origin: iPSC | P10301 | RRAS | 6 | Homo sapien | 1.46135717 | 0.39930209 | 2.52341224 | -1.1184088 | 2.93232229 | 0.01028005 | 0.32685781 | -2.8870723 up |
| KI - WT, cell_origin: iPSC | Q6NY19 | KANK3 | 12 | Homo sapien | 2.61143667 | 0.71134897 | 4.51152437 | -0.8339396 | 2.92892479 | 0.01035127 | 0.32685781 | -2.8935394 up |
| KI - WT, cell_origin: iPSC | Q9BZL4 | PPP1R12C | 8 | Homo sapien | -2.8592418 | -4.9523434 | -0.7661402 | -0.6123866 | -2.9111391 | 0.01073206 | 0.33585584 | -2.9273642 down |
| KI - WT, cell_origin: iPSC | Q14203 | DCTN1 | 5 | Homo sapien | 0.8902467 | 0.23425355 | 1.54623985 | -2.192842 | 2.89035209 | 0.0111317 | 0.34527965 | -2.9635886 up |
| KI - WT, cell_origin: iPSC | Q02818 | NUCB1 | 4 | Homo sapien | 3.84744278 | 1.00251075 | 6.6923748 | -2.6317816 | 2.88205751 | 0.01138437 | 0.34925904 | -2.9825603 up |
| KI - WT, cell_origin: iPSC | Q8TDZ2 | MICAL1 | 6 | Homo sapien | 1.41545971 | 0.36753633 | 2.4633831 | -0.7020113 | 2.87852727 | 0.01146614 | 0.34925904 | -2.989251 up |
| KI - WT, cell_origin: iPSC | P06756 | ITGAV | 24 | Homo sapien | 1.71467005 | 0.44347299 | 2.98586711 | -1.6076655 | 2.87455121 | 0.01155893 | 0.34925904 | -2.9967841 up |
| KI - WT, cell_origin: iPSC | P20645 | M6PR | 7 | Homo sapien | 2.2622534 | 0.58144284 | 3.94306395 | 0.31404447 | 2.86830183 | 0.01170625 | 0.35068728 | -3.0086188 up |
| KI - WT, cell_origin: iPSC | Q16537 | PPP2R5E | 2 | Homo sapien | 1.25768778 | 0.31615591 | 2.19921965 | -1.5046729 | 2.84533557 | 0.01221164 | 0.36272717 | -3.0496447 up |
| KI - WT, cell_origin: iPSC | Q9UPQ0 | LIMCH1 | 19 | Homo sapien | 2.91357456 | 0.70788279 | 5.11926632 | 0.32229266 | 2.81503424 | 0.01303854 | 0.38403429 | -3.1092119 up |
| KI - WT, cell_origin: iPSC | O75348 | ATP6V1G1 | 3 | Homo sapien | 1.36035481 | 0.31573247 | 2.40497714 | 0.11048717 | 2.77520619 | 0.01412996 | 0.41271265 | -3.1840793 up |
| KI - WT, cell_origin: iPSC | Q07954 | LRP1 | 66 | Homo sapien | 1.6054993 | 0.37043559 | 2.84056301 | 0.43410895 | 2.77027656 | 0.01427107 | 0.4133893 | -3.1933243 up |
| KI - WT, cell_origin: iPSC | P09601 | HMOX1 | 2 | Homo sapien | 1.2538363 | 0.28768128 | 2.21999132 | -1.0030651 | 2.7650484 | 0.01439659 | 0.41360683 | -3.2021448 up |
| KI - WT, cell_origin: iPSC | P33527 | ABCC1 | 24 | Homo sapien | 3.46623197 | 0.78519002 | 6.14727392 | -3.8913436 | 2.75522064 | 0.01471056 | 0.41919114 | -3.2215297 up |
| KI - WT, cell_origin: iPSC | P50148 | GNAQ | 9 | Homo sapien | 1.41734233 | 0.31625639 | 2.51842828 | 0.59508272 | 2.74319012 | 0.01507114 | 0.42325205 | -3.2440341 up |
| KI - WT, cell_origin: iPSC | P61225 | RAP2B | 6 | Homo sapien | 5.28527492 | 1.17816484 | 9.392385 | -0.8794123 | 2.74241772 | 0.01509458 | 0.42325205 | -3.2454779 up |
| KI - WT, cell_origin: iPSC | P13796 | LCP1 | 25 | Homo sapien | -4.2102929 | -7.5193247 | -0.9012611 | -0.8702006 | -2.7115258 | 0.01606159 | 0.44679252 | -3.3031205 down |
| KI - WT, cell_origin: iPSC | P04839 | CYBB | 3 | Homo sapien | -2.4927564 | -4.4650683 | -0.5204445 | -1.9145766 | -2.6934356 | 0.0166554 | 0.45649323 | -3.33678 down |
| KI - WT, cell_origin: iPSC | O15040 | TECPR2 | 2 | Homo sapien | 0.89691046 | 0.18695697 | 1.60686394 | -1.258725 | 2.6906603 | 0.01667079 | 0.45649323 | -3.3394616 up |
| KI - WT, cell_origin: iPSC | P13473 | LAMP2 | 2 | Homo sapien | 2.6165739 | 0.53902181 | 4.694126 | -1.3760202 | 2.6840058 | 0.01697329 | 0.4570514 | -3.3542969 up |
| KI - WT, cell_origin: iPSC | Q9BTU6 | PI4K2A | 12 | Homo sapien | 2.92545863 | 0.59879756 | 5.25211969 | -2.2531717 | 2.67955827 | 0.01712524 | 0.4570514 | -3.3625517 up |
| KI - WT, cell_origin: iPSC | Q9P2N2 | ARHGAP28 | 6 | Homo sapien | 2.74432148 | 0.56037434 | 4.92826862 | -0.6919631 | 2.67790539 | 0.01718205 | 0.4570514 | -3.3656184 up |
| KI - WT, cell_origin: iPSC | P07332 | FES | 5 | Homo sapien | -0.976688 | -1.7543704 | -0.1990055 | -0.5893001 | -2.675004 | 0.01721278 | 0.4570514 | -3.3688677 down |
| KI - WT, cell_origin: iPSC | Q9UHD9 | UBQLN2 | 5 | Homo sapien | 1.33610501 | 0.26934925 | 2.40286077 | -1.4727681 | 2.66844375 | 0.01747454 | 0.45767193 | -3.382068 up |
| KI - WT, cell_origin: iPSC | Q9C0B0 | UNK | 10 | Homo sapien | 0.83915223 | 0.16891537 | 1.50938909 | -0.8628087 | 2.66656472 | 0.0174973 | 0.45767193 | -3.384252 up |
| KI - WT, cell_origin: iPSC | P08648 | ITGA5 | 2 | Homo sapien | 2.92623659 | 0.5859639 | 5.26650927 | -3.048906 | 2.66468171 | 0.01764312 | 0.45806767 | -3.3901305 up |
| KI - WT, cell_origin: iPSC | P48431 | SOX2 | 2 | Homo sapien | -0.9573259 | -1.7250355 | -0.1896163 | 1.29385967 | -2.6562542 | 0.0178834 | 0.46089216 | -3.4039791 down |
| KI - WT, cell_origin: iPSC | P25686 | DNAJB2 | 4 | Homo sapien | -1.6985128 | -3.0665413 | -0.3304843 | -1.7305625 | -2.645916 | 0.01831795 | 0.46670104 | -3.4248463 down |

| | | | | | | | | | | | | |
|---|---|---|---|---|---|---|---|---|---|---|---|---|
| KI - WT, cell_origin: iPSC | Q12959 | DLG1 | 14 | Homo sapien | 1.398686 | 0.27148221 | 2.52588979 | -0.4976748 | 2.64435743 | 0.01837511 | 0.46670104 | -3.4277257 up |
| KI - WT, cell_origin: iPSC | P11169 | SLC2A3 | 6 | Homo sapien | 4.6969096 | 0.89633497 | 8.49748422 | -0.6018784 | 2.63369394 | 0.01877077 | 0.47332041 | -3.4474117 up |
| KI - WT, cell_origin: iPSC | Q86WG5 | SBF2 | 5 | Homo sapien | 1.41031012 | 0.26540466 | 2.55521557 | -1.3337564 | 2.62510915 | 0.01909524 | 0.47806308 | -3.4632404 up |
| KI - WT, cell_origin: iPSC | Q16584 | MAP3K11 | 3 | Homo sapien | 1.46104173 | 0.27298226 | 2.6491012 | -0.4968341 | 2.62075733 | 0.01926178 | 0.47881241 | -3.4712574 up |
| KI - WT, cell_origin: iPSC | O43493 | TGOLN2 | 4 | Homo sapien | 1.90036971 | 0.35109005 | 3.44964936 | -1.4688914 | 2.61403049 | 0.01952196 | 0.48186252 | -3.4836407 up |
| KI - WT, cell_origin: iPSC | Q8WVT3 | TRAPPC12 | 2 | Homo sapien | 2.36475291 | 0.43328829 | 4.29621753 | -3.8757443 | 2.60805602 | 0.01969682 | 0.48277864 | -3.4930929 up |
| KI - WT, cell_origin: iPSC | Q9H0E2 | TOLLIP | 3 | Homo sapien | 1.20461863 | 0.21887827 | 2.19035898 | -1.3947876 | 2.60318445 | 0.01988926 | 0.48411018 | -3.5020581 up |
| KI - WT, cell_origin: iPSC | O43169 | CYB5B | 2 | Homo sapien | 1.81648003 | 0.32719349 | 3.30576657 | -1.7607803 | 2.59928998 | 0.02010395 | 0.48596108 | -3.5107371 up |
| KI - WT, cell_origin: iPSC | Q15392 | DHCR24 | 3 | Homo sapien | 1.28354077 | 0.22858873 | 2.33849281 | -0.8052846 | 2.5928615 | 0.02036295 | 0.48662787 | -3.522537 up |
| KI - WT, cell_origin: iPSC | P21926 | CD9 | 3 | Homo sapien | 1.14118566 | 0.20280029 | 2.07957103 | 0.10643977 | 2.59146639 | 0.02040921 | 0.48662787 | -3.524838 up |
| KI - WT, cell_origin: iPSC | O00560 | SDCBP | 3 | Homo sapien | 1.52154243 | 0.26838354 | 2.77470131 | -1.2437952 | 2.58749963 | 0.02058142 | 0.48741812 | -3.5323712 up |
| KI - WT, cell_origin: iPSC | P12931 | SRC | 9 | Homo sapien | 1.11227933 | 0.18313314 | 2.04142551 | -0.5761644 | 2.55071482 | 0.02211824 | 0.52029811 | -3.5991032 up |
| KI - WT, cell_origin: iPSC | P56377 | AP1S2 | 3 | Homo sapien | 1.39206155 | 0.22496824 | 2.55915486 | -1.7039377 | 2.54151025 | 0.02252826 | 0.52383799 | -3.6159392 up |
| KI - WT, cell_origin: iPSC | Q16864 | ATP6V1F | 3 | Homo sapien | 0.92815585 | 0.14965015 | 1.70666155 | -0.0575071 | 2.5395384 | 0.02256763 | 0.52383799 | -3.6184589 up |
| KI - WT, cell_origin: iPSC | P04626 | ERBB2 | 12 | Homo sapien | 2.50872389 | 0.39422649 | 4.62322128 | -2.1648651 | 2.52841337 | 0.02314258 | 0.53056882 | -3.6402454 up |
| KI - WT, cell_origin: iPSC | Q14118 | DAG1 | 7 | Homo sapien | 2.55228932 | 0.39864486 | 4.70593378 | -0.4829128 | 2.52556333 | 0.02327353 | 0.53056882 | -3.6454251 up |
| KI - WT, cell_origin: iPSC | P41217 | CD200 | 5 | Homo sapien | 2.15703327 | 0.33631213 | 3.97775441 | -1.7692549 | 2.52473486 | 0.02331173 | 0.53056882 | -3.6469304 up |
| KI - WT, cell_origin: iPSC | O00192 | ARVCF | 11 | Homo sapien | 2.50489035 | 0.38673816 | 4.62304253 | -2.6895438 | 2.52019373 | 0.02352216 | 0.53129587 | -3.655178 up |
| KI - WT, cell_origin: iPSC | Q9ULL8 | SHROOM4 | 4 | Homo sapien | 2.49948687 | 0.38366054 | 4.61531321 | -2.1685192 | 2.51752163 | 0.02364683 | 0.53129587 | -3.6600284 up |
| KI - WT, cell_origin: iPSC | Q8IXV7 | KLHDC8B | 4 | Homo sapien | -2.541626 | -4.7038682 | -0.3793838 | 4.4512288 | -2.5050112 | 0.02423897 | 0.53214908 | -3.682711 down |
| KI - WT, cell_origin: iPSC | P39656 | DDOST | 5 | Homo sapien | 1.41179683 | 0.21051947 | 2.61307419 | -1.080703 | 2.50455907 | 0.02426063 | 0.53214908 | -3.6835299 up |
| KI - WT, cell_origin: iPSC | Q9BQE5 | APOL2 | 3 | Homo sapien | 1.56781849 | 0.2343391 | 2.90129789 | -2.5975618 | 2.50453657 | 0.02419677 | 0.53214908 | -3.6822676 up |
| KI - WT, cell_origin: iPSC | Q96AJ9 | VTI1A | 2 | Homo sapien | 2.14863586 | 0.31991369 | 3.97735804 | -2.2747637 | 2.50390275 | 0.02429211 | 0.53214908 | -3.6847186 up |
| KI - WT, cell_origin: iPSC | Q93050 | ATP6V0A1 | 12 | Homo sapien | 2.74094056 | 0.40211888 | 5.07976223 | -1.3543223 | 2.49749642 | 0.02460143 | 0.53557768 | -3.6963149 up |
| KI - WT, cell_origin: iPSC | Q13561 | DCTN2 | 9 | Homo sapien | 0.84437696 | 0.12246178 | 1.56629214 | -0.996878 | 2.49109267 | 0.02482081 | 0.53701807 | -3.7060794 up |
| KI - WT, cell_origin: iPSC | P54578 | USP14 | 2 | Homo sapien | -1.3516523 | -2.5123173 | -0.1909873 | -2.8706628 | -2.4802632 | 0.02535779 | 0.54527026 | -3.7256687 down |
| KI - WT, cell_origin: iPSC | P61769 | B2M | 2 | Homo sapien | 1.35923299 | 0.18188526 | 2.53658072 | -0.6244108 | 2.46031959 | 0.02647183 | 0.56575472 | -3.763378 up |
| KI - WT, cell_origin: iPSC | P10071 | GLI3 | 3 | Homo sapien | 1.76228252 | 0.22241783 | 3.3021472 | -2.9243481 | 2.43823427 | 0.02760151 | 0.58632291 | -3.8022693 up |
| KI - WT, cell_origin: iPSC | O75995 | SASH3 | 3 | Homo sapien | 2.7845956 | 0.32574324 | 5.24344795 | -3.7958551 | 2.41341523 | 0.02902489 | 0.60800811 | -3.8474047 up |
| KI - WT, cell_origin: iPSC | Q13557 | CAMK2D | 14 | Homo sapien | -0.8582607 | -1.6166314 | -0.0998901 | 0.73989208 | -2.4107075 | 0.02910463 | 0.60800811 | -3.8510593 down |
| KI - WT, cell_origin: iPSC | O00186 | STXBP3 | 14 | Homo sapien | 4.20762951 | 0.4877947 | 7.92746432 | -3.6702002 | 2.41055034 | 0.02918816 | 0.60800811 | -3.8525152 up |
| KI - WT, cell_origin: iPSC | P49748 | ACADVL | 3 | Homo sapien | -2.0570897 | -3.8775172 | -0.2366623 | -2.6035029 | -2.4081428 | 0.02932604 | 0.60800811 | -3.8568079 down |
| KI - WT, cell_origin: iPSC | O94763 | URI1 | 2 | Homo sapien | 0.92231076 | 0.10504848 | 1.73957303 | -1.1439369 | 2.40359601 | 0.02948969 | 0.60800811 | -3.8633857 up |
| KI - WT, cell_origin: iPSC | Q6DD88 | ATL3 | 5 | Homo sapien | 2.10849396 | 0.23608979 | 3.98089812 | -2.6567265 | 2.39980045 | 0.0298086 | 0.61098917 | -3.8716684 up |
| KI - WT, cell_origin: iPSC | Q6PIU2 | NCEH1 | 8 | Homo sapien | 1.79382712 | 0.19830814 | 3.38934611 | -0.1476819 | 2.39596781 | 0.03003281 | 0.6118197 | -3.8784883 up |
| KI - WT, cell_origin: iPSC | Q9UI12 | ATP6V1H | 6 | Homo sapien | 0.84292692 | 0.09222139 | 1.59363246 | -0.443507 | 2.39147562 | 0.03019823 | 0.6118197 | -3.8849847 up |
| KI - WT, cell_origin: iPSC | P00167 | CYB5A | 2 | Homo sapien | 1.6342161 | 0.17368235 | 3.09474985 | -2.2355129 | 2.38451608 | 0.03071227 | 0.61612641 | -3.898838 up |
| KI - WT, cell_origin: iPSC | P17947 | SPI1 | 3 | Homo sapien | -1.2925837 | -2.448198 | -0.1369694 | -0.9169713 | -2.3836811 | 0.03076237 | 0.61612641 | -3.9003201 down |
| KI - WT, cell_origin: iPSC | P51114 | FXR1 | 21 | Homo sapien | -0.7274347 | -1.3801572 | -0.0747122 | 2.91426754 | -2.3735868 | 0.03127109 | 0.61930283 | -3.9167433 down |
| KI - WT, cell_origin: iPSC | P78310 | CXADR | 9 | Homo sapien | 0.7101423 | 0.07292187 | 1.34736273 | 1.11578627 | 2.37353352 | 0.03127435 | 0.61930283 | -3.9168379 up |
| KI - WT, cell_origin: iPSC | O00264 | PGRMC1 | 3 | Homo sapien | 1.83499162 | 0.18381498 | 3.48616826 | -0.6274804 | 2.36833382 | 0.03169721 | 0.62415003 | -3.9275221 up |
| KI - WT, cell_origin: iPSC | Q9UIQ6 | LNPEP | 8 | Homo sapien | 2.61767861 | 0.24059814 | 4.99475908 | -2.5059379 | 2.34679313 | 0.03305459 | 0.64724213 | -3.9655717 up |
| KI - WT, cell_origin: iPSC | Q8N128 | FAM177A1 | 6 | Homo sapien | 0.73704294 | 0.06512296 | 1.40896292 | -2.4049959 | 2.33622636 | 0.03363412 | 0.65493114 | -3.9828038 up |
| KI - WT, cell_origin: iPSC | P11216 | PYGB | 12 | Homo sapien | -0.9613615 | -1.8405985 | -0.0821245 | -0.3584517 | -2.3293545 | 0.03413346 | 0.65977938 | -3.9954997 down |
| KI - WT, cell_origin: iPSC | Q8WUI4 | HDAC7 | 3 | Homo sapien | -3.0989519 | -5.9353516 | -0.2625521 | 0.62062092 | -2.328358 | 0.03425959 | 0.65977938 | -3.998013 down |
| KI - WT, cell_origin: iPSC | P21283 | ATP6V1C1 | 6 | Homo sapien | 1.99058743 | 0.16264009 | 3.81853477 | -2.1015842 | 2.32070504 | 0.03477186 | 0.66302014 | -4.0114467 up |

| | | | | | | | | | | | | | |
|---|---|---|---|---|---|---|---|---|---|---|---|---|---|
| KI - WT, cell_origin: iPSC | P13861 | PRKAR2A | 11 | Homo sapien | 1.01618449 | 0.08278692 | 1.94958207 | -0.9253346 | 2.31946064 | 0.03480619 | 0.66302014 | -4.0130143 | up |
| KI - WT, cell_origin: iPSC | P51148 | RAB5C | 6 | Homo sapien | 1.19290495 | 0.09431627 | 2.29149363 | 2.03626011 | 2.31405186 | 0.03522304 | 0.66374608 | -4.023109 | up |
| KI - WT, cell_origin: iPSC | Q9P0K7 | RAI14 | 16 | Homo sapien | 0.98000741 | 0.07746535 | 1.88254946 | -0.0920409 | 2.3135499 | 0.03522219 | 0.66374608 | -4.0235603 | up |
| KI - WT, cell_origin: iPSC | Q6UX71 | PLXDC2 | 6 | Homo sapien | 1.41101011 | 0.10972052 | 2.71229969 | -1.6628473 | 2.31077968 | 0.03544696 | 0.66439357 | -4.0288392 | up |
| KI - WT, cell_origin: iPSC | Q9UBT2 | UBA2 | 3 | Homo sapien | -0.9848146 | -1.895452 | -0.0741773 | -1.7950236 | -2.3032906 | 0.03585566 | 0.66847921 | -4.0406483 | down |
| KI - WT, cell_origin: iPSC | Q9UMX0 | UBQLN1 | 10 | Homo sapien | 1.0013191 | 0.06770705 | 1.93493115 | 2.49764075 | 2.28564313 | 0.03721216 | 0.69009845 | -4.0727338 | up |
| KI - WT, cell_origin: iPSC | Q8IZA0 | KIAA0319L | 2 | Homo sapien | 0.94856552 | 0.05257296 | 1.84455808 | -2.0725436 | 2.2547719 | 0.03938012 | 0.72645954 | -4.1251639 | up |
| KI - WT, cell_origin: iPSC | O43166 | SIPA1L1 | 7 | Homo sapien | 0.68510581 | 0.03496395 | 1.33524767 | -1.3302779 | 2.24434287 | 0.04017883 | 0.73475475 | -4.1432187 | up |
| KI - WT, cell_origin: iPSC | O75365 | PTP4A3 | 8 | Homo sapien | 2.65497865 | 0.13054521 | 5.17941209 | -2.1822959 | 2.24129735 | 0.0405286 | 0.73475475 | -4.1496231 | up |
| KI - WT, cell_origin: iPSC | P51116 | FXR2 | 16 | Homo sapien | -0.6913574 | -1.3493489 | -0.0333659 | 2.99405756 | -2.2378038 | 0.04068731 | 0.73475475 | -4.1545187 | down |
| KI - WT, cell_origin: iPSC | P62873 | GNB1 | 7 | Homo sapien | 0.78776144 | 0.03737631 | 1.53814657 | 1.79429464 | 2.23617698 | 0.04083906 | 0.73475475 | -4.157568 | up |
| KI - WT, cell_origin: iPSC | P42694 | HELZ | 10 | Homo sapien | -1.1961345 | -2.3362782 | -0.0559908 | -0.9188905 | -2.2357477 | 0.04096255 | 0.73475475 | -4.1591948 | down |
| KI - WT, cell_origin: iPSC | Q9HCD6 | TANC2 | 4 | Homo sapien | 0.99769894 | 0.04601715 | 1.94938073 | -0.5903695 | 2.23385899 | 0.04108757 | 0.73475475 | -4.1622178 | up |
| KI - WT, cell_origin: iPSC | Q9H3N1 | TMX1 | 2 | Homo sapien | 0.94547796 | 0.04217464 | 1.84878128 | -0.9908219 | 2.22964492 | 0.04136428 | 0.73594824 | -4.1689268 | up |
| KI - WT, cell_origin: iPSC | Q99569 | PKP4 | 5 | Homo sapien | 1.64812636 | 0.0610154 | 3.23523732 | -2.9026788 | 2.21267413 | 0.04278357 | 0.75374321 | -4.1985894 | up |
| KI - WT, cell_origin: iPSC | Q9C0B2 | CFAP74 | 2 | Homo sapien | -0.9055133 | -1.7781175 | -0.032909 | 1.2279553 | -2.211106 | 0.04291117 | 0.75374321 | -4.2012704 | down |
| KI - WT, cell_origin: iPSC | P15498 | VAV1 | 13 | Homo sapien | 1.89950842 | 0.06805813 | 3.7309587 | -0.5061731 | 2.21028591 | 0.0430096 | 0.75374321 | -4.202962 | up |
| KI - WT, cell_origin: iPSC | Q6NZY7 | CDC42EP5 | 3 | Homo sapien | 2.10645318 | 0.06936307 | 4.14354329 | -2.3490163 | 2.2036567 | 0.04355798 | 0.75955581 | -4.2143169 | up |
| KI - WT, cell_origin: iPSC | Q14147 | DHX34 | 3 | Homo sapien | -1.3766067 | -2.7096452 | -0.0435681 | -1.9333896 | -2.200744 | 0.04380098 | 0.75977731 | -4.2193006 | down |
| KI - WT, cell_origin: iPSC | Q92556 | ELMO1 | 20 | Homo sapien | 0.82213021 | 0.02479209 | 1.61946833 | 0.36100684 | 2.19642301 | 0.04408154 | 0.75977731 | -4.225961 | up |
| KI - WT, cell_origin: iPSC | Q15758 | SLC1A5 | 9 | Homo sapien | 4.54207197 | 0.12858582 | 8.95555812 | -3.6166451 | 2.19318169 | 0.04443776 | 0.75977731 | -4.2322246 | up |
| KI - WT, cell_origin: iPSC | Q8TD26 | CHD6 | 2 | Homo sapien | 1.56547423 | 0.0452229 | 3.08572556 | -4.1770226 | 2.19315948 | 0.04432229 | 0.75977731 | -4.2312343 | up |
| KI - WT, cell_origin: iPSC | Q9HC98 | NEK6 | 3 | Homo sapien | -1.5585005 | -3.0761769 | -0.0408242 | -1.2353171 | -2.1884174 | 0.04484334 | 0.7629899 | -4.2403555 | down |
| KI - WT, cell_origin: iPSC | Q5VT52 | RPRD2 | 3 | Homo sapien | 2.63925203 | 0.06030393 | 5.21820014 | -4.4169796 | 2.18092449 | 0.04548817 | 0.77022239 | -4.2531252 | up |
| KI - WT, cell_origin: iPSC | Q9UNE7 | STUB1 | 5 | Homo sapien | 1.96202052 | 0.02492158 | 3.89911945 | -2.8788384 | 2.15851022 | 0.0474688 | 0.79989493 | -4.2911926 | up |
| KI - WT, cell_origin: iPSC | Q86TI0 | TBC1D1 | 3 | Homo sapien | 1.82201803 | 0.01831233 | 3.62572373 | -1.7022114 | 2.15272899 | 0.04799246 | 0.80484958 | -4.3009788 | up |
| KI - WT, cell_origin: iPSC | Q562E7 | WDR81 | 11 | Homo sapien | 0.68421192 | 0.00593013 | 1.36249372 | -1.023017 | 2.14842482 | 0.04826514 | 0.8055682 | -4.3073256 | up |
| KI - WT, cell_origin: iPSC | Q3MIN7 | RGL3 | 4 | Homo sapien | 1.75387342 | 0.00894535 | 3.49880148 | -2.5275891 | 2.14201785 | 0.04897681 | 0.81161062 | -4.3190744 | up |
| KI - WT, cell_origin: iPSC | P31146 | CORO1A | 7 | Homo sapien | -2.5123102 | -5.0132334 | -0.0113869 | -1.9753121 | -2.1407958 | 0.04909029 | 0.81161062 | -4.321136 | down |
| KI - WT, cell_origin: iPSC | O00764 | PDXK | 2 | Homo sapien | 1.27728859 | 0.00184803 | 2.55272916 | -1.7507366 | 2.13418065 | 0.04970882 | 0.81797853 | -4.3322853 | up |
| KI - WT, cell_origin: iPSC | P51572 | BCAP31 | 7 | Homo sapien | 1.01096083 | 0.0002656 | 2.02165606 | 0.40483092 | 2.13165286 | 0.04994707 | 0.81805838 | -4.336541 | up |
| KI - WT, cell_origin: iPSC | Q8WXG6 | MADD | 10 | Homo sapien | 0.69727386 | -0.00196 | 1.39650775 | -1.2478036 | 2.12383416 | 0.05056829 | 0.82398266 | -4.3488089 | up |
| KI - WT, cell_origin: iPSC | P52948 | NUP98 | 3 | Homo sapien | 1.37558861 | -0.0053002 | 2.75647745 | -1.3457099 | 2.12291309 | 0.05077896 | 0.82398266 | -4.3512346 | up |
| KI - WT, cell_origin: iPSC | Q86YV0 | RASAL3 | 28 | Homo sapien | 2.19805853 | -0.0143324 | 4.41044952 | -0.0255819 | 2.11728705 | 0.05132121 | 0.82691319 | -4.3606766 | up |
| KI - WT, cell_origin: iPSC | Q14677 | CLINT1 | 10 | Homo sapien | 1.33954589 | -0.0094587 | 2.68855047 | 0.36241626 | 2.11615044 | 0.05143141 | 0.82691319 | -4.3625825 | up |
| KI - WT, cell_origin: iPSC | Q9HD45 | TM9SF3 | 3 | Homo sapien | 2.03054334 | -0.0316076 | 4.09269424 | -2.1656865 | 2.09842856 | 0.05317798 | 0.85109053 | -4.39223 | up |
| KI - WT, cell_origin: iPSC | P62070 | RRAS2 | 11 | Homo sapien | 1.052013 | -0.0260306 | 2.13005663 | -0.0795543 | 2.07963508 | 0.05508964 | 0.87767814 | -4.4235254 | up |
| KI - WT, cell_origin: iPSC | A3KN83 | SBNO1 | 5 | Homo sapien | 1.88062514 | -0.0634065 | 3.82465682 | 4.43544291 | 2.0615851 | 0.05698486 | 0.90376438 | -4.4534399 | up |
| KI - WT, cell_origin: iPSC | P22090 | RPS4Y1 | 8 | Homo sapien | 0.48418446 | -0.018941 | 0.98730989 | 2.14306547 | 2.04962427 | 0.05814602 | 0.91802614 | -4.4724528 | up |
| KI - WT, cell_origin: iPSC | Q5VZ89 | DENND4C | 19 | Homo sapien | 1.15056581 | -0.0519887 | 2.35312026 | -0.8118701 | 2.03896175 | 0.05944455 | 0.93014798 | -4.4907332 | up |
| KI - WT, cell_origin: iPSC | P04004 | VTN | 2 | Homo sapien | -0.744541 | -1.5223638 | 0.03328184 | -1.3107881 | -2.0386732 | 0.05934843 | 0.93014798 | -4.490496 | down |
| KI - WT, cell_origin: iPSC | P38646 | HSPA9 | 12 | Homo sapien | 0.82524948 | -0.0397071 | 1.69020604 | 4.24368536 | 2.03326196 | 0.06007937 | 0.93590307 | -4.5000933 | up |
| KI - WT, cell_origin: iPSC | O75140 | DEPDC5 | 2 | Homo sapien | -0.970621 | -1.9936382 | 0.05239615 | -0.1915866 | -2.0219441 | 0.06135829 | 0.9483105 | -4.5186363 | down |
| KI - WT, cell_origin: iPSC | Q8IWB7 | WDFY1 | 10 | Homo sapien | 1.67688426 | -0.0909712 | 3.44473975 | -0.6402051 | 2.02142994 | 0.06141697 | 0.9483105 | -4.5194772 | up |
| KI - WT, cell_origin: iPSC | O75970 | MPDZ | 4 | Homo sapien | 1.3355636 | -0.0768234 | 2.7479506 | -2.9609795 | 2.01426988 | 0.06214304 | 0.95531296 | -4.5306802 | up |
| KI - WT, cell_origin: iPSC | P55160 | NCKAP1L | 13 | Homo sapien | 0.85194024 | -0.0518956 | 1.75577609 | 0.7986375 | 2.00858434 | 0.06288409 | 0.96248351 | -4.5403705 | up |

| | | | | | | | | | | | | | |
|---|---|---|---|---|---|---|---|---|---|---|---|---|---|
| KI - WT, cell_origin: iPSC | Q15008 | PSMD6 | 4 | Homo sapien | 0.64887032 | -0.0445263 | 1.34226689 | -1.2943281 | 1.99303944 | 0.06460698 | 0.98455426 | -4.5651 | up |
| KI - WT, cell_origin: iPSC | Q8TCU6 | PREX1 | 11 | Homo sapien | 2.71990603 | -0.193162 | 5.63297404 | -2.5539045 | 1.98978267 | 0.06512909 | 0.98821413 | -4.5710116 | up |
| KI - WT, cell_origin: iPSC | O43278 | SPINT1 | 12 | Homo sapien | 2.42236512 | -0.1834105 | 5.02814071 | -1.3702078 | 1.98109344 | 0.06618339 | 0.9998827 | -4.5850804 | up |
| KI - WT, cell_origin: iPSC | O14745 | SLC9A3R1 | 6 | Homo sapien | 1.88172943 | -0.1465307 | 3.90998953 | -0.8981286 | 1.97713307 | 0.06666904 | 0.99999785 | -4.5914811 | up |
| KI - WT, cell_origin: iPSC | Q8ND71 | GIMAP8 | 30 | Homo sapien | -1.2049101 | -2.5046992 | 0.09487898 | 0.10471237 | -1.9755323 | 0.06686625 | 0.99999785 | -4.5940661 | down |
| KI - WT, cell_origin: iPSC | Q9Y5K8 | ATP6V1D | 3 | Homo sapien | 0.6573646 | -0.0569482 | 1.37167745 | -0.516336 | 1.9600066 | 0.06867383 | 0.99999785 | -4.6185022 | up |
| KI - WT, cell_origin: iPSC | O95864 | FADS2 | 7 | Homo sapien | 1.30310523 | -0.1143785 | 2.72058891 | -1.6001098 | 1.9591324 | 0.06891735 | 0.99999785 | -4.6204803 | up |
| KI - WT, cell_origin: iPSC | Q9H270 | VPS11 | 7 | Homo sapien | 0.50656158 | -0.0460406 | 1.05916378 | -0.5437213 | 1.95235742 | 0.06964799 | 0.99999785 | -4.6307945 | up |
| KI - WT, cell_origin: iPSC | Q8IZD4 | DCP1B | 2 | Homo sapien | 0.75714846 | -0.0759666 | 1.5902635 | -1.9386358 | 1.9356006 | 0.07182582 | 0.99999785 | -4.6576244 | up |
| KI - WT, cell_origin: iPSC | P52888 | THOP1 | 4 | Homo sapien | 0.75142141 | -0.0772839 | 1.58012673 | -1.5553515 | 1.93118165 | 0.07241028 | 0.99999785 | -4.6646771 | up |
| KI - WT, cell_origin: iPSC | P21980 | TGM2 | 19 | Homo sapien | -1.2127697 | -2.5528647 | 0.12732531 | 0.30124676 | -1.9286131 | 0.07288641 | 0.99999785 | -4.6692944 | down |
| KI - WT, cell_origin: iPSC | Q9NZN4 | EHD2 | 11 | Homo sapien | 1.26730901 | -0.134029 | 2.66864704 | -2.6409376 | 1.92658325 | 0.07307859 | 0.99999785 | -4.6722206 | up |
| KI - WT, cell_origin: iPSC | O75976 | CPD | 10 | Homo sapien | 1.62186567 | -0.1723605 | 3.4160918 | -2.5693093 | 1.92637163 | 0.07318589 | 0.99999785 | -4.6728619 | up |
| KI - WT, cell_origin: iPSC | P46459 | NSF | 22 | Homo sapien | 0.8791467 | -0.0937247 | 1.8520181 | 0.71321485 | 1.92578712 | 0.07326416 | 0.99999785 | -4.6737917 | up |
| KI - WT, cell_origin: iPSC | Q641Q2;Q9Y | WASHC2A;W | 10 | Homo sapien | -1.1833744 | -2.4995236 | 0.13277488 | -0.8383935 | -1.9161054 | 0.07457175 | 0.99999785 | -4.6891697 | down |
| KI - WT, cell_origin: iPSC | P15170 | GSPT1 | 3 | Homo sapien | 0.72781919 | -0.0823959 | 1.53803429 | -1.5467963 | 1.9132109 | 0.07483151 | 0.99999785 | -4.6932599 | up |
| KI - WT, cell_origin: iPSC | Q8WUP2 | FBLIM1 | 7 | Homo sapien | 0.45566079 | -0.0543551 | 0.96567665 | 0.35577752 | 1.90281979 | 0.0762645 | 0.99999785 | -4.7097144 | up |
| KI - WT, cell_origin: iPSC | Q86XP1 | DGKH | 9 | Homo sapien | 1.09756956 | -0.1355334 | 2.33067251 | -1.9608299 | 1.89655224 | 0.07724033 | 0.99999785 | -4.7199595 | up |
| KI - WT, cell_origin: iPSC | Q9NP72 | RAB18 | 9 | Homo sapien | 1.20942705 | -0.1541929 | 2.57304695 | 1.4704268 | 1.8901171 | 0.07818601 | 0.99999785 | -4.7302191 | up |
| KI - WT, cell_origin: iPSC | P42566 | EPS15 | 5 | Homo sapien | 0.91832577 | -0.1208001 | 1.95745158 | -2.4955145 | 1.88221104 | 0.07917961 | 0.99999785 | -4.7421887 | up |
| KI - WT, cell_origin: iPSC | P13284 | IFI30 | 2 | Homo sapien | 2.31808568 | -0.3161495 | 4.9523209 | -3.332556 | 1.87532826 | 0.08031216 | 0.99999785 | -4.7534278 | up |
| KI - WT, cell_origin: iPSC | Q13503 | MED21 | 2 | Homo sapien | 0.69689516 | -0.0957745 | 1.48956483 | -0.7704939 | 1.87262 | 0.08058812 | 0.99999785 | -4.7572866 | up |
| KI - WT, cell_origin: iPSC | P05026 | ATP1B1 | 7 | Homo sapien | 5.48871281 | -0.7753705 | 11.7527961 | 0.27779627 | 1.86730539 | 0.08148711 | 0.99999785 | -4.765972 | up |
| KI - WT, cell_origin: iPSC | Q13615 | MTMR3 | 5 | Homo sapien | 1.2007132 | -0.1748561 | 2.57628254 | -1.4318911 | 1.86019798 | 0.08254079 | 0.99999785 | -4.7770574 | up |
| KI - WT, cell_origin: iPSC | Q10567 | AP1B1 | 13 | Homo sapien | 0.7585285 | -0.1133561 | 1.63041306 | 1.09426522 | 1.85372374 | 0.08347428 | 0.99999785 | -4.7870231 | up |
| KI - WT, cell_origin: iPSC | O75815 | BCAR3 | 3 | Homo sapien | 1.37192361 | -0.2055899 | 2.94943715 | -1.6152377 | 1.85335752 | 0.08356636 | 0.99999785 | -4.787702 | up |
| KI - WT, cell_origin: iPSC | O15234 | CASC3 | 5 | Homo sapien | -0.7884527 | -1.6990353 | 0.12212977 | -1.8067507 | -1.8441492 | 0.08482659 | 0.99999785 | -4.8015962 | down |
| KI - WT, cell_origin: iPSC | Q96MP8 | KCTD7 | 3 | Homo sapien | 0.65640597 | -0.1027859 | 1.41559787 | 0.44074423 | 1.84169046 | 0.08523293 | 0.99999785 | -4.805492 | up |
| KI - WT, cell_origin: iPSC | Q96F07 | CYFIP2 | 9 | Homo sapien | 0.54833428 | -0.0875764 | 1.18424496 | -0.9758139 | 1.83649167 | 0.08600512 | 0.99999785 | -4.8134575 | up |
| KI - WT, cell_origin: iPSC | P07686 | HEXB | 3 | Homo sapien | 1.53798546 | -0.2496829 | 3.3256538 | -2.1167764 | 1.83344399 | 0.08661693 | 0.99999785 | -4.8185519 | up |
| KI - WT, cell_origin: iPSC | Q86SZ2 | TRAPPC6B | 3 | Homo sapien | 0.39105533 | -0.0633779 | 0.84548854 | 1.18752246 | 1.83276943 | 0.08658323 | 0.99999785 | -4.819212 | up |
| KI - WT, cell_origin: iPSC | Q9Y478 | PRKAB1 | 5 | Homo sapien | 0.96446879 | -0.1569168 | 2.08585442 | -1.9835888 | 1.83210809 | 0.08672764 | 0.99999785 | -4.8203471 | up |
| KI - WT, cell_origin: iPSC | Q9NQG5 | RPRD1B | 3 | Homo sapien | 4.34965986 | -0.7277894 | 9.42710914 | -6.9331399 | 1.82562709 | 0.08784122 | 0.99999785 | -4.8306052 | up |
| KI - WT, cell_origin: iPSC | P31323 | PRKAR2B | 7 | Homo sapien | 0.92056459 | -0.1553386 | 1.99646782 | -0.0642548 | 1.8234062 | 0.08819186 | 0.99999785 | -4.8340238 | up |
| KI - WT, cell_origin: iPSC | Q86WX3 | RPS19BP1 | 3 | Homo sapien | -1.6063778 | -3.486764 | 0.27400833 | 0.57077229 | -1.8205517 | 0.08864434 | 0.99999785 | -4.8384139 | down |
| KI - WT, cell_origin: iPSC | Q13206 | DDX10 | 7 | Homo sapien | 0.59143102 | -0.1011801 | 1.28404213 | -2.2753995 | 1.81867179 | 0.0888042 | 0.99999785 | -4.8409405 | up |
| KI - WT, cell_origin: iPSC | Q5JSL3 | DOCK11 | 5 | Homo sapien | 1.21993915 | -0.2135313 | 2.6534096 | -0.7824671 | 1.81364295 | 0.08974805 | 0.99999785 | -4.8490216 | up |
| KI - WT, cell_origin: iPSC | P98196 | ATP11A | 7 | Homo sapien | 1.10231582 | -0.194982 | 2.39961361 | -2.2863131 | 1.81028717 | 0.09022412 | 0.99999785 | -4.8540028 | up |
| KI - WT, cell_origin: iPSC | P78549 | NTHL1 | 2 | Homo sapien | -1.106003 | -2.4089842 | 0.19697814 | -2.3952916 | -1.8081568 | 0.09053511 | 0.99999785 | -4.8571823 | down |
| KI - WT, cell_origin: iPSC | Q9UK59 | DBR1 | 5 | Homo sapien | -0.5546249 | -1.2126427 | 0.10339293 | -0.4637372 | -1.7951526 | 0.09262204 | 0.99999785 | -4.8769544 | down |
| KI - WT, cell_origin: iPSC | Q9P000 | COMMD9 | 5 | Homo sapien | 1.57958855 | -0.2956571 | 3.45483419 | -1.1921966 | 1.79509806 | 0.09277114 | 0.99999785 | -4.8773693 | up |
| KI - WT, cell_origin: iPSC | Q9Y5S9 | RBM8A | 4 | Homo sapien | -0.4740643 | -1.0382041 | 0.09007549 | 0.35507731 | -1.7897411 | 0.09352076 | 0.99999785 | -4.8851988 | down |
| KI - WT, cell_origin: iPSC | P45985 | MAP2K4 | 4 | Homo sapien | -2.0151986 | -4.4192432 | 0.38884599 | -2.6641059 | -1.7863959 | 0.09422052 | 0.99999785 | -4.8906077 | down |
| KI - WT, cell_origin: iPSC | P25774 | CTSS | 2 | Homo sapien | 1.70856727 | -0.3336238 | 3.75075837 | -1.5397975 | 1.78294551 | 0.0948007 | 0.99999785 | -4.8958453 | up |
| KI - WT, cell_origin: iPSC | P08779 | KRT16 | 15 | Homo sapien | -2.0540391 | -4.509176 | 0.40109777 | -0.0328019 | -1.7829344 | 0.09480257 | 0.99999785 | -4.8958621 | down |
| KI - WT, cell_origin: iPSC | P62879 | GNB2 | 14 | Homo sapien | 0.75943964 | -0.1492836 | 1.66816291 | 3.87324942 | 1.78100025 | 0.09512918 | 0.99999785 | -4.8987953 | up |

| | | | | | | | | | | | | | |
|---|---|---|---|---|---|---|---|---|---|---|---|---|---|
| KI - WT, cell_origin: iPSC | O95721 | SNAP29 | 5 | Homo sapien | 1.17350733 | -0.2345128 | 2.58152746 | -0.9755164 | 1.77614866 | 0.09595282 | 0.99999785 | -4.9061438 | up |
| KI - WT, cell_origin: iPSC | Q99523 | SORT1 | 12 | Homo sapien | 2.7320781 | -0.5578184 | 6.02197464 | -1.7208247 | 1.76975536 | 0.09704779 | 0.99999785 | -4.9158079 | up |
| KI - WT, cell_origin: iPSC | Q9NYB9 | ABI2 | 4 | Homo sapien | 0.53892754 | -0.1134818 | 1.19133689 | 0.69750692 | 1.75934045 | 0.09871397 | 0.99999785 | -4.9312154 | up |
| KI - WT, cell_origin: iPSC | O00193 | SMAP | 4 | Homo sapien | -0.6226717 | -1.3771426 | 0.13179911 | -0.3786416 | -1.757836 | 0.09898925 | 0.99999785 | -4.9335034 | down |
| KI - WT, cell_origin: iPSC | Q9Y3A6 | TMED5 | 5 | Homo sapien | 2.57883555 | -0.5507241 | 5.70839521 | 0.31325308 | 1.75607387 | 0.09942803 | 0.99999785 | -4.9364132 | up |
| KI - WT, cell_origin: iPSC | P52926 | HMGA2 | 4 | Homo sapien | -1.5233321 | -3.3739996 | 0.32733549 | 0.88197114 | -1.7541573 | 0.09976553 | 0.99999785 | -4.9392914 | down |
| KI - WT, cell_origin: iPSC | P46020 | PHKA1 | 10 | Homo sapien | -1.1850973 | -2.6311053 | 0.26091061 | -1.420906 | -1.7465688 | 0.10111172 | 0.99999785 | -4.9506674 | down |
| KI - WT, cell_origin: iPSC | Q9Y651 | SOX21 | 2 | Homo sapien | -0.7958 | -1.7671086 | 0.1755086 | 0.96479749 | -1.7460194 | 0.10120981 | 0.99999785 | -4.9514898 | down |
| KI - WT, cell_origin: iPSC | P10253 | GAA | 4 | Homo sapien | 1.51974377 | -0.3368306 | 3.37631813 | -0.4422938 | 1.7444575 | 0.1014891 | 0.99999785 | -4.9538268 | up |
| KI - WT, cell_origin: iPSC | P49917 | LIG4 | 3 | Homo sapien | 1.56430083 | -0.3637171 | 3.49231873 | -3.0197376 | 1.72906605 | 0.10427764 | 0.99999785 | -4.9767832 | up |
| KI - WT, cell_origin: iPSC | Q9H0A8 | COMMD4 | 3 | Homo sapien | 1.4599357 | -0.3482366 | 3.26810805 | -2.093486 | 1.7206648 | 0.10582784 | 0.99999785 | -4.9892573 | up |
| KI - WT, cell_origin: iPSC | Q86U44 | METTL3 | 4 | Homo sapien | -1.5654049 | -3.5158599 | 0.38505015 | -2.2615148 | -1.710382 | 0.10775259 | 0.99999785 | -5.0044705 | down |
| KI - WT, cell_origin: iPSC | Q8IV50 | LYSMD2 | 6 | Homo sapien | -1.0165251 | -2.2843324 | 0.25128225 | 0.10897608 | -1.7087055 | 0.10806929 | 0.99999785 | -5.0069451 | down |
| KI - WT, cell_origin: iPSC | Q9Y4K0 | LOXL2 | 9 | Homo sapien | 1.2115911 | -0.3008038 | 2.72398597 | -2.4512323 | 1.70723477 | 0.10834777 | 0.99999785 | -5.0091146 | up |
| KI - WT, cell_origin: iPSC | Q15738 | NSDHL | 2 | Homo sapien | 0.90592407 | -0.2247593 | 2.03660742 | -2.4399943 | 1.70645675 | 0.10835562 | 0.99999785 | -5.0100387 | up |
| KI - WT, cell_origin: iPSC | Q96SQ9 | CYP2S1 | 5 | Homo sapien | 1.18141087 | -0.2977295 | 2.66055126 | -0.1046481 | 1.70213473 | 0.10931832 | 0.99999785 | -5.0166283 | up |
| KI - WT, cell_origin: iPSC | Q2NKX8 | ERCC6L | 3 | Homo sapien | 0.69844261 | -0.1766377 | 1.57352296 | -2.7861291 | 1.69989648 | 0.10960422 | 0.99999785 | -5.019701 | up |
| KI - WT, cell_origin: iPSC | Q8WV41 | SNX33 | 3 | Homo sapien | 1.26674151 | -0.324995 | 2.85847803 | -2.6366312 | 1.695974 | 0.11050083 | 0.99999785 | -5.0256848 | up |
| KI - WT, cell_origin: iPSC | P19823 | ITIH2 | 2 | Homo sapien | -1.2633033 | -2.8588502 | 0.33224369 | -0.2296824 | -1.6873314 | 0.11217844 | 0.99999785 | -5.0383525 | down |
| KI - WT, cell_origin: iPSC | P38919 | EIF4A3 | 11 | Homo sapien | -0.4520708 | -1.0231843 | 0.1190427 | 0.55116616 | -1.6858686 | 0.11232193 | 0.99999785 | -5.040287 | down |
| KI - WT, cell_origin: iPSC | P33121 | ACSL1 | 27 | Homo sapien | 1.97406373 | -0.5287459 | 4.47687331 | 0.17779752 | 1.6808762 | 0.11344587 | 0.99999785 | -5.0477859 | up |
| KI - WT, cell_origin: iPSC | Q9UBK9 | UXT | 2 | Homo sapien | 0.65922409 | -0.179529 | 1.49797722 | -1.5342591 | 1.67393501 | 0.11467974 | 0.99999785 | -5.0577091 | up |
| KI - WT, cell_origin: iPSC | Q8NHY2 | COP1 | 3 | Homo sapien | 1.30764585 | -0.3610169 | 2.97630859 | -1.6320719 | 1.67002873 | 0.11560365 | 0.99999785 | -5.063583 | up |
| KI - WT, cell_origin: iPSC | Q92917 | GPKOW | 2 | Homo sapien | -1.1735577 | -2.6708534 | 0.32373803 | -3.3596943 | -1.6694568 | 0.11559656 | 0.99999785 | -5.0642531 | down |
| KI - WT, cell_origin: iPSC | Q15286 | RAB35 | 5 | Homo sapien | 0.92163289 | -0.2580892 | 2.10135499 | 4.29550835 | 1.66487111 | 0.11664202 | 0.99999785 | -5.0710697 | up |
| KI - WT, cell_origin: iPSC | Q8NCE2 | MTMR14 | 3 | Homo sapien | 1.32821046 | -0.3722294 | 3.02865029 | -3.627736 | 1.6638433 | 0.11674351 | 0.99999785 | -5.0724235 | up |
| KI - WT, cell_origin: iPSC | Q08AG7 | MZT1 | 3 | Homo sapien | 0.40425978 | -0.1140766 | 0.92259618 | 0.82597529 | 1.66107218 | 0.11726905 | 0.99999785 | -5.0763938 | up |
| KI - WT, cell_origin: iPSC | O14639 | ABLIM1 | 2 | Homo sapien | 2.62151686 | -0.7431626 | 5.98619629 | -4.1201932 | 1.66039467 | 0.11754978 | 0.99999785 | -5.0775549 | up |
| KI - WT, cell_origin: iPSC | Q07960 | ARHGAP1 | 11 | Homo sapien | 1.55824492 | -0.4454981 | 3.56198796 | -1.8820449 | 1.65728065 | 0.11818485 | 0.99999785 | -5.0820593 | up |
| KI - WT, cell_origin: iPSC | Q5VV52 | ZNF691 | 2 | Homo sapien | -0.6855905 | -1.5688049 | 0.19762402 | -1.3809824 | -1.6533641 | 0.11886125 | 0.99999785 | -5.0875634 | down |
| KI - WT, cell_origin: iPSC | P0C2W1 | FBXO45 | 4 | Homo sapien | -0.7935602 | -1.8164573 | 0.2293368 | -1.0859428 | -1.6531737 | 0.11900962 | 0.99999785 | -5.0879702 | down |
| KI - WT, cell_origin: iPSC | O15231 | ZNF185 | 4 | Homo sapien | 1.45820296 | -0.4277841 | 3.34419001 | -2.8001508 | 1.64771328 | 0.12015461 | 0.99999785 | -5.0958623 | up |
| KI - WT, cell_origin: iPSC | P10586 | PTPRF | 8 | Homo sapien | 2.64421226 | -0.7905636 | 6.07898813 | -3.2751586 | 1.64059083 | 0.12163932 | 0.99999785 | -5.1061025 | up |
| KI - WT, cell_origin: iPSC | Q6ZNL6 | FGD5 | 26 | Homo sapien | -0.7290015 | -1.6767932 | 0.21879012 | 0.61058183 | -1.6391471 | 0.1219422 | 0.99999785 | -5.1081745 | down |
| KI - WT, cell_origin: iPSC | Q9NV70 | EXOC1 | 16 | Homo sapien | 1.63790423 | -0.5000676 | 3.7758761 | -1.6528211 | 1.63263418 | 0.12331659 | 0.99999785 | -5.117506 | up |
| KI - WT, cell_origin: iPSC | Q17RY0 | CPEB4 | 13 | Homo sapien | 0.74020969 | -0.22895 | 1.70936935 | -0.5864737 | 1.62753351 | 0.12438489 | 0.99999785 | -5.1247782 | up |
| KI - WT, cell_origin: iPSC | Q9NZI8 | IGF2BP1 | 24 | Homo sapien | -0.456316 | -1.0538344 | 0.14120247 | 4.93624859 | -1.6267096 | 0.12446574 | 0.99999785 | -5.1258562 | down |
| KI - WT, cell_origin: iPSC | Q12926 | ELAVL2 | 3 | Homo sapien | -0.5590278 | -1.2912467 | 0.17319103 | -0.5884817 | -1.6260437 | 0.12457773 | 0.99999785 | -5.1267756 | down |
| KI - WT, cell_origin: iPSC | Q96PE2 | ARHGEF17 | 15 | Homo sapien | 0.85458891 | -0.2655328 | 1.97471061 | -1.5768336 | 1.62563909 | 0.12476926 | 0.99999785 | -5.1274606 | up |
| KI - WT, cell_origin: iPSC | O75695 | RP2 | 8 | Homo sapien | 0.68727245 | -0.218137 | 1.59268193 | 1.05060246 | 1.61753766 | 0.1265363 | 0.99999785 | -5.1390203 | up |
| KI - WT, cell_origin: iPSC | Q9ULV4 | CORO1C | 6 | Homo sapien | -0.408555 | -0.9467042 | 0.12959407 | -0.8271285 | -1.6169166 | 0.12654518 | 0.99999785 | -5.1397812 | down |
| KI - WT, cell_origin: iPSC | P04844 | RPN2 | 6 | Homo sapien | 2.64563875 | -0.8528346 | 6.14411214 | -4.0565176 | 1.61158915 | 0.12784905 | 0.99999785 | -5.1474831 | up |
| KI - WT, cell_origin: iPSC | Q92609 | TBC1D5 | 11 | Homo sapien | 0.44104739 | -0.1426858 | 1.02478055 | -0.6121795 | 1.60920201 | 0.12822887 | 0.99999785 | -5.1507344 | up |
| KI - WT, cell_origin: iPSC | Q9Y296 | TRAPPC4 | 7 | Homo sapien | 0.45626378 | -0.1483576 | 1.0608852 | -0.2657885 | 1.60720821 | 0.12866712 | 0.99999785 | -5.1535593 | up |
| KI - WT, cell_origin: iPSC | Q9NRR5 | UBQLN4 | 3 | Homo sapien | 0.92265058 | -0.3038935 | 2.14919469 | -0.0826114 | 1.60308466 | 0.12972076 | 0.99999785 | -5.1595204 | up |
| KI - WT, cell_origin: iPSC | O43491 | EPB41L2 | 9 | Homo sapien | 1.20557603 | -0.4002524 | 2.81140447 | -1.9616932 | 1.59991838 | 0.13042357 | 0.99999785 | -5.1639906 | up |

| | | | | | | | | | | | | | |
|---|---|---|---|---|---|---|---|---|---|---|---|---|---|
| KI - WT, cell_origin: iPSC | P51452 | DUSP3 | 2 | Homo sapien | -0.8427643 | -1.9651661 | 0.27963755 | -1.8706801 | -1.5994684 | 0.13042318 | 0.99999785 | -5.1645388 | down |
| KI - WT, cell_origin: iPSC | Q6NVY1 | HIBCH | 13 | Homo sapien | 0.67424674 | -0.2241564 | 1.57264986 | -0.053454 | 1.59905927 | 0.13056828 | 0.99999785 | -5.1651624 | up |
| KI - WT, cell_origin: iPSC | Q9Y6D5 | ARFGEF2 | 4 | Homo sapien | -1.1414169 | -2.6696896 | 0.38685586 | -2.9739241 | -1.5910784 | 0.13231897 | 0.99999785 | -5.17637 | down |
| KI - WT, cell_origin: iPSC | Q8IW19 | APLF | 8 | Homo sapien | 1.96061376 | -0.6710669 | 4.59229446 | -3.2252483 | 1.58767359 | 0.13317223 | 0.99999785 | -5.1812197 | up |
| KI - WT, cell_origin: iPSC | P37268 | FDFT1 | 4 | Homo sapien | 0.48109316 | -0.1649316 | 1.1271179 | -1.1208422 | 1.58606035 | 0.13339487 | 0.99999785 | -5.1833706 | up |
| KI - WT, cell_origin: iPSC | P14735 | IDE | 6 | Homo sapien | 0.49503723 | -0.1699571 | 1.16003156 | -1.2421019 | 1.58547575 | 0.13352764 | 0.99999785 | -5.1841908 | up |
| KI - WT, cell_origin: iPSC | P33176 | KIF5B | 26 | Homo sapien | 0.3763029 | -0.1293535 | 0.88195932 | -0.1292027 | 1.58497246 | 0.13364203 | 0.99999785 | -5.1848967 | up |
| KI - WT, cell_origin: iPSC | Q9H9H4 | VPS37B | 7 | Homo sapien | -0.5575314 | -1.3090488 | 0.19398594 | 0.0870654 | -1.5801606 | 0.13475694 | 0.99999785 | -5.1916505 | down |
| KI - WT, cell_origin: iPSC | Q02413 | DSG1 | 8 | Homo sapien | 1.43332797 | -0.5051537 | 3.37180965 | -1.3181481 | 1.5757461 | 0.135897 | 0.99999785 | -5.1979127 | up |
| KI - WT, cell_origin: iPSC | Q99704 | DOK1 | 3 | Homo sapien | -0.9430847 | -2.2183176 | 0.33214828 | -2.5981021 | -1.575317 | 0.13588938 | 0.99999785 | -5.1984348 | down |
| KI - WT, cell_origin: iPSC | Q8IUX4 | APOBEC3F | 4 | Homo sapien | -0.8956642 | -2.1174523 | 0.32612385 | -1.1801476 | -1.5622542 | 0.13903616 | 0.99999785 | -5.2166876 | down |
| KI - WT, cell_origin: iPSC | Q9NXF1 | TEX10 | 6 | Homo sapien | 0.45316402 | -0.165967 | 1.07229507 | -1.9344455 | 1.55887941 | 0.13968797 | 0.99999785 | -5.221276 | up |
| KI - WT, cell_origin: iPSC | Q08431 | MFGE8 | 14 | Homo sapien | 2.04536581 | -0.7520952 | 4.84282684 | -1.7606709 | 1.55815018 | 0.14000317 | 0.99999785 | -5.2223759 | up |
| KI - WT, cell_origin: iPSC | P37235 | HPCAL1 | 7 | Homo sapien | -1.1834098 | -2.8026207 | 0.43580117 | -1.9933459 | -1.5575216 | 0.14015178 | 0.99999785 | -5.2232462 | down |
| KI - WT, cell_origin: iPSC | P17677 | GAP43 | 5 | Homo sapien | 1.24051445 | -0.4629966 | 2.94402551 | -2.229272 | 1.55188393 | 0.14149065 | 0.99999785 | -5.2310405 | up |
| KI - WT, cell_origin: iPSC | Q9GZT4 | SRR | 5 | Homo sapien | 1.47349219 | -0.5591061 | 3.50609048 | -2.677419 | 1.54489386 | 0.1431657 | 0.99999785 | -5.2406765 | up |
| KI - WT, cell_origin: iPSC | O95716 | RAB3D | 3 | Homo sapien | -0.9250202 | -2.2016206 | 0.35158022 | -0.7261554 | -1.5441824 | 0.14333712 | 0.99999785 | -5.2416555 | down |
| KI - WT, cell_origin: iPSC | P49189 | ALDH9A1 | 7 | Homo sapien | 0.92430295 | -0.3517576 | 2.20036351 | -0.9713958 | 1.54363784 | 0.14346846 | 0.99999785 | -5.2424046 | up |
| KI - WT, cell_origin: iPSC | Q9H0T7 | RAB17 | 7 | Homo sapien | 2.8929478 | -1.1054462 | 6.89134179 | -2.6450103 | 1.54190416 | 0.14388724 | 0.99999785 | -5.2447883 | up |
| KI - WT, cell_origin: iPSC | Q63HR2 | TNS2 | 27 | Homo sapien | -0.9415605 | -2.2486834 | 0.36556231 | 0.44896642 | -1.5350913 | 0.14554294 | 0.99999785 | -5.254137 | down |
| KI - WT, cell_origin: iPSC | O60716 | CTNND1 | 33 | Homo sapien | 0.7509582 | -0.2963994 | 1.7983158 | 1.81910363 | 1.52799925 | 0.14728353 | 0.99999785 | -5.263837 | up |
| KI - WT, cell_origin: iPSC | Q9Y312 | AAR2 | 4 | Homo sapien | 0.74007024 | -0.2923085 | 1.77244896 | -0.8776354 | 1.52769361 | 0.14735893 | 0.99999785 | -5.2642543 | up |
| KI - WT, cell_origin: iPSC | Q9NZL6 | RGL1 | 4 | Homo sapien | -1.5198566 | -3.6417639 | 0.60205084 | -1.8471372 | -1.5264358 | 0.14766959 | 0.99999785 | -5.265971 | down |
| KI - WT, cell_origin: iPSC | O43347 | MSI1 | 8 | Homo sapien | -0.6312814 | -1.5123308 | 0.2497679 | -0.4289315 | -1.5264325 | 0.14759035 | 0.99999785 | -5.2659386 | down |
| KI - WT, cell_origin: iPSC | P16435 | POR | 3 | Homo sapien | 1.57160238 | -0.6239802 | 3.76718499 | -1.8864171 | 1.52544047 | 0.14791581 | 0.99999785 | -5.2673287 | up |
| KI - WT, cell_origin: iPSC | Q52LW3 | ARHGAP29 | 35 | Homo sapien | -0.5974372 | -1.4369238 | 0.24204937 | -0.0183072 | -1.5159903 | 0.15017087 | 0.99999785 | -5.2801471 | down |
| KI - WT, cell_origin: iPSC | Q9Y2H1 | STK38L | 4 | Homo sapien | 1.46777835 | -0.5966748 | 3.53223154 | -3.1029552 | 1.51515759 | 0.15047985 | 0.99999785 | -5.2813181 | up |
| KI - WT, cell_origin: iPSC | Q96RU3 | FNBP1 | 7 | Homo sapien | 0.58891373 | -0.2392323 | 1.41705972 | -0.5411576 | 1.51481065 | 0.15046471 | 0.99999785 | -5.2817478 | up |
| KI - WT, cell_origin: iPSC | O00425 | IGF2BP3 | 29 | Homo sapien | -0.6286003 | -1.5131431 | 0.25594256 | 5.12121804 | -1.5144609 | 0.15065491 | 0.99999785 | -5.2822635 | down |
| KI - WT, cell_origin: iPSC | Q9GZV5 | WWTR1 | 2 | Homo sapien | -0.4839992 | -1.1647718 | 0.19677328 | -2.0512693 | -1.5141969 | 0.15057896 | 0.99999785 | -5.2825649 | down |
| KI - WT, cell_origin: iPSC | Q9NZB2 | FAM120A | 31 | Homo sapien | -0.5426689 | -1.3065575 | 0.22121971 | 1.58417958 | -1.5132507 | 0.15085302 | 0.99999785 | -5.2838629 | down |
| KI - WT, cell_origin: iPSC | P30566 | ADSL | 3 | Homo sapien | -0.5257767 | -1.26605 | 0.21449663 | -1.4948772 | -1.5126864 | 0.15095926 | 0.99999785 | -5.2846141 | down |
| KI - WT, cell_origin: iPSC | Q14112 | NID2 | 12 | Homo sapien | 0.96974064 | -0.3975413 | 2.33702256 | -1.2646142 | 1.51147125 | 0.15140808 | 0.99999785 | -5.2863165 | up |
| KI - WT, cell_origin: iPSC | P02768 | ALB | 13 | Homo sapien | -0.7722797 | -1.8649955 | 0.32043612 | 2.91130147 | -1.5061553 | 0.15275511 | 0.99999785 | -5.2935089 | down |
| KI - WT, cell_origin: iPSC | Q9UJX5 | ANAPC4 | 2 | Homo sapien | 1.07821125 | -0.4465689 | 2.60299138 | -3.6208457 | 1.50603935 | 0.15264242 | 0.99999785 | -5.2936144 | up |
| KI - WT, cell_origin: iPSC | Q14160 | SCRIB | 19 | Homo sapien | 0.81601841 | -0.3410595 | 1.97309635 | -0.9665921 | 1.5029333 | 0.15357643 | 0.99999785 | -5.2978592 | up |
| KI - WT, cell_origin: iPSC | P62273 | RPS29 | 4 | Homo sapien | -0.3270912 | -0.7912645 | 0.13708199 | 4.34027992 | -1.5008196 | 0.15397516 | 0.99999785 | -5.3006618 | down |
| KI - WT, cell_origin: iPSC | Q15717 | ELAVL1 | 12 | Homo sapien | -0.5209816 | -1.2662536 | 0.22429041 | 2.91975081 | -1.4890907 | 0.1570453 | 0.99999785 | -5.3164432 | down |
| KI - WT, cell_origin: iPSC | P29350 | PTPN6 | 31 | Homo sapien | -1.3923301 | -3.3856457 | 0.60098541 | -0.7826688 | -1.4885675 | 0.15728353 | 0.99999785 | -5.3171728 | down |
| KI - WT, cell_origin: iPSC | P63165 | SUMO1 | 2 | Homo sapien | -0.7920876 | -1.9284258 | 0.3442506 | -1.6624703 | -1.4852436 | 0.15811392 | 0.99999785 | -5.3216121 | down |
| KI - WT, cell_origin: iPSC | P06858 | LPL | 8 | Homo sapien | -0.5179998 | -1.2617914 | 0.22579189 | -0.6703076 | -1.4832622 | 0.15852964 | 0.99999785 | -5.3242348 | down |
| KI - WT, cell_origin: iPSC | Q9UPY3 | DICER1 | 27 | Homo sapien | 0.77690443 | -0.3395685 | 1.89337736 | -1.3035661 | 1.48293381 | 0.15875757 | 0.99999785 | -5.3247096 | up |
| KI - WT, cell_origin: iPSC | Q9UP95 | SLC12A4 | 9 | Homo sapien | 1.24151082 | -0.5453963 | 3.02841795 | -2.8280859 | 1.48064483 | 0.15935976 | 0.99999785 | -5.3277658 | up |
| KI - WT, cell_origin: iPSC | Q9Y5X2 | SNX8 | 6 | Homo sapien | 0.48138948 | -0.212093 | 1.17487195 | -1.5109764 | 1.47843009 | 0.15980267 | 0.99999785 | -5.3306867 | up |
| KI - WT, cell_origin: iPSC | Q14764 | MVP | 9 | Homo sapien | 0.78597747 | -0.3525436 | 1.92449849 | -1.2842436 | 1.47119895 | 0.16186496 | 0.99999785 | -5.3403409 | up |
| KI - WT, cell_origin: iPSC | Q9NUL3 | STAU2 | 15 | Homo sapien | -0.4036571 | -0.9880512 | 0.18073688 | 0.6455826 | -1.4711148 | 0.16174606 | 0.99999785 | -5.3404244 | down |

| | | | | | | | | | | | | | |
|---|---|---|---|---|---|---|---|---|---|---|---|---|---|
| KI - WT, cell_origin: iPSC | Q16637 | SMN1 | 5 | Homo sapien | -0.5835473 | -1.4305266 | 0.26343206 | 0.0533622 | -1.4676454 | 0.16271667 | 0.99999785 | -5.3450379 | down |
| KI - WT, cell_origin: iPSC | Q14671 | PUM1 | 24 | Homo sapien | -0.4440738 | -1.0889038 | 0.20075623 | 0.71319201 | -1.466728 | 0.16292087 | 0.99999785 | -5.3462467 | down |
| KI - WT, cell_origin: iPSC | Q96CP2 | FLYWCH2 | 3 | Homo sapien | 1.09549675 | -0.4962843 | 2.6872778 | -1.7333802 | 1.4666623 | 0.16307975 | 0.99999785 | -5.3463593 | up |
| KI - WT, cell_origin: iPSC | O95219 | SNX4 | 4 | Homo sapien | 0.79678951 | -0.3608549 | 1.95443391 | -1.9340896 | 1.46627896 | 0.16309985 | 0.99999785 | -5.3468524 | up |
| KI - WT, cell_origin: iPSC | Q92625 | ANKS1A | 11 | Homo sapien | 1.37330462 | -0.6280853 | 3.37469451 | -1.6423947 | 1.46230359 | 0.16425401 | 0.99999785 | -5.3521286 | up |
| KI - WT, cell_origin: iPSC | Q9ULE0 | WWC3 | 4 | Homo sapien | 1.32937272 | -0.6161798 | 3.27492525 | -2.0944813 | 1.45615017 | 0.16592373 | 0.99999785 | -5.3602517 | up |
| KI - WT, cell_origin: iPSC | Q9NUD5 | ZCCHC3 | 17 | Homo sapien | -0.5059537 | -1.2464932 | 0.23458576 | 0.71261102 | -1.4552987 | 0.16604173 | 0.99999785 | -5.3613587 | down |
| KI - WT, cell_origin: iPSC | O00767 | SCD | 2 | Homo sapien | 1.36090524 | -0.6381325 | 3.359943 | -2.6376569 | 1.4508057 | 0.16738533 | 0.99999785 | -5.3672862 | up |
| KI - WT, cell_origin: iPSC | P06702 | S100A9 | 2 | Homo sapien | 0.75126425 | -0.3561349 | 1.85866338 | -0.169133 | 1.44574238 | 0.16877985 | 0.99999785 | -5.3739329 | up |
| KI - WT, cell_origin: iPSC | Q8TAT6 | NPLOC4 | 4 | Homo sapien | 0.61535233 | -0.2930405 | 1.52374512 | -1.8058358 | 1.4427459 | 0.16946913 | 0.99999785 | -5.3778469 | up |
| KI - WT, cell_origin: iPSC | Q2M1Z3 | ARHGAP31 | 12 | Homo sapien | -1.1651042 | -2.8870121 | 0.55680358 | -0.9968082 | -1.4419734 | 0.16982412 | 0.99999785 | -5.3788691 | down |
| KI - WT, cell_origin: iPSC | Q9UI08 | EVL | 14 | Homo sapien | -0.9032323 | -2.2386822 | 0.43221767 | -0.453478 | -1.4413658 | 0.16999298 | 0.99999785 | -5.3796641 | down |
| KI - WT, cell_origin: iPSC | Q96RY7 | IFT140 | 2 | Homo sapien | -0.4576203 | -1.1339871 | 0.21874645 | -0.7948898 | -1.4409958 | 0.16995538 | 0.99999785 | -5.3801377 | down |
| KI - WT, cell_origin: iPSC | Q9Y3E1 | HDGFL3 | 4 | Homo sapien | -1.7018179 | -4.2230183 | 0.8193825 | -1.1210887 | -1.4384941 | 0.17079286 | 0.99999785 | -5.3834174 | down |
| KI - WT, cell_origin: iPSC | P29083 | GTF2E1 | 3 | Homo sapien | 0.7675117 | -0.3758685 | 1.91089192 | -1.3954531 | 1.43052909 | 0.17302772 | 0.99999785 | -5.3937986 | up |
| KI - WT, cell_origin: iPSC | P50151 | GNG10 | 2 | Homo sapien | 1.13323036 | -0.5572239 | 2.82368462 | 0.78952581 | 1.42862138 | 0.17356655 | 0.99999785 | -5.3962785 | up |
| KI - WT, cell_origin: iPSC | Q02241 | KIF23 | 10 | Homo sapien | -0.3013942 | -0.7546715 | 0.15188315 | -0.3930189 | -1.4161542 | 0.17698244 | 0.99999785 | -5.4124275 | down |
| KI - WT, cell_origin: iPSC | Q9HAV0 | GNB4 | 7 | Homo sapien | 0.78023363 | -0.3944936 | 1.95496082 | 0.92001045 | 1.41543527 | 0.17732886 | 0.99999785 | -5.4133517 | up |
| KI - WT, cell_origin: iPSC | Q96LD4 | TRIM47 | 16 | Homo sapien | 0.54341412 | -0.27444 | 1.3612682 | 0.00148302 | 1.41537981 | 0.17724684 | 0.99999785 | -5.413426 | up |
| KI - WT, cell_origin: iPSC | Q5JVF3 | PCID2 | 5 | Homo sapien | -0.4576474 | -1.1490719 | 0.23377705 | -0.8293907 | -1.4096976 | 0.17884739 | 0.99999785 | -5.4207502 | down |
| KI - WT, cell_origin: iPSC | Q96BY6 | DOCK10 | 6 | Homo sapien | 0.49871535 | -0.2548056 | 1.25223626 | -1.6308299 | 1.40960396 | 0.17887456 | 0.99999785 | -5.4208707 | up |
| KI - WT, cell_origin: iPSC | Q12899 | TRIM26 | 5 | Homo sapien | 0.56872125 | -0.2948886 | 1.43233112 | -1.3263723 | 1.40262879 | 0.18091883 | 0.99999785 | -5.4298281 | up |
| KI - WT, cell_origin: iPSC | Q9UPN7 | PPP6R1 | 2 | Homo sapien | -0.8013258 | -2.0253734 | 0.42272181 | -1.0045845 | -1.3951252 | 0.18325447 | 0.99999785 | -5.4394138 | down |
| KI - WT, cell_origin: iPSC | P67809 | YBX1 | 7 | Homo sapien | -0.3922604 | -0.9923046 | 0.20778374 | 6.26201582 | -1.3927475 | 0.18389465 | 0.99999785 | -5.4424532 | down |
| KI - WT, cell_origin: iPSC | Q9H0J9 | PARP12 | 13 | Homo sapien | 0.75860574 | -0.4026096 | 1.91982107 | -0.0626159 | 1.39221315 | 0.18411719 | 0.99999785 | -5.443127 | up |
| KI - WT, cell_origin: iPSC | P61163 | ACTR1A | 5 | Homo sapien | 0.63613427 | -0.3375502 | 1.6098187 | -1.4340784 | 1.39170912 | 0.18416953 | 0.99999785 | -5.4437803 | up |
| KI - WT, cell_origin: iPSC | P63092;Q5JV | GNAS | 12 | Homo sapien | 0.91139311 | -0.4843498 | 2.30713599 | -0.5097687 | 1.39156242 | 0.18431043 | 0.99999785 | -5.443956 | up |
| KI - WT, cell_origin: iPSC | Q5EBL8 | PDZD11 | 3 | Homo sapien | 0.61767889 | -0.3282925 | 1.56365029 | -2.3873655 | 1.39067111 | 0.18443682 | 0.99999785 | -5.4451074 | up |
| KI - WT, cell_origin: iPSC | Q6AI08 | HEATR6 | 3 | Homo sapien | 0.75168089 | -0.3998791 | 1.90324086 | -2.5779043 | 1.39022991 | 0.1845681 | 0.99999785 | -5.4456692 | up |
| KI - WT, cell_origin: iPSC | O60711 | LPXN | 3 | Homo sapien | -0.6240733 | -1.5806832 | 0.33253664 | -1.2608879 | -1.3896922 | 0.18476947 | 0.99999785 | -5.4463487 | down |
| KI - WT, cell_origin: iPSC | Q96S59 | RANBP9 | 4 | Homo sapien | 1.01977909 | -0.5442766 | 2.58383478 | -2.0323311 | 1.38949266 | 0.18492615 | 0.99999785 | -5.4465906 | up |
| KI - WT, cell_origin: iPSC | P21964 | COMT | 7 | Homo sapien | 1.17824251 | -0.6313563 | 2.98784133 | -1.9024418 | 1.38756952 | 0.18549977 | 0.99999785 | -5.4490359 | up |
| KI - WT, cell_origin: iPSC | P14543 | NID1 | 11 | Homo sapien | 1.36172784 | -0.7299356 | 3.45339128 | -1.1358286 | 1.3873974 | 0.18555117 | 0.99999785 | -5.4492547 | up |
| KI - WT, cell_origin: iPSC | Q13546 | RIPK1 | 9 | Homo sapien | 0.45077085 | -0.2412527 | 1.14279442 | -0.9009872 | 1.38731354 | 0.18543781 | 0.99999785 | -5.4493793 | up |
| KI - WT, cell_origin: iPSC | Q96CD2 | PPCDC | 2 | Homo sapien | 1.1956651 | -0.6421596 | 3.03348983 | -4.0701626 | 1.38580186 | 0.18591943 | 0.99999785 | -5.4512961 | up |
| KI - WT, cell_origin: iPSC | O15226 | NKRF | 4 | Homo sapien | 0.53464525 | -0.2890276 | 1.3583181 | -0.8373984 | 1.38263368 | 0.18687012 | 0.99999785 | -5.4553161 | up |
| KI - WT, cell_origin: iPSC | Q9Y450 | HBS1L | 6 | Homo sapien | 0.51045914 | -0.2763167 | 1.29723494 | -1.0730734 | 1.38181428 | 0.18708686 | 0.99999785 | -5.4563591 | up |
| KI - WT, cell_origin: iPSC | Q8NFA0 | USP32 | 14 | Homo sapien | 0.66561929 | -0.3612982 | 1.6925368 | -0.5209673 | 1.38131493 | 0.18737533 | 0.99999785 | -5.4569709 | up |
| KI - WT, cell_origin: iPSC | Q8N3D4 | EHBP1L1 | 26 | Homo sapien | -0.5229955 | -1.3322901 | 0.28629901 | 7.59297268 | -1.3771896 | 0.18862082 | 0.99999785 | -5.4621895 | down |
| KI - WT, cell_origin: iPSC | Q13131 | PRKAA1 | 7 | Homo sapien | -0.5463788 | -1.392098 | 0.29934045 | -0.282794 | -1.3762126 | 0.18881995 | 0.99999785 | -5.4634401 | down |
| KI - WT, cell_origin: iPSC | Q6F5E8 | CARMIL2 | 3 | Homo sapien | 1.85126318 | -1.0155258 | 4.71805219 | -3.1268006 | 1.37617859 | 0.18892709 | 0.99999785 | -5.4634666 | up |
| KI - WT, cell_origin: iPSC | Q969F9 | HPS3 | 7 | Homo sapien | 0.42954343 | -0.2372173 | 1.09630419 | -0.3119914 | 1.37207147 | 0.19003767 | 0.99999785 | -5.4686725 | up |
| KI - WT, cell_origin: iPSC | Q06587 | RING1 | 3 | Homo sapien | 0.73527857 | -0.4062787 | 1.87683583 | -2.1254181 | 1.37202179 | 0.19008801 | 0.99999785 | -5.4687287 | up |
| KI - WT, cell_origin: iPSC | P23142 | FBLN1 | 9 | Homo sapien | -1.0114799 | -2.5876139 | 0.56465415 | -2.2613964 | -1.3676232 | 0.19153493 | 0.99999785 | -5.4742448 | down |
| KI - WT, cell_origin: iPSC | Q5T0D9 | TPRG1L | 6 | Homo sapien | 1.05299117 | -0.5888918 | 2.69487415 | -0.9469513 | 1.36673683 | 0.19180678 | 0.99999785 | -5.4753585 | up |
| KI - WT, cell_origin: iPSC | Q08AF3 | SLFN5 | 12 | Homo sapien | 0.65774662 | -0.3684418 | 1.68393508 | 0.20347575 | 1.36594707 | 0.19204926 | 0.99999785 | -5.4763503 | up |

| | | | | | | | | | | | | | |
|---|---|---|---|---|---|---|---|---|---|---|---|---|---|
| KI - WT, cell_origin: iPSC | Q2PPJ7 | RALGAPA2 | 9 | Homo sapien | 0.90505682 | -0.5096879 | 2.31980157 | -2.1490912 | 1.36332727 | 0.19285538 | 0.99999785 | -5.4796373 | up |
| KI - WT, cell_origin: iPSC | Q14257 | RCN2 | 2 | Homo sapien | -0.6544795 | -1.6782794 | 0.36932034 | -1.4823096 | -1.3617547 | 0.19324421 | 0.99999785 | -5.481629 | down |
| KI - WT, cell_origin: iPSC | Q96K83 | ZNF521 | 9 | Homo sapien | -0.5525342 | -1.417178 | 0.31210968 | -0.9968787 | -1.3612211 | 0.1934033 | 0.99999785 | -5.4822988 | down |
| KI - WT, cell_origin: iPSC | Q9NXC2 | GFOD1 | 4 | Homo sapien | 1.01333651 | -0.5746064 | 2.60127937 | -2.0942316 | 1.35994451 | 0.19390033 | 0.99999785 | -5.4838743 | up |
| KI - WT, cell_origin: iPSC | O14976 | GAK | 23 | Homo sapien | 0.9112077 | -0.5182313 | 2.34064672 | -0.9071571 | 1.35848272 | 0.19435329 | 0.99999785 | -5.4857028 | up |
| KI - WT, cell_origin: iPSC | Q8TDB6 | DTX3L | 39 | Homo sapien | 0.43022204 | -0.2453771 | 1.1058212 | 3.34310575 | 1.35639833 | 0.19488665 | 0.99999785 | -5.4883342 | up |
| KI - WT, cell_origin: iPSC | Q96MI6 | PPM1M | 8 | Homo sapien | 0.74427636 | -0.4254982 | 1.91405096 | 0.38734676 | 1.35592105 | 0.19514914 | 0.99999785 | -5.4889032 | up |
| KI - WT, cell_origin: iPSC | O43663 | PRC1 | 5 | Homo sapien | 0.59304867 | -0.3433472 | 1.52944452 | -0.1753054 | 1.34951856 | 0.19712149 | 0.99999785 | -5.4968892 | up |
| KI - WT, cell_origin: iPSC | P32119 | PRDX2 | 10 | Homo sapien | -0.4265739 | -1.1012804 | 0.24813255 | 1.57924261 | -1.3465395 | 0.19794977 | 0.99999785 | -5.5006208 | down |
| KI - WT, cell_origin: iPSC | Q9NRA8 | EIF4ENIF1 | 7 | Homo sapien | -0.5725496 | -1.478685 | 0.3335858 | -0.9593411 | -1.3459779 | 0.19816741 | 0.99999785 | -5.5013074 | down |
| KI - WT, cell_origin: iPSC | Q5SW79 | CEP170 | 19 | Homo sapien | -0.4993548 | -1.2908846 | 0.29217503 | -0.9425621 | -1.3437003 | 0.1988566 | 0.99999785 | -5.5041417 | down |
| KI - WT, cell_origin: iPSC | O75683 | SURF6 | 6 | Homo sapien | 0.66895901 | -0.3927916 | 1.73070964 | -0.9274468 | 1.34270111 | 0.19929819 | 0.99999785 | -5.5053453 | up |
| KI - WT, cell_origin: iPSC | P37840 | SNCA | 4 | Homo sapien | 0.68353767 | -0.4032204 | 1.77029569 | -0.584746 | 1.34039242 | 0.20003001 | 0.99999785 | -5.5082038 | up |
| KI - WT, cell_origin: iPSC | Q9UPZ3 | HPS5 | 9 | Homo sapien | 0.43759681 | -0.2603478 | 1.13554147 | -0.0883685 | 1.33534303 | 0.20150222 | 0.99999785 | -5.5144835 | up |
| KI - WT, cell_origin: iPSC | O75925 | PIAS1 | 2 | Homo sapien | 1.22915954 | -0.7375811 | 3.1959002 | -2.4745737 | 1.33187519 | 0.20274853 | 0.99999785 | -5.5187163 | up |
| KI - WT, cell_origin: iPSC | Q9UKM9 | RALY | 6 | Homo sapien | -0.3887268 | -1.0111396 | 0.23368586 | 0.16785144 | -1.3301658 | 0.2031621 | 0.99999785 | -5.5208631 | down |
| KI - WT, cell_origin: iPSC | P52435;Q9G: | POLR2J;POLR | 2 | Homo sapien | 0.91916391 | -0.5541262 | 2.39245399 | -0.6855112 | 1.32955734 | 0.20349346 | 0.99999785 | -5.5215681 | up |
| KI - WT, cell_origin: iPSC | P61221 | ABCE1 | 15 | Homo sapien | 0.50188599 | -0.3024652 | 1.30623721 | -0.8096543 | 1.32905529 | 0.20354227 | 0.99999785 | -5.5222216 | up |
| KI - WT, cell_origin: iPSC | Q9HCE0 | EPG5 | 3 | Homo sapien | 1.16781476 | -0.7050978 | 3.04072732 | -1.1275561 | 1.32879757 | 0.20373811 | 0.99999785 | -5.522502 | up |
| KI - WT, cell_origin: iPSC | Q8IX90 | SKA3 | 8 | Homo sapien | 0.66739746 | -0.4052034 | 1.73999828 | -1.7860244 | 1.32545297 | 0.20472289 | 0.99999785 | -5.52664 | up |
| KI - WT, cell_origin: iPSC | P42356 | PI4KA | 12 | Homo sapien | 0.56450094 | -0.3432587 | 1.47226062 | -0.7661401 | 1.32486097 | 0.20494472 | 0.99999785 | -5.527356 | up |
| KI - WT, cell_origin: iPSC | O43307 | ARHGEF9 | 2 | Homo sapien | 0.48374318 | -0.2948279 | 1.26231421 | -2.7494812 | 1.32329384 | 0.20538221 | 0.99999785 | -5.5293008 | up |
| KI - WT, cell_origin: iPSC | O43719 | HTATSF1 | 3 | Homo sapien | -0.4548862 | -1.1874001 | 0.27762771 | -0.6207699 | -1.322594 | 0.20560938 | 0.99999785 | -5.5301581 | down |
| KI - WT, cell_origin: iPSC | P14621 | ACYP2 | 3 | Homo sapien | -0.6043988 | -1.5795104 | 0.37071291 | -1.6443199 | -1.3202765 | 0.20639195 | 0.99999785 | -5.5329847 | down |
| KI - WT, cell_origin: iPSC | Q9BVQ7 | SPATA5L1 | 3 | Homo sapien | -0.5856338 | -1.5304638 | 0.35919624 | -2.3734913 | -1.3201161 | 0.2064154 | 0.99999785 | -5.533191 | down |
| KI - WT, cell_origin: iPSC | Q08379 | GOLGA2 | 3 | Homo sapien | 1.58339602 | -0.9744593 | 4.14125138 | -6.3791079 | 1.31841837 | 0.20696907 | 0.99999785 | -5.5352663 | up |
| KI - WT, cell_origin: iPSC | P22897 | MRC1 | 10 | Homo sapien | -2.2452189 | -5.8857426 | 1.39530479 | -2.849005 | -1.3143081 | 0.20844919 | 0.99999785 | -5.5402327 | down |
| KI - WT, cell_origin: iPSC | P31483 | TIA1 | 2 | Homo sapien | -0.5157755 | -1.3530296 | 0.32147868 | -2.5908338 | -1.3120278 | 0.20906388 | 0.99999785 | -5.5430593 | down |
| KI - WT, cell_origin: iPSC | Q02833 | RASSF7 | 2 | Homo sapien | 0.45536352 | -0.2840223 | 1.19474936 | -1.8617659 | 1.31167665 | 0.20917949 | 0.99999785 | -5.5434867 | up |
| KI - WT, cell_origin: iPSC | Q8IUD6 | RNF135 | 2 | Homo sapien | -1.6339185 | -4.2913261 | 1.02348907 | -1.1569324 | -1.3103116 | 0.20976381 | 0.99999785 | -5.5450963 | down |
| KI - WT, cell_origin: iPSC | P46778 | RPL21 | 6 | Homo sapien | -0.2877833 | -0.7573173 | 0.18175076 | 1.74153284 | -1.3053836 | 0.21125975 | 0.99999785 | -5.5511299 | down |
| KI - WT, cell_origin: iPSC | Q8N9N7 | LRRC57 | 9 | Homo sapien | 1.17600769 | -0.7473036 | 3.09931896 | -0.1241745 | 1.30305562 | 0.21216752 | 0.99999785 | -5.5538966 | up |
| KI - WT, cell_origin: iPSC | P51178 | PLCD1 | 2 | Homo sapien | 2.21707434 | -1.4181752 | 5.85232385 | -3.2841477 | 1.2997158 | 0.21328125 | 0.99999785 | -5.5579343 | up |
| KI - WT, cell_origin: iPSC | Q9NQS7 | INCENP | 4 | Homo sapien | -0.3310501 | -0.8743261 | 0.21222589 | -0.7924982 | -1.2978153 | 0.21378333 | 0.99999785 | -5.5602835 | down |
| KI - WT, cell_origin: iPSC | Q9UNM6 | PSMD13 | 8 | Homo sapien | 0.50718426 | -0.3265568 | 1.34092528 | -1.1349601 | 1.29570019 | 0.21450827 | 0.99999785 | -5.5628277 | up |
| KI - WT, cell_origin: iPSC | Q96EK4 | THAP11 | 2 | Homo sapien | -0.3060349 | -0.8106022 | 0.19853237 | -0.7382927 | -1.291789 | 0.21580986 | 0.99999785 | -5.567542 | down |
| KI - WT, cell_origin: iPSC | P53814 | SMTN | 8 | Homo sapien | -0.5320266 | -1.4106358 | 0.3465825 | -1.3808426 | -1.2897386 | 0.21651518 | 0.99999785 | -5.5700001 | down |
| KI - WT, cell_origin: iPSC | Q68CZ2 | TNS3 | 19 | Homo sapien | -0.7836923 | -2.0803739 | 0.51298918 | 0.72587266 | -1.2879964 | 0.21722614 | 0.99999785 | -5.5720375 | down |
| KI - WT, cell_origin: iPSC | O95999 | BCL10 | 2 | Homo sapien | 0.66706157 | -0.4370366 | 1.77115976 | -0.9969399 | 1.28753958 | 0.21738107 | 0.99999785 | -5.5725852 | up |
| KI - WT, cell_origin: iPSC | P35269 | GTF2F1 | 14 | Homo sapien | -1.0087075 | -2.6796281 | 0.66221308 | -1.04733 | -1.286506 | 0.21773195 | 0.99999785 | -5.5738238 | down |
| KI - WT, cell_origin: iPSC | Q9C005 | DPY30 | 2 | Homo sapien | -1.1883892 | -3.162726 | 0.7859476 | -0.7910201 | -1.2827435 | 0.219013 | 0.99999785 | -5.578326 | down |
| KI - WT, cell_origin: iPSC | Q15293 | RCN1 | 5 | Homo sapien | 1.0631965 | -0.7039819 | 2.83037495 | -3.016424 | 1.28214015 | 0.21921898 | 0.99999785 | -5.5790469 | up |
| KI - WT, cell_origin: iPSC | Q6ZSR9 | | 4 | Homo sapien | -0.6318302 | -1.6826065 | 0.41894618 | -1.6273965 | -1.2808786 | 0.219557 | 0.99999785 | -5.5805964 | down |
| KI - WT, cell_origin: iPSC | Q86WR7 | PROSER2 | 3 | Homo sapien | -0.7455496 | -1.9860427 | 0.49494352 | -2.0057907 | -1.2806023 | 0.21970922 | 0.99999785 | -5.5808998 | down |
| KI - WT, cell_origin: iPSC | Q13685 | AAMP | 3 | Homo sapien | 0.46623347 | -0.3102867 | 1.24275367 | -1.7348045 | 1.2787639 | 0.22024191 | 0.99999785 | -5.5831381 | up |
| KI - WT, cell_origin: iPSC | Q53GL0 | PLEKHO1 | 3 | Homo sapien | 1.65955356 | -1.1155941 | 4.43470119 | -3.5079678 | 1.27440524 | 0.22187326 | 0.99999785 | -5.5882659 | up |

| | | | | | | | | | | | | | |
|---|---|---|---|---|---|---|---|---|---|---|---|---|---|
| KI - WT, cell_origin: iPSC | Q8IYU2 | HACE1 | 4 | Homo sapien | -0.6566993 | -1.7549207 | 0.44152206 | -2.1592469 | -1.2736801 | 0.22201332 | 0.99999785 | -5.5891805 | down |
| KI - WT, cell_origin: iPSC | Q92900 | UPF1 | 41 | Homo sapien | -0.5858847 | -1.5664173 | 0.39464797 | 2.25731587 | -1.2733636 | 0.22223262 | 0.99999785 | -5.5895039 | down |
| KI - WT, cell_origin: iPSC | Q9UK53 | ING1 | 2 | Homo sapien | 1.7951363 | -1.2146109 | 4.80488347 | -2.295415 | 1.27107092 | 0.22302525 | 0.99999785 | -5.5922262 | up |
| KI - WT, cell_origin: iPSC | O43237 | DYNC1LI2 | 4 | Homo sapien | 0.57364946 | -0.3889931 | 1.53629201 | -1.168613 | 1.26949122 | 0.22349523 | 0.99999785 | -5.5941375 | up |
| KI - WT, cell_origin: iPSC | O43374 | RASA4 | 21 | Homo sapien | 0.84066805 | -0.5709765 | 2.25231256 | 0.28155794 | 1.26911671 | 0.2237026 | 0.99999785 | -5.5945434 | up |
| KI - WT, cell_origin: iPSC | Q8TF46 | DIS3L | 5 | Homo sapien | 0.64163376 | -0.438935 | 1.72220247 | -1.4902492 | 1.26515031 | 0.22503477 | 0.99999785 | -5.5992617 | up |
| KI - WT, cell_origin: iPSC | O95544 | NADK | 4 | Homo sapien | 0.51241698 | -0.3512992 | 1.37613312 | -1.4498831 | 1.26354921 | 0.22550972 | 0.99999785 | -5.6011959 | up |
| KI - WT, cell_origin: iPSC | O00443 | PIK3C2A | 12 | Homo sapien | 0.74636659 | -0.5151544 | 2.00788762 | -1.6609396 | 1.26084025 | 0.22658933 | 0.99999785 | -5.6043257 | up |
| KI - WT, cell_origin: iPSC | Q9Y5B0 | CTDP1 | 3 | Homo sapien | -0.5536827 | -1.4925018 | 0.38513645 | 0.43887506 | -1.256844 | 0.22799362 | 0.99999785 | -5.6090306 | down |
| KI - WT, cell_origin: iPSC | P52565 | ARHGDIA | 5 | Homo sapien | -0.7134673 | -1.9249048 | 0.49797012 | -0.7565918 | -1.2550917 | 0.22861153 | 0.99999785 | -5.6110899 | down |
| KI - WT, cell_origin: iPSC | P05109 | S100A8 | 3 | Homo sapien | 0.95841164 | -0.6697079 | 2.58653118 | -0.412864 | 1.25449276 | 0.22882303 | 0.99999785 | -5.6117932 | up |
| KI - WT, cell_origin: iPSC | P25054 | APC | 7 | Homo sapien | 0.99727543 | -0.6979755 | 2.69252633 | -2.4256741 | 1.25367079 | 0.22911354 | 0.99999785 | -5.612758 | up |
| KI - WT, cell_origin: iPSC | Q16881 | TXNRD1 | 5 | Homo sapien | -0.9191227 | -2.4815276 | 0.64328226 | -1.6952768 | -1.2536672 | 0.22911483 | 0.99999785 | -5.6127623 | down |
| KI - WT, cell_origin: iPSC | P21266 | GSTM3 | 5 | Homo sapien | 0.66362797 | -0.464437 | 1.79169299 | -2.3530905 | 1.25302022 | 0.22922672 | 0.99999785 | -5.6135834 | up |
| KI - WT, cell_origin: iPSC | P36507 | MAP2K2 | 6 | Homo sapien | -0.9010632 | -2.436087 | 0.6339607 | -0.7332718 | -1.2509573 | 0.23007466 | 0.99999785 | -5.6159394 | down |
| KI - WT, cell_origin: iPSC | O94915 | FRYL | 9 | Homo sapien | 0.71120154 | -0.5005616 | 1.92296463 | -1.7774752 | 1.25067788 | 0.23015794 | 0.99999785 | -5.6162752 | up |
| KI - WT, cell_origin: iPSC | Q92879 | CELF1 | 9 | Homo sapien | -0.4754286 | -1.285684 | 0.33482688 | 0.35428324 | -1.2499171 | 0.23035219 | 0.99999785 | -5.6172068 | down |
| KI - WT, cell_origin: iPSC | Q9Y237 | PIN4 | 3 | Homo sapien | -1.6015934 | -4.3325037 | 1.12931682 | -1.3552043 | -1.2498193 | 0.23047866 | 0.99999785 | -5.617272 | down |
| KI - WT, cell_origin: iPSC | Q13588 | GRAP | 5 | Homo sapien | 1.18643739 | -0.8375108 | 3.21038555 | -2.4004955 | 1.24924555 | 0.23068256 | 0.99999785 | -5.6179434 | up |
| KI - WT, cell_origin: iPSC | P46100 | ATRX | 5 | Homo sapien | -0.713922 | -1.9314125 | 0.50356856 | -2.2339621 | -1.2491168 | 0.23063665 | 0.99999785 | -5.6181436 | down |
| KI - WT, cell_origin: iPSC | Q86YV9 | HPS6 | 5 | Homo sapien | 0.47688906 | -0.3365861 | 1.29036423 | -0.1145306 | 1.24878202 | 0.23075358 | 0.99999785 | -5.6185365 | up |
| KI - WT, cell_origin: iPSC | Q96I24 | FUBP3 | 20 | Homo sapien | -0.4208834 | -1.1390026 | 0.29723592 | 2.3880818 | -1.2483463 | 0.23088693 | 0.99999785 | -5.6190579 | down |
| KI - WT, cell_origin: iPSC | Q9UK61 | TASOR | 2 | Homo sapien | 0.36268906 | -0.2567531 | 0.98213119 | -0.7618137 | 1.24701994 | 0.23134442 | 0.99999785 | -5.6206169 | up |
| KI - WT, cell_origin: iPSC | Q8N3Y1 | FBXW8 | 3 | Homo sapien | 0.59886466 | -0.4256122 | 1.62334153 | -0.9961136 | 1.24574426 | 0.23192992 | 0.99999785 | -5.6220357 | up |
| KI - WT, cell_origin: iPSC | Q99717 | SMAD5 | 2 | Homo sapien | -0.9316619 | -2.5256974 | 0.66237364 | -0.938114 | -1.2455544 | 0.23199772 | 0.99999785 | -5.6222573 | down |
| KI - WT, cell_origin: iPSC | P98155 | VLDLR | 10 | Homo sapien | 1.043652 | -0.7428229 | 2.83012691 | -3.7318409 | 1.24444783 | 0.23230168 | 0.99999785 | -5.623599 | up |
| KI - WT, cell_origin: iPSC | Q9UHY8 | FEZ2 | 2 | Homo sapien | 1.78699981 | -1.276053 | 4.85005256 | -4.322365 | 1.24328987 | 0.23280746 | 0.99999785 | -5.6248988 | up |
| KI - WT, cell_origin: iPSC | O94905 | ERLIN2 | 4 | Homo sapien | 0.53757261 | -0.3889267 | 1.46407191 | -1.3799199 | 1.23592424 | 0.2353566 | 0.99999785 | -5.6335208 | up |
| KI - WT, cell_origin: iPSC | O43164 | PJA2 | 5 | Homo sapien | 0.50489342 | -0.3654459 | 1.37523271 | -2.0187244 | 1.23552291 | 0.23547197 | 0.99999785 | -5.6340034 | up |
| KI - WT, cell_origin: iPSC | Q9BU61 | NDUFAF3 | 2 | Homo sapien | 1.0405808 | -0.7555535 | 2.83671508 | -1.4589749 | 1.23463725 | 0.23592183 | 0.99999785 | -5.6349559 | up |
| KI - WT, cell_origin: iPSC | Q9NZI7 | UBP1 | 2 | Homo sapien | -0.7550849 | -2.0597501 | 0.54958028 | -2.4105965 | -1.2328624 | 0.23647378 | 0.99999785 | -5.6370645 | down |
| KI - WT, cell_origin: iPSC | Q12974 | PTP4A2 | 5 | Homo sapien | 0.84886927 | -0.6215147 | 2.31925319 | -1.0839081 | 1.23030399 | 0.23749367 | 0.99999785 | -5.6399712 | up |
| KI - WT, cell_origin: iPSC | Q96BM9 | ARL8A | 6 | Homo sapien | 1.97554138 | -1.4512029 | 5.40228567 | -1.3978254 | 1.2285895 | 0.23811783 | 0.99999785 | -5.6419516 | up |
| KI - WT, cell_origin: iPSC | P02790 | HPX | 4 | Homo sapien | -0.4449191 | -1.2173542 | 0.32751601 | -0.1497312 | -1.2268818 | 0.23863343 | 0.99999785 | -5.6439856 | down |
| KI - WT, cell_origin: iPSC | Q96JJ3 | ELMO2 | 31 | Homo sapien | 0.44246453 | -0.3260797 | 1.21100875 | 1.59211643 | 1.22635556 | 0.23883703 | 0.99999785 | -5.6445857 | up |
| KI - WT, cell_origin: iPSC | Q9H2H9 | SLC38A1 | 4 | Homo sapien | 0.69301792 | -0.5128635 | 1.89889936 | -1.3152455 | 1.22473528 | 0.23952562 | 0.99999785 | -5.6463954 | up |
| KI - WT, cell_origin: iPSC | Q9UPX8 | SHANK2 | 14 | Homo sapien | 0.77025559 | -0.5724444 | 2.1129556 | 0.62356465 | 1.22252637 | 0.24033536 | 0.99999785 | -5.648937 | up |
| KI - WT, cell_origin: iPSC | Q96P47 | AGAP3 | 5 | Homo sapien | 1.19227275 | -0.8897011 | 3.27424657 | -1.6178058 | 1.22040146 | 0.2411163 | 0.99999785 | -5.6513785 | up |
| KI - WT, cell_origin: iPSC | Q00537 | CDK17 | 18 | Homo sapien | 0.45972946 | -0.343133 | 1.26259188 | 0.93996943 | 1.21977324 | 0.24125737 | 0.99999785 | -5.6521546 | up |
| KI - WT, cell_origin: iPSC | P57723 | PCBP4 | 2 | Homo sapien | -1.9269978 | -5.2944032 | 1.44040755 | -1.0159162 | -1.2195179 | 0.24144159 | 0.99999785 | -5.6523927 | down |
| KI - WT, cell_origin: iPSC | P05997 | COL5A2 | 2 | Homo sapien | 1.10444968 | -0.8277327 | 3.03663205 | -2.6222907 | 1.21814836 | 0.24194651 | 0.99999785 | -5.6539635 | up |
| KI - WT, cell_origin: iPSC | Q9NWH9 | SLTM | 9 | Homo sapien | -0.4759142 | -1.3116084 | 0.35977999 | -0.7592552 | -1.2130762 | 0.24372834 | 0.99999785 | -5.6598281 | down |
| KI - WT, cell_origin: iPSC | Q9HCE1 | MOV10 | 34 | Homo sapien | -0.7206694 | -1.988402 | 0.54706322 | 0.74927908 | -1.2114647 | 0.24442229 | 0.99999785 | -5.6616088 | down |
| KI - WT, cell_origin: iPSC | Q7L5N7 | LPCAT2 | 4 | Homo sapien | 1.21465275 | -0.9293622 | 3.35866766 | -2.1710314 | 1.20733198 | 0.24596294 | 0.99999785 | -5.666319 | up |
| KI - WT, cell_origin: iPSC | Q96PV7 | FAM193B | 3 | Homo sapien | -0.5643066 | -1.5610197 | 0.43240654 | -1.7268199 | -1.2059814 | 0.24636781 | 0.99999785 | -5.6679195 | down |
| KI - WT, cell_origin: iPSC | Q01432 | AMPD3 | 6 | Homo sapien | 0.38125621 | -0.2920635 | 1.05457593 | -1.3445063 | 1.20596657 | 0.2463462 | 0.99999785 | -5.6679538 | up |

| | | | | | | | | | | | | | |
|---|---|---|---|---|---|---|---|---|---|---|---|---|---|
| KI - WT, cell_origin: iPSC | P17948 | FLT1 | 15 | Homo sapien | 0.90414405 | -0.694385 | 2.50267307 | -0.1415597 | 1.20536748 | 0.24669792 | 0.99999785 | -5.6685533 | up |
| KI - WT, cell_origin: iPSC | Q9Y263 | PLAA | 4 | Homo sapien | -0.460283 | -1.2759755 | 0.35540945 | -1.722385 | -1.2018166 | 0.24790364 | 0.99999785 | -5.6726669 | down |
| KI - WT, cell_origin: iPSC | P62333 | PSMC6 | 6 | Homo sapien | 0.40664648 | -0.3141653 | 1.12745831 | -0.05543 | 1.20153049 | 0.24801128 | 0.99999785 | -5.6729913 | up |
| KI - WT, cell_origin: iPSC | Q15369 | ELOC | 5 | Homo sapien | -0.2998282 | -0.8319008 | 0.23224444 | 2.48099941 | -1.2001657 | 0.24852528 | 0.99999785 | -5.6745381 | down |
| KI - WT, cell_origin: iPSC | Q13310 | PABPC4 | 14 | Homo sapien | -0.3289899 | -0.9135976 | 0.25561778 | 4.00831929 | -1.1985544 | 0.24913322 | 0.99999785 | -5.6763624 | down |
| KI - WT, cell_origin: iPSC | Q06787 | FMR1 | 10 | Homo sapien | -0.3811831 | -1.0624069 | 0.30004068 | 0.0228733 | -1.1917455 | 0.25171469 | 0.99999785 | -5.6840491 | down |
| KI - WT, cell_origin: iPSC | O14773 | TPP1 | 3 | Homo sapien | -0.6365886 | -1.7777365 | 0.50455942 | -0.6218433 | -1.1888286 | 0.25295292 | 0.99999785 | -5.6872458 | down |
| KI - WT, cell_origin: iPSC | Q9Y3A5 | SBDS | 5 | Homo sapien | 0.83959988 | -0.6661718 | 2.34537152 | -2.1383352 | 1.18827133 | 0.25316577 | 0.99999785 | -5.6878719 | up |
| KI - WT, cell_origin: iPSC | Q5SRE5 | NUP188 | 3 | Homo sapien | -0.3097213 | -0.86505 | 0.24560748 | -1.3118567 | -1.1878472 | 0.25320192 | 0.99999785 | -5.6884338 | down |
| KI - WT, cell_origin: iPSC | P49796 | RGS3 | 8 | Homo sapien | 0.47862599 | -0.381116 | 1.33836793 | -0.9294205 | 1.18586421 | 0.25399315 | 0.99999785 | -5.6906376 | up |
| KI - WT, cell_origin: iPSC | Q92896 | GLG1 | 13 | Homo sapien | 0.43538407 | -0.3481565 | 1.21892465 | -0.9051393 | 1.18346849 | 0.25488327 | 0.99999785 | -5.6933426 | up |
| KI - WT, cell_origin: iPSC | Q03135 | CAV1 | 5 | Homo sapien | 0.6098558 | -0.4892818 | 1.70899345 | 0.02313336 | 1.18243545 | 0.25540318 | 0.99999785 | -5.6944145 | up |
| KI - WT, cell_origin: iPSC | P23508 | MCC | 6 | Homo sapien | -0.45076 | -1.2642189 | 0.36269887 | -1.208267 | -1.1801833 | 0.25614533 | 0.99999785 | -5.6970192 | down |
| KI - WT, cell_origin: iPSC | Q96JM2 | ZNF462 | 10 | Homo sapien | -0.7225641 | -2.0278086 | 0.58268045 | -1.9118019 | -1.1797415 | 0.25644111 | 0.99999785 | -5.6974257 | down |
| KI - WT, cell_origin: iPSC | Q53H96 | PYCR3 | 4 | Homo sapien | -0.3781596 | -1.0644545 | 0.30813532 | -0.8573363 | -1.1735565 | 0.25871146 | 0.99999785 | -5.7044058 | down |
| KI - WT, cell_origin: iPSC | P09132 | SRP19 | 4 | Homo sapien | 0.33659036 | -0.2745398 | 0.94772055 | 0.34592569 | 1.17302595 | 0.25891777 | 0.99999785 | -5.7049957 | up |
| KI - WT, cell_origin: iPSC | Q9C0D7 | ZC3H12C | 4 | Homo sapien | -1.1848858 | -3.3381146 | 0.96834306 | -2.7310737 | -1.1727047 | 0.25916748 | 0.99999785 | -5.7052645 | down |
| KI - WT, cell_origin: iPSC | O75376 | NCOR1 | 3 | Homo sapien | 1.02511283 | -0.8397207 | 2.88994639 | -3.4334762 | 1.1713309 | 0.25967654 | 0.99999785 | -5.7068086 | up |
| KI - WT, cell_origin: iPSC | P07099 | EPHX1 | 3 | Homo sapien | 2.12762638 | -1.7445598 | 5.99981255 | -5.0750466 | 1.1709585 | 0.25984745 | 0.99999785 | -5.7072037 | up |
| KI - WT, cell_origin: iPSC | P53999 | SUB1 | 5 | Homo sapien | -1.0827486 | -3.0568708 | 0.89137353 | 4.89588208 | -1.1688425 | 0.26067326 | 0.99999785 | -5.7095503 | down |
| KI - WT, cell_origin: iPSC | Q9Y6M1 | IGF2BP2 | 19 | Homo sapien | -0.4604993 | -1.3002303 | 0.37923167 | 3.70289429 | -1.1686681 | 0.26074141 | 0.99999785 | -5.7097436 | down |
| KI - WT, cell_origin: iPSC | Q9NQC3 | RTN4 | 6 | Homo sapien | 0.80320616 | -0.6626198 | 2.2690321 | 0.91492076 | 1.16774226 | 0.26110342 | 0.99999785 | -5.7107691 | up |
| KI - WT, cell_origin: iPSC | Q5VWQ8 | DAB2IP | 2 | Homo sapien | 0.46003146 | -0.3792417 | 1.29930467 | -1.1392287 | 1.16749508 | 0.2610906 | 0.99999785 | -5.7111215 | up |
| KI - WT, cell_origin: iPSC | Q9BRX8 | PRXL2A | 5 | Homo sapien | 1.12830714 | -0.9322652 | 3.18887949 | -1.8832938 | 1.16692203 | 0.26142446 | 0.99999785 | -5.7116771 | up |
| KI - WT, cell_origin: iPSC | O15211 | RGL2 | 23 | Homo sapien | 1.0103317 | -0.8349007 | 2.85556416 | 0.67279973 | 1.16685063 | 0.26145242 | 0.99999785 | -5.7117561 | up |
| KI - WT, cell_origin: iPSC | P61513 | RPL37A | 5 | Homo sapien | -0.3622208 | -1.0234245 | 0.29898283 | 1.89952439 | -1.1667501 | 0.26136755 | 0.99999785 | -5.7119568 | down |
| KI - WT, cell_origin: iPSC | O43242 | PSMD3 | 8 | Homo sapien | 0.40970007 | -0.3383653 | 1.15776545 | -1.2225268 | 1.1664501 | 0.2614851 | 0.99999785 | -5.7122888 | up |
| KI - WT, cell_origin: iPSC | P56945 | BCAR1 | 6 | Homo sapien | 0.83617714 | -0.6919934 | 2.36434767 | -1.0473213 | 1.16608132 | 0.26175384 | 0.99999785 | -5.7126072 | up |
| KI - WT, cell_origin: iPSC | Q6UB35 | MTHFD1L | 4 | Homo sapien | -2.1049314 | -5.9556226 | 1.7457598 | -4.879851 | -1.1649348 | 0.26220352 | 0.99999785 | -5.7138747 | down |
| KI - WT, cell_origin: iPSC | Q96Q89 | KIF20B | 3 | Homo sapien | 0.35553014 | -0.2950874 | 1.00614765 | -1.9055599 | 1.16383216 | 0.26251257 | 0.99999785 | -5.7151828 | up |
| KI - WT, cell_origin: iPSC | O15155 | BET1 | 2 | Homo sapien | -1.5764426 | -4.4647514 | 1.31186621 | -1.9340616 | -1.1631532 | 0.2629035 | 0.99999785 | -5.7158424 | down |
| KI - WT, cell_origin: iPSC | Q9Y2X0 | MED16 | 6 | Homo sapien | 0.35807815 | -0.2996612 | 1.01581751 | -0.8015153 | 1.1594811 | 0.26422703 | 0.99999785 | -5.7199807 | up |
| KI - WT, cell_origin: iPSC | Q8NEW0 | SLC30A7 | 4 | Homo sapien | 0.32091478 | -0.2686628 | 0.91049233 | 0.95101914 | 1.15928034 | 0.26430634 | 0.99999785 | -5.7202017 | up |
| KI - WT, cell_origin: iPSC | Q9BZI7 | UPF3B | 8 | Homo sapien | -0.4117178 | -1.1703659 | 0.34693033 | -1.0035465 | -1.1558432 | 0.265667 | 0.99999785 | -5.7239807 | down |
| KI - WT, cell_origin: iPSC | Q9BRK4 | LZTS2 | 11 | Homo sapien | 0.81501232 | -0.6880171 | 2.31804178 | -1.4484842 | 1.15557743 | 0.26589572 | 0.99999785 | -5.7241811 | up |
| KI - WT, cell_origin: iPSC | P31949 | S100A11 | 4 | Homo sapien | -0.6084984 | -1.732733 | 0.51573619 | 0.3746429 | -1.1534662 | 0.26673419 | 0.99999785 | -5.7264969 | down |
| KI - WT, cell_origin: iPSC | Q9Y2A7 | NCKAP1 | 33 | Homo sapien | 0.39667099 | -0.3363338 | 1.12967575 | 1.50399643 | 1.15264223 | 0.26695355 | 0.99999785 | -5.7274807 | up |
| KI - WT, cell_origin: iPSC | Q07617 | SPAG1 | 6 | Homo sapien | 0.95866736 | -0.8160826 | 2.73341733 | -2.7065173 | 1.15115322 | 0.26765509 | 0.99999785 | -5.72903 | up |
| KI - WT, cell_origin: iPSC | O00231 | PSMD11 | 13 | Homo sapien | 0.5567934 | -0.4747507 | 1.58833747 | -1.2460406 | 1.15010737 | 0.26803942 | 0.99999785 | -5.7301986 | up |
| KI - WT, cell_origin: iPSC | Q9NZ63 | C9orf78 | 2 | Homo sapien | -0.4918377 | -1.4043267 | 0.42065122 | -1.4313011 | -1.1480952 | 0.2687742 | 0.99999785 | -5.7324495 | down |
| KI - WT, cell_origin: iPSC | P55899 | FCGRT | 3 | Homo sapien | -2.1949022 | -6.2890566 | 1.89925222 | -2.351773 | -1.1424924 | 0.27112472 | 0.99999785 | -5.7384772 | down |
| KI - WT, cell_origin: iPSC | Q5TAX3 | TUT4 | 7 | Homo sapien | 0.39945873 | -0.3484749 | 1.14739239 | -1.471111 | 1.13749244 | 0.27302144 | 0.99999785 | -5.7439981 | up |
| KI - WT, cell_origin: iPSC | P29597 | TYK2 | 5 | Homo sapien | 0.76008806 | -0.664688 | 2.18486408 | -2.0955904 | 1.13689323 | 0.27338582 | 0.99999785 | -5.744553 | up |
| KI - WT, cell_origin: iPSC | O60784 | TOM1 | 3 | Homo sapien | 0.86315036 | -0.7548885 | 2.48118926 | -2.2189236 | 1.13684136 | 0.27340684 | 0.99999785 | -5.7446092 | up |
| KI - WT, cell_origin: iPSC | Q96SZ5 | ADO | 3 | Homo sapien | -0.3740697 | -1.0751675 | 0.32702805 | -1.070878 | -1.136354 | 0.27348269 | 0.99999785 | -5.7452311 | down |
| KI - WT, cell_origin: iPSC | Q9H2M9 | RAB3GAP2 | 21 | Homo sapien | 0.5696121 | -0.501253 | 1.64047716 | 0.07749693 | 1.13356604 | 0.27473615 | 0.99999785 | -5.7481516 | up |

| | | | | | | | | | | | | | |
|---|---|---|---|---|---|---|---|---|---|---|---|---|---|
| KI - WT, cell_origin: iPSC | Q9Y6I4 | USP3 | 4 | Homo sapien | -1.641825 | -4.7298913 | 1.44624132 | -3.140396 | -1.1330331 | 0.27495289 | 0.99999785 | -5.7487271 | down |
| KI - WT, cell_origin: iPSC | O15014 | ZNF609 | 2 | Homo sapien | -0.5248948 | -1.5121735 | 0.46238386 | -0.9529807 | -1.1328477 | 0.27499908 | 0.99999785 | -5.7489502 | down |
| KI - WT, cell_origin: iPSC | P52564 | MAP2K6 | 4 | Homo sapien | -0.5274773 | -1.5196406 | 0.46468599 | -1.8087459 | -1.1324116 | 0.2751049 | 0.99999785 | -5.7494771 | down |
| KI - WT, cell_origin: iPSC | P35237 | SERPINB6 | 9 | Homo sapien | -0.3124063 | -0.9016545 | 0.27684191 | 1.27956054 | -1.1291748 | 0.27640502 | 0.99999785 | -5.7529829 | down |
| KI - WT, cell_origin: iPSC | Q92890 | UFD1 | 3 | Homo sapien | -0.5421184 | -1.5707027 | 0.48646597 | -0.3700526 | -1.1231987 | 0.2789759 | 0.99999785 | -5.7593079 | down |
| KI - WT, cell_origin: iPSC | P17900 | GM2A | 2 | Homo sapien | -0.5195422 | -1.5060315 | 0.46694717 | -0.2739899 | -1.1223564 | 0.2793225 | 0.99999785 | -5.7602106 | down |
| KI - WT, cell_origin: iPSC | P48556 | PSMD8 | 4 | Homo sapien | 0.44316343 | -0.3981037 | 1.2844306 | -1.1011218 | 1.12205376 | 0.27934692 | 0.99999785 | -5.7606148 | up |
| KI - WT, cell_origin: iPSC | P07477 | PRSS1 | 2 | Homo sapien | 0.45132971 | -0.4069987 | 1.30965815 | 1.42559448 | 1.12027044 | 0.28012739 | 0.99999785 | -5.7624875 | up |
| KI - WT, cell_origin: iPSC | P46379 | BAG6 | 21 | Homo sapien | 0.86732978 | -0.7866658 | 2.52132532 | 0.48089788 | 1.11751224 | 0.2813221 | 0.99999785 | -5.7653905 | up |
| KI - WT, cell_origin: iPSC | Q6NYC8 | PPP1R18 | 2 | Homo sapien | 0.96927244 | -0.8802265 | 2.81877142 | -0.2315322 | 1.11684816 | 0.28159705 | 0.99999785 | -5.7660992 | up |
| KI - WT, cell_origin: iPSC | P63218 | GNG5 | 3 | Homo sapien | 0.62732335 | -0.5769009 | 1.83154758 | 2.56023184 | 1.11016227 | 0.2843765 | 0.99999785 | -5.773214 | up |
| KI - WT, cell_origin: iPSC | Q9HDC9 | APMAP | 4 | Homo sapien | 0.48373445 | -0.445246 | 1.41271485 | 0.20573968 | 1.10961154 | 0.2845919 | 0.99999785 | -5.7738103 | up |
| KI - WT, cell_origin: iPSC | Q9BQG0 | MYBBP1A | 23 | Homo sapien | 0.55828088 | -0.516429 | 1.63299072 | -1.7694178 | 1.10657136 | 0.28579422 | 0.99999785 | -5.7770893 | up |
| KI - WT, cell_origin: iPSC | Q9NPA3 | MID1IP1 | 3 | Homo sapien | 0.44077097 | -0.4099709 | 1.29151282 | -0.7682635 | 1.10365439 | 0.28701823 | 0.99999785 | -5.7801739 | up |
| KI - WT, cell_origin: iPSC | Q8N6T3 | ARFGAP1 | 10 | Homo sapien | 0.34369262 | -0.3199397 | 1.00732494 | 0.31698237 | 1.10301738 | 0.28725072 | 0.99999785 | -5.7808759 | up |
| KI - WT, cell_origin: iPSC | Q2VPB7 | AP5B1 | 4 | Homo sapien | 0.61343723 | -0.5742408 | 1.8011153 | -1.8922529 | 1.10046965 | 0.28839904 | 0.99999785 | -5.7835004 | up |
| KI - WT, cell_origin: iPSC | Q03113 | GNA12 | 7 | Homo sapien | 0.73632889 | -0.6963722 | 2.16903003 | -1.1050705 | 1.09526347 | 0.29064367 | 0.99999785 | -5.7889387 | up |
| KI - WT, cell_origin: iPSC | P56211 | ARPP19 | 2 | Homo sapien | 1.45847106 | -1.3797379 | 4.29668003 | -2.4028135 | 1.09510513 | 0.29071082 | 0.99999785 | -5.7891049 | up |
| KI - WT, cell_origin: iPSC | O75962 | TRIO | 29 | Homo sapien | 0.35471298 | -0.3355356 | 1.04496154 | -0.621737 | 1.09448861 | 0.2908546 | 0.99999785 | -5.7898511 | up |
| KI - WT, cell_origin: iPSC | Q9Y490 | TLN1 | 44 | Homo sapien | 0.45611347 | -0.4323378 | 1.34456474 | 0.13919309 | 1.09382538 | 0.29121202 | 0.99999785 | -5.7904825 | up |
| KI - WT, cell_origin: iPSC | Q13501 | SQSTM1 | 2 | Homo sapien | 0.96434281 | -0.9161118 | 2.84479746 | -1.2915037 | 1.09287616 | 0.29165735 | 0.99999785 | -5.7914417 | up |
| KI - WT, cell_origin: iPSC | Q8WWN8 | ARAP3 | 15 | Homo sapien | 0.35521383 | -0.3372259 | 1.04765358 | -0.7022704 | 1.09256568 | 0.29167176 | 0.99999785 | -5.7918665 | up |
| KI - WT, cell_origin: iPSC | Q3KQU3 | MAP7D1 | 10 | Homo sapien | 0.30561012 | -0.2909197 | 0.90213991 | 0.41578113 | 1.09112694 | 0.29228427 | 0.99999785 | -5.7933724 | up |
| KI - WT, cell_origin: iPSC | Q15024 | EXOSC7 | 4 | Homo sapien | 0.40901685 | -0.3893925 | 1.20742617 | -1.2057828 | 1.09107673 | 0.29230566 | 0.99999785 | -5.793425 | up |
| KI - WT, cell_origin: iPSC | Q96HC4 | PDLIM5 | 21 | Homo sapien | -0.3073834 | -0.907863 | 0.29309622 | 0.06198299 | -1.0902392 | 0.29266266 | 0.99999785 | -5.7943008 | down |
| KI - WT, cell_origin: iPSC | Q9P206 | KIAA1522 | 5 | Homo sapien | 0.86382162 | -0.82583 | 2.5534732 | -1.2398174 | 1.08950513 | 0.29309317 | 0.99999785 | -5.7949681 | up |
| KI - WT, cell_origin: iPSC | Q9Y5A6 | ZSCAN21 | 2 | Homo sapien | -0.6242188 | -1.845008 | 0.59657044 | -3.4623114 | -1.0890199 | 0.29318299 | 0.99999785 | -5.7955748 | down |
| KI - WT, cell_origin: iPSC | P13984 | GTF2F2 | 10 | Homo sapien | -0.7707255 | -2.2790916 | 0.73764055 | -0.4759672 | -1.0889185 | 0.29334359 | 0.99999785 | -5.7955808 | down |
| KI - WT, cell_origin: iPSC | Q5D1E8 | ZC3H12A | 2 | Homo sapien | 1.8719544 | -1.7916891 | 5.53559792 | -5.7385954 | 1.08889104 | 0.2933553 | 0.99999785 | -5.7956095 | up |
| KI - WT, cell_origin: iPSC | P49815 | TSC2 | 4 | Homo sapien | 0.41596749 | -0.3977168 | 1.2296518 | -1.656274 | 1.08878751 | 0.29328225 | 0.99999785 | -5.7958176 | up |
| KI - WT, cell_origin: iPSC | P02538 | KRT6A | 2 | Homo sapien | -0.9967995 | -2.9497296 | 0.9561305 | -0.7981029 | -1.087736 | 0.29384879 | 0.99999785 | -5.796815 | down |
| KI - WT, cell_origin: iPSC | Q9ULR3 | PPM1H | 13 | Homo sapien | 0.58040084 | -0.5582231 | 1.71902481 | 0.60741528 | 1.08630075 | 0.29446286 | 0.99999785 | -5.7983116 | up |
| KI - WT, cell_origin: iPSC | O75436 | VPS26A | 2 | Homo sapien | 0.71657605 | -0.6895935 | 2.12274563 | -0.6249807 | 1.08599283 | 0.29459473 | 0.99999785 | -5.7986324 | up |
| KI - WT, cell_origin: iPSC | Q13614 | MTMR2 | 4 | Homo sapien | 0.27992296 | -0.2690533 | 0.82889917 | -1.0211812 | 1.08598711 | 0.29448019 | 0.99999785 | -5.7987386 | up |
| KI - WT, cell_origin: iPSC | Q6GYQ0 | RALGAPA1 | 4 | Homo sapien | 0.55912025 | -0.5381379 | 1.65637842 | -3.280715 | 1.08526571 | 0.29478937 | 0.99999785 | -5.79949 | up |
| KI - WT, cell_origin: iPSC | A1A4S6 | ARHGAP10 | 4 | Homo sapien | 1.21275972 | -1.1702396 | 3.59575908 | 0.55047484 | 1.08455905 | 0.29520932 | 0.99999785 | -5.8001254 | up |
| KI - WT, cell_origin: iPSC | Q8WX92 | NELFB | 4 | Homo sapien | 0.37338768 | -0.3605381 | 1.10731347 | -0.9543517 | 1.08354643 | 0.2955272 | 0.99999785 | -5.8012792 | up |
| KI - WT, cell_origin: iPSC | P30086 | PEBP1 | 5 | Homo sapien | -0.8106996 | -2.4066446 | 0.7852454 | -1.7260069 | -1.0825411 | 0.2960759 | 0.99999785 | -5.8022237 | down |
| KI - WT, cell_origin: iPSC | Q96BH1 | RNF25 | 4 | Homo sapien | 0.709926 | -0.6897646 | 2.10961662 | -1.2864883 | 1.08089473 | 0.29678433 | 0.99999785 | -5.8039333 | up |
| KI - WT, cell_origin: iPSC | O43765 | SGTA | 8 | Homo sapien | 0.38424643 | -0.3734838 | 1.14197669 | -0.1128059 | 1.08010529 | 0.29702187 | 0.99999785 | -5.804841 | up |
| KI - WT, cell_origin: iPSC | O95248 | SBF1 | 31 | Homo sapien | 0.26693935 | -0.2597271 | 0.79360581 | 0.92692494 | 1.07948502 | 0.29727555 | 0.99999785 | -5.8054961 | up |
| KI - WT, cell_origin: iPSC | Q15003 | NCAPH | 2 | Homo sapien | 0.55461452 | -0.5400467 | 1.64927573 | -1.7142718 | 1.0793435 | 0.29738466 | 0.99999785 | -5.8056012 | up |
| KI - WT, cell_origin: iPSC | P11940 | PABPC1 | 33 | Homo sapien | -0.3443139 | -1.0245873 | 0.33595951 | 5.64923886 | -1.0785531 | 0.29777994 | 0.99999785 | -5.8063731 | down |
| KI - WT, cell_origin: iPSC | Q86XA9 | HEATR5A | 3 | Homo sapien | 0.89978746 | -0.8784891 | 2.67806402 | -1.1834025 | 1.07830843 | 0.29789971 | 0.99999785 | -5.8066143 | up |
| KI - WT, cell_origin: iPSC | Q86XN8 | MEX3D | 2 | Homo sapien | 0.46871752 | -0.4580937 | 1.39552871 | -1.1045457 | 1.07730288 | 0.29825264 | 0.99999785 | -5.8077262 | up |
| KI - WT, cell_origin: iPSC | P20292 | ALOX5AP | 2 | Homo sapien | 0.73451249 | -0.7207555 | 2.18978053 | -0.7918819 | 1.07561925 | 0.29906272 | 0.99999785 | -5.8093962 | up |

| | | | | | | | | | | | | | |
|---|---|---|---|---|---|---|---|---|---|---|---|---|---|
| KI - WT, cell_origin: iPSC | Q8NFD5 | ARID1B | 4 | Homo sapien | -0.4932059 | -1.4740409 | 0.48762913 | -2.1533835 | -1.0709569 | 0.30097148 | 0.99999785 | -5.8143068 | down |
| KI - WT, cell_origin: iPSC | P00488 | F13A1 | 9 | Homo sapien | 1.13783294 | -1.1365608 | 3.41222667 | -0.767026 | 1.06614241 | 0.30318787 | 0.99999785 | -5.8191525 | up |
| KI - WT, cell_origin: iPSC | Q9Y6W5 | WASF2 | 9 | Homo sapien | 0.3200785 | -0.3202839 | 0.96044094 | 0.7665304 | 1.06456045 | 0.30376561 | 0.99999785 | -5.8208759 | up |
| KI - WT, cell_origin: iPSC | Q96LI5 | CNOT6L | 8 | Homo sapien | -0.28934 | -0.8699345 | 0.29125448 | -0.2898978 | -1.0613907 | 0.30515726 | 0.99999785 | -5.8241187 | down |
| KI - WT, cell_origin: iPSC | Q96RT8 | TUBGCP5 | 4 | Homo sapien | 0.31086584 | -0.3144434 | 0.93617509 | -1.2596082 | 1.05880951 | 0.30629395 | 0.99999785 | -5.8267533 | up |
| KI - WT, cell_origin: iPSC | P24666 | ACP1 | 2 | Homo sapien | -0.4023279 | -1.2143398 | 0.40968398 | -0.9016712 | -1.055362 | 0.30783604 | 0.99999785 | -5.8302463 | down |
| KI - WT, cell_origin: iPSC | P04275 | VWF | 30 | Homo sapien | -1.1457739 | -3.4598159 | 1.16826801 | -2.6243615 | -1.0551886 | 0.30800769 | 0.99999785 | -5.8303373 | down |
| KI - WT, cell_origin: iPSC | Q6IA86 | ELP2 | 7 | Homo sapien | 0.36974653 | -0.3780327 | 1.11752576 | -1.8405214 | 1.05310191 | 0.30881838 | 0.99999785 | -5.8325594 | up |
| KI - WT, cell_origin: iPSC | Q96PY5 | FMNL2 | 36 | Homo sapien | 0.62243968 | -0.6372075 | 1.88208689 | 0.64180894 | 1.05305416 | 0.30895333 | 0.99999785 | -5.8325052 | up |
| KI - WT, cell_origin: iPSC | P12277 | CKB | 7 | Homo sapien | -1.726408 | -5.2262798 | 1.77346386 | -2.828625 | -1.0512201 | 0.30976758 | 0.99999785 | -5.834365 | down |
| KI - WT, cell_origin: iPSC | Q5T1V6 | DDX59 | 8 | Homo sapien | 0.64594199 | -0.664723 | 1.956607 | -1.6629686 | 1.0502778 | 0.31018655 | 0.99999785 | -5.8353194 | up |
| KI - WT, cell_origin: iPSC | Q96QR8 | PURB | 5 | Homo sapien | -0.3707483 | -1.122777 | 0.38128039 | 0.40409221 | -1.0500947 | 0.31017355 | 0.99999785 | -5.8355905 | down |
| KI - WT, cell_origin: iPSC | Q02218 | OGDH | 8 | Homo sapien | -0.8199939 | -2.4874515 | 0.84746375 | -2.1615247 | -1.0479925 | 0.31120433 | 0.99999785 | -5.8376311 | down |
| KI - WT, cell_origin: iPSC | P41567 | EIF1 | 4 | Homo sapien | -0.5354986 | -1.6260526 | 0.55505533 | 0.71128918 | -1.0464382 | 0.31189795 | 0.99999785 | -5.839201 | down |
| KI - WT, cell_origin: iPSC | Q13492 | PICALM | 3 | Homo sapien | 0.37569834 | -0.3895402 | 1.14093687 | -1.9873342 | 1.04563986 | 0.3121416 | 0.99999785 | -5.8401095 | up |
| KI - WT, cell_origin: iPSC | Q6IE81 | JADE1 | 2 | Homo sapien | 0.50251175 | -0.5229164 | 1.52793985 | -1.2777174 | 1.04410641 | 0.31289824 | 0.99999785 | -5.8415909 | up |
| KI - WT, cell_origin: iPSC | Q92614 | MYO18A | 8 | Homo sapien | -0.472845 | -1.4394869 | 0.49379686 | -1.0312184 | -1.0421999 | 0.31375014 | 0.99999785 | -5.8435123 | down |
| KI - WT, cell_origin: iPSC | Q9BT78 | COPS4 | 8 | Homo sapien | 0.30528803 | -0.3188826 | 0.92945863 | -1.3454143 | 1.04170835 | 0.3139029 | 0.99999785 | -5.8440689 | up |
| KI - WT, cell_origin: iPSC | Q86SQ0 | PHLDB2 | 9 | Homo sapien | 0.79511055 | -0.8400724 | 2.43029345 | -1.7759271 | 1.03624762 | 0.31647329 | 0.99999785 | -5.8494433 | up |
| KI - WT, cell_origin: iPSC | Q13459 | MYO9B | 16 | Homo sapien | -0.2937679 | -0.8976583 | 0.31012246 | -0.1651632 | -1.0360625 | 0.31644483 | 0.99999785 | -5.8497322 | down |
| KI - WT, cell_origin: iPSC | Q08209 | PPP3CA | 6 | Homo sapien | 0.44993464 | -0.4754246 | 1.37529388 | -0.5545268 | 1.0360272 | 0.31654282 | 0.99999785 | -5.8496916 | up |
| KI - WT, cell_origin: iPSC | P51511 | MMP15 | 5 | Homo sapien | 0.67604681 | -0.7146782 | 2.06677183 | -1.4288276 | 1.03594778 | 0.31660865 | 0.99999785 | -5.8497434 | up |
| KI - WT, cell_origin: iPSC | Q5VZ18 | SHE | 13 | Homo sapien | 0.41286667 | -0.4364224 | 1.26215578 | 0.71472007 | 1.03570657 | 0.31666644 | 0.99999785 | -5.850032 | up |
| KI - WT, cell_origin: iPSC | Q86YV5 | PRAG1 | 14 | Homo sapien | 0.64568548 | -0.6829419 | 1.97431291 | -2.3298245 | 1.03538118 | 0.31681343 | 0.99999785 | -5.8503574 | up |
| KI - WT, cell_origin: iPSC | P69905 | HBA1 | 5 | Homo sapien | -1.2997383 | -3.9794905 | 1.380014 | -1.4896365 | -1.0336265 | 0.31765794 | 0.99999785 | -5.8520638 | down |
| KI - WT, cell_origin: iPSC | Q9Y4E6 | WDR7 | 12 | Homo sapien | 0.30780348 | -0.3284253 | 0.94403226 | 0.14497039 | 1.03038585 | 0.31901554 | 0.99999785 | -5.8553995 | up |
| KI - WT, cell_origin: iPSC | Q86UU1 | PHLDB1 | 8 | Homo sapien | 1.34871968 | -1.4428367 | 4.140276 | -2.1394402 | 1.02962165 | 0.31947415 | 0.99999785 | -5.8560566 | up |
| KI - WT, cell_origin: iPSC | Q96AJ1 | CLUAP1 | 4 | Homo sapien | -0.4301071 | -1.3209083 | 0.46069402 | -1.6480672 | -1.0283372 | 0.31994696 | 0.99999785 | -5.8574381 | down |
| KI - WT, cell_origin: iPSC | Q14019 | COTL1 | 3 | Homo sapien | -0.6245116 | -1.9188098 | 0.66978669 | -0.7806814 | -1.0282731 | 0.3200874 | 0.99999785 | -5.857398 | down |
| KI - WT, cell_origin: iPSC | P62714 | PPP2CB | 2 | Homo sapien | -0.5182074 | -1.5940744 | 0.55765957 | -1.5520297 | -1.0261893 | 0.32098599 | 0.99999785 | -5.8595153 | down |
| KI - WT, cell_origin: iPSC | Q00577 | PURA | 5 | Homo sapien | 0.70377081 | -0.7595251 | 2.16706671 | -0.3784569 | 1.02494712 | 0.32160353 | 0.99999785 | -5.8607001 | up |
| KI - WT, cell_origin: iPSC | Q9HBH0 | RHOF | 2 | Homo sapien | 0.38240688 | -0.4122547 | 1.17706841 | -1.9564409 | 1.02490399 | 0.32151232 | 0.99999785 | -5.8608468 | up |
| KI - WT, cell_origin: iPSC | Q12980 | NPRL3 | 3 | Homo sapien | 0.349201 | -0.376905 | 1.075307 | -1.3023214 | 1.02427159 | 0.32180125 | 0.99999785 | -5.8614736 | up |
| KI - WT, cell_origin: iPSC | Q14161 | GIT2 | 28 | Homo sapien | 0.8027038 | -0.8683615 | 2.4737691 | 1.11233644 | 1.02368011 | 0.32218244 | 0.99999785 | -5.8619555 | up |
| KI - WT, cell_origin: iPSC | Q9Y3D0 | CIAO2B | 3 | Homo sapien | 0.41788149 | -0.4519058 | 1.28766881 | -1.8031819 | 1.02324527 | 0.32227056 | 0.99999785 | -5.8624901 | up |
| KI - WT, cell_origin: iPSC | P06576 | ATP5F1B | 12 | Homo sapien | 1.5346496 | -1.6615667 | 4.73086593 | -1.4101447 | 1.02323511 | 0.32238594 | 0.99999785 | -5.8623961 | up |
| KI - WT, cell_origin: iPSC | O15446 | POLR1G | 2 | Homo sapien | 0.8295517 | -0.8988919 | 2.55799533 | -2.0691342 | 1.02279973 | 0.32258514 | 0.99999785 | -5.8628271 | up |
| KI - WT, cell_origin: iPSC | P61927 | RPL37 | 4 | Homo sapien | -0.7172029 | -2.2120722 | 0.77766638 | 0.60470712 | -1.0224479 | 0.32274616 | 0.99999785 | -5.8631752 | down |
| KI - WT, cell_origin: iPSC | Q4G0J3 | LARP7 | 6 | Homo sapien | 0.37064859 | -0.4025878 | 1.14388493 | -0.5588277 | 1.02098878 | 0.32331726 | 0.99999785 | -5.8647096 | up |
| KI - WT, cell_origin: iPSC | Q8NEG4 | FAM83F | 4 | Homo sapien | 0.66882216 | -0.7294449 | 2.0670892 | -0.4021317 | 1.019349 | 0.32416704 | 0.99999785 | -5.8662369 | up |
| KI - WT, cell_origin: iPSC | Q96R06 | SPAG5 | 2 | Homo sapien | 0.82709337 | -0.9021989 | 2.55638562 | -1.477452 | 1.01926828 | 0.32420411 | 0.99999785 | -5.8663166 | up |
| KI - WT, cell_origin: iPSC | Q96QH2 | PRAM1 | 3 | Homo sapien | 0.36049641 | -0.3929221 | 1.11391487 | 1.92300354 | 1.01920221 | 0.32414766 | 0.99999785 | -5.8664637 | up |
| KI - WT, cell_origin: iPSC | O43741 | PRKAB2 | 2 | Homo sapien | 0.7589037 | -0.8288189 | 2.34662629 | -3.2918439 | 1.01819263 | 0.32462101 | 0.99999785 | -5.8674506 | up |
| KI - WT, cell_origin: iPSC | Q9UHD2 | TBK1 | 11 | Homo sapien | 0.31973208 | -0.349152 | 0.98861612 | -0.4228992 | 1.01806397 | 0.32464735 | 0.99999785 | -5.8676084 | up |
| KI - WT, cell_origin: iPSC | Q9BR76 | CORO1B | 2 | Homo sapien | -0.5633659 | -1.7427729 | 0.6160411 | -1.6114309 | -1.0179565 | 0.32480696 | 0.99999785 | -5.8676101 | down |
| KI - WT, cell_origin: iPSC | O75663 | TIPRL | 5 | Homo sapien | 0.49692637 | -0.5431937 | 1.53704645 | -1.3230634 | 1.01792373 | 0.32478191 | 0.99999785 | -5.8676804 | up |

| | | | | | | | | | | | | | |
|---|---|---|---|---|---|---|---|---|---|---|---|---|---|
| KI - WT, cell_origin: iPSC | Q9H792 | PEAK1 | 4 | Homo sapien | 0.30577972 | -0.3372236 | 0.94878306 | -1.8857795 | 1.01282668 | 0.32706257 | 0.99999785 | -5.8727591 | up |
| KI - WT, cell_origin: iPSC | Q92888 | ARHGEF1 | 17 | Homo sapien | 0.53299865 | -0.5892234 | 1.65522066 | 0.12681895 | 1.01216123 | 0.32747995 | 0.99999785 | -5.8733075 | up |
| KI - WT, cell_origin: iPSC | P49419 | ALDH7A1 | 22 | Homo sapien | 0.41962878 | -0.464009 | 1.30326657 | 1.54746164 | 1.01186618 | 0.32758715 | 0.99999785 | -5.8736247 | up |
| KI - WT, cell_origin: iPSC | Q13418 | ILK | 13 | Homo sapien | 0.36586968 | -0.4044382 | 1.13617759 | -0.6760139 | 1.01164019 | 0.32762166 | 0.99999785 | -5.8739131 | up |
| KI - WT, cell_origin: iPSC | O43399 | TPD52L2 | 7 | Homo sapien | 0.63089334 | -0.6992348 | 1.96102151 | -0.6094289 | 1.01079903 | 0.32811052 | 0.99999785 | -5.8746426 | up |
| KI - WT, cell_origin: iPSC | Q9P2D1 | CHD7 | 14 | Homo sapien | 0.21857579 | -0.2419925 | 0.67914409 | -0.7349547 | 1.01075915 | 0.32801956 | 0.99999785 | -5.8747861 | up |
| KI - WT, cell_origin: iPSC | Q92759 | GTF2H4 | 4 | Homo sapien | 0.39594674 | -0.4389727 | 1.23086618 | -2.3765489 | 1.01002445 | 0.32836011 | 0.99999785 | -5.8755055 | up |
| KI - WT, cell_origin: iPSC | Q9NP61 | ARFGAP3 | 9 | Homo sapien | -1.2619887 | -3.9265753 | 1.40259802 | -1.1179574 | -1.0093179 | 0.32879714 | 0.99999785 | -5.8760925 | down |
| KI - WT, cell_origin: iPSC | Q8IZP0 | ABI1 | 5 | Homo sapien | 0.42019766 | -0.4670307 | 1.30742601 | 0.51881238 | 1.00915349 | 0.32884706 | 0.99999785 | -5.8762785 | up |
| KI - WT, cell_origin: iPSC | Q9NP97 | DYNLRB1 | 2 | Homo sapien | 0.48669278 | -0.5426453 | 1.51603086 | -0.1451971 | 1.00762571 | 0.32958284 | 0.99999785 | -5.8777467 | up |
| KI - WT, cell_origin: iPSC | Q9HD42 | CHMP1A | 5 | Homo sapien | -0.3226745 | -1.0047463 | 0.35939734 | 0.78058164 | -1.0075676 | 0.32950074 | 0.99999785 | -5.8779079 | down |
| KI - WT, cell_origin: iPSC | P62310 | LSM3 | 2 | Homo sapien | 0.23188193 | -0.2583889 | 0.72215274 | 1.05296564 | 1.00732719 | 0.32961251 | 0.99999785 | -5.8781428 | up |
| KI - WT, cell_origin: iPSC | O75569 | PRKRA | 7 | Homo sapien | -0.3120741 | -0.9719217 | 0.34777351 | -0.4309726 | -1.0072882 | 0.32963063 | 0.99999785 | -5.8781808 | down |
| KI - WT, cell_origin: iPSC | Q0PNE2 | ELP6 | 2 | Homo sapien | 0.40039239 | -0.4472945 | 1.24807928 | -0.7255612 | 1.00616278 | 0.33018668 | 0.99999785 | -5.8792483 | up |
| KI - WT, cell_origin: iPSC | P01137 | TGFB1 | 3 | Homo sapien | -1.5728973 | -4.9065678 | 1.7607732 | -3.1775322 | -1.0054953 | 0.3305739 | 0.99999785 | -5.8798259 | down |
| KI - WT, cell_origin: iPSC | O00273 | DFFA | 3 | Homo sapien | -0.5526729 | -1.7246358 | 0.61929007 | -0.4080138 | -1.0049782 | 0.33081481 | 0.99999785 | -5.88033 | down |
| KI - WT, cell_origin: iPSC | Q7L576 | CYFIP1 | 39 | Homo sapien | 0.33919905 | -0.3805166 | 1.05891464 | 2.58981982 | 1.00386019 | 0.33124381 | 0.99999785 | -5.8815078 | up |
| KI - WT, cell_origin: iPSC | Q96D71 | REPS1 | 4 | Homo sapien | 0.46183545 | -0.5182978 | 1.44196867 | -1.3384283 | 1.00373728 | 0.33131704 | 0.99999785 | -5.8816122 | up |
| KI - WT, cell_origin: iPSC | Q7Z628 | NET1 | 2 | Homo sapien | 0.57214135 | -0.6437771 | 1.78805982 | -2.7306685 | 1.00216712 | 0.3320186 | 0.99999785 | -5.8831703 | up |
| KI - WT, cell_origin: iPSC | Q9Y5V0 | ZNF706 | 3 | Homo sapien | -0.698782 | -2.1856861 | 0.78812202 | 3.21484848 | -1.0015235 | 0.33242729 | 0.99999785 | -5.883692 | down |
| KI - WT, cell_origin: iPSC | P17655 | CAPN2 | 18 | Homo sapien | 1.15473989 | -1.3024209 | 3.61190066 | -1.0177722 | 1.00150463 | 0.33243613 | 0.99999785 | -5.8837103 | up |
| KI - WT, cell_origin: iPSC | P18669 | PGAM1 | 6 | Homo sapien | -0.5344995 | -1.6725082 | 0.60350918 | -0.4085396 | -1.0009309 | 0.33270447 | 0.99999785 | -5.8842677 | down |
| KI - WT, cell_origin: iPSC | Q9UBI1 | COMMD3 | 4 | Homo sapien | -0.3773419 | -1.1815674 | 0.42688365 | -0.6323933 | -0.999431 | 0.33332153 | 0.99999785 | -5.8858057 | down |
| KI - WT, cell_origin: iPSC | O43294 | TGFB1I1 | 10 | Homo sapien | 0.21160524 | -0.2403978 | 0.66360827 | 0.46326555 | 0.99706796 | 0.33440722 | 0.99999785 | -5.8881177 | up |
| KI - WT, cell_origin: iPSC | Q12967 | RALGDS | 7 | Homo sapien | -0.6414519 | -2.0127033 | 0.72979947 | -1.4056173 | -0.996895 | 0.33459647 | 0.99999785 | -5.8881805 | down |
| KI - WT, cell_origin: iPSC | Q96NW4 | ANKRD27 | 3 | Homo sapien | 0.74731695 | -0.850555 | 2.34518885 | -3.8261014 | 0.99609911 | 0.33486257 | 0.99999785 | -5.8890551 | up |
| KI - WT, cell_origin: iPSC | Q96E14 | RMI2 | 2 | Homo sapien | 0.4755564 | -0.5419396 | 1.49305237 | -0.2862426 | 0.99602836 | 0.33500371 | 0.99999785 | -5.8890189 | up |
| KI - WT, cell_origin: iPSC | P05455 | SSB | 23 | Homo sapien | -0.1933958 | -0.6069373 | 0.22014566 | 3.2408222 | -0.9960192 | 0.33490016 | 0.99999785 | -5.8891324 | down |
| KI - WT, cell_origin: iPSC | P38117 | ETFB | 3 | Homo sapien | 0.67040834 | -0.7641934 | 2.10501007 | -0.4006426 | 0.99588783 | 0.33506979 | 0.99999785 | -5.8891548 | up |
| KI - WT, cell_origin: iPSC | Q06830 | PRDX1 | 16 | Homo sapien | -0.3296821 | -1.0350631 | 0.37569893 | 3.90195168 | -0.9956105 | 0.33512449 | 0.99999785 | -5.8894963 | down |
| KI - WT, cell_origin: iPSC | Q14191 | WRN | 6 | Homo sapien | 0.58707935 | -0.6724053 | 1.84656398 | -2.124102 | 0.99309184 | 0.33633845 | 0.99999785 | -5.8919015 | up |
| KI - WT, cell_origin: iPSC | O75818 | RPP40 | 3 | Homo sapien | 0.25129402 | -0.2886607 | 0.79124873 | -0.4629172 | 0.99120734 | 0.33716835 | 0.99999785 | -5.8937758 | up |
| KI - WT, cell_origin: iPSC | Q02535 | ID3 | 2 | Homo sapien | 0.41749771 | -0.4800081 | 1.31500354 | -0.9013928 | 0.99091115 | 0.33734029 | 0.99999785 | -5.8940299 | up |
| KI - WT, cell_origin: iPSC | P49720 | PSMB3 | 4 | Homo sapien | 0.38513184 | -0.4437275 | 1.21399117 | -1.4570836 | 0.98961959 | 0.33791916 | 0.99999785 | -5.8953037 | up |
| KI - WT, cell_origin: iPSC | Q8WXF1 | PSPC1 | 17 | Homo sapien | -0.5640369 | -1.7790756 | 0.6510018 | 1.55848041 | -0.9892813 | 0.33818645 | 0.99999785 | -5.8955244 | down |
| KI - WT, cell_origin: iPSC | Q6UW02 | CYP20A1 | 2 | Homo sapien | 0.89158894 | -1.0316148 | 2.81479271 | -2.3239633 | 0.9879654 | 0.33880966 | 0.99999785 | -5.8967886 | up |
| KI - WT, cell_origin: iPSC | Q9H1E3 | NUCKS1 | 3 | Homo sapien | -1.2527729 | -3.9568003 | 1.45125453 | -0.810233 | -0.9873329 | 0.33910949 | 0.99999785 | -5.8973957 | down |
| KI - WT, cell_origin: iPSC | Q8NF37 | LPCAT1 | 6 | Homo sapien | 1.6746269 | -1.9399914 | 5.28924522 | -2.786121 | 0.98732012 | 0.33911557 | 0.99999785 | -5.897408 | up |
| KI - WT, cell_origin: iPSC | P61006 | RAB8A | 12 | Homo sapien | 0.36997873 | -0.4286826 | 1.16864005 | 5.34924013 | 0.98722577 | 0.33916031 | 0.99999785 | -5.8974986 | up |
| KI - WT, cell_origin: iPSC | Q9NZW5 | PALS2 | 15 | Homo sapien | 0.42962736 | -0.4983217 | 1.35757643 | 0.48011405 | 0.98666598 | 0.33942588 | 0.99999785 | -5.8980356 | up |
| KI - WT, cell_origin: iPSC | Q9BU76 | MMTAG2 | 6 | Homo sapien | 0.28022977 | -0.3247966 | 0.88525609 | -1.0810898 | 0.98646046 | 0.33941656 | 0.99999785 | -5.8983374 | up |
| KI - WT, cell_origin: iPSC | Q13425 | SNTB2 | 2 | Homo sapien | 0.86219703 | -1.0038063 | 2.72820031 | -3.197967 | 0.98468311 | 0.34036772 | 0.99999785 | -5.8999357 | up |
| KI - WT, cell_origin: iPSC | P15428 | HPGD | 10 | Homo sapien | -2.6237754 | -8.3091608 | 3.06160998 | -1.8708497 | -0.9834881 | 0.34093623 | 0.99999785 | -5.9010792 | down |
| KI - WT, cell_origin: iPSC | A6NI28 | ARHGAP42 | 4 | Homo sapien | -0.6558505 | -2.0776825 | 0.76598153 | -2.7987112 | -0.9825948 | 0.34128692 | 0.99999785 | -5.9020067 | down |
| KI - WT, cell_origin: iPSC | Q9BXB4 | OSBPL11 | 2 | Homo sapien | 0.61611856 | -0.7234927 | 1.95572983 | 0.77448099 | 0.98013944 | 0.34253289 | 0.99999785 | -5.9042771 | up |
| KI - WT, cell_origin: iPSC | Q14587 | ZNF268 | 5 | Homo sapien | -0.4735283 | -1.5025855 | 0.55552894 | -2.2675909 | -0.9800451 | 0.34247177 | 0.99999785 | -5.9044717 | down |

| | | | | | | | | | | | | |
|---|---|---|---|---|---|---|---|---|---|---|---|---|
| KI - WT, cell_origin: iPSC | P19367 | HK1 | 6 | Homo sapien | 1.67033289 | -1.9638921 | 5.30455787 | -2.8686235 | 0.97947553 | 0.34285007 | 0.99999785 | -5.90491 up |
| KI - WT, cell_origin: iPSC | Q16666 | IFI16 | 25 | Homo sapien | 0.55207886 | -0.6495977 | 1.7537554 | 1.01103847 | 0.97907488 | 0.34304158 | 0.99999785 | -5.9052918 up |
| KI - WT, cell_origin: iPSC | Q9NXW9 | ALKBH4 | 3 | Homo sapien | 0.5926731 | -0.6995615 | 1.88490773 | -1.2345124 | 0.97740871 | 0.34383881 | 0.99999785 | -5.9068778 up |
| KI - WT, cell_origin: iPSC | Q13099 | IFT88 | 9 | Homo sapien | 0.36213457 | -0.4272827 | 1.15155185 | -1.9423778 | 0.97701908 | 0.34391959 | 0.99999785 | -5.907353 up |
| KI - WT, cell_origin: iPSC | A1A5D9 | BICDL2 | 10 | Homo sapien | 0.78623392 | -0.9291091 | 2.50157694 | -2.0859535 | 0.97679441 | 0.34413307 | 0.99999785 | -5.907462 up |
| KI - WT, cell_origin: iPSC | A2RUS2 | DENND3 | 6 | Homo sapien | -0.4228677 | -1.3471263 | 0.5013909 | -1.293978 | -0.9745807 | 0.34511634 | 0.99999785 | -5.9096424 down |
| KI - WT, cell_origin: iPSC | Q9UHR4 | BAIAP2L1 | 4 | Homo sapien | -0.2455425 | -0.7821738 | 0.29108886 | -0.9846586 | -0.9745189 | 0.34511903 | 0.99999785 | -5.9097277 down |
| KI - WT, cell_origin: iPSC | Q8TDN4 | CABLES1 | 2 | Homo sapien | 0.80525565 | -0.9563759 | 2.56688721 | -2.8507259 | 0.97413931 | 0.34540694 | 0.99999785 | -5.9099832 up |
| KI - WT, cell_origin: iPSC | Q9Y244 | POMP | 3 | Homo sapien | 0.69071317 | -0.8205271 | 2.2019534 | -3.031428 | 0.97360351 | 0.34559035 | 0.99999785 | -5.9105647 up |
| KI - WT, cell_origin: iPSC | Q8TA86 | RP9 | 6 | Homo sapien | 0.63049786 | -0.7507337 | 2.01172947 | -0.2566372 | 0.97279085 | 0.34605517 | 0.99999785 | -5.9112613 up |
| KI - WT, cell_origin: iPSC | P13647 | KRT5 | 19 | Homo sapien | -0.57506 | -1.83505 | 0.68492992 | 0.62635398 | -0.9726318 | 0.34613169 | 0.99999785 | -5.911412 down |
| KI - WT, cell_origin: iPSC | Q9GZZ9 | UBA5 | 6 | Homo sapien | 0.67296378 | -0.804029 | 2.14995657 | -0.3881044 | 0.97099206 | 0.34692122 | 0.99999785 | -5.9129639 up |
| KI - WT, cell_origin: iPSC | O15037 | KHNYN | 18 | Homo sapien | 0.29246148 | -0.35099 | 0.935913 | 0.10841359 | 0.96803824 | 0.34824181 | 0.99999785 | -5.9158583 up |
| KI - WT, cell_origin: iPSC | P27448 | MARK3 | 2 | Homo sapien | 0.27075704 | -0.3251682 | 0.86668233 | -0.9734234 | 0.96767079 | 0.34841946 | 0.99999785 | -5.9162048 up |
| KI - WT, cell_origin: iPSC | P27361 | MAPK3 | 8 | Homo sapien | 0.27226957 | -0.3273092 | 0.87184831 | 0.21132846 | 0.96714715 | 0.34867273 | 0.99999785 | -5.9166984 up |
| KI - WT, cell_origin: iPSC | P54652 | HSPA2 | 17 | Homo sapien | 0.30716326 | -0.369749 | 0.98407556 | 5.17022003 | 0.96687192 | 0.34888262 | 0.99999785 | -5.9168812 up |
| KI - WT, cell_origin: iPSC | Q9UPM8 | AP4E1 | 2 | Homo sapien | 0.44677392 | -0.538172 | 1.43171984 | -1.4575066 | 0.96625852 | 0.34913401 | 0.99999785 | -5.9175044 up |
| KI - WT, cell_origin: iPSC | Q9BZZ5 | API5 | 2 | Homo sapien | 0.4016577 | -0.4837446 | 1.28705999 | -1.4572411 | 0.96618683 | 0.34913993 | 0.99999785 | -5.9176007 up |
| KI - WT, cell_origin: iPSC | Q9NX58 | LYAR | 11 | Homo sapien | 0.32663991 | -0.3951096 | 1.04838937 | -0.8184056 | 0.96387886 | 0.35025643 | 0.99999785 | -5.919774 up |
| KI - WT, cell_origin: iPSC | Q5U651 | RASIP1 | 30 | Homo sapien | -1.2542656 | -4.0302448 | 1.52171366 | 1.54837054 | -0.9628877 | 0.35084198 | 0.99999785 | -5.9206003 down |
| KI - WT, cell_origin: iPSC | Q8IWW6 | ARHGAP12 | 3 | Homo sapien | 0.32658099 | -0.3958051 | 1.04896704 | -1.8783153 | 0.96285575 | 0.35075323 | 0.99999785 | -5.9207349 up |
| KI - WT, cell_origin: iPSC | Q9UI10 | EIF2B4 | 3 | Homo sapien | -0.3991139 | -1.2823036 | 0.48407579 | -1.4078872 | -0.9625136 | 0.35092912 | 0.99999785 | -5.9210463 down |
| KI - WT, cell_origin: iPSC | Q8N3F8 | MICALL1 | 5 | Homo sapien | 0.35159995 | -0.4268342 | 1.1300341 | -1.1450114 | 0.96198126 | 0.35117825 | 0.99999785 | -5.9215555 up |
| KI - WT, cell_origin: iPSC | Q9NSC5 | HOMER3 | 11 | Homo sapien | -1.3921452 | -4.482387 | 1.69809657 | -1.4035227 | -0.9600513 | 0.35222149 | 0.99999785 | -5.9232597 down |
| KI - WT, cell_origin: iPSC | P48668 | KRT6C | 30 | Homo sapien | -0.6805044 | -2.1911498 | 0.8301409 | 2.23801352 | -0.9599991 | 0.35224694 | 0.99999785 | -5.9233086 down |
| KI - WT, cell_origin: iPSC | Q96GY0 | ZC2HC1A | 5 | Homo sapien | -0.2849036 | -0.9171326 | 0.34732537 | 0.18843359 | -0.9597613 | 0.35225881 | 0.99999785 | -5.9236356 down |
| KI - WT, cell_origin: iPSC | P20618 | PSMB1 | 5 | Homo sapien | 0.40650772 | -0.4961741 | 1.30918953 | 0.07395757 | 0.95954688 | 0.35243943 | 0.99999785 | -5.9237598 up |
| KI - WT, cell_origin: iPSC | P06730 | EIF4E | 7 | Homo sapien | -0.2413727 | -0.7786647 | 0.29591926 | 1.83681372 | -0.9567919 | 0.35370778 | 0.99999785 | -5.9264115 down |
| KI - WT, cell_origin: iPSC | Q9NZU5 | LMCD1 | 5 | Homo sapien | -0.7005414 | -2.2611584 | 0.86007567 | -2.9649667 | -0.9563567 | 0.3539768 | 0.99999785 | -5.926761 down |
| KI - WT, cell_origin: iPSC | O60828 | PQBP1 | 3 | Homo sapien | -0.7678464 | -2.4845766 | 0.94888371 | -0.5385209 | -0.9531795 | 0.3555793 | 0.99999785 | -5.9296738 down |
| KI - WT, cell_origin: iPSC | Q96PU8 | QKI | 8 | Homo sapien | -0.3319469 | -1.0746817 | 0.41078789 | 1.20056905 | -0.9519597 | 0.35609187 | 0.99999785 | -5.930895 down |
| KI - WT, cell_origin: iPSC | Q8TB03 | CXorf38 | 10 | Homo sapien | 0.42608384 | -0.5278772 | 1.38004488 | 0.46368748 | 0.95184621 | 0.35623338 | 0.99999785 | -5.9309136 up |
| KI - WT, cell_origin: iPSC | Q04206 | RELA | 2 | Homo sapien | -0.6609015 | -2.1427416 | 0.82093861 | -2.4698319 | -0.9504686 | 0.35691008 | 0.99999785 | -5.9321931 down |
| KI - WT, cell_origin: iPSC | Q9BTT6 | LRRC1 | 5 | Homo sapien | 0.69240285 | -0.8617842 | 2.24658985 | -2.8009789 | 0.94928054 | 0.35746957 | 0.99999785 | -5.9333202 up |
| KI - WT, cell_origin: iPSC | P50851 | LRBA | 12 | Homo sapien | 0.4985281 | -0.6217464 | 1.61880256 | -1.8110878 | 0.94801099 | 0.35805922 | 0.99999785 | -5.9345325 up |
| KI - WT, cell_origin: iPSC | P10644 | PRKAR1A | 3 | Homo sapien | 0.45654762 | -0.5701755 | 1.48327078 | -0.338142 | 0.9476219 | 0.3583112 | 0.99999785 | -5.9348316 up |
| KI - WT, cell_origin: iPSC | P62330 | ARF6 | 8 | Homo sapien | 0.59390416 | -0.7442618 | 1.93207007 | 1.98025137 | 0.9458206 | 0.35919976 | 0.99999785 | -5.9364975 up |
| KI - WT, cell_origin: iPSC | Q16401 | PSMD5 | 4 | Homo sapien | 0.54992098 | -0.6903783 | 1.7902203 | -1.6494146 | 0.9448789 | 0.35966489 | 0.99999785 | -5.9373674 up |
| KI - WT, cell_origin: iPSC | O00567 | NOP56 | 21 | Homo sapien | -0.3905479 | -1.2734069 | 0.49231104 | 0.64490959 | -0.9425882 | 0.36077347 | 0.99999785 | -5.9395051 down |
| KI - WT, cell_origin: iPSC | P62068 | USP46 | 2 | Homo sapien | 0.44809981 | -0.5657041 | 1.46190376 | -1.1476993 | 0.94179207 | 0.36116607 | 0.99999785 | -5.9402402 up |
| KI - WT, cell_origin: iPSC | Q66K14 | TBC1D9B | 5 | Homo sapien | 0.65496081 | -0.8300417 | 2.13996335 | -1.7278612 | 0.93991913 | 0.36212156 | 0.99999785 | -5.941936 up |
| KI - WT, cell_origin: iPSC | P04080 | CSTB | 7 | Homo sapien | -0.5394852 | -1.7642245 | 0.6852541 | 3.72721402 | -0.9387247 | 0.36271491 | 0.99999785 | -5.9430331 down |
| KI - WT, cell_origin: iPSC | Q9H6I2 | SOX17 | 4 | Homo sapien | -0.3230115 | -1.0571896 | 0.4111666 | -1.5587721 | -0.9370359 | 0.36345356 | 0.99999785 | -5.9446859 down |
| KI - WT, cell_origin: iPSC | P04049 | RAF1 | 12 | Homo sapien | 0.38263567 | -0.4896755 | 1.25494684 | 0.15706144 | 0.93453696 | 0.36475438 | 0.99999785 | -5.946917 up |
| KI - WT, cell_origin: iPSC | P21741 | MDK | 2 | Homo sapien | -0.9429742 | -3.0965883 | 1.21063989 | -1.6868007 | -0.9331131 | 0.36551153 | 0.99999785 | -5.9481706 down |
| KI - WT, cell_origin: iPSC | Q16643 | DBN1 | 4 | Homo sapien | -0.5517105 | -1.8118561 | 0.70843504 | -1.5921572 | -0.9330242 | 0.36555595 | 0.99999785 | -5.9482518 down |

| | | | | | | | | | | | | |
|---|---|---|---|---|---|---|---|---|---|---|---|---|
| KI - WT, cell_origin: iPSC | Q4G0F5 | VPS26B | 3 | Homo sapien | 0.36918488 | -0.4782089 | 1.21657871 | -1.0501719 | 0.92801348 | 0.36798685 | 0.99999785 | -5.9528971 up |
| KI - WT, cell_origin: iPSC | Q96CX2 | KCTD12 | 9 | Homo sapien | 0.23027361 | -0.2988244 | 0.75937165 | 0.12251209 | 0.92693162 | 0.36850902 | 0.99999785 | -5.9539017 up |
| KI - WT, cell_origin: iPSC | P60900 | PSMA6 | 4 | Homo sapien | 0.32207905 | -0.418061 | 1.0622191 | -1.1413126 | 0.92680477 | 0.36857279 | 0.99999785 | -5.9540168 up |
| KI - WT, cell_origin: iPSC | Q8N2M8 | CLASRP | 4 | Homo sapien | -0.4873153 | -1.6084891 | 0.6338584 | -0.4974858 | -0.9262741 | 0.36893987 | 0.99999785 | -5.9543949 down |
| KI - WT, cell_origin: iPSC | Q9H2U2 | PPA2 | 6 | Homo sapien | -0.7332165 | -2.4218615 | 0.95542851 | -2.634433 | -0.9253291 | 0.36941531 | 0.99999785 | -5.9552517 down |
| KI - WT, cell_origin: iPSC | A0AVF1 | TTC26 | 4 | Homo sapien | -0.3454751 | -1.1412442 | 0.45029396 | -1.3363528 | -0.924633 | 0.36966578 | 0.99999785 | -5.9559858 down |
| KI - WT, cell_origin: iPSC | P67812 | SEC11A | 3 | Homo sapien | 1.23267354 | -1.6114407 | 4.07678774 | -3.4300313 | 0.92364144 | 0.37026546 | 0.99999785 | -5.9567801 up |
| KI - WT, cell_origin: iPSC | Q9NPA8 | ENY2 | 6 | Homo sapien | -0.4567059 | -1.510494 | 0.59708211 | -0.2559033 | -0.9236039 | 0.37028438 | 0.99999785 | -5.956814 down |
| KI - WT, cell_origin: iPSC | Q9UNH7 | SNX6 | 16 | Homo sapien | 0.22881822 | -0.2995418 | 0.75717827 | 1.08573191 | 0.92235965 | 0.37081227 | 0.99999785 | -5.9580424 up |
| KI - WT, cell_origin: iPSC | O14777 | NDC80 | 2 | Homo sapien | 0.59293786 | -0.7827219 | 1.96859759 | 3.48397933 | 0.9185452 | 0.37284076 | 0.99999785 | -5.9613801 up |
| KI - WT, cell_origin: iPSC | Q9H7P6 | MVB12B | 2 | Homo sapien | 0.80659985 | -1.0675875 | 2.68078723 | -1.3714899 | 0.91716506 | 0.37354029 | 0.99999785 | -5.9626219 up |
| KI - WT, cell_origin: iPSC | Q66PJ3 | ARL6IP4 | 8 | Homo sapien | -0.4088664 | -1.3604677 | 0.54273487 | 3.72626377 | -0.9156485 | 0.37431001 | 0.99999785 | -5.9639846 down |
| KI - WT, cell_origin: iPSC | Q5T5P2 | KIAA1217 | 2 | Homo sapien | 0.68842552 | -0.91356 | 2.29041102 | -3.044559 | 0.9155474 | 0.37431617 | 0.99999785 | -5.9641224 up |
| KI - WT, cell_origin: iPSC | Q9NUP9 | LIN7C | 4 | Homo sapien | 0.49895934 | -0.6638884 | 1.66180709 | -0.1904075 | 0.9144178 | 0.37493542 | 0.99999785 | -5.965089 up |
| KI - WT, cell_origin: iPSC | Q9Y4R8 | TELO2 | 8 | Homo sapien | 0.31788417 | -0.4247184 | 1.06048677 | -1.91438 | 0.91170034 | 0.37622034 | 0.99999785 | -5.9676255 up |
| KI - WT, cell_origin: iPSC | Q14103 | HNRNPD | 14 | Homo sapien | -0.2509644 | -0.8372871 | 0.3353583 | 3.89269665 | -0.9116226 | 0.37625997 | 0.99999785 | -5.967695 down |
| KI - WT, cell_origin: iPSC | Q15434 | RBMS2 | 2 | Homo sapien | 0.37148185 | -0.4984172 | 1.24138092 | -0.0553439 | 0.90978863 | 0.37724536 | 0.99999785 | -5.969282 up |
| KI - WT, cell_origin: iPSC | Q5SQI0 | ATAT1 | 3 | Homo sapien | 0.49461108 | -0.6639319 | 1.65315408 | -1.7677212 | 0.9096381 | 0.37733916 | 0.99999785 | -5.9693988 up |
| KI - WT, cell_origin: iPSC | Q6P0Q8 | MAST2 | 2 | Homo sapien | -1.4779236 | -4.9410971 | 1.9852498 | -4.4833127 | -0.9094527 | 0.37746581 | 0.99999785 | -5.9695309 down |
| KI - WT, cell_origin: iPSC | Q15633 | TARBP2 | 3 | Homo sapien | 0.33916402 | -0.4553 | 1.13362809 | -0.5777314 | 0.90935025 | 0.37744081 | 0.99999785 | -5.969703 up |
| KI - WT, cell_origin: iPSC | Q13485 | SMAD4 | 4 | Homo sapien | -0.2549255 | -0.8521228 | 0.34227187 | -1.0964898 | -0.909149 | 0.37752272 | 0.99999785 | -5.9699045 down |
| KI - WT, cell_origin: iPSC | Q9BZL6 | PRKD2 | 6 | Homo sapien | 0.2867628 | -0.3857506 | 0.95927618 | -0.7515842 | 0.90815833 | 0.37802921 | 0.99999785 | -5.9707878 up |
| KI - WT, cell_origin: iPSC | Q9UL54 | TAOK2 | 5 | Homo sapien | -0.7178335 | -2.402577 | 0.96690995 | -2.8873868 | -0.9080135 | 0.37820143 | 0.99999785 | -5.9708144 down |
| KI - WT, cell_origin: iPSC | Q99614 | TTC1 | 6 | Homo sapien | 0.34535598 | -0.4653005 | 1.15601244 | 0.17933407 | 0.90750294 | 0.37839378 | 0.99999785 | -5.9713412 up |
| KI - WT, cell_origin: iPSC | Q6UXK5 | LRRN1 | 6 | Homo sapien | -0.2882985 | -0.9650352 | 0.38843817 | 2.29357539 | -0.9073239 | 0.3784562 | 0.99999785 | -5.9715312 down |
| KI - WT, cell_origin: iPSC | P16070 | CD44 | 4 | Homo sapien | -0.8638587 | -2.8992356 | 1.17151828 | -1.8699679 | -0.9044826 | 0.38001035 | 0.99999785 | -5.9739557 down |
| KI - WT, cell_origin: iPSC | P38432 | COIL | 5 | Homo sapien | -0.3735503 | -1.2548702 | 0.50776965 | -1.2156411 | -0.9028407 | 0.38077673 | 0.99999785 | -5.975493 down |
| KI - WT, cell_origin: iPSC | O75817 | POP7 | 2 | Homo sapien | 0.22869515 | -0.3118208 | 0.76921115 | 0.52712261 | 0.90113132 | 0.38163529 | 0.99999785 | -5.9770291 up |
| KI - WT, cell_origin: iPSC | Q8NEL9 | DDHD1 | 4 | Homo sapien | 0.46132481 | -0.6298367 | 1.55248637 | -1.2946155 | 0.90092422 | 0.38182745 | 0.99999785 | -5.9771227 up |
| KI - WT, cell_origin: iPSC | Q9P289 | STK26 | 2 | Homo sapien | 0.35443555 | -0.4863624 | 1.19523347 | -2.0949123 | 0.89781183 | 0.38334683 | 0.99999785 | -5.9799623 up |
| KI - WT, cell_origin: iPSC | P14618 | PKM | 26 | Homo sapien | -0.3933253 | -1.3275443 | 0.5408937 | 2.07018547 | -0.8972337 | 0.38374235 | 0.99999785 | -5.9803703 down |
| KI - WT, cell_origin: iPSC | Q658Y4 | FAM91A1 | 15 | Homo sapien | 0.24878748 | -0.3422266 | 0.83980157 | 0.60811068 | 0.89654142 | 0.38400322 | 0.99999785 | -5.9810823 up |
| KI - WT, cell_origin: iPSC | P98171 | ARHGAP4 | 2 | Homo sapien | 0.51684819 | -0.7123581 | 1.74605444 | -2.3274341 | 0.89568674 | 0.38447405 | 0.99999785 | -5.9818047 up |
| KI - WT, cell_origin: iPSC | Q8N1S5 | SLC39A11 | 3 | Homo sapien | 0.56970047 | -0.7859199 | 1.92532086 | -0.7994598 | 0.89559333 | 0.38459029 | 0.99999785 | -5.9818155 up |
| KI - WT, cell_origin: iPSC | Q8TBQ9 | TMEM167A | 2 | Homo sapien | -0.8988235 | -3.0390354 | 1.24138836 | 0.54836086 | -0.8949938 | 0.38490051 | 0.99999785 | -5.9823431 down |
| KI - WT, cell_origin: iPSC | P32321 | DCTD | 4 | Homo sapien | 0.47853794 | -0.6610307 | 1.61810657 | -1.4838659 | 0.89484208 | 0.38496733 | 0.99999785 | -5.9824889 up |
| KI - WT, cell_origin: iPSC | Q9NVR0 | KLHL11 | 7 | Homo sapien | -0.2423419 | -0.820487 | 0.33580318 | -1.0050318 | -0.8927531 | 0.38596505 | 0.99999785 | -5.9844137 down |
| KI - WT, cell_origin: iPSC | Q12774 | ARHGEF5 | 14 | Homo sapien | 0.40948233 | -0.5697049 | 1.38866954 | -1.7437608 | 0.89074446 | 0.38702412 | 0.99999785 | -5.986158 up |
| KI - WT, cell_origin: iPSC | Q6FI81 | CIAPIN1 | 6 | Homo sapien | 0.3649042 | -0.5084048 | 1.2382132 | -0.0771101 | 0.8901997 | 0.38734124 | 0.99999785 | -5.9865992 up |
| KI - WT, cell_origin: iPSC | Q9Y4K3 | TRAF6 | 2 | Homo sapien | -0.5621718 | -1.9114807 | 0.78713705 | -1.7487141 | -0.8878918 | 0.38858824 | 0.99999785 | -5.9885689 down |
| KI - WT, cell_origin: iPSC | O14602 | EIF1AY | 4 | Homo sapien | -0.7320757 | -2.4904028 | 1.0262514 | 2.18971604 | -0.8872759 | 0.38890917 | 0.99999785 | -5.9891067 down |
| KI - WT, cell_origin: iPSC | P42684 | ABL2 | 9 | Homo sapien | 0.50406654 | -0.7067691 | 1.71490214 | -0.9708837 | 0.88716634 | 0.38896628 | 0.99999785 | -5.9892023 up |
| KI - WT, cell_origin: iPSC | Q96QK1 | VPS35 | 18 | Homo sapien | 0.36933656 | -0.5185021 | 1.25717524 | 0.6163909 | 0.88639491 | 0.38934547 | 0.99999785 | -5.9898998 up |
| KI - WT, cell_origin: iPSC | O43683 | BUB1 | 9 | Homo sapien | 0.2813963 | -0.395316 | 0.95810858 | -2.2206868 | 0.88563344 | 0.3896703 | 0.99999785 | -5.9906404 up |
| KI - WT, cell_origin: iPSC | Q9H9G7 | AGO3 | 14 | Homo sapien | -0.4638648 | -1.5835526 | 0.65582303 | -0.0899169 | -0.8828701 | 0.39121003 | 0.99999785 | -5.9929441 down |
| KI - WT, cell_origin: iPSC | Q9UKZ1 | CNOT11 | 6 | Homo sapien | 0.39706905 | -0.5617715 | 1.35590958 | -0.1619807 | 0.88251484 | 0.39139598 | 0.99999785 | -5.9932528 up |

| | | | | | | | | | | | | | |
|---|---|---|---|---|---|---|---|---|---|---|---|---|---|
| KI - WT, cell_origin: iPSC | P30153 | PPP2R1A | 28 | Homo sapien | 0.18341602 | -0.2592417 | 0.62607373 | 1.99111126 | 0.88248818 | 0.39131473 | 0.99999785 | -5.9933769 | up |
| KI - WT, cell_origin: iPSC | Q15042 | RAB3GAP1 | 13 | Homo sapien | 0.49840562 | -0.7053129 | 1.70212413 | -0.9749879 | 0.88238955 | 0.39146156 | 0.99999785 | -5.9933617 | up |
| KI - WT, cell_origin: iPSC | Q05519 | SRSF11 | 10 | Homo sapien | -0.7380628 | -2.5212955 | 1.04516986 | 2.12723484 | -0.8820388 | 0.39164523 | 0.99999785 | -5.9936663 | down |
| KI - WT, cell_origin: iPSC | Q6P2E9 | EDC4 | 18 | Homo sapien | -0.3137961 | -1.0718976 | 0.44430541 | -1.3409244 | -0.8815762 | 0.39179239 | 0.99999785 | -5.9941687 | down |
| KI - WT, cell_origin: iPSC | Q07812 | BAX | 2 | Homo sapien | -0.250047 | -0.855318 | 0.35522393 | 0.0074182 | -0.8798559 | 0.39269455 | 0.99999785 | -5.9956604 | down |
| KI - WT, cell_origin: iPSC | O60508 | CDC40 | 3 | Homo sapien | 0.37306691 | -0.5306478 | 1.27678163 | -2.3539951 | 0.87921494 | 0.393031 | 0.99999785 | -5.9962155 | up |
| KI - WT, cell_origin: iPSC | Q9Y383 | LUC7L2 | 23 | Homo sapien | -0.6951763 | -2.3805962 | 0.99024355 | 3.42078596 | -0.8790007 | 0.39323829 | 0.99999785 | -5.9963002 | down |
| KI - WT, cell_origin: iPSC | Q8IZQ1 | WDFY3 | 21 | Homo sapien | -0.7482784 | -2.5627516 | 1.06619477 | -0.9794016 | -0.8788506 | 0.39331715 | 0.99999785 | -5.9964302 | down |
| KI - WT, cell_origin: iPSC | Q9NSI2 | SLX9 | 4 | Homo sapien | 0.2346912 | -0.3341599 | 0.8035423 | -0.4362465 | 0.87869445 | 0.39330437 | 0.99999785 | -5.9966659 | up |
| KI - WT, cell_origin: iPSC | Q14108 | SCARB2 | 3 | Homo sapien | 1.01832364 | -1.451704 | 3.48835124 | -3.0044867 | 0.87859026 | 0.39345386 | 0.99999785 | -5.9966555 | up |
| KI - WT, cell_origin: iPSC | Q969S3 | ZNF622 | 3 | Homo sapien | 0.26383052 | -0.3771978 | 0.90485881 | -0.4965034 | 0.87657187 | 0.39442051 | 0.99999785 | -5.9985006 | up |
| KI - WT, cell_origin: iPSC | Q5T011 | SZT2 | 4 | Homo sapien | 0.26036111 | -0.3727257 | 0.89344794 | -0.7853206 | 0.87589595 | 0.39477638 | 0.99999785 | -5.999084 | up |
| KI - WT, cell_origin: iPSC | P49757 | NUMB | 11 | Homo sapien | -0.3711062 | -1.2748914 | 0.53267908 | 0.42161597 | -0.8749454 | 0.39535203 | 0.99999785 | -5.9998242 | down |
| KI - WT, cell_origin: iPSC | Q5BKY9 | FAM133B | 5 | Homo sapien | -0.6060799 | -2.0826806 | 0.87052078 | 1.92654605 | -0.8747203 | 0.39549017 | 0.99999785 | -5.9999974 | down |
| KI - WT, cell_origin: iPSC | P37108 | SRP14 | 6 | Homo sapien | -0.2755374 | -0.9469862 | 0.39591137 | 3.25820031 | -0.8740546 | 0.3957581 | 0.99999785 | -6.0006593 | down |
| KI - WT, cell_origin: iPSC | Q9NRY5 | FAM114A2 | 19 | Homo sapien | 0.73845645 | -1.0625838 | 2.53949673 | 0.04314461 | 0.87378349 | 0.39598415 | 0.99999785 | -6.0008044 | up |
| KI - WT, cell_origin: iPSC | Q5SSJ5 | HP1BP3 | 3 | Homo sapien | 0.88192291 | -1.2723992 | 3.03624501 | -2.5221792 | 0.87241346 | 0.39670731 | 0.99999785 | -6.0019832 | up |
| KI - WT, cell_origin: iPSC | Q9Y371 | SH3GLB1 | 8 | Homo sapien | 0.42660773 | -0.6170062 | 1.47022169 | 0.53241518 | 0.87114652 | 0.39737684 | 0.99999785 | -6.0030717 | up |
| KI - WT, cell_origin: iPSC | Q2TAA2 | IAH1 | 4 | Homo sapien | 0.36803934 | -0.5322167 | 1.26829533 | 0.14137184 | 0.87108454 | 0.3973845 | 0.99999785 | -6.0031517 | up |
| KI - WT, cell_origin: iPSC | P23246 | SFPQ | 37 | Homo sapien | -0.6282355 | -2.1653815 | 0.90891052 | 4.39791711 | -0.8709831 | 0.39746327 | 0.99999785 | -6.0032121 | down |
| KI - WT, cell_origin: iPSC | Q8IZL8 | PELP1 | 6 | Homo sapien | 0.51814563 | -0.7495426 | 1.78583382 | -2.0813813 | 0.8709064 | 0.39747872 | 0.99999785 | -6.0033047 | up |
| KI - WT, cell_origin: iPSC | Q9BST9 | RTKN | 5 | Homo sapien | 0.4006934 | -0.580099 | 1.38148585 | -0.6056941 | 0.87063766 | 0.39764597 | 0.99999785 | -6.0035086 | up |
| KI - WT, cell_origin: iPSC | Q9BTK6 | PAGR1 | 2 | Homo sapien | -0.9156493 | -3.1556983 | 1.32439965 | -6.2709478 | -0.8705853 | 0.39757983 | 0.99999785 | -6.0036535 | down |
| KI - WT, cell_origin: iPSC | Q13572 | ITPK1 | 2 | Homo sapien | 0.40344282 | -0.5841742 | 1.3910598 | -0.3117313 | 0.8705542 | 0.39769012 | 0.99999785 | -6.0035802 | up |
| KI - WT, cell_origin: iPSC | P20930 | FLG | 3 | Homo sapien | 1.10807486 | -1.6055335 | 3.82168319 | -3.4315774 | 0.87021047 | 0.39787199 | 0.99999785 | -6.0038751 | up |
| KI - WT, cell_origin: iPSC | Q9C0B9 | ZCCHC2 | 4 | Homo sapien | -0.6339422 | -2.187942 | 0.9200575 | -2.7842619 | -0.8692361 | 0.39836526 | 0.99999785 | -6.0047346 | down |
| KI - WT, cell_origin: iPSC | Q9H857 | NT5DC2 | 5 | Homo sapien | 0.63410746 | -0.9207845 | 2.18899944 | -2.0381 | 0.86909052 | 0.39846496 | 0.99999785 | -6.0048353 | up |
| KI - WT, cell_origin: iPSC | Q9UNS2 | COPS3 | 6 | Homo sapien | 0.30650322 | -0.4448345 | 1.05784093 | -0.9539676 | 0.86883946 | 0.39850434 | 0.99999785 | -6.0051502 | up |
| KI - WT, cell_origin: iPSC | Q07866 | KLC1 | 18 | Homo sapien | 0.30111886 | -0.4371475 | 1.03938521 | -0.7207412 | 0.8686895 | 0.39858382 | 0.99999785 | -6.0052787 | up |
| KI - WT, cell_origin: iPSC | O43299 | AP5Z1 | 6 | Homo sapien | 0.40010608 | -0.581289 | 1.38150111 | -1.9580975 | 0.86830234 | 0.39878906 | 0.99999785 | -6.0056102 | up |
| KI - WT, cell_origin: iPSC | Q9UPT9 | USP22 | 5 | Homo sapien | -0.3546413 | -1.2256576 | 0.51637506 | -1.0183974 | -0.8673008 | 0.39934418 | 0.99999785 | -6.0064416 | down |
| KI - WT, cell_origin: iPSC | P35998 | PSMC2 | 10 | Homo sapien | 0.35490522 | -0.5167941 | 1.22660455 | -0.6890771 | 0.8672885 | 0.39935468 | 0.99999785 | -6.0064479 | up |
| KI - WT, cell_origin: iPSC | Q99536 | VAT1 | 14 | Homo sapien | 0.7277577 | -1.0618206 | 2.51733604 | 1.51722325 | 0.86663946 | 0.39976473 | 0.99999785 | -6.0069328 | up |
| KI - WT, cell_origin: iPSC | P31151 | S100A7 | 3 | Homo sapien | 0.35511622 | -0.5203672 | 1.23059962 | -0.6332261 | 0.86405322 | 0.40107391 | 0.99999785 | -6.00921 | up |
| KI - WT, cell_origin: iPSC | Q9BUP3 | HTATIP2 | 6 | Homo sapien | 0.33364179 | -0.4894161 | 1.15669968 | -0.370567 | 0.86350256 | 0.40136548 | 0.99999785 | -6.0096808 | up |
| KI - WT, cell_origin: iPSC | O43414 | ERI3 | 2 | Homo sapien | -0.2724597 | -0.9450864 | 0.4001671 | -0.9940847 | -0.8627158 | 0.40175836 | 0.99999785 | -6.0103787 | down |
| KI - WT, cell_origin: iPSC | Q8WUH2 | TGFBRAP1 | 4 | Homo sapien | 0.3872086 | -0.569494 | 1.34391117 | -2.0141217 | 0.86200094 | 0.40213937 | 0.99999785 | -6.0109869 | up |
| KI - WT, cell_origin: iPSC | Q9UHR5 | SAP30BP | 3 | Homo sapien | -0.7072862 | -2.4565212 | 1.04194876 | -1.2673963 | -0.8616867 | 0.4023997 | 0.99999785 | -6.0111548 | down |
| KI - WT, cell_origin: iPSC | P23284 | PPIB | 15 | Homo sapien | -1.0347256 | -3.5954152 | 1.52596396 | 3.93499001 | -0.8611338 | 0.40269459 | 0.99999785 | -6.0116248 | down |
| KI - WT, cell_origin: iPSC | Q6UB99 | ANKRD11 | 3 | Homo sapien | 0.60425936 | -0.8918992 | 2.10041794 | -2.6152966 | 0.86058499 | 0.40296824 | 0.99999785 | -6.0121115 | up |
| KI - WT, cell_origin: iPSC | Q9Y4F9 | RIPOR2 | 16 | Homo sapien | 0.66708015 | -0.9858628 | 2.32002312 | -1.1026638 | 0.86004766 | 0.40327425 | 0.99999785 | -6.0125472 | up |
| KI - WT, cell_origin: iPSC | Q8N5A5 | ZGPAT | 3 | Homo sapien | -0.4567378 | -1.5884738 | 0.67499826 | -1.4648521 | -0.8599878 | 0.40329498 | 0.99999785 | -6.01261 | down |
| KI - WT, cell_origin: iPSC | Q5T5Y3 | CAMSAP1 | 2 | Homo sapien | -0.9129354 | -3.1754851 | 1.34961426 | -2.4160396 | -0.8598928 | 0.40335695 | 0.99999785 | -6.0126786 | down |
| KI - WT, cell_origin: iPSC | Q8ND25 | ZNRF1 | 4 | Homo sapien | 1.12928896 | -1.6701619 | 3.92873979 | -3.0094157 | 0.85967562 | 0.40347293 | 0.99999785 | -6.0128629 | up |
| KI - WT, cell_origin: iPSC | P11308 | ERG | 13 | Homo sapien | 0.33002231 | -0.4886805 | 1.14872513 | -0.6679389 | 0.8586651 | 0.4039441 | 0.99999785 | -6.0137936 | up |
| KI - WT, cell_origin: iPSC | P26599 | PTBP1 | 12 | Homo sapien | -0.2932254 | -1.0211495 | 0.43469866 | 3.27743453 | -0.8580912 | 0.40425495 | 0.99999785 | -6.0142756 | down |

| | | | | | | | | | | | | | |
|---|---|---|---|---|---|---|---|---|---|---|---|---|---|
| KI - WT, cell_origin: iPSC | P33992 | MCM5 | 43 | Homo sapien | -0.3666545 | -1.2787176 | 0.54540855 | 3.33975216 | -0.8567114 | 0.40505822 | 0.99999785 | -6.015374 | down |
| KI - WT, cell_origin: iPSC | O14818 | PSMA7 | 6 | Homo sapien | 0.3226004 | -0.4798118 | 1.12501262 | 0.21114046 | 0.85641699 | 0.40515118 | 0.99999785 | -6.0156924 | up |
| KI - WT, cell_origin: iPSC | Q9H5X1 | CIAO2A | 3 | Homo sapien | 0.4704335 | -0.704 | 1.64486705 | 0.60101562 | 0.8536349 | 0.40670789 | 0.99999785 | -6.0179719 | up |
| KI - WT, cell_origin: iPSC | Q14554 | PDIA5 | 19 | Homo sapien | 2.17239317 | -3.2554969 | 7.6002832 | -1.6603259 | 0.85292286 | 0.40709033 | 0.99999785 | -6.018572 | up |
| KI - WT, cell_origin: iPSC | O75223 | GGCT | 3 | Homo sapien | 0.26778339 | -0.4009404 | 0.93650721 | 0.7038734 | 0.85285759 | 0.40703366 | 0.99999785 | -6.0187255 | up |
| KI - WT, cell_origin: iPSC | Q7Z7H5 | TMED4 | 9 | Homo sapien | 0.80489119 | -1.2089954 | 2.81877781 | 1.20539733 | 0.85173507 | 0.40772883 | 0.99999785 | -6.019572 | up |
| KI - WT, cell_origin: iPSC | Q7Z4L5 | TTC21B | 4 | Homo sapien | -0.3423924 | -1.1997402 | 0.51495536 | -0.3495664 | -0.8507168 | 0.40821245 | 0.99999785 | -6.0204973 | down |
| KI - WT, cell_origin: iPSC | P61326;Q96/ | MAGOH;MA( | 7 | Homo sapien | -0.5619249 | -1.9727381 | 0.84888817 | 0.15171298 | -0.8488114 | 0.40930324 | 0.99999785 | -6.0220281 | down |
| KI - WT, cell_origin: iPSC | Q9BTC0 | DIDO1 | 6 | Homo sapien | -0.4411552 | -1.5487927 | 0.66648235 | -1.7668726 | -0.8484805 | 0.4094281 | 0.99999785 | -6.0223631 | down |
| KI - WT, cell_origin: iPSC | O95644;Q12! | NFATC1;NFA | 2 | Homo sapien | 1.74927657 | -2.6452581 | 6.14381128 | -3.937416 | 0.84829703 | 0.40958063 | 0.99999785 | -6.0224593 | up |
| KI - WT, cell_origin: iPSC | Q9NVJ2 | ARL8B | 3 | Homo sapien | -0.4548094 | -1.599232 | 0.68961317 | -0.8652129 | -0.8469259 | 0.41032071 | 0.99999785 | -6.0236079 | down |
| KI - WT, cell_origin: iPSC | P0DMV8;P0D | HSPA1A;HSP/ | 27 | Homo sapien | 0.17002054 | -0.257973 | 0.59801408 | 5.48588307 | 0.84606526 | 0.41069471 | 0.99999785 | -6.0244259 | up |
| KI - WT, cell_origin: iPSC | Q15397 | PUM3 | 8 | Homo sapien | 0.36542588 | -0.5549986 | 1.2858504 | -1.7972624 | 0.84557242 | 0.41096119 | 0.99999785 | -6.0248379 | up |
| KI - WT, cell_origin: iPSC | Q00688 | FKBP3 | 7 | Homo sapien | -0.8130058 | -2.8624831 | 1.2364716 | -0.3143494 | -0.8453817 | 0.41115518 | 0.99999785 | -6.0248994 | down |
| KI - WT, cell_origin: iPSC | Q00765 | REEP5 | 2 | Homo sapien | 0.60418732 | -0.9192626 | 2.12763728 | -1.454077 | 0.84517332 | 0.4112679 | 0.99999785 | -6.0250736 | up |
| KI - WT, cell_origin: iPSC | Q9Y221 | NIP7 | 5 | Homo sapien | -0.2588277 | -0.9115884 | 0.39393291 | -1.3596826 | -0.8444939 | 0.41154471 | 0.99999785 | -6.0257388 | down |
| KI - WT, cell_origin: iPSC | Q8WUW1 | BRK1 | 3 | Homo sapien | 0.350131 | -0.5340621 | 1.23432414 | 0.77021047 | 0.84376681 | 0.41200725 | 0.99999785 | -6.0262714 | up |
| KI - WT, cell_origin: iPSC | Q13618 | CUL3 | 9 | Homo sapien | 0.22880508 | -0.3488003 | 0.80641048 | -1.0838475 | 0.84367292 | 0.4119893 | 0.99999785 | -6.0264238 | up |
| KI - WT, cell_origin: iPSC | Q86YL5 | TDRP | 3 | Homo sapien | 0.43585377 | -0.6657204 | 1.53742798 | -3.3909533 | 0.84268783 | 0.41252314 | 0.99999785 | -6.027245 | up |
| KI - WT, cell_origin: iPSC | P20936 | RASA1 | 9 | Homo sapien | 0.32286176 | -0.4948871 | 1.14061063 | -0.2601412 | 0.84102755 | 0.41344933 | 0.99999785 | -6.0285997 | up |
| KI - WT, cell_origin: iPSC | P57740 | NUP107 | 9 | Homo sapien | 0.25876351 | -0.397003 | 0.91453005 | -0.7586097 | 0.84041434 | 0.41375692 | 0.99999785 | -6.0291369 | up |
| KI - WT, cell_origin: iPSC | Q9NUQ3 | TXLNG | 9 | Homo sapien | -0.3907238 | -1.3813562 | 0.59990858 | -1.9341743 | -0.8400459 | 0.41395913 | 0.99999785 | -6.0294408 | down |
| KI - WT, cell_origin: iPSC | O43865 | AHCYL1 | 14 | Homo sapien | 0.37266171 | -0.5729864 | 1.31830981 | 0.23126642 | 0.83982266 | 0.41416861 | 0.99999785 | -6.0295311 | up |
| KI - WT, cell_origin: iPSC | P60763 | RAC3 | 3 | Homo sapien | 0.33000068 | -0.5076319 | 1.16763326 | 1.79166465 | 0.83928495 | 0.41440795 | 0.99999785 | -6.0300348 | up |
| KI - WT, cell_origin: iPSC | Q9UKK9 | NUDT5 | 5 | Homo sapien | -0.4457587 | -1.5792052 | 0.68768771 | -1.748942 | -0.837879 | 0.41518454 | 0.99999785 | -6.0311884 | down |
| KI - WT, cell_origin: iPSC | P08243 | ASNS | 2 | Homo sapien | -0.2651898 | -0.9399099 | 0.40953032 | -1.988204 | -0.8370913 | 0.41556461 | 0.99999785 | -6.0318938 | down |
| KI - WT, cell_origin: iPSC | Q9NZN5 | ARHGEF12 | 62 | Homo sapien | 0.40410487 | -0.6268677 | 1.43507741 | 2.38521983 | 0.83531323 | 0.41662362 | 0.99999785 | -6.033268 | up |
| KI - WT, cell_origin: iPSC | P48729 | CSNK1A1 | 9 | Homo sapien | 0.31669312 | -0.4912404 | 1.12462659 | 2.2177441 | 0.83498933 | 0.41673737 | 0.99999785 | -6.0336037 | up |
| KI - WT, cell_origin: iPSC | Q15075 | EEA1 | 59 | Homo sapien | -0.6669163 | -2.3696376 | 1.03580506 | 1.45317219 | -0.8346994 | 0.41695853 | 0.99999785 | -6.0337752 | down |
| KI - WT, cell_origin: iPSC | Q86UX7 | FERMT3 | 11 | Homo sapien | 0.58924561 | -0.9160212 | 2.09451242 | -1.5571714 | 0.83422893 | 0.41721533 | 0.99999785 | -6.0341638 | up |
| KI - WT, cell_origin: iPSC | Q9NW64 | RBM22 | 10 | Homo sapien | -0.2880474 | -1.0238433 | 0.44774847 | 0.99507467 | -0.8338546 | 0.41734526 | 0.99999785 | -6.0345534 | down |
| KI - WT, cell_origin: iPSC | Q9H223 | EHD4 | 11 | Homo sapien | 0.31871183 | -0.4964179 | 1.13384153 | -0.0255134 | 0.83289331 | 0.41788218 | 0.99999785 | -6.0353336 | up |
| KI - WT, cell_origin: iPSC | Q8NDI1 | EHBP1 | 6 | Homo sapien | 0.5549933 | -0.8651001 | 1.97508666 | -0.4417198 | 0.8328623 | 0.4179619 | 0.99999785 | -6.0352913 | up |
| KI - WT, cell_origin: iPSC | Q86Y07 | VRK2 | 5 | Homo sapien | 0.56498411 | -0.8808072 | 2.01077545 | -1.527719 | 0.83278517 | 0.41800406 | 0.99999785 | -6.0353549 | up |
| KI - WT, cell_origin: iPSC | Q13136 | PPFIA1 | 4 | Homo sapien | -0.3699899 | -1.317355 | 0.57737521 | -1.5105862 | -0.8318943 | 0.41842098 | 0.99999785 | -6.0361649 | down |
| KI - WT, cell_origin: iPSC | Q9BT25 | HAUS8 | 2 | Homo sapien | -0.3340851 | -1.1914163 | 0.52324612 | -1.1629852 | -0.8300269 | 0.41943949 | 0.99999785 | -6.0377056 | down |
| KI - WT, cell_origin: iPSC | Q96RL7 | VPS13A | 49 | Homo sapien | 0.85002397 | -1.3369709 | 3.03701887 | -0.5830682 | 0.8282964 | 0.42046246 | 0.99999785 | -6.0390462 | up |
| KI - WT, cell_origin: iPSC | Q8NHP8 | PLBD2 | 3 | Homo sapien | 0.40254236 | -0.6333037 | 1.43838845 | -2.1277084 | 0.82767078 | 0.4207176 | 0.99999785 | -6.0396548 | up |
| KI - WT, cell_origin: iPSC | P48426 | PIP4K2A | 6 | Homo sapien | -0.7338685 | -2.624641 | 1.1569039 | -1.8734319 | -0.8271445 | 0.42109484 | 0.99999785 | -6.0399906 | down |
| KI - WT, cell_origin: iPSC | O43251;Q9N | RBFOX2;RBF( | 2 | Homo sapien | -0.2679028 | -0.9577764 | 0.4219709 | 0.1433217 | -0.8270796 | 0.42104177 | 0.99999785 | -6.0401398 | down |
| KI - WT, cell_origin: iPSC | Q709C8 | VPS13C | 168 | Homo sapien | 0.61505927 | -0.9703987 | 2.20051723 | 1.88357189 | 0.82673173 | 0.42132158 | 0.99999785 | -6.0403286 | up |
| KI - WT, cell_origin: iPSC | Q8N0X7 | SPART | 12 | Homo sapien | 0.62401696 | -0.9848407 | 2.2328746 | -0.5722709 | 0.82657287 | 0.42140888 | 0.99999785 | -6.0404587 | up |
| KI - WT, cell_origin: iPSC | Q13610 | PWP1 | 10 | Homo sapien | 0.25721638 | -0.4059404 | 0.9203732 | -0.1248562 | 0.8260799 | 0.42159124 | 0.99999785 | -6.0409583 | up |
| KI - WT, cell_origin: iPSC | P62263 | RPS14 | 6 | Homo sapien | -0.1587302 | -0.5680832 | 0.25062286 | 5.05499496 | -0.82585 | 0.42171768 | 0.99999785 | -6.0411464 | down |
| KI - WT, cell_origin: iPSC | Q96MG8 | PCMTD1 | 5 | Homo sapien | -0.5700147 | -2.041855 | 0.90182561 | 1.48108013 | -0.8253302 | 0.42209211 | 0.99999785 | -6.0414755 | down |
| KI - WT, cell_origin: iPSC | P0CAP2 | POLR2M | 2 | Homo sapien | -0.2418875 | -0.8665727 | 0.38279764 | -1.079014 | -0.8246924 | 0.42235465 | 0.99999785 | -6.0420927 | down |

| | | | | | | | | | | | | | |
|---|---|---|---|---|---|---|---|---|---|---|---|---|---|
| KI - WT, cell_origin: iPSC | Q9NRV9 | HEBP1 | 3 | Homo sapien | 0.38296498 | -0.6068699 | 1.3727999 | -0.8917438 | 0.82445468 | 0.42256317 | 0.99999785 | -6.0422026 | up |
| KI - WT, cell_origin: iPSC | P56962 | STX17 | 5 | Homo sapien | 0.56108186 | -0.8896852 | 2.01184894 | 0.11228169 | 0.82419676 | 0.42271589 | 0.99999785 | -6.0424017 | up |
| KI - WT, cell_origin: iPSC | Q86X10 | RALGAPB | 8 | Homo sapien | 0.31732105 | -0.5063584 | 1.1410005 | -0.688919 | 0.82063011 | 0.42461724 | 0.99999785 | -6.0453796 | up |
| KI - WT, cell_origin: iPSC | O15068 | MCF2L | 12 | Homo sapien | 0.46484059 | -0.7426751 | 1.67235628 | -1.2744956 | 0.82037729 | 0.42482234 | 0.99999785 | -6.0455141 | up |
| KI - WT, cell_origin: iPSC | P59780 | AP3S2 | 2 | Homo sapien | 0.48657227 | -0.7777805 | 1.750925 | -1.5976689 | 0.82012768 | 0.42496023 | 0.99999785 | -6.0457171 | up |
| KI - WT, cell_origin: iPSC | Q13445 | TMED1 | 5 | Homo sapien | -0.733713 | -2.6502478 | 1.18282187 | -0.4525283 | -0.8158529 | 0.42732629 | 0.99999785 | -6.049184 | down |
| KI - WT, cell_origin: iPSC | O60934 | NBN | 3 | Homo sapien | 1.06139784 | -1.7136287 | 3.83642438 | -1.8653976 | 0.81510476 | 0.42774123 | 0.99999785 | -6.0497891 | up |
| KI - WT, cell_origin: iPSC | Q8TDJ6 | DMXL2 | 25 | Homo sapien | -0.6447903 | -2.3351607 | 1.0455801 | -1.0901729 | -0.8129035 | 0.42896366 | 0.99999785 | -6.0515664 | down |
| KI - WT, cell_origin: iPSC | Q06323 | PSME1 | 15 | Homo sapien | -0.219867 | -0.796124 | 0.35639002 | 1.85134572 | -0.8126125 | 0.42903843 | 0.99999785 | -6.0518956 | down |
| KI - WT, cell_origin: iPSC | O95391 | SLU7 | 3 | Homo sapien | -0.4433172 | -1.6062528 | 0.71961851 | -1.2283342 | -0.8123837 | 0.42925262 | 0.99999785 | -6.0519854 | down |
| KI - WT, cell_origin: iPSC | O95817 | BAG3 | 9 | Homo sapien | 0.34837151 | -0.5654651 | 1.26220807 | -0.6400743 | 0.81218797 | 0.42932182 | 0.99999785 | -6.0521863 | up |
| KI - WT, cell_origin: iPSC | P60866 | RPS20 | 5 | Homo sapien | -0.1494385 | -0.5414752 | 0.24259824 | 6.09592089 | -0.8118492 | 0.42946302 | 0.99999785 | -6.0525106 | down |
| KI - WT, cell_origin: iPSC | O43813 | LANCL1 | 21 | Homo sapien | 0.24181742 | -0.3929269 | 0.87656176 | 5.09935047 | 0.81153388 | 0.42966439 | 0.99999785 | -6.0527364 | up |
| KI - WT, cell_origin: iPSC | Q8NCB2 | CAMKV | 4 | Homo sapien | 0.4863912 | -0.7912311 | 1.76401355 | -3.2127645 | 0.81081708 | 0.43003761 | 0.99999785 | -6.0533414 | up |
| KI - WT, cell_origin: iPSC | P16615 | ATP2A2 | 8 | Homo sapien | 0.43092963 | -0.7026609 | 1.56452011 | -1.0679328 | 0.81012593 | 0.43050932 | 0.99999785 | -6.0538028 | up |
| KI - WT, cell_origin: iPSC | Q8IUF1;Q9BI | CBWD2;CBW | 2 | Homo sapien | 0.5184951 | -0.8489264 | 1.88591656 | -3.053345 | 0.80763132 | 0.43182443 | 0.99999785 | -6.0558882 | up |
| KI - WT, cell_origin: iPSC | Q8WXX5 | DNAJC9 | 9 | Homo sapien | 0.34994001 | -0.5735658 | 1.27344579 | 0.64321863 | 0.80752569 | 0.43195952 | 0.99999785 | -6.05589 | up |
| KI - WT, cell_origin: iPSC | Q9NQ88 | TIGAR | 5 | Homo sapien | -0.2570737 | -0.9381448 | 0.42399749 | -0.3612754 | -0.8039051 | 0.43389804 | 0.99999785 | -6.0588797 | down |
| KI - WT, cell_origin: iPSC | Q9Y2I8 | WDR37 | 3 | Homo sapien | 0.57200775 | -0.9475729 | 2.09158839 | -3.1474009 | 0.80185541 | 0.43507258 | 0.99999785 | -6.0604859 | up |
| KI - WT, cell_origin: iPSC | Q15233 | NONO | 28 | Homo sapien | -0.5846736 | -2.1409904 | 0.9716431 | 3.38910249 | -0.8006042 | 0.43583493 | 0.99999785 | -6.061416 | down |
| KI - WT, cell_origin: iPSC | Q9P2Y4 | ZNF219 | 5 | Homo sapien | -0.3624812 | -1.3272877 | 0.60232525 | -2.3542723 | -0.800175 | 0.43599048 | 0.99999785 | -6.0618505 | down |
| KI - WT, cell_origin: iPSC | P11233 | RALA | 8 | Homo sapien | 0.59091313 | -0.9851609 | 2.16698719 | 2.79892628 | 0.79900479 | 0.43673362 | 0.99999785 | -6.0626868 | up |
| KI - WT, cell_origin: iPSC | O60869 | EDF1 | 10 | Homo sapien | -0.8084239 | -2.9666946 | 1.34984676 | 0.90986703 | -0.7982439 | 0.43716154 | 0.99999785 | -6.0632905 | down |
| KI - WT, cell_origin: iPSC | A6NI72;A8M | NCF1B;NCF1( | 3 | Homo sapien | -0.7414453 | -2.7220801 | 1.23918941 | 0.02593783 | -0.7977689 | 0.43742883 | 0.99999785 | -6.0636672 | down |
| KI - WT, cell_origin: iPSC | O60524 | NEMF | 8 | Homo sapien | 0.22479456 | -0.3756702 | 0.82525937 | -0.5785773 | 0.79732966 | 0.43759093 | 0.99999785 | -6.0641082 | up |
| KI - WT, cell_origin: iPSC | Q9Y6Q2 | STON1 | 6 | Homo sapien | -0.3658009 | -1.3459293 | 0.61432746 | -1.662406 | -0.7949824 | 0.43893211 | 0.99999785 | -6.0659453 | down |
| KI - WT, cell_origin: iPSC | Q14155 | ARHGEF7 | 26 | Homo sapien | 0.57815904 | -0.9732639 | 2.12958203 | 1.0284916 | 0.79418096 | 0.43945112 | 0.99999785 | -6.0665053 | up |
| KI - WT, cell_origin: iPSC | Q9HBG6 | IFT122 | 2 | Homo sapien | -0.4583127 | -1.6904201 | 0.77379484 | -2.0750572 | -0.7924935 | 0.44036567 | 0.99999785 | -6.0678783 | down |
| KI - WT, cell_origin: iPSC | Q3L8U1 | CHD9 | 12 | Homo sapien | 0.62512281 | -1.0563656 | 2.30661121 | -1.2834078 | 0.79227115 | 0.44052996 | 0.99999785 | -6.0680113 | up |
| KI - WT, cell_origin: iPSC | Q9ULC3 | RAB23 | 9 | Homo sapien | 0.32518448 | -0.5498246 | 1.20019358 | 2.7832811 | 0.79193174 | 0.44071161 | 0.99999785 | -6.0682898 | up |
| KI - WT, cell_origin: iPSC | P36915 | GNL1 | 14 | Homo sapien | 0.77217739 | -1.3064886 | 2.8508434 | -1.9290326 | 0.79165277 | 0.44087963 | 0.99999785 | -6.0684982 | up |
| KI - WT, cell_origin: iPSC | Q5BKZ1 | ZNF326 | 7 | Homo sapien | -0.2563684 | -0.9462022 | 0.43346542 | -0.3794299 | -0.791516 | 0.44087254 | 0.99999785 | -6.0686981 | down |
| KI - WT, cell_origin: iPSC | Q6IBS0 | TWF2 | 6 | Homo sapien | -0.3878782 | -1.4336187 | 0.65786237 | -1.0904768 | -0.7904489 | 0.44156092 | 0.99999785 | -6.0694451 | down |
| KI - WT, cell_origin: iPSC | Q15464 | SHB | 2 | Homo sapien | -1.2381882 | -4.5784143 | 2.10203802 | -4.5195782 | -0.7899746 | 0.44182947 | 0.99999785 | -6.0698177 | down |
| KI - WT, cell_origin: iPSC | P62424 | RPL7A | 12 | Homo sapien | -0.1838556 | -0.6795852 | 0.31187389 | 5.12053 | -0.7898995 | 0.44178776 | 0.99999785 | -6.0699688 | down |
| KI - WT, cell_origin: iPSC | P21333 | FLNA | 89 | Homo sapien | -0.2937088 | -1.0862299 | 0.49881221 | 2.19473628 | -0.789449 | 0.44206808 | 0.99999785 | -6.0702951 | down |
| KI - WT, cell_origin: iPSC | P11441 | UBL4A | 7 | Homo sapien | 0.44257529 | -0.7523018 | 1.63745238 | -0.413216 | 0.78934396 | 0.44218675 | 0.99999785 | -6.070313 | up |
| KI - WT, cell_origin: iPSC | Q92616 | GCN1 | 45 | Homo sapien | 0.25134543 | -0.4268431 | 0.92953393 | -0.4146873 | 0.78933298 | 0.44210878 | 0.99999785 | -6.0704136 | up |
| KI - WT, cell_origin: iPSC | P08133 | ANXA6 | 8 | Homo sapien | 0.52284737 | -0.8919931 | 1.93768785 | -1.593576 | 0.78753491 | 0.44321265 | 0.99999785 | -6.0717316 | up |
| KI - WT, cell_origin: iPSC | Q15366 | PCBP2 | 13 | Homo sapien | -0.2165762 | -0.8025737 | 0.36942139 | 4.33266672 | -0.7871475 | 0.4433491 | 0.99999785 | -6.0721262 | down |
| KI - WT, cell_origin: iPSC | O15116 | LSM1 | 2 | Homo sapien | 0.22327445 | -0.3812179 | 0.82776684 | -0.0660251 | 0.78666144 | 0.44362464 | 0.99999785 | -6.0725071 | up |
| KI - WT, cell_origin: iPSC | Q86VY4 | TSPYL5 | 4 | Homo sapien | 0.26428656 | -0.4517646 | 0.98033768 | -1.4089544 | 0.78608721 | 0.44395089 | 0.99999785 | -6.0729563 | up |
| KI - WT, cell_origin: iPSC | Q96A49 | SYAP1 | 14 | Homo sapien | 0.54662463 | -0.9360824 | 2.02933168 | 1.40826097 | 0.78566283 | 0.44427585 | 0.99999785 | -6.0731964 | up |
| KI - WT, cell_origin: iPSC | P59768 | GNG2 | 3 | Homo sapien | 0.30417896 | -0.5208357 | 1.12919364 | 2.89980127 | 0.78555063 | 0.44430913 | 0.99999785 | -6.0733175 | up |
| KI - WT, cell_origin: iPSC | O75391 | SPAG7 | 5 | Homo sapien | -0.2495831 | -0.9263117 | 0.42714555 | -0.262609 | -0.7854893 | 0.44429074 | 0.99999785 | -6.0734236 | down |
| KI - WT, cell_origin: iPSC | Q08AM6 | VAC14 | 11 | Homo sapien | 0.19652151 | -0.3364891 | 0.72953208 | -0.0339542 | 0.78526083 | 0.44442066 | 0.99999785 | -6.0736021 | up |

| | | | | | | | | | | | | | |
|---|---|---|---|---|---|---|---|---|---|---|---|---|---|
| KI - WT, cell_origin: iPSC | Q13643 | FHL3 | 5 | Homo sapien | -0.2854114 | -1.0596411 | 0.48881837 | 1.09693738 | -0.7852505 | 0.44444793 | 0.99999785 | -6.0735868 | down |
| KI - WT, cell_origin: iPSC | O60884 | DNAJA2 | 11 | Homo sapien | 0.20830693 | -0.3568024 | 0.7734163 | 1.4108819 | 0.78507453 | 0.44452661 | 0.99999785 | -6.0737476 | up |
| KI - WT, cell_origin: iPSC | P09211 | GSTP1 | 6 | Homo sapien | -0.3912217 | -1.4552848 | 0.67284146 | 0.62296024 | -0.7835341 | 0.44548679 | 0.99999785 | -6.0748582 | down |
| KI - WT, cell_origin: iPSC | Q9UL46 | PSME2 | 12 | Homo sapien | -0.2168402 | -0.806505 | 0.37282467 | 0.96353788 | -0.7832027 | 0.44559201 | 0.99999785 | -6.0752078 | down |
| KI - WT, cell_origin: iPSC | Q5JU85 | IQSEC2 | 5 | Homo sapien | -0.4346491 | -1.6167903 | 0.74749208 | -2.3674456 | -0.7831864 | 0.44561912 | 0.99999785 | -6.0752011 | down |
| KI - WT, cell_origin: iPSC | Q7L014 | DDX46 | 47 | Homo sapien | -0.6036436 | -2.2464041 | 1.03911688 | 1.9829136 | -0.7830847 | 0.4457427 | 0.99999785 | -6.0752085 | down |
| KI - WT, cell_origin: iPSC | O60925 | PFDN1 | 4 | Homo sapien | 0.35094488 | -0.6051935 | 1.30708326 | 0.79493024 | 0.7822049 | 0.44624393 | 0.99999785 | -6.0758937 | up |
| KI - WT, cell_origin: iPSC | Q96SY0 | INTS14 | 6 | Homo sapien | 0.57331027 | -0.9902168 | 2.13683737 | -2.5679971 | 0.78142388 | 0.44668921 | 0.99999785 | -6.0765015 | up |
| KI - WT, cell_origin: iPSC | Q03001 | DST | 8 | Homo sapien | -0.2910512 | -1.0844738 | 0.50237154 | -1.3448576 | -0.7812758 | 0.44669042 | 0.99999785 | -6.0767077 | down |
| KI - WT, cell_origin: iPSC | P05204 | HMGN2 | 3 | Homo sapien | -0.8834351 | -3.2936425 | 1.52677226 | 4.66451755 | -0.7811288 | 0.44685754 | 0.99999785 | -6.076731 | down |
| KI - WT, cell_origin: iPSC | Q9HBD1 | RC3H2 | 4 | Homo sapien | -0.452301 | -1.686098 | 0.78149594 | -2.0075382 | -0.7810896 | 0.44685284 | 0.99999785 | -6.076791 | down |
| KI - WT, cell_origin: iPSC | Q9H444 | CHMP4B | 3 | Homo sapien | 0.52340315 | -0.905552 | 1.9523583 | -1.4527498 | 0.78058481 | 0.44716789 | 0.99999785 | -6.0771537 | up |
| KI - WT, cell_origin: iPSC | P41223 | BUD31 | 10 | Homo sapien | -0.9305723 | -3.4736559 | 1.61251142 | -1.0740561 | -0.7798154 | 0.4476071 | 0.99999785 | -6.0777512 | down |
| KI - WT, cell_origin: iPSC | Q659C4 | LARP1B | 3 | Homo sapien | -0.290642 | -1.0845341 | 0.5032501 | -1.9955307 | -0.7797162 | 0.44758072 | 0.99999785 | -6.0779191 | down |
| KI - WT, cell_origin: iPSC | P48506 | GCLC | 14 | Homo sapien | 0.28718528 | -0.4976348 | 1.07200535 | -0.9783041 | 0.77935142 | 0.44778961 | 0.99999785 | -6.0782016 | up |
| KI - WT, cell_origin: iPSC | Q14457 | BECN1 | 8 | Homo sapien | 0.19860829 | -0.3442371 | 0.7414537 | 0.01219071 | 0.7792214 | 0.4478634 | 0.99999785 | -6.078303 | up |
| KI - WT, cell_origin: iPSC | O00151 | PDLIM1 | 5 | Homo sapien | -0.4479643 | -1.6730616 | 0.77713297 | -2.9419131 | -0.7787759 | 0.448118 | 0.99999785 | -6.0786484 | down |
| KI - WT, cell_origin: iPSC | Q9Y6A4 | CFAP20 | 5 | Homo sapien | -0.3714733 | -1.3884124 | 0.64546592 | 0.27226658 | -0.7784576 | 0.4483829 | 0.99999785 | -6.0788045 | down |
| KI - WT, cell_origin: iPSC | Q99829 | CPNE1 | 3 | Homo sapien | -0.4081848 | -1.5250537 | 0.70868408 | -2.4733704 | -0.7783847 | 0.44834164 | 0.99999785 | -6.0789516 | down |
| KI - WT, cell_origin: iPSC | P39748 | FEN1 | 10 | Homo sapien | -0.656304 | -2.4538068 | 1.14119885 | -0.6905976 | -0.7781043 | 0.44858487 | 0.99999785 | -6.0790782 | down |
| KI - WT, cell_origin: iPSC | P51606 | RENBP | 6 | Homo sapien | -0.3597158 | -1.3451867 | 0.62575521 | -0.8181934 | -0.7778326 | 0.44873021 | 0.99999785 | -6.0792996 | down |
| KI - WT, cell_origin: iPSC | Q9BV68 | RNF126 | 5 | Homo sapien | 0.30013997 | -0.5222793 | 1.12255927 | -0.1896607 | 0.777407 | 0.44892559 | 0.99999785 | -6.0796818 | up |
| KI - WT, cell_origin: iPSC | O95602 | POLR1A | 9 | Homo sapien | 0.31324515 | -0.5455772 | 1.17206754 | -1.3020406 | 0.77685844 | 0.4492216 | 0.99999785 | -6.0801259 | up |
| KI - WT, cell_origin: iPSC | P28290 | ITPRID2 | 9 | Homo sapien | 1.32680328 | -2.3131821 | 4.96678863 | -2.6602627 | 0.7768001 | 0.44933101 | 0.99999785 | -6.0800878 | up |
| KI - WT, cell_origin: iPSC | O14618 | CCS | 6 | Homo sapien | -0.3842453 | -1.4388622 | 0.67037166 | 0.12954148 | -0.7764547 | 0.44952872 | 0.99999785 | -6.0803549 | down |
| KI - WT, cell_origin: iPSC | P05129 | PRKCG | 2 | Homo sapien | 0.36219235 | -0.6320782 | 1.3564629 | -1.8633364 | 0.7758997 | 0.44977382 | 0.99999785 | -6.0808635 | up |
| KI - WT, cell_origin: iPSC | Q13114 | TRAF3 | 3 | Homo sapien | 0.49889814 | -0.8716881 | 1.86948439 | -3.0326507 | 0.77532723 | 0.45010459 | 0.99999785 | -6.0813027 | up |
| KI - WT, cell_origin: iPSC | Q9UM82 | SPATA2 | 4 | Homo sapien | -0.5083518 | -1.9067333 | 0.89002965 | -2.2843065 | -0.7746566 | 0.45054901 | 0.99999785 | -6.0817546 | down |
| KI - WT, cell_origin: iPSC | Q9UKY7 | CDV3 | 3 | Homo sapien | 0.55217623 | -0.9685252 | 2.07287762 | -1.0495906 | 0.7738132 | 0.45104277 | 0.99999785 | -6.0823941 | up |
| KI - WT, cell_origin: iPSC | Q96L92 | SNX27 | 4 | Homo sapien | 0.52824644 | -0.9269673 | 1.98346017 | -2.0412178 | 0.77359234 | 0.4511695 | 0.99999785 | -6.0825643 | up |
| KI - WT, cell_origin: iPSC | P50552 | VASP | 12 | Homo sapien | 0.25796936 | -0.4528469 | 0.96878562 | 0.5569874 | 0.77295894 | 0.45145271 | 0.99999785 | -6.0831402 | up |
| KI - WT, cell_origin: iPSC | Q14004 | CDK13 | 4 | Homo sapien | 0.37444856 | -0.6583162 | 1.40721336 | -1.9576501 | 0.77227936 | 0.45185499 | 0.99999785 | -6.0836501 | up |
| KI - WT, cell_origin: iPSC | Q9Y2D4 | EXOC6B | 9 | Homo sapien | -0.3430413 | -1.2899643 | 0.6038818 | -0.9619867 | -0.7718166 | 0.45215178 | 0.99999785 | -6.0839722 | down |
| KI - WT, cell_origin: iPSC | P17980 | PSMC3 | 9 | Homo sapien | 0.30212305 | -0.5327003 | 1.1369464 | -0.4772467 | 0.77091623 | 0.4526493 | 0.99999785 | -6.0846862 | up |
| KI - WT, cell_origin: iPSC | Q96DH6 | MSI2 | 6 | Homo sapien | -0.2971811 | -1.1185519 | 0.52418967 | 1.40583692 | -0.7707258 | 0.45275886 | 0.99999785 | -6.0848324 | down |
| KI - WT, cell_origin: iPSC | Q8IVF7 | FMNL3 | 23 | Homo sapien | -0.873531 | -3.2892662 | 1.54220416 | -1.4577905 | -0.7706042 | 0.45288632 | 0.99999785 | -6.0848627 | down |
| KI - WT, cell_origin: iPSC | P30291 | WEE1 | 2 | Homo sapien | 0.31258447 | -0.5525605 | 1.1777294 | -2.470023 | 0.76951699 | 0.45343017 | 0.99999785 | -6.0857866 | up |
| KI - WT, cell_origin: iPSC | Q9Y3D2 | MSRB2 | 3 | Homo sapien | -0.2622015 | -0.9879556 | 0.46355265 | -1.3787457 | -0.7694587 | 0.45346374 | 0.99999785 | -6.0858313 | down |
| KI - WT, cell_origin: iPSC | Q66K74 | MAP1S | 14 | Homo sapien | 0.20542499 | -0.3636357 | 0.7744857 | -0.3658399 | 0.76883716 | 0.45382175 | 0.99999785 | -6.0863076 | up |
| KI - WT, cell_origin: iPSC | Q86WT1 | TTC30A | 5 | Homo sapien | 0.20484025 | -0.363101 | 0.77278149 | -1.3798939 | 0.76815982 | 0.45421211 | 0.99999785 | -6.0868263 | up |
| KI - WT, cell_origin: iPSC | O60293 | ZFC3H1 | 2 | Homo sapien | 0.442778 | -0.7852764 | 1.67083241 | -3.483203 | 0.76790609 | 0.45435839 | 0.99999785 | -6.0870205 | up |
| KI - WT, cell_origin: iPSC | Q8TCT9 | HM13 | 6 | Homo sapien | -0.3653313 | -1.3808809 | 0.65021819 | -0.4865391 | -0.7666342 | 0.45517357 | 0.99999785 | -6.0879038 | down |
| KI - WT, cell_origin: iPSC | Q9BV38 | WDR18 | 5 | Homo sapien | -0.4761179 | -1.800325 | 0.84808916 | -0.7102667 | -0.7662332 | 0.45540497 | 0.99999785 | -6.0882101 | down |
| KI - WT, cell_origin: iPSC | P33240 | CSTF2 | 3 | Homo sapien | -0.4109272 | -1.5558814 | 0.73402693 | -1.4252436 | -0.7648551 | 0.45620086 | 0.99999785 | -6.0892618 | down |
| KI - WT, cell_origin: iPSC | Q53H47 | SETMAR | 4 | Homo sapien | -0.6233735 | -2.361772 | 1.115025 | -2.0934816 | -0.76419 | 0.45658529 | 0.99999785 | -6.0897688 | down |
| KI - WT, cell_origin: iPSC | P50452 | SERPINB8 | 7 | Homo sapien | 0.30300773 | -0.5432801 | 1.14929559 | -0.1665789 | 0.76269957 | 0.45739061 | 0.99999785 | -6.0909657 | up |

| | | | | | | | | | | | | | |
|---|---|---|---|---|---|---|---|---|---|---|---|---|---|
| KI - WT, cell_origin: iPSC | Q01543 | FLI1 | 5 | Homo sapien | -0.206687 | -0.7838571 | 0.37048313 | -0.888815 | -0.7626916 | 0.45737111 | 0.99999785 | -6.0909982 | down |
| KI - WT, cell_origin: iPSC | Q86YP4 | GATAD2A | 2 | Homo sapien | 0.33989429 | -0.6095975 | 1.28938605 | -2.0214759 | 0.76241661 | 0.45753032 | 0.99999785 | -6.0912073 | up |
| KI - WT, cell_origin: iPSC | Q96MW1 | CCDC43 | 3 | Homo sapien | -0.6574902 | -2.496408 | 1.18142761 | -0.8353633 | -0.761955 | 0.45787857 | 0.99999785 | -6.0914694 | down |
| KI - WT, cell_origin: iPSC | P60891 | PRPS1 | 8 | Homo sapien | 0.35810125 | -0.645322 | 1.3615245 | 1.09211848 | 0.76054346 | 0.45869653 | 0.99999785 | -6.0925411 | up |
| KI - WT, cell_origin: iPSC | Q92626 | PXDN | 51 | Homo sapien | -0.8338562 | -3.1723241 | 1.50461172 | -1.7030443 | -0.7599099 | 0.45906395 | 0.99999785 | -6.0930215 | down |
| KI - WT, cell_origin: iPSC | O15523 | DDX3Y | 11 | Homo sapien | -0.2261032 | -0.860056 | 0.40784958 | 3.64815083 | -0.7596505 | 0.45914128 | 0.99999785 | -6.0932983 | down |
| KI - WT, cell_origin: iPSC | O14929 | HAT1 | 6 | Homo sapien | -0.2350837 | -0.8956083 | 0.42544084 | 0.59522843 | -0.7580071 | 0.4600881 | 0.99999785 | -6.0945503 | down |
| KI - WT, cell_origin: iPSC | P00441 | SOD1 | 2 | Homo sapien | 0.42738116 | -0.7763481 | 1.63111038 | -1.1444021 | 0.75663937 | 0.46096355 | 0.99999785 | -6.0954956 | up |
| KI - WT, cell_origin: iPSC | P26583 | HMGB2 | 7 | Homo sapien | -0.7602868 | -2.9050057 | 1.38443213 | 1.5617569 | -0.7554564 | 0.4616518 | 0.99999785 | -6.096388 | down |
| KI - WT, cell_origin: iPSC | P05114 | HMGN1 | 5 | Homo sapien | -0.6644061 | -2.5391574 | 1.21034515 | 3.00159449 | -0.7552528 | 0.46177036 | 0.99999785 | -6.0965415 | down |
| KI - WT, cell_origin: iPSC | O60645 | EXOC3 | 12 | Homo sapien | 0.80712354 | -1.4725721 | 3.08681916 | -2.4877712 | 0.7545109 | 0.4622024 | 0.99999785 | -6.0971004 | up |
| KI - WT, cell_origin: iPSC | P09486 | SPARC | 2 | Homo sapien | -0.2779679 | -1.0631584 | 0.50722268 | -1.4412505 | -0.7539789 | 0.46243242 | 0.99999785 | -6.0975885 | down |
| KI - WT, cell_origin: iPSC | P62888 | RPL30 | 7 | Homo sapien | -0.1784521 | -0.6827419 | 0.32583767 | 3.82899644 | -0.75367 | 0.46261249 | 0.99999785 | -6.0978209 | down |
| KI - WT, cell_origin: iPSC | P02747 | C1QC | 2 | Homo sapien | -0.2107212 | -0.8062754 | 0.384833 | 0.35949338 | -0.7535752 | 0.46266774 | 0.99999785 | -6.0978922 | down |
| KI - WT, cell_origin: iPSC | P48736 | PIK3CG | 15 | Homo sapien | -0.3874587 | -1.4838691 | 0.70895166 | 0.84788814 | -0.7531035 | 0.46302269 | 0.99999785 | -6.0981593 | down |
| KI - WT, cell_origin: iPSC | Q16576 | RBBP7 | 3 | Homo sapien | -0.4485122 | -1.7183082 | 0.82128384 | -1.0225437 | -0.7527359 | 0.46323707 | 0.99999785 | -6.0984355 | down |
| KI - WT, cell_origin: iPSC | P18077 | RPL35A | 9 | Homo sapien | -0.1869816 | -0.7162393 | 0.34227602 | 3.83224984 | -0.7524393 | 0.46333034 | 0.99999785 | -6.0987458 | down |
| KI - WT, cell_origin: iPSC | O00750 | PIK3C2B | 4 | Homo sapien | -0.394283 | -1.5107305 | 0.72216442 | -1.8413969 | -0.7522942 | 0.46343881 | 0.99999785 | -6.0988287 | down |
| KI - WT, cell_origin: iPSC | P23219 | PTGS1 | 6 | Homo sapien | 0.57758179 | -1.0620542 | 2.2172178 | -1.8508836 | 0.75070345 | 0.4644236 | 0.99999785 | -6.0999608 | up |
| KI - WT, cell_origin: iPSC | Q8TBB5 | KLHDC4 | 3 | Homo sapien | -0.4258808 | -1.6351161 | 0.7833544 | -2.5562928 | -0.7501651 | 0.46467078 | 0.99999785 | -6.100438 | down |
| KI - WT, cell_origin: iPSC | Q460N5 | PARP14 | 37 | Homo sapien | -0.5104952 | -1.9617374 | 0.94074706 | -0.3550676 | -0.7496423 | 0.46504382 | 0.99999785 | -6.1007556 | down |
| KI - WT, cell_origin: iPSC | P60228 | EIF3E | 23 | Homo sapien | -0.1915736 | -0.7359162 | 0.35276898 | 2.73866367 | -0.7495542 | 0.46501591 | 0.99999785 | -6.1009085 | down |
| KI - WT, cell_origin: iPSC | Q15154 | PCM1 | 10 | Homo sapien | 0.33839891 | -0.6243335 | 1.30113131 | -1.1395278 | 0.74883256 | 0.46547486 | 0.99999785 | -6.101408 | up |
| KI - WT, cell_origin: iPSC | P61353 | RPL27 | 5 | Homo sapien | -0.1519126 | -0.5840346 | 0.28020929 | 3.18456964 | -0.7487336 | 0.46549605 | 0.99999785 | -6.1015223 | down |
| KI - WT, cell_origin: iPSC | Q96PV6 | LENG8 | 3 | Homo sapien | 0.22288028 | -0.4113993 | 0.8571599 | -0.2019875 | 0.74839446 | 0.46569453 | 0.99999785 | -6.1017758 | up |
| KI - WT, cell_origin: iPSC | Q14451 | GRB7 | 6 | Homo sapien | 0.42108648 | -0.7780462 | 1.62021917 | -1.5619434 | 0.74835287 | 0.46579814 | 0.99999785 | -6.1017199 | up |
| KI - WT, cell_origin: iPSC | P49841 | GSK3B | 2 | Homo sapien | -0.4051999 | -1.5593569 | 0.74895715 | -2.6090134 | -0.7477288 | 0.46608434 | 0.99999785 | -6.102273 | down |
| KI - WT, cell_origin: iPSC | P25705 | ATP5F1A | 13 | Homo sapien | 0.80588944 | -1.4928535 | 3.10463237 | -0.8722566 | 0.74711495 | 0.46652305 | 0.99999785 | -6.1026444 | up |
| KI - WT, cell_origin: iPSC | Q9UJX6 | ANAPC2 | 3 | Homo sapien | 0.42290523 | -0.7843415 | 1.63015192 | -3.6174043 | 0.74608227 | 0.4670493 | 0.99999785 | -6.103501 | up |
| KI - WT, cell_origin: iPSC | Q8TB72 | PUM2 | 9 | Homo sapien | -0.2479027 | -0.9557775 | 0.45997208 | -0.2572848 | -0.7458723 | 0.46717242 | 0.99999785 | -6.1036574 | down |
| KI - WT, cell_origin: iPSC | P18085 | ARF4 | 6 | Homo sapien | 0.38398425 | -0.7138071 | 1.48177557 | 2.75965454 | 0.74541133 | 0.46752178 | 0.99999785 | -6.1039142 | up |
| KI - WT, cell_origin: iPSC | Q93074 | MED12 | 4 | Homo sapien | 0.3219743 | -0.5979904 | 1.24193901 | -1.8714031 | 0.74540058 | 0.46744917 | 0.99999785 | -6.1040087 | up |
| KI - WT, cell_origin: iPSC | O15111 | CHUK | 2 | Homo sapien | -0.3242067 | -1.2508201 | 0.60240677 | -1.8239746 | -0.7451832 | 0.46757675 | 0.99999785 | -6.1041705 | down |
| KI - WT, cell_origin: iPSC | P13797 | PLS3 | 22 | Homo sapien | -0.2894026 | -1.1167712 | 0.53796604 | 1.48542239 | -0.7451115 | 0.4676423 | 0.99999785 | -6.1041981 | down |
| KI - WT, cell_origin: iPSC | Q13541 | EIF4EBP1 | 3 | Homo sapien | -0.3770609 | -1.455504 | 0.70138228 | 1.19953681 | -0.7451035 | 0.4677024 | 0.99999785 | -6.1041434 | down |
| KI - WT, cell_origin: iPSC | P63010 | AP2B1 | 38 | Homo sapien | 0.23991238 | -0.4461964 | 0.92602112 | 2.64827947 | 0.74473091 | 0.4678422 | 0.99999785 | -6.104507 | up |
| KI - WT, cell_origin: iPSC | Q969S8 | HDAC10 | 6 | Homo sapien | 0.34281137 | -0.6383621 | 1.3239848 | -1.0901454 | 0.74441672 | 0.46807677 | 0.99999785 | -6.1046858 | up |
| KI - WT, cell_origin: iPSC | P61158 | ACTR3 | 11 | Homo sapien | 0.21675937 | -0.4036272 | 0.83714592 | 0.07843566 | 0.74414093 | 0.46818862 | 0.99999785 | -6.1049457 | up |
| KI - WT, cell_origin: iPSC | Q9UHX1 | PUF60 | 19 | Homo sapien | -0.5586711 | -2.1601034 | 1.04276109 | 2.20805078 | -0.743447 | 0.46867494 | 0.99999785 | -6.105375 | down |
| KI - WT, cell_origin: iPSC | Q8N5J2 | MINDY1 | 2 | Homo sapien | 0.59346155 | -1.1072406 | 2.29416366 | -3.5505332 | 0.74333099 | 0.4686879 | 0.99999785 | -6.1055217 | up |
| KI - WT, cell_origin: iPSC | P25205 | MCM3 | 49 | Homo sapien | -0.3434518 | -1.3281459 | 0.64124223 | 2.27369179 | -0.7433047 | 0.46875858 | 0.99999785 | -6.1054807 | down |
| KI - WT, cell_origin: iPSC | Q8TDD1 | DDX54 | 13 | Homo sapien | -0.3064205 | -1.1851363 | 0.57229522 | 0.16306199 | -0.7429369 | 0.46893886 | 0.99999785 | -6.1057932 | down |
| KI - WT, cell_origin: iPSC | Q9UL25 | RAB21 | 6 | Homo sapien | 0.42786071 | -0.7995833 | 1.65530472 | -1.7102022 | 0.74285335 | 0.46902382 | 0.99999785 | -6.1058158 | up |
| KI - WT, cell_origin: iPSC | P01023 | A2M | 5 | Homo sapien | -0.6564496 | -2.5411311 | 1.22823179 | 0.19867757 | -0.7422767 | 0.46936283 | 0.99999785 | -6.1062437 | down |
| KI - WT, cell_origin: iPSC | Q9BRL6 | SRSF8 | 3 | Homo sapien | -0.3782761 | -1.4660745 | 0.70952241 | 0.97722648 | -0.741076 | 0.47006918 | 0.99999785 | -6.1071335 | down |
| KI - WT, cell_origin: iPSC | P35232 | PHB | 6 | Homo sapien | 1.24273658 | -2.3315121 | 4.81698521 | -2.700911 | 0.74096329 | 0.47013555 | 0.99999785 | -6.107217 | up |

| | | | | | | | | | | | | | |
|---|---|---|---|---|---|---|---|---|---|---|---|---|---|
| KI - WT, cell_origin: iPSC | Q86XP3 | DDX42 | 23 | Homo sapien | -0.4943088 | -1.91646 | 0.92784231 | 0.32867252 | -0.7407216 | 0.47027784 | 0.99999785 | -6.107396 | down |
| KI - WT, cell_origin: iPSC | O15498 | YKT6 | 8 | Homo sapien | 0.33186256 | -0.6231008 | 1.28682589 | 0.23936569 | 0.74058332 | 0.47035924 | 0.99999785 | -6.1074983 | up |
| KI - WT, cell_origin: iPSC | Q96S55 | WRNIP1 | 11 | Homo sapien | -0.2852822 | -1.1072201 | 0.53665577 | -0.8334254 | -0.739318 | 0.4710431 | 0.99999785 | -6.1085015 | down |
| KI - WT, cell_origin: iPSC | Q8IXJ6 | SIRT2 | 15 | Homo sapien | 0.54032497 | -1.0172 | 2.09784989 | 1.10254323 | 0.73930288 | 0.47111352 | 0.99999785 | -6.1084452 | up |
| KI - WT, cell_origin: iPSC | Q8NHV4 | NEDD1 | 12 | Homo sapien | -0.4252619 | -1.651326 | 0.80080207 | 0.13316741 | -0.7391724 | 0.47119042 | 0.99999785 | -6.1085416 | down |
| KI - WT, cell_origin: iPSC | Q5MNZ6 | WDR45B | 3 | Homo sapien | 0.34396355 | -0.6484798 | 1.33640693 | -0.845152 | 0.73854537 | 0.47155063 | 0.99999785 | -6.1090151 | up |
| KI - WT, cell_origin: iPSC | Q9H773 | DCTPP1 | 3 | Homo sapien | -0.2019193 | -0.7847704 | 0.38093175 | 0.62802421 | -0.7378362 | 0.47190042 | 0.99999785 | -6.1096135 | down |
| KI - WT, cell_origin: iPSC | P51665 | PSMD7 | 4 | Homo sapien | 0.40798794 | -0.7717167 | 1.58769262 | -1.9961541 | 0.73675349 | 0.4725718 | 0.99999785 | -6.1103761 | up |
| KI - WT, cell_origin: iPSC | Q63HN8 | RNF213 | 108 | Homo sapien | 0.18421644 | -0.3484622 | 0.71689505 | 1.22647766 | 0.73655096 | 0.47265926 | 0.99999785 | -6.1105605 | up |
| KI - WT, cell_origin: iPSC | P01024 | C3 | 9 | Homo sapien | -0.5767378 | -2.2494585 | 1.0959829 | -0.9882443 | -0.73478 | 0.47378371 | 0.99999785 | -6.1117778 | down |
| KI - WT, cell_origin: iPSC | Q8NEZ3 | WDR19 | 4 | Homo sapien | -0.1949894 | -0.7603864 | 0.37040757 | -1.1440388 | -0.7345091 | 0.47386634 | 0.99999785 | -6.1120619 | down |
| KI - WT, cell_origin: iPSC | Q9NR31 | SAR1A | 9 | Homo sapien | 0.35644436 | -0.6778212 | 1.39070989 | 5.10727355 | 0.73444972 | 0.47397908 | 0.99999785 | -6.1120205 | up |
| KI - WT, cell_origin: iPSC | P11142 | HSPA8 | 23 | Homo sapien | 0.21885812 | -0.4161597 | 0.85387589 | 5.34444854 | 0.73421787 | 0.47407072 | 0.99999785 | -6.1122406 | up |
| KI - WT, cell_origin: iPSC | Q8WVK2 | SNRNP27 | 6 | Homo sapien | -0.3875608 | -1.5136397 | 0.73851813 | 2.90430599 | -0.7334548 | 0.47456782 | 0.99999785 | -6.1127507 | down |
| KI - WT, cell_origin: iPSC | P14317 | HCLS1 | 7 | Homo sapien | 0.4755562 | -0.9065225 | 1.85763491 | -1.7705979 | 0.7332827 | 0.47466973 | 0.99999785 | -6.1128769 | up |
| KI - WT, cell_origin: iPSC | O95801 | TTC4 | 2 | Homo sapien | -0.4322034 | -1.6883034 | 0.82389667 | -1.793852 | -0.733274 | 0.47467488 | 0.99999785 | -6.1128833 | down |
| KI - WT, cell_origin: iPSC | Q92844 | TANK | 3 | Homo sapien | -0.2117676 | -0.8270458 | 0.40351064 | -0.3334705 | -0.7330399 | 0.47473606 | 0.99999785 | -6.1131398 | down |
| KI - WT, cell_origin: iPSC | Q9UEG4 | ZNF629 | 6 | Homo sapien | 0.93866611 | -1.7903466 | 3.66767878 | -3.2692644 | 0.73300672 | 0.47483314 | 0.99999785 | -6.1130793 | up |
| KI - WT, cell_origin: iPSC | O60313 | OPA1 | 25 | Homo sapien | -0.5270377 | -2.0607847 | 1.00670925 | -1.2254617 | -0.7323022 | 0.47525044 | 0.99999785 | -6.1135955 | down |
| KI - WT, cell_origin: iPSC | P67936 | TPM4 | 15 | Homo sapien | -0.2703262 | -1.0567742 | 0.51612182 | 2.15717565 | -0.7322106 | 0.47525042 | 0.99999785 | -6.1137221 | down |
| KI - WT, cell_origin: iPSC | O14730 | RIOK3 | 5 | Homo sapien | 0.32780378 | -0.6288362 | 1.28444376 | -0.9882674 | 0.73005025 | 0.47655213 | 0.99999785 | -6.1152796 | up |
| KI - WT, cell_origin: iPSC | Q15424 | SAFB | 7 | Homo sapien | -0.2598386 | -1.0178899 | 0.49821269 | -1.0064297 | -0.7300368 | 0.47651678 | 0.99999785 | -6.1153368 | down |
| KI - WT, cell_origin: iPSC | P61088 | UBE2N | 7 | Homo sapien | -0.2693403 | -1.0567529 | 0.51807235 | 0.41353525 | -0.7286278 | 0.47737335 | 0.99999785 | -6.1163433 | down |
| KI - WT, cell_origin: iPSC | P30048 | PRDX3 | 6 | Homo sapien | 0.7794234 | -1.5023611 | 3.06120784 | -0.6954563 | 0.7279494 | 0.47783372 | 0.99999785 | -6.1167749 | up |
| KI - WT, cell_origin: iPSC | Q96RE7 | NACC1 | 11 | Homo sapien | -0.211464 | -0.8301733 | 0.40724523 | 0.95255865 | -0.7279299 | 0.47776849 | 0.99999785 | -6.1168732 | down |
| KI - WT, cell_origin: iPSC | O60343 | TBC1D4 | 20 | Homo sapien | 0.2798358 | -0.5396056 | 1.09927723 | 0.09832547 | 0.72745017 | 0.47807666 | 0.99999785 | -6.1171974 | up |
| KI - WT, cell_origin: iPSC | Q9BYB0 | SHANK3 | 61 | Homo sapien | -0.3716566 | -1.4624297 | 0.71911658 | 3.09806119 | -0.7261223 | 0.47892059 | 0.99999785 | -6.1181042 | down |
| KI - WT, cell_origin: iPSC | Q9BPX5 | ARPC5L | 3 | Homo sapien | 0.34314228 | -0.6643957 | 1.35068026 | -0.6841373 | 0.72579699 | 0.47911424 | 0.99999785 | -6.1183405 | up |
| KI - WT, cell_origin: iPSC | P83876 | TXNL4A | 3 | Homo sapien | 0.27617434 | -0.5346172 | 1.08696583 | -1.2268474 | 0.72546058 | 0.47923806 | 0.99999785 | -6.1186686 | up |
| KI - WT, cell_origin: iPSC | Q9BYM8 | RBCK1 | 5 | Homo sapien | 0.43301241 | -0.8393617 | 1.70538654 | -0.8078901 | 0.72525023 | 0.47943985 | 0.99999785 | -6.1187375 | up |
| KI - WT, cell_origin: iPSC | Q9H1J1 | UPF3A | 3 | Homo sapien | -0.4212763 | -1.6592502 | 0.81669763 | -2.2482958 | -0.7248922 | 0.47959949 | 0.99999785 | -6.1190561 | down |
| KI - WT, cell_origin: iPSC | P63000 | RAC1 | 12 | Homo sapien | 0.24725577 | -0.4797417 | 0.97425324 | 5.86000515 | 0.72479619 | 0.47971034 | 0.99999785 | -6.119067 | up |
| KI - WT, cell_origin: iPSC | P01111 | NRAS | 4 | Homo sapien | 0.46579226 | -0.9041218 | 1.8357063 | 2.00305366 | 0.724605 | 0.47982427 | 0.99999785 | -6.1192057 | up |
| KI - WT, cell_origin: iPSC | Q8TBC3 | SHKBP1 | 13 | Homo sapien | 0.24453149 | -0.4753862 | 0.96444918 | 1.51808908 | 0.72345848 | 0.48043793 | 0.99999785 | -6.1201131 | up |
| KI - WT, cell_origin: iPSC | P50897 | PPT1 | 5 | Homo sapien | 0.27402011 | -0.5335664 | 1.0816066 | -0.3550117 | 0.72277013 | 0.48086176 | 0.99999785 | -6.120597 | up |
| KI - WT, cell_origin: iPSC | Q9Y6A5 | TACC3 | 4 | Homo sapien | -0.2226755 | -0.8788792 | 0.43352828 | -1.1589499 | -0.7227254 | 0.48086897 | 0.99999785 | -6.1206506 | down |
| KI - WT, cell_origin: iPSC | P17028 | ZNF24 | 3 | Homo sapien | -0.3043854 | -1.2015536 | 0.59278273 | -1.0098495 | -0.7227164 | 0.48089706 | 0.99999785 | -6.1206323 | down |
| KI - WT, cell_origin: iPSC | Q14697 | GANAB | 7 | Homo sapien | -0.5054691 | -1.9971449 | 0.98620657 | -2.5555225 | -0.722089 | 0.48131581 | 0.99999785 | -6.1210376 | down |
| KI - WT, cell_origin: iPSC | Q7KZF4 | SND1 | 10 | Homo sapien | -0.2260771 | -0.8930054 | 0.44085126 | -0.469598 | -0.7219664 | 0.48132213 | 0.99999785 | -6.1211994 | down |
| KI - WT, cell_origin: iPSC | Q9P032 | NDUFAF4 | 3 | Homo sapien | 0.74669972 | -1.4584211 | 2.95182056 | -2.5904642 | 0.72163229 | 0.48159777 | 0.99999785 | -6.1213576 | up |
| KI - WT, cell_origin: iPSC | Q9NVC6 | MED17 | 6 | Homo sapien | 0.28603054 | -0.5582619 | 1.13032299 | -1.3250385 | 0.72153796 | 0.48157808 | 0.99999785 | -6.121509 | up |
| KI - WT, cell_origin: iPSC | P84101 | SERF2 | 3 | Homo sapien | -0.794791 | -3.1427202 | 1.55313819 | 3.04450623 | -0.7213903 | 0.48174229 | 0.99999785 | -6.1215324 | down |
| KI - WT, cell_origin: iPSC | Q9NZQ3 | NCKIPSD | 2 | Homo sapien | 0.36604515 | -0.7149131 | 1.44700338 | -1.7058545 | 0.72134604 | 0.48171538 | 0.99999785 | -6.1216228 | up |
| KI - WT, cell_origin: iPSC | Q14185 | DOCK1 | 6 | Homo sapien | 0.43007483 | -0.8425522 | 1.70270181 | -0.9109589 | 0.72018698 | 0.48246145 | 0.99999785 | -6.1224009 | up |
| KI - WT, cell_origin: iPSC | Q96K58 | ZNF668 | 2 | Homo sapien | -0.3048699 | -1.2085436 | 0.59880386 | -1.9178255 | -0.718526 | 0.48337946 | 0.99999785 | -6.1236802 | down |
| KI - WT, cell_origin: iPSC | Q96NC0 | ZMAT2 | 5 | Homo sapien | -0.7794531 | -3.0944952 | 1.53558893 | -1.5315174 | -0.7175191 | 0.48405812 | 0.99999785 | -6.1243215 | down |

| | | | | | | | | | | | | |
|---|---|---|---|---|---|---|---|---|---|---|---|---|
| KI - WT, cell_origin: iPSC | Q14978 | NOLC1 | 11 | Homo sapien | -0.5799161 | -2.3034914 | 1.14365912 | 0.01739575 | -0.71703 | 0.48435122 | 0.99999785 | -6.1246729 down |
| KI - WT, cell_origin: iPSC | Q14331 | FRG1 | 14 | Homo sapien | -0.5021326 | -1.9961816 | 0.9919165 | 2.17213794 | -0.7162356 | 0.48482743 | 0.99999785 | -6.1252432 down |
| KI - WT, cell_origin: iPSC | Q96M27 | PRRC1 | 8 | Homo sapien | 0.53224188 | -1.0520349 | 2.11651862 | -0.0584036 | 0.71594617 | 0.485001 | 0.99999785 | -6.1254508 up |
| KI - WT, cell_origin: iPSC | O60502 | OGA | 4 | Homo sapien | -0.2385218 | -0.9480977 | 0.47105413 | -0.8328625 | -0.7159272 | 0.48493701 | 0.99999785 | -6.1255468 down |
| KI - WT, cell_origin: iPSC | Q02539 | H1-1 | 2 | Homo sapien | -0.3638008 | -1.4473841 | 0.71978244 | -1.077574 | -0.7154903 | 0.48527446 | 0.99999785 | -6.1257776 down |
| KI - WT, cell_origin: iPSC | Q32MZ4 | LRRFIP1 | 10 | Homo sapien | 0.53346598 | -1.0568074 | 2.12373935 | -1.4554609 | 0.71488686 | 0.4856366 | 0.99999785 | -6.12621 up |
| KI - WT, cell_origin: iPSC | Q92835 | INPP5D | 5 | Homo sapien | 0.34679606 | -0.687158 | 1.38075009 | -0.5918467 | 0.71478478 | 0.48569787 | 0.99999785 | -6.1262831 up |
| KI - WT, cell_origin: iPSC | Q9Y4G8 | RAPGEF2 | 48 | Homo sapien | 0.2992118 | -0.5931395 | 1.19156308 | 0.70621351 | 0.71448129 | 0.48586452 | 0.99999785 | -6.1265174 up |
| KI - WT, cell_origin: iPSC | Q9NWQ9 | C14orf119 | 2 | Homo sapien | 0.37884188 | -0.7512061 | 1.50888982 | -1.9386415 | 0.71413287 | 0.48603654 | 0.99999785 | -6.1268074 up |
| KI - WT, cell_origin: iPSC | P12268 | IMPDH2 | 15 | Homo sapien | -0.3201954 | -1.2762476 | 0.63585675 | 0.98704814 | -0.7137332 | 0.48632938 | 0.99999785 | -6.1270356 down |
| KI - WT, cell_origin: iPSC | Q70Z53 | FRA10AC1 | 6 | Homo sapien | -0.8121973 | -3.2380048 | 1.61361031 | -1.5854828 | -0.7135223 | 0.48645609 | 0.99999785 | -6.1271864 down |
| KI - WT, cell_origin: iPSC | Q6PJT7 | ZC3H14 | 6 | Homo sapien | -0.2632274 | -1.049096 | 0.52264112 | -0.1972553 | -0.713472 | 0.48642728 | 0.99999785 | -6.127287 down |
| KI - WT, cell_origin: iPSC | P46939 | UTRN | 27 | Homo sapien | 0.29682814 | -0.5903303 | 1.18398653 | -0.4519116 | 0.7128209 | 0.48684171 | 0.99999785 | -6.1277269 up |
| KI - WT, cell_origin: iPSC | P47224 | RABIF | 2 | Homo sapien | 0.46135193 | -0.9181844 | 1.84088828 | 0.42254729 | 0.71269148 | 0.48695543 | 0.99999785 | -6.1277801 up |
| KI - WT, cell_origin: iPSC | Q8WUA2 | PPIL4 | 21 | Homo sapien | -0.5220974 | -2.0868358 | 1.04264106 | 4.07885779 | -0.7110696 | 0.48793109 | 0.99999785 | -6.1289371 down |
| KI - WT, cell_origin: iPSC | Q9Y4F5 | CEP170B | 7 | Homo sapien | 0.29701727 | -0.5958494 | 1.1898839 | -2.0018525 | 0.70849174 | 0.48940982 | 0.99999785 | -6.1308525 up |
| KI - WT, cell_origin: iPSC | P48200 | IREB2 | 2 | Homo sapien | 0.42237837 | -0.8484068 | 1.69316351 | -1.5288882 | 0.70832393 | 0.48958549 | 0.99999785 | -6.1308903 up |
| KI - WT, cell_origin: iPSC | Q53SF7 | COBLL1 | 3 | Homo sapien | 0.47737365 | -0.9596549 | 1.91440222 | -2.2623136 | 0.70793831 | 0.48981811 | 0.99999785 | -6.1311641 up |
| KI - WT, cell_origin: iPSC | Q01780 | EXOSC10 | 12 | Homo sapien | -0.3112984 | -1.2486309 | 0.62603412 | -0.6966308 | -0.7076447 | 0.48997536 | 0.99999785 | -6.1313941 down |
| KI - WT, cell_origin: iPSC | O43318 | MAP3K7 | 2 | Homo sapien | 0.40793049 | -0.8206587 | 1.63651966 | -0.9817021 | 0.70759027 | 0.49002811 | 0.99999785 | -6.131411 up |
| KI - WT, cell_origin: iPSC | Q2Q1W2 | TRIM71 | 36 | Homo sapien | -0.4018742 | -1.6132621 | 0.8095137 | 1.51482761 | -0.7069835 | 0.49039436 | 0.99999785 | -6.1318413 down |
| KI - WT, cell_origin: iPSC | Q14653 | IRF3 | 2 | Homo sapien | 0.57957985 | -1.1693208 | 2.3284805 | -0.5378104 | 0.70623706 | 0.49084513 | 0.99999785 | -6.1323701 up |
| KI - WT, cell_origin: iPSC | Q92558 | WASF1 | 2 | Homo sapien | 0.26083463 | -0.5262472 | 1.04791649 | -1.2080869 | 0.70580547 | 0.49103175 | 0.99999785 | -6.1327566 up |
| KI - WT, cell_origin: iPSC | P61626 | LYZ | 2 | Homo sapien | -0.3301594 | -1.3276856 | 0.6673669 | 2.64347808 | -0.7053451 | 0.49138412 | 0.99999785 | -6.1330013 down |
| KI - WT, cell_origin: iPSC | P63208 | SKP1 | 10 | Homo sapien | 0.15165067 | -0.3068982 | 0.6101995 | 2.10958727 | 0.70436607 | 0.49190214 | 0.99999785 | -6.1337741 up |
| KI - WT, cell_origin: iPSC | P52272 | HNRNPM | 34 | Homo sapien | -0.2480189 | -0.9989655 | 0.5029277 | 2.55475735 | -0.7035298 | 0.49242713 | 0.99999785 | -6.1343438 down |
| KI - WT, cell_origin: iPSC | Q99081 | TCF12 | 6 | Homo sapien | 0.48326875 | -0.9818848 | 1.94842228 | -0.62106 | 0.7029233 | 0.49284927 | 0.99999785 | -6.1347115 up |
| KI - WT, cell_origin: iPSC | Q9HBF4 | ZFYVE1 | 3 | Homo sapien | 0.22519516 | -0.4576423 | 0.9080326 | -0.8463267 | 0.70239498 | 0.49309552 | 0.99999785 | -6.1351643 up |
| KI - WT, cell_origin: iPSC | P54646 | PRKAA2 | 4 | Homo sapien | -0.6071392 | -2.4529166 | 1.23863818 | -2.7321281 | -0.7009892 | 0.49402124 | 0.99999785 | -6.1360733 down |
| KI - WT, cell_origin: iPSC | Q5D862 | FLG2 | 6 | Homo sapien | 0.46491944 | -0.9486662 | 1.87850508 | -0.4426524 | 0.70090304 | 0.49407348 | 0.99999785 | -6.1361339 up |
| KI - WT, cell_origin: iPSC | Q8IYI6 | EXOC8 | 16 | Homo sapien | 0.58397901 | -1.1941901 | 2.36214808 | -0.5298262 | 0.69988478 | 0.49469119 | 0.99999785 | -6.1368494 up |
| KI - WT, cell_origin: iPSC | P37802 | TAGLN2 | 8 | Homo sapien | 0.25902165 | -0.529959 | 1.04800235 | 0.3225759 | 0.69932087 | 0.49497887 | 0.99999785 | -6.1373048 up |
| KI - WT, cell_origin: iPSC | Q9NQ29 | LUC7L | 9 | Homo sapien | -0.702989 | -2.8465707 | 1.44059282 | 1.59337744 | -0.6988932 | 0.49529315 | 0.99999785 | -6.1375452 down |
| KI - WT, cell_origin: iPSC | Q9UNZ5 | C19orf53 | 4 | Homo sapien | -0.6177145 | -2.5024604 | 1.26703141 | 1.68837023 | -0.6984533 | 0.49556036 | 0.99999785 | -6.1378536 down |
| KI - WT, cell_origin: iPSC | O00442 | RTCA | 8 | Homo sapien | -0.3020794 | -1.2255806 | 0.62142188 | -0.3797555 | -0.6969726 | 0.49644081 | 0.99999785 | -6.1389116 down |
| KI - WT, cell_origin: iPSC | Q16543 | CDC37 | 13 | Homo sapien | 0.25855406 | -0.5320898 | 1.04919787 | -0.0112556 | 0.69659007 | 0.49663863 | 0.99999785 | -6.139217 up |
| KI - WT, cell_origin: iPSC | Q13796 | SHROOM2 | 28 | Homo sapien | 0.30547525 | -0.6286125 | 1.23956305 | -1.6076687 | 0.69656109 | 0.49664627 | 0.99999785 | -6.1392481 up |
| KI - WT, cell_origin: iPSC | Q53G59 | KLHL12 | 3 | Homo sapien | -0.2429326 | -0.9858104 | 0.49994521 | -0.9946176 | -0.6964791 | 0.49668751 | 0.99999785 | -6.1393149 down |
| KI - WT, cell_origin: iPSC | Q8IXQ6 | PARP9 | 30 | Homo sapien | -0.3136833 | -1.2736132 | 0.64624651 | 0.21982001 | -0.6963929 | 0.49681295 | 0.99999785 | -6.1392956 down |
| KI - WT, cell_origin: iPSC | Q96TA1 | NIBAN2 | 5 | Homo sapien | 0.52103756 | -1.0748098 | 2.11688492 | -2.1742232 | 0.695793 | 0.49717798 | 0.99999785 | -6.1397146 up |
| KI - WT, cell_origin: iPSC | Q2M2I8 | AAK1 | 7 | Homo sapien | -0.4386267 | -1.7822829 | 0.90502947 | -1.2113341 | -0.6956796 | 0.49724698 | 0.99999785 | -6.1397938 down |
| KI - WT, cell_origin: iPSC | Q8IXW5 | RPAP2 | 5 | Homo sapien | -0.2520046 | -1.024144 | 0.52013473 | -1.9786482 | -0.6951084 | 0.49752193 | 0.99999785 | -6.1402719 down |
| KI - WT, cell_origin: iPSC | Q5SYE7 | NHSL1 | 11 | Homo sapien | 0.28956472 | -0.5983914 | 1.1775208 | -0.5560241 | 0.69471869 | 0.49779124 | 0.99999785 | -6.1405089 up |
| KI - WT, cell_origin: iPSC | Q9Y613 | FHOD1 | 4 | Homo sapien | 0.33014615 | -0.6832808 | 1.34357315 | -1.6707101 | 0.69393785 | 0.49825375 | 0.99999785 | -6.1410675 up |
| KI - WT, cell_origin: iPSC | Q12996 | CSTF3 | 12 | Homo sapien | -0.2842998 | -1.1570273 | 0.58842768 | -1.0027599 | -0.6939233 | 0.49826447 | 0.99999785 | -6.1410756 down |
| KI - WT, cell_origin: iPSC | Q8NAV1 | PRPF38A | 10 | Homo sapien | -0.6093981 | -2.4813516 | 1.26255538 | 0.83060131 | -0.6937586 | 0.4984171 | 0.99999785 | -6.1411333 down |

| | | | | | | | | | | | | | |
|---|---|---|---|---|---|---|---|---|---|---|---|---|---|
| KI - WT, cell_origin: iPSC | P62280 | RPS11 | 14 | Homo sapien | -0.1310751 | -0.5335321 | 0.27138201 | 4.63614331 | -0.6936497 | 0.49841087 | 0.99999785 | -6.1412884 | down |
| KI - WT, cell_origin: iPSC | Q7L5D6 | GET4 | 5 | Homo sapien | 0.37135145 | -0.7696895 | 1.51239237 | -0.9997521 | 0.69356358 | 0.49853601 | 0.99999785 | -6.1412692 | up |
| KI - WT, cell_origin: iPSC | Q9P021 | CRIPT | 2 | Homo sapien | -0.5998459 | -2.4438551 | 1.2441633 | 2.29175326 | -0.6932326 | 0.4987378 | 0.99999785 | -6.1414995 | down |
| KI - WT, cell_origin: iPSC | Q9C037 | TRIM4 | 8 | Homo sapien | 0.35725984 | -0.7410589 | 1.45557864 | 0.02025195 | 0.69319937 | 0.49875807 | 0.99999785 | -6.1415227 | up |
| KI - WT, cell_origin: iPSC | P10589;P244 | NR2F1;NR2F2 | 2 | Homo sapien | 0.76360675 | -1.5847392 | 3.11195265 | -3.5946073 | 0.69296302 | 0.4989022 | 0.99999785 | -6.1416871 | up |
| KI - WT, cell_origin: iPSC | Q96EP5 | DAZAP1 | 7 | Homo sapien | -0.4206944 | -1.7166341 | 0.87524541 | 0.72373565 | -0.6918059 | 0.49960822 | 0.99999785 | -6.1424915 | down |
| KI - WT, cell_origin: iPSC | Q14012 | CAMK1 | 2 | Homo sapien | 0.51117784 | -1.0640274 | 2.08638304 | -2.0807043 | 0.69157176 | 0.49975113 | 0.99999785 | -6.1426541 | up |
| KI - WT, cell_origin: iPSC | Q9NY65 | TUBA8 | 2 | Homo sapien | -0.221418 | -0.9034537 | 0.4606177 | 4.00848448 | -0.6915503 | 0.49971341 | 0.99999785 | -6.1427244 | down |
| KI - WT, cell_origin: iPSC | Q9BQ70 | TCF25 | 4 | Homo sapien | -0.2332315 | -0.9518062 | 0.48534314 | -0.4696078 | -0.6912817 | 0.49985586 | 0.99999785 | -6.1429343 | down |
| KI - WT, cell_origin: iPSC | Q99541 | PLIN2 | 15 | Homo sapien | -0.7149204 | -2.9193308 | 1.48948997 | -0.6037222 | -0.6911425 | 0.50001322 | 0.99999785 | -6.142952 | down |
| KI - WT, cell_origin: iPSC | Q9UIG0 | BAZ1B | 11 | Homo sapien | -0.3950582 | -1.6129601 | 0.82284369 | -2.57491 | -0.6909275 | 0.50008434 | 0.99999785 | -6.1431668 | down |
| KI - WT, cell_origin: iPSC | Q96AE4 | FUBP1 | 17 | Homo sapien | -0.2278699 | -0.9304933 | 0.47475353 | 3.8210056 | -0.6908475 | 0.50014262 | 0.99999785 | -6.143212 | down |
| KI - WT, cell_origin: iPSC | Q96HD1 | CRELD1 | 2 | Homo sapien | 0.55733715 | -1.1628339 | 2.2775082 | -0.9161694 | 0.69047623 | 0.50042021 | 0.99999785 | -6.1434142 | up |
| KI - WT, cell_origin: iPSC | Q9H5H4 | ZNF768 | 4 | Homo sapien | -0.272326 | -1.1129775 | 0.56832544 | -0.9454269 | -0.690038 | 0.50063232 | 0.99999785 | -6.1437786 | down |
| KI - WT, cell_origin: iPSC | Q96EU6 | RRP36 | 4 | Homo sapien | 0.8744044 | -1.8293219 | 3.57813069 | -2.0146247 | 0.68921065 | 0.50119379 | 0.99999785 | -6.1442909 | up |
| KI - WT, cell_origin: iPSC | P57737 | CORO7 | 4 | Homo sapien | 0.20467265 | -0.4278525 | 0.83719775 | 0.40710511 | 0.68916263 | 0.50115103 | 0.99999785 | -6.1444027 | up |
| KI - WT, cell_origin: iPSC | Q7L1Q6 | BZW1 | 6 | Homo sapien | -0.2835941 | -1.1617839 | 0.59459574 | -0.5616933 | -0.6879386 | 0.50192778 | 0.99999785 | -6.1452187 | down |
| KI - WT, cell_origin: iPSC | P50613 | CDK7 | 3 | Homo sapien | 0.23119509 | -0.4851887 | 0.94757889 | -0.6579244 | 0.68734144 | 0.50226571 | 0.99999785 | -6.1456613 | up |
| KI - WT, cell_origin: iPSC | P98170 | XIAP | 5 | Homo sapien | -0.1758714 | -0.7210529 | 0.36931006 | -0.0894763 | -0.6870587 | 0.50243891 | 0.99999785 | -6.1458564 | down |
| KI - WT, cell_origin: iPSC | Q13148 | TARDBP | 11 | Homo sapien | -0.2904172 | -1.1920831 | 0.61124867 | 1.57474177 | -0.686317 | 0.50295032 | 0.99999785 | -6.1463059 | down |
| KI - WT, cell_origin: iPSC | Q9UMX1 | SUFU | 2 | Homo sapien | 0.61489924 | -1.2983045 | 2.52810301 | -0.8952713 | 0.68492827 | 0.50381656 | 0.99999785 | -6.1472463 | up |
| KI - WT, cell_origin: iPSC | O14936 | CASK | 10 | Homo sapien | 0.45127896 | -0.9532182 | 1.85577612 | -1.6887139 | 0.68474141 | 0.50393119 | 0.99999785 | -6.1473748 | up |
| KI - WT, cell_origin: iPSC | O75410 | TACC1 | 32 | Homo sapien | 0.4613373 | -0.9769651 | 1.89963966 | 1.81082207 | 0.68355072 | 0.50466194 | 0.99999785 | -6.1481933 | up |
| KI - WT, cell_origin: iPSC | Q9H4G4 | GLIPR2 | 2 | Homo sapien | 1.61595165 | -3.4260265 | 6.65792975 | -2.6615488 | 0.68301427 | 0.50499138 | 0.99999785 | -6.1485616 | up |
| KI - WT, cell_origin: iPSC | P14866 | HNRNPL | 15 | Homo sapien | -0.3208809 | -1.3222554 | 0.68049371 | 2.17734487 | -0.6828882 | 0.5050688 | 0.99999785 | -6.1486481 | down |
| KI - WT, cell_origin: iPSC | Q9UIA0 | CYTH4 | 2 | Homo sapien | -0.3402797 | -1.4021823 | 0.72162279 | -1.5998021 | -0.6826524 | 0.50517181 | 0.99999785 | -6.1488554 | down |
| KI - WT, cell_origin: iPSC | O60232 | ZNRD2 | 3 | Homo sapien | -0.2531067 | -1.043065 | 0.53685159 | -1.1489895 | -0.6824002 | 0.50529731 | 0.99999785 | -6.1490603 | down |
| KI - WT, cell_origin: iPSC | P45973 | CBX5 | 7 | Homo sapien | -0.5087569 | -2.0977871 | 1.08027339 | -0.2938747 | -0.682308 | 0.50542527 | 0.99999785 | -6.149046 | down |
| KI - WT, cell_origin: iPSC | O75531 | BANF1 | 8 | Homo sapien | -0.5323998 | -2.1959526 | 1.13115301 | 1.70481708 | -0.6820303 | 0.50559598 | 0.99999785 | -6.1492364 | down |
| KI - WT, cell_origin: iPSC | P13489 | RNH1 | 5 | Homo sapien | -0.3033973 | -1.2516492 | 0.64485466 | 0.02807733 | -0.6818523 | 0.5057054 | 0.99999785 | -6.1493584 | down |
| KI - WT, cell_origin: iPSC | O14531 | DPYSL4 | 9 | Homo sapien | -0.2205091 | -0.910522 | 0.46950369 | -0.6681034 | -0.6806269 | 0.50638789 | 0.99999785 | -6.1502746 | down |
| KI - WT, cell_origin: iPSC | Q9Y421 | FAM32A | 4 | Homo sapien | 0.44006183 | -0.938352 | 1.81847566 | 0.79527874 | 0.68035636 | 0.50662544 | 0.99999785 | -6.1503824 | up |
| KI - WT, cell_origin: iPSC | P49761 | CLK3 | 8 | Homo sapien | -0.2888621 | -1.1935927 | 0.61586844 | -0.3361287 | -0.6802638 | 0.50665637 | 0.99999785 | -6.150474 | down |
| KI - WT, cell_origin: iPSC | Q9GZM8 | NDEL1 | 6 | Homo sapien | 0.32787948 | -0.7003314 | 1.35609037 | -0.5311765 | 0.67957033 | 0.50710928 | 0.99999785 | -6.1509196 | up |
| KI - WT, cell_origin: iPSC | Q9NTK5 | OLA1 | 4 | Homo sapien | 0.28084664 | -0.5996632 | 1.16135644 | -1.6408065 | 0.67932049 | 0.50719218 | 0.99999785 | -6.1511672 | up |
| KI - WT, cell_origin: iPSC | Q9UIC8 | LCMT1 | 3 | Homo sapien | 0.27645931 | -0.5918214 | 1.14474004 | -1.3473793 | 0.67815964 | 0.5079132 | 0.99999785 | -6.1519529 | up |
| KI - WT, cell_origin: iPSC | Q8N684 | CPSF7 | 13 | Homo sapien | -0.4506657 | -1.8675217 | 0.96619028 | 0.9612392 | -0.6778462 | 0.50817147 | 0.99999785 | -6.1520958 | down |
| KI - WT, cell_origin: iPSC | Q13625 | TP53BP2 | 2 | Homo sapien | 0.26591561 | -0.5709353 | 1.10276655 | -0.3690664 | 0.67688302 | 0.50871584 | 0.99999785 | -6.1528056 | up |
| KI - WT, cell_origin: iPSC | Q7L4I2 | RSRC2 | 7 | Homo sapien | -0.3961916 | -1.644919 | 0.85253577 | 1.29326253 | -0.6761453 | 0.50922066 | 0.99999785 | -6.1532535 | down |
| KI - WT, cell_origin: iPSC | P51692 | STAT5B | 6 | Homo sapien | 0.35007336 | -0.7544747 | 1.45462146 | -0.7323425 | 0.67542449 | 0.50966563 | 0.99999785 | -6.1537433 | up |
| KI - WT, cell_origin: iPSC | Q5VT25 | CDC42BPA | 17 | Homo sapien | -0.5656914 | -2.3583639 | 1.22698112 | -0.8100647 | -0.6724825 | 0.5114842 | 0.99999785 | -6.1557372 | down |
| KI - WT, cell_origin: iPSC | Q9BZG1 | RAB34 | 10 | Homo sapien | 0.60195583 | -1.3061558 | 2.51006748 | 0.66732117 | 0.67230015 | 0.51159703 | 0.99999785 | -6.1558605 | up |
| KI - WT, cell_origin: iPSC | Q7Z3K3 | POGZ | 7 | Homo sapien | 0.27021115 | -0.587318 | 1.1277403 | -1.3080202 | 0.67113063 | 0.51225459 | 0.99999785 | -6.1567228 | up |
| KI - WT, cell_origin: iPSC | Q13322 | GRB10 | 4 | Homo sapien | -0.2360095 | -0.9854878 | 0.51346886 | -0.6049593 | -0.6706718 | 0.5125354 | 0.99999785 | -6.1570361 | down |
| KI - WT, cell_origin: iPSC | P55036 | PSMD4 | 4 | Homo sapien | 0.23369548 | -0.5084632 | 0.97585415 | -0.6259009 | 0.67064584 | 0.51255149 | 0.99999785 | -6.1570536 | up |
| KI - WT, cell_origin: iPSC | O95197 | RTN3 | 2 | Homo sapien | 0.42140195 | -0.9187225 | 1.76152635 | -1.3710788 | 0.67012187 | 0.51294608 | 0.99999785 | -6.1573312 | up |

| | | | | | | | | | | | | | |
|---|---|---|---|---|---|---|---|---|---|---|---|---|---|
| KI - WT, cell_origin: iPSC | O96007 | MOCS2 | 2 | Homo sapien | 0.56734878 | -1.2370626 | 2.37176014 | -2.5621068 | 0.67006502 | 0.51298132 | 0.99999785 | -6.1573696 | up |
| KI - WT, cell_origin: iPSC | Q96J02 | ITCH | 4 | Homo sapien | 0.30391413 | -0.6640028 | 1.27183109 | -2.0744976 | 0.66873256 | 0.51373791 | 0.99999785 | -6.1583423 | up |
| KI - WT, cell_origin: iPSC | O75563 | SKAP2 | 5 | Homo sapien | -0.1719908 | -0.7201906 | 0.37620901 | -0.0342463 | -0.6681993 | 0.51406886 | 0.99999785 | -6.1587009 | down |
| KI - WT, cell_origin: iPSC | Q05823 | RNASEL | 4 | Homo sapien | 0.45464123 | -0.9961566 | 1.90543901 | -3.0094339 | 0.66753725 | 0.51449947 | 0.99999785 | -6.1591245 | up |
| KI - WT, cell_origin: iPSC | Q6NZI2 | CAVIN1 | 5 | Homo sapien | 0.27109137 | -0.5942913 | 1.13647403 | -0.4358528 | 0.66730657 | 0.51464388 | 0.99999785 | -6.1592782 | up |
| KI - WT, cell_origin: iPSC | Q5VW36 | FOCAD | 27 | Homo sapien | 0.13251151 | -0.2905016 | 0.55552464 | 0.44341607 | 0.66717447 | 0.51470523 | 0.99999785 | -6.1593893 | up |
| KI - WT, cell_origin: iPSC | Q7Z6I6 | ARHGAP30 | 17 | Homo sapien | -0.2318009 | -0.9723153 | 0.50871349 | 1.02260784 | -0.6667626 | 0.51497432 | 0.99999785 | -6.1596514 | down |
| KI - WT, cell_origin: iPSC | O43583 | DENR | 2 | Homo sapien | -0.3793629 | -1.5950835 | 0.8363577 | -1.7187468 | -0.6650028 | 0.51612447 | 0.99999785 | -6.1607698 | down |
| KI - WT, cell_origin: iPSC | Q9BZH6 | WDR11 | 14 | Homo sapien | 0.19539632 | -0.4309113 | 0.82170396 | 0.60299529 | 0.66445926 | 0.51639343 | 0.99999785 | -6.1612082 | up |
| KI - WT, cell_origin: iPSC | Q13905 | RAPGEF1 | 3 | Homo sapien | 0.56778011 | -1.2534733 | 2.38903353 | -3.0816531 | 0.66437329 | 0.51651608 | 0.99999785 | -6.1611909 | up |
| KI - WT, cell_origin: iPSC | Q9BZ29 | DOCK9 | 48 | Homo sapien | -0.4547374 | -1.9136492 | 1.00417429 | 0.41205633 | -0.6642538 | 0.51659042 | 0.99999785 | -6.1612708 | down |
| KI - WT, cell_origin: iPSC | O43929 | ORC4 | 7 | Homo sapien | 0.24333203 | -0.5370587 | 1.02372275 | -1.9105231 | 0.66408989 | 0.51662334 | 0.99999785 | -6.1614551 | up |
| KI - WT, cell_origin: iPSC | P49354 | FNTA | 5 | Homo sapien | -0.2164297 | -0.9109534 | 0.47809399 | -1.9252853 | -0.6636964 | 0.5168683 | 0.99999785 | -6.1617179 | down |
| KI - WT, cell_origin: iPSC | Q13595 | TRA2A | 6 | Homo sapien | -0.3177821 | -1.338946 | 0.70338176 | 0.40802569 | -0.6631876 | 0.51725421 | 0.99999785 | -6.1619831 | down |
| KI - WT, cell_origin: iPSC | O60841 | EIF5B | 40 | Homo sapien | -0.5718059 | -2.409297 | 1.26568513 | 0.86104628 | -0.6631714 | 0.51726427 | 0.99999785 | -6.1619939 | down |
| KI - WT, cell_origin: iPSC | Q96CW5 | TUBGCP3 | 29 | Homo sapien | 0.15573981 | -0.3446275 | 0.65610708 | 0.66587255 | 0.66290369 | 0.51736204 | 0.99999785 | -6.1622471 | up |
| KI - WT, cell_origin: iPSC | Q6ZN17 | LIN28B | 10 | Homo sapien | -0.1955555 | -0.8239947 | 0.43288368 | 0.77844278 | -0.662745 | 0.5174609 | 0.99999785 | -6.162353 | down |
| KI - WT, cell_origin: iPSC | Q6P2C8 | MED27 | 4 | Homo sapien | 0.20674357 | -0.4597389 | 0.87322604 | -1.3776478 | 0.66066754 | 0.51875625 | 0.99999785 | -6.1637366 | up |
| KI - WT, cell_origin: iPSC | Q9BTV4 | TMEM43 | 2 | Homo sapien | -0.2773223 | -1.1725982 | 0.6179536 | -1.8940288 | -0.6597321 | 0.51934014 | 0.99999785 | -6.1643583 | down |
| KI - WT, cell_origin: iPSC | Q8N8A6 | DDX51 | 3 | Homo sapien | 0.30501135 | -0.68 | 1.29002274 | -3.0330016 | 0.65949945 | 0.51948538 | 0.99999785 | -6.1645127 | up |
| KI - WT, cell_origin: iPSC | Q9Y3D6 | FIS1 | 5 | Homo sapien | 0.49850217 | -1.1128313 | 2.10983562 | 1.17916919 | 0.65930139 | 0.5196776 | 0.99999785 | -6.1645702 | up |
| KI - WT, cell_origin: iPSC | Q96F24 | NRBF2 | 2 | Homo sapien | 0.30348173 | -0.6783667 | 1.28533015 | -0.7994644 | 0.65865594 | 0.52007242 | 0.99999785 | -6.1650075 | up |
| KI - WT, cell_origin: iPSC | Q86WB0 | ZC3HC1 | 5 | Homo sapien | 0.30615655 | -0.684815 | 1.29712812 | -2.4560955 | 0.65799416 | 0.52042586 | 0.99999785 | -6.165511 | up |
| KI - WT, cell_origin: iPSC | Q9P2R3 | ANKFY1 | 4 | Homo sapien | -0.2179062 | -0.9235053 | 0.48769303 | -0.4002132 | -0.6577352 | 0.52058772 | 0.99999785 | -6.1656825 | down |
| KI - WT, cell_origin: iPSC | Q01085 | TIAL1 | 7 | Homo sapien | -0.2212694 | -0.9393531 | 0.49681439 | 0.01908174 | -0.656275 | 0.52150113 | 0.99999785 | -6.1666485 | down |
| KI - WT, cell_origin: iPSC | P37198 | NUP62 | 4 | Homo sapien | -0.3157592 | -1.3410499 | 0.70953147 | -1.9647549 | -0.6559639 | 0.52170397 | 0.99999785 | -6.1668454 | down |
| KI - WT, cell_origin: iPSC | Q9Y5Y2 | NUBP2 | 6 | Homo sapien | 0.21075952 | -0.4735859 | 0.89510494 | 0.63703597 | 0.65592096 | 0.52172273 | 0.99999785 | -6.1668824 | up |
| KI - WT, cell_origin: iPSC | P39019 | RPS19 | 12 | Homo sapien | -0.1423008 | -0.6049368 | 0.32033516 | 6.79666616 | -0.6551002 | 0.52223667 | 0.99999785 | -6.1674243 | down |
| KI - WT, cell_origin: iPSC | Q8NEY8 | PPHLN1 | 6 | Homo sapien | -0.3232049 | -1.375769 | 0.72935917 | -1.4768219 | -0.6542 | 0.52283725 | 0.99999785 | -6.1679784 | down |
| KI - WT, cell_origin: iPSC | Q96B97 | SH3KBP1 | 3 | Homo sapien | 0.71357313 | -1.6125263 | 3.03967253 | -2.8776235 | 0.65375133 | 0.52314976 | 0.99999785 | -6.1682401 | up |
| KI - WT, cell_origin: iPSC | P35789 | ZNF93 | 3 | Homo sapien | -0.2632913 | -1.1225132 | 0.59593055 | -1.0191216 | -0.6527367 | 0.52373552 | 0.99999785 | -6.1689622 | down |
| KI - WT, cell_origin: iPSC | Q13442 | PDAP1 | 9 | Homo sapien | -0.7191044 | -3.0689072 | 1.63069842 | 1.72469238 | -0.6521731 | 0.5241395 | 0.99999785 | -6.1692784 | down |
| KI - WT, cell_origin: iPSC | P49023 | PXN | 9 | Homo sapien | -0.2115325 | -0.9028759 | 0.47981088 | 2.23398737 | -0.6516628 | 0.5243921 | 0.99999785 | -6.1696864 | down |
| KI - WT, cell_origin: iPSC | P84095 | RHOG | 13 | Homo sapien | 0.1964612 | -0.4470547 | 0.83997713 | 6.1914477 | 0.65056096 | 0.52514342 | 0.99999785 | -6.1703453 | up |
| KI - WT, cell_origin: iPSC | O14656 | TOR1A | 3 | Homo sapien | 0.65520742 | -1.4942909 | 2.80470577 | -3.1788985 | 0.64959708 | 0.52575725 | 0.99999785 | -6.170968 | up |
| KI - WT, cell_origin: iPSC | Q7Z739 | YTHDF3 | 15 | Homo sapien | -0.1768498 | -0.7567434 | 0.40304376 | 1.5555043 | -0.6495252 | 0.5257351 | 0.99999785 | -6.1710875 | down |
| KI - WT, cell_origin: iPSC | P63241 | EIF5A | 4 | Homo sapien | -0.325835 | -1.394981 | 0.74331099 | 0.95295661 | -0.649476 | 0.52583335 | 0.99999785 | -6.1710472 | down |
| KI - WT, cell_origin: iPSC | Q04917 | YWHAH | 8 | Homo sapien | 0.1885603 | -0.4303405 | 0.8074611 | 2.07330177 | 0.64888672 | 0.52613657 | 0.99999785 | -6.1715052 | up |
| KI - WT, cell_origin: iPSC | Q9NQT5 | EXOSC3 | 4 | Homo sapien | 0.21212943 | -0.485406 | 0.90966482 | -0.6980854 | 0.64770071 | 0.52688284 | 0.99999785 | -6.1722799 | up |
| KI - WT, cell_origin: iPSC | O15371 | EIF3D | 16 | Homo sapien | -0.1812623 | -0.7777492 | 0.41522456 | 2.61409461 | -0.6472116 | 0.52719075 | 0.99999785 | -6.172599 | down |
| KI - WT, cell_origin: iPSC | P60903 | S100A10 | 5 | Homo sapien | 0.311173 | -0.7138965 | 1.33624249 | 2.32265898 | 0.64692058 | 0.52744108 | 0.99999785 | -6.1727168 | up |
| KI - WT, cell_origin: iPSC | Q9UBQ0 | VPS29 | 5 | Homo sapien | 0.37075497 | -0.8508366 | 1.59234655 | -0.2164583 | 0.64679004 | 0.52752328 | 0.99999785 | -6.1728019 | up |
| KI - WT, cell_origin: iPSC | Q9NYU2 | UGGT1 | 21 | Homo sapien | 0.60602902 | -1.3911471 | 2.60320509 | -1.2591514 | 0.64666511 | 0.52760196 | 0.99999785 | -6.1728833 | up |
| KI - WT, cell_origin: iPSC | Q9C0F1 | CEP44 | 2 | Homo sapien | 0.36077735 | -0.8286534 | 1.55020812 | -2.0732763 | 0.64612716 | 0.52789381 | 0.99999785 | -6.1732844 | up |
| KI - WT, cell_origin: iPSC | Q2TAZ0 | ATG2A | 4 | Homo sapien | 0.31328228 | -0.7210146 | 1.34757919 | -1.9642432 | 0.64517019 | 0.52848829 | 0.99999785 | -6.1739165 | up |
| KI - WT, cell_origin: iPSC | Q9P2R6 | RERE | 3 | Homo sapien | 0.37634164 | -0.8673287 | 1.62001197 | -2.4933943 | 0.64459035 | 0.52885983 | 0.99999785 | -6.1742871 | up |

| | | | | | | | | | | | | |
|---|---|---|---|---|---|---|---|---|---|---|---|---|
| KI - WT, cell_origin: iPSC | O75874 | IDH1 | 8 | Homo sapien | 0.40481065 | -0.9353684 | 1.74498972 | -0.742924 | 0.64371178 | 0.52946378 | 0.99999785 | -6.1748044 up |
| KI - WT, cell_origin: iPSC | Q9BXW9 | FANCD2 | 4 | Homo sapien | 0.39415447 | -0.9104396 | 1.69874852 | -3.3383827 | 0.64347361 | 0.52954745 | 0.99999785 | -6.1750304 up |
| KI - WT, cell_origin: iPSC | Q9P2K8 | EIF2AK4 | 10 | Homo sapien | 0.41667057 | -0.9637248 | 1.79706598 | -1.776265 | 0.64326763 | 0.5297441 | 0.99999785 | -6.1750925 up |
| KI - WT, cell_origin: iPSC | Q7RTV0 | PHF5A | 7 | Homo sapien | -0.3206715 | -1.3837207 | 0.74237778 | 0.93367721 | -0.6428495 | 0.53000809 | 0.99999785 | -6.1753637 down |
| KI - WT, cell_origin: iPSC | P46734 | MAP2K3 | 4 | Homo sapien | -0.2672091 | -1.1530734 | 0.61865523 | 0.69918519 | -0.6427218 | 0.53007265 | 0.99999785 | -6.1754637 down |
| KI - WT, cell_origin: iPSC | Q8TEQ0 | SNX29 | 3 | Homo sapien | 0.40534583 | -0.9416053 | 1.75229694 | -3.6633521 | 0.64093437 | 0.53115173 | 0.99999785 | -6.1766745 up |
| KI - WT, cell_origin: iPSC | P98175 | RBM10 | 23 | Homo sapien | -0.2327705 | -1.0077426 | 0.54220171 | 0.69896392 | -0.6398064 | 0.53188217 | 0.99999785 | -6.1773847 down |
| KI - WT, cell_origin: iPSC | Q96LT7 | C9orf72 | 10 | Homo sapien | 0.4116822 | -0.960213 | 1.78357743 | 0.02822142 | 0.63950437 | 0.53212254 | 0.99999785 | -6.1775268 up |
| KI - WT, cell_origin: iPSC | Q9NX05 | FAM120C | 2 | Homo sapien | -0.6508539 | -2.8201541 | 1.51844636 | -2.338951 | -0.6393905 | 0.53219458 | 0.99999785 | -6.1776002 down |
| KI - WT, cell_origin: iPSC | Q96PN7 | TRERF1 | 2 | Homo sapien | -0.4941947 | -2.142 | 1.15361068 | -2.8727919 | -0.6391378 | 0.53235451 | 0.99999785 | -6.1777631 down |
| KI - WT, cell_origin: iPSC | O60244 | MED14 | 7 | Homo sapien | 0.1800711 | -0.4200035 | 0.78014574 | -1.3418413 | 0.63911414 | 0.5323034 | 0.99999785 | -6.1778493 up |
| KI - WT, cell_origin: iPSC | Q9Y606 | PUS1 | 4 | Homo sapien | 0.30055474 | -0.702713 | 1.30382245 | -1.4534099 | 0.63815276 | 0.53293189 | 0.99999785 | -6.1784474 up |
| KI - WT, cell_origin: iPSC | Q9BZ67 | FRMD8 | 2 | Homo sapien | -0.3749069 | -1.6294628 | 0.87964907 | -0.8580235 | -0.6368479 | 0.53380505 | 0.99999785 | -6.1792369 down |
| KI - WT, cell_origin: iPSC | P53634 | CTSC | 7 | Homo sapien | 0.51789184 | -1.2153743 | 2.25115793 | -1.8442556 | 0.63676061 | 0.53386039 | 0.99999785 | -6.179293 up |
| KI - WT, cell_origin: iPSC | Q8ND56 | LSM14A | 9 | Homo sapien | -0.1446384 | -0.6286445 | 0.33936769 | 2.88757954 | -0.636462 | 0.53398395 | 0.99999785 | -6.1795553 down |
| KI - WT, cell_origin: iPSC | Q9ULW0 | TPX2 | 16 | Homo sapien | -0.2685864 | -1.168103 | 0.6309303 | 0.79811773 | -0.6362756 | 0.53415996 | 0.99999785 | -6.1796131 down |
| KI - WT, cell_origin: iPSC | P49591 | SARS1 | 3 | Homo sapien | -0.4116094 | -1.7903394 | 0.96712072 | -2.8783413 | -0.6359351 | 0.53433495 | 0.99999785 | -6.1798754 down |
| KI - WT, cell_origin: iPSC | P46019 | PHKA2 | 24 | Homo sapien | 0.16908875 | -0.397217 | 0.73539446 | 0.87624834 | 0.6359214 | 0.53432684 | 0.99999785 | -6.1799022 up |
| KI - WT, cell_origin: iPSC | Q6XZF7 | DNMBP | 28 | Homo sapien | 0.31132426 | -0.7328103 | 1.35545879 | -0.4891527 | 0.63541707 | 0.53471252 | 0.99999785 | -6.1801553 up |
| KI - WT, cell_origin: iPSC | Q9H8H0 | NOL11 | 6 | Homo sapien | 0.40935655 | -0.9629919 | 1.78170503 | -3.750849 | 0.63529731 | 0.53472286 | 0.99999785 | -6.1803023 up |
| KI - WT, cell_origin: iPSC | O14795 | UNC13B | 12 | Homo sapien | -0.2268515 | -0.9878767 | 0.53417372 | -0.690751 | -0.634875 | 0.53499244 | 0.99999785 | -6.1805713 down |
| KI - WT, cell_origin: iPSC | Q9UEY8 | ADD3 | 12 | Homo sapien | 0.46454546 | -1.0951438 | 2.02423477 | -0.2993448 | 0.63473507 | 0.53514536 | 0.99999785 | -6.1805923 up |
| KI - WT, cell_origin: iPSC | O95199 | RCBTB2 | 8 | Homo sapien | 0.14951495 | -0.3521876 | 0.6512175 | 0.14526595 | 0.63471386 | 0.53509324 | 0.99999785 | -6.1806761 up |
| KI - WT, cell_origin: iPSC | Q9BR61 | ACBD6 | 3 | Homo sapien | 0.30633699 | -0.722854 | 1.33552803 | -2.5815788 | 0.63393267 | 0.53558938 | 0.99999785 | -6.181176 up |
| KI - WT, cell_origin: iPSC | P84098 | RPL19 | 6 | Homo sapien | -0.1912299 | -0.8341841 | 0.45172425 | 5.1091991 | -0.6335688 | 0.53584008 | 0.99999785 | -6.1813878 down |
| KI - WT, cell_origin: iPSC | Q9H0A0 | NAT10 | 24 | Homo sapien | -0.4614342 | -2.0143891 | 1.09152077 | -0.7395413 | -0.633218 | 0.53610888 | 0.99999785 | -6.1815629 down |
| KI - WT, cell_origin: iPSC | P02794 | FTH1 | 3 | Homo sapien | 0.31578109 | -0.7474866 | 1.37904877 | -0.5851454 | 0.63291571 | 0.536301 | 0.99999785 | -6.1817561 up |
| KI - WT, cell_origin: iPSC | Q96B26 | EXOSC8 | 7 | Homo sapien | -0.205364 | -0.8968645 | 0.48613645 | -0.4908215 | -0.632516 | 0.53648973 | 0.99999785 | -6.1820811 down |
| KI - WT, cell_origin: iPSC | Q8NE71 | ABCF1 | 23 | Homo sapien | -0.3368713 | -1.4731601 | 0.79941744 | 1.8253511 | -0.6317972 | 0.53701214 | 0.99999785 | -6.1824699 down |
| KI - WT, cell_origin: iPSC | Q9P0L0 | VAPA | 5 | Homo sapien | 0.41408141 | -0.9828852 | 1.81104798 | -0.6533376 | 0.63168722 | 0.53708211 | 0.99999785 | -6.18254 up |
| KI - WT, cell_origin: iPSC | Q52LJ0 | FAM98B | 4 | Homo sapien | 0.22269277 | -0.5294782 | 0.9748637 | 0.9657048 | 0.63064697 | 0.53769306 | 0.99999785 | -6.1832572 up |
| KI - WT, cell_origin: iPSC | Q6NZY4 | ZCCHC8 | 5 | Homo sapien | -0.1552112 | -0.680986 | 0.37056356 | 0.80032482 | -0.6287284 | 0.53890112 | 0.99999785 | -6.1844916 down |
| KI - WT, cell_origin: iPSC | Q8TF01 | PNISR | 8 | Homo sapien | -0.5060144 | -2.221227 | 1.20919826 | 0.39336675 | -0.6287055 | 0.53898059 | 0.99999785 | -6.1844369 down |
| KI - WT, cell_origin: iPSC | P35659 | DEK | 10 | Homo sapien | -0.5261626 | -2.3102489 | 1.2579238 | 1.70716265 | -0.6285017 | 0.53911049 | 0.99999785 | -6.1845662 down |
| KI - WT, cell_origin: iPSC | O15397 | IPO8 | 5 | Homo sapien | 0.16225868 | -0.3878422 | 0.7123596 | -1.4472347 | 0.62821058 | 0.53923125 | 0.99999785 | -6.18482 up |
| KI - WT, cell_origin: iPSC | Q5T200 | ZC3H13 | 10 | Homo sapien | -0.2760857 | -1.2131529 | 0.66098141 | 0.20273821 | -0.6278427 | 0.53952452 | 0.99999785 | -6.1849907 down |
| KI - WT, cell_origin: iPSC | Q9UIV1 | CNOT7 | 8 | Homo sapien | 0.15446375 | -0.3696323 | 0.67855978 | 1.01460548 | 0.62770473 | 0.53955386 | 0.99999785 | -6.1851407 up |
| KI - WT, cell_origin: iPSC | Q96T37 | RBM15 | 14 | Homo sapien | -0.3521046 | -1.5485705 | 0.84436139 | -1.4105028 | -0.6271532 | 0.53997038 | 0.99999785 | -6.185421 down |
| KI - WT, cell_origin: iPSC | Q9Y2M0 | FAN1 | 7 | Homo sapien | -0.4199607 | -1.8466824 | 1.00676108 | -3.6310521 | -0.6269155 | 0.5400574 | 0.99999785 | -6.1856404 down |
| KI - WT, cell_origin: iPSC | Q15269 | PWP2 | 6 | Homo sapien | 0.14053759 | -0.3371316 | 0.61820674 | -0.3506009 | 0.62662104 | 0.54024534 | 0.99999785 | -6.1858267 up |
| KI - WT, cell_origin: iPSC | Q9Y262 | EIF3L | 24 | Homo sapien | -0.1576594 | -0.6936264 | 0.37830765 | 2.07499814 | -0.6265005 | 0.5403223 | 0.99999785 | -6.185903 down |
| KI - WT, cell_origin: iPSC | Q13107 | USP4 | 9 | Homo sapien | 0.25983392 | -0.6239923 | 1.14366014 | 0.75661266 | 0.62642338 | 0.54042054 | 0.99999785 | -6.1858995 up |
| KI - WT, cell_origin: iPSC | P04259 | KRT6B | 3 | Homo sapien | -0.6543752 | -2.8805632 | 1.57181279 | 2.54089812 | -0.6264225 | 0.54043666 | 0.99999785 | -6.1858835 down |
| KI - WT, cell_origin: iPSC | Q13330 | MTA1 | 5 | Homo sapien | 0.29643633 | -0.7119274 | 1.30480005 | -2.0123338 | 0.62612331 | 0.54056457 | 0.99999785 | -6.1861399 up |
| KI - WT, cell_origin: iPSC | Q9Y6R4 | MAP3K4 | 7 | Homo sapien | 0.41540894 | -0.999062 | 1.82987992 | -2.3679693 | 0.62582413 | 0.54081085 | 0.99999785 | -6.1862702 up |
| KI - WT, cell_origin: iPSC | Q9BPZ7 | MAPKAP1 | 5 | Homo sapien | 0.41158414 | -0.9910945 | 1.81426276 | -1.189326 | 0.62532073 | 0.54114016 | 0.99999785 | -6.1865798 up |

| | | | | | | | | | | | | |
|---|---|---|---|---|---|---|---|---|---|---|---|---|
| KI - WT, cell_origin: iPSC | O75822 | EIF3J | 7 | Homo sapien | 0.20381664 | -0.4908276 | 0.89846085 | 0.42730533 | 0.62490919 | 0.54133865 | 0.99999785 | -6.1869082 up |
| KI - WT, cell_origin: iPSC | Q96AB3 | ISOC2 | 2 | Homo sapien | 0.34923313 | -0.8413262 | 1.53979242 | -2.2708157 | 0.62484343 | 0.54139713 | 0.99999785 | -6.1869322 up |
| KI - WT, cell_origin: iPSC | O43679 | LDB2 | 3 | Homo sapien | -0.412174 | -1.8179392 | 0.99359113 | -0.4211125 | -0.624842 | 0.54144598 | 0.99999785 | -6.186882 down |
| KI - WT, cell_origin: iPSC | P51659 | HSD17B4 | 39 | Homo sapien | -0.2896336 | -1.27967 | 0.70040273 | 3.97020163 | -0.623448 | 0.54233707 | 0.99999785 | -6.1877608 down |
| KI - WT, cell_origin: iPSC | Q9Y5L0 | TNPO3 | 14 | Homo sapien | -0.3090473 | -1.3655864 | 0.74749174 | 0.72899377 | -0.6233641 | 0.54239069 | 0.99999785 | -6.1878136 down |
| KI - WT, cell_origin: iPSC | Q9NRW4 | DUSP22 | 6 | Homo sapien | 0.30727315 | -0.7434526 | 1.35799891 | 0.27336314 | 0.62321456 | 0.54248636 | 0.99999785 | -6.1879077 up |
| KI - WT, cell_origin: iPSC | Q9H0V9 | LMAN2L | 9 | Homo sapien | 0.7533006 | -1.8307571 | 3.33735829 | -0.6583926 | 0.62125297 | 0.54374181 | 0.99999785 | -6.1891407 up |
| KI - WT, cell_origin: iPSC | Q9H1I8 | ASCC2 | 15 | Homo sapien | 0.1582304 | -0.3850735 | 0.70153433 | 1.20896061 | 0.62027855 | 0.54430217 | 0.99999785 | -6.1898196 up |
| KI - WT, cell_origin: iPSC | Q92930 | RAB8B | 6 | Homo sapien | 0.24009566 | -0.5853651 | 1.06555643 | 3.04306547 | 0.61975487 | 0.54468461 | 0.99999785 | -6.1900979 up |
| KI - WT, cell_origin: iPSC | O75821 | EIF3G | 19 | Homo sapien | -0.1308705 | -0.5815472 | 0.31980622 | 2.9576489 | -0.6184667 | 0.54546416 | 0.99999785 | -6.1909532 down |
| KI - WT, cell_origin: iPSC | Q13490 | BIRC2 | 14 | Homo sapien | 0.13957901 | -0.3419462 | 0.62110426 | -0.03116 | 0.61736321 | 0.54617249 | 0.99999785 | -6.191642 up |
| KI - WT, cell_origin: iPSC | P07437 | TUBB | 4 | Homo sapien | -0.1551371 | -0.6904436 | 0.38016952 | 7.24999384 | -0.6173492 | 0.54620042 | 0.99999785 | -6.1916307 down |
| KI - WT, cell_origin: iPSC | Q6ICG6 | KIAA0930 | 8 | Homo sapien | 0.27049941 | -0.6636645 | 1.20466334 | -1.2963809 | 0.61681766 | 0.54654073 | 0.99999785 | -6.1919632 up |
| KI - WT, cell_origin: iPSC | Q12979 | ABR | 6 | Homo sapien | -0.3545168 | -1.5811282 | 0.87209466 | -1.4681049 | -0.6159311 | 0.54715588 | 0.99999785 | -6.1924671 down |
| KI - WT, cell_origin: iPSC | Q9Y4F1 | FARP1 | 10 | Homo sapien | 0.33774966 | -0.8312595 | 1.50675878 | -0.8871323 | 0.61571452 | 0.54729507 | 0.99999785 | -6.1926019 up |
| KI - WT, cell_origin: iPSC | Q9H910 | JPT2 | 4 | Homo sapien | -0.2315488 | -1.0326533 | 0.56955563 | 0.32586986 | -0.6157031 | 0.54725797 | 0.99999785 | -6.1926562 down |
| KI - WT, cell_origin: iPSC | Q9Y2X7 | GIT1 | 35 | Homo sapien | 0.45468056 | -1.1210889 | 2.03045003 | 1.90546704 | 0.61491639 | 0.54780816 | 0.99999785 | -6.1930983 up |
| KI - WT, cell_origin: iPSC | Q01804 | OTUD4 | 13 | Homo sapien | -0.1996832 | -0.89273 | 0.49336374 | 0.4815139 | -0.6136468 | 0.54856179 | 0.99999785 | -6.1939534 down |
| KI - WT, cell_origin: iPSC | Q8TEQ6 | GEMIN5 | 37 | Homo sapien | -0.256476 | -1.1473544 | 0.63440229 | 0.90399818 | -0.6134461 | 0.548741 | 0.99999785 | -6.1940249 down |
| KI - WT, cell_origin: iPSC | Q13144 | EIF2B5 | 3 | Homo sapien | 0.22578152 | -0.5581214 | 1.00968444 | -1.1496884 | 0.61343111 | 0.54870065 | 0.99999785 | -6.1940871 up |
| KI - WT, cell_origin: iPSC | Q8ND04 | SMG8 | 8 | Homo sapien | -0.1739481 | -0.7797899 | 0.4318937 | -0.5699857 | -0.6115051 | 0.54994129 | 0.99999785 | -6.1952793 down |
| KI - WT, cell_origin: iPSC | Q86YZ3 | HRNR | 9 | Homo sapien | -0.5089703 | -2.2851243 | 1.26718374 | -0.1446435 | -0.6106807 | 0.55053555 | 0.99999785 | -6.1957223 down |
| KI - WT, cell_origin: iPSC | P62979 | RPS27A | 7 | Homo sapien | 0.14802228 | -0.3684291 | 0.66447364 | 3.93423945 | 0.61043206 | 0.55063317 | 0.99999785 | -6.195942 up |
| KI - WT, cell_origin: iPSC | Q5JR12 | PPM1J | 6 | Homo sapien | 0.28747317 | -0.7186456 | 1.29359198 | 0.78631854 | 0.60890623 | 0.5516803 | 0.99999785 | -6.1968164 up |
| KI - WT, cell_origin: iPSC | Q9Y547 | HSPB11 | 2 | Homo sapien | 0.17022616 | -0.4263534 | 0.76680569 | -0.8361537 | 0.60771175 | 0.55238926 | 0.99999785 | -6.197617 up |
| KI - WT, cell_origin: iPSC | P51617 | IRAK1 | 8 | Homo sapien | 0.22967115 | -0.5753569 | 1.0346992 | -0.9659082 | 0.60766659 | 0.55242562 | 0.99999785 | -6.1976372 up |
| KI - WT, cell_origin: iPSC | O60282 | KIF5C | 15 | Homo sapien | 0.21530819 | -0.5393886 | 0.97000496 | -0.4609935 | 0.60763752 | 0.55244122 | 0.99999785 | -6.1976584 up |
| KI - WT, cell_origin: iPSC | Q96GA3 | LTV1 | 8 | Homo sapien | 0.2259047 | -0.5660667 | 1.01787608 | 0.57814071 | 0.60762227 | 0.55246565 | 0.99999785 | -6.1976524 up |
| KI - WT, cell_origin: iPSC | Q9UKV3 | ACIN1 | 17 | Homo sapien | -0.3345828 | -1.5087361 | 0.83957053 | -0.6543579 | -0.607269 | 0.55273765 | 0.99999785 | -6.1978232 down |
| KI - WT, cell_origin: iPSC | Q6DN90 | IQSEC1 | 19 | Homo sapien | -0.289151 | -1.3039063 | 0.72560438 | -0.394046 | -0.6072474 | 0.55275162 | 0.99999785 | -6.1978365 down |
| KI - WT, cell_origin: iPSC | Q96EY4 | TMA16 | 8 | Homo sapien | -0.2739567 | -1.2365047 | 0.68859134 | 0.1151534 | -0.6065434 | 0.55320665 | 0.99999785 | -6.1982686 down |
| KI - WT, cell_origin: iPSC | P39687 | ANP32A | 5 | Homo sapien | -0.3244613 | -1.4646237 | 0.81570102 | -1.3494334 | -0.606455 | 0.55326377 | 0.99999785 | -6.1983228 down |
| KI - WT, cell_origin: iPSC | O75190 | DNAJB6 | 9 | Homo sapien | 0.18427004 | -0.4639277 | 0.83246775 | 0.79602194 | 0.60546204 | 0.55384384 | 0.99999785 | -6.1989969 up |
| KI - WT, cell_origin: iPSC | Q16875 | PFKFB3 | 2 | Homo sapien | -0.3029113 | -1.3689432 | 0.7631207 | -1.2980913 | -0.6054582 | 0.55389343 | 0.99999785 | -6.1989496 down |
| KI - WT, cell_origin: iPSC | P11274 | BCR | 15 | Homo sapien | 0.25928737 | -0.6547877 | 1.17336241 | 0.3155294 | 0.60443211 | 0.5545596 | 0.99999785 | -6.1995752 up |
| KI - WT, cell_origin: iPSC | Q9Y617 | PSAT1 | 7 | Homo sapien | -0.4457275 | -2.0178695 | 1.12641462 | -1.7620738 | -0.604199 | 0.55472337 | 0.99999785 | -6.1997042 down |
| KI - WT, cell_origin: iPSC | Q13895 | BYSL | 17 | Homo sapien | 0.15648745 | -0.3953895 | 0.70836445 | 0.83190925 | 0.60391652 | 0.55484431 | 0.99999785 | -6.199942 up |
| KI - WT, cell_origin: iPSC | Q9UJQ4 | SALL4 | 9 | Homo sapien | -0.2608124 | -1.1813346 | 0.65970991 | -0.5892561 | -0.6036742 | 0.55504112 | 0.99999785 | -6.2000481 down |
| KI - WT, cell_origin: iPSC | P05386 | RPLP1 | 3 | Homo sapien | -0.1410179 | -0.6386361 | 0.35660032 | 5.34385819 | -0.6035561 | 0.55507779 | 0.99999785 | -6.2001621 down |
| KI - WT, cell_origin: iPSC | Q5T3I0 | GPATCH4 | 8 | Homo sapien | -0.4833516 | -2.1908489 | 1.22414576 | -1.7072152 | -0.6032613 | 0.5553306 | 0.99999785 | -6.2002769 down |
| KI - WT, cell_origin: iPSC | P49770 | EIF2B2 | 5 | Homo sapien | -0.1825102 | -0.8279592 | 0.46293882 | -0.4285931 | -0.6022335 | 0.55593494 | 0.99999785 | -6.2009687 down |
| KI - WT, cell_origin: iPSC | Q15642 | TRIP10 | 17 | Homo sapien | 0.39856059 | -1.0134403 | 1.81056152 | 0.04205842 | 0.60153615 | 0.55644876 | 0.99999785 | -6.2013284 up |
| KI - WT, cell_origin: iPSC | Q9UL40 | ZNF346 | 3 | Homo sapien | 0.28240505 | -0.7177574 | 1.28256748 | -2.9988476 | 0.60136979 | 0.55649504 | 0.99999785 | -6.2014944 up |
| KI - WT, cell_origin: iPSC | Q13287 | NMI | 9 | Homo sapien | -0.2874971 | -1.3069038 | 0.73190968 | -1.447654 | -0.6007639 | 0.5569065 | 0.99999785 | -6.2018436 down |
| KI - WT, cell_origin: iPSC | Q13200 | PSMD2 | 12 | Homo sapien | 0.29913337 | -0.7627769 | 1.36104363 | -0.8629085 | 0.60031532 | 0.55724078 | 0.99999785 | -6.2020707 up |
| KI - WT, cell_origin: iPSC | P27635 | RPL10 | 11 | Homo sapien | -0.1406381 | -0.6397356 | 0.35845933 | 3.2526117 | -0.6001467 | 0.55728876 | 0.99999785 | -6.2022378 down |

| | | | | | | | | | | | | | |
|---|---|---|---|---|---|---|---|---|---|---|---|---|---|
| KI - WT, cell_origin: iPSC | Q14258 | TRIM25 | 21 | Homo sapien | -0.131395 | -0.5977997 | 0.3350096 | 1.66073578 | -0.6000063 | 0.55737991 | 0.99999785 | -6.202323 | down |
| KI - WT, cell_origin: iPSC | A6NDG6 | PGP | 3 | Homo sapien | -0.3323235 | -1.512785 | 0.84813802 | -1.3017657 | -0.5999452 | 0.557481 | 0.99999785 | -6.2022955 | down |
| KI - WT, cell_origin: iPSC | Q6NXG1 | ESRP1 | 10 | Homo sapien | -0.1342842 | -0.6115136 | 0.34294529 | 1.2251031 | -0.5992903 | 0.55784488 | 0.99999785 | -6.2027574 | down |
| KI - WT, cell_origin: iPSC | Q9BRJ7 | NUDT16L1 | 12 | Homo sapien | -0.2483769 | -1.132342 | 0.63558814 | 1.73213958 | -0.5986986 | 0.55827418 | 0.99999785 | -6.2030689 | down |
| KI - WT, cell_origin: iPSC | P23528 | CFL1 | 17 | Homo sapien | -0.1638267 | -0.7471489 | 0.41949558 | 3.8476903 | -0.5981577 | 0.55858076 | 0.99999785 | -6.2034435 | down |
| KI - WT, cell_origin: iPSC | Q92608 | DOCK2 | 79 | Homo sapien | -0.4721133 | -2.1560326 | 1.21180599 | 0.1839641 | -0.5974854 | 0.55907899 | 0.99999785 | -6.203786 | down |
| KI - WT, cell_origin: iPSC | Q9H0C8 | ILKAP | 10 | Homo sapien | -0.2865966 | -1.3091806 | 0.7359874 | 2.78455164 | -0.5972751 | 0.55921576 | 0.99999785 | -6.2039132 | down |
| KI - WT, cell_origin: iPSC | Q92747 | ARPC1A | 3 | Homo sapien | -0.2482838 | -1.1341413 | 0.63757375 | -1.0943302 | -0.5970233 | 0.55933401 | 0.99999785 | -6.2041131 | down |
| KI - WT, cell_origin: iPSC | O75940 | SMNDC1 | 6 | Homo sapien | -0.4114957 | -1.8817859 | 1.05879443 | -0.6333972 | -0.5964371 | 0.55976079 | 0.99999785 | -6.2044195 | down |
| KI - WT, cell_origin: iPSC | Q8WWM7 | ATXN2L | 26 | Homo sapien | -0.2933863 | -1.3426885 | 0.75591583 | 2.54664851 | -0.5958565 | 0.56013859 | 0.99999785 | -6.2047699 | down |
| KI - WT, cell_origin: iPSC | Q8IWZ3 | ANKHD1 | 27 | Homo sapien | -0.1909167 | -0.8737684 | 0.491935 | -0.043239 | -0.5954663 | 0.56033164 | 0.99999785 | -6.205069 | down |
| KI - WT, cell_origin: iPSC | Q4KMP7 | TBC1D10B | 8 | Homo sapien | -0.7149745 | -3.2764725 | 1.84652349 | -2.3520016 | -0.5948383 | 0.56080144 | 0.99999785 | -6.2053835 | down |
| KI - WT, cell_origin: iPSC | P02751 | FN1 | 44 | Homo sapien | 0.66467351 | -1.7231105 | 3.05245757 | -0.4568192 | 0.59321987 | 0.5618559 | 0.99999785 | -6.2063569 | up |
| KI - WT, cell_origin: iPSC | P09874 | PARP1 | 50 | Homo sapien | -0.3643724 | -1.674522 | 0.94577729 | 3.04735036 | -0.592689 | 0.562202 | 0.99999785 | -6.2066756 | down |
| KI - WT, cell_origin: iPSC | P04075 | ALDOA | 14 | Homo sapien | -0.3278008 | -1.5073325 | 0.8517309 | 0.32836108 | -0.5922469 | 0.56249035 | 0.99999785 | -6.2069408 | down |
| KI - WT, cell_origin: iPSC | Q9Y5L4 | TIMM13 | 2 | Homo sapien | -0.2264729 | -1.0413648 | 0.58841911 | -0.2989045 | -0.5920017 | 0.56260527 | 0.99999785 | -6.207135 | down |
| KI - WT, cell_origin: iPSC | P49721 | PSMB2 | 4 | Homo sapien | 0.22565827 | -0.5865441 | 1.03786068 | -0.7767122 | 0.59181048 | 0.56272745 | 0.99999785 | -6.2072522 | up |
| KI - WT, cell_origin: iPSC | Q86U06 | RBM23 | 2 | Homo sapien | -0.5403271 | -2.4860513 | 1.40539703 | -2.7677963 | -0.591804 | 0.5627793 | 0.99999785 | -6.2072064 | down |
| KI - WT, cell_origin: iPSC | P52789 | HK2 | 23 | Homo sapien | 0.24724947 | -0.643998 | 1.13849696 | -0.4622607 | 0.59104176 | 0.56324882 | 0.99999785 | -6.207692 | up |
| KI - WT, cell_origin: iPSC | P23469 | PTPRE | 18 | Homo sapien | 0.58961202 | -1.5363407 | 2.71556477 | -0.6596747 | 0.59103757 | 0.56327943 | 0.99999785 | -6.2076653 | up |
| KI - WT, cell_origin: iPSC | P17096 | HMGA1 | 5 | Homo sapien | -0.6698885 | -3.0873515 | 1.74757439 | 3.59032409 | -0.5905343 | 0.56360802 | 0.99999785 | -6.2079665 | down |
| KI - WT, cell_origin: iPSC | Q8WXA9 | SREK1 | 6 | Homo sapien | -0.5426896 | -2.5035386 | 1.41815941 | 0.86107484 | -0.5898067 | 0.56408318 | 0.99999785 | -6.2084013 | down |
| KI - WT, cell_origin: iPSC | O75179 | ANKRD17 | 15 | Homo sapien | -0.2145628 | -0.9911637 | 0.56203814 | -0.2485141 | -0.5884915 | 0.56489266 | 0.99999785 | -6.2092383 | down |
| KI - WT, cell_origin: iPSC | Q9P0V3 | SH3BP4 | 3 | Homo sapien | -0.4393373 | -2.0317436 | 1.15306889 | -1.3562137 | -0.5879584 | 0.56529124 | 0.99999785 | -6.2095038 | down |
| KI - WT, cell_origin: iPSC | Q9H9Y6 | POLR1B | 6 | Homo sapien | 0.33572892 | -0.8816713 | 1.55312912 | -2.6313431 | 0.58738833 | 0.56561108 | 0.99999785 | -6.2098985 | up |
| KI - WT, cell_origin: iPSC | P35612 | ADD2 | 4 | Homo sapien | 0.22416314 | -0.5890084 | 1.03733473 | -2.026052 | 0.58711299 | 0.56578435 | 0.99999785 | -6.2100694 | up |
| KI - WT, cell_origin: iPSC | P49959 | MRE11 | 9 | Homo sapien | 0.16326712 | -0.4290615 | 0.75559569 | -0.2557126 | 0.58705085 | 0.56582502 | 0.99999785 | -6.2101064 | up |
| KI - WT, cell_origin: iPSC | Q96RS6 | NUDCD1 | 9 | Homo sapien | -0.2423197 | -1.1225645 | 0.63792501 | -0.9931028 | -0.5864125 | 0.56626069 | 0.99999785 | -6.2104671 | down |
| KI - WT, cell_origin: iPSC | Q96JY6 | PDLIM2 | 5 | Homo sapien | -0.2347686 | -1.088204 | 0.61866678 | 0.20127923 | -0.5860269 | 0.56652008 | 0.99999785 | -6.2106889 | down |
| KI - WT, cell_origin: iPSC | P52298 | NCBP2 | 7 | Homo sapien | -0.3172043 | -1.4732417 | 0.83883312 | 1.52673387 | -0.5847491 | 0.56739216 | 0.99999785 | -6.2114102 | down |
| KI - WT, cell_origin: iPSC | P42768 | WAS | 10 | Homo sapien | -0.7133814 | -3.3133609 | 1.88659821 | 0.17399852 | -0.5847284 | 0.56740571 | 0.99999785 | -6.2114224 | down |
| KI - WT, cell_origin: iPSC | Q9NUQ9 | CYRIB | 14 | Homo sapien | 0.34230246 | -0.9081005 | 1.59270543 | 0.87042297 | 0.58339458 | 0.56828008 | 0.99999785 | -6.2122118 | up |
| KI - WT, cell_origin: iPSC | Q6YN16 | HSDL2 | 2 | Homo sapien | -0.3150001 | -1.4670754 | 0.83707515 | 0.13815781 | -0.5826828 | 0.56874695 | 0.99999785 | -6.2126323 | down |
| KI - WT, cell_origin: iPSC | Q6PD62 | CTR9 | 8 | Homo sapien | 0.31539713 | -0.8418091 | 1.47260335 | -1.3496077 | 0.5808304 | 0.56996301 | 0.99999785 | -6.2137244 | up |
| KI - WT, cell_origin: iPSC | Q8NFH4 | NUP37 | 6 | Homo sapien | -0.2429987 | -1.1347906 | 0.64879326 | -1.035753 | -0.5804412 | 0.57017712 | 0.99999785 | -6.2139965 | down |
| KI - WT, cell_origin: iPSC | Q96EE3 | SEH1L | 10 | Homo sapien | 0.16951241 | -0.4527991 | 0.79182396 | 0.40816698 | 0.58014068 | 0.57035702 | 0.99999785 | -6.2141915 | up |
| KI - WT, cell_origin: iPSC | Q9NWB6 | ARGLU1 | 16 | Homo sapien | -0.4908264 | -2.2950687 | 1.31341598 | 3.5239964 | -0.5797428 | 0.57067758 | 0.99999785 | -6.214364 | down |
| KI - WT, cell_origin: iPSC | O00505 | KPNA3 | 7 | Homo sapien | 0.19685675 | -0.5265246 | 0.92023808 | -0.2265121 | 0.57959243 | 0.5707174 | 0.99999785 | -6.2145137 | up |
| KI - WT, cell_origin: iPSC | Q96HY6 | DDRGK1 | 2 | Homo sapien | -0.5618058 | -2.6281085 | 1.5044969 | -2.8099191 | -0.5794215 | 0.57088878 | 0.99999785 | -6.2145528 | down |
| KI - WT, cell_origin: iPSC | Q6PIY7 | TENT2 | 3 | Homo sapien | -0.2418922 | -1.1313704 | 0.64758604 | -0.9765831 | -0.5793011 | 0.57092649 | 0.99999785 | -6.2146665 | down |
| KI - WT, cell_origin: iPSC | Q9H336 | CRISPLD1 | 2 | Homo sapien | -0.3837281 | -1.7966519 | 1.02919573 | -2.845439 | -0.5785258 | 0.57143642 | 0.99999785 | -6.2151214 | down |
| KI - WT, cell_origin: iPSC | Q99623 | PHB2 | 4 | Homo sapien | 0.51998543 | -1.3972967 | 2.43726752 | 3.78167593 | 0.57797297 | 0.57184147 | 0.99999785 | -6.2154025 | up |
| KI - WT, cell_origin: iPSC | P61289 | PSME3 | 10 | Homo sapien | 0.26396431 | -0.7097337 | 1.23766237 | -0.2166551 | 0.57772783 | 0.57200278 | 0.99999785 | -6.2155461 | up |
| KI - WT, cell_origin: iPSC | Q15311 | RALBP1 | 2 | Homo sapien | -0.3367139 | -1.5804765 | 0.90704866 | 0.35073501 | -0.5769338 | 0.57252543 | 0.99999785 | -6.2160109 | down |
| KI - WT, cell_origin: iPSC | Q8TED0 | UTP15 | 4 | Homo sapien | 0.39273707 | -1.058691 | 1.84416518 | -2.9739787 | 0.5764004 | 0.57283547 | 0.99999785 | -6.2163654 | up |
| KI - WT, cell_origin: iPSC | Q86XZ4 | SPATS2 | 5 | Homo sapien | -0.3527508 | -1.6572074 | 0.95170583 | -1.487671 | -0.5762895 | 0.57294968 | 0.99999785 | -6.2163875 | down |

| | | | | | | | | | | | | | |
|---|---|---|---|---|---|---|---|---|---|---|---|---|---|
| KI - WT, cell_origin: iPSC | O43823 | AKAP8 | 5 | Homo sapien | -0.1318851 | -0.6195462 | 0.35577589 | 0.42333431 | -0.5759934 | 0.57308614 | 0.99999785 | -6.2166212 | down |
| KI - WT, cell_origin: iPSC | Q9UK39 | NOCT | 15 | Homo sapien | 0.3663321 | -0.989345 | 1.72200921 | 0.50768406 | 0.57586553 | 0.57322898 | 0.99999785 | -6.2166351 | up |
| KI - WT, cell_origin: iPSC | O94842 | TOX4 | 5 | Homo sapien | -0.3658207 | -1.7197977 | 0.98815632 | -1.8906115 | -0.5757837 | 0.5732829 | 0.99999785 | -6.2166829 | down |
| KI - WT, cell_origin: iPSC | Q5PRF9 | SAMD4B | 6 | Homo sapien | -0.2222937 | -1.0450411 | 0.60045373 | -0.7691517 | -0.5755293 | 0.57340691 | 0.99999785 | -6.2168765 | down |
| KI - WT, cell_origin: iPSC | O75717 | WDHD1 | 13 | Homo sapien | -0.2950218 | -1.3881867 | 0.79814316 | -2.0984085 | -0.5748626 | 0.57384389 | 0.99999785 | -6.2172678 | down |
| KI - WT, cell_origin: iPSC | Q9P2H3 | IFT80 | 2 | Homo sapien | 0.16837614 | -0.4554718 | 0.79222408 | -0.9803217 | 0.57483271 | 0.57385112 | 0.99999785 | -6.2172981 | up |
| KI - WT, cell_origin: iPSC | P62191 | PSMC1 | 8 | Homo sapien | 0.2855607 | -0.7731989 | 1.34432032 | -1.101213 | 0.57478235 | 0.57394281 | 0.99999785 | -6.217267 | up |
| KI - WT, cell_origin: iPSC | Q15643 | TRIP11 | 30 | Homo sapien | 0.44853918 | -1.2148725 | 2.11195088 | -1.3286244 | 0.57464946 | 0.57403042 | 0.99999785 | -6.2173445 | up |
| KI - WT, cell_origin: iPSC | Q9Y2H5 | PLEKHA6 | 2 | Homo sapien | -0.2791294 | -1.3139884 | 0.75572957 | -2.1432302 | -0.5744657 | 0.57409312 | 0.99999785 | -6.2175119 | down |
| KI - WT, cell_origin: iPSC | P55196 | AFDN | 47 | Homo sapien | 0.26917914 | -0.7313428 | 1.26970112 | 0.89833542 | 0.57334645 | 0.5748898 | 0.99999785 | -6.2181029 | up |
| KI - WT, cell_origin: iPSC | O75368 | SH3BGRL | 2 | Homo sapien | -0.4516585 | -2.1330453 | 1.22972826 | -1.7230499 | -0.5724597 | 0.57547501 | 0.99999785 | -6.2186181 | down |
| KI - WT, cell_origin: iPSC | Q9H4Z3 | PCIF1 | 2 | Homo sapien | 0.35764337 | -0.9746118 | 1.68989859 | -2.7800693 | 0.57183344 | 0.57584526 | 0.99999785 | -6.2190261 | up |
| KI - WT, cell_origin: iPSC | Q9BPW9 | DHRS9 | 5 | Homo sapien | -0.5730131 | -2.7090385 | 1.56301222 | -2.4488176 | -0.5716899 | 0.57598332 | 0.99999785 | -6.2190647 | down |
| KI - WT, cell_origin: iPSC | P13010 | XRCC5 | 30 | Homo sapien | -0.3024941 | -1.4301655 | 0.82517738 | 3.24098834 | -0.5716585 | 0.57600408 | 0.99999785 | -6.2190829 | down |
| KI - WT, cell_origin: iPSC | Q9UBP9 | GULP1 | 2 | Homo sapien | 0.27066272 | -0.7413073 | 1.28263272 | -2.8151821 | 0.56964 | 0.57728014 | 0.99999785 | -6.2203109 | up |
| KI - WT, cell_origin: iPSC | P61978 | HNRNPK | 22 | Homo sapien | -0.2303124 | -1.0922782 | 0.6316535 | 4.32080761 | -0.5694159 | 0.57748625 | 0.99999785 | -6.2203807 | down |
| KI - WT, cell_origin: iPSC | O00571 | DDX3X | 47 | Homo sapien | -0.1416693 | -0.6725504 | 0.38921192 | 5.29862261 | -0.5683528 | 0.57813181 | 0.99999785 | -6.2210537 | down |
| KI - WT, cell_origin: iPSC | O43427 | FIBP | 9 | Homo sapien | 0.25036299 | -0.6880255 | 1.18875151 | -1.2320241 | 0.56833624 | 0.57815994 | 0.99999785 | -6.2210455 | up |
| KI - WT, cell_origin: iPSC | Q9NXR1 | NDE1 | 2 | Homo sapien | -0.2361228 | -1.1215944 | 0.64934869 | -1.2431019 | -0.5680141 | 0.57836832 | 0.99999785 | -6.2212362 | down |
| KI - WT, cell_origin: iPSC | P31939 | ATIC | 9 | Homo sapien | 0.35022049 | -0.9639908 | 1.66443179 | -1.9622713 | 0.56790897 | 0.57848326 | 0.99999785 | -6.22125 | up |
| KI - WT, cell_origin: iPSC | Q8IWA0 | WDR75 | 7 | Homo sapien | -0.3189152 | -1.5159199 | 0.87808959 | -0.2696429 | -0.5677821 | 0.57856728 | 0.99999785 | -6.2213231 | down |
| KI - WT, cell_origin: iPSC | Q9NWK9 | ZNHIT6 | 3 | Homo sapien | 0.55717298 | -1.534309 | 2.64865495 | -2.7969375 | 0.56772535 | 0.57860481 | 0.99999785 | -6.2213558 | up |
| KI - WT, cell_origin: iPSC | Q15370 | ELOB | 5 | Homo sapien | -0.1534279 | -0.7295894 | 0.42273359 | 1.68357545 | -0.5671524 | 0.5789266 | 0.99999785 | -6.2217449 | down |
| KI - WT, cell_origin: iPSC | Q9H2D6 | TRIOBP | 20 | Homo sapien | -0.1863909 | -0.8864275 | 0.51364582 | 0.57479588 | -0.5670789 | 0.57897528 | 0.99999785 | -6.2217871 | down |
| KI - WT, cell_origin: iPSC | Q8N201 | INTS1 | 24 | Homo sapien | 0.13054598 | -0.3599039 | 0.62099583 | -0.0929534 | 0.56690279 | 0.57909193 | 0.99999785 | -6.2218884 | up |
| KI - WT, cell_origin: iPSC | Q13043 | STK4 | 2 | Homo sapien | -0.3171219 | -1.509903 | 0.87565925 | -1.3849155 | -0.5665886 | 0.57935763 | 0.99999785 | -6.2220099 | down |
| KI - WT, cell_origin: iPSC | Q6DD87 | ZNF787 | 4 | Homo sapien | -0.2171372 | -1.036752 | 0.60247758 | -0.7738562 | -0.5643185 | 0.58081829 | 0.99999785 | -6.2233577 | down |
| KI - WT, cell_origin: iPSC | P61011 | SRP54 | 20 | Homo sapien | 0.33407283 | -0.9289021 | 1.59704781 | 1.30820461 | 0.56370096 | 0.58127217 | 0.99999785 | -6.2236659 | up |
| KI - WT, cell_origin: iPSC | Q9H0W8 | SMG9 | 7 | Homo sapien | 0.62300897 | -1.7334895 | 2.97950746 | -1.0718702 | 0.56341643 | 0.581461 | 0.99999785 | -6.2238286 | up |
| KI - WT, cell_origin: iPSC | Q5TFE4 | NT5DC1 | 3 | Homo sapien | -0.2710272 | -1.2986116 | 0.75655713 | -2.0137976 | -0.561777 | 0.58249888 | 0.99999785 | -6.2248166 | down |
| KI - WT, cell_origin: iPSC | Q3MHD2 | LSM12 | 4 | Homo sapien | -0.3075989 | -1.4765618 | 0.86136402 | 1.84402497 | -0.5607721 | 0.58321737 | 0.99999785 | -6.2253372 | down |
| KI - WT, cell_origin: iPSC | Q8WUY9 | DEPDC1B | 3 | Homo sapien | 0.21354844 | -0.5985798 | 1.02567664 | -1.5565499 | 0.56003026 | 0.58365387 | 0.99999785 | -6.2258172 | up |
| KI - WT, cell_origin: iPSC | Q13162 | PRDX4 | 7 | Homo sapien | 0.61915548 | -1.7374672 | 2.9757782 | 0.26048271 | 0.55990201 | 0.58379591 | 0.99999785 | -6.2258321 | up |
| KI - WT, cell_origin: iPSC | Q9UGJ1 | TUBGCP4 | 13 | Homo sapien | 0.12021899 | -0.3371764 | 0.57761436 | 0.208117 | 0.55978467 | 0.58381723 | 0.99999785 | -6.2259568 | up |
| KI - WT, cell_origin: iPSC | P30044 | PRDX5 | 2 | Homo sapien | -0.2722119 | -1.3090167 | 0.76459283 | -0.5755012 | -0.559516 | 0.58405265 | 0.99999785 | -6.2260514 | down |
| KI - WT, cell_origin: iPSC | P00813 | ADA | 9 | Homo sapien | 0.27553921 | -0.7739673 | 1.32504571 | -0.4218998 | 0.55950071 | 0.58406283 | 0.99999785 | -6.2260601 | up |
| KI - WT, cell_origin: iPSC | Q8TD19 | NEK9 | 16 | Homo sapien | 0.21509057 | -0.6039784 | 1.03415952 | -0.0741723 | 0.55939528 | 0.58409317 | 0.99999785 | -6.2261607 | up |
| KI - WT, cell_origin: iPSC | P41134 | ID1 | 2 | Homo sapien | -0.5314775 | -2.5595022 | 1.49654723 | -1.7224471 | -0.5584882 | 0.58473659 | 0.99999785 | -6.2266347 | down |
| KI - WT, cell_origin: iPSC | Q13573 | SNW1 | 8 | Homo sapien | -0.3063499 | -1.4763274 | 0.86362766 | -0.0041207 | -0.5580107 | 0.58505444 | 0.99999785 | -6.2269053 | down |
| KI - WT, cell_origin: iPSC | Q16513 | PKN2 | 59 | Homo sapien | 0.33063824 | -0.933167 | 1.59444346 | 6.89419893 | 0.55753906 | 0.58536852 | 0.99999785 | -6.2271724 | up |
| KI - WT, cell_origin: iPSC | Q96HP0 | DOCK6 | 56 | Homo sapien | 0.27326762 | -0.771283 | 1.31781823 | -0.442897 | 0.55752078 | 0.58538068 | 0.99999785 | -6.2271827 | up |
| KI - WT, cell_origin: iPSC | O76071 | CIAO1 | 10 | Homo sapien | 0.29090789 | -0.8239101 | 1.40572584 | 0.31825349 | 0.55610131 | 0.58632644 | 0.99999785 | -6.2279853 | up |
| KI - WT, cell_origin: iPSC | A7E2V4 | ZSWIM8 | 35 | Homo sapien | 0.19090592 | -0.5406222 | 0.92243407 | 1.69715994 | 0.55585208 | 0.58644299 | 0.99999785 | -6.2281765 | up |
| KI - WT, cell_origin: iPSC | Q8IY37 | DHX37 | 8 | Homo sapien | -0.2171249 | -1.0509157 | 0.61666588 | -1.6995563 | -0.5546158 | 0.58726091 | 0.99999785 | -6.2288802 | down |
| KI - WT, cell_origin: iPSC | P50570 | DNM2 | 14 | Homo sapien | 0.28625185 | -0.8139064 | 1.38641014 | -0.0955333 | 0.55449227 | 0.58739943 | 0.99999785 | -6.2288926 | up |
| KI - WT, cell_origin: iPSC | Q9UL15 | BAG5 | 5 | Homo sapien | 0.18560434 | -0.5280801 | 0.8992888 | -1.2429608 | 0.5538875 | 0.58774694 | 0.99999785 | -6.22929 | up |

| | | | | | | | | | | | | | |
|---|---|---|---|---|---|---|---|---|---|---|---|---|---|
| KI - WT, cell_origin: iPSC | P29558 | RBMS1 | 5 | Homo sapien | 0.16665414 | -0.4744966 | 0.80780489 | 1.96615282 | 0.55359942 | 0.58793925 | 0.99999785 | -6.229452 | up |
| KI - WT, cell_origin: iPSC | Q8NDT2 | RBM15B | 5 | Homo sapien | -0.2489805 | -1.2088725 | 0.71091139 | -1.6431679 | -0.552493 | 0.58868755 | 0.99999785 | -6.2300638 | down |
| KI - WT, cell_origin: iPSC | Q6PJ69 | TRIM65 | 14 | Homo sapien | 0.20235654 | -0.5792767 | 0.98398975 | 2.18235164 | 0.55148303 | 0.58936968 | 0.99999785 | -6.2306225 | up |
| KI - WT, cell_origin: iPSC | P61204;P840 | ARF3;ARF1 | 11 | Homo sapien | 0.27277942 | -0.783523 | 1.32908185 | 5.64785841 | 0.55033317 | 0.59017757 | 0.99999785 | -6.2312262 | up |
| KI - WT, cell_origin: iPSC | Q96JI7 | SPG11 | 18 | Homo sapien | 0.17254065 | -0.4959149 | 0.8409962 | -0.5775096 | 0.54974155 | 0.59051774 | 0.99999785 | -6.2316131 | up |
| KI - WT, cell_origin: iPSC | Q9Y2I7 | PIKFYVE | 4 | Homo sapien | 0.34186184 | -0.9833823 | 1.66710597 | -2.3336226 | 0.54958794 | 0.59065072 | 0.99999785 | -6.2316682 | up |
| KI - WT, cell_origin: iPSC | O95071 | UBR5 | 22 | Homo sapien | 0.24601429 | -0.710057 | 1.20208557 | -0.9350642 | 0.54825016 | 0.59155171 | 0.99999785 | -6.2324086 | up |
| KI - WT, cell_origin: iPSC | Q6P6C2 | ALKBH5 | 3 | Homo sapien | -0.4993061 | -2.4407435 | 1.44213133 | -2.5113655 | -0.5480824 | 0.59168381 | 0.99999785 | -6.232482 | down |
| KI - WT, cell_origin: iPSC | Q1ED39 | KNOP1 | 11 | Homo sapien | -0.1473265 | -0.7200648 | 0.42541179 | 0.45929192 | -0.5478534 | 0.5917818 | 0.99999785 | -6.2326656 | down |
| KI - WT, cell_origin: iPSC | Q9NPF4 | OSGEP | 3 | Homo sapien | -0.1778215 | -0.8693665 | 0.51372353 | -0.1217509 | -0.5476504 | 0.59191778 | 0.99999785 | -6.2327785 | down |
| KI - WT, cell_origin: iPSC | Q9P016 | THYN1 | 9 | Homo sapien | -0.2775845 | -1.359521 | 0.804352 | -0.1102939 | -0.5467588 | 0.59257043 | 0.99999785 | -6.2332182 | down |
| KI - WT, cell_origin: iPSC | Q96EB6 | SIRT1 | 5 | Homo sapien | -0.2953207 | -1.4465721 | 0.85593065 | -2.0052357 | -0.546439 | 0.59274607 | 0.99999785 | -6.2334351 | down |
| KI - WT, cell_origin: iPSC | Q93034 | CUL5 | 6 | Homo sapien | 0.26232272 | -0.7616739 | 1.28631935 | -1.1350367 | 0.54585997 | 0.59316067 | 0.99999785 | -6.2337296 | up |
| KI - WT, cell_origin: iPSC | P60953 | CDC42 | 10 | Homo sapien | 0.23040135 | -0.6692766 | 1.13007929 | 6.93049191 | 0.54575827 | 0.59324114 | 0.99999785 | -6.2337736 | up |
| KI - WT, cell_origin: iPSC | Q13510 | ASAH1 | 9 | Homo sapien | 0.35346714 | -1.0273161 | 1.73425038 | -0.2908363 | 0.54553913 | 0.59338808 | 0.99999785 | -6.2338951 | up |
| KI - WT, cell_origin: iPSC | Q8IV48 | ERI1 | 4 | Homo sapien | 0.23753718 | -0.6904807 | 1.16555508 | -1.9501144 | 0.54514864 | 0.59359496 | 0.99999785 | -6.2341671 | up |
| KI - WT, cell_origin: iPSC | O94973 | AP2A2 | 27 | Homo sapien | 0.18460613 | -0.5367259 | 0.90593812 | 1.77590701 | 0.54509844 | 0.59363373 | 0.99999785 | -6.2341898 | up |
| KI - WT, cell_origin: iPSC | P62140 | PPP1CB | 4 | Homo sapien | 0.18743223 | -0.5452386 | 0.92010304 | 0.08271325 | 0.54485513 | 0.59379313 | 0.99999785 | -6.2343284 | up |
| KI - WT, cell_origin: iPSC | P60842 | EIF4A1 | 21 | Homo sapien | -0.178039 | -0.8743728 | 0.51829473 | 3.48508429 | -0.5446338 | 0.59395445 | 0.99999785 | -6.2344379 | down |
| KI - WT, cell_origin: iPSC | P38935 | IGHMBP2 | 3 | Homo sapien | 0.45043421 | -1.312707 | 2.21357544 | -2.8484554 | 0.54443575 | 0.59412825 | 0.99999785 | -6.2345062 | up |
| KI - WT, cell_origin: iPSC | O00244 | ATOX1 | 4 | Homo sapien | -0.1860303 | -0.9145231 | 0.54246246 | 0.71206073 | -0.5439169 | 0.59442876 | 0.99999785 | -6.2348413 | down |
| KI - WT, cell_origin: iPSC | Q9Y485 | DMXL1 | 22 | Homo sapien | 0.17304904 | -0.504695 | 0.85079313 | -0.5657647 | 0.54380493 | 0.59449676 | 0.99999785 | -6.2349104 | up |
| KI - WT, cell_origin: iPSC | Q8NDA8 | MROH1 | 11 | Homo sapien | 0.4108553 | -1.2015259 | 2.02323655 | -1.8123374 | 0.54302963 | 0.59507218 | 0.99999785 | -6.2352832 | up |
| KI - WT, cell_origin: iPSC | O00161 | SNAP23 | 4 | Homo sapien | -0.324859 | -1.6008701 | 0.9511521 | 1.44443341 | -0.5425538 | 0.59539176 | 0.99999785 | -6.2355457 | down |
| KI - WT, cell_origin: iPSC | P04183 | TK1 | 8 | Homo sapien | 0.29326358 | -0.8597662 | 1.44629338 | 0.4158491 | 0.54202582 | 0.5957465 | 0.99999785 | -6.2358368 | up |
| KI - WT, cell_origin: iPSC | Q9Y3L5 | RAP2C | 2 | Homo sapien | 0.35917572 | -1.0530507 | 1.77140209 | 2.68376719 | 0.54200716 | 0.59575904 | 0.99999785 | -6.2358471 | up |
| KI - WT, cell_origin: iPSC | O43739 | CYTH3 | 2 | Homo sapien | -0.204297 | -1.00788 | 0.59928601 | 0.37358726 | -0.5415632 | 0.59601905 | 0.99999785 | -6.2361302 | down |
| KI - WT, cell_origin: iPSC | A6NIZ1 | | 4 | Homo sapien | 0.28486438 | -0.8384124 | 1.40814114 | 6.17378921 | 0.54044777 | 0.59680734 | 0.99999785 | -6.236705 | up |
| KI - WT, cell_origin: iPSC | P56270 | MAZ | 5 | Homo sapien | -0.4323525 | -2.137462 | 1.27275694 | -1.0011564 | -0.5403661 | 0.59686225 | 0.99999785 | -6.2367498 | down |
| KI - WT, cell_origin: iPSC | P02533 | KRT14 | 25 | Homo sapien | -0.444969 | -2.2025959 | 1.31265796 | 3.94360079 | -0.5395173 | 0.59743328 | 0.99999785 | -6.2372158 | down |
| KI - WT, cell_origin: iPSC | Q9Y4B5 | MTCL1 | 7 | Homo sapien | -0.3566703 | -1.7659404 | 1.05259972 | -1.9418312 | -0.5393555 | 0.59754216 | 0.99999785 | -6.2373045 | down |
| KI - WT, cell_origin: iPSC | Q13427 | PPIG | 9 | Homo sapien | -0.3612469 | -1.7889242 | 1.0664304 | -0.3351715 | -0.539233 | 0.59762464 | 0.99999785 | -6.2373717 | down |
| KI - WT, cell_origin: iPSC | Q16342 | PDCD2 | 7 | Homo sapien | 0.19559776 | -0.5775583 | 0.96875377 | 0.30467399 | 0.53888556 | 0.59781661 | 0.99999785 | -6.2376042 | up |
| KI - WT, cell_origin: iPSC | P62861 | FAU | 2 | Homo sapien | 0.32324074 | -0.9551772 | 1.60165872 | 6.33291418 | 0.53883474 | 0.59789268 | 0.99999785 | -6.2375899 | up |
| KI - WT, cell_origin: iPSC | Q96CB8 | INTS12 | 2 | Homo sapien | 0.36261635 | -1.0719547 | 1.79718738 | -1.0320484 | 0.53867609 | 0.59799948 | 0.99999785 | -6.2376768 | up |
| KI - WT, cell_origin: iPSC | P50748 | KNTC1 | 7 | Homo sapien | 0.23021967 | -0.6802881 | 1.14072741 | -2.1257006 | 0.53851581 | 0.59805314 | 0.99999785 | -6.2378191 | up |
| KI - WT, cell_origin: iPSC | Q9UNW9 | NOVA2 | 3 | Homo sapien | 0.19315842 | -0.5709138 | 0.95723063 | -1.3944744 | 0.53841719 | 0.59811955 | 0.99999785 | -6.2378731 | up |
| KI - WT, cell_origin: iPSC | Q96MY1 | NOL4L | 7 | Homo sapien | -0.6608271 | -3.2772811 | 1.95562696 | -2.721791 | -0.5382414 | 0.59829216 | 0.99999785 | -6.2379148 | down |
| KI - WT, cell_origin: iPSC | P38571 | LIPA | 3 | Homo sapien | 0.34910964 | -1.0351232 | 1.73334245 | -0.2775128 | 0.53747104 | 0.59881102 | 0.99999785 | -6.2383361 | up |
| KI - WT, cell_origin: iPSC | P12270 | TPR | 2 | Homo sapien | 0.34701743 | -1.0281151 | 1.72214993 | -3.3269908 | 0.53746035 | 0.59876409 | 0.99999785 | -6.2383963 | up |
| KI - WT, cell_origin: iPSC | P54277 | PMS1 | 5 | Homo sapien | 0.50913281 | -1.5099955 | 2.52826117 | -1.3015097 | 0.53736519 | 0.59888233 | 0.99999785 | -6.2383939 | up |
| KI - WT, cell_origin: iPSC | Q92974 | ARHGEF2 | 55 | Homo sapien | 0.47249162 | -1.402035 | 2.34701824 | 3.76876858 | 0.53716149 | 0.59901958 | 0.99999785 | -6.2385052 | up |
| KI - WT, cell_origin: iPSC | Q15436 | SEC23A | 24 | Homo sapien | -0.1419323 | -0.705165 | 0.42130031 | 2.47852464 | -0.536702 | 0.5992752 | 0.99999785 | -6.2388103 | down |
| KI - WT, cell_origin: iPSC | Q9Y5K6 | CD2AP | 9 | Homo sapien | 0.19705185 | -0.5852583 | 0.97936199 | -0.4891617 | 0.53651917 | 0.59940748 | 0.99999785 | -6.238901 | up |
| KI - WT, cell_origin: iPSC | Q12756 | KIF1A | 27 | Homo sapien | 0.14539936 | -0.4320648 | 0.72286348 | -0.3060981 | 0.5362622 | 0.59957168 | 0.99999785 | -6.2390502 | up |
| KI - WT, cell_origin: iPSC | O75554 | WBP4 | 6 | Homo sapien | -0.123795 | -0.6154785 | 0.3678885 | -0.8879416 | -0.5362374 | 0.59958839 | 0.99999785 | -6.2390637 | down |

| | | | | | | | | | | | | | |
|---|---|---|---|---|---|---|---|---|---|---|---|---|---|
| KI - WT, cell_origin: iPSC | Q86YM7 | HOMER1 | 11 | Homo sapien | 0.49025525 | -1.4592743 | 2.43978476 | -1.2903463 | 0.53591364 | 0.59986067 | 0.99999785 | -6.239186 | up |
| KI - WT, cell_origin: iPSC | P00492 | HPRT1 | 7 | Homo sapien | -0.1738145 | -0.8658328 | 0.51820376 | 0.36407872 | -0.5349438 | 0.60046096 | 0.99999785 | -6.2397681 | down |
| KI - WT, cell_origin: iPSC | P42679 | MATK | 3 | Homo sapien | -0.3772486 | -1.8808979 | 1.12640069 | -1.2773692 | -0.5346671 | 0.60070147 | 0.99999785 | -6.2398646 | down |
| KI - WT, cell_origin: iPSC | Q8N122 | RPTOR | 12 | Homo sapien | 0.15979052 | -0.4772444 | 0.79682545 | -0.1415628 | 0.53422897 | 0.60094338 | 0.99999785 | -6.2401566 | up |
| KI - WT, cell_origin: iPSC | Q9BQ61 | TRIR | 3 | Homo sapien | -0.3059365 | -1.5274079 | 0.91553493 | 2.58074636 | -0.5337653 | 0.60131012 | 0.99999785 | -6.2403546 | down |
| KI - WT, cell_origin: iPSC | Q5M775 | SPECC1 | 17 | Homo sapien | 0.31318626 | -0.940434 | 1.56680649 | -0.3041415 | 0.53240127 | 0.60223128 | 0.99999785 | -6.2410942 | up |
| KI - WT, cell_origin: iPSC | Q13190 | STX5 | 6 | Homo sapien | 0.44882071 | -1.3481098 | 2.2457512 | -0.2764431 | 0.5322847 | 0.60231004 | 0.99999785 | -6.2411573 | up |
| KI - WT, cell_origin: iPSC | Q8IVI9 | NOSTRIN | 3 | Homo sapien | -0.2186113 | -1.0947275 | 0.65750482 | -2.0596701 | -0.5314356 | 0.60283041 | 0.99999785 | -6.24167 | down |
| KI - WT, cell_origin: iPSC | P62834 | RAP1A | 13 | Homo sapien | 0.20631772 | -0.6211942 | 1.03382966 | 8.19256891 | 0.53133036 | 0.602955 | 0.99999785 | -6.2416736 | up |
| KI - WT, cell_origin: iPSC | Q9Y3A4 | RRP7A | 5 | Homo sapien | 0.30911103 | -0.930941 | 1.54916308 | -2.4059227 | 0.53099222 | 0.60314524 | 0.99999785 | -6.2418946 | up |
| KI - WT, cell_origin: iPSC | Q9ULK4 | MED23 | 10 | Homo sapien | -0.1864292 | -0.9344414 | 0.56158301 | -0.8860487 | -0.5308172 | 0.60324861 | 0.99999785 | -6.2420041 | down |
| KI - WT, cell_origin: iPSC | Q9Y4A5 | TRRAP | 11 | Homo sapien | -0.2061717 | -1.0346446 | 0.6223012 | -2.4293023 | -0.5300177 | 0.6037894 | 0.99999785 | -6.2424353 | down |
| KI - WT, cell_origin: iPSC | P50281 | MMP14 | 4 | Homo sapien | 0.25826878 | -0.780327 | 1.29686451 | 0.39759399 | 0.52994127 | 0.60389439 | 0.99999785 | -6.2424234 | up |
| KI - WT, cell_origin: iPSC | Q9H2H8 | PPIL3 | 4 | Homo sapien | -0.4354562 | -2.188289 | 1.31737667 | -1.0436736 | -0.5294273 | 0.60424216 | 0.99999785 | -6.2427004 | down |
| KI - WT, cell_origin: iPSC | Q9Y3D9 | MRPS23 | 2 | Homo sapien | -0.7424234 | -3.7311772 | 2.24633047 | -3.5287384 | -0.5293755 | 0.60427718 | 0.99999785 | -6.2427283 | down |
| KI - WT, cell_origin: iPSC | P47897 | QARS1 | 25 | Homo sapien | 0.17047476 | -0.5155602 | 0.85650966 | 0.29940054 | 0.52924108 | 0.60431499 | 0.99999785 | -6.2428537 | up |
| KI - WT, cell_origin: iPSC | P49321 | NASP | 18 | Homo sapien | -0.270796 | -1.3618824 | 0.8202905 | -0.2664477 | -0.5289144 | 0.60458929 | 0.99999785 | -6.2429765 | down |
| KI - WT, cell_origin: iPSC | Q9NUJ7 | PLCXD1 | 5 | Homo sapien | 0.18044988 | -0.5461963 | 0.90709608 | -1.3164832 | 0.52889965 | 0.60454612 | 0.99999785 | -6.2430374 | up |
| KI - WT, cell_origin: iPSC | P84085 | ARF5 | 6 | Homo sapien | 0.26025414 | -0.7884655 | 1.30897382 | 1.6288136 | 0.52885986 | 0.6046262 | 0.99999785 | -6.2430059 | up |
| KI - WT, cell_origin: iPSC | Q9NQC7 | CYLD | 5 | Homo sapien | -0.25696 | -1.2929928 | 0.77907292 | -0.8759131 | -0.52856 | 0.6048292 | 0.99999785 | -6.2431672 | down |
| KI - WT, cell_origin: iPSC | Q969M3 | YIPF5 | 2 | Homo sapien | -0.5758378 | -2.9021091 | 1.7504336 | 1.77947519 | -0.5275239 | 0.60553088 | 0.99999785 | -6.2437238 | down |
| KI - WT, cell_origin: iPSC | Q5T7W0 | ZNF618 | 4 | Homo sapien | -0.3281426 | -1.6540331 | 0.9977478 | -1.2844197 | -0.527421 | 0.60560062 | 0.99999785 | -6.2437791 | down |
| KI - WT, cell_origin: iPSC | Q96ST3 | SIN3A | 12 | Homo sapien | 0.80078459 | -2.435984 | 4.0375532 | -3.2892697 | 0.52723766 | 0.6057248 | 0.99999785 | -6.2438774 | up |
| KI - WT, cell_origin: iPSC | Q9Y2P8 | RCL1 | 6 | Homo sapien | 0.1413535 | -0.4297753 | 0.71248228 | -1.0504527 | 0.52712326 | 0.60574937 | 0.99999785 | -6.2439915 | up |
| KI - WT, cell_origin: iPSC | P49005 | POLD2 | 5 | Homo sapien | -0.1637886 | -0.826721 | 0.49914377 | -0.0684243 | -0.526204 | 0.6063725 | 0.99999785 | -6.244484 | down |
| KI - WT, cell_origin: iPSC | Q8TEV9 | SMCR8 | 23 | Homo sapien | 0.31481341 | -0.9614864 | 1.59111323 | 0.6177177 | 0.52565751 | 0.60679587 | 0.99999785 | -6.2447239 | up |
| KI - WT, cell_origin: iPSC | P82979 | SARNP | 6 | Homo sapien | -0.4623972 | -2.3397166 | 1.41492212 | -0.5977752 | -0.5249035 | 0.60730731 | 0.99999785 | -6.245127 | down |
| KI - WT, cell_origin: iPSC | Q9NXG2 | THUMPD1 | 4 | Homo sapien | 0.1282111 | -0.3920689 | 0.64849107 | -1.1329498 | 0.52484155 | 0.60729664 | 0.99999785 | -6.2452124 | up |
| KI - WT, cell_origin: iPSC | O14545 | TRAFD1 | 4 | Homo sapien | 0.2236987 | -0.6842633 | 1.13166069 | -1.6149031 | 0.52473142 | 0.60737169 | 0.99999785 | -6.2452709 | up |
| KI - WT, cell_origin: iPSC | P62633 | CNBP | 10 | Homo sapien | -0.2516992 | -1.2745465 | 0.7711481 | 2.02024417 | -0.5244129 | 0.60764013 | 0.99999785 | -6.245389 | down |
| KI - WT, cell_origin: iPSC | Q96P70 | IPO9 | 8 | Homo sapien | 0.33731945 | -1.0343172 | 1.7089561 | -1.3345779 | 0.52408856 | 0.60786027 | 0.99999785 | -6.245562 | up |
| KI - WT, cell_origin: iPSC | Q9UNX3 | RPL26L1 | 8 | Homo sapien | -0.119682 | -0.6060913 | 0.36672729 | 3.72666648 | -0.5240426 | 0.60783886 | 0.99999785 | -6.2456387 | down |
| KI - WT, cell_origin: iPSC | Q99426 | TBCB | 4 | Homo sapien | -0.1895306 | -0.9621348 | 0.5830736 | -0.8166574 | -0.5224779 | 0.60890269 | 0.99999785 | -6.2464706 | down |
| KI - WT, cell_origin: iPSC | P08134 | RHOC | 9 | Homo sapien | -0.2511569 | -1.2760344 | 0.77372053 | 5.02906789 | -0.5222466 | 0.60911108 | 0.99999785 | -6.2465429 | down |
| KI - WT, cell_origin: iPSC | Q8IZH2 | XRN1 | 25 | Homo sapien | 0.15565539 | -0.4796497 | 0.79096051 | 0.07647993 | 0.52182091 | 0.60934797 | 0.99999785 | -6.2468209 | up |
| KI - WT, cell_origin: iPSC | Q00325 | SLC25A3 | 3 | Homo sapien | 0.49428575 | -1.524388 | 2.51295953 | -0.8872906 | 0.52181231 | 0.60940615 | 0.99999785 | -6.2467737 | up |
| KI - WT, cell_origin: iPSC | Q9Y2X3 | NOP58 | 18 | Homo sapien | -0.2379625 | -1.2106416 | 0.73471663 | -0.25788 | -0.5213643 | 0.60971064 | 0.99999785 | -6.2470115 | down |
| KI - WT, cell_origin: iPSC | Q9P1F3 | ABRACL | 3 | Homo sapien | -0.1700666 | -0.8661005 | 0.5259672 | 0.79710734 | -0.5203894 | 0.61032131 | 0.99999785 | -6.24758 | down |
| KI - WT, cell_origin: iPSC | Q8NEY1 | NAV1 | 9 | Homo sapien | 0.31010353 | -0.961711 | 1.58191807 | -1.2635069 | 0.51961932 | 0.61089729 | 0.99999785 | -6.2479361 | up |
| KI - WT, cell_origin: iPSC | Q15287 | RNPS1 | 5 | Homo sapien | -0.1980535 | -1.0110576 | 0.61495064 | 0.87467519 | -0.5189287 | 0.61133076 | 0.99999785 | -6.2483373 | down |
| KI - WT, cell_origin: iPSC | Q15814 | TBCC | 3 | Homo sapien | -0.4308814 | -2.2005057 | 1.33874297 | -1.8822754 | -0.5188944 | 0.61139059 | 0.99999785 | -6.2483194 | down |
| KI - WT, cell_origin: iPSC | P07602 | PSAP | 11 | Homo sapien | -0.2658336 | -1.3584898 | 0.82682256 | -0.8136546 | -0.5184761 | 0.61167536 | 0.99999785 | -6.2485403 | down |
| KI - WT, cell_origin: iPSC | Q9BXS6 | NUSAP1 | 4 | Homo sapien | 0.22862592 | -0.7116007 | 1.16885252 | -0.9904979 | 0.51805283 | 0.61193959 | 0.99999785 | -6.2487873 | up |
| KI - WT, cell_origin: iPSC | Q9UGI8 | TES | 9 | Homo sapien | 0.36536049 | -1.1406979 | 1.87141888 | -1.6632369 | 0.51698999 | 0.61268748 | 0.99999785 | -6.2493237 | up |
| KI - WT, cell_origin: iPSC | Q9BZR9 | TRIM8 | 6 | Homo sapien | -0.6115124 | -3.1309929 | 1.90796809 | -6.1323192 | -0.5169326 | 0.6126748 | 0.99999785 | -6.2494049 | down |
| KI - WT, cell_origin: iPSC | O15085 | ARHGEF11 | 55 | Homo sapien | 0.25104596 | -0.7854663 | 1.28755819 | 2.65484072 | 0.51615624 | 0.61325568 | 0.99999785 | -6.2497623 | up |

| | | | | | | | | | | | | | |
|---|---|---|---|---|---|---|---|---|---|---|---|---|---|
| KI - WT, cell_origin: iPSC | P62495 | ETF1 | 3 | Homo sapien | -0.3290907 | -1.6879723 | 1.02979084 | -1.8639612 | -0.516103 | 0.61329196 | 0.99999785 | -6.2497903 | down |
| KI - WT, cell_origin: iPSC | P52566 | ARHGDIB | 3 | Homo sapien | -0.5368925 | -2.7551692 | 1.68138419 | -1.0642935 | -0.5157913 | 0.61350448 | 0.99999785 | -6.2499541 | down |
| KI - WT, cell_origin: iPSC | Q02952 | AKAP12 | 83 | Homo sapien | 0.30602817 | -0.9593312 | 1.5713875 | 2.16814069 | 0.51540651 | 0.61376684 | 0.99999785 | -6.2501561 | up |
| KI - WT, cell_origin: iPSC | Q9Y3I0 | RTCB | 24 | Homo sapien | -0.2135502 | -1.0965424 | 0.66944205 | 3.86104642 | -0.5154012 | 0.61377048 | 0.99999785 | -6.2501589 | down |
| KI - WT, cell_origin: iPSC | Q9NVM9 | INTS13 | 10 | Homo sapien | 0.23657555 | -0.7428892 | 1.21604031 | -0.9383971 | 0.51465955 | 0.61426389 | 0.99999785 | -6.2505601 | up |
| KI - WT, cell_origin: iPSC | Q14195 | DPYSL3 | 18 | Homo sapien | -0.1812557 | -0.932774 | 0.57026264 | 1.06880184 | -0.5137452 | 0.61485982 | 0.99999785 | -6.2510665 | down |
| KI - WT, cell_origin: iPSC | P62258 | YWHAE | 11 | Homo sapien | 0.12126175 | -0.3816016 | 0.62412506 | 2.08530228 | 0.51358648 | 0.61495719 | 0.99999785 | -6.2511602 | up |
| KI - WT, cell_origin: iPSC | Q9Y3Z3 | SAMHD1 | 31 | Homo sapien | -0.2170975 | -1.1179768 | 0.68378187 | 1.61102419 | -0.5134949 | 0.61506043 | 0.99999785 | -6.2511681 | down |
| KI - WT, cell_origin: iPSC | Q9BVI4 | NOC4L | 8 | Homo sapien | 0.35618704 | -1.1241766 | 1.83655066 | -2.699079 | 0.51265674 | 0.6156267 | 0.99999785 | -6.251612 | up |
| KI - WT, cell_origin: iPSC | Q99873 | PRMT1 | 10 | Homo sapien | -0.2061983 | -1.0637875 | 0.65139099 | 1.44634433 | -0.5122573 | 0.61589283 | 0.99999785 | -6.2518272 | down |
| KI - WT, cell_origin: iPSC | Q9ULX3 | NOB1 | 17 | Homo sapien | 0.17026574 | -0.538595 | 0.87912645 | 0.35687548 | 0.51157117 | 0.61633381 | 0.99999785 | -6.2522121 | up |
| KI - WT, cell_origin: iPSC | Q8N474 | SFRP1 | 6 | Homo sapien | -0.3101389 | -1.6050058 | 0.98472789 | -1.6448292 | -0.5104269 | 0.61716711 | 0.99999785 | -6.2527576 | down |
| KI - WT, cell_origin: iPSC | P15104 | GLUL | 7 | Homo sapien | 0.38087311 | -1.2108206 | 1.97256685 | -0.1875312 | 0.5099448 | 0.6174968 | 0.99999785 | -6.2530082 | up |
| KI - WT, cell_origin: iPSC | Q13201 | MMRN1 | 28 | Homo sapien | -0.7191951 | -3.7247601 | 2.28636996 | -0.9051369 | -0.5099445 | 0.61749697 | 0.99999785 | -6.2530083 | down |
| KI - WT, cell_origin: iPSC | Q99832 | CCT7 | 19 | Homo sapien | -0.1868669 | -0.9681729 | 0.59443915 | 0.21966309 | -0.5094692 | 0.61778418 | 0.99999785 | -6.2532922 | down |
| KI - WT, cell_origin: iPSC | Q13263 | TRIM28 | 27 | Homo sapien | -0.2235379 | -1.1595101 | 0.71243435 | 3.94018457 | -0.5089681 | 0.61816499 | 0.99999785 | -6.2535152 | down |
| KI - WT, cell_origin: iPSC | P25786 | PSMA1 | 9 | Homo sapien | 0.15843956 | -0.5047237 | 0.82160281 | -0.032821 | 0.50884189 | 0.61820049 | 0.99999785 | -6.2536303 | up |
| KI - WT, cell_origin: iPSC | O75676 | RPS6KA4 | 4 | Homo sapien | -0.2052403 | -1.0644953 | 0.65401465 | -1.0714886 | -0.5087922 | 0.61824613 | 0.99999785 | -6.2536447 | down |
| KI - WT, cell_origin: iPSC | Q9UBB9 | TFIP11 | 19 | Homo sapien | 0.25581681 | -0.815972 | 1.32760564 | -0.1071389 | 0.50865371 | 0.61838011 | 0.99999785 | -6.2536781 | up |
| KI - WT, cell_origin: iPSC | Q8NCJ5 | SPRYD3 | 6 | Homo sapien | -0.2585179 | -1.3416661 | 0.8246303 | -0.4157794 | -0.5086337 | 0.61839382 | 0.99999785 | -6.2536885 | down |
| KI - WT, cell_origin: iPSC | Q92572 | AP3S1 | 5 | Homo sapien | 0.20403393 | -0.6521816 | 1.06024947 | 0.15599557 | 0.50765359 | 0.61903502 | 0.99999785 | -6.254225 | up |
| KI - WT, cell_origin: iPSC | P08579 | SNRPB2 | 5 | Homo sapien | -0.2361895 | -1.2279019 | 0.7555228 | 1.0546741 | -0.5075482 | 0.61913693 | 0.99999785 | -6.2542504 | down |
| KI - WT, cell_origin: iPSC | Q9NYF8 | BCLAF1 | 33 | Homo sapien | -0.2812981 | -1.4624437 | 0.89984755 | 2.77471915 | -0.5075347 | 0.61914622 | 0.99999785 | -6.2542575 | down |
| KI - WT, cell_origin: iPSC | P04350 | TUBB4A | 2 | Homo sapien | -0.1451135 | -0.7543308 | 0.46410376 | -0.4436847 | -0.5073123 | 0.61924787 | 0.99999785 | -6.2544219 | down |
| KI - WT, cell_origin: iPSC | Q15796 | SMAD2 | 7 | Homo sapien | -0.1534757 | -0.798789 | 0.49183755 | -0.9879589 | -0.5065342 | 0.61978096 | 0.99999785 | -6.2548237 | down |
| KI - WT, cell_origin: iPSC | Q9BVJ7 | DUSP23 | 8 | Homo sapien | -0.3306602 | -1.7230354 | 1.06171496 | 0.10589665 | -0.5060904 | 0.62013567 | 0.99999785 | -6.2550034 | down |
| KI - WT, cell_origin: iPSC | Q9UI26 | IPO11 | 7 | Homo sapien | -0.2076942 | -1.0831378 | 0.66774953 | 1.01985277 | -0.5055081 | 0.62052125 | 0.99999785 | -6.2553166 | down |
| KI - WT, cell_origin: iPSC | Q5VTL8 | PRPF38B | 16 | Homo sapien | -0.4063844 | -2.1201086 | 1.30733982 | 2.49953333 | -0.5053572 | 0.62063822 | 0.99999785 | -6.2553812 | down |
| KI - WT, cell_origin: iPSC | Q6Y7W6 | GIGYF2 | 36 | Homo sapien | 0.31489709 | -1.0132953 | 1.64308952 | 1.93413743 | 0.5052543 | 0.62070878 | 0.99999785 | -6.2554342 | up |
| KI - WT, cell_origin: iPSC | Q9BVA1 | TUBB2B | 5 | Homo sapien | -0.138091 | -0.7204909 | 0.44430896 | 5.9437163 | -0.505082 | 0.6207915 | 0.99999785 | -6.2555573 | down |
| KI - WT, cell_origin: iPSC | Q71UM5 | RPS27L | 3 | Homo sapien | -0.1453097 | -0.7582076 | 0.46758818 | 4.7776754 | -0.5050255 | 0.62082811 | 0.99999785 | -6.2555885 | down |
| KI - WT, cell_origin: iPSC | Q13416 | ORC2 | 5 | Homo sapien | 0.2535233 | -0.8163084 | 1.32335504 | -2.1500257 | 0.50474534 | 0.62101322 | 0.99999785 | -6.2557395 | up |
| KI - WT, cell_origin: iPSC | Q75QN2 | INTS8 | 3 | Homo sapien | 0.25533293 | -0.8239654 | 1.33463127 | -1.9234569 | 0.50393222 | 0.62157814 | 0.99999785 | -6.2561505 | up |
| KI - WT, cell_origin: iPSC | O60563 | CCNT1 | 8 | Homo sapien | 0.1810954 | -0.5845778 | 0.94676857 | -1.2692535 | 0.50373678 | 0.62169941 | 0.99999785 | -6.2562633 | up |
| KI - WT, cell_origin: iPSC | Q9NQY0 | BIN3 | 7 | Homo sapien | -0.2742552 | -1.4357045 | 0.88719416 | -0.5692596 | -0.5032188 | 0.62210511 | 0.99999785 | -6.2564802 | down |
| KI - WT, cell_origin: iPSC | Q5T7N2 | L1TD1 | 34 | Homo sapien | -0.3653976 | -1.9129228 | 1.18212754 | 0.01384921 | -0.5031881 | 0.62212621 | 0.99999785 | -6.256496 | down |
| KI - WT, cell_origin: iPSC | Q9UN81 | L1RE1 | 15 | Homo sapien | -0.2747054 | -1.438704 | 0.88929308 | -0.5437593 | -0.5029412 | 0.6222957 | 0.99999785 | -6.2566226 | down |
| KI - WT, cell_origin: iPSC | Q9NRR4 | DROSHA | 2 | Homo sapien | 0.30018239 | -0.972103 | 1.5724678 | -2.0729323 | 0.50272763 | 0.62242891 | 0.99999785 | -6.2567451 | up |
| KI - WT, cell_origin: iPSC | P29474 | NOS3 | 41 | Homo sapien | 0.30864604 | -1.0046747 | 1.62196675 | -0.00576 | 0.50083225 | 0.62374425 | 0.99999785 | -6.2577015 | up |
| KI - WT, cell_origin: iPSC | Q8IYB3 | SRRM1 | 10 | Homo sapien | -0.3336407 | -1.7536653 | 1.08638397 | 0.96055486 | -0.5007091 | 0.6238289 | 0.99999785 | -6.2577644 | down |
| KI - WT, cell_origin: iPSC | P67775 | PPP2CA | 13 | Homo sapien | 0.11327409 | -0.3686962 | 0.59524441 | 1.16174754 | 0.50055287 | 0.62388637 | 0.99999785 | -6.2578924 | up |
| KI - WT, cell_origin: iPSC | Q96D46 | NMD3 | 8 | Homo sapien | 0.18843249 | -0.6133648 | 0.99022981 | -1.4396186 | 0.50053087 | 0.6239015 | 0.99999785 | -6.2579037 | up |
| KI - WT, cell_origin: iPSC | P57678 | GEMIN4 | 22 | Homo sapien | -0.2643162 | -1.3899625 | 0.86133023 | -0.6149279 | -0.5004078 | 0.62403597 | 0.99999785 | -6.2579181 | down |
| KI - WT, cell_origin: iPSC | Q01081 | U2AF1 | 10 | Homo sapien | -0.4466887 | -2.3562139 | 1.46283637 | 1.62254383 | -0.4985193 | 0.62533475 | 0.99999785 | -6.2588797 | down |
| KI - WT, cell_origin: iPSC | Q9Y6B6 | SAR1B | 10 | Homo sapien | 0.25797302 | -0.8449318 | 1.36087785 | 5.28163249 | 0.49846952 | 0.62536902 | 0.99999785 | -6.258905 | up |
| KI - WT, cell_origin: iPSC | Q96C57 | CUSTOS | 2 | Homo sapien | -0.2408738 | -1.2705638 | 0.78881623 | -1.934042 | -0.4982723 | 0.62546342 | 0.99999785 | -6.2590452 | down |

| | | | | | | | | | | | | | |
|---|---|---|---|---|---|---|---|---|---|---|---|---|---|
| KI - WT, cell_origin: iPSC | P0DP23;P0DI | CALM1;CALM | 7 | Homo sapien | 0.24292819 | -0.796812 | 1.28266839 | 1.77090349 | 0.49791526 | 0.62575046 | 0.99999785 | -6.2591866 | up |
| KI - WT, cell_origin: iPSC | P15291 | B4GALT1 | 2 | Homo sapien | 0.3277102 | -1.0762201 | 1.73164053 | -2.351186 | 0.49738895 | 0.62610322 | 0.99999785 | -6.2594629 | up |
| KI - WT, cell_origin: iPSC | O75044 | SRGAP2 | 8 | Homo sapien | -0.1525925 | -0.8061814 | 0.50099633 | -1.3203762 | -0.4972426 | 0.626164 | 0.99999785 | -6.2595757 | down |
| KI - WT, cell_origin: iPSC | P06733 | ENO1 | 19 | Homo sapien | -0.2671699 | -1.4123909 | 0.8780511 | 3.23323337 | -0.4971651 | 0.6262669 | 0.99999785 | -6.2595671 | down |
| KI - WT, cell_origin: iPSC | Q9BYW2 | SETD2 | 10 | Homo sapien | -0.2555539 | -1.3515503 | 0.84044246 | -1.675313 | -0.4967638 | 0.62651955 | 0.99999785 | -6.2597934 | down |
| KI - WT, cell_origin: iPSC | A6NMB9 | FIGNL2 | 7 | Homo sapien | 0.35195421 | -1.1591162 | 1.86302465 | -1.4230969 | 0.49636806 | 0.62681581 | 0.99999785 | -6.2599709 | up |
| KI - WT, cell_origin: iPSC | Q9H2G2 | SLK | 43 | Homo sapien | 0.33508835 | -1.1084418 | 1.77861848 | 1.13963465 | 0.49469309 | 0.62797011 | 0.99999785 | -6.2608173 | up |
| KI - WT, cell_origin: iPSC | P61247 | RPS3A | 24 | Homo sapien | -0.0921831 | -0.4892569 | 0.30489061 | 7.70899698 | -0.4944472 | 0.62809044 | 0.99999785 | -6.2609887 | down |
| KI - WT, cell_origin: iPSC | O95352 | ATG7 | 4 | Homo sapien | -0.2517839 | -1.3373501 | 0.83378224 | -1.1555273 | -0.4942812 | 0.62825411 | 0.99999785 | -6.261025 | down |
| KI - WT, cell_origin: iPSC | O75592 | MYCBP2 | 87 | Homo sapien | -0.2478742 | -1.3185622 | 0.82281393 | 0.87084507 | -0.4933676 | 0.62888426 | 0.99999785 | -6.2614851 | down |
| KI - WT, cell_origin: iPSC | Q8WYA0 | IFT81 | 11 | Homo sapien | -0.1197979 | -0.6385297 | 0.39893387 | -0.91666 | -0.4918652 | 0.62987235 | 0.99999785 | -6.262287 | down |
| KI - WT, cell_origin: iPSC | Q9H3S7 | PTPN23 | 22 | Homo sapien | -0.155953 | -0.8352545 | 0.52334853 | -0.9547347 | -0.4889571 | 0.63188206 | 0.99999785 | -6.2637413 | down |
| KI - WT, cell_origin: iPSC | Q6P996 | PDXDC1 | 18 | Homo sapien | 0.25337649 | -0.8516262 | 1.35837913 | 1.22838008 | 0.48865839 | 0.63213722 | 0.99999785 | -6.2638438 | up |
| KI - WT, cell_origin: iPSC | Q13613 | MTMR1 | 4 | Homo sapien | 0.19130206 | -0.6427421 | 1.02534623 | 0.40392154 | 0.4885944 | 0.63214739 | 0.99999785 | -6.2639083 | up |
| KI - WT, cell_origin: iPSC | O00429 | DNM1L | 13 | Homo sapien | 0.37969123 | -1.2778984 | 2.03728084 | -1.647911 | 0.48815295 | 0.63248683 | 0.99999785 | -6.2640957 | up |
| KI - WT, cell_origin: iPSC | Q12800 | TFCP2 | 4 | Homo sapien | -0.1987744 | -1.0669461 | 0.66939736 | -0.4091769 | -0.4877224 | 0.63275071 | 0.99999785 | -6.2643425 | down |
| KI - WT, cell_origin: iPSC | O00459 | PIK3R2 | 4 | Homo sapien | -0.1836688 | -0.9858505 | 0.61851294 | -2.4460028 | -0.4876433 | 0.63279099 | 0.99999785 | -6.2643956 | down |
| KI - WT, cell_origin: iPSC | P40616 | ARL1 | 9 | Homo sapien | 0.21670734 | -0.7304195 | 1.16383417 | 2.1158488 | 0.48760468 | 0.63286616 | 0.99999785 | -6.2643686 | up |
| KI - WT, cell_origin: iPSC | Q96T51 | RUFY1 | 7 | Homo sapien | -0.1907754 | -1.0252336 | 0.64368281 | 1.96381996 | -0.4870415 | 0.63322754 | 0.99999785 | -6.2646757 | down |
| KI - WT, cell_origin: iPSC | Q8N1G4 | LRRC47 | 18 | Homo sapien | -0.2757866 | -1.4831701 | 0.93159682 | 1.05518624 | -0.4867773 | 0.63343877 | 0.99999785 | -6.2647798 | down |
| KI - WT, cell_origin: iPSC | P31327 | CPS1 | 3 | Homo sapien | 0.30104566 | -1.0175591 | 1.61965041 | 0.0324992 | 0.48654173 | 0.63360189 | 0.99999785 | -6.2648968 | up |
| KI - WT, cell_origin: iPSC | Q9NZJ0 | DTL | 2 | Homo sapien | -0.1896162 | -1.0199629 | 0.6407305 | -2.6692019 | -0.4863575 | 0.63368115 | 0.99999785 | -6.2650344 | down |
| KI - WT, cell_origin: iPSC | Q8N1N4 | KRT78 | 5 | Homo sapien | -0.179325 | -0.9650258 | 0.60637584 | -0.0440549 | -0.4861647 | 0.63382573 | 0.99999785 | -6.2651194 | down |
| KI - WT, cell_origin: iPSC | Q27J81 | INF2 | 21 | Homo sapien | 0.23477113 | -0.7961015 | 1.26564374 | -0.353841 | 0.4853355 | 0.63443728 | 0.99999785 | -6.265495 | up |
| KI - WT, cell_origin: iPSC | Q14498 | RBM39 | 19 | Homo sapien | -0.4307825 | -2.3225351 | 1.4609702 | 4.05192945 | -0.4852841 | 0.63447291 | 0.99999785 | -6.2655204 | down |
| KI - WT, cell_origin: iPSC | Q9BVK6 | TMED9 | 7 | Homo sapien | 0.50879551 | -1.7273228 | 2.74491386 | 1.9155504 | 0.48489851 | 0.63474005 | 0.99999785 | -6.2657113 | up |
| KI - WT, cell_origin: iPSC | P51530 | DNA2 | 4 | Homo sapien | 0.33919732 | -1.1521829 | 1.83057752 | -2.3671847 | 0.48469262 | 0.63488272 | 0.99999785 | -6.2658132 | up |
| KI - WT, cell_origin: iPSC | P51570 | GALK1 | 3 | Homo sapien | 0.32464261 | -1.103006 | 1.75229118 | -0.0566507 | 0.48460354 | 0.63494446 | 0.99999785 | -6.2658572 | up |
| KI - WT, cell_origin: iPSC | Q9BVC4 | MLST8 | 4 | Homo sapien | 0.19102079 | -0.6488184 | 1.03085994 | -0.6312376 | 0.48450964 | 0.63497578 | 0.99999785 | -6.2659358 | up |
| KI - WT, cell_origin: iPSC | Q96EB1 | ELP4 | 2 | Homo sapien | 0.13430455 | -0.4563734 | 0.72498255 | -0.9566857 | 0.48426113 | 0.63513373 | 0.99999785 | -6.2660722 | up |
| KI - WT, cell_origin: iPSC | Q8N8R5 | C2orf69 | 2 | Homo sapien | 0.15861074 | -0.5391344 | 0.85635587 | -1.5886071 | 0.48414499 | 0.63521425 | 0.99999785 | -6.2661296 | up |
| KI - WT, cell_origin: iPSC | P62195 | PSMC5 | 6 | Homo sapien | 0.21099957 | -0.7174309 | 1.13943004 | -1.1819597 | 0.48411672 | 0.63524816 | 0.99999785 | -6.2661299 | up |
| KI - WT, cell_origin: iPSC | Q13642 | FHL1 | 3 | Homo sapien | -0.2036036 | -1.1005479 | 0.69334079 | -0.4020507 | -0.4836179 | 0.63560584 | 0.99999785 | -6.266365 | down |
| KI - WT, cell_origin: iPSC | Q15649 | ZNHIT3 | 2 | Homo sapien | -0.1639184 | -0.8863833 | 0.55854642 | -0.1137646 | -0.4832265 | 0.63585121 | 0.99999785 | -6.2665829 | down |
| KI - WT, cell_origin: iPSC | Q9HAS0 | C17orf75 | 6 | Homo sapien | 0.13955209 | -0.4760377 | 0.75514186 | -0.3824986 | 0.4828193 | 0.63613367 | 0.99999785 | -6.2667835 | up |
| KI - WT, cell_origin: iPSC | P51812 | RPS6KA3 | 4 | Homo sapien | -0.2449491 | -1.3268447 | 0.83694643 | -1.1503064 | -0.4824951 | 0.63640643 | 0.99999785 | -6.2668977 | down |
| KI - WT, cell_origin: iPSC | P54727 | RAD23B | 5 | Homo sapien | -0.1897067 | -1.027335 | 0.64792166 | -0.8411338 | -0.4824342 | 0.63641311 | 0.99999785 | -6.2669616 | down |
| KI - WT, cell_origin: iPSC | O95613 | PCNT | 3 | Homo sapien | -0.56662 | -3.069991 | 1.93675097 | -2.6061748 | -0.4823575 | 0.6365019 | 0.99999785 | -6.2669655 | down |
| KI - WT, cell_origin: iPSC | P0C0L4;P0C0 | C4A;C4B | 2 | Homo sapien | 0.29780061 | -1.0197005 | 1.61530168 | -1.279401 | 0.48170037 | 0.63695793 | 0.99999785 | -6.2672888 | up |
| KI - WT, cell_origin: iPSC | P49755 | TMED10 | 9 | Homo sapien | 0.51487087 | -1.7639283 | 2.79367 | 1.55486194 | 0.48149818 | 0.63709827 | 0.99999785 | -6.2673882 | up |
| KI - WT, cell_origin: iPSC | Q6P3W7 | SCYL2 | 10 | Homo sapien | 0.58508167 | -2.0044892 | 3.17465254 | -0.1268439 | 0.4814942 | 0.63710103 | 0.99999785 | -6.2673901 | up |
| KI - WT, cell_origin: iPSC | Q8N8S7 | ENAH | 8 | Homo sapien | 0.18584091 | -0.636336 | 1.00801786 | -1.5761802 | 0.48141067 | 0.63711129 | 0.99999785 | -6.2674765 | up |
| KI - WT, cell_origin: iPSC | O43707 | ACTN4 | 16 | Homo sapien | 0.23638143 | -0.8103117 | 1.28307456 | -0.7576673 | 0.48127837 | 0.63725086 | 0.99999785 | -6.2674962 | up |
| KI - WT, cell_origin: iPSC | O43900 | PRICKLE3 | 2 | Homo sapien | -0.4383821 | -2.3816822 | 1.50491814 | -1.0805162 | -0.4807455 | 0.63762081 | 0.99999785 | -6.2677578 | down |
| KI - WT, cell_origin: iPSC | Q9NV06 | DCAF13 | 4 | Homo sapien | 0.28917811 | -0.9952905 | 1.57364673 | -1.146118 | 0.47978238 | 0.63828977 | 0.99999785 | -6.26823 | up |
| KI - WT, cell_origin: iPSC | Q969X5 | ERGIC1 | 8 | Homo sapien | -0.5640294 | -3.0693449 | 1.94128615 | 0.1026152 | -0.4797795 | 0.63829179 | 0.99999785 | -6.2682314 | down |

| | | | | | | | | | | | | |
|---|---|---|---|---|---|---|---|---|---|---|---|---|
| KI - WT, cell_origin: iPSC | O15287 | FANCG | 7 | Homo sapien | 0.208978 | -0.718931 | 1.13688697 | -1.2057631 | 0.47974792 | 0.63828033 | 0.99999785 | -6.2682785 up |
| KI - WT, cell_origin: iPSC | P19623 | SRM | 4 | Homo sapien | -0.4447808 | -2.4205676 | 1.53100608 | -1.7383864 | -0.4797426 | 0.6383174 | 0.99999785 | -6.2682495 down |
| KI - WT, cell_origin: iPSC | Q9NUV9 | GIMAP4 | 9 | Homo sapien | -0.707666 | -3.8517109 | 2.43637883 | -1.5095318 | -0.4796694 | 0.63836828 | 0.99999785 | -6.2682853 down |
| KI - WT, cell_origin: iPSC | O95232 | LUC7L3 | 18 | Homo sapien | -0.4007747 | -2.1840619 | 1.3825125 | 2.90145201 | -0.4789403 | 0.63887492 | 0.99999785 | -6.2686421 down |
| KI - WT, cell_origin: iPSC | Q9NZC9 | SMARCAL1 | 2 | Homo sapien | -0.457235 | -2.4934141 | 1.57894399 | -2.3596186 | -0.4785484 | 0.63914728 | 0.99999785 | -6.2688336 down |
| KI - WT, cell_origin: iPSC | Q13596 | SNX1 | 14 | Homo sapien | 0.13839439 | -0.4778841 | 0.7546729 | 1.46668734 | 0.47827883 | 0.63928735 | 0.99999785 | -6.26901 up |
| KI - WT, cell_origin: iPSC | P00973 | OAS1 | 7 | Homo sapien | 0.72270876 | -2.49753 | 3.94294753 | -1.0363947 | 0.47827493 | 0.63933743 | 0.99999785 | -6.2689672 up |
| KI - WT, cell_origin: iPSC | Q14011 | CIRBP | 3 | Homo sapien | -0.1737756 | -0.9483303 | 0.60077912 | 1.2050638 | -0.4779071 | 0.63955796 | 0.99999785 | -6.26918 down |
| KI - WT, cell_origin: iPSC | O60231 | DHX16 | 10 | Homo sapien | -0.3103861 | -1.6962797 | 1.07550764 | -1.4859077 | -0.4772816 | 0.64002825 | 0.99999785 | -6.2694517 down |
| KI - WT, cell_origin: iPSC | Q8NFH5 | NUP35 | 2 | Homo sapien | 0.26920813 | -0.9352387 | 1.4736549 | -1.7987721 | 0.47626488 | 0.64072587 | 0.99999785 | -6.2699558 up |
| KI - WT, cell_origin: iPSC | Q9BTE7 | DCUN1D5 | 5 | Homo sapien | -0.2545221 | -1.3937109 | 0.88466663 | -1.2333758 | -0.4761373 | 0.64082442 | 0.99999785 | -6.2700087 down |
| KI - WT, cell_origin: iPSC | Q8TAQ2 | SMARCC2 | 6 | Homo sapien | 0.31051606 | -1.0824168 | 1.70344897 | -1.3505843 | 0.4750685 | 0.6415685 | 0.99999785 | -6.2705277 up |
| KI - WT, cell_origin: iPSC | Q96G25 | MED8 | 4 | Homo sapien | 0.15061392 | -0.5249876 | 0.8262154 | -1.2559725 | 0.47480381 | 0.64170586 | 0.99999785 | -6.2707003 up |
| KI - WT, cell_origin: iPSC | Q9H6R4 | NOL6 | 5 | Homo sapien | 0.28047739 | -0.9784902 | 1.53944494 | -2.6200903 | 0.47454675 | 0.64189494 | 0.99999785 | -6.2708154 up |
| KI - WT, cell_origin: iPSC | Q16644 | MAPKAPK3 | 3 | Homo sapien | -0.2400239 | -1.3188058 | 0.83875804 | -1.0354318 | -0.4741581 | 0.64220263 | 0.99999785 | -6.2709689 down |
| KI - WT, cell_origin: iPSC | Q9UJY5 | GGA1 | 4 | Homo sapien | -0.4228871 | -2.3265749 | 1.48080069 | -1.4813323 | -0.4734031 | 0.64272875 | 0.99999785 | -6.2713343 down |
| KI - WT, cell_origin: iPSC | Q5U5Q3 | MEX3C | 6 | Homo sapien | 0.16600357 | -0.5815263 | 0.91353345 | -0.3712257 | 0.472971 | 0.64298419 | 0.99999785 | -6.271586 up |
| KI - WT, cell_origin: iPSC | O75083 | WDR1 | 5 | Homo sapien | -0.2204603 | -1.2142067 | 0.77328611 | -1.0204454 | -0.472709 | 0.64320131 | 0.99999785 | -6.2716802 down |
| KI - WT, cell_origin: iPSC | Q96MU7 | YTHDC1 | 9 | Homo sapien | 0.26064942 | -0.9140155 | 1.43531434 | -2.4916986 | 0.47262065 | 0.6432328 | 0.99999785 | -6.271751 up |
| KI - WT, cell_origin: iPSC | P40121 | CAPG | 7 | Homo sapien | 0.38688622 | -1.3585456 | 2.13231805 | -0.3493514 | 0.47237047 | 0.6434486 | 0.99999785 | -6.2718329 up |
| KI - WT, cell_origin: iPSC | Q9NWU2 | GID8 | 6 | Homo sapien | 0.14643543 | -0.5147063 | 0.80757716 | -0.2706579 | 0.47172758 | 0.64385032 | 0.99999785 | -6.2721866 up |
| KI - WT, cell_origin: iPSC | P80217 | IFI35 | 8 | Homo sapien | -0.3141852 | -1.7350455 | 1.10667522 | -2.1170839 | -0.471234 | 0.64424128 | 0.99999785 | -6.2723806 down |
| KI - WT, cell_origin: iPSC | Q05655 | PRKCD | 7 | Homo sapien | 0.41392507 | -1.4588813 | 2.28673141 | -1.5195145 | 0.47101119 | 0.64439675 | 0.99999785 | -6.2724878 up |
| KI - WT, cell_origin: iPSC | P82909 | MRPS36 | 2 | Homo sapien | -0.5687088 | -3.1449018 | 2.00748413 | -2.0344525 | -0.4704505 | 0.64478804 | 0.99999785 | -6.2727574 down |
| KI - WT, cell_origin: iPSC | Q14669 | TRIP12 | 21 | Homo sapien | 0.18863208 | -0.6657411 | 1.04300521 | -0.3151639 | 0.4703118 | 0.64485222 | 0.99999785 | -6.2728546 up |
| KI - WT, cell_origin: iPSC | P10588 | NR2F6 | 3 | Homo sapien | 0.25824586 | -0.9128196 | 1.42931136 | -3.4924369 | 0.46966896 | 0.64528723 | 0.99999785 | -6.273176 up |
| KI - WT, cell_origin: iPSC | Q9NY15 | STAB1 | 14 | Homo sapien | -0.6376278 | -3.5317257 | 2.25647005 | -1.7152979 | -0.4695225 | 0.64543591 | 0.99999785 | -6.2732029 down |
| KI - WT, cell_origin: iPSC | Q9GZR7 | DDX24 | 10 | Homo sapien | -0.2294874 | -1.2723988 | 0.81342399 | -0.3731035 | -0.4689363 | 0.64584532 | 0.99999785 | -6.2734839 down |
| KI - WT, cell_origin: iPSC | Q8WVM8 | SCFD1 | 12 | Homo sapien | 0.47727905 | -1.6947434 | 2.64930149 | -0.7782903 | 0.4682852 | 0.64630016 | 0.99999785 | -6.2737955 up |
| KI - WT, cell_origin: iPSC | Q15417 | CNN3 | 17 | Homo sapien | -0.2558426 | -1.4210488 | 0.90936361 | 2.27869784 | -0.4679209 | 0.64655472 | 0.99999785 | -6.2739698 down |
| KI - WT, cell_origin: iPSC | Q69YQ0 | SPECC1L | 2 | Homo sapien | -0.2262536 | -1.2582842 | 0.80577697 | -1.515187 | -0.4670043 | 0.64716305 | 0.99999785 | -6.2744377 down |
| KI - WT, cell_origin: iPSC | O00182 | LGALS9 | 5 | Homo sapien | 0.34232651 | -1.2206315 | 1.90528456 | -0.0979956 | 0.4667621 | 0.64736478 | 0.99999785 | -6.274523 up |
| KI - WT, cell_origin: iPSC | Q8IX18 | DHX40 | 6 | Homo sapien | -0.3277886 | -1.8238387 | 1.16826151 | -2.9413351 | -0.4667299 | 0.64735496 | 0.99999785 | -6.2745685 down |
| KI - WT, cell_origin: iPSC | P63173 | RPL38 | 3 | Homo sapien | -0.2415637 | -1.3450697 | 0.86194239 | 6.94500124 | -0.4665082 | 0.64754233 | 0.99999785 | -6.2746441 down |
| KI - WT, cell_origin: iPSC | O60573 | EIF4E2 | 9 | Homo sapien | 0.29407873 | -1.0508205 | 1.63897795 | 2.09889324 | 0.46598962 | 0.64790503 | 0.99999785 | -6.2748911 up |
| KI - WT, cell_origin: iPSC | P36873 | PPP1CC | 3 | Homo sapien | 0.18293472 | -0.6539466 | 1.01981604 | 0.18464662 | 0.46563988 | 0.64811743 | 0.99999785 | -6.2750876 up |
| KI - WT, cell_origin: iPSC | O60504 | SORBS3 | 13 | Homo sapien | 0.25412505 | -0.9100994 | 1.41834945 | -0.5401205 | 0.46517154 | 0.6484774 | 0.99999785 | -6.2752803 up |
| KI - WT, cell_origin: iPSC | Q14739 | LBR | 3 | Homo sapien | 0.55644847 | -1.9976711 | 3.11056805 | -0.9419646 | 0.46428654 | 0.64909684 | 0.99999785 | -6.2757005 up |
| KI - WT, cell_origin: iPSC | Q8IWX8 | CHERP | 19 | Homo sapien | -0.1777395 | -0.9935933 | 0.63811429 | 0.75011815 | -0.4640765 | 0.64921176 | 0.99999785 | -6.27583 down |
| KI - WT, cell_origin: iPSC | Q9UBE8 | NLK | 3 | Homo sapien | 0.35904584 | -1.2901135 | 2.0082052 | -2.5187196 | 0.46396973 | 0.64931865 | 0.99999785 | -6.2758508 up |
| KI - WT, cell_origin: iPSC | Q93009 | USP7 | 36 | Homo sapien | 0.23435949 | -0.8449057 | 1.31362469 | 0.84662101 | 0.462761 | 0.65016525 | 0.99999785 | -6.2764231 up |
| KI - WT, cell_origin: iPSC | Q9P2E3 | ZNFX1 | 9 | Homo sapien | -0.2215929 | -1.2426427 | 0.79945695 | -1.5568806 | -0.462303 | 0.65045414 | 0.99999785 | -6.2766693 down |
| KI - WT, cell_origin: iPSC | Q9UBI9 | HECA | 6 | Homo sapien | 0.32520731 | -1.1747237 | 1.82513832 | -0.5533066 | 0.46205256 | 0.65066168 | 0.99999785 | -6.2767579 up |
| KI - WT, cell_origin: iPSC | Q9Y5B8 | NME7 | 4 | Homo sapien | 0.20970747 | -0.757469 | 1.17688391 | -0.7311629 | 0.46201513 | 0.65067848 | 0.99999785 | -6.2767843 up |
| KI - WT, cell_origin: iPSC | O95816 | BAG2 | 4 | Homo sapien | 0.18158519 | -0.6592349 | 1.02240523 | -0.8257769 | 0.46002789 | 0.65204762 | 0.99999785 | -6.2777431 up |
| KI - WT, cell_origin: iPSC | Q8N183 | NDUFAF2 | 9 | Homo sapien | 0.32670436 | -1.1890326 | 1.84244127 | 0.39793929 | 0.45933915 | 0.65256462 | 0.99999785 | -6.2780355 up |

| | | | | | | | | | | | | | |
|---|---|---|---|---|---|---|---|---|---|---|---|---|---|
| KI - WT, cell_origin: iPSC | Q96K17 | BTF3L4 | 2 | Homo sapien | -0.3715215 | -2.0952683 | 1.35222532 | -0.5091452 | -0.4593173 | 0.65257993 | 0.99999785 | -6.2780458 | down |
| KI - WT, cell_origin: iPSC | P26196 | DDX6 | 18 | Homo sapien | -0.1578642 | -0.890182 | 0.5744536 | 2.27797784 | -0.4591639 | 0.65264996 | 0.99999785 | -6.2781525 | down |
| KI - WT, cell_origin: iPSC | Q15363 | TMED2 | 7 | Homo sapien | 0.47064019 | -1.7138514 | 2.65513182 | 1.76524539 | 0.45913563 | 0.65270745 | 0.99999785 | -6.2781311 | up |
| KI - WT, cell_origin: iPSC | Q9UNX4 | WDR3 | 5 | Homo sapien | 0.13040861 | -0.4747929 | 0.73561017 | -1.6530669 | 0.45892944 | 0.65280696 | 0.99999785 | -6.2782695 | up |
| KI - WT, cell_origin: iPSC | P15735 | PHKG2 | 16 | Homo sapien | 0.1381678 | -0.5036974 | 0.78003298 | 0.93235928 | 0.45846134 | 0.65313561 | 0.99999785 | -6.2784889 | up |
| KI - WT, cell_origin: iPSC | Q13439 | GOLGA4 | 97 | Homo sapien | -0.1659544 | -0.9373193 | 0.60541038 | 1.19275612 | -0.4582852 | 0.65327083 | 0.99999785 | -6.2785608 | down |
| KI - WT, cell_origin: iPSC | A1L020 | MEX3A | 11 | Homo sapien | -0.1178694 | -0.6658854 | 0.43014653 | -0.0363951 | -0.4580867 | 0.65339869 | 0.99999785 | -6.2786644 | down |
| KI - WT, cell_origin: iPSC | Q5F1R6 | DNAJC21 | 5 | Homo sapien | 0.14271706 | -0.520952 | 0.80638612 | -0.4274048 | 0.45799843 | 0.6534607 | 0.99999785 | -6.2787057 | up |
| KI - WT, cell_origin: iPSC | Q6P5R6 | RPL22L1 | 3 | Homo sapien | -0.4052478 | -2.2970783 | 1.48658281 | 0.52759837 | -0.4565 | 0.65455844 | 0.99999785 | -6.2793646 | down |
| KI - WT, cell_origin: iPSC | Q9NTI5 | PDS5B | 6 | Homo sapien | -0.3199959 | -1.8164361 | 1.17644429 | -1.5290532 | -0.4557088 | 0.65511451 | 0.99999785 | -6.2797335 | down |
| KI - WT, cell_origin: iPSC | O75937 | DNAJC8 | 9 | Homo sapien | -0.2321897 | -1.3182486 | 0.85386923 | -0.4866011 | -0.4556086 | 0.65518496 | 0.99999785 | -6.2797802 | down |
| KI - WT, cell_origin: iPSC | Q15365 | PCBP1 | 10 | Homo sapien | -0.1369772 | -0.7784008 | 0.50444638 | 4.31488632 | -0.454894 | 0.65565412 | 0.99999785 | -6.2801433 | down |
| KI - WT, cell_origin: iPSC | P60174 | TPI1 | 13 | Homo sapien | -0.3103243 | -1.7669652 | 1.14631654 | 0.48970927 | -0.4540103 | 0.65630904 | 0.99999785 | -6.2805234 | down |
| KI - WT, cell_origin: iPSC | O95619 | YEATS4 | 8 | Homo sapien | -0.1050952 | -0.598289 | 0.38809861 | 0.57486788 | -0.4538422 | 0.65638269 | 0.99999785 | -6.2806423 | down |
| KI - WT, cell_origin: iPSC | P45974 | USP5 | 16 | Homo sapien | -0.1338302 | -0.7622004 | 0.49454007 | 0.34510582 | -0.4536053 | 0.65654942 | 0.99999785 | -6.2807522 | down |
| KI - WT, cell_origin: iPSC | P51858 | HDGF | 7 | Homo sapien | -0.1963768 | -1.1201498 | 0.72739608 | 0.39835278 | -0.4529972 | 0.65701659 | 0.99999785 | -6.2809981 | down |
| KI - WT, cell_origin: iPSC | P10909 | CLU | 2 | Homo sapien | -0.2307455 | -1.3165257 | 0.85503481 | -0.9336768 | -0.4528909 | 0.65709685 | 0.99999785 | -6.2810424 | down |
| KI - WT, cell_origin: iPSC | Q92541 | RTF1 | 17 | Homo sapien | -0.2080369 | -1.18733 | 0.77125623 | 0.38147224 | -0.4527204 | 0.65721689 | 0.99999785 | -6.2811214 | down |
| KI - WT, cell_origin: iPSC | Q8IWT6 | LRRC8A | 13 | Homo sapien | 0.27760228 | -1.0283989 | 1.58360351 | -2.9672806 | 0.45271522 | 0.65717701 | 0.99999785 | -6.2811636 | up |
| KI - WT, cell_origin: iPSC | Q8TEA1 | NSUN6 | 2 | Homo sapien | 0.47637357 | -1.7683096 | 2.72105674 | -3.7208547 | 0.45226708 | 0.65753604 | 0.99999785 | -6.2813311 | up |
| KI - WT, cell_origin: iPSC | Q96HR8 | NAF1 | 4 | Homo sapien | 0.16865446 | -0.6261933 | 0.96350223 | -0.5331614 | 0.45196995 | 0.65771033 | 0.99999785 | -6.2815004 | up |
| KI - WT, cell_origin: iPSC | P52732 | KIF11 | 19 | Homo sapien | 0.25421842 | -0.9445483 | 1.4529851 | -0.8179976 | 0.45193369 | 0.65777082 | 0.99999785 | -6.2814852 | up |
| KI - WT, cell_origin: iPSC | Q8NBR6 | MINDY2 | 5 | Homo sapien | -0.3074555 | -1.7582488 | 1.14333771 | -2.4092331 | -0.4516005 | 0.65800132 | 0.99999785 | -6.281643 | down |
| KI - WT, cell_origin: iPSC | Q9UGU5 | HMGXB4 | 4 | Homo sapien | -0.1171112 | -0.6696264 | 0.435404 | -0.2296202 | -0.4514336 | 0.65807867 | 0.99999785 | -6.2817567 | down |
| KI - WT, cell_origin: iPSC | Q86TB9 | PATL1 | 12 | Homo sapien | 0.17878111 | -0.6653936 | 1.02295583 | 0.27906646 | 0.45116487 | 0.65828598 | 0.99999785 | -6.2818644 | up |
| KI - WT, cell_origin: iPSC | Q5BJH7 | YIF1B | 4 | Homo sapien | -0.5816143 | -3.3292131 | 2.16598449 | -1.0021262 | -0.4511117 | 0.65834981 | 0.99999785 | -6.2818648 | down |
| KI - WT, cell_origin: iPSC | Q9Y5U9 | IER3IP1 | 2 | Homo sapien | -0.7206095 | -4.1266843 | 2.6854654 | -0.4656911 | -0.4508667 | 0.65852247 | 0.99999785 | -6.2819778 | down |
| KI - WT, cell_origin: iPSC | P02545 | LMNA | 15 | Homo sapien | 0.32219315 | -1.201055 | 1.84544129 | -0.286563 | 0.45076274 | 0.65859571 | 0.99999785 | -6.2820257 | up |
| KI - WT, cell_origin: iPSC | Q13151 | HNRNPA0 | 9 | Homo sapien | -0.2016317 | -1.1552687 | 0.75200523 | 3.37261137 | -0.4505865 | 0.65871991 | 0.99999785 | -6.2821069 | down |
| KI - WT, cell_origin: iPSC | Q8NI27 | THOC2 | 8 | Homo sapien | -0.260507 | -1.4926004 | 0.97158648 | 4.43626508 | -0.4505864 | 0.65872 | 0.99999785 | -6.282107 | down |
| KI - WT, cell_origin: iPSC | Q13509 | TUBB3 | 5 | Homo sapien | 0.21771808 | -0.8129068 | 1.24834298 | 4.11890176 | 0.4501904 | 0.65899907 | 0.99999785 | -6.2822894 | up |
| KI - WT, cell_origin: iPSC | Q6YHU6 | THADA | 12 | Homo sapien | 0.32319987 | -1.2067664 | 1.85316615 | -2.1421072 | 0.4501857 | 0.65900238 | 0.99999785 | -6.2822916 | up |
| KI - WT, cell_origin: iPSC | Q15059 | BRD3 | 5 | Homo sapien | -0.3995352 | -2.2941071 | 1.49503661 | -1.5587546 | -0.4494138 | 0.65954658 | 0.99999785 | -6.2826467 | down |
| KI - WT, cell_origin: iPSC | Q16630 | CPSF6 | 15 | Homo sapien | -0.4243776 | -2.4372052 | 1.58845009 | 4.04267155 | -0.4493122 | 0.65961821 | 0.99999785 | -6.2826934 | down |
| KI - WT, cell_origin: iPSC | Q86W42 | THOC6 | 5 | Homo sapien | 0.12784345 | -0.4795697 | 0.73525658 | -1.6035971 | 0.44826412 | 0.6603134 | 0.99999785 | -6.2832145 | up |
| KI - WT, cell_origin: iPSC | O14950;P191 | MYL12B;MYL | 5 | Homo sapien | -0.140111 | -0.8063263 | 0.52610424 | 0.96835236 | -0.4479169 | 0.66055841 | 0.99999785 | -6.2833736 | down |
| KI - WT, cell_origin: iPSC | Q9BSJ2 | TUBGCP2 | 30 | Homo sapien | 0.09150709 | -0.3439028 | 0.526917 | 1.55359317 | 0.44760625 | 0.66077767 | 0.99999785 | -6.2835158 | up |
| KI - WT, cell_origin: iPSC | Q1KMD3 | HNRNPUL2 | 16 | Homo sapien | -0.1170001 | -0.6737149 | 0.43971462 | 0.77089924 | -0.4476033 | 0.66077977 | 0.99999785 | -6.2835172 | down |
| KI - WT, cell_origin: iPSC | O43432 | EIF4G3 | 14 | Homo sapien | -0.170751 | -0.9836632 | 0.64216124 | -0.3200478 | -0.4474313 | 0.66091238 | 0.99999785 | -6.2835856 | down |
| KI - WT, cell_origin: iPSC | Q13547 | HDAC1 | 7 | Homo sapien | 0.15542716 | -0.5851081 | 0.89596245 | 9.6644738 | 0.44728417 | 0.66104893 | 0.99999785 | -6.2836233 | up |
| KI - WT, cell_origin: iPSC | Q14204 | DYNC1H1 | 108 | Homo sapien | 0.21960129 | -0.8275685 | 1.26677109 | 0.23002329 | 0.44691007 | 0.66131301 | 0.99999785 | -6.2837944 | up |
| KI - WT, cell_origin: iPSC | Q9H2K8 | TAOK3 | 5 | Homo sapien | 0.52351782 | -1.9744408 | 3.02147648 | -1.8264604 | 0.44663072 | 0.66151023 | 0.99999785 | -6.283922 | up |
| KI - WT, cell_origin: iPSC | Q13637 | RAB32 | 2 | Homo sapien | 0.19270383 | -0.7271456 | 1.11255326 | -1.5664703 | 0.44621435 | 0.66176547 | 0.99999785 | -6.2841473 | up |
| KI - WT, cell_origin: iPSC | P30101 | PDIA3 | 22 | Homo sapien | -0.402545 | -2.3319877 | 1.52689777 | 0.95989655 | -0.4446158 | 0.66293351 | 0.99999785 | -6.2848406 | down |
| KI - WT, cell_origin: iPSC | Q9HCE7 | SMURF1 | 2 | Homo sapien | 0.26588856 | -1.011229 | 1.54300616 | -1.5855884 | 0.44368131 | 0.66359405 | 0.99999785 | -6.2852652 | up |
| KI - WT, cell_origin: iPSC | P16383 | GCFC2 | 11 | Homo sapien | 0.3035313 | -1.1551823 | 1.76224494 | -1.1474815 | 0.44344096 | 0.66376399 | 0.99999785 | -6.2853743 | up |

| | | | | | | | | | | | | |
|---|---|---|---|---|---|---|---|---|---|---|---|---|
| KI - WT, cell_origin: iPSC | Q96P48 | ARAP1 | 17 | Homo sapien | -0.204161 | -1.1862562 | 0.7779341 | -0.1296342 | -0.4430183 | 0.66406288 | 0.99999785 | -6.285566 down |
| KI - WT, cell_origin: iPSC | Q9NVZ3 | NECAP2 | 2 | Homo sapien | 0.15254824 | -0.5812349 | 0.8863314 | -0.9483525 | 0.44277097 | 0.66419439 | 0.99999785 | -6.2857172 up |
| KI - WT, cell_origin: iPSC | Q15750 | TAB1 | 5 | Homo sapien | 0.22118944 | -0.8439956 | 1.28637449 | -0.2797637 | 0.44252895 | 0.664409 | 0.99999785 | -6.2857876 up |
| KI - WT, cell_origin: iPSC | P41229;Q9B\ | KDM5C;KDM | 2 | Homo sapien | -0.1514432 | -0.8805036 | 0.57761728 | -2.0227564 | -0.4424109 | 0.66444913 | 0.99999785 | -6.2858802 down |
| KI - WT, cell_origin: iPSC | Q16512 | PKN1 | 30 | Homo sapien | 0.30277416 | -1.1559837 | 1.76153198 | 2.51051777 | 0.44232143 | 0.66455581 | 0.99999785 | -6.2858816 up |
| KI - WT, cell_origin: iPSC | Q8N9R8 | SCAI | 2 | Homo sapien | 0.10034045 | -0.3831307 | 0.58381156 | -0.419604 | 0.44202334 | 0.66472337 | 0.99999785 | -6.2860555 up |
| KI - WT, cell_origin: iPSC | Q14157 | UBAP2L | 16 | Homo sapien | -0.1802478 | -1.0499113 | 0.68941565 | 2.01460239 | -0.4416102 | 0.66504552 | 0.99999785 | -6.2862154 down |
| KI - WT, cell_origin: iPSC | P08865 | RPSA | 14 | Homo sapien | 0.0986566 | -0.3777346 | 0.57504776 | 5.90683707 | 0.44106451 | 0.66540206 | 0.99999785 | -6.2864886 up |
| KI - WT, cell_origin: iPSC | Q9Y4C2 | TCAF1 | 8 | Homo sapien | 0.14482962 | -0.5546627 | 0.8443219 | -0.5777838 | 0.44097518 | 0.6654653 | 0.99999785 | -6.2865289 up |
| KI - WT, cell_origin: iPSC | Q8NB16 | MLKL | 11 | Homo sapien | 0.64188933 | -2.4621462 | 3.74592488 | -2.1518209 | 0.44069268 | 0.66570852 | 0.99999785 | -6.2866174 up |
| KI - WT, cell_origin: iPSC | Q8NF50 | DOCK8 | 16 | Homo sapien | -0.1580064 | -0.9219203 | 0.60590759 | -0.3706782 | -0.440555 | 0.66576783 | 0.99999785 | -6.2867139 down |
| KI - WT, cell_origin: iPSC | Q01201 | RELB | 8 | Homo sapien | 0.30377583 | -1.1694089 | 1.77696055 | -1.2268811 | 0.43943878 | 0.66659654 | 0.99999785 | -6.2871821 up |
| KI - WT, cell_origin: iPSC | Q9H299 | SH3BGRL3 | 3 | Homo sapien | 0.63642335 | -2.4508032 | 3.72364986 | -2.0236648 | 0.43931899 | 0.6666814 | 0.99999785 | -6.287236 up |
| KI - WT, cell_origin: iPSC | Q92828 | CORO2A | 7 | Homo sapien | -0.3053916 | -1.7912884 | 1.18050526 | -1.978508 | -0.4379966 | 0.66761851 | 0.99999785 | -6.2878296 down |
| KI - WT, cell_origin: iPSC | Q9BZJ0 | CRNKL1 | 6 | Homo sapien | 0.191346 | -0.7393969 | 1.1220889 | -2.260679 | 0.43785401 | 0.66767673 | 0.99999785 | -6.287932 up |
| KI - WT, cell_origin: iPSC | O14613 | CDC42EP2 | 5 | Homo sapien | 0.4191078 | -1.6248201 | 2.46303569 | -0.6942023 | 0.43698098 | 0.66833865 | 0.99999785 | -6.2882844 up |
| KI - WT, cell_origin: iPSC | P62753 | RPS6 | 11 | Homo sapien | -0.0874052 | -0.5137978 | 0.33898746 | 6.08810294 | -0.4365834 | 0.66857792 | 0.99999785 | -6.2885004 down |
| KI - WT, cell_origin: iPSC | Q15404 | RSU1 | 10 | Homo sapien | 0.23750615 | -0.9228549 | 1.39786716 | 0.05618028 | 0.43619844 | 0.66889374 | 0.99999785 | -6.2886341 up |
| KI - WT, cell_origin: iPSC | P46013 | MKI67 | 16 | Homo sapien | 0.11246856 | -0.436775 | 0.66171206 | -0.0306193 | 0.43611986 | 0.6689068 | 0.99999785 | -6.2887073 up |
| KI - WT, cell_origin: iPSC | Q15398 | DLGAP5 | 6 | Homo sapien | 0.15537354 | -0.6049946 | 0.91574165 | -1.374606 | 0.43520395 | 0.66955687 | 0.99999785 | -6.2891156 up |
| KI - WT, cell_origin: iPSC | P50479 | PDLIM4 | 4 | Homo sapien | -0.2665074 | -1.5716161 | 1.03860127 | -1.2100204 | -0.4351761 | 0.66961924 | 0.99999785 | -6.28909 down |
| KI - WT, cell_origin: iPSC | Q9NPE3 | NOP10 | 2 | Homo sapien | 0.11751631 | -0.4577683 | 0.6928009 | 0.18948207 | 0.43506595 | 0.66965484 | 0.99999785 | -6.2891771 up |
| KI - WT, cell_origin: iPSC | O95057 | DIRAS1 | 4 | Homo sapien | -0.6523375 | -3.8491783 | 2.54450328 | -2.8314008 | -0.4348643 | 0.66984058 | 0.99999785 | -6.2892289 down |
| KI - WT, cell_origin: iPSC | Q07021 | C1QBP | 3 | Homo sapien | 0.38796548 | -1.513649 | 2.28957994 | -1.0736916 | 0.43478343 | 0.66989797 | 0.99999785 | -6.2892649 up |
| KI - WT, cell_origin: iPSC | P61960 | UFM1 | 3 | Homo sapien | 0.25360252 | -0.9903874 | 1.49759241 | 0.54842721 | 0.43444928 | 0.6701352 | 0.99999785 | -6.2894135 up |
| KI - WT, cell_origin: iPSC | Q96IZ7 | RSRC1 | 4 | Homo sapien | -0.2977987 | -1.7586698 | 1.16307242 | -0.5805067 | -0.4344234 | 0.67015356 | 0.99999785 | -6.289425 down |
| KI - WT, cell_origin: iPSC | Q9UK45 | LSM7 | 3 | Homo sapien | 0.14405997 | -0.5629459 | 0.85106581 | -0.02399 | 0.43397029 | 0.67043289 | 0.99999785 | -6.2896643 up |
| KI - WT, cell_origin: iPSC | Q9Y3B4 | SF3B6 | 5 | Homo sapien | -0.2108317 | -1.2466131 | 0.82494975 | 0.69027217 | -0.4337806 | 0.67061005 | 0.99999785 | -6.2897107 down |
| KI - WT, cell_origin: iPSC | Q9BRA0 | NAA38 | 3 | Homo sapien | -0.4175967 | -2.4745638 | 1.6393704 | -3.1834361 | -0.4326454 | 0.67141656 | 0.99999785 | -6.2902142 down |
| KI - WT, cell_origin: iPSC | P53816 | PLAAT3 | 6 | Homo sapien | 0.52824403 | -2.0743748 | 3.13086282 | -1.7284786 | 0.43254012 | 0.67149136 | 0.99999785 | -6.2902608 up |
| KI - WT, cell_origin: iPSC | P22681 | CBL | 8 | Homo sapien | -0.1648815 | -0.9782958 | 0.64853277 | -0.3229989 | -0.4317844 | 0.67199706 | 0.99999785 | -6.290623 down |
| KI - WT, cell_origin: iPSC | Q9Y2L5 | TRAPPC8 | 7 | Homo sapien | 0.16524772 | -0.6507693 | 0.98126471 | -2.2065266 | 0.43129652 | 0.67233323 | 0.99999785 | -6.2908481 up |
| KI - WT, cell_origin: iPSC | P55199 | ELL | 2 | Homo sapien | 0.28299167 | -1.1148541 | 1.68083748 | -2.6469824 | 0.43128321 | 0.67236011 | 0.99999785 | -6.2908386 up |
| KI - WT, cell_origin: iPSC | Q0VF96 | CGNL1 | 2 | Homo sapien | -0.2311587 | -1.3737403 | 0.911423 | -2.099198 | -0.4309638 | 0.67258239 | 0.99999785 | -6.2909838 down |
| KI - WT, cell_origin: iPSC | Q9UQ80 | PA2G4 | 4 | Homo sapien | 0.20138354 | -0.7944856 | 1.19725266 | -1.2711437 | 0.43080365 | 0.6727027 | 0.99999785 | -6.2910488 up |
| KI - WT, cell_origin: iPSC | Q9H074 | PAIP1 | 7 | Homo sapien | -0.1526745 | -0.9089093 | 0.60356024 | -0.3813513 | -0.4300055 | 0.67325553 | 0.99999785 | -6.2914138 down |
| KI - WT, cell_origin: iPSC | Q8NBT2 | SPC24 | 2 | Homo sapien | 0.31522773 | -1.2472288 | 1.87768423 | -3.2133618 | 0.4297683 | 0.67343293 | 0.99999785 | -6.2915105 up |
| KI - WT, cell_origin: iPSC | Q96CT7 | CCDC124 | 14 | Homo sapien | -0.3056244 | -1.8227589 | 1.21151004 | 2.38218012 | -0.4293054 | 0.67379183 | 0.99999785 | -6.291688 down |
| KI - WT, cell_origin: iPSC | P07858 | CTSB | 5 | Homo sapien | -0.4416909 | -2.6352063 | 1.7518245 | -0.1876442 | -0.4291214 | 0.67392281 | 0.99999785 | -6.2917689 down |
| KI - WT, cell_origin: iPSC | O95466 | FMNL1 | 54 | Homo sapien | -0.3031156 | -1.8105858 | 1.20435468 | 2.16410288 | -0.4285109 | 0.67435737 | 0.99999785 | -6.2920369 down |
| KI - WT, cell_origin: iPSC | Q8IVH8 | MAP4K3 | 5 | Homo sapien | -0.163639 | -0.9775811 | 0.6503031 | -0.6840418 | -0.4282458 | 0.67451387 | 0.99999785 | -6.2921817 down |
| KI - WT, cell_origin: iPSC | O43809 | NUDT21 | 17 | Homo sapien | -0.437385 | -2.6158691 | 1.74109902 | 2.90686015 | -0.4278701 | 0.67481369 | 0.99999785 | -6.292318 down |
| KI - WT, cell_origin: iPSC | O00541 | PES1 | 7 | Homo sapien | -0.3150917 | -1.8845845 | 1.25440097 | -1.4011738 | -0.4278387 | 0.67483601 | 0.99999785 | -6.2923317 down |
| KI - WT, cell_origin: iPSC | Q15942 | ZYX | 12 | Homo sapien | 0.18590333 | -0.7408132 | 1.11261988 | 0.49779517 | 0.42750639 | 0.6750727 | 0.99999785 | -6.2924772 up |
| KI - WT, cell_origin: iPSC | P07737 | PFN1 | 7 | Homo sapien | -0.2403483 | -1.4395459 | 0.95884928 | 2.30437678 | -0.4271227 | 0.675346 | 0.99999785 | -6.2926451 down |
| KI - WT, cell_origin: iPSC | O95373 | IPO7 | 9 | Homo sapien | 0.26394596 | -1.0532277 | 1.58111966 | -0.369437 | 0.42704568 | 0.67540089 | 0.99999785 | -6.2926788 up |

| | | | | | | | | | | | | | |
|---|---|---|---|---|---|---|---|---|---|---|---|---|---|
| KI - WT, cell_origin: iPSC | P50750 | CDK9 | 11 | Homo sapien | 0.22151754 | -0.8851397 | 1.32817473 | 0.30613356 | 0.42657694 | 0.67573487 | 0.99999785 | -6.2928837 | up |
| KI - WT, cell_origin: iPSC | Q9UHB7 | AFF4 | 14 | Homo sapien | 0.28439743 | -1.1376565 | 1.70645134 | -1.5253435 | 0.42619856 | 0.67600452 | 0.99999785 | -6.293049 | up |
| KI - WT, cell_origin: iPSC | Q9BX40 | LSM14B | 5 | Homo sapien | -0.3339147 | -2.00433 | 1.33650067 | -1.8515751 | -0.4260037 | 0.67614339 | 0.99999785 | -6.293134 | down |
| KI - WT, cell_origin: iPSC | P00390 | GSR | 14 | Homo sapien | -0.1605817 | -0.9638358 | 0.64267238 | 2.53063512 | -0.4258543 | 0.67622073 | 0.99999785 | -6.2932249 | down |
| KI - WT, cell_origin: iPSC | Q92540 | SMG7 | 7 | Homo sapien | 0.38420807 | -1.5431296 | 2.31154576 | -1.6263906 | 0.42482596 | 0.67698307 | 0.99999785 | -6.2936472 | up |
| KI - WT, cell_origin: iPSC | O94804 | STK10 | 11 | Homo sapien | 0.15870435 | -0.6370518 | 0.95446054 | -0.4864347 | 0.42481955 | 0.67695503 | 0.99999785 | -6.2936786 | up |
| KI - WT, cell_origin: iPSC | Q13015 | MLLT11 | 2 | Homo sapien | 0.49079603 | -1.9736223 | 2.95521433 | -1.3147745 | 0.4244133 | 0.67727738 | 0.99999785 | -6.2938266 | up |
| KI - WT, cell_origin: iPSC | O00178 | GTPBP1 | 12 | Homo sapien | 0.11017135 | -0.4427585 | 0.66310125 | 1.0222557 | 0.42436375 | 0.67727132 | 0.99999785 | -6.2938845 | up |
| KI - WT, cell_origin: iPSC | Q86VM9 | ZC3H18 | 12 | Homo sapien | -0.1605572 | -0.9670727 | 0.64595828 | 1.34912403 | -0.4240675 | 0.67749501 | 0.99999785 | -6.2940024 | down |
| KI - WT, cell_origin: iPSC | Q9NYV4 | CDK12 | 13 | Homo sapien | -0.3354807 | -2.0214235 | 1.35046204 | -1.5865871 | -0.4240598 | 0.67752952 | 0.99999785 | -6.2939802 | down |
| KI - WT, cell_origin: iPSC | P26639 | TARS1 | 8 | Homo sapien | -0.2370344 | -1.4286337 | 0.95456492 | -0.6726143 | -0.4239196 | 0.67762959 | 0.99999785 | -6.2940411 | down |
| KI - WT, cell_origin: iPSC | Q8IUQ4 | SIAH1 | 8 | Homo sapien | 0.11196175 | -0.4509587 | 0.67488217 | 2.67619485 | 0.42360623 | 0.67781182 | 0.99999785 | -6.2942134 | up |
| KI - WT, cell_origin: iPSC | Q13363 | CTBP1 | 3 | Homo sapien | -0.196899 | -1.1875488 | 0.79375074 | -1.6271255 | -0.4233799 | 0.67798393 | 0.99999785 | -6.2943023 | down |
| KI - WT, cell_origin: iPSC | Q9NQS1 | AVEN | 8 | Homo sapien | 0.09740366 | -0.3929031 | 0.58771043 | 0.92963145 | 0.42310392 | 0.67817033 | 0.99999785 | -6.2944312 | up |
| KI - WT, cell_origin: iPSC | Q99627 | COPS8 | 2 | Homo sapien | 0.15475134 | -0.6256712 | 0.93517386 | -0.8728212 | 0.4223284 | 0.67872492 | 0.99999785 | -6.294766 | up |
| KI - WT, cell_origin: iPSC | P62736 | ACTA2 | 6 | Homo sapien | -0.205897 | -1.2454414 | 0.83364745 | 4.6924865 | -0.4220941 | 0.67893246 | 0.99999785 | -6.2948321 | down |
| KI - WT, cell_origin: iPSC | P55081 | MFAP1 | 9 | Homo sapien | -0.3602523 | -2.1791384 | 1.45863391 | 0.3105727 | -0.4220885 | 0.67893641 | 0.99999785 | -6.2948345 | down |
| KI - WT, cell_origin: iPSC | P07195 | LDHB | 9 | Homo sapien | -0.2739425 | -1.6574044 | 1.10951936 | 0.27796387 | -0.4219827 | 0.67901199 | 0.99999785 | -6.2948803 | down |
| KI - WT, cell_origin: iPSC | Q147X3 | NAA30 | 9 | Homo sapien | 0.22383509 | -0.9071351 | 1.35480532 | 0.05767386 | 0.42177357 | 0.67916132 | 0.99999785 | -6.2949707 | up |
| KI - WT, cell_origin: iPSC | P62277 | RPS13 | 10 | Homo sapien | 0.08823887 | -0.3575395 | 0.53401728 | 5.31049487 | 0.42158058 | 0.67925805 | 0.99999785 | -6.29509 | up |
| KI - WT, cell_origin: iPSC | Q9BSD7 | NTPCR | 11 | Homo sapien | 0.19191312 | -0.7783105 | 1.16213678 | 1.58989015 | 0.42153649 | 0.67933063 | 0.99999785 | -6.2950731 | up |
| KI - WT, cell_origin: iPSC | P26368 | U2AF2 | 14 | Homo sapien | -0.4059051 | -2.4629466 | 1.65113636 | 3.16585439 | -0.4205173 | 0.6800587 | 0.99999785 | -6.2955127 | down |
| KI - WT, cell_origin: iPSC | Q00403 | GTF2B | 4 | Homo sapien | 0.18505718 | -0.7524549 | 1.12256922 | -1.4310561 | 0.42046407 | 0.68006513 | 0.99999785 | -6.2955632 | up |
| KI - WT, cell_origin: iPSC | P12955 | PEPD | 7 | Homo sapien | -0.1724288 | -1.0459265 | 0.70106883 | -1.4928793 | -0.4204243 | 0.68008415 | 0.99999785 | -6.2955885 | down |
| KI - WT, cell_origin: iPSC | P21359 | NF1 | 24 | Homo sapien | 0.14329117 | -0.583639 | 0.87022134 | -0.951522 | 0.41982319 | 0.6805138 | 0.99999785 | -6.2958472 | up |
| KI - WT, cell_origin: iPSC | Q9Y573 | IPP | 3 | Homo sapien | 0.27312811 | -1.116444 | 1.66270018 | -1.8602425 | 0.41887814 | 0.6812303 | 0.99999785 | -6.2962175 | up |
| KI - WT, cell_origin: iPSC | Q9P2E2 | KIF17 | 6 | Homo sapien | 0.12976413 | -0.5302206 | 0.78974882 | -0.643806 | 0.41875544 | 0.68127725 | 0.99999785 | -6.2963057 | up |
| KI - WT, cell_origin: iPSC | Q9BVS4 | RIOK2 | 8 | Homo sapien | 0.23321588 | -0.9551839 | 1.42161564 | -1.356012 | 0.41821339 | 0.68170568 | 0.99999785 | -6.2965026 | up |
| KI - WT, cell_origin: iPSC | P22626 | HNRNPA2B1 | 21 | Homo sapien | -0.2690583 | -1.6404079 | 1.10229131 | 3.74605712 | -0.4181196 | 0.68177273 | 0.99999785 | -6.2965428 | down |
| KI - WT, cell_origin: iPSC | Q8NB91 | FANCB | 7 | Homo sapien | 0.25759807 | -1.0553854 | 1.57058152 | -1.7582249 | 0.41810534 | 0.68178296 | 0.99999785 | -6.2965489 | up |
| KI - WT, cell_origin: iPSC | P32969 | RPL9 | 9 | Homo sapien | -0.0934069 | -0.5695545 | 0.38274071 | 3.85515065 | -0.4178082 | 0.68195484 | 0.99999785 | -6.2967115 | down |
| KI - WT, cell_origin: iPSC | P21127 | CDK11B | 13 | Homo sapien | -0.2846499 | -1.7368647 | 1.16756492 | -0.101347 | -0.4177174 | 0.6820605 | 0.99999785 | -6.2967151 | down |
| KI - WT, cell_origin: iPSC | Q9UGP4 | LIMD1 | 11 | Homo sapien | -0.1816978 | -1.1086568 | 0.74526117 | -1.4021376 | -0.4175273 | 0.6821645 | 0.99999785 | -6.2968242 | down |
| KI - WT, cell_origin: iPSC | Q14999 | CUL7 | 16 | Homo sapien | 0.20226158 | -0.830796 | 1.23531915 | 0.06066459 | 0.41724509 | 0.68239839 | 0.99999785 | -6.2969171 | up |
| KI - WT, cell_origin: iPSC | Q12907 | LMAN2 | 13 | Homo sapien | 0.41331069 | -1.6999747 | 2.52659612 | 2.4620973 | 0.41679341 | 0.68272161 | 0.99999785 | -6.2971101 | up |
| KI - WT, cell_origin: iPSC | P58317 | ZNF121 | 4 | Homo sapien | -0.164265 | -1.0038044 | 0.6752744 | -1.974299 | -0.4167193 | 0.6827341 | 0.99999785 | -6.297177 | down |
| KI - WT, cell_origin: iPSC | O60493 | SNX3 | 4 | Homo sapien | 0.28490286 | -1.1737048 | 1.74351053 | -0.9955013 | 0.41625618 | 0.68310615 | 0.99999785 | -6.2973394 | up |
| KI - WT, cell_origin: iPSC | Q8IZC4 | RTKN2 | 5 | Homo sapien | 0.41270022 | -1.7008646 | 2.526265 | -1.9653634 | 0.41612279 | 0.68320164 | 0.99999785 | -6.2973963 | up |
| KI - WT, cell_origin: iPSC | P12830 | CDH1 | 14 | Homo sapien | 0.17439331 | -0.7197039 | 1.06849051 | 0.67468958 | 0.41561679 | 0.68355555 | 0.99999785 | -6.2976192 | up |
| KI - WT, cell_origin: iPSC | P33991 | MCM4 | 11 | Homo sapien | 0.16005377 | -0.6603885 | 0.98049605 | 0.18625663 | 0.41556196 | 0.68357478 | 0.99999785 | -6.2976599 | up |
| KI - WT, cell_origin: iPSC | P35080 | PFN2 | 6 | Homo sapien | -0.1442314 | -0.884073 | 0.59561012 | 3.33307254 | -0.4152781 | 0.68377806 | 0.99999785 | -6.2977807 | down |
| KI - WT, cell_origin: iPSC | P18074 | ERCC2 | 9 | Homo sapien | -0.1788877 | -1.0966269 | 0.73885142 | -1.271918 | -0.4152103 | 0.68382496 | 0.99999785 | -6.297811 | down |
| KI - WT, cell_origin: iPSC | P25490 | YY1 | 6 | Homo sapien | -0.2254637 | -1.3838928 | 0.93296545 | -0.5992662 | -0.4147721 | 0.6841689 | 0.99999785 | -6.2979714 | down |
| KI - WT, cell_origin: iPSC | O14737 | PDCD5 | 2 | Homo sapien | 0.31400305 | -1.2997262 | 1.9277323 | -2.075891 | 0.41467282 | 0.68423999 | 0.99999785 | -6.2980135 | up |
| KI - WT, cell_origin: iPSC | P07910 | HNRNPC | 17 | Homo sapien | -0.2124164 | -1.3049103 | 0.88007748 | 3.18826973 | -0.4143539 | 0.68446848 | 0.99999785 | -6.298149 | down |
| KI - WT, cell_origin: iPSC | Q6P1M3 | LLGL2 | 6 | Homo sapien | 0.23381747 | -0.970281 | 1.43791598 | -2.1897717 | 0.41364982 | 0.68494473 | 0.99999785 | -6.2984722 | up |

| | | | | | | | | | | | | | |
|---|---|---|---|---|---|---|---|---|---|---|---|---|---|
| KI - WT, cell_origin: iPSC | Q9NQC8 | IFT46 | 4 | Homo sapien | 0.11063453 | -0.4591306 | 0.68039969 | -1.1422155 | 0.41355616 | 0.68499988 | 0.99999785 | -6.2985222 | up |
| KI - WT, cell_origin: iPSC | Q96HR3 | MED30 | 3 | Homo sapien | 0.22390228 | -0.9299107 | 1.37771531 | -1.8486584 | 0.41343338 | 0.68510975 | 0.99999785 | -6.2985553 | up |
| KI - WT, cell_origin: iPSC | Q8N1I0 | DOCK4 | 28 | Homo sapien | 0.24029531 | -0.9984904 | 1.47908106 | 0.35425995 | 0.4133819 | 0.68516499 | 0.99999785 | -6.2985613 | up |
| KI - WT, cell_origin: iPSC | Q9ULX6 | AKAP8L | 13 | Homo sapien | 0.20441258 | -0.852313 | 1.26113813 | -0.6266208 | 0.41223776 | 0.68598526 | 0.99999785 | -6.2990454 | up |
| KI - WT, cell_origin: iPSC | Q96MX3 | ZNF48 | 6 | Homo sapien | -0.1134682 | -0.7002774 | 0.47334091 | -1.5962261 | -0.4118292 | 0.68623823 | 0.99999785 | -6.2992524 | down |
| KI - WT, cell_origin: iPSC | Q9H8H3 | METTL7A | 2 | Homo sapien | -0.1524879 | -0.9414127 | 0.636437 | -1.0790279 | -0.4116606 | 0.6863592 | 0.99999785 | -6.2993236 | down |
| KI - WT, cell_origin: iPSC | Q8IZ69 | TRMT2A | 2 | Homo sapien | 0.13146998 | -0.5493525 | 0.81229246 | -1.9219731 | 0.41127507 | 0.68663579 | 0.99999785 | -6.2994861 | up |
| KI - WT, cell_origin: iPSC | Q9HAD4 | WDR41 | 19 | Homo sapien | 0.28581229 | -1.1953809 | 1.76700544 | 2.36878251 | 0.41121749 | 0.68671707 | 0.99999785 | -6.299476 | up |
| KI - WT, cell_origin: iPSC | Q8IZQ5 | SELENOH | 5 | Homo sapien | -0.2369458 | -1.4671162 | 0.99322455 | -0.3658822 | -0.4104745 | 0.68725022 | 0.99999785 | -6.2997889 | down |
| KI - WT, cell_origin: iPSC | Q9Y2T2 | AP3M1 | 4 | Homo sapien | -0.1772107 | -1.0979918 | 0.74357036 | 0.08403974 | -0.4100923 | 0.68751626 | 0.99999785 | -6.2999567 | down |
| KI - WT, cell_origin: iPSC | P00338 | LDHA | 9 | Homo sapien | -0.3404205 | -2.1102257 | 1.42938469 | 0.9448357 | -0.4099139 | 0.68765259 | 0.99999785 | -6.3000246 | down |
| KI - WT, cell_origin: iPSC | O00299 | CLIC1 | 7 | Homo sapien | -0.1411189 | -0.8747011 | 0.59246329 | 0.60243348 | -0.409746 | 0.68773916 | 0.99999785 | -6.3001243 | down |
| KI - WT, cell_origin: iPSC | Q9Y6G9 | DYNC1LI1 | 11 | Homo sapien | 0.30454052 | -1.279532 | 1.88861301 | -1.4501044 | 0.40970608 | 0.68780175 | 0.99999785 | -6.3001119 | up |
| KI - WT, cell_origin: iPSC | P27797 | CALR | 9 | Homo sapien | 0.25047134 | -1.0530176 | 1.55396032 | -0.6026692 | 0.40949919 | 0.68795028 | 0.99999785 | -6.3001988 | up |
| KI - WT, cell_origin: iPSC | P35637 | FUS | 10 | Homo sapien | -0.1987761 | -1.233597 | 0.83604488 | 5.89025775 | -0.4093561 | 0.68805302 | 0.99999785 | -6.3002589 | down |
| KI - WT, cell_origin: iPSC | Q99816 | TSG101 | 18 | Homo sapien | 0.09748071 | -0.4101485 | 0.60510989 | 2.38505872 | 0.40898915 | 0.68827679 | 0.99999785 | -6.3004468 | up |
| KI - WT, cell_origin: iPSC | P29373 | CRABP2 | 3 | Homo sapien | -0.4726167 | -2.9355699 | 1.99033642 | -2.3968399 | -0.408936 | 0.6883547 | 0.99999785 | -6.3004351 | down |
| KI - WT, cell_origin: iPSC | O95747 | OXSR1 | 2 | Homo sapien | -0.2048801 | -1.2729766 | 0.86321645 | -1.607042 | -0.4086367 | 0.6885463 | 0.99999785 | -6.3005804 | down |
| KI - WT, cell_origin: iPSC | A5D8V6 | VPS37C | 4 | Homo sapien | 0.11890683 | -0.5008854 | 0.73869907 | 0.99784617 | 0.40860185 | 0.68855497 | 0.99999785 | -6.3006091 | up |
| KI - WT, cell_origin: iPSC | P49674 | CSNK1E | 5 | Homo sapien | -0.0996981 | -0.6195897 | 0.42019347 | 0.09817402 | -0.4084264 | 0.68868104 | 0.99999785 | -6.3006826 | down |
| KI - WT, cell_origin: iPSC | P23193 | TCEA1 | 7 | Homo sapien | -0.3026294 | -1.8825138 | 1.2772551 | -0.9081816 | -0.4082142 | 0.68887311 | 0.99999785 | -6.3007374 | down |
| KI - WT, cell_origin: iPSC | Q9UBC3 | DNMT3B | 17 | Homo sapien | -0.2905677 | -1.8075193 | 1.22638388 | 0.18366039 | -0.4082047 | 0.68887993 | 0.99999785 | -6.3007414 | down |
| KI - WT, cell_origin: iPSC | Q9Y3B3 | TMED7 | 8 | Homo sapien | 0.36010982 | -1.5216666 | 2.24188622 | 0.36719852 | 0.40782075 | 0.68915577 | 0.99999785 | -6.300902 | up |
| KI - WT, cell_origin: iPSC | Q9UKA9 | PTBP2 | 3 | Homo sapien | -0.1352561 | -0.841784 | 0.57127177 | 3.4686249 | -0.4077984 | 0.68914403 | 0.99999785 | -6.3009351 | down |
| KI - WT, cell_origin: iPSC | Q5T0Z8 | C6orf132 | 7 | Homo sapien | 0.20233213 | -0.8547849 | 1.25944916 | -2.219357 | 0.40764437 | 0.68924291 | 0.99999785 | -6.3010096 | up |
| KI - WT, cell_origin: iPSC | P56182 | RRP1 | 4 | Homo sapien | -0.2773836 | -1.7291098 | 1.17434273 | -2.5343392 | -0.4071318 | 0.68964134 | 0.99999785 | -6.301198 | down |
| KI - WT, cell_origin: iPSC | Q96RQ9 | IL4I1 | 7 | Homo sapien | 0.32191124 | -1.3632222 | 2.00704463 | 0.59361568 | 0.40710292 | 0.68967159 | 0.99999785 | -6.3012019 | up |
| KI - WT, cell_origin: iPSC | O95319 | CELF2 | 7 | Homo sapien | -0.3590101 | -2.2400631 | 1.52204285 | 0.81619294 | -0.4067317 | 0.6899384 | 0.99999785 | -6.3013567 | down |
| KI - WT, cell_origin: iPSC | Q96Q05 | TRAPPC9 | 2 | Homo sapien | 0.25911083 | -1.0992999 | 1.61752154 | -1.0640907 | 0.40649652 | 0.69010747 | 0.99999785 | -6.3014548 | up |
| KI - WT, cell_origin: iPSC | P35527 | KRT9 | 25 | Homo sapien | -0.4479101 | -2.801021 | 1.90520074 | 2.81430246 | -0.4056494 | 0.69071656 | 0.99999785 | -6.3018075 | down |
| KI - WT, cell_origin: iPSC | P50453 | SERPINB9 | 19 | Homo sapien | -0.3008335 | -1.8813075 | 1.27964049 | 1.92497084 | -0.4056404 | 0.69072301 | 0.99999785 | -6.3018112 | down |
| KI - WT, cell_origin: iPSC | Q96II8 | LRCH3 | 13 | Homo sapien | -0.1287929 | -0.8050345 | 0.54744879 | -0.5020498 | -0.4056295 | 0.6906915 | 0.99999785 | -6.3018493 | down |
| KI - WT, cell_origin: iPSC | O75340 | PDCD6 | 9 | Homo sapien | -0.1286925 | -0.8047258 | 0.54734075 | 4.10776034 | -0.4055115 | 0.69078814 | 0.99999785 | -6.3018884 | down |
| KI - WT, cell_origin: iPSC | Q9NP66 | HMG20A | 3 | Homo sapien | 0.1080752 | -0.4599544 | 0.67610477 | -1.2905943 | 0.40522366 | 0.69098344 | 0.99999785 | -6.302018 | up |
| KI - WT, cell_origin: iPSC | Q8IZ40 | RCOR2 | 5 | Homo sapien | 0.1129984 | -0.4824515 | 0.70844825 | -0.8510085 | 0.4041725 | 0.6917398 | 0.99999785 | -6.302454 | up |
| KI - WT, cell_origin: iPSC | Q8NEZ2 | VPS37A | 11 | Homo sapien | 0.11832983 | -0.5053796 | 0.74203921 | 2.0319957 | 0.40406537 | 0.69181691 | 0.99999785 | -6.3024984 | up |
| KI - WT, cell_origin: iPSC | Q8TAF3 | WDR48 | 7 | Homo sapien | 0.19530924 | -0.8348787 | 1.22549722 | -0.5854316 | 0.40402541 | 0.69188486 | 0.99999785 | -6.3024816 | up |
| KI - WT, cell_origin: iPSC | Q9H7D0 | DOCK5 | 10 | Homo sapien | -0.4879149 | -3.0624461 | 2.08661628 | -1.0633368 | -0.4038762 | 0.69199222 | 0.99999785 | -6.3025434 | down |
| KI - WT, cell_origin: iPSC | P16152 | CBR1 | 9 | Homo sapien | -0.1676113 | -1.0529643 | 0.7177417 | -0.1392956 | -0.4033381 | 0.69236171 | 0.99999785 | -6.3027813 | down |
| KI - WT, cell_origin: iPSC | Q04724;Q04 | TLE1;TLE2;TL | 3 | Homo sapien | -0.1917756 | -1.2051717 | 0.82162048 | -0.9154816 | -0.4032891 | 0.6924148 | 0.99999785 | -6.3027864 | down |
| KI - WT, cell_origin: iPSC | Q9H9A5 | CNOT10 | 15 | Homo sapien | 0.14469984 | -0.61981 | 0.90920973 | 1.27047072 | 0.40317329 | 0.69246909 | 0.99999785 | -6.302859 | up |
| KI - WT, cell_origin: iPSC | P04264 | KRT1 | 29 | Homo sapien | -0.3158408 | -1.9860774 | 1.35439576 | 5.37510654 | -0.4029885 | 0.69263126 | 0.99999785 | -6.3029107 | down |
| KI - WT, cell_origin: iPSC | P62854 | RPS26 | 4 | Homo sapien | 0.09111002 | -0.3904848 | 0.57270481 | 4.88676441 | 0.40292482 | 0.69263801 | 0.99999785 | -6.3029702 | up |
| KI - WT, cell_origin: iPSC | P06454 | PTMA | 5 | Homo sapien | -0.3635968 | -2.2889915 | 1.56179796 | -1.8518923 | -0.4024413 | 0.69302522 | 0.99999785 | -6.3031367 | down |
| KI - WT, cell_origin: iPSC | Q96PK6 | RBM14 | 19 | Homo sapien | -0.3060504 | -1.9271611 | 1.31506034 | 2.45107497 | -0.4023302 | 0.69310527 | 0.99999785 | -6.3031826 | down |
| KI - WT, cell_origin: iPSC | Q9BT23 | LIMD2 | 3 | Homo sapien | 0.23216079 | -0.9981883 | 1.46250986 | 0.64101568 | 0.40212668 | 0.69325183 | 0.99999785 | -6.3032665 | up |

| | | | | | | | | | | | | | |
|---|---|---|---|---|---|---|---|---|---|---|---|---|---|
| KI - WT, cell_origin: iPSC | P61981 | YWHAG | 9 | Homo sapien | 0.09376432 | -0.4030049 | 0.59053357 | 2.86000973 | 0.40199681 | 0.6933064 | 0.99999785 | -6.3033531 | up |
| KI - WT, cell_origin: iPSC | Q6P1N0 | CC2D1A | 6 | Homo sapien | 0.19175009 | -0.8253334 | 1.20883362 | -1.6835795 | 0.40159265 | 0.69360752 | 0.99999785 | -6.3035111 | up |
| KI - WT, cell_origin: iPSC | Q8WUA4 | GTF3C2 | 3 | Homo sapien | 0.08614273 | -0.3707181 | 0.54300359 | -0.8423757 | 0.4015821 | 0.69360517 | 0.99999785 | -6.3035239 | up |
| KI - WT, cell_origin: iPSC | O00311 | CDC7 | 5 | Homo sapien | -0.2851003 | -1.7993282 | 1.22912767 | -4.1460454 | -0.4010016 | 0.6940235 | 0.99999785 | -6.3037627 | down |
| KI - WT, cell_origin: iPSC | Q5RKV6 | EXOSC6 | 5 | Homo sapien | -0.1565881 | -0.9898792 | 0.67670296 | -0.3899203 | -0.4002948 | 0.69454445 | 0.99999785 | -6.3040433 | down |
| KI - WT, cell_origin: iPSC | P62847 | RPS24 | 5 | Homo sapien | 0.09712953 | -0.4207007 | 0.61495976 | 4.88406247 | 0.39948784 | 0.69511476 | 0.99999785 | -6.3043839 | up |
| KI - WT, cell_origin: iPSC | Q7Z6E9 | RBBP6 | 11 | Homo sapien | -0.323595 | -2.0565119 | 1.40932199 | -1.756315 | -0.397948 | 0.69626406 | 0.99999785 | -6.304981 | down |
| KI - WT, cell_origin: iPSC | Q8N5F7 | NKAP | 7 | Homo sapien | 0.18041155 | -0.7858049 | 1.14662799 | -1.2630895 | 0.39774783 | 0.69638149 | 0.99999785 | -6.3050855 | up |
| KI - WT, cell_origin: iPSC | Q96KP1 | EXOC2 | 20 | Homo sapien | 0.30365488 | -1.3247956 | 1.9321054 | -1.3656785 | 0.39738189 | 0.69667259 | 0.99999785 | -6.305212 | up |
| KI - WT, cell_origin: iPSC | Q00653 | NFKB2 | 16 | Homo sapien | 0.47434286 | -2.0725958 | 3.02128153 | -0.7963479 | 0.39689557 | 0.6970236 | 0.99999785 | -6.3054101 | up |
| KI - WT, cell_origin: iPSC | P35568 | IRS1 | 4 | Homo sapien | -0.3361592 | -2.1409502 | 1.46863184 | -3.3656047 | -0.396878 | 0.697027 | 0.99999785 | -6.305425 | down |
| KI - WT, cell_origin: iPSC | Q6UN15 | FIP1L1 | 7 | Homo sapien | -0.2072218 | -1.321158 | 0.90671444 | -0.0793481 | -0.39644 | 0.69735247 | 0.99999785 | -6.3055955 | down |
| KI - WT, cell_origin: iPSC | P09936 | UCHL1 | 4 | Homo sapien | -0.2701239 | -1.722272 | 1.18202415 | -1.8606359 | -0.3964191 | 0.69736757 | 0.99999785 | -6.305604 | down |
| KI - WT, cell_origin: iPSC | Q8N1G2 | CMTR1 | 18 | Homo sapien | -0.2042285 | -1.3025631 | 0.89410601 | -0.91067 | -0.3962636 | 0.69747983 | 0.99999785 | -6.3056672 | down |
| KI - WT, cell_origin: iPSC | P46060 | RANGAP1 | 12 | Homo sapien | 0.1976582 | -0.865491 | 1.26080743 | -0.3395681 | 0.39620777 | 0.69752014 | 0.99999785 | -6.3056899 | up |
| KI - WT, cell_origin: iPSC | Q9BQA1 | WDR77 | 8 | Homo sapien | -0.2211152 | -1.4116896 | 0.96945912 | 0.56088881 | -0.3957897 | 0.69782202 | 0.99999785 | -6.3058597 | down |
| KI - WT, cell_origin: iPSC | O75369 | FLNB | 54 | Homo sapien | -0.264371 | -1.688765 | 1.16002294 | 0.86357007 | -0.3955361 | 0.69800518 | 0.99999785 | -6.3059627 | down |
| KI - WT, cell_origin: iPSC | Q9NWS0 | PIH1D1 | 4 | Homo sapien | 0.13943784 | -0.6123023 | 0.89117801 | 1.52660573 | 0.39510141 | 0.69828906 | 0.99999785 | -6.3061641 | up |
| KI - WT, cell_origin: iPSC | Q96T76 | MMS19 | 10 | Homo sapien | 0.21082831 | -0.9264101 | 1.34806674 | -1.6326068 | 0.39501323 | 0.69837293 | 0.99999785 | -6.306183 | up |
| KI - WT, cell_origin: iPSC | Q96S15 | WDR24 | 15 | Homo sapien | -0.0997035 | -0.6385691 | 0.43916219 | -0.0463628 | -0.3940664 | 0.69902883 | 0.99999785 | -6.3065898 | down |
| KI - WT, cell_origin: iPSC | Q6GMV2 | SMYD5 | 3 | Homo sapien | 0.15508241 | -0.6849377 | 0.99510254 | -2.5509387 | 0.39319911 | 0.69965581 | 0.99999785 | -6.30694 | up |
| KI - WT, cell_origin: iPSC | P07339 | CTSD | 11 | Homo sapien | 0.31595769 | -1.3966843 | 2.02859973 | 0.03226991 | 0.39315582 | 0.69972513 | 0.99999785 | -6.3069257 | up |
| KI - WT, cell_origin: iPSC | Q86V48 | LUZP1 | 26 | Homo sapien | -0.1477113 | -0.9483432 | 0.65292054 | 0.58892454 | -0.3930057 | 0.69980701 | 0.99999785 | -6.3070085 | down |
| KI - WT, cell_origin: iPSC | P62937 | PPIA | 11 | Homo sapien | -0.1200999 | -0.7717017 | 0.53150197 | 5.3489409 | -0.3926836 | 0.70004924 | 0.99999785 | -6.3071306 | down |
| KI - WT, cell_origin: iPSC | O75152 | ZC3H11A | 4 | Homo sapien | -0.1977497 | -1.2704942 | 0.87499485 | -2.9887057 | -0.392608 | 0.70008325 | 0.99999785 | -6.3071782 | down |
| KI - WT, cell_origin: iPSC | P15924 | DSP | 115 | Homo sapien | 0.11728099 | -0.5191637 | 0.75372572 | 1.74362251 | 0.39247012 | 0.70018298 | 0.99999785 | -6.3072337 | up |
| KI - WT, cell_origin: iPSC | Q14315 | FLNC | 25 | Homo sapien | 0.15440125 | -0.6863036 | 0.99510605 | -0.7847515 | 0.39121372 | 0.70110157 | 0.99999785 | -6.3077307 | up |
| KI - WT, cell_origin: iPSC | Q9H944 | MED20 | 2 | Homo sapien | 0.17104108 | -0.7612847 | 1.10336681 | -1.3499912 | 0.3907865 | 0.70141074 | 0.99999785 | -6.307902 | up |
| KI - WT, cell_origin: iPSC | Q01469 | FABP5 | 8 | Homo sapien | -0.3002365 | -1.9407378 | 1.3402648 | 0.53074144 | -0.3900222 | 0.70199206 | 0.99999785 | -6.3081849 | down |
| KI - WT, cell_origin: iPSC | P31943 | HNRNPH1 | 18 | Homo sapien | -0.2343424 | -1.5159302 | 1.04724528 | 4.49591583 | -0.3896772 | 0.70224181 | 0.99999785 | -6.3083229 | down |
| KI - WT, cell_origin: iPSC | Q9Y5R8 | TRAPPC1 | 4 | Homo sapien | 0.16037253 | -0.7165787 | 1.03732375 | -0.3568958 | 0.38958542 | 0.70228615 | 0.99999785 | -6.3083779 | up |
| KI - WT, cell_origin: iPSC | P19838 | NFKB1 | 10 | Homo sapien | 0.41948721 | -1.875719 | 2.71469339 | -1.8687156 | 0.38949275 | 0.70237533 | 0.99999785 | -6.3083967 | up |
| KI - WT, cell_origin: iPSC | Q9UHV9 | PFDN2 | 4 | Homo sapien | 0.14815711 | -0.6622436 | 0.95855779 | 0.31907994 | 0.38944005 | 0.70238708 | 0.99999785 | -6.3084396 | up |
| KI - WT, cell_origin: iPSC | P12814 | ACTN1 | 7 | Homo sapien | -0.2499681 | -1.6197696 | 1.11983338 | -1.0396459 | -0.3888923 | 0.70281016 | 0.99999785 | -6.3086365 | down |
| KI - WT, cell_origin: iPSC | Q12906 | ILF3 | 36 | Homo sapien | -0.1049844 | -0.680093 | 0.47012415 | 2.5074631 | -0.3887897 | 0.70284693 | 0.99999785 | -6.3087085 | down |
| KI - WT, cell_origin: iPSC | Q6KC79 | NIPBL | 6 | Homo sapien | -0.2183698 | -1.4157559 | 0.97901626 | -2.0761459 | -0.3885074 | 0.70306586 | 0.99999785 | -6.3088091 | down |
| KI - WT, cell_origin: iPSC | Q9UET6 | FTSJ1 | 5 | Homo sapien | 0.10583329 | -0.4744467 | 0.68611327 | -0.144622 | 0.38844039 | 0.70309993 | 0.99999785 | -6.3088478 | up |
| KI - WT, cell_origin: iPSC | Q9H2Y7 | ZNF106 | 4 | Homo sapien | -0.1636737 | -1.0623338 | 0.73498644 | -0.0082195 | -0.3880613 | 0.70339986 | 0.99999785 | -6.3089779 | down |
| KI - WT, cell_origin: iPSC | P47914 | RPL29 | 3 | Homo sapien | 0.14613514 | -0.6601837 | 0.95245401 | 4.74446304 | 0.38623373 | 0.70473658 | 0.99999785 | -6.309694 | up |
| KI - WT, cell_origin: iPSC | P78346 | RPP30 | 10 | Homo sapien | -0.1789218 | -1.1660833 | 0.80823965 | -0.9086451 | -0.3862097 | 0.70474632 | 0.99999785 | -6.3097099 | down |
| KI - WT, cell_origin: iPSC | Q96J01 | THOC3 | 7 | Homo sapien | 0.15474441 | -0.699015 | 1.00850377 | -0.9018433 | 0.38608798 | 0.70481451 | 0.99999785 | -6.3097746 | up |
| KI - WT, cell_origin: iPSC | P37837 | TALDO1 | 3 | Homo sapien | 0.26494704 | -1.1978303 | 1.72772438 | -0.9843932 | 0.38599636 | 0.70490868 | 0.99999785 | -6.3097881 | up |
| KI - WT, cell_origin: iPSC | P02452 | COL1A1 | 3 | Homo sapien | -0.2134896 | -1.392586 | 0.96560673 | -0.235285 | -0.3858601 | 0.70500751 | 0.99999785 | -6.3098421 | down |
| KI - WT, cell_origin: iPSC | P78527 | PRKDC | 154 | Homo sapien | -0.1853519 | -1.2092826 | 0.83857873 | 2.41672144 | -0.3857704 | 0.70507251 | 0.99999785 | -6.3098776 | down |
| KI - WT, cell_origin: iPSC | P49643 | PRIM2 | 3 | Homo sapien | -0.4183456 | -2.7306469 | 1.8939557 | -3.2633104 | -0.3855611 | 0.70522433 | 0.99999785 | -6.3099604 | down |
| KI - WT, cell_origin: iPSC | Q14738 | PPP2R5D | 8 | Homo sapien | 0.18347482 | -0.8317319 | 1.19868154 | -1.3035241 | 0.38503757 | 0.70558687 | 0.99999785 | -6.3101816 | up |

| | | | | | | | | | | | | | |
|---|---|---|---|---|---|---|---|---|---|---|---|---|---|
| KI - WT, cell_origin: iPSC | Q8WVD3 | RNF138 | 4 | Homo sapien | -0.2410256 | -1.5752387 | 1.09318748 | -2.503854 | -0.3848185 | 0.70573687 | 0.99999785 | -6.3102755 | down |
| KI - WT, cell_origin: iPSC | P12956 | XRCC6 | 38 | Homo sapien | -0.210202 | -1.3743649 | 0.95396091 | 3.29865245 | -0.3847915 | 0.70578251 | 0.99999785 | -6.3102647 | down |
| KI - WT, cell_origin: iPSC | Q16555 | DPYSL2 | 23 | Homo sapien | -0.1344743 | -0.8791626 | 0.61021399 | 1.33268046 | -0.3846348 | 0.70586532 | 0.99999785 | -6.310352 | down |
| KI - WT, cell_origin: iPSC | Q5SZL2 | CEP85L | 4 | Homo sapien | -0.5691976 | -3.7254718 | 2.58707662 | -1.8057217 | -0.384318 | 0.70612605 | 0.99999785 | -6.3104516 | down |
| KI - WT, cell_origin: iPSC | P50222 | MEOX2 | 2 | Homo sapien | -0.2500372 | -1.6387443 | 1.13866993 | -2.4163533 | -0.3836419 | 0.70660676 | 0.99999785 | -6.3107262 | down |
| KI - WT, cell_origin: iPSC | P54274 | TERF1 | 7 | Homo sapien | -0.1984376 | -1.3011552 | 0.90427998 | -1.9412705 | -0.3833342 | 0.70681402 | 0.99999785 | -6.3108605 | down |
| KI - WT, cell_origin: iPSC | Q99613 | EIF3C | 31 | Homo sapien | -0.0858549 | -0.5635366 | 0.39182683 | 3.37158978 | -0.3827948 | 0.70719464 | 0.99999785 | -6.3110816 | down |
| KI - WT, cell_origin: iPSC | Q9BY44 | EIF2A | 18 | Homo sapien | 0.11529312 | -0.5274707 | 0.75805696 | 0.73940169 | 0.38202488 | 0.70775381 | 0.99999785 | -6.3113837 | up |
| KI - WT, cell_origin: iPSC | Q15382 | RHEB | 4 | Homo sapien | 0.2380946 | -1.091806 | 1.56799519 | -1.0779545 | 0.38153356 | 0.70814749 | 0.99999785 | -6.3115462 | up |
| KI - WT, cell_origin: iPSC | Q9C0D5 | TANC1 | 8 | Homo sapien | 0.15155886 | -0.6950671 | 0.9981848 | -1.7829963 | 0.38126721 | 0.70830423 | 0.99999785 | -6.3116805 | up |
| KI - WT, cell_origin: iPSC | O14893 | GEMIN2 | 3 | Homo sapien | -0.2103232 | -1.3869687 | 0.9663223 | -1.1526521 | -0.3809289 | 0.70858678 | 0.99999785 | -6.3117829 | down |
| KI - WT, cell_origin: iPSC | Q9BV44 | THUMPD3 | 7 | Homo sapien | 0.21860327 | -1.0039579 | 1.44116446 | -2.665457 | 0.38085055 | 0.70861102 | 0.99999785 | -6.3118402 | up |
| KI - WT, cell_origin: iPSC | Q99497 | PARK7 | 12 | Homo sapien | -0.1205614 | -0.7951005 | 0.5539777 | 3.69851228 | -0.3807219 | 0.70870982 | 0.99999785 | -6.3118861 | down |
| KI - WT, cell_origin: iPSC | Q13568 | IRF5 | 2 | Homo sapien | 0.50017504 | -2.2999605 | 3.3003106 | -1.6214598 | 0.38066708 | 0.708777 | 0.99999785 | -6.3118852 | up |
| KI - WT, cell_origin: iPSC | Q7Z4H7 | HAUS6 | 5 | Homo sapien | 0.21946984 | -1.0114759 | 1.45041561 | -2.637783 | 0.379761 | 0.70940378 | 0.99999785 | -6.3122648 | up |
| KI - WT, cell_origin: iPSC | Q9NNW5 | WDR6 | 10 | Homo sapien | 0.13893811 | -0.6407586 | 0.91863486 | 0.67184576 | 0.37958517 | 0.70953706 | 0.99999785 | -6.3123289 | up |
| KI - WT, cell_origin: iPSC | Q149N8 | SHPRH | 2 | Homo sapien | -0.1353348 | -0.8953533 | 0.62468374 | -2.7456709 | -0.3792494 | 0.70977091 | 0.99999785 | -6.312468 | down |
| KI - WT, cell_origin: iPSC | P62995 | TRA2B | 9 | Homo sapien | -0.204283 | -1.3525688 | 0.94400274 | 2.13091286 | -0.3791269 | 0.70989649 | 0.99999785 | -6.312486 | down |
| KI - WT, cell_origin: iPSC | P40763 | STAT3 | 31 | Homo sapien | 0.1298423 | -0.5999737 | 0.85965826 | 1.50046213 | 0.37894202 | 0.70999865 | 0.99999785 | -6.3125842 | up |
| KI - WT, cell_origin: iPSC | O94992 | HEXIM1 | 5 | Homo sapien | -0.1789193 | -1.1864762 | 0.8286376 | -0.4471761 | -0.3784339 | 0.71040048 | 0.99999785 | -6.3127556 | down |
| KI - WT, cell_origin: iPSC | Q9BV94 | EDEM2 | 2 | Homo sapien | -0.2021733 | -1.3413939 | 0.9370472 | 2.76437995 | -0.3781973 | 0.71057254 | 0.99999785 | -6.3128475 | down |
| KI - WT, cell_origin: iPSC | O00422 | SAP18 | 10 | Homo sapien | -0.1444205 | -0.9579661 | 0.66912519 | 0.61957626 | -0.3781505 | 0.71058101 | 0.99999785 | -6.3128864 | down |
| KI - WT, cell_origin: iPSC | Q9UBF2 | COPG2 | 13 | Homo sapien | 0.19975234 | -0.926779 | 1.32628371 | -1.009298 | 0.37787744 | 0.71080522 | 0.99999785 | -6.3129716 | up |
| KI - WT, cell_origin: iPSC | P34947 | GRK5 | 4 | Homo sapien | -0.1769078 | -1.1752412 | 0.82142567 | -2.2440417 | -0.3774079 | 0.71111046 | 0.99999785 | -6.3131832 | down |
| KI - WT, cell_origin: iPSC | Q8NEB9 | PIK3C3 | 13 | Homo sapien | 0.15757395 | -0.7325537 | 1.04770161 | -0.2081844 | 0.37715042 | 0.7113176 | 0.99999785 | -6.3132669 | up |
| KI - WT, cell_origin: iPSC | Q9BYG3 | NIFK | 4 | Homo sapien | 0.21569659 | -1.0038138 | 1.43520695 | 0.1872597 | 0.37692951 | 0.71149492 | 0.99999785 | -6.313339 | up |
| KI - WT, cell_origin: iPSC | Q9H501 | ESF1 | 8 | Homo sapien | -0.1751535 | -1.1652301 | 0.81492307 | -1.1950696 | -0.3769055 | 0.71149583 | 0.99999785 | -6.3133617 | down |
| KI - WT, cell_origin: iPSC | O75147 | OBSL1 | 29 | Homo sapien | 0.22159414 | -1.0314624 | 1.47465068 | -1.1940961 | 0.37686861 | 0.71153924 | 0.99999785 | -6.3133626 | up |
| KI - WT, cell_origin: iPSC | Q9BRS2 | RIOK1 | 14 | Homo sapien | -0.2416637 | -1.6086729 | 1.12534551 | -0.0284832 | -0.3767405 | 0.71163246 | 0.99999785 | -6.3134121 | down |
| KI - WT, cell_origin: iPSC | Q9UG01 | IFT172 | 16 | Homo sapien | 0.10088243 | -0.4712912 | 0.67305603 | -1.9076474 | 0.37551508 | 0.7124884 | 0.99999785 | -6.3139147 | up |
| KI - WT, cell_origin: iPSC | O95639 | CPSF4 | 5 | Homo sapien | -0.1820119 | -1.2148708 | 0.85084693 | -1.4506795 | -0.375411 | 0.71257911 | 0.99999785 | -6.3139427 | down |
| KI - WT, cell_origin: iPSC | P36578 | RPL4 | 22 | Homo sapien | -0.0872485 | -0.5830348 | 0.40853784 | 5.46782499 | -0.3748029 | 0.71300712 | 0.99999785 | -6.314189 | down |
| KI - WT, cell_origin: iPSC | P11413 | G6PD | 17 | Homo sapien | -0.1409116 | -0.9424671 | 0.66064386 | 0.53829685 | -0.3744822 | 0.7132515 | 0.99999785 | -6.3143037 | down |
| KI - WT, cell_origin: iPSC | P20290 | BTF3 | 7 | Homo sapien | -0.1406698 | -0.9413474 | 0.66000773 | 1.297073 | -0.3742494 | 0.71342105 | 0.99999785 | -6.3143931 | down |
| KI - WT, cell_origin: iPSC | P31948 | STIP1 | 22 | Homo sapien | 0.16918511 | -0.7947024 | 1.13307256 | 0.263029 | 0.37405733 | 0.7135863 | 0.99999785 | -6.3144466 | up |
| KI - WT, cell_origin: iPSC | Q9UBW7 | ZMYM2 | 14 | Homo sapien | 0.24073897 | -1.1310117 | 1.61248961 | 0.12504112 | 0.37400171 | 0.71362682 | 0.99999785 | -6.314468 | up |
| KI - WT, cell_origin: iPSC | P62306 | SNRPF | 5 | Homo sapien | -0.2806762 | -1.8808074 | 1.31945509 | 1.95608559 | -0.3738112 | 0.71376564 | 0.99999785 | -6.3145411 | down |
| KI - WT, cell_origin: iPSC | Q07020 | RPL18 | 7 | Homo sapien | -0.093957 | -0.6312276 | 0.44331355 | 5.4263399 | -0.3724581 | 0.71471623 | 0.99999785 | -6.3150883 | down |
| KI - WT, cell_origin: iPSC | P50402 | EMD | 2 | Homo sapien | -0.2693905 | -1.8114532 | 1.27267213 | -0.4590078 | -0.3722911 | 0.71487354 | 0.99999785 | -6.3151236 | down |
| KI - WT, cell_origin: iPSC | P27708 | CAD | 86 | Homo sapien | 0.12600976 | -0.5954399 | 0.84745939 | 4.44352043 | 0.3721605 | 0.7149592 | 0.99999785 | -6.3151812 | up |
| KI - WT, cell_origin: iPSC | Q00013 | MPP1 | 8 | Homo sapien | 0.24339671 | -1.1517287 | 1.63852214 | -0.3072362 | 0.37179524 | 0.7152351 | 0.99999785 | -6.3153131 | up |
| KI - WT, cell_origin: iPSC | P02686 | MBP | 4 | Homo sapien | 0.30527454 | -1.4446197 | 2.05516874 | 0.69165347 | 0.37177584 | 0.71524925 | 0.99999785 | -6.3153205 | up |
| KI - WT, cell_origin: iPSC | P46783 | RPS10 | 14 | Homo sapien | -0.0746541 | -0.5036111 | 0.354303 | 5.79551387 | -0.3706631 | 0.71602522 | 0.99999785 | -6.3157733 | down |
| KI - WT, cell_origin: iPSC | Q7Z7K6 | CENPV | 4 | Homo sapien | -0.2872599 | -1.9394499 | 1.36493005 | -1.6748478 | -0.3705249 | 0.71616165 | 0.99999785 | -6.3157974 | down |
| KI - WT, cell_origin: iPSC | P50238 | CRIP1 | 4 | Homo sapien | 0.34527021 | -1.6456704 | 2.33621084 | -0.1670657 | 0.3695755 | 0.71685447 | 0.99999785 | -6.3161583 | up |
| KI - WT, cell_origin: iPSC | Q14527 | HLTF | 11 | Homo sapien | 0.11965261 | -0.5698898 | 0.80919502 | -1.0626231 | 0.36957353 | 0.71682039 | 0.99999785 | -6.3161874 | up |

| | | | | | | | | | | | | | |
|---|---|---|---|---|---|---|---|---|---|---|---|---|---|
| KI - WT, cell_origin: iPSC | P46777 | RPL5 | 15 | Homo sapien | -0.0910327 | -0.6156831 | 0.43361765 | 3.95537283 | -0.3695449 | 0.71684127 | 0.99999785 | -6.3161983 | down |
| KI - WT, cell_origin: iPSC | Q9P2I0 | CPSF2 | 16 | Homo sapien | -0.1529418 | -1.0345292 | 0.7286455 | -1.3018802 | -0.3695257 | 0.71686124 | 0.99999785 | -6.3162008 | down |
| KI - WT, cell_origin: iPSC | Q5THJ4 | VPS13D | 17 | Homo sapien | 0.14085878 | -0.671785 | 0.95350255 | -0.5178528 | 0.36922448 | 0.71708426 | 0.99999785 | -6.3163126 | up |
| KI - WT, cell_origin: iPSC | Q92538 | GBF1 | 34 | Homo sapien | 0.178594 | -0.8528329 | 1.21002086 | 0.56022404 | 0.36900377 | 0.71727179 | 0.99999785 | -6.3163752 | up |
| KI - WT, cell_origin: iPSC | P78362 | SRPK2 | 11 | Homo sapien | 0.09136206 | -0.4369538 | 0.61967786 | 0.32564052 | 0.36830867 | 0.71774391 | 0.99999785 | -6.3166666 | up |
| KI - WT, cell_origin: iPSC | Q86W50 | METTL16 | 4 | Homo sapien | -0.2075262 | -1.4084949 | 0.99344248 | -0.2704591 | -0.3682507 | 0.71782159 | 0.99999785 | -6.3166604 | down |
| KI - WT, cell_origin: iPSC | P16403 | H1-2 | 6 | Homo sapien | -0.218493 | -1.4836921 | 1.04670597 | 4.22692912 | -0.3680282 | 0.71798408 | 0.99999785 | -6.3167446 | down |
| KI - WT, cell_origin: iPSC | Q9HCG8 | CWC22 | 21 | Homo sapien | -0.1735334 | -1.178832 | 0.83176514 | 0.2184678 | -0.3678667 | 0.71810204 | 0.99999785 | -6.3168056 | down |
| KI - WT, cell_origin: iPSC | Q9Y508 | RNF114 | 7 | Homo sapien | -0.1943613 | -1.3203932 | 0.93167052 | -0.0665568 | -0.3678422 | 0.71811992 | 0.99999785 | -6.3168149 | down |
| KI - WT, cell_origin: iPSC | Q92945 | KHSRP | 29 | Homo sapien | -0.1941601 | -1.3196456 | 0.93132535 | 4.45673156 | -0.3676398 | 0.71826778 | 0.99999785 | -6.3168914 | down |
| KI - WT, cell_origin: iPSC | O43684 | BUB3 | 13 | Homo sapien | -0.185477 | -1.2626548 | 0.89170086 | 2.39333057 | -0.3669484 | 0.71877285 | 0.99999785 | -6.3171523 | down |
| KI - WT, cell_origin: iPSC | Q9BW19 | KIFC1 | 18 | Homo sapien | -0.193216 | -1.3173249 | 0.93089286 | -0.8280022 | -0.3663002 | 0.71924646 | 0.99999785 | -6.3173964 | down |
| KI - WT, cell_origin: iPSC | P16949 | STMN1 | 6 | Homo sapien | -0.1450312 | -0.9888642 | 0.69880166 | 0.77355312 | -0.3661741 | 0.7193226 | 0.99999785 | -6.3174566 | down |
| KI - WT, cell_origin: iPSC | P38159 | RBMX | 22 | Homo sapien | -0.2290354 | -1.56278 | 1.10470909 | 2.91695613 | -0.365959 | 0.71949583 | 0.99999785 | -6.3175248 | down |
| KI - WT, cell_origin: iPSC | P45880 | VDAC2 | 7 | Homo sapien | 0.53110556 | -2.56469 | 3.62690113 | -2.1480011 | 0.365604 | 0.71975529 | 0.99999785 | -6.3176582 | up |
| KI - WT, cell_origin: iPSC | Q9HCH0 | NCKAP5L | 3 | Homo sapien | -0.9391361 | -6.4168578 | 4.53858556 | 3.90005494 | -0.3653684 | 0.71992754 | 0.99999785 | -6.3177467 | down |
| KI - WT, cell_origin: iPSC | Q9NQZ5 | STARD7 | 6 | Homo sapien | -0.1619086 | -1.1064049 | 0.78258763 | 1.10896729 | -0.3653189 | 0.71996369 | 0.99999785 | -6.3177653 | down |
| KI - WT, cell_origin: iPSC | P26447 | S100A4 | 2 | Homo sapien | -0.4045642 | -2.7646116 | 1.95548311 | -3.1344493 | -0.3653164 | 0.71996555 | 0.99999785 | -6.3177662 | down |
| KI - WT, cell_origin: iPSC | P52943 | CRIP2 | 11 | Homo sapien | 0.25817769 | -1.247935 | 1.76429041 | 1.88520973 | 0.36531172 | 0.71996895 | 0.99999785 | -6.317768 | up |
| KI - WT, cell_origin: iPSC | Q9Y3Y2 | CHTOP | 2 | Homo sapien | 0.18839032 | -0.9114119 | 1.28819259 | -1.7468364 | 0.36491536 | 0.72023818 | 0.99999785 | -6.3179329 | up |
| KI - WT, cell_origin: iPSC | Q96SI1 | KCTD15 | 3 | Homo sapien | 0.31263144 | -1.5131711 | 2.138434 | -2.4002125 | 0.36490617 | 0.72026546 | 0.99999785 | -6.3179201 | up |
| KI - WT, cell_origin: iPSC | Q7KZI7 | MARK2 | 4 | Homo sapien | -0.1397379 | -0.9567811 | 0.67730534 | -1.0333497 | -0.3642831 | 0.72069012 | 0.99999785 | -6.3181779 | down |
| KI - WT, cell_origin: iPSC | Q8TEW0 | PARD3 | 15 | Homo sapien | 0.16869391 | -0.8182258 | 1.15561366 | -0.7365206 | 0.36425304 | 0.72074083 | 0.99999785 | -6.3181665 | up |
| KI - WT, cell_origin: iPSC | P09086 | POU2F2 | 3 | Homo sapien | 0.10582589 | -0.5129829 | 0.72463466 | -1.7867816 | 0.36422952 | 0.72072532 | 0.99999785 | -6.3182012 | up |
| KI - WT, cell_origin: iPSC | P01116 | KRAS | 11 | Homo sapien | 0.23389162 | -1.1372299 | 1.60501315 | 3.67398202 | 0.36353068 | 0.72127144 | 0.99999785 | -6.3184349 | up |
| KI - WT, cell_origin: iPSC | Q96KG9 | SCYL1 | 22 | Homo sapien | 0.24448067 | -1.1896592 | 1.67862055 | 0.54386944 | 0.36329163 | 0.72144633 | 0.99999785 | -6.3185242 | up |
| KI - WT, cell_origin: iPSC | Q6UWE0 | LRSAM1 | 7 | Homo sapien | 0.08886363 | -0.4324706 | 0.61019783 | -0.7995974 | 0.36303419 | 0.72159986 | 0.99999785 | -6.3186477 | up |
| KI - WT, cell_origin: iPSC | P35908 | KRT2 | 35 | Homo sapien | -0.2428334 | -1.66858 | 1.1829132 | 4.66530628 | -0.3629681 | 0.72168305 | 0.99999785 | -6.3186449 | down |
| KI - WT, cell_origin: iPSC | Q9BUA3 | SPINDOC | 3 | Homo sapien | 0.1943047 | -0.9463973 | 1.33500666 | -1.8926822 | 0.36287686 | 0.72172938 | 0.99999785 | -6.318695 | up |
| KI - WT, cell_origin: iPSC | Q86X95 | CIR1 | 6 | Homo sapien | -0.2280001 | -1.5686276 | 1.11262735 | -0.9767252 | -0.3624344 | 0.72207362 | 0.99999785 | -6.3188438 | down |
| KI - WT, cell_origin: iPSC | Q9UHI6 | DDX20 | 17 | Homo sapien | -0.2199272 | -1.5135333 | 1.07367881 | 3.33171175 | -0.3623092 | 0.72216525 | 0.99999785 | -6.3188905 | down |
| KI - WT, cell_origin: iPSC | O15212 | PFDN6 | 6 | Homo sapien | 0.12042988 | -0.5877352 | 0.828595 | 0.15742124 | 0.36219245 | 0.72221595 | 0.99999785 | -6.3189612 | up |
| KI - WT, cell_origin: iPSC | Q16629 | SRSF7 | 10 | Homo sapien | -0.2979506 | -2.0546266 | 1.45872533 | 2.54829833 | -0.3614557 | 0.72279004 | 0.99999785 | -6.3192079 | down |
| KI - WT, cell_origin: iPSC | O96013 | PAK4 | 20 | Homo sapien | 0.20164295 | -0.9876672 | 1.39095314 | 1.21986408 | 0.36131857 | 0.72289041 | 0.99999785 | -6.3192588 | up |
| KI - WT, cell_origin: iPSC | Q15631 | TSN | 4 | Homo sapien | 0.13321007 | -0.6529108 | 0.91933095 | -1.3319102 | 0.36090045 | 0.72316198 | 0.99999785 | -6.3194411 | up |
| KI - WT, cell_origin: iPSC | Q13555 | CAMK2G | 3 | Homo sapien | 0.10468104 | -0.5138448 | 0.72320683 | -0.7857486 | 0.36045403 | 0.72348896 | 0.99999785 | -6.3196065 | up |
| KI - WT, cell_origin: iPSC | Q92797 | SYMPK | 12 | Homo sapien | -0.2638594 | -1.8251471 | 1.29742835 | -1.6528431 | -0.3601571 | 0.72374103 | 0.99999785 | -6.3196894 | down |
| KI - WT, cell_origin: iPSC | Q9Y597 | KCTD3 | 10 | Homo sapien | -0.1113329 | -0.7711634 | 0.54849763 | 2.12170341 | -0.3593609 | 0.7242899 | 0.99999785 | -6.3200107 | down |
| KI - WT, cell_origin: iPSC | Q9Y5X1 | SNX9 | 19 | Homo sapien | 0.17059841 | -0.841474 | 1.1826708 | 2.35769264 | 0.35922435 | 0.72442437 | 0.99999785 | -6.3200342 | up |
| KI - WT, cell_origin: iPSC | P15153 | RAC2 | 6 | Homo sapien | 0.21118533 | -1.0434772 | 1.46584787 | 4.43913967 | 0.35870645 | 0.7248039 | 0.99999785 | -6.3202253 | up |
| KI - WT, cell_origin: iPSC | Q9BTE3 | MCMBP | 22 | Homo sapien | -0.1426182 | -0.9910076 | 0.70577133 | 0.90393753 | -0.3581476 | 0.7251979 | 0.99999785 | -6.3204434 | down |
| KI - WT, cell_origin: iPSC | Q8WWK9 | CKAP2 | 6 | Homo sapien | 0.16043045 | -0.7939887 | 1.11484962 | -2.1148156 | 0.35800355 | 0.72528483 | 0.99999785 | -6.320511 | up |
| KI - WT, cell_origin: iPSC | Q96H79 | ZC3HAV1L | 10 | Homo sapien | -0.0884891 | -0.6158254 | 0.43884715 | 0.32284531 | -0.3573896 | 0.72573501 | 0.99999785 | -6.3207366 | down |
| KI - WT, cell_origin: iPSC | Q3B726 | POLR1F | 6 | Homo sapien | -0.3769275 | -2.6258477 | 1.8719928 | -1.2684173 | -0.3571791 | 0.72592359 | 0.99999785 | -6.3207873 | down |
| KI - WT, cell_origin: iPSC | Q9NY12 | GAR1 | 6 | Homo sapien | 0.15867045 | -0.7885047 | 1.10584558 | 0.9918906 | 0.35699993 | 0.72605502 | 0.99999785 | -6.320853 | up |
| KI - WT, cell_origin: iPSC | Q9H0Q0 | CYRIA | 10 | Homo sapien | 0.19301373 | -0.9603942 | 1.34642162 | 1.73878544 | 0.3566216 | 0.7263325 | 0.99999785 | -6.3209918 | up |

| | | | | | | | | | | | | |
|---|---|---|---|---|---|---|---|---|---|---|---|---|
| KI - WT, cell_origin: iPSC | Q9P270 | SLAIN2 | 5 | Homo sapien | 0.15403401 | -0.7662013 | 1.07426932 | -0.7507298 | 0.35661549 | 0.7263214 | 0.99999785 | -6.3210062 up |
| KI - WT, cell_origin: iPSC | Q9UPU5 | USP24 | 46 | Homo sapien | 0.12622266 | -0.6283129 | 0.88075822 | 0.57640639 | 0.35633073 | 0.72651902 | 0.99999785 | -6.3211192 up |
| KI - WT, cell_origin: iPSC | P07203 | GPX1 | 10 | Homo sapien | 0.12135141 | -0.6047575 | 0.84746033 | 0.94085494 | 0.35597044 | 0.72678014 | 0.99999785 | -6.3212536 up |
| KI - WT, cell_origin: iPSC | Q15418 | RPS6KA1 | 25 | Homo sapien | -0.1346359 | -0.9414112 | 0.67213943 | -0.4040825 | -0.3554799 | 0.72714479 | 0.99999785 | -6.3214293 down |
| KI - WT, cell_origin: iPSC | Q9NRZ9 | HELLS | 20 | Homo sapien | 0.19761032 | -0.9873449 | 1.38256552 | -1.1354857 | 0.35539398 | 0.72723315 | 0.99999785 | -6.3214411 up |
| KI - WT, cell_origin: iPSC | Q8WVY7 | UBLCP1 | 3 | Homo sapien | -0.1857622 | -1.3003079 | 0.92878343 | -1.4312387 | -0.3551649 | 0.72739712 | 0.99999785 | -6.3215279 down |
| KI - WT, cell_origin: iPSC | Q9H8Y5 | ANKZF1 | 5 | Homo sapien | 0.2323106 | -1.1623972 | 1.62701839 | -1.7948166 | 0.35496716 | 0.72754639 | 0.99999785 | -6.3215969 up |
| KI - WT, cell_origin: iPSC | O75362 | ZNF217 | 9 | Homo sapien | 0.31673796 | -1.5853673 | 2.21884326 | -2.4225479 | 0.35486889 | 0.72761851 | 0.99999785 | -6.3216328 up |
| KI - WT, cell_origin: iPSC | P28482 | MAPK1 | 10 | Homo sapien | -0.0806499 | -0.5646911 | 0.40339137 | 1.12485763 | -0.3548633 | 0.72758867 | 0.99999785 | -6.3216611 down |
| KI - WT, cell_origin: iPSC | P62875 | POLR2L | 2 | Homo sapien | -0.1592615 | -1.1158307 | 0.79730775 | -0.5303734 | -0.3547846 | 0.72767624 | 0.99999785 | -6.3216667 down |
| KI - WT, cell_origin: iPSC | O95278 | EPM2A | 9 | Homo sapien | 0.23501851 | -1.1766886 | 1.64672565 | -0.1559339 | 0.35478056 | 0.72768334 | 0.99999785 | -6.321665 up |
| KI - WT, cell_origin: iPSC | P40429 | RPL13A | 11 | Homo sapien | -0.0855789 | -0.5999348 | 0.42877696 | 2.44836999 | -0.3543584 | 0.72795933 | 0.99999785 | -6.3218451 down |
| KI - WT, cell_origin: iPSC | Q8NFP7;Q96 | NUDT10;NUI | 3 | Homo sapien | -0.1252473 | -0.8795205 | 0.62902601 | -0.9278794 | -0.3536545 | 0.72847618 | 0.99999785 | -6.3221012 down |
| KI - WT, cell_origin: iPSC | P46063 | RECQL | 16 | Homo sapien | -0.1290274 | -0.9065134 | 0.64845857 | -0.0181462 | -0.3534965 | 0.72859946 | 0.99999785 | -6.3221531 down |
| KI - WT, cell_origin: iPSC | Q9NRN7 | AASDHPPT | 6 | Homo sapien | 0.09775638 | -0.4916104 | 0.6871232 | -1.5897673 | 0.35326379 | 0.72876316 | 0.99999785 | -6.3222431 up |
| KI - WT, cell_origin: iPSC | O15144 | ARPC2 | 5 | Homo sapien | -0.1357485 | -0.9544286 | 0.68293146 | -1.0437386 | -0.3531758 | 0.72883169 | 0.99999785 | -6.3222721 down |
| KI - WT, cell_origin: iPSC | Q9NZ09 | UBAP1 | 9 | Homo sapien | 0.13876034 | -0.6998148 | 0.9773355 | -0.3634813 | 0.3524855 | 0.72934495 | 0.99999785 | -6.3225177 up |
| KI - WT, cell_origin: iPSC | P33981 | TTK | 23 | Homo sapien | 0.28641114 | -1.4454765 | 2.01829874 | -0.8459028 | 0.35242976 | 0.72940957 | 0.99999785 | -6.3225196 up |
| KI - WT, cell_origin: iPSC | P22314 | UBA1 | 23 | Homo sapien | -0.1187875 | -0.8389194 | 0.60134447 | 0.78518218 | -0.3513408 | 0.73018003 | 0.99999785 | -6.3229365 down |
| KI - WT, cell_origin: iPSC | P06744 | GPI | 6 | Homo sapien | -0.2236203 | -1.5806615 | 1.13342079 | -0.5521651 | -0.3511726 | 0.73033332 | 0.99999785 | -6.3229744 down |
| KI - WT, cell_origin: iPSC | O15160 | POLR1C | 6 | Homo sapien | 0.20281749 | -1.0277037 | 1.43333866 | -2.046058 | 0.35115517 | 0.73033087 | 0.99999785 | -6.3229924 up |
| KI - WT, cell_origin: iPSC | Q15007 | WTAP | 7 | Homo sapien | -0.1500472 | -1.0607871 | 0.76069274 | -0.9449976 | -0.350955 | 0.73046974 | 0.99999785 | -6.323071 down |
| KI - WT, cell_origin: iPSC | Q9Y3B2 | EXOSC1 | 4 | Homo sapien | -0.1196332 | -0.8477853 | 0.60851894 | -1.0939245 | -0.3499204 | 0.73122049 | 0.99999785 | -6.3234514 down |
| KI - WT, cell_origin: iPSC | P23381 | WARS1 | 5 | Homo sapien | -0.2116873 | -1.5014361 | 1.07806158 | -0.9928327 | -0.3497776 | 0.73135894 | 0.99999785 | -6.3234771 down |
| KI - WT, cell_origin: iPSC | Q01130 | SRSF2 | 3 | Homo sapien | 0.13217683 | -0.6729229 | 0.93727654 | 1.21941867 | 0.34972249 | 0.73137597 | 0.99999785 | -6.3235149 up |
| KI - WT, cell_origin: iPSC | Q12788 | TBL3 | 12 | Homo sapien | 0.24488847 | -1.2476077 | 1.73738462 | -0.5276134 | 0.34966928 | 0.73143857 | 0.99999785 | -6.3235161 up |
| KI - WT, cell_origin: iPSC | P06737 | PYGL | 24 | Homo sapien | 0.12442368 | -0.635628 | 0.8844754 | 0.83743101 | 0.34871184 | 0.73211798 | 0.99999785 | -6.3238788 up |
| KI - WT, cell_origin: iPSC | Q9NQT8 | KIF13B | 8 | Homo sapien | 0.16799806 | -0.858549 | 1.19454513 | -1.5114686 | 0.34861278 | 0.73219227 | 0.99999785 | -6.3239132 up |
| KI - WT, cell_origin: iPSC | P46776 | RPL27A | 5 | Homo sapien | -0.0819643 | -0.5831488 | 0.41922023 | 5.13384514 | -0.3483105 | 0.73240476 | 0.99999785 | -6.3240291 down |
| KI - WT, cell_origin: iPSC | Q9H8V3 | ECT2 | 20 | Homo sapien | 0.25757945 | -1.3208117 | 1.83597061 | -1.7299973 | 0.34777547 | 0.73283175 | 0.99999785 | -6.3241952 up |
| KI - WT, cell_origin: iPSC | Q02543 | RPL18A | 12 | Homo sapien | -0.0886486 | -0.631554 | 0.45425671 | 3.39626059 | -0.3477664 | 0.73280522 | 0.99999785 | -6.3242238 down |
| KI - WT, cell_origin: iPSC | P62244 | RPS15A | 11 | Homo sapien | 0.06853031 | -0.3513416 | 0.48840221 | 5.12494275 | 0.34762062 | 0.7329125 | 0.99999785 | -6.3242759 up |
| KI - WT, cell_origin: iPSC | Q13472 | TOP3A | 6 | Homo sapien | -0.150859 | -1.077234 | 0.77551612 | 0.04045396 | -0.3470022 | 0.73339402 | 0.99999785 | -6.3244767 down |
| KI - WT, cell_origin: iPSC | P43686 | PSMC4 | 5 | Homo sapien | 0.12294152 | -0.6318415 | 0.87772451 | -0.2241697 | 0.34693252 | 0.73342268 | 0.99999785 | -6.3245188 up |
| KI - WT, cell_origin: iPSC | P61604 | HSPE1 | 3 | Homo sapien | 0.29500534 | -1.5173929 | 2.10740353 | 0.04596988 | 0.34687949 | 0.73349121 | 0.99999785 | -6.3245152 up |
| KI - WT, cell_origin: iPSC | Q9Y3Q3 | TMED3 | 3 | Homo sapien | 0.17604711 | -0.9057138 | 1.25780806 | -0.1431134 | 0.34681668 | 0.73353745 | 0.99999785 | -6.3245376 up |
| KI - WT, cell_origin: iPSC | Q9Y484 | WDR45 | 2 | Homo sapien | 0.15314851 | -0.7875067 | 1.09380377 | -1.7158108 | 0.34676503 | 0.73354404 | 0.99999785 | -6.32458 up |
| KI - WT, cell_origin: iPSC | Q8N6H7 | ARFGAP2 | 4 | Homo sapien | -0.1653017 | -1.1815229 | 0.8509196 | -0.8357495 | -0.3466501 | 0.7336601 | 0.99999785 | -6.324597 down |
| KI - WT, cell_origin: iPSC | P53041 | PPP5C | 2 | Homo sapien | 0.22857131 | -1.1776961 | 1.63483872 | -2.4105711 | 0.34633215 | 0.7338862 | 0.99999785 | -6.3247164 up |
| KI - WT, cell_origin: iPSC | Q504Q3 | PAN2 | 23 | Homo sapien | 0.11072801 | -0.5711818 | 0.79263783 | 0.73913339 | 0.34583604 | 0.73422647 | 0.99999785 | -6.324912 up |
| KI - WT, cell_origin: iPSC | Q13098 | GPS1 | 6 | Homo sapien | 0.12714759 | -0.6562139 | 0.91050906 | -1.8134737 | 0.34568904 | 0.73433475 | 0.99999785 | -6.3249642 up |
| KI - WT, cell_origin: iPSC | Q9BXP5 | SRRT | 23 | Homo sapien | -0.1584328 | -1.1359564 | 0.81909075 | 0.60466969 | -0.3453984 | 0.73458184 | 0.99999785 | -6.3250425 down |
| KI - WT, cell_origin: iPSC | P05107 | ITGB2 | 13 | Homo sapien | -0.166377 | -1.194254 | 0.86150003 | 0.04789938 | -0.3449486 | 0.73491311 | 0.99999785 | -6.3252022 down |
| KI - WT, cell_origin: iPSC | Q8IXH7 | NELFCD | 2 | Homo sapien | 0.12234058 | -0.633169 | 0.87785016 | -1.5810959 | 0.34488176 | 0.73492946 | 0.99999785 | -6.3252508 up |
| KI - WT, cell_origin: iPSC | P25787 | PSMA2 | 6 | Homo sapien | 0.2067239 | -1.0713172 | 1.48476504 | -1.0617456 | 0.3447055 | 0.73509222 | 0.99999785 | -6.3252884 up |
| KI - WT, cell_origin: iPSC | Q9NW13 | RBM28 | 9 | Homo sapien | -0.1993572 | -1.4318293 | 1.03311499 | -2.0854853 | -0.3446174 | 0.73514213 | 0.99999785 | -6.325331 down |

| | | | | | | | | | | | | |
|---|---|---|---|---|---|---|---|---|---|---|---|---|
| KI - WT, cell_origin: iPSC | Q8IYM9 | TRIM22 | 7 | Homo sapien | -0.1787316 | -1.2850475 | 0.9275843 | 0.69036004 | -0.3442901 | 0.7353983 | 0.99999785 | -6.3254356 down |
| KI - WT, cell_origin: iPSC | Q7Z4V5 | HDGFL2 | 10 | Homo sapien | -0.187818 | -1.3513102 | 0.97567415 | -0.1081517 | -0.3440141 | 0.73560169 | 0.99999785 | -6.3255333 down |
| KI - WT, cell_origin: iPSC | O43379 | WDR62 | 2 | Homo sapien | 0.18789832 | -0.9773026 | 1.35309925 | -1.3854104 | 0.34365641 | 0.73586528 | 0.99999785 | -6.3256598 up |
| KI - WT, cell_origin: iPSC | Q92963 | RIT1 | 2 | Homo sapien | -0.2032856 | -1.465337 | 1.05876578 | -1.9446295 | -0.3432418 | 0.73616693 | 0.99999785 | -6.3258093 down |
| KI - WT, cell_origin: iPSC | P26038 | MSN | 21 | Homo sapien | -0.1847107 | -1.3341451 | 0.96472372 | -0.7766961 | -0.3424603 | 0.73674704 | 0.99999785 | -6.3260819 down |
| KI - WT, cell_origin: iPSC | Q8WVP5 | TNFAIP8L1 | 5 | Homo sapien | 0.2229058 | -1.1642657 | 1.61007733 | -1.34737 | 0.34244717 | 0.7367567 | 0.99999785 | -6.3260865 up |
| KI - WT, cell_origin: iPSC | P18031 | PTPN1 | 8 | Homo sapien | -0.2737291 | -1.9777094 | 1.43025123 | -1.2801046 | -0.3423408 | 0.7368351 | 0.99999785 | -6.3261239 down |
| KI - WT, cell_origin: iPSC | Q8IUC4 | RHPN2 | 21 | Homo sapien | 0.21444491 | -1.1228362 | 1.55172606 | 0.44828077 | 0.34173967 | 0.73727844 | 0.99999785 | -6.3263355 up |
| KI - WT, cell_origin: iPSC | P19447 | ERCC3 | 3 | Homo sapien | 0.15236735 | -0.7977038 | 1.10243852 | -0.9674298 | 0.34167897 | 0.73730833 | 0.99999785 | -6.326368 up |
| KI - WT, cell_origin: iPSC | Q86VV8 | RTTN | 8 | Homo sapien | 0.17109207 | -0.8967264 | 1.23891058 | 1.00736192 | 0.34145605 | 0.73748762 | 0.99999785 | -6.3264351 up |
| KI - WT, cell_origin: iPSC | Q15637 | SF1 | 10 | Homo sapien | -0.1760384 | -1.2752684 | 0.92319164 | 1.11392959 | -0.3412881 | 0.73761148 | 0.99999785 | -6.3264941 down |
| KI - WT, cell_origin: iPSC | Q05209 | PTPN12 | 14 | Homo sapien | 0.12002931 | -0.6300671 | 0.8701257 | -0.1358406 | 0.34082996 | 0.73792042 | 0.99999785 | -6.3266766 up |
| KI - WT, cell_origin: iPSC | P98160 | HSPG2 | 69 | Homo sapien | 0.50922909 | -2.6758075 | 3.69426572 | -1.7481178 | 0.34072276 | 0.73802857 | 0.99999785 | -6.3266924 up |
| KI - WT, cell_origin: iPSC | Q15582 | TGFBI | 12 | Homo sapien | -0.2862083 | -2.077465 | 1.50504827 | -0.8856445 | -0.3405076 | 0.7381873 | 0.99999785 | -6.3267678 down |
| KI - WT, cell_origin: iPSC | Q9NVD7 | PARVA | 8 | Homo sapien | -0.1788473 | -1.298325 | 0.94063033 | -2.0200997 | -0.340318 | 0.73830444 | 0.99999785 | -6.3268513 down |
| KI - WT, cell_origin: iPSC | Q9NR46 | SH3GLB2 | 8 | Homo sapien | -0.1073243 | -0.7793813 | 0.56473278 | -0.6161674 | -0.3401194 | 0.73844133 | 0.99999785 | -6.326928 down |
| KI - WT, cell_origin: iPSC | P11021 | HSPA5 | 20 | Homo sapien | 0.21810707 | -1.1487358 | 1.58494995 | 1.3305111 | 0.34005841 | 0.73851879 | 0.99999785 | -6.3269251 up |
| KI - WT, cell_origin: iPSC | Q92734 | TFG | 4 | Homo sapien | 0.20930602 | -1.1031855 | 1.52179753 | -0.6893467 | 0.33985025 | 0.73867241 | 0.99999785 | -6.3269979 up |
| KI - WT, cell_origin: iPSC | Q9H0H0 | INTS2 | 4 | Homo sapien | 0.17162179 | -0.9041446 | 1.2473882 | -2.5615967 | 0.33977713 | 0.73869399 | 0.99999785 | -6.3270477 up |
| KI - WT, cell_origin: iPSC | P35241 | RDX | 16 | Homo sapien | 0.18326455 | -0.9663961 | 1.33292521 | 0.99619619 | 0.33971221 | 0.73877429 | 0.99999785 | -6.3270461 up |
| KI - WT, cell_origin: iPSC | Q9BUL8 | PDCD10 | 4 | Homo sapien | 0.12700227 | -0.6699677 | 0.92397225 | -1.0944092 | 0.33939794 | 0.73897391 | 0.99999785 | -6.3271801 up |
| KI - WT, cell_origin: iPSC | P10155 | RO60 | 10 | Homo sapien | -0.122233 | -0.8901937 | 0.64572758 | -0.5177616 | -0.3390155 | 0.73925997 | 0.99999785 | -6.3273107 down |
| KI - WT, cell_origin: iPSC | P55209 | NAP1L1 | 6 | Homo sapien | 0.13582069 | -0.7190645 | 0.99070586 | 2.03992933 | 0.33848591 | 0.73966486 | 0.99999785 | -6.3274849 up |
| KI - WT, cell_origin: iPSC | P12004 | PCNA | 6 | Homo sapien | 0.2314661 | -1.2277812 | 1.69071344 | -1.1529392 | 0.33803436 | 0.74001303 | 0.99999785 | -6.3276311 up |
| KI - WT, cell_origin: iPSC | P30533 | LRPAP1 | 10 | Homo sapien | -0.3140588 | -2.2942165 | 1.66609902 | -1.6090694 | -0.3379975 | 0.74004025 | 0.99999785 | -6.3276439 down |
| KI - WT, cell_origin: iPSC | P09651 | HNRNPA1 | 24 | Homo sapien | -0.1758874 | -1.2873857 | 0.93561081 | 4.49519938 | -0.3372317 | 0.7406059 | 0.99999785 | -6.3279099 down |
| KI - WT, cell_origin: iPSC | O43156 | TTI1 | 6 | Homo sapien | 0.13886973 | -0.7382976 | 1.01603702 | -1.8076634 | 0.33718235 | 0.74061022 | 0.99999785 | -6.3279509 up |
| KI - WT, cell_origin: iPSC | Q7Z6Z7 | HUWE1 | 20 | Homo sapien | 0.15602144 | -0.8310849 | 1.1431278 | 1.51304901 | 0.33683925 | 0.74089582 | 0.99999785 | -6.328046 up |
| KI - WT, cell_origin: iPSC | Q9BQ39 | DDX50 | 14 | Homo sapien | 0.2392383 | -1.2754783 | 1.75395493 | -1.7999712 | 0.33659036 | 0.74107971 | 0.99999785 | -6.3281322 up |
| KI - WT, cell_origin: iPSC | Q9H3P7 | ACBD3 | 5 | Homo sapien | 0.2233974 | -1.1941728 | 1.64096756 | -1.1438862 | 0.33584271 | 0.74163222 | 0.99999785 | -6.3283909 up |
| KI - WT, cell_origin: iPSC | Q9Y366 | IFT52 | 4 | Homo sapien | 0.08737508 | -0.4675722 | 0.64232232 | -0.6697259 | 0.33533245 | 0.74197745 | 0.99999785 | -6.3285907 up |
| KI - WT, cell_origin: iPSC | Q9Y285 | FARSA | 20 | Homo sapien | -0.1382865 | -1.0173426 | 0.74076963 | 1.30728554 | -0.3351932 | 0.74210378 | 0.99999785 | -6.3286214 down |
| KI - WT, cell_origin: iPSC | P05783 | KRT18 | 21 | Homo sapien | 0.25871255 | -1.3881526 | 1.90557774 | 1.07203482 | 0.33478178 | 0.74241649 | 0.99999785 | -6.3287569 up |
| KI - WT, cell_origin: iPSC | Q9UBR2 | CTSZ | 2 | Homo sapien | 0.23533604 | -1.265272 | 1.73594405 | -1.8377698 | 0.33421317 | 0.74283695 | 0.99999785 | -6.3289526 up |
| KI - WT, cell_origin: iPSC | Q5VWQ0 | RSBN1 | 2 | Homo sapien | 0.18998519 | -1.021575 | 1.40154536 | -2.0218828 | 0.33408519 | 0.74291708 | 0.99999785 | -6.3290074 up |
| KI - WT, cell_origin: iPSC | Q9BRX2 | PELO | 15 | Homo sapien | -0.1018388 | -0.751097 | 0.54741944 | 2.58485202 | -0.3340684 | 0.74291218 | 0.99999785 | -6.3290259 down |
| KI - WT, cell_origin: iPSC | Q5T280 | SPOUT1 | 10 | Homo sapien | 0.12377113 | -0.6662147 | 0.91375691 | -0.9636251 | 0.33369194 | 0.7431914 | 0.99999785 | -6.3291547 up |
| KI - WT, cell_origin: iPSC | Q9HC44 | GPBP1L1 | 4 | Homo sapien | -0.1778555 | -1.3142111 | 0.95850007 | -2.4323189 | -0.3333487 | 0.74344532 | 0.99999785 | -6.3292724 down |
| KI - WT, cell_origin: iPSC | Q9H5V9 | STEEP1 | 7 | Homo sapien | -0.2324793 | -1.7211602 | 1.25620167 | -1.2993586 | -0.3328013 | 0.74388131 | 0.99999785 | -6.3294372 down |
| KI - WT, cell_origin: iPSC | O75934 | BCAS2 | 5 | Homo sapien | -0.2128579 | -1.5759959 | 1.15028002 | -0.8992651 | -0.3327763 | 0.7438998 | 0.99999785 | -6.3294458 down |
| KI - WT, cell_origin: iPSC | O75928 | PIAS2 | 3 | Homo sapien | -0.0942132 | -0.6973187 | 0.50889225 | -0.6885037 | -0.3327042 | 0.74392146 | 0.99999785 | -6.3294938 down |
| KI - WT, cell_origin: iPSC | O60664 | PLIN3 | 14 | Homo sapien | 0.18039106 | -0.9766357 | 1.3374178 | -0.0870908 | 0.33225688 | 0.74428416 | 0.99999785 | -6.3296235 up |
| KI - WT, cell_origin: iPSC | P30740 | SERPINB1 | 13 | Homo sapien | -0.1802075 | -1.3370526 | 0.97663755 | -0.5832201 | -0.331971 | 0.74449573 | 0.99999785 | -6.3297212 down |
| KI - WT, cell_origin: iPSC | O43516 | WIPF1 | 8 | Homo sapien | -0.3267506 | -2.4244315 | 1.77093035 | 0.49348917 | -0.3319551 | 0.74450751 | 0.99999785 | -6.3297267 down |
| KI - WT, cell_origin: iPSC | P28070 | PSMB4 | 4 | Homo sapien | 0.19048323 | -1.0324532 | 1.41341964 | -0.8479969 | 0.33193668 | 0.74452112 | 0.99999785 | -6.3297329 up |
| KI - WT, cell_origin: iPSC | Q9BY32 | ITPA | 5 | Homo sapien | 0.11304999 | -0.6125602 | 0.83866015 | -0.1003786 | 0.33182327 | 0.74457349 | 0.99999785 | -6.3297949 up |

| | | | | | | | | | | | | | |
|---|---|---|---|---|---|---|---|---|---|---|---|---|---|
| KI - WT, cell_origin: iPSC | Q9H6T3 | RPAP3 | 8 | Homo sapien | 0.10823949 | -0.5867648 | 0.80324377 | -1.1696735 | 0.33169425 | 0.744669 | 0.99999785 | -6.3298389 | up |
| KI - WT, cell_origin: iPSC | Q9NSD9 | FARSB | 23 | Homo sapien | -0.1504915 | -1.1182498 | 0.81726669 | 1.73568402 | -0.3313962 | 0.74492115 | 0.99999785 | -6.3299174 | down |
| KI - WT, cell_origin: iPSC | O15360 | FANCA | 6 | Homo sapien | 0.24233709 | -1.3169162 | 1.80159041 | -1.8331247 | 0.33121163 | 0.74505781 | 0.99999785 | -6.3299804 | up |
| KI - WT, cell_origin: iPSC | P78312 | FAM193A | 6 | Homo sapien | 0.18355722 | -0.9978587 | 1.36497317 | -2.1462595 | 0.33096842 | 0.74521576 | 0.99999785 | -6.3300795 | up |
| KI - WT, cell_origin: iPSC | O95714 | HERC2 | 5 | Homo sapien | 0.21938186 | -1.1938953 | 1.63265906 | -1.7475968 | 0.33080779 | 0.74535679 | 0.99999785 | -6.3301179 | up |
| KI - WT, cell_origin: iPSC | P78537 | BLOC1S1 | 2 | Homo sapien | 0.15667807 | -0.8524039 | 1.16576003 | -1.3696898 | 0.33077292 | 0.74536415 | 0.99999785 | -6.3301434 | up |
| KI - WT, cell_origin: iPSC | P48059 | LIMS1 | 5 | Homo sapien | -0.1487759 | -1.1087216 | 0.81116973 | 1.32732478 | -0.3302846 | 0.74574418 | 0.99999785 | -6.3302959 | down |
| KI - WT, cell_origin: iPSC | Q16539 | MAPK14 | 6 | Homo sapien | -0.0889409 | -0.6635023 | 0.48562049 | -1.1577465 | -0.3296892 | 0.7461538 | 0.99999785 | -6.3305211 | down |
| KI - WT, cell_origin: iPSC | O75915 | ARL6IP5 | 2 | Homo sapien | -0.2055792 | -1.5354631 | 1.12430478 | 0.03702044 | -0.3294335 | 0.74637457 | 0.99999785 | -6.3305849 | down |
| KI - WT, cell_origin: iPSC | P58546 | MTPN | 5 | Homo sapien | -0.1140022 | -0.8511641 | 0.62315971 | 1.01513 | -0.3294079 | 0.74636745 | 0.99999785 | -6.3306126 | down |
| KI - WT, cell_origin: iPSC | P49916 | LIG3 | 19 | Homo sapien | -0.3294461 | -2.4615401 | 1.80264794 | -1.4529916 | -0.3292914 | 0.74647984 | 0.99999785 | -6.330633 | down |
| KI - WT, cell_origin: iPSC | Q9UDY8 | MALT1 | 8 | Homo sapien | -0.1719713 | -1.2880392 | 0.94409667 | -1.3807294 | -0.3283544 | 0.74717113 | 0.99999785 | -6.3309523 | down |
| KI - WT, cell_origin: iPSC | P54619 | PRKAG1 | 3 | Homo sapien | 0.14563854 | -0.8017252 | 1.09300226 | -0.2729252 | 0.32758957 | 0.74773713 | 0.99999785 | -6.3312111 | up |
| KI - WT, cell_origin: iPSC | P40222 | TXLNA | 6 | Homo sapien | -0.1683806 | -1.2641331 | 0.92737182 | -1.7841334 | -0.3273389 | 0.74790488 | 0.99999785 | -6.3313088 | down |
| KI - WT, cell_origin: iPSC | P49411 | TUFM | 8 | Homo sapien | -0.3600913 | -2.7087383 | 1.98855564 | -1.9533951 | -0.3267362 | 0.74837355 | 0.99999785 | -6.3314958 | down |
| KI - WT, cell_origin: iPSC | P49006 | MARCKSL1 | 4 | Homo sapien | 0.30773302 | -1.6995501 | 2.31501616 | 4.18765212 | 0.32671407 | 0.74838996 | 0.99999785 | -6.3315032 | up |
| KI - WT, cell_origin: iPSC | Q0VD83 | APOBR | 8 | Homo sapien | -0.2745317 | -2.0734615 | 1.52439808 | -1.6148163 | -0.3252226 | 0.74949613 | 0.99999785 | -6.3320037 | down |
| KI - WT, cell_origin: iPSC | Q13356 | PPIL2 | 15 | Homo sapien | -0.1968114 | -1.4868461 | 1.0932233 | -0.743634 | -0.3251257 | 0.74956803 | 0.99999785 | -6.3320361 | down |
| KI - WT, cell_origin: iPSC | Q8NFW8 | CMAS | 8 | Homo sapien | 0.17726882 | -0.9849254 | 1.33946307 | 0.02192656 | 0.32505437 | 0.74962092 | 0.99999785 | -6.33206 | up |
| KI - WT, cell_origin: iPSC | Q8TAD8 | SNIP1 | 2 | Homo sapien | -0.2451352 | -1.8558678 | 1.36559725 | -2.9840673 | -0.3242757 | 0.75019042 | 0.99999785 | -6.3323262 | down |
| KI - WT, cell_origin: iPSC | Q969X6 | UTP4 | 6 | Homo sapien | 0.22993582 | -1.2831718 | 1.74304341 | -2.8137882 | 0.32376853 | 0.75056284 | 0.99999785 | -6.3324983 | up |
| KI - WT, cell_origin: iPSC | Q9NPJ6 | MED4 | 6 | Homo sapien | 0.16111585 | -0.9023466 | 1.22457829 | -2.1496134 | 0.322686 | 0.7513509 | 0.99999785 | -6.3328697 | up |
| KI - WT, cell_origin: iPSC | Q9Y546 | LRRC42 | 2 | Homo sapien | 0.43823581 | -2.4583078 | 3.33477946 | -1.871568 | 0.32242607 | 0.75157169 | 0.99999785 | -6.332936 | up |
| KI - WT, cell_origin: iPSC | Q9Y6V7 | DDX49 | 5 | Homo sapien | -0.1229406 | -0.935168 | 0.68928684 | -0.7656033 | -0.3224135 | 0.75155693 | 0.99999785 | -6.3329575 | down |
| KI - WT, cell_origin: iPSC | Q9ULZ3 | PYCARD | 9 | Homo sapien | -0.1103429 | -0.8396758 | 0.61898993 | -0.2861474 | -0.3222244 | 0.75169086 | 0.99999785 | -6.3330249 | down |
| KI - WT, cell_origin: iPSC | Q8WYQ5 | DGCR8 | 6 | Homo sapien | 0.24207667 | -1.3593674 | 1.84352071 | -3.2549242 | 0.32200238 | 0.75186487 | 0.99999785 | -6.333092 | up |
| KI - WT, cell_origin: iPSC | Q9UN86 | G3BP2 | 19 | Homo sapien | -0.1783255 | -1.3587377 | 1.00208674 | 2.73041761 | -0.3219453 | 0.7519287 | 0.99999785 | -6.3330955 | down |
| KI - WT, cell_origin: iPSC | Q9HCM4 | EPB41L5 | 11 | Homo sapien | -0.1437361 | -1.0978936 | 0.8104213 | 0.03466913 | -0.321032 | 0.75260714 | 0.99999785 | -6.3333978 | down |
| KI - WT, cell_origin: iPSC | P49750 | YLPM1 | 15 | Homo sapien | -0.1167337 | -0.8912563 | 0.65778894 | -1.285093 | -0.3209976 | 0.75260222 | 0.99999785 | -6.333431 | down |
| KI - WT, cell_origin: iPSC | O75607 | NPM3 | 4 | Homo sapien | 0.11984225 | -0.676126 | 0.91581054 | -1.2745526 | 0.32066671 | 0.75284808 | 0.99999785 | -6.3335402 | up |
| KI - WT, cell_origin: iPSC | Q9Y467 | SALL2 | 11 | Homo sapien | 0.07131989 | -0.4026446 | 0.54528439 | 1.35790379 | 0.32048267 | 0.75298484 | 0.99999785 | -6.333601 | up |
| KI - WT, cell_origin: iPSC | Q15052 | ARHGEF6 | 34 | Homo sapien | 0.20651083 | -1.1678207 | 1.58084241 | 2.84141062 | 0.32022385 | 0.75320756 | 0.99999785 | -6.3336646 | up |
| KI - WT, cell_origin: iPSC | Q12802 | AKAP13 | 53 | Homo sapien | 0.23369813 | -1.3228275 | 1.79022377 | 0.42682873 | 0.31996416 | 0.75340056 | 0.99999785 | -6.3337502 | up |
| KI - WT, cell_origin: iPSC | P35268 | RPL22 | 4 | Homo sapien | -0.1300699 | -0.9968762 | 0.7367363 | 6.25450006 | -0.3197844 | 0.75353415 | 0.99999785 | -6.3338094 | down |
| KI - WT, cell_origin: iPSC | P29966 | MARCKS | 16 | Homo sapien | 0.29953034 | -1.6979707 | 2.29703138 | 6.44850746 | 0.31956277 | 0.75369888 | 0.99999785 | -6.3338823 | up |
| KI - WT, cell_origin: iPSC | Q9BUJ2 | HNRNPUL1 | 28 | Homo sapien | -0.1901833 | -1.4594344 | 1.07906778 | 3.37163072 | -0.3193208 | 0.75387871 | 0.99999785 | -6.3339619 | down |
| KI - WT, cell_origin: iPSC | Q9HAU5 | UPF2 | 12 | Homo sapien | -0.0966082 | -0.7414562 | 0.54823979 | 0.24638275 | -0.3190777 | 0.75402921 | 0.99999785 | -6.3340634 | down |
| KI - WT, cell_origin: iPSC | Q15654 | TRIP6 | 7 | Homo sapien | -0.1686397 | -1.2953409 | 0.95806138 | -0.704946 | -0.3189727 | 0.75413748 | 0.99999785 | -6.3340763 | down |
| KI - WT, cell_origin: iPSC | P27695 | APEX1 | 11 | Homo sapien | -0.2643927 | -2.0325151 | 1.50372979 | 0.50845983 | -0.3186687 | 0.7543635 | 0.99999785 | -6.3341761 | down |
| KI - WT, cell_origin: iPSC | Q5T8P6 | RBM26 | 11 | Homo sapien | -0.2711398 | -2.084489 | 1.5422094 | -0.5917653 | -0.3186502 | 0.75437729 | 0.99999785 | -6.3341822 | down |
| KI - WT, cell_origin: iPSC | O95359 | TACC2 | 6 | Homo sapien | 0.71742319 | -4.0833814 | 5.51822783 | -2.8458961 | 0.31846649 | 0.75451386 | 0.99999785 | -6.3342425 | up |
| KI - WT, cell_origin: iPSC | P84090 | ERH | 5 | Homo sapien | -0.1665771 | -1.2817776 | 0.94862341 | 3.28731718 | -0.3183205 | 0.7546224 | 0.99999785 | -6.3342903 | down |
| KI - WT, cell_origin: iPSC | O00267 | SUPT5H | 19 | Homo sapien | 0.15910115 | -0.9063713 | 1.22457361 | -0.3646037 | 0.3182244 | 0.75469388 | 0.99999785 | -6.3343218 | up |
| KI - WT, cell_origin: iPSC | Q09161 | NCBP1 | 16 | Homo sapien | -0.1383241 | -1.0650017 | 0.7883535 | 0.15286726 | -0.3180824 | 0.75479581 | 0.99999785 | -6.3343709 | down |
| KI - WT, cell_origin: iPSC | Q9BSL1 | UBAC1 | 4 | Homo sapien | -0.107796 | -0.8303653 | 0.61477333 | -0.7509689 | -0.3177333 | 0.75502895 | 0.99999785 | -6.334504 | down |
| KI - WT, cell_origin: iPSC | Q9BZK7 | TBL1XR1 | 5 | Homo sapien | -0.1661561 | -1.2812345 | 0.94892238 | -0.5089122 | -0.3175508 | 0.75519486 | 0.99999785 | -6.3345423 | down |

| | | | | | | | | | | | | | |
|---|---|---|---|---|---|---|---|---|---|---|---|---|---|
| KI - WT, cell_origin: iPSC | Q9H3Q1 | CDC42EP4 | 9 | Homo sapien | 0.19366047 | -1.1067047 | 1.49402567 | 1.15303547 | 0.31737887 | 0.75532272 | 0.99999785 | -6.3345985 | up |
| KI - WT, cell_origin: iPSC | P05141 | SLC25A5 | 2 | Homo sapien | 0.51355512 | -2.9366608 | 3.96377101 | -2.1904886 | 0.31720729 | 0.75545035 | 0.99999785 | -6.3346546 | up |
| KI - WT, cell_origin: iPSC | Q15029 | EFTUD2 | 40 | Homo sapien | -0.1078362 | -0.8320222 | 0.61634979 | 1.5511017 | -0.3171626 | 0.75545668 | 0.99999785 | -6.3346883 | down |
| KI - WT, cell_origin: iPSC | Q07157 | TJP1 | 59 | Homo sapien | -0.1878786 | -1.45052 | 1.07476275 | 1.84860491 | -0.3171025 | 0.75552827 | 0.99999785 | -6.3346888 | down |
| KI - WT, cell_origin: iPSC | P63313 | TMSB10 | 2 | Homo sapien | 0.18251396 | -1.0456606 | 1.41068856 | 1.92040926 | 0.31669291 | 0.75583301 | 0.99999785 | -6.3348225 | up |
| KI - WT, cell_origin: iPSC | Q9P2A4 | ABI3 | 2 | Homo sapien | -0.3806247 | -2.9473433 | 2.18609397 | -1.917871 | -0.3160247 | 0.75633021 | 0.99999785 | -6.3350402 | down |
| KI - WT, cell_origin: iPSC | P11908 | PRPS2 | 3 | Homo sapien | 0.28439404 | -1.6366463 | 2.20543435 | -2.7030636 | 0.31549057 | 0.75672773 | 0.99999785 | -6.3352139 | up |
| KI - WT, cell_origin: iPSC | Q9Y315 | DERA | 4 | Homo sapien | -0.1254303 | -0.9725269 | 0.72166638 | -0.7912722 | -0.3154155 | 0.75676216 | 0.99999785 | -6.3352535 | down |
| KI - WT, cell_origin: iPSC | Q13243 | SRSF5 | 9 | Homo sapien | 0.12100969 | -0.6976812 | 0.93970055 | 2.77298452 | 0.31490628 | 0.75714888 | 0.99999785 | -6.3354133 | up |
| KI - WT, cell_origin: iPSC | Q8IY31 | IFT20 | 2 | Homo sapien | 0.08375966 | -0.4828318 | 0.65035111 | -0.8727435 | 0.31485064 | 0.75717425 | 0.99999785 | -6.3354427 | up |
| KI - WT, cell_origin: iPSC | Q8N573 | OXR1 | 16 | Homo sapien | 0.29422466 | -1.7007872 | 2.28923656 | -0.3625277 | 0.3142939 | 0.75761858 | 0.99999785 | -6.335602 | up |
| KI - WT, cell_origin: iPSC | Q8TED1 | GPX8 | 5 | Homo sapien | 0.33049149 | -1.9125216 | 2.57350459 | -1.4378583 | 0.31400086 | 0.75783678 | 0.99999785 | -6.3356968 | up |
| KI - WT, cell_origin: iPSC | P06396 | GSN | 22 | Homo sapien | 0.1985936 | -1.150577 | 1.54776422 | 2.03446386 | 0.31369004 | 0.75806825 | 0.99999785 | -6.3357973 | up |
| KI - WT, cell_origin: iPSC | O60506 | SYNCRIP | 26 | Homo sapien | -0.053836 | -0.4197271 | 0.31205505 | 3.81619583 | -0.3133724 | 0.75827514 | 0.99999785 | -6.3359207 | down |
| KI - WT, cell_origin: iPSC | Q99496 | RNF2 | 5 | Homo sapien | 0.08742348 | -0.507185 | 0.682032 | -1.0711174 | 0.31313864 | 0.7584493 | 0.99999785 | -6.3359961 | up |
| KI - WT, cell_origin: iPSC | Q14145 | KEAP1 | 9 | Homo sapien | -0.1699118 | -1.3277689 | 0.98794534 | -0.7721997 | -0.312731 | 0.75878263 | 0.99999785 | -6.3361067 | down |
| KI - WT, cell_origin: iPSC | P18887 | XRCC1 | 9 | Homo sapien | 0.2772408 | -1.6120435 | 2.16652506 | -0.573741 | 0.31272471 | 0.75878729 | 0.99999785 | -6.3361087 | up |
| KI - WT, cell_origin: iPSC | Q0VG06 | FAAP100 | 4 | Homo sapien | 0.33833359 | -1.9732238 | 2.64989093 | -1.3178745 | 0.31191971 | 0.75938708 | 0.99999785 | -6.3363677 | up |
| KI - WT, cell_origin: iPSC | Q9H4A4 | RNPEP | 12 | Homo sapien | -0.1242118 | -0.9734966 | 0.72507295 | -0.2084781 | -0.3115498 | 0.75964201 | 0.99999785 | -6.3365009 | down |
| KI - WT, cell_origin: iPSC | Q9Y224 | RTRAF | 17 | Homo sapien | -0.1479665 | -1.1602999 | 0.86436696 | 3.8035091 | -0.3114886 | 0.75970839 | 0.99999785 | -6.3365061 | down |
| KI - WT, cell_origin: iPSC | Q7Z7C8 | TAF8 | 2 | Homo sapien | 0.13096981 | -0.7650975 | 1.02703717 | -1.4932003 | 0.31130853 | 0.75981543 | 0.99999785 | -6.3365828 | up |
| KI - WT, cell_origin: iPSC | Q8WUH6 | TMEM263 | 5 | Homo sapien | 0.28601538 | -1.6720549 | 2.24408561 | 0.52717933 | 0.31128879 | 0.75985728 | 0.99999785 | -6.3365702 | up |
| KI - WT, cell_origin: iPSC | P20810 | CAST | 19 | Homo sapien | 0.2951749 | -1.7276044 | 2.3179542 | -0.8549058 | 0.31098059 | 0.760087 | 0.99999785 | -6.3366689 | up |
| KI - WT, cell_origin: iPSC | Q8NHQ9 | DDX55 | 5 | Homo sapien | -0.1442366 | -1.1325991 | 0.84412603 | -1.8453975 | -0.3108343 | 0.76017001 | 0.99999785 | -6.336734 | down |
| KI - WT, cell_origin: iPSC | Q6P1X5 | TAF2 | 3 | Homo sapien | -0.1489525 | -1.1727815 | 0.87487651 | -0.9661777 | -0.3100435 | 0.7607856 | 0.99999785 | -6.3369687 | down |
| KI - WT, cell_origin: iPSC | P50454 | SERPINH1 | 19 | Homo sapien | -0.2724801 | -2.147636 | 1.60267575 | 0.1556221 | -0.3096705 | 0.76106379 | 0.99999785 | -6.3370877 | down |
| KI - WT, cell_origin: iPSC | Q9BYK8 | HELZ2 | 6 | Homo sapien | 0.14096516 | -0.828723 | 1.11065335 | -2.4363255 | 0.30961311 | 0.76107727 | 0.99999785 | -6.3371264 | up |
| KI - WT, cell_origin: iPSC | Q7Z3U7 | MON2 | 8 | Homo sapien | 0.11328949 | -0.6662157 | 0.89279466 | -1.5139664 | 0.30953537 | 0.76113525 | 0.99999785 | -6.3371512 | up |
| KI - WT, cell_origin: iPSC | Q9H000 | MKRN2 | 9 | Homo sapien | 0.09174037 | -0.5430323 | 0.72651302 | -0.1253039 | 0.30780946 | 0.76242292 | 0.99999785 | -6.3376998 | up |
| KI - WT, cell_origin: iPSC | Q16527 | CSRP2 | 10 | Homo sapien | -0.1998366 | -1.5835302 | 1.18385688 | 3.12698639 | -0.307778 | 0.76247551 | 0.99999785 | -6.3376896 | down |
| KI - WT, cell_origin: iPSC | P0DOX8 | | 2 | Homo sapien | -0.7434058 | -5.8980221 | 4.41121055 | -0.0306333 | -0.3073491 | 0.76279557 | 0.99999785 | -6.3378255 | down |
| KI - WT, cell_origin: iPSC | O14497 | ARID1A | 12 | Homo sapien | -0.1601614 | -1.27093 | 0.95060725 | -0.7181565 | -0.3072816 | 0.76284592 | 0.99999785 | -6.3378469 | down |
| KI - WT, cell_origin: iPSC | P46977 | STT3A | 4 | Homo sapien | 0.35546043 | -2.1110787 | 2.82199953 | -3.0504754 | 0.30711825 | 0.76296786 | 0.99999785 | -6.3378986 | up |
| KI - WT, cell_origin: iPSC | Q969T9 | WBP2 | 3 | Homo sapien | 0.13821451 | -0.821148 | 1.09757703 | 0.04463822 | 0.30702467 | 0.7630377 | 0.99999785 | -6.3379282 | up |
| KI - WT, cell_origin: iPSC | Q2KHR3 | QSER1 | 4 | Homo sapien | 0.1611497 | -0.9576214 | 1.27992084 | -2.9341998 | 0.30678063 | 0.76319085 | 0.99999785 | -6.3380254 | up |
| KI - WT, cell_origin: iPSC | P24941 | CDK2 | 4 | Homo sapien | -0.0939148 | -0.7462124 | 0.55838282 | -1.2036919 | -0.3066393 | 0.76329636 | 0.99999785 | -6.3380701 | down |
| KI - WT, cell_origin: iPSC | Q70CQ2 | USP34 | 9 | Homo sapien | 0.12509732 | -0.7443421 | 0.99453676 | -1.3087521 | 0.3064634 | 0.76343103 | 0.99999785 | -6.3381233 | up |
| KI - WT, cell_origin: iPSC | O75896 | TUSC2 | 2 | Homo sapien | 0.19719297 | -1.1749433 | 1.56932924 | -0.4662565 | 0.30626442 | 0.7636052 | 0.99999785 | -6.3381684 | up |
| KI - WT, cell_origin: iPSC | P50502 | ST13 | 5 | Homo sapien | -0.1262873 | -1.0054016 | 0.75282697 | 1.0115449 | -0.306093 | 0.76372622 | 0.99999785 | -6.3382273 | down |
| KI - WT, cell_origin: iPSC | P46821 | MAP1B | 35 | Homo sapien | -0.1519475 | -1.210048 | 0.90615305 | -1.0643052 | -0.3060335 | 0.7637776 | 0.99999785 | -6.3382412 | down |
| KI - WT, cell_origin: iPSC | O60684 | KPNA6 | 8 | Homo sapien | 0.16698483 | -0.9966314 | 1.33060105 | 0.02768272 | 0.30582262 | 0.76393506 | 0.99999785 | -6.3383077 | up |
| KI - WT, cell_origin: iPSC | O60610 | DIAPH1 | 25 | Homo sapien | -0.1996844 | -1.5912756 | 1.19190676 | -0.5798928 | -0.3057981 | 0.76395333 | 0.99999785 | -6.3383154 | down |
| KI - WT, cell_origin: iPSC | Q99460 | PSMD1 | 15 | Homo sapien | 0.17203724 | -1.0299118 | 1.37398629 | -1.4308967 | 0.30502734 | 0.76452895 | 0.99999785 | -6.338558 | up |
| KI - WT, cell_origin: iPSC | O00139 | KIF2A | 17 | Homo sapien | 0.07846469 | -0.4697142 | 0.62664358 | 0.77953001 | 0.30485384 | 0.76462972 | 0.99999785 | -6.3386323 | up |
| KI - WT, cell_origin: iPSC | Q02790 | FKBP4 | 7 | Homo sapien | -0.1773485 | -1.4171792 | 1.06248222 | -1.1047604 | -0.3048369 | 0.76467119 | 0.99999785 | -6.3386178 | down |
| KI - WT, cell_origin: iPSC | Q13428 | TCOF1 | 32 | Homo sapien | -0.2068287 | -1.6532283 | 1.23957095 | 0.98980551 | -0.3047367 | 0.76474601 | 0.99999785 | -6.3386493 | down |

| | | | | | | | | | | | | | |
|---|---|---|---|---|---|---|---|---|---|---|---|---|---|
| KI - WT, cell_origin: iPSC | O95983 | MBD3 | 2 | Homo sapien | 0.13535335 | -0.8117934 | 1.08250009 | -2.2067011 | 0.30436269 | 0.76499664 | 0.99999785 | -6.3387864 | up |
| KI - WT, cell_origin: iPSC | P09429 | HMGB1 | 10 | Homo sapien | -0.2740159 | -2.1928744 | 1.64484269 | 0.76949497 | -0.3043232 | 0.76505487 | 0.99999785 | -6.338779 | down |
| KI - WT, cell_origin: iPSC | O95433 | AHSA1 | 2 | Homo sapien | 0.08857395 | -0.5318491 | 0.70899701 | -0.3082935 | 0.30405892 | 0.76522361 | 0.99999785 | -6.3388816 | up |
| KI - WT, cell_origin: iPSC | Q9NUJ1 | ABHD10 | 7 | Homo sapien | -0.4316511 | -3.4575193 | 2.59421717 | -1.1922542 | -0.3040081 | 0.76529028 | 0.99999785 | -6.3388778 | down |
| KI - WT, cell_origin: iPSC | Q9Y2L9 | LRCH1 | 8 | Homo sapien | 0.12523717 | -0.7551441 | 1.00561842 | -0.2585564 | 0.30306722 | 0.76597963 | 0.99999785 | -6.3391814 | up |
| KI - WT, cell_origin: iPSC | Q7Z3E5 | ARMC9 | 3 | Homo sapien | 0.12540172 | -0.7562103 | 1.00701379 | -1.26944 | 0.30297701 | 0.76603694 | 0.99999785 | -6.3392165 | up |
| KI - WT, cell_origin: iPSC | Q9HD67 | MYO10 | 26 | Homo sapien | 0.10038613 | -0.6053288 | 0.80610107 | -0.9503328 | 0.30295917 | 0.76604548 | 0.99999785 | -6.3392254 | up |
| KI - WT, cell_origin: iPSC | Q9NZN8 | CNOT2 | 9 | Homo sapien | -0.0706428 | -0.5676425 | 0.42635682 | 0.37963421 | -0.3027274 | 0.76621872 | 0.99999785 | -6.3392976 | down |
| KI - WT, cell_origin: iPSC | Q9Y6E2 | BZW2 | 7 | Homo sapien | -0.1377978 | -1.1080543 | 0.83245866 | -1.4738937 | -0.3025259 | 0.76637665 | 0.99999785 | -6.3393555 | down |
| KI - WT, cell_origin: iPSC | P11586 | MTHFD1 | 37 | Homo sapien | 0.10000905 | -0.604318 | 0.80433612 | 1.28627452 | 0.30241589 | 0.7664516 | 0.99999785 | -6.3393947 | up |
| KI - WT, cell_origin: iPSC | O95985 | TOP3B | 4 | Homo sapien | 0.11762466 | -0.7109797 | 0.946229 | -1.0385419 | 0.30236734 | 0.76649268 | 0.99999785 | -6.3394066 | up |
| KI - WT, cell_origin: iPSC | Q9P253 | VPS18 | 16 | Homo sapien | -0.0709225 | -0.5705653 | 0.42872029 | 0.61615203 | -0.3023181 | 0.76652467 | 0.99999785 | -6.3394252 | down |
| KI - WT, cell_origin: iPSC | Q99733 | NAP1L4 | 6 | Homo sapien | 0.0979897 | -0.593399 | 0.78937836 | 0.00083679 | 0.30185465 | 0.76687121 | 0.99999785 | -6.3395694 | up |
| KI - WT, cell_origin: iPSC | E9PAV3;Q13 | NACA | 5 | Homo sapien | -0.1557264 | -1.2563941 | 0.94494141 | 1.93036101 | -0.3015145 | 0.76715401 | 0.99999785 | -6.3396557 | down |
| KI - WT, cell_origin: iPSC | P18583 | SON | 12 | Homo sapien | 0.12863226 | -0.7802741 | 1.03753866 | -1.0506185 | 0.30147315 | 0.76716496 | 0.99999785 | -6.3396822 | up |
| KI - WT, cell_origin: iPSC | Q70E73 | RAPH1 | 5 | Homo sapien | 0.10164226 | -0.6165989 | 0.81988343 | -0.2484096 | 0.3014003 | 0.76721096 | 0.99999785 | -6.3397106 | up |
| KI - WT, cell_origin: iPSC | P04632 | CAPNS1 | 9 | Homo sapien | 0.21991284 | -1.3352496 | 1.77507526 | 2.08994348 | 0.30135416 | 0.76727392 | 0.99999785 | -6.3397055 | up |
| KI - WT, cell_origin: iPSC | Q9Y230 | RUVBL2 | 23 | Homo sapien | -0.087994 | -0.7100807 | 0.53409255 | 1.85542182 | -0.3012605 | 0.76731554 | 0.99999785 | -6.339754 | down |
| KI - WT, cell_origin: iPSC | P52209 | PGD | 9 | Homo sapien | -0.1398096 | -1.1295893 | 0.84997006 | -0.4220209 | -0.3010238 | 0.76752093 | 0.99999785 | -6.339808 | down |
| KI - WT, cell_origin: iPSC | O75064 | DENND4B | 2 | Homo sapien | 0.41315204 | -2.5136399 | 3.33994395 | 1.09902621 | 0.3008295 | 0.76766626 | 0.99999785 | -6.3398683 | up |
| KI - WT, cell_origin: iPSC | Q9P2F8 | SIPA1L2 | 27 | Homo sapien | -0.3574355 | -2.8928452 | 2.17797421 | -0.9883473 | -0.3004359 | 0.76796062 | 0.99999785 | -6.3399902 | down |
| KI - WT, cell_origin: iPSC | P62241 | RPS8 | 10 | Homo sapien | 0.06289467 | -0.3841218 | 0.50991116 | 5.94447117 | 0.29966085 | 0.76851214 | 0.99999785 | -6.3402491 | up |
| KI - WT, cell_origin: iPSC | Q9NVA2 | SEPTIN11 | 8 | Homo sapien | -0.1186193 | -0.9619351 | 0.72469648 | -0.9990739 | -0.2996131 | 0.76855387 | 0.99999785 | -6.3402597 | down |
| KI - WT, cell_origin: iPSC | P61962 | DCAF7 | 11 | Homo sapien | -0.0717157 | -0.5815621 | 0.43813066 | 0.69378485 | -0.2995814 | 0.76857161 | 0.99999785 | -6.3402736 | down |
| KI - WT, cell_origin: iPSC | Q63ZY3 | KANK2 | 5 | Homo sapien | 0.19167245 | -1.1716404 | 1.55498534 | -2.6792988 | 0.29948982 | 0.76864854 | 0.99999785 | -6.3402962 | up |
| KI - WT, cell_origin: iPSC | Q16531 | DDB1 | 35 | Homo sapien | 0.09521647 | -0.582069 | 0.77250194 | 0.73278712 | 0.29941945 | 0.76869278 | 0.99999785 | -6.3403236 | up |
| KI - WT, cell_origin: iPSC | Q9Y678 | COPG1 | 29 | Homo sapien | -0.1392103 | -1.1301509 | 0.85173019 | 1.09611325 | -0.2993824 | 0.76874876 | 0.99999785 | -6.3403159 | down |
| KI - WT, cell_origin: iPSC | Q14839 | CHD4 | 17 | Homo sapien | 0.13701806 | -0.8378689 | 1.11190503 | -1.6767157 | 0.29937754 | 0.76873016 | 0.99999785 | -6.3403324 | up |
| KI - WT, cell_origin: iPSC | P08727 | KRT19 | 7 | Homo sapien | 0.17275857 | -1.0571601 | 1.40267722 | -0.281327 | 0.29934057 | 0.76878005 | 0.99999785 | -6.3403288 | up |
| KI - WT, cell_origin: iPSC | Q8TF39 | ZNF483 | 6 | Homo sapien | 0.13136856 | -0.8044264 | 1.06716354 | -2.128091 | 0.29898568 | 0.76901739 | 0.99999785 | -6.3404573 | up |
| KI - WT, cell_origin: iPSC | Q9UQE7 | SMC3 | 42 | Homo sapien | -0.2060511 | -1.6749608 | 1.26285868 | -0.3609931 | -0.2989387 | 0.76908075 | 0.99999785 | -6.3404527 | down |
| KI - WT, cell_origin: iPSC | Q14247 | CTTN | 26 | Homo sapien | -0.1280578 | -1.0436563 | 0.78754066 | 1.03879567 | -0.2980598 | 0.76973856 | 0.99999785 | -6.340723 | down |
| KI - WT, cell_origin: iPSC | Q13823 | GNL2 | 15 | Homo sapien | -0.099531 | -0.8113163 | 0.61225435 | -0.9041942 | -0.2978167 | 0.76989245 | 0.99999785 | -6.3408166 | down |
| KI - WT, cell_origin: iPSC | Q9P2K5 | MYEF2 | 8 | Homo sapien | 0.13388961 | -0.8239604 | 1.09173963 | -2.195919 | 0.297707 | 0.76997456 | 0.99999785 | -6.3408503 | up |
| KI - WT, cell_origin: iPSC | P05161 | ISG15 | 4 | Homo sapien | -0.2513614 | -2.0514856 | 1.54876268 | -2.2917202 | -0.2975765 | 0.77010036 | 0.99999785 | -6.3408714 | down |
| KI - WT, cell_origin: iPSC | O95487 | SEC24B | 21 | Homo sapien | 0.10182239 | -0.6278717 | 0.83151654 | 1.18892437 | 0.29721678 | 0.77034495 | 0.99999785 | -6.3409983 | up |
| KI - WT, cell_origin: iPSC | P23258 | TUBG1 | 18 | Homo sapien | 0.06752454 | -0.4165079 | 0.55155695 | 2.24823039 | 0.29711658 | 0.77041666 | 0.99999785 | -6.3410312 | up |
| KI - WT, cell_origin: iPSC | Q8NDN9 | RCBTB1 | 5 | Homo sapien | -0.0640922 | -0.5239009 | 0.3957165 | -0.6613366 | -0.2968709 | 0.77060067 | 0.99999785 | -6.3411064 | down |
| KI - WT, cell_origin: iPSC | O75390 | CS | 2 | Homo sapien | 0.35752354 | -2.209274 | 2.92432112 | -3.154182 | 0.29683519 | 0.77065537 | 0.99999785 | -6.3410985 | up |
| KI - WT, cell_origin: iPSC | P00558 | PGK1 | 10 | Homo sapien | -0.1999976 | -1.6373714 | 1.23737619 | 0.50251145 | -0.2965224 | 0.77088963 | 0.99999785 | -6.3411941 | down |
| KI - WT, cell_origin: iPSC | Q92979 | EMG1 | 3 | Homo sapien | 0.20580305 | -1.2747654 | 1.68637153 | -1.00022 | 0.29622771 | 0.77111031 | 0.99999785 | -6.3412841 | up |
| KI - WT, cell_origin: iPSC | P02649 | APOE | 17 | Homo sapien | 0.11728123 | -0.7267208 | 0.96128328 | 1.5053461 | 0.29604759 | 0.77123185 | 0.99999785 | -6.3413481 | up |
| KI - WT, cell_origin: iPSC | Q12769 | NUP160 | 8 | Homo sapien | 0.13765085 | -0.8570649 | 1.13236661 | -0.8873331 | 0.29488344 | 0.77211395 | 0.99999785 | -6.3416959 | up |
| KI - WT, cell_origin: iPSC | P49756 | RBM25 | 11 | Homo sapien | -0.1256054 | -1.0331882 | 0.7819775 | -1.0750064 | -0.2948083 | 0.77215408 | 0.99999785 | -6.3417295 | down |
| KI - WT, cell_origin: iPSC | Q8WXE1 | ATRIP | 9 | Homo sapien | 0.20155062 | -1.2567863 | 1.65988754 | -1.0295552 | 0.2945294 | 0.77238261 | 0.99999785 | -6.3418012 | up |
| KI - WT, cell_origin: iPSC | Q00587 | CDC42EP1 | 17 | Homo sapien | -0.1795468 | -1.4803985 | 1.12130483 | 1.26824594 | -0.2941388 | 0.7726753 | 0.99999785 | -6.3419197 | down |

| | | | | | | | | | | | | |
|---|---|---|---|---|---|---|---|---|---|---|---|---|
| KI - WT, cell_origin: iPSC | Q13426 | XRCC4 | 3 | Homo sapien | 0.35781758 | -2.2347567 | 2.95039182 | -2.4809493 | 0.29412561 | 0.77268521 | 0.99999785 | -6.3419237 up |
| KI - WT, cell_origin: iPSC | Q6P3X3 | TTC27 | 3 | Homo sapien | 0.10105349 | -0.6321932 | 0.83430016 | -1.2789572 | 0.29352216 | 0.77310985 | 0.99999785 | -6.3421249 up |
| KI - WT, cell_origin: iPSC | Q9P266 | JCAD | 7 | Homo sapien | 0.33099406 | -2.0724275 | 2.73441565 | -2.2799505 | 0.29348953 | 0.77316197 | 0.99999785 | -6.3421164 up |
| KI - WT, cell_origin: iPSC | Q9NUL5 | SHFL | 2 | Homo sapien | -0.1934421 | -1.598427 | 1.21154291 | -0.5587337 | -0.2934145 | 0.77321819 | 0.99999785 | -6.342139 down |
| KI - WT, cell_origin: iPSC | P30419 | NMT1 | 7 | Homo sapien | -0.2083424 | -1.7221718 | 1.30548701 | -0.8567104 | -0.2932939 | 0.77330858 | 0.99999785 | -6.3421755 down |
| KI - WT, cell_origin: iPSC | Q9H9A6 | LRRC40 | 26 | Homo sapien | 0.10408178 | -0.6526618 | 0.86082537 | 0.72917773 | 0.29297594 | 0.77352635 | 0.99999785 | -6.3422853 up |
| KI - WT, cell_origin: iPSC | P52294 | KPNA1 | 10 | Homo sapien | 0.09379074 | -0.5884257 | 0.77600717 | -0.259139 | 0.29280431 | 0.77364808 | 0.99999785 | -6.3423418 up |
| KI - WT, cell_origin: iPSC | Q14164 | IKBKE | 4 | Homo sapien | 0.20427255 | -1.2836597 | 1.69220484 | -1.8285964 | 0.2925696 | 0.77385165 | 0.99999785 | -6.3423942 up |
| KI - WT, cell_origin: iPSC | Q13576 | IQGAP2 | 2 | Homo sapien | -0.2199767 | -1.8270498 | 1.38709653 | -2.5081838 | -0.2917046 | 0.77450031 | 0.99999785 | -6.3426547 down |
| KI - WT, cell_origin: iPSC | P68104 | EEF1A1 | 22 | Homo sapien | -0.0834842 | -0.6933275 | 0.52635924 | 5.28788483 | -0.2916108 | 0.77455138 | 0.99999785 | -6.3426957 down |
| KI - WT, cell_origin: iPSC | Q9UJF2 | RASAL2 | 5 | Homo sapien | 0.15818024 | -0.9990021 | 1.31536255 | -0.9815328 | 0.29130828 | 0.77479759 | 0.99999785 | -6.3427738 up |
| KI - WT, cell_origin: iPSC | Q06203 | PPAT | 8 | Homo sapien | 0.16131398 | -1.0204166 | 1.34304451 | -1.280248 | 0.29090816 | 0.77509774 | 0.99999785 | -6.3428939 up |
| KI - WT, cell_origin: iPSC | O14979 | HNRNPDL | 9 | Homo sapien | -0.1330022 | -1.1074532 | 0.84144879 | 2.82952588 | -0.2908715 | 0.77512525 | 0.99999785 | -6.3429049 down |
| KI - WT, cell_origin: iPSC | Q32P44 | EML3 | 7 | Homo sapien | 0.27003473 | -1.7088838 | 2.24895322 | -2.6923264 | 0.29079978 | 0.77517904 | 0.99999785 | -6.3429264 up |
| KI - WT, cell_origin: iPSC | Q9H1A4 | ANAPC1 | 10 | Homo sapien | -0.1384877 | -1.1536116 | 0.87663615 | -0.9915759 | -0.2907118 | 0.77524172 | 0.99999785 | -6.342955 down |
| KI - WT, cell_origin: iPSC | P25685 | DNAJB1 | 4 | Homo sapien | -0.0953325 | -0.7944044 | 0.60373949 | -0.8162283 | -0.2904415 | 0.77542051 | 0.99999785 | -6.3430518 down |
| KI - WT, cell_origin: iPSC | P24557 | TBXAS1 | 2 | Homo sapien | -1.3001126 | -10.840518 | 8.24029234 | -3.8561583 | -0.2904133 | 0.775469 | 0.99999785 | -6.3430422 down |
| KI - WT, cell_origin: iPSC | Q16204 | CCDC6 | 7 | Homo sapien | 0.14276812 | -0.9050161 | 1.19055232 | -1.0275079 | 0.29037669 | 0.77549648 | 0.99999785 | -6.3430532 up |
| KI - WT, cell_origin: iPSC | Q14807 | KIF22 | 13 | Homo sapien | -0.1566705 | -1.3076818 | 0.99434079 | -1.1309807 | -0.2900748 | 0.77572298 | 0.99999785 | -6.3431435 down |
| KI - WT, cell_origin: iPSC | O43353 | RIPK2 | 5 | Homo sapien | 0.11578524 | -0.7349862 | 0.96655665 | -1.2834881 | 0.28985844 | 0.77585869 | 0.99999785 | -6.3432258 up |
| KI - WT, cell_origin: iPSC | P46940 | IQGAP1 | 42 | Homo sapien | -0.1251247 | -1.0480615 | 0.79781217 | 0.70128747 | -0.2889172 | 0.77659183 | 0.99999785 | -6.343489 down |
| KI - WT, cell_origin: iPSC | Q03923 | ZNF85 | 2 | Homo sapien | 0.09931668 | -0.633081 | 0.83171434 | -1.2012779 | 0.28881179 | 0.77664376 | 0.99999785 | -6.3435383 up |
| KI - WT, cell_origin: iPSC | P60981 | DSTN | 8 | Homo sapien | -0.1152907 | -0.9657806 | 0.7351991 | 1.56472033 | -0.2888061 | 0.77666263 | 0.99999785 | -6.3435303 down |
| KI - WT, cell_origin: iPSC | Q86U70 | LDB1 | 5 | Homo sapien | -0.143547 | -1.2041971 | 0.91710308 | 0.01564064 | -0.2884193 | 0.77696559 | 0.99999785 | -6.3436372 down |
| KI - WT, cell_origin: iPSC | Q14847 | LASP1 | 11 | Homo sapien | 0.09746792 | -0.622896 | 0.81783182 | 3.58078982 | 0.28822237 | 0.7770944 | 0.99999785 | -6.3437082 up |
| KI - WT, cell_origin: iPSC | O75688 | PPM1B | 3 | Homo sapien | 0.18757104 | -1.1991745 | 1.57431658 | -4.5784663 | 0.28807706 | 0.77719544 | 0.99999785 | -6.3437567 up |
| KI - WT, cell_origin: iPSC | Q9NXH9 | TRMT1 | 3 | Homo sapien | 0.13130844 | -0.8416756 | 1.10429246 | -1.9393927 | 0.28742636 | 0.77768413 | 0.99999785 | -6.3439497 up |
| KI - WT, cell_origin: iPSC | O00203 | AP3B1 | 24 | Homo sapien | 0.10736723 | -0.6901864 | 0.90492087 | 0.30511186 | 0.28676453 | 0.77818887 | 0.99999785 | -6.3441405 up |
| KI - WT, cell_origin: iPSC | P55010 | EIF5 | 3 | Homo sapien | -0.1453843 | -1.2276034 | 0.93683474 | -1.763658 | -0.2861675 | 0.77863785 | 0.99999785 | -6.3443165 down |
| KI - WT, cell_origin: iPSC | Q5VUJ6 | LRCH2 | 3 | Homo sapien | 0.10757626 | -0.6930996 | 0.90825208 | -1.1683407 | 0.28615374 | 0.77864019 | 0.99999785 | -6.3443258 up |
| KI - WT, cell_origin: iPSC | Q2TBE0 | CWF19L2 | 21 | Homo sapien | -0.2385821 | -2.015794 | 1.53862977 | -0.4430746 | -0.2860889 | 0.77871581 | 0.99999785 | -6.3443274 down |
| KI - WT, cell_origin: iPSC | P13667 | PDIA4 | 21 | Homo sapien | -0.2899596 | -2.4514358 | 1.87151663 | -0.539417 | -0.2858837 | 0.77886998 | 0.99999785 | -6.3443879 down |
| KI - WT, cell_origin: iPSC | Q96MG7 | NSMCE3 | 5 | Homo sapien | -0.127168 | -1.0750761 | 0.82074019 | -1.9832204 | -0.2857269 | 0.77896092 | 0.99999785 | -6.3444516 down |
| KI - WT, cell_origin: iPSC | P07900 | HSP90AA1 | 14 | Homo sapien | -0.1155765 | -0.9778519 | 0.74669897 | 1.14196676 | -0.2855819 | 0.77908703 | 0.99999785 | -6.3444832 down |
| KI - WT, cell_origin: iPSC | O94906 | PRPF6 | 36 | Homo sapien | -0.1339551 | -1.1343424 | 0.86643214 | 0.88796531 | -0.2853603 | 0.77926326 | 0.99999785 | -6.3445421 down |
| KI - WT, cell_origin: iPSC | Q6PKG0 | LARP1 | 37 | Homo sapien | -0.0601299 | -0.5089552 | 0.38869536 | 2.75526045 | -0.2853335 | 0.77925655 | 0.99999785 | -6.3445674 down |
| KI - WT, cell_origin: iPSC | Q15477 | SKIV2L | 17 | Homo sapien | -0.1212245 | -1.0266553 | 0.7842064 | -0.7226353 | -0.2852407 | 0.77934032 | 0.99999785 | -6.3445856 down |
| KI - WT, cell_origin: iPSC | Q0VDF9 | HSPA14 | 13 | Homo sapien | -0.109807 | -0.9308801 | 0.71126622 | 0.64221161 | -0.2848826 | 0.77960347 | 0.99999785 | -6.3446947 down |
| KI - WT, cell_origin: iPSC | P05198 | EIF2S1 | 13 | Homo sapien | 0.09632526 | -0.6240914 | 0.81674193 | 2.44050298 | 0.28479169 | 0.77966701 | 0.99999785 | -6.3447246 up |
| KI - WT, cell_origin: iPSC | P52630 | STAT2 | 6 | Homo sapien | 0.1365563 | -0.8879462 | 1.1610588 | -0.7756425 | 0.28405411 | 0.78024504 | 0.99999785 | -6.3449255 up |
| KI - WT, cell_origin: iPSC | Q9Y5P4 | CERT1 | 5 | Homo sapien | -0.396798 | -3.3753194 | 2.58172343 | -3.083932 | -0.2839037 | 0.7803581 | 0.99999785 | -6.3449696 down |
| KI - WT, cell_origin: iPSC | O75533 | SF3B1 | 40 | Homo sapien | -0.1517934 | -1.2920055 | 0.98841877 | 4.08810654 | -0.2837067 | 0.78050625 | 0.99999785 | -6.3450273 down |
| KI - WT, cell_origin: iPSC | P55735 | SEC13 | 12 | Homo sapien | 0.08462185 | -0.5511369 | 0.7203806 | 2.8288311 | 0.28348484 | 0.78064639 | 0.99999785 | -6.3451094 up |
| KI - WT, cell_origin: iPSC | P50749 | RASSF2 | 3 | Homo sapien | -0.1183879 | -1.0089555 | 0.77217965 | -1.1781623 | -0.2831701 | 0.7808899 | 0.99999785 | -6.3451969 down |
| KI - WT, cell_origin: iPSC | Q9UKV8 | AGO2 | 21 | Homo sapien | -0.1467371 | -1.2513802 | 0.95790611 | 1.37477051 | -0.2830871 | 0.78097208 | 0.99999785 | -6.3452083 down |
| KI - WT, cell_origin: iPSC | Q96P16 | RPRD1A | 5 | Homo sapien | 0.10551455 | -0.6890439 | 0.90007298 | -0.3168257 | 0.28286948 | 0.78111524 | 0.99999785 | -6.3452851 up |

| | | | | | | | | | | | | | |
|---|---|---|---|---|---|---|---|---|---|---|---|---|---|
| KI - WT, cell_origin: iPSC | P02792 | FTL | 2 | Homo sapien | 0.15490903 | -1.0122896 | 1.32210763 | -1.3080536 | 0.28283578 | 0.78116111 | 0.99999785 | -6.3452817 | up |
| KI - WT, cell_origin: iPSC | Q9ULC4 | MCTS1 | 4 | Homo sapien | -0.1071572 | -0.9150572 | 0.70074275 | -0.1845424 | -0.2825339 | 0.78136835 | 0.99999785 | -6.3453824 | down |
| KI - WT, cell_origin: iPSC | P04792 | HSPB1 | 8 | Homo sapien | -0.135287 | -1.1567848 | 0.88621092 | 1.65153398 | -0.2822415 | 0.78160809 | 0.99999785 | -6.3454548 | down |
| KI - WT, cell_origin: iPSC | P29401 | TKT | 14 | Homo sapien | -0.1531957 | -1.3103808 | 1.00398939 | 0.06234089 | -0.2821279 | 0.78169349 | 0.99999785 | -6.3454879 | down |
| KI - WT, cell_origin: iPSC | Q8IWE2 | FAM114A1 | 4 | Homo sapien | -0.3609688 | -3.0879418 | 2.36600414 | -1.7999047 | -0.2820923 | 0.7817203 | 0.99999785 | -6.3454982 | down |
| KI - WT, cell_origin: iPSC | Q86Y56 | DNAAF5 | 28 | Homo sapien | -0.1453248 | -1.2432383 | 0.95258872 | 0.50066757 | -0.2820811 | 0.78172872 | 0.99999785 | -6.3455015 | down |
| KI - WT, cell_origin: iPSC | P62917 | RPL8 | 10 | Homo sapien | -0.0742352 | -0.6352767 | 0.48680626 | 4.08563017 | -0.2818089 | 0.781907 | 0.99999785 | -6.3455977 | down |
| KI - WT, cell_origin: iPSC | Q99836 | MYD88 | 11 | Homo sapien | -0.1011108 | -0.8658502 | 0.66362864 | 1.73618346 | -0.2816376 | 0.78204262 | 0.99999785 | -6.3456431 | down |
| KI - WT, cell_origin: iPSC | O94855 | SEC24D | 8 | Homo sapien | 0.08032107 | -0.5276852 | 0.68832737 | 0.02831838 | 0.28135919 | 0.78224538 | 0.99999785 | -6.3457282 | up |
| KI - WT, cell_origin: iPSC | Q8NDV7 | TNRC6A | 19 | Homo sapien | -0.1537462 | -1.3185671 | 1.01107472 | -0.1398709 | -0.2812856 | 0.78232717 | 0.99999785 | -6.3457326 | down |
| KI - WT, cell_origin: iPSC | P56537 | EIF6 | 5 | Homo sapien | -0.0810184 | -0.6949828 | 0.53294611 | 1.36570324 | -0.2810476 | 0.78247986 | 0.99999785 | -6.3458186 | down |
| KI - WT, cell_origin: iPSC | P00387 | CYB5R3 | 6 | Homo sapien | 0.15266826 | -1.0051809 | 1.3105174 | -0.9877061 | 0.28099536 | 0.78254557 | 0.99999785 | -6.3458167 | up |
| KI - WT, cell_origin: iPSC | Q13347 | EIF3I | 19 | Homo sapien | -0.0678716 | -0.5826972 | 0.44695412 | 2.66493233 | -0.2807808 | 0.78268067 | 0.99999785 | -6.3458958 | down |
| KI - WT, cell_origin: iPSC | Q9Y4D8 | HECTD4 | 6 | Homo sapien | 0.09586399 | -0.6313226 | 0.82305054 | -1.3867197 | 0.28076912 | 0.78268943 | 0.99999785 | -6.3458992 | up |
| KI - WT, cell_origin: iPSC | P53367 | ARFIP1 | 10 | Homo sapien | 0.10804534 | -0.7120016 | 0.92809226 | 0.58574964 | 0.28066305 | 0.78277712 | 0.99999785 | -6.3459249 | up |
| KI - WT, cell_origin: iPSC | O00479 | HMGN4 | 3 | Homo sapien | -0.2685008 | -2.3107656 | 1.77376397 | 2.38748725 | -0.2801792 | 0.78315975 | 0.99999785 | -6.3460529 | down |
| KI - WT, cell_origin: iPSC | P0C0S5;Q71U | H2AZ1;H2AZ2 | 2 | Homo sapien | -0.1377396 | -1.1880344 | 0.91255519 | 0.364167 | -0.2794795 | 0.78368641 | 0.99999785 | -6.3462548 | down |
| KI - WT, cell_origin: iPSC | P09038 | FGF2 | 4 | Homo sapien | -0.2380199 | -2.0532866 | 1.57724679 | -1.1491531 | -0.2794314 | 0.78372267 | 0.99999785 | -6.3462687 | down |
| KI - WT, cell_origin: iPSC | P47755 | CAPZA2 | 10 | Homo sapien | 0.0937428 | -0.620844 | 0.80832962 | 2.25313895 | 0.27941319 | 0.78371255 | 0.99999785 | -6.3462892 | up |
| KI - WT, cell_origin: iPSC | Q9UL18 | AGO1 | 27 | Homo sapien | -0.1216102 | -1.0496058 | 0.80638546 | 1.80738614 | -0.2792713 | 0.78384318 | 0.99999785 | -6.3463148 | down |
| KI - WT, cell_origin: iPSC | Q9GZV8 | PRDM14 | 8 | Homo sapien | 0.09644556 | -0.639145 | 0.83203615 | -0.2928679 | 0.27924524 | 0.78383658 | 0.99999785 | -6.3463391 | up |
| KI - WT, cell_origin: iPSC | Q99729 | HNRNPAB | 8 | Homo sapien | -0.1111912 | -0.959553 | 0.73717049 | 2.35911251 | -0.2792365 | 0.78385743 | 0.99999785 | -6.3463325 | down |
| KI - WT, cell_origin: iPSC | Q96A65 | EXOC4 | 27 | Homo sapien | 0.19010337 | -1.2625369 | 1.64274366 | -0.9010722 | 0.27889074 | 0.78412971 | 0.99999785 | -6.3464244 | up |
| KI - WT, cell_origin: iPSC | P07108 | DBI | 2 | Homo sapien | -0.206022 | -1.7815401 | 1.36949599 | -1.153273 | -0.2786716 | 0.78429473 | 0.99999785 | -6.3464874 | down |
| KI - WT, cell_origin: iPSC | A5YKK6 | CNOT1 | 62 | Homo sapien | 0.07670014 | -0.5097729 | 0.66317313 | 1.13404918 | 0.27854015 | 0.78436753 | 0.99999785 | -6.3465419 | up |
| KI - WT, cell_origin: iPSC | P51532 | SMARCA4 | 20 | Homo sapien | -0.1670293 | -1.446282 | 1.11222342 | -1.0505096 | -0.2782522 | 0.78461051 | 0.99999785 | -6.3466079 | down |
| KI - WT, cell_origin: iPSC | Q9UBU8 | MORF4L1 | 8 | Homo sapien | -0.1547166 | -1.3397007 | 1.03026749 | -1.1791236 | -0.2782446 | 0.78461621 | 0.99999785 | -6.34661 | down |
| KI - WT, cell_origin: iPSC | Q9NQA3 | WASH6P | 4 | Homo sapien | -0.1560156 | -1.3509535 | 1.03892242 | -1.6987981 | -0.278223 | 0.78462933 | 0.99999785 | -6.3466183 | down |
| KI - WT, cell_origin: iPSC | Q9UBD5 | ORC3 | 10 | Homo sapien | -0.1847132 | -1.5997014 | 1.23027503 | -1.211314 | -0.2781938 | 0.78465452 | 0.99999785 | -6.3466246 | down |
| KI - WT, cell_origin: iPSC | Q9BS26 | ERP44 | 15 | Homo sapien | -0.0931982 | -0.8070345 | 0.62063799 | 2.70442124 | -0.2780921 | 0.78470894 | 0.99999785 | -6.3466679 | down |
| KI - WT, cell_origin: iPSC | O95163 | ELP1 | 24 | Homo sapien | 0.08992332 | -0.5992227 | 0.77906932 | -0.2853941 | 0.27790784 | 0.78484377 | 0.99999785 | -6.3467232 | up |
| KI - WT, cell_origin: iPSC | Q8N9N2 | ASCC1 | 6 | Homo sapien | -0.0934349 | -0.8095537 | 0.6226839 | 0.46135934 | -0.277888 | 0.78485932 | 0.99999785 | -6.3467286 | down |
| KI - WT, cell_origin: iPSC | Q96Q15 | SMG1 | 13 | Homo sapien | 0.11110341 | -0.7421553 | 0.96436212 | -1.6124295 | 0.2773233 | 0.78528412 | 0.99999785 | -6.3468906 | up |
| KI - WT, cell_origin: iPSC | O95049 | TJP3 | 5 | Homo sapien | -0.3743547 | -3.2513507 | 2.50264126 | -4.1325816 | -0.2772978 | 0.78532935 | 0.99999785 | -6.3468813 | down |
| KI - WT, cell_origin: iPSC | Q14240 | EIF4A2 | 6 | Homo sapien | -0.1202844 | -1.0448548 | 0.80428598 | -0.4260627 | -0.2772052 | 0.78539214 | 0.99999785 | -6.3469122 | down |
| KI - WT, cell_origin: iPSC | Q6DKJ4 | NXN | 3 | Homo sapien | -0.1737712 | -1.5109468 | 1.16340447 | -1.5367528 | -0.2769438 | 0.78559603 | 0.99999785 | -6.3469825 | down |
| KI - WT, cell_origin: iPSC | Q13627 | DYRK1A | 9 | Homo sapien | 0.07651152 | -0.512973 | 0.66599607 | -0.2483136 | 0.27643569 | 0.78595291 | 0.99999785 | -6.347144 | up |
| KI - WT, cell_origin: iPSC | O14974 | PPP1R12A | 21 | Homo sapien | -0.0929321 | -0.8099723 | 0.62410814 | 1.52449326 | -0.2760417 | 0.78625108 | 0.99999785 | -6.3472554 | down |
| KI - WT, cell_origin: iPSC | P12081 | HARS1 | 5 | Homo sapien | -0.2513238 | -2.1914563 | 1.68880875 | -3.6053502 | -0.2760204 | 0.78628552 | 0.99999785 | -6.3472498 | down |
| KI - WT, cell_origin: iPSC | Q14241 | ELOA | 8 | Homo sapien | -0.2528097 | -2.2081553 | 1.70253582 | -0.8732792 | -0.2755324 | 0.78665956 | 0.99999785 | -6.3473847 | down |
| KI - WT, cell_origin: iPSC | Q14137 | BOP1 | 4 | Homo sapien | -0.1266741 | -1.1062383 | 0.85289009 | -1.6811906 | -0.275455 | 0.78669757 | 0.99999785 | -6.3474195 | down |
| KI - WT, cell_origin: iPSC | Q9Y5S2 | CDC42BPB | 81 | Homo sapien | 0.20679233 | -1.3934017 | 1.80698635 | 2.40448337 | 0.27540013 | 0.78675923 | 0.99999785 | -6.3474223 | up |
| KI - WT, cell_origin: iPSC | P30626 | SRI | 2 | Homo sapien | -0.3432706 | -3.001214 | 2.31467273 | -2.0526959 | -0.2752284 | 0.78688865 | 0.99999785 | -6.347471 | down |
| KI - WT, cell_origin: iPSC | Q15645 | TRIP13 | 8 | Homo sapien | 0.17744173 | -1.1971315 | 1.55201498 | -0.860682 | 0.27509978 | 0.78698562 | 0.99999785 | -6.3475075 | up |
| KI - WT, cell_origin: iPSC | Q9NR09 | BIRC6 | 9 | Homo sapien | 0.10586309 | -0.7138937 | 0.92561992 | -0.7506135 | 0.27508022 | 0.78698047 | 0.99999785 | -6.3475256 | up |
| KI - WT, cell_origin: iPSC | O60832 | DKC1 | 14 | Homo sapien | -0.1507207 | -1.3187995 | 1.01735815 | 0.56530887 | -0.2749812 | 0.78707497 | 0.99999785 | -6.3475412 | down |

| | | | | | | | | | | | | | |
|---|---|---|---|---|---|---|---|---|---|---|---|---|---|
| KI - WT, cell_origin: iPSC | Q9UGR2 | ZC3H7B | 9 | Homo sapien | -0.0675062 | -0.5907869 | 0.45577443 | 0.25286882 | -0.2747571 | 0.78721814 | 0.99999785 | -6.347621 | down |
| KI - WT, cell_origin: iPSC | Q9NZL4 | HSPBP1 | 7 | Homo sapien | 0.13500115 | -0.9116431 | 1.18164537 | -2.4330368 | 0.27471229 | 0.78725194 | 0.99999785 | -6.3476337 | up |
| KI - WT, cell_origin: iPSC | P09234 | SNRPC | 3 | Homo sapien | -0.2065487 | -1.8090253 | 1.39592794 | 1.84577564 | -0.2746838 | 0.78729919 | 0.99999785 | -6.3476255 | down |
| KI - WT, cell_origin: iPSC | P51153 | RAB13 | 10 | Homo sapien | 0.13631302 | -0.9228253 | 1.19545132 | 4.30192768 | 0.27427551 | 0.78760701 | 0.99999785 | -6.3477411 | up |
| KI - WT, cell_origin: iPSC | P55265 | ADAR | 33 | Homo sapien | -0.0827865 | -0.7264386 | 0.56086557 | 0.27043028 | -0.2739353 | 0.78783782 | 0.99999785 | -6.3478535 | down |
| KI - WT, cell_origin: iPSC | Q9Y2W1 | THRAP3 | 40 | Homo sapien | -0.1419722 | -1.2470955 | 0.96315115 | 3.1089941 | -0.2737757 | 0.7879839 | 0.99999785 | -6.3478824 | down |
| KI - WT, cell_origin: iPSC | Q9H3F6 | KCTD10 | 2 | Homo sapien | -0.2001255 | -1.7571058 | 1.35685487 | -3.7593526 | -0.273753 | 0.78797529 | 0.99999785 | -6.3479049 | down |
| KI - WT, cell_origin: iPSC | P19474 | TRIM21 | 6 | Homo sapien | 0.10247381 | -0.6951112 | 0.90005886 | -0.4099394 | 0.27367509 | 0.7880399 | 0.99999785 | -6.3479233 | up |
| KI - WT, cell_origin: iPSC | P84103 | SRSF3 | 10 | Homo sapien | -0.1654926 | -1.4548603 | 1.12387512 | 3.64849044 | -0.2735294 | 0.78816956 | 0.99999785 | -6.3479519 | down |
| KI - WT, cell_origin: iPSC | Q4VC05 | BCL7A | 2 | Homo sapien | 0.2149475 | -1.4604313 | 1.89032626 | -1.7505838 | 0.27341464 | 0.78825614 | 0.99999785 | -6.3479843 | up |
| KI - WT, cell_origin: iPSC | P28066 | PSMA5 | 8 | Homo sapien | 0.12545742 | -0.8546572 | 1.105572 | -0.8921001 | 0.27274964 | 0.78875207 | 0.99999785 | -6.3481752 | up |
| KI - WT, cell_origin: iPSC | Q96KQ4 | PPP1R13B | 2 | Homo sapien | -0.1401364 | -1.2346541 | 0.95438116 | -2.7187466 | -0.2726893 | 0.78877766 | 0.99999785 | -6.3482047 | down |
| KI - WT, cell_origin: iPSC | Q86VW0 | SESTD1 | 5 | Homo sapien | -0.1165132 | -1.0271867 | 0.7941604 | -0.8492174 | -0.2725589 | 0.78888653 | 0.99999785 | -6.3482347 | down |
| KI - WT, cell_origin: iPSC | Q8IZ83 | ALDH16A1 | 3 | Homo sapien | 0.14291854 | -0.9748598 | 1.2606969 | -1.3924061 | 0.2724618 | 0.78897193 | 0.99999785 | -6.3482544 | up |
| KI - WT, cell_origin: iPSC | Q9UKX7 | NUP50 | 4 | Homo sapien | 0.20524666 | -1.4004968 | 1.8109901 | -1.2602317 | 0.27239699 | 0.7890237 | 0.99999785 | -6.3482708 | up |
| KI - WT, cell_origin: iPSC | P17812 | CTPS1 | 26 | Homo sapien | -0.1300123 | -1.1475516 | 0.8875271 | 1.61216631 | -0.2722923 | 0.78910264 | 0.99999785 | -6.3483002 | down |
| KI - WT, cell_origin: iPSC | Q04323 | UBXN1 | 6 | Homo sapien | 0.12286155 | -0.841156 | 1.08687907 | 0.35715275 | 0.2716023 | 0.78962325 | 0.99999785 | -6.3484938 | up |
| KI - WT, cell_origin: iPSC | P29372 | MPG | 6 | Homo sapien | 0.1387164 | -0.9504251 | 1.22785793 | -0.5148318 | 0.2714225 | 0.78975892 | 0.99999785 | -6.3485442 | up |
| KI - WT, cell_origin: iPSC | P14174 | MIF | 4 | Homo sapien | -0.0927785 | -0.8234391 | 0.63788205 | 6.95251959 | -0.270604 | 0.7903766 | 0.99999785 | -6.348773 | down |
| KI - WT, cell_origin: iPSC | P10809 | HSPD1 | 16 | Homo sapien | 0.16076898 | -1.1058102 | 1.42734812 | 0.44633392 | 0.27050313 | 0.79045274 | 0.99999785 | -6.3488012 | up |
| KI - WT, cell_origin: iPSC | Q8N5P1 | ZC3H8 | 3 | Homo sapien | 0.10873383 | -0.7487553 | 0.96622299 | -1.5936091 | 0.2700696 | 0.79075467 | 0.99999785 | -6.3489378 | up |
| KI - WT, cell_origin: iPSC | Q99959 | PKP2 | 11 | Homo sapien | -0.2376824 | -2.1138024 | 1.63843757 | -0.8230986 | -0.2699845 | 0.79084423 | 0.99999785 | -6.3489458 | down |
| KI - WT, cell_origin: iPSC | O95881 | TXNDC12 | 8 | Homo sapien | -0.3314288 | -2.9488273 | 2.28596961 | 0.93893102 | -0.2698502 | 0.79094557 | 0.99999785 | -6.3489832 | down |
| KI - WT, cell_origin: iPSC | Q14149 | MORC3 | 2 | Homo sapien | 0.1948028 | -1.3489105 | 1.7385161 | -1.3676409 | 0.26892483 | 0.79164427 | 0.99999785 | -6.3492404 | up |
| KI - WT, cell_origin: iPSC | Q8WX93 | PALLD | 28 | Homo sapien | 0.18396806 | -1.278974 | 1.64691017 | 0.83732523 | 0.26798943 | 0.7923507 | 0.99999785 | -6.3494995 | up |
| KI - WT, cell_origin: iPSC | A0MZ66 | SHTN1 | 2 | Homo sapien | -0.2554931 | -2.2881404 | 1.77715416 | 5.48688201 | -0.2678672 | 0.79244302 | 0.99999785 | -6.3495333 | down |
| KI - WT, cell_origin: iPSC | Q5VZK9 | CARMIL1 | 25 | Homo sapien | 0.31694938 | -2.2096932 | 2.84359196 | 0.33844515 | 0.26733047 | 0.79284846 | 0.99999785 | -6.3496815 | up |
| KI - WT, cell_origin: iPSC | O60547 | GMDS | 13 | Homo sapien | -0.1316382 | -1.1820952 | 0.91881877 | 0.31861255 | -0.2670583 | 0.7930541 | 0.99999785 | -6.3497565 | down |
| KI - WT, cell_origin: iPSC | P42166 | TMPO | 16 | Homo sapien | -0.0916998 | -0.8230854 | 0.63968582 | -0.0404611 | -0.2670319 | 0.79304917 | 0.99999785 | -6.3497791 | down |
| KI - WT, cell_origin: iPSC | O43148 | RNMT | 8 | Homo sapien | -0.2027281 | -1.8235473 | 1.41809103 | 0.26160122 | -0.2665519 | 0.79343668 | 0.99999785 | -6.3498959 | down |
| KI - WT, cell_origin: iPSC | Q92567 | FAM168A | 2 | Homo sapien | 0.13226682 | -0.9255179 | 1.19005151 | -0.4387243 | 0.26647471 | 0.79349503 | 0.99999785 | -6.3499172 | up |
| KI - WT, cell_origin: iPSC | Q15056 | EIF4H | 6 | Homo sapien | 0.12667467 | -0.8883565 | 1.14170583 | 0.26771788 | 0.26595783 | 0.79388564 | 0.99999785 | -6.3500592 | up |
| KI - WT, cell_origin: iPSC | A0AVT1 | UBA6 | 21 | Homo sapien | 0.08993608 | -0.6306157 | 0.8104879 | 0.62133676 | 0.26584975 | 0.79394506 | 0.99999785 | -6.3501025 | up |
| KI - WT, cell_origin: iPSC | Q5T7P8 | SYT6 | 7 | Homo sapien | 0.07739761 | -0.5441487 | 0.69894388 | -1.9335653 | 0.26521238 | 0.79442424 | 0.99999785 | -6.3502787 | up |
| KI - WT, cell_origin: iPSC | Q96AX1 | VPS33A | 14 | Homo sapien | -0.0620838 | -0.5606806 | 0.43651312 | 0.66709612 | -0.2651967 | 0.79443609 | 0.99999785 | -6.350283 | down |
| KI - WT, cell_origin: iPSC | O75494 | SRSF10 | 10 | Homo sapien | -0.1144948 | -1.0346178 | 0.80562809 | 1.25538175 | -0.265181 | 0.79447277 | 0.99999785 | -6.3502721 | down |
| KI - WT, cell_origin: iPSC | Q8IW93 | ARHGEF19 | 13 | Homo sapien | 0.23002029 | -1.619048 | 2.07908856 | 5.48772135 | 0.26510356 | 0.79453133 | 0.99999785 | -6.3502933 | up |
| KI - WT, cell_origin: iPSC | Q9Y4E8 | USP15 | 27 | Homo sapien | 0.08381401 | -0.5919368 | 0.75956477 | 1.24557967 | 0.26416164 | 0.79521872 | 0.99999785 | -6.3505656 | up |
| KI - WT, cell_origin: iPSC | Q13868 | EXOSC2 | 6 | Homo sapien | 0.13164662 | -0.9325198 | 1.1958131 | -1.3404875 | 0.26359201 | 0.7956676 | 0.99999785 | -6.3507096 | up |
| KI - WT, cell_origin: iPSC | Q14444 | CAPRIN1 | 17 | Homo sapien | -0.1393932 | -1.266621 | 0.9878347 | 4.46726989 | -0.2635313 | 0.79572014 | 0.99999785 | -6.3507221 | down |
| KI - WT, cell_origin: iPSC | Q5HYI8 | RABL3 | 2 | Homo sapien | -0.0958523 | -0.8730816 | 0.68137706 | -1.1371695 | -0.2626593 | 0.79635505 | 0.99999785 | -6.3509738 | down |
| KI - WT, cell_origin: iPSC | Q9H6W3 | RIOX1 | 4 | Homo sapien | -0.0612929 | -0.5589722 | 0.4363864 | -0.2957103 | -0.2623011 | 0.79662606 | 0.99999785 | -6.3510708 | down |
| KI - WT, cell_origin: iPSC | P63151 | PPP2R2A | 14 | Homo sapien | 0.07508166 | -0.5366799 | 0.68684325 | 0.28432479 | 0.26139144 | 0.79731445 | 0.99999785 | -6.3513165 | up |
| KI - WT, cell_origin: iPSC | Q8TC07 | TBC1D15 | 12 | Homo sapien | 0.10810508 | -0.7732337 | 0.9894438 | 0.44720825 | 0.26135771 | 0.79735788 | 0.99999785 | -6.3513148 | up |
| KI - WT, cell_origin: iPSC | Q13435 | SF3B2 | 28 | Homo sapien | -0.1083667 | -0.9923523 | 0.7756189 | 0.55051893 | -0.2612093 | 0.79747077 | 0.99999785 | -6.3513545 | down |
| KI - WT, cell_origin: iPSC | Q8N5H3 | FAM89B | 3 | Homo sapien | -0.1984736 | -1.8188427 | 1.42189551 | -1.0653596 | -0.2610304 | 0.79761209 | 0.99999785 | -6.3513991 | down |

| | | | | | | | | | | | | | |
|---|---|---|---|---|---|---|---|---|---|---|---|---|---|
| KI - WT, cell_origin: iPSC | Q15181 | PPA1 | 15 | Homo sapien | 0.12167563 | -0.8729405 | 1.11629178 | 2.02408035 | 0.26070566 | 0.79785789 | 0.99999785 | -6.3514865 | up |
| KI - WT, cell_origin: iPSC | Q9GZP4 | PITHD1 | 13 | Homo sapien | 0.09595039 | -0.688748 | 0.88064879 | 0.62125158 | 0.26047251 | 0.79801725 | 0.99999785 | -6.3515595 | up |
| KI - WT, cell_origin: iPSC | O60763 | USO1 | 15 | Homo sapien | 0.15635073 | -1.1239335 | 1.43663495 | 0.29435659 | 0.26025308 | 0.79820045 | 0.99999785 | -6.3516082 | up |
| KI - WT, cell_origin: iPSC | Q9HAV4 | XPO5 | 27 | Homo sapien | -0.116693 | -1.0721402 | 0.83875414 | -1.263364 | -0.2601694 | 0.79824674 | 0.99999785 | -6.3516409 | down |
| KI - WT, cell_origin: iPSC | O95336 | PGLS | 5 | Homo sapien | 0.08436208 | -0.6078868 | 0.77661097 | -0.566641 | 0.2595522 | 0.79870676 | 0.99999785 | -6.3518108 | up |
| KI - WT, cell_origin: iPSC | Q8N9T8 | KRI1 | 7 | Homo sapien | 0.1104639 | -0.7968347 | 1.01776254 | -1.846809 | 0.25930435 | 0.79889445 | 0.99999785 | -6.3518771 | up |
| KI - WT, cell_origin: iPSC | Q92600 | CNOT9 | 13 | Homo sapien | 0.05999516 | -0.4333848 | 0.55337509 | 1.33208641 | 0.25898489 | 0.79913636 | 0.99999785 | -6.3519625 | up |
| KI - WT, cell_origin: iPSC | Q9H7D7 | WDR26 | 9 | Homo sapien | 0.13104462 | -0.9474773 | 1.20956657 | -1.8104392 | 0.25882612 | 0.79926381 | 0.99999785 | -6.3520006 | up |
| KI - WT, cell_origin: iPSC | P23921 | RRM1 | 11 | Homo sapien | 0.15920467 | -1.1545679 | 1.47297727 | -1.2378723 | 0.25824859 | 0.79971817 | 0.99999785 | -6.3521445 | up |
| KI - WT, cell_origin: iPSC | Q09028 | RBBP4 | 9 | Homo sapien | -0.1202578 | -1.1138641 | 0.87334855 | 0.05236748 | -0.2579296 | 0.79995978 | 0.99999785 | -6.3522295 | down |
| KI - WT, cell_origin: iPSC | Q96QG7 | MTMR9 | 4 | Homo sapien | 0.29785789 | -2.1631869 | 2.75890269 | -4.2207504 | 0.25792412 | 0.79996393 | 0.99999785 | -6.352231 | up |
| KI - WT, cell_origin: iPSC | Q9Y3B9 | RRP15 | 2 | Homo sapien | 0.13241077 | -0.9615659 | 1.22638745 | -2.1168311 | 0.25781663 | 0.80002639 | 0.99999785 | -6.3522709 | up |
| KI - WT, cell_origin: iPSC | Q9UBV8 | PEF1 | 7 | Homo sapien | -0.0874543 | -0.810742 | 0.63583349 | 3.79310407 | -0.2575835 | 0.80020782 | 0.99999785 | -6.35233 | down |
| KI - WT, cell_origin: iPSC | Q9UJV9 | DDX41 | 15 | Homo sapien | -0.0924953 | -0.8576009 | 0.67261031 | 0.42273351 | -0.2575121 | 0.80025743 | 0.99999785 | -6.3523517 | down |
| KI - WT, cell_origin: iPSC | P35579 | MYH9 | 63 | Homo sapien | -0.0684458 | -0.6356099 | 0.49871835 | 1.08726489 | -0.2570262 | 0.80062009 | 0.99999785 | -6.3524839 | down |
| KI - WT, cell_origin: iPSC | O95671 | ASMTL | 8 | Homo sapien | -0.1411758 | -1.3117497 | 1.02939812 | -1.9991766 | -0.2569222 | 0.80070813 | 0.99999785 | -6.352506 | down |
| KI - WT, cell_origin: iPSC | Q6UX04 | CWC27 | 10 | Homo sapien | -0.164469 | -1.5296239 | 1.20068586 | 0.14277391 | -0.2567465 | 0.80085605 | 0.99999785 | -6.3525438 | down |
| KI - WT, cell_origin: iPSC | Q9Y2W2 | WBP11 | 6 | Homo sapien | -0.3563663 | -3.3144532 | 2.6017206 | -0.6033178 | -0.2567368 | 0.80086343 | 0.99999785 | -6.3525464 | down |
| KI - WT, cell_origin: iPSC | P15880 | RPS2 | 19 | Homo sapien | -0.0468165 | -0.4366023 | 0.34296932 | 5.67051102 | -0.2558071 | 0.80154404 | 0.99999785 | -6.3528065 | down |
| KI - WT, cell_origin: iPSC | Q7Z2W4 | ZC3HAV1 | 27 | Homo sapien | 0.07576525 | -0.5565735 | 0.70810398 | 2.07872832 | 0.25518785 | 0.80201343 | 0.99999785 | -6.3529697 | up |
| KI - WT, cell_origin: iPSC | Q9Y2H2 | INPP5F | 3 | Homo sapien | 0.085213 | -0.6264498 | 0.79687582 | -2.3247124 | 0.25501825 | 0.802142 | 0.99999785 | -6.3530144 | up |
| KI - WT, cell_origin: iPSC | Q7Z417 | NUFIP2 | 19 | Homo sapien | -0.1633241 | -1.5342036 | 1.2075554 | 2.34874641 | -0.2538946 | 0.80301776 | 0.99999785 | -6.3532955 | down |
| KI - WT, cell_origin: iPSC | Q9P0V9 | SEPTIN10 | 4 | Homo sapien | 0.15744543 | -1.1641969 | 1.47908775 | -1.4965892 | 0.25387415 | 0.80303323 | 0.99999785 | -6.3533008 | up |
| KI - WT, cell_origin: iPSC | Q9Y2L1 | DIS3 | 6 | Homo sapien | -0.1334315 | -1.2553869 | 0.98852397 | -1.670872 | -0.2533895 | 0.80339204 | 0.99999785 | -6.3534328 | down |
| KI - WT, cell_origin: iPSC | Q93008 | USP9X | 74 | Homo sapien | -0.063689 | -0.5992405 | 0.47186241 | 3.19137607 | -0.2532813 | 0.80345916 | 0.99999785 | -6.3534699 | down |
| KI - WT, cell_origin: iPSC | P13611 | VCAN | 2 | Homo sapien | 0.42241242 | -3.1349334 | 3.97975822 | 0.74797016 | 0.25305386 | 0.8036553 | 0.99999785 | -6.3535155 | up |
| KI - WT, cell_origin: iPSC | Q6NXR4 | TTI2 | 2 | Homo sapien | -0.0840063 | -0.7930084 | 0.62499569 | -1.0999762 | -0.2523506 | 0.80416521 | 0.99999785 | -6.3537128 | down |
| KI - WT, cell_origin: iPSC | Q8IXB1 | DNAJC10 | 9 | Homo sapien | -0.1502314 | -1.4196933 | 1.11923055 | -1.0853066 | -0.252199 | 0.80430377 | 0.99999785 | -6.3537384 | down |
| KI - WT, cell_origin: iPSC | Q9UPN3 | MACF1 | 71 | Homo sapien | 0.08835549 | -0.659088 | 0.83579903 | -1.1349546 | 0.2517647 | 0.80460972 | 0.99999785 | -6.3538651 | up |
| KI - WT, cell_origin: iPSC | P16333 | NCK1 | 17 | Homo sapien | -0.1316269 | -1.2458767 | 0.98262298 | 0.2582654 | -0.251747 | 0.80464662 | 0.99999785 | -6.353856 | down |
| KI - WT, cell_origin: iPSC | Q96IJ6 | GMPPA | 4 | Homo sapien | 0.13074755 | -0.978009 | 1.23950413 | -1.9803828 | 0.25119747 | 0.80504716 | 0.99999785 | -6.3540083 | up |
| KI - WT, cell_origin: iPSC | Q4L235 | AASDH | 2 | Homo sapien | -0.1588537 | -1.5057592 | 1.18805187 | -4.0548247 | -0.2511885 | 0.80504696 | 0.99999785 | -6.3540147 | down |
| KI - WT, cell_origin: iPSC | Q6PGN9 | PSRC1 | 3 | Homo sapien | -0.1164761 | -1.1042727 | 0.8713206 | -1.3147955 | -0.2511811 | 0.80505954 | 0.99999785 | -6.3540125 | down |
| KI - WT, cell_origin: iPSC | Q13247 | SRSF6 | 11 | Homo sapien | -0.0681282 | -0.6463885 | 0.51013215 | 2.66580634 | -0.2509244 | 0.80524739 | 0.99999785 | -6.3540831 | down |
| KI - WT, cell_origin: iPSC | O00743 | PPP6C | 7 | Homo sapien | 0.09276644 | -0.694713 | 0.88024589 | -0.8781475 | 0.25090867 | 0.80526152 | 0.99999785 | -6.3540859 | up |
| KI - WT, cell_origin: iPSC | Q9NZM5 | NOP53 | 2 | Homo sapien | -0.1895176 | -1.7999119 | 1.42087673 | -2.3682275 | -0.2507955 | 0.80536869 | 0.99999785 | -6.3541029 | down |
| KI - WT, cell_origin: iPSC | Q15652 | JMJD1C | 7 | Homo sapien | -0.2317543 | -2.2004135 | 1.73690486 | -4.7252906 | -0.2507246 | 0.80539905 | 0.99999785 | -6.3541348 | down |
| KI - WT, cell_origin: iPSC | O95382 | MAP3K6 | 8 | Homo sapien | -0.0963417 | -0.9154356 | 0.72275209 | -1.1390898 | -0.2505108 | 0.80556185 | 0.99999785 | -6.3541898 | down |
| KI - WT, cell_origin: iPSC | Q9P258 | RCC2 | 25 | Homo sapien | -0.1617436 | -1.5379721 | 1.21448493 | 3.64439409 | -0.2504603 | 0.80562305 | 0.99999785 | -6.3541896 | down |
| KI - WT, cell_origin: iPSC | Q9H1B7 | IRF2BPL | 11 | Homo sapien | -0.1830414 | -1.7417833 | 1.37570049 | -0.8297296 | -0.250252 | 0.80578118 | 0.99999785 | -6.3542434 | down |
| KI - WT, cell_origin: iPSC | Q8N567 | ZCCHC9 | 3 | Homo sapien | 0.1438931 | -1.0822636 | 1.37004975 | -1.2241674 | 0.25009003 | 0.80590409 | 0.99999785 | -6.3542853 | up |
| KI - WT, cell_origin: iPSC | Q04695 | KRT17 | 9 | Homo sapien | -0.2244631 | -2.1372338 | 1.6883076 | -1.8250069 | -0.2500831 | 0.80590933 | 0.99999785 | -6.354287 | down |
| KI - WT, cell_origin: iPSC | Q8TBF4 | ZCRB1 | 2 | Homo sapien | -0.1036234 | -0.9863196 | 0.77907271 | -1.0470463 | -0.2500693 | 0.80590305 | 0.99999785 | -6.3543004 | down |
| KI - WT, cell_origin: iPSC | Q3YEC7 | RABL6 | 12 | Homo sapien | 0.15216919 | -1.1454684 | 1.44980673 | 0.57499199 | 0.24990543 | 0.80604421 | 0.99999785 | -6.3543329 | up |
| KI - WT, cell_origin: iPSC | Q8N6R0 | METTL13 | 3 | Homo sapien | -0.0843543 | -0.8034332 | 0.63472469 | -1.8151862 | -0.2498447 | 0.806067 | 0.99999785 | -6.3543621 | down |
| KI - WT, cell_origin: iPSC | Q13838 | DDX39B | 9 | Homo sapien | -0.2249002 | -2.1451741 | 1.69537372 | -0.9564672 | -0.2495911 | 0.80628285 | 0.99999785 | -6.354414 | down |

| | | | | | | | | | | | | | |
|---|---|---|---|---|---|---|---|---|---|---|---|---|---|
| KI - WT, cell_origin: iPSC | Q03252 | LMNB2 | 33 | Homo sapien | 0.22705355 | -1.7117744 | 2.16588154 | 1.22362971 | 0.24956943 | 0.80629927 | 0.99999785 | -6.3544195 | up |
| KI - WT, cell_origin: iPSC | Q14684 | RRP1B | 11 | Homo sapien | -0.1183554 | -1.1302066 | 0.89349576 | -0.9919622 | -0.2492539 | 0.80653599 | 0.99999785 | -6.3545024 | down |
| KI - WT, cell_origin: iPSC | Q7Z478 | DHX29 | 13 | Homo sapien | -0.0732852 | -0.7000636 | 0.55349314 | -0.9550171 | -0.2490245 | 0.80668973 | 0.99999785 | -6.3545733 | down |
| KI - WT, cell_origin: iPSC | Q8NHM5 | KDM2B | 6 | Homo sapien | 0.14551651 | -1.1008567 | 1.39188971 | -1.2911255 | 0.24880925 | 0.80687642 | 0.99999785 | -6.3546151 | up |
| KI - WT, cell_origin: iPSC | P52756 | RBM5 | 7 | Homo sapien | 0.1442815 | -1.0950036 | 1.38356664 | -1.4058141 | 0.24810858 | 0.80740849 | 0.99999785 | -6.3547949 | up |
| KI - WT, cell_origin: iPSC | P08238 | HSP90AB1 | 30 | Homo sapien | -0.0952123 | -0.9130407 | 0.72261607 | 3.35592597 | -0.2480856 | 0.80742315 | 0.99999785 | -6.3548024 | down |
| KI - WT, cell_origin: iPSC | Q8NEZ5 | FBXO22 | 7 | Homo sapien | -0.1367722 | -1.3117215 | 1.03817712 | -0.6345839 | -0.2480739 | 0.80743482 | 0.99999785 | -6.3548038 | down |
| KI - WT, cell_origin: iPSC | Q9BZF1 | OSBPL8 | 2 | Homo sapien | -0.3129534 | -3.0064241 | 2.38051739 | -2.3439119 | -0.2476109 | 0.80778648 | 0.99999785 | -6.3549223 | down |
| KI - WT, cell_origin: iPSC | Q8TCG1 | CIP2A | 11 | Homo sapien | 0.10605421 | -0.8078203 | 1.01992869 | -0.6263602 | 0.24724306 | 0.80805537 | 0.99999785 | -6.3550223 | up |
| KI - WT, cell_origin: iPSC | P41250 | GARS1 | 6 | Homo sapien | -0.087774 | -0.8441988 | 0.66865068 | -1.6157442 | -0.2471383 | 0.80812239 | 0.99999785 | -6.3550562 | down |
| KI - WT, cell_origin: iPSC | Q96LB3 | IFT74 | 11 | Homo sapien | 0.05850356 | -0.4459399 | 0.56294697 | -0.8104233 | 0.24700712 | 0.80822206 | 0.99999785 | -6.3550897 | up |
| KI - WT, cell_origin: iPSC | O75330 | HMMR | 22 | Homo sapien | 0.20777898 | -1.5860219 | 2.00157986 | -0.8905534 | 0.24684807 | 0.80836593 | 0.99999785 | -6.355117 | up |
| KI - WT, cell_origin: iPSC | Q9NPH2 | ISYNA1 | 9 | Homo sapien | -0.1268813 | -1.2260689 | 0.97230636 | -0.143761 | -0.245996 | 0.80901332 | 0.99999785 | -6.3553338 | down |
| KI - WT, cell_origin: iPSC | P98179 | RBM3 | 6 | Homo sapien | -0.0971359 | -0.9391721 | 0.74490023 | 1.04734833 | -0.2457708 | 0.8091739 | 0.99999785 | -6.3553971 | down |
| KI - WT, cell_origin: iPSC | P53680 | AP2S1 | 4 | Homo sapien | -0.0795292 | -0.769813 | 0.61075457 | 0.83675643 | -0.2453798 | 0.80945872 | 0.99999785 | -6.3555032 | down |
| KI - WT, cell_origin: iPSC | O75608 | LYPLA1 | 2 | Homo sapien | 0.06563315 | -0.5057034 | 0.63696966 | -0.4527047 | 0.24466449 | 0.81000245 | 0.99999785 | -6.3556841 | up |
| KI - WT, cell_origin: iPSC | Q9UKF6 | CPSF3 | 8 | Homo sapien | -0.1266741 | -1.2319896 | 0.97864141 | -1.1185637 | -0.2442328 | 0.81035341 | 0.99999785 | -6.3557801 | down |
| KI - WT, cell_origin: iPSC | P09917 | ALOX5 | 5 | Homo sapien | -0.1007225 | -0.9819983 | 0.78055325 | -0.9239771 | -0.2434629 | 0.81092285 | 0.99999785 | -6.355983 | down |
| KI - WT, cell_origin: iPSC | O60942 | RNGTT | 6 | Homo sapien | -0.146349 | -1.427704 | 1.13500606 | -1.3386744 | -0.2434011 | 0.81098574 | 0.99999785 | -6.3559895 | down |
| KI - WT, cell_origin: iPSC | Q08554 | DSC1 | 2 | Homo sapien | 0.17461512 | -1.355655 | 1.70488529 | 0.04074951 | 0.24317342 | 0.81115887 | 0.99999785 | -6.3560467 | up |
| KI - WT, cell_origin: iPSC | Q9NP64 | ZCCHC17 | 3 | Homo sapien | -0.1401286 | -1.3691455 | 1.08888823 | -0.3000559 | -0.2429805 | 0.81130558 | 0.99999785 | -6.3560952 | down |
| KI - WT, cell_origin: iPSC | Q9H3K6 | BOLA2 | 4 | Homo sapien | -0.0804492 | -0.7863463 | 0.62544795 | 2.007049 | -0.2427289 | 0.81147444 | 0.99999785 | -6.356171 | down |
| KI - WT, cell_origin: iPSC | P22102 | GART | 29 | Homo sapien | -0.1333926 | -1.3045789 | 1.03779367 | 0.26248908 | -0.2427215 | 0.81150256 | 0.99999785 | -6.3561601 | down |
| KI - WT, cell_origin: iPSC | O14744 | PRMT5 | 10 | Homo sapien | -0.1572662 | -1.5388725 | 1.22434006 | 0.33639808 | -0.2425792 | 0.81161078 | 0.99999785 | -6.3561958 | down |
| KI - WT, cell_origin: iPSC | P06753 | TPM3 | 5 | Homo sapien | -0.1104139 | -1.08186 | 0.86103229 | 1.85129087 | -0.2422185 | 0.81188511 | 0.99999785 | -6.3562861 | down |
| KI - WT, cell_origin: iPSC | Q99700 | ATXN2 | 15 | Homo sapien | 0.11351677 | -0.8861133 | 1.11314684 | 1.66176854 | 0.24200431 | 0.81204802 | 0.99999785 | -6.3563396 | up |
| KI - WT, cell_origin: iPSC | O95429 | BAG4 | 4 | Homo sapien | 0.13129348 | -1.0261849 | 1.28877187 | -0.1542537 | 0.24173116 | 0.8122558 | 0.99999785 | -6.3564079 | up |
| KI - WT, cell_origin: iPSC | P35520 | CBS | 20 | Homo sapien | -0.2439943 | -2.3986396 | 1.91065087 | 0.98031341 | -0.2413272 | 0.81256307 | 0.99999785 | -6.3565086 | down |
| KI - WT, cell_origin: iPSC | Q96RF0 | SNX18 | 8 | Homo sapien | -0.0926819 | -0.910983 | 0.72561925 | -0.6807813 | -0.2412618 | 0.81259613 | 0.99999785 | -6.3565344 | down |
| KI - WT, cell_origin: iPSC | P48643 | CCT5 | 20 | Homo sapien | 0.06911739 | -0.5410548 | 0.67928959 | 0.86055989 | 0.24125404 | 0.81259629 | 0.99999785 | -6.3565395 | up |
| KI - WT, cell_origin: iPSC | P16402 | H1-3 | 2 | Homo sapien | -0.3648066 | -3.5973325 | 2.8677193 | -2.9851683 | -0.2405044 | 0.81318914 | 0.99999785 | -6.3567134 | down |
| KI - WT, cell_origin: iPSC | O95782 | AP2A1 | 40 | Homo sapien | 0.07014633 | -0.5519881 | 0.6922808 | 2.19044503 | 0.24013773 | 0.81344581 | 0.99999785 | -6.3568169 | up |
| KI - WT, cell_origin: iPSC | Q04637 | EIF4G1 | 33 | Homo sapien | 0.07491567 | -0.5900946 | 0.73992594 | 1.67990061 | 0.23992969 | 0.81360415 | 0.99999785 | -6.3568685 | up |
| KI - WT, cell_origin: iPSC | P14373 | TRIM27 | 3 | Homo sapien | 0.10206372 | -0.8052449 | 1.00937234 | 0.65679338 | 0.23971039 | 0.81379068 | 0.99999785 | -6.3569118 | up |
| KI - WT, cell_origin: iPSC | Q9NV88 | INTS9 | 3 | Homo sapien | 0.11777796 | -0.9301485 | 1.16570438 | -1.954087 | 0.23940264 | 0.8140101 | 0.99999785 | -6.3569962 | up |
| KI - WT, cell_origin: iPSC | A0AV96 | RBM47 | 12 | Homo sapien | -0.1144694 | -1.1342365 | 0.90529767 | -0.1258991 | -0.2392164 | 0.8141694 | 0.99999785 | -6.3570324 | down |
| KI - WT, cell_origin: iPSC | Q15058 | KIF14 | 22 | Homo sapien | -0.1120176 | -1.1102776 | 0.88624253 | -1.0982069 | -0.2390972 | 0.81425419 | 0.99999785 | -6.3570652 | down |
| KI - WT, cell_origin: iPSC | P23588 | EIF4B | 18 | Homo sapien | -0.0794227 | -0.7887543 | 0.62990895 | 1.22406193 | -0.2384739 | 0.81471291 | 0.99999785 | -6.3572277 | down |
| KI - WT, cell_origin: iPSC | Q8N163 | CCAR2 | 11 | Homo sapien | -0.1200997 | -1.1946077 | 0.95440826 | -0.0130395 | -0.2381961 | 0.81494612 | 0.99999785 | -6.357284 | down |
| KI - WT, cell_origin: iPSC | Q01082 | SPTBN1 | 44 | Homo sapien | -0.0778862 | -0.7744492 | 0.61867679 | 0.11000927 | -0.238144 | 0.81496361 | 0.99999785 | -6.3573092 | down |
| KI - WT, cell_origin: iPSC | P07996 | THBS1 | 36 | Homo sapien | 0.22410247 | -1.7879919 | 2.2361968 | 0.33169765 | 0.23735625 | 0.81558565 | 0.99999785 | -6.3574903 | up |
| KI - WT, cell_origin: iPSC | Q9UQB8 | BAIAP2 | 9 | Homo sapien | 0.18343935 | -1.4649652 | 1.83184392 | -1.4574899 | 0.23715433 | 0.81573943 | 0.99999785 | -6.3575397 | up |
| KI - WT, cell_origin: iPSC | Q9NPI1 | BRD7 | 5 | Homo sapien | -0.1572144 | -1.5759825 | 1.26155376 | -1.7244781 | -0.2361474 | 0.81650637 | 0.99999785 | -6.3577859 | down |
| KI - WT, cell_origin: iPSC | Q9P107 | GMIP | 4 | Homo sapien | -0.1539367 | -1.5443739 | 1.23650047 | -2.5432982 | -0.2358696 | 0.81670789 | 0.99999785 | -6.3578592 | down |
| KI - WT, cell_origin: iPSC | Q96GM8 | TOE1 | 12 | Homo sapien | 0.12810941 | -1.0311945 | 1.28741333 | -0.7525746 | 0.2354974 | 0.81700159 | 0.99999785 | -6.3579442 | up |
| KI - WT, cell_origin: iPSC | A4D1P6 | WDR91 | 11 | Homo sapien | -0.0776713 | -0.780741 | 0.62539836 | -0.1056277 | -0.2352892 | 0.81713831 | 0.99999785 | -6.3580069 | down |

| | | | | | | | | | | | | |
|---|---|---|---|---|---|---|---|---|---|---|---|---|
| KI - WT, cell_origin: iPSC | Q13813 | SPTAN1 | 51 | Homo sapien | 0.06498118 | -0.5233557 | 0.65331808 | -0.3890573 | 0.23523458 | 0.81717995 | 0.99999785 | -6.3580202 up |
| KI - WT, cell_origin: iPSC | Q09472 | EP300 | 8 | Homo sapien | -0.1015032 | -1.0207468 | 0.8177405 | -1.4585295 | -0.2351975 | 0.8172119 | 0.99999785 | -6.3580272 down |
| KI - WT, cell_origin: iPSC | Q9BXJ9 | NAA15 | 9 | Homo sapien | 0.15462979 | -1.246668 | 1.55592755 | -0.8963469 | 0.23516089 | 0.81725799 | 0.99999785 | -6.358026 up |
| KI - WT, cell_origin: iPSC | Q96KP4 | CNDP2 | 6 | Homo sapien | 0.07078111 | -0.5704802 | 0.71204237 | -0.3990883 | 0.23508343 | 0.81729515 | 0.99999785 | -6.3580569 up |
| KI - WT, cell_origin: iPSC | Q6GQQ9 | OTUD7B | 9 | Homo sapien | 0.10152562 | -0.8191428 | 1.02219405 | -1.2815468 | 0.23489794 | 0.81744209 | 0.99999785 | -6.3580988 up |
| KI - WT, cell_origin: iPSC | Q92522 | H1-10 | 6 | Homo sapien | 0.16201689 | -1.3080413 | 1.63207506 | 0.76767289 | 0.23487033 | 0.8174794 | 0.99999785 | -6.3580966 up |
| KI - WT, cell_origin: iPSC | Q92598 | HSPH1 | 27 | Homo sapien | 0.12286871 | -0.992575 | 1.23831247 | -0.0937209 | 0.23474481 | 0.81757504 | 0.99999785 | -6.358127 up |
| KI - WT, cell_origin: iPSC | Q6P1J9 | CDC73 | 14 | Homo sapien | -0.1297055 | -1.310512 | 1.05110109 | -0.1470857 | -0.2340895 | 0.8180745 | 0.99999785 | -6.3582857 down |
| KI - WT, cell_origin: iPSC | O43439 | CBFA2T2 | 3 | Homo sapien | -0.0800592 | -0.8087551 | 0.64863676 | -1.4095015 | -0.2339939 | 0.81812559 | 0.99999785 | -6.3583208 down |
| KI - WT, cell_origin: iPSC | O14578 | CIT | 37 | Homo sapien | -0.0995511 | -1.0071891 | 0.8080869 | -0.3479861 | -0.233691 | 0.81837043 | 0.99999785 | -6.3583862 down |
| KI - WT, cell_origin: iPSC | Q86X55 | CARM1 | 11 | Homo sapien | 0.10740449 | -0.8726552 | 1.0874642 | 0.64013548 | 0.23354591 | 0.8184888 | 0.99999785 | -6.358417 up |
| KI - WT, cell_origin: iPSC | Q9UKJ3 | GPATCH8 | 22 | Homo sapien | -0.1394573 | -1.4120051 | 1.13309042 | -0.994578 | -0.2335445 | 0.81848989 | 0.99999785 | -6.3584173 down |
| KI - WT, cell_origin: iPSC | Q00341 | HDLBP | 37 | Homo sapien | -0.0695621 | -0.7041677 | 0.5650436 | 0.94582437 | -0.2334577 | 0.81853434 | 0.99999785 | -6.3584502 down |
| KI - WT, cell_origin: iPSC | P11234 | RALB | 14 | Homo sapien | 0.14559939 | -1.1876941 | 1.4788929 | 6.21705167 | 0.23272131 | 0.81911744 | 0.99999785 | -6.3586156 up |
| KI - WT, cell_origin: iPSC | Q00839 | HNRNPU | 29 | Homo sapien | -0.0937368 | -0.9524655 | 0.76499194 | 2.41790853 | -0.2325848 | 0.81921532 | 0.99999785 | -6.3586518 down |
| KI - WT, cell_origin: iPSC | P16284 | PECAM1 | 2 | Homo sapien | -0.1580231 | -1.6071554 | 1.29110911 | -1.4024401 | -0.2323887 | 0.81937104 | 0.99999785 | -6.3586955 down |
| KI - WT, cell_origin: iPSC | P56192 | MARS1 | 16 | Homo sapien | 0.07503113 | -0.6126228 | 0.7626851 | 1.10632243 | 0.23238667 | 0.81935098 | 0.99999785 | -6.3587078 up |
| KI - WT, cell_origin: iPSC | P09382 | LGALS1 | 6 | Homo sapien | -0.1218807 | -1.2402993 | 0.99653794 | 0.27847651 | -0.2322378 | 0.81948611 | 0.99999785 | -6.3587317 down |
| KI - WT, cell_origin: iPSC | P51991 | HNRNPA3 | 16 | Homo sapien | -0.1090063 | -1.1095118 | 0.89149916 | 1.59386516 | -0.2321852 | 0.81952622 | 0.99999785 | -6.3587443 down |
| KI - WT, cell_origin: iPSC | O00160 | MYO1F | 6 | Homo sapien | 0.11231927 | -0.9197877 | 1.14442621 | -1.3720238 | 0.23185192 | 0.81977041 | 0.99999785 | -6.3588296 up |
| KI - WT, cell_origin: iPSC | P53671 | LIMK2 | 16 | Homo sapien | -0.0970175 | -0.9888497 | 0.79481473 | -0.9100131 | -0.2317313 | 0.8198572 | 0.99999785 | -6.3588613 down |
| KI - WT, cell_origin: iPSC | Q13574 | DGKZ | 12 | Homo sapien | 0.06823058 | -0.5598755 | 0.69633662 | 0.22182308 | 0.23135867 | 0.82013503 | 0.99999785 | -6.3589539 up |
| KI - WT, cell_origin: iPSC | P48507 | GCLM | 9 | Homo sapien | -0.0962778 | -0.9835332 | 0.79097748 | -0.8278745 | -0.2311509 | 0.82029988 | 0.99999785 | -6.3590001 down |
| KI - WT, cell_origin: iPSC | Q7Z3G6 | PRICKLE2 | 4 | Homo sapien | 0.23285894 | -1.9175073 | 2.38322518 | -2.3169494 | 0.23077186 | 0.82060411 | 0.99999785 | -6.3590823 up |
| KI - WT, cell_origin: iPSC | Q9NTJ3 | SMC4 | 11 | Homo sapien | -0.1279529 | -1.3096054 | 1.05369952 | -1.8478868 | -0.2307158 | 0.82063988 | 0.99999785 | -6.3590995 down |
| KI - WT, cell_origin: iPSC | O43396 | TXNL1 | 5 | Homo sapien | 0.08448196 | -0.6959541 | 0.86491797 | 0.31540839 | 0.23058628 | 0.82072973 | 0.99999785 | -6.3591352 up |
| KI - WT, cell_origin: iPSC | O60749 | SNX2 | 15 | Homo sapien | 0.06988959 | -0.5756546 | 0.71543374 | 0.27875768 | 0.23058243 | 0.82072719 | 0.99999785 | -6.3591391 up |
| KI - WT, cell_origin: iPSC | P21291 | CSRP1 | 9 | Homo sapien | -0.0972803 | -0.9973847 | 0.80282403 | 0.53737352 | -0.2302927 | 0.82096522 | 0.99999785 | -6.3591989 down |
| KI - WT, cell_origin: iPSC | O75808 | CAPN15 | 3 | Homo sapien | -0.1340463 | -1.3755654 | 1.10747283 | -1.8917022 | -0.2300763 | 0.82113214 | 0.99999785 | -6.3592493 down |
| KI - WT, cell_origin: iPSC | Q96PU4 | UHRF2 | 3 | Homo sapien | -0.1261766 | -1.2944992 | 1.0421459 | -2.7320618 | -0.2300148 | 0.82116025 | 0.99999785 | -6.3592741 down |
| KI - WT, cell_origin: iPSC | Q9BPX3 | NCAPG | 8 | Homo sapien | 0.09609283 | -0.7960129 | 0.98819855 | -1.3881786 | 0.22942754 | 0.82161094 | 0.99999785 | -6.359412 up |
| KI - WT, cell_origin: iPSC | O00401 | WASL | 3 | Homo sapien | 0.11540622 | -0.9566734 | 1.18748585 | -0.726648 | 0.22940587 | 0.82164626 | 0.99999785 | -6.359407 up |
| KI - WT, cell_origin: iPSC | Q92499 | DDX1 | 39 | Homo sapien | -0.1127685 | -1.1650373 | 0.93950037 | 3.40596865 | -0.2283828 | 0.82242702 | 0.99999785 | -6.359649 down |
| KI - WT, cell_origin: iPSC | O15357 | INPPL1 | 15 | Homo sapien | 0.06924215 | -0.5771114 | 0.71559567 | -0.5866847 | 0.2281603 | 0.82257565 | 0.99999785 | -6.3597128 up |
| KI - WT, cell_origin: iPSC | P04908;P0C0 | H2AC4;H2AC | 4 | Homo sapien | -0.1395936 | -1.4438913 | 1.16470403 | 3.55380162 | -0.2280821 | 0.82265652 | 0.99999785 | -6.3597199 down |
| KI - WT, cell_origin: iPSC | P42331 | ARHGAP25 | 4 | Homo sapien | 0.05731962 | -0.4793308 | 0.59397002 | -1.1213245 | 0.22748435 | 0.82309169 | 0.99999785 | -6.3598719 up |
| KI - WT, cell_origin: iPSC | O75396 | SEC22B | 10 | Homo sapien | 0.23060952 | -1.9307025 | 2.3919215 | 0.82778033 | 0.22738517 | 0.82318854 | 0.99999785 | -6.3598839 up |
| KI - WT, cell_origin: iPSC | P41227 | NAA10 | 4 | Homo sapien | -0.141724 | -1.4735199 | 1.190072 | -2.0498684 | -0.2267817 | 0.8236493 | 0.99999785 | -6.3600255 down |
| KI - WT, cell_origin: iPSC | P53384 | NUBP1 | 5 | Homo sapien | 0.09390674 | -0.7885211 | 0.97633463 | 0.69863211 | 0.22675484 | 0.82366471 | 0.99999785 | -6.3600345 up |
| KI - WT, cell_origin: iPSC | Q8TAV0 | FAM76A | 3 | Homo sapien | -0.2340121 | -2.4355115 | 1.96748728 | -2.2023482 | -0.2265281 | 0.82384291 | 0.99999785 | -6.3600849 down |
| KI - WT, cell_origin: iPSC | Q15084 | PDIA6 | 21 | Homo sapien | -0.3032581 | -3.1569689 | 2.55045283 | 3.52019829 | -0.2264669 | 0.82388965 | 0.99999785 | -6.3600992 down |
| KI - WT, cell_origin: iPSC | P0DME0;Q01 | SETSIP;SET | 4 | Homo sapien | -0.1434213 | -1.494528 | 1.20768535 | 0.5218551 | -0.2262176 | 0.82408001 | 0.99999785 | -6.3601575 down |
| KI - WT, cell_origin: iPSC | Q9UFC0 | LRWD1 | 3 | Homo sapien | -0.1249619 | -1.304372 | 1.05444818 | -1.2125243 | -0.2257954 | 0.82440241 | 0.99999785 | -6.3602561 down |
| KI - WT, cell_origin: iPSC | P62906 | RPL10A | 13 | Homo sapien | -0.0477242 | -0.4979371 | 0.4024887 | 3.87386256 | -0.225767 | 0.82440314 | 0.99999785 | -6.3602739 down |
| KI - WT, cell_origin: iPSC | Q7Z7G8 | VPS13B | 6 | Homo sapien | 0.11630371 | -0.9835554 | 1.21616285 | -2.0875937 | 0.22524538 | 0.82480638 | 0.99999785 | -6.3603928 up |
| KI - WT, cell_origin: iPSC | P62760 | VSNL1 | 3 | Homo sapien | 0.05586081 | -0.4723983 | 0.58411996 | -0.6758835 | 0.22521634 | 0.82482381 | 0.99999785 | -6.3604021 up |

| | | | | | | | | | | | | |
|---|---|---|---|---|---|---|---|---|---|---|---|---|
| KI - WT, cell_origin: iPSC | O43670 | ZNF207 | 7 | Homo sapien | -0.1385285 | -1.4508434 | 1.17378638 | 2.73333159 | -0.2249591 | 0.82504122 | 0.99999785 | -6.3604509 down |
| KI - WT, cell_origin: iPSC | Q9Y5X3 | SNX5 | 14 | Homo sapien | 0.06241953 | -0.5286005 | 0.65343955 | 0.89622394 | 0.22493548 | 0.82503837 | 0.99999785 | -6.3604674 up |
| KI - WT, cell_origin: iPSC | Q9UQ35 | SRRM2 | 51 | Homo sapien | -0.1321287 | -1.3859542 | 1.12169681 | 4.31643145 | -0.2245755 | 0.82533423 | 0.99999785 | -6.36054 down |
| KI - WT, cell_origin: iPSC | O43852 | CALU | 7 | Homo sapien | 0.15495888 | -1.3168827 | 1.62680048 | -1.5076363 | 0.22436638 | 0.82549401 | 0.99999785 | -6.3605885 up |
| KI - WT, cell_origin: iPSC | O75150 | RNF40 | 9 | Homo sapien | 0.14190853 | -1.2096817 | 1.49349872 | -1.399991 | 0.22375144 | 0.82596383 | 0.99999785 | -6.3607308 up |
| KI - WT, cell_origin: iPSC | Q92973 | TNPO1 | 28 | Homo sapien | -0.0554296 | -0.5830826 | 0.47222347 | 2.86552481 | -0.2237344 | 0.82595611 | 0.99999785 | -6.3607457 down |
| KI - WT, cell_origin: iPSC | Q8NB90 | SPATA5 | 6 | Homo sapien | -0.068723 | -0.7231158 | 0.5856697 | 0.04968604 | -0.2236679 | 0.82600694 | 0.99999785 | -6.3607611 down |
| KI - WT, cell_origin: iPSC | Q9Y446 | PKP3 | 4 | Homo sapien | 0.18104756 | -1.5477587 | 1.90985386 | -1.5393156 | 0.22317663 | 0.82640308 | 0.99999785 | -6.3608636 up |
| KI - WT, cell_origin: iPSC | P46779 | RPL28 | 10 | Homo sapien | -0.0611629 | -0.6452305 | 0.52290479 | 3.8663719 | -0.2230306 | 0.82649401 | 0.99999785 | -6.3609081 down |
| KI - WT, cell_origin: iPSC | P62266 | RPS23 | 5 | Homo sapien | 0.05745859 | -0.4915081 | 0.60642526 | 4.55702221 | 0.22291984 | 0.82657864 | 0.99999785 | -6.3609336 up |
| KI - WT, cell_origin: iPSC | Q8N3U4 | STAG2 | 5 | Homo sapien | 0.11042801 | -0.9450166 | 1.1658726 | -2.5045847 | 0.22283504 | 0.82664345 | 0.99999785 | -6.3609532 up |
| KI - WT, cell_origin: iPSC | Q96GQ7 | DDX27 | 6 | Homo sapien | -0.0860366 | -0.9087722 | 0.73669897 | 0.35521488 | -0.2227619 | 0.82670551 | 0.99999785 | -6.3609668 down |
| KI - WT, cell_origin: iPSC | Q9NZJ4 | SACS | 48 | Homo sapien | 0.13133912 | -1.1275043 | 1.39018256 | -0.8621512 | 0.22234366 | 0.82703969 | 0.99999785 | -6.3610553 up |
| KI - WT, cell_origin: iPSC | Q69YH5 | CDCA2 | 2 | Homo sapien | 0.07445748 | -0.6391999 | 0.78811484 | -1.9825891 | 0.22220726 | 0.82712333 | 0.99999785 | -6.3610975 up |
| KI - WT, cell_origin: iPSC | Q99848 | EBNA1BP2 | 6 | Homo sapien | 0.14096516 | -1.2129983 | 1.49492858 | -1.8000503 | 0.22187442 | 0.82739838 | 0.99999785 | -6.361163 up |
| KI - WT, cell_origin: iPSC | Q969G3 | SMARCE1 | 7 | Homo sapien | 0.1184782 | -1.0206913 | 1.25764771 | -0.5822373 | 0.2216422 | 0.82757589 | 0.99999785 | -6.3612163 up |
| KI - WT, cell_origin: iPSC | Q8TER5 | ARHGEF40 | 24 | Homo sapien | -0.1067158 | -1.1329835 | 0.91955192 | 4.14105025 | -0.2216003 | 0.82760791 | 0.99999785 | -6.3612258 down |
| KI - WT, cell_origin: iPSC | P43378 | PTPN9 | 9 | Homo sapien | -0.0977228 | -1.0383983 | 0.84295278 | -0.4446829 | -0.2213579 | 0.82778832 | 0.99999785 | -6.3612839 down |
| KI - WT, cell_origin: iPSC | P49585 | PCYT1A | 17 | Homo sapien | 0.09227734 | -0.795989 | 0.98054372 | 1.08031646 | 0.22135578 | 0.82778991 | 0.99999785 | -6.3612844 up |
| KI - WT, cell_origin: iPSC | Q9HAU0 | PLEKHA5 | 7 | Homo sapien | 0.12064946 | -1.0415739 | 1.28287284 | -1.2729959 | 0.22122701 | 0.82789331 | 0.99999785 | -6.3613113 up |
| KI - WT, cell_origin: iPSC | Q8TBX8 | PIP4K2C | 5 | Homo sapien | -0.1061477 | -1.1297889 | 0.91749358 | -1.7443932 | -0.2208866 | 0.82813831 | 0.99999785 | -6.361397 down |
| KI - WT, cell_origin: iPSC | A6NDU8 | C5orf51 | 2 | Homo sapien | -0.1266948 | -1.3498101 | 1.09642062 | -1.4270659 | -0.2207464 | 0.82826081 | 0.99999785 | -6.361421 down |
| KI - WT, cell_origin: iPSC | Q9ULJ7 | ANKRD50 | 8 | Homo sapien | 0.15681274 | -1.3589934 | 1.67261885 | -2.3059058 | 0.22046521 | 0.8284758 | 0.99999785 | -6.3614851 up |
| KI - WT, cell_origin: iPSC | O95104 | SCAF4 | 6 | Homo sapien | -0.1828932 | -1.955333 | 1.58954653 | -0.4703202 | -0.2199017 | 0.82890677 | 0.99999785 | -6.3616134 down |
| KI - WT, cell_origin: iPSC | P31942 | HNRNPH3 | 8 | Homo sapien | -0.1393177 | -1.4901985 | 1.21156305 | 1.49323137 | -0.2197818 | 0.82899845 | 0.99999785 | -6.3616406 down |
| KI - WT, cell_origin: iPSC | P09661 | SNRPA1 | 14 | Homo sapien | 0.14706434 | -1.2794193 | 1.57354794 | 2.87508234 | 0.21970652 | 0.829056 | 0.99999785 | -6.3616577 up |
| KI - WT, cell_origin: iPSC | Q01581 | HMGCS1 | 4 | Homo sapien | 0.0658174 | -0.5736765 | 0.70531127 | -1.465421 | 0.21920177 | 0.82942175 | 0.99999785 | -6.3617827 up |
| KI - WT, cell_origin: iPSC | Q9UL03 | INTS6 | 10 | Homo sapien | 0.13503219 | -1.1801312 | 1.45019557 | -1.4346153 | 0.21880637 | 0.82974453 | 0.99999785 | -6.3618617 up |
| KI - WT, cell_origin: iPSC | Q8TAG9 | EXOC6 | 11 | Homo sapien | -0.1069685 | -1.1493307 | 0.93539367 | -0.6456185 | -0.2186954 | 0.8298294 | 0.99999785 | -6.3618868 down |
| KI - WT, cell_origin: iPSC | P39023 | RPL3 | 24 | Homo sapien | -0.0762268 | -0.8191805 | 0.66672692 | 3.08930222 | -0.2185565 | 0.82992144 | 0.99999785 | -6.3619255 down |
| KI - WT, cell_origin: iPSC | P35244 | RPA3 | 4 | Homo sapien | -0.0672938 | -0.7239108 | 0.58932326 | -0.5521345 | -0.2182741 | 0.83013146 | 0.99999785 | -6.3619923 down |
| KI - WT, cell_origin: iPSC | Q9H9Z2 | LIN28A | 12 | Homo sapien | -0.1674075 | -1.8023158 | 1.4675008 | 2.50350493 | -0.2182146 | 0.83019725 | 0.99999785 | -6.3619953 down |
| KI - WT, cell_origin: iPSC | P36957 | DLST | 11 | Homo sapien | 0.2104393 | -1.8449649 | 2.26584347 | -0.0641395 | 0.21818856 | 0.83021718 | 0.99999785 | -6.3620012 up |
| KI - WT, cell_origin: iPSC | Q9NQW6 | ANLN | 14 | Homo sapien | -0.1092633 | -1.1764762 | 0.95794966 | -1.5773732 | -0.2181112 | 0.83026499 | 0.99999785 | -6.3620245 down |
| KI - WT, cell_origin: iPSC | Q9Y3F4 | STRAP | 18 | Homo sapien | -0.0510341 | -0.5493909 | 0.44732268 | 1.10773137 | -0.2181022 | 0.83026307 | 0.99999785 | -6.3620311 down |
| KI - WT, cell_origin: iPSC | P68400 | CSNK2A1 | 16 | Homo sapien | 0.08096457 | -0.7110068 | 0.87293589 | 1.76300624 | 0.2177727 | 0.83052119 | 0.99999785 | -6.3621022 up |
| KI - WT, cell_origin: iPSC | P08758 | ANXA5 | 9 | Homo sapien | -0.1830853 | -1.9751712 | 1.60900058 | -0.0773991 | -0.2177194 | 0.83057616 | 0.99999785 | -6.3621069 down |
| KI - WT, cell_origin: iPSC | Q9Y6Y8 | SEC23IP | 18 | Homo sapien | -0.072437 | -0.7810621 | 0.63618807 | 0.74916104 | -0.2177157 | 0.83055927 | 0.99999785 | -6.3621179 down |
| KI - WT, cell_origin: iPSC | P49792 | RANBP2 | 45 | Homo sapien | 0.07731609 | -0.6800425 | 0.83467469 | -0.8378075 | 0.21742426 | 0.83078184 | 0.99999785 | -6.3621836 up |
| KI - WT, cell_origin: iPSC | O43390 | HNRNPR | 17 | Homo sapien | 0.06593856 | -0.5810897 | 0.71296685 | 1.54645262 | 0.21704803 | 0.83106979 | 0.99999785 | -6.362268 up |
| KI - WT, cell_origin: iPSC | Q96PZ0 | PUS7 | 9 | Homo sapien | -0.0547615 | -0.5922356 | 0.48271261 | -0.6079418 | -0.2169989 | 0.83110742 | 0.99999785 | -6.3622791 down |
| KI - WT, cell_origin: iPSC | P47974 | ZFP36L2 | 4 | Homo sapien | -0.0970802 | -1.0499112 | 0.85575075 | -2.1058245 | -0.2169974 | 0.83110856 | 0.99999785 | -6.3622794 down |
| KI - WT, cell_origin: iPSC | O60341 | KDM1A | 13 | Homo sapien | 0.11142832 | -0.9843523 | 1.20720898 | -0.8567701 | 0.21670769 | 0.83135037 | 0.99999785 | -6.362334 up |
| KI - WT, cell_origin: iPSC | Q05682 | CALD1 | 15 | Homo sapien | -0.1033335 | -1.1205969 | 0.91392992 | 0.18257116 | -0.2164762 | 0.83152758 | 0.99999785 | -6.3623858 down |
| KI - WT, cell_origin: iPSC | Q8IVW6 | ARID3B | 6 | Homo sapien | -0.1784401 | -1.939297 | 1.58241678 | 1.57646553 | -0.2159587 | 0.83192365 | 0.99999785 | -6.3625015 down |
| KI - WT, cell_origin: iPSC | Q9H7X7 | IFT22 | 3 | Homo sapien | -0.0756583 | -0.8218155 | 0.67049883 | -1.3661576 | -0.2159564 | 0.83190542 | 0.99999785 | -6.3625122 down |

| | | | | | | | | | | | | | |
|---|---|---|---|---|---|---|---|---|---|---|---|---|---|
| KI - WT, cell_origin: iPSC | P18858 | LIG1 | 5 | Homo sapien | 0.11817186 | -1.0474789 | 1.28382264 | -2.3750508 | 0.21593811 | 0.83192277 | 0.99999785 | -6.3625146 | up |
| KI - WT, cell_origin: iPSC | Q96RW7 | HMCN1 | 28 | Homo sapien | -0.1232175 | -1.3402119 | 1.09377687 | 2.41179987 | -0.2157676 | 0.83206993 | 0.99999785 | -6.3625441 | down |
| KI - WT, cell_origin: iPSC | Q9Y6K5 | OAS3 | 12 | Homo sapien | 0.09525006 | -0.8454908 | 1.03599092 | 0.02350999 | 0.21575744 | 0.83207531 | 0.99999785 | -6.3625476 | up |
| KI - WT, cell_origin: iPSC | P43246 | MSH2 | 19 | Homo sapien | -0.1207588 | -1.3137781 | 1.0722605 | -0.1083332 | -0.2157117 | 0.83211277 | 0.99999785 | -6.3625566 | down |
| KI - WT, cell_origin: iPSC | Q9P1Y6 | PHRF1 | 6 | Homo sapien | -0.1584849 | -1.7252647 | 1.40829498 | -0.8733136 | -0.215567 | 0.83222354 | 0.99999785 | -6.3625888 | down |
| KI - WT, cell_origin: iPSC | O60879 | DIAPH2 | 27 | Homo sapien | -0.1154928 | -1.2576087 | 1.02662306 | 1.1783803 | -0.2154999 | 0.83227485 | 0.99999785 | -6.3626038 | down |
| KI - WT, cell_origin: iPSC | Q9BWD1 | ACAT2 | 3 | Homo sapien | 0.11537037 | -1.0264556 | 1.2571963 | -1.3236365 | 0.21532614 | 0.83240791 | 0.99999785 | -6.3626425 | up |
| KI - WT, cell_origin: iPSC | P49589 | CARS1 | 26 | Homo sapien | -0.1092872 | -1.1912292 | 0.97265476 | -0.0356631 | -0.2152622 | 0.83245686 | 0.99999785 | -6.3626567 | down |
| KI - WT, cell_origin: iPSC | Q15050 | RRS1 | 10 | Homo sapien | 0.11012578 | -0.9802654 | 1.20051694 | -1.190736 | 0.21523309 | 0.83247915 | 0.99999785 | -6.3626632 | up |
| KI - WT, cell_origin: iPSC | P11388 | TOP2A | 35 | Homo sapien | 0.15927263 | -1.4179759 | 1.7365212 | -1.3316702 | 0.21520054 | 0.83250407 | 0.99999785 | -6.3626704 | up |
| KI - WT, cell_origin: iPSC | P19174 | PLCG1 | 34 | Homo sapien | 0.06932248 | -0.6171334 | 0.75577835 | 0.90548318 | 0.21508057 | 0.832576 | 0.99999785 | -6.3627072 | up |
| KI - WT, cell_origin: iPSC | Q8IWR0 | ZC3H7A | 14 | Homo sapien | -0.0890388 | -0.9711387 | 0.79306115 | 0.45352925 | -0.2150767 | 0.83259356 | 0.99999785 | -6.3627007 | down |
| KI - WT, cell_origin: iPSC | Q14152 | EIF3A | 79 | Homo sapien | -0.0482454 | -0.5263672 | 0.42987646 | 3.63696092 | -0.2149101 | 0.83270651 | 0.99999785 | -6.3627451 | down |
| KI - WT, cell_origin: iPSC | O14980 | XPO1 | 28 | Homo sapien | -0.093802 | -1.0245794 | 0.8369754 | 0.42964518 | -0.2147675 | 0.83283563 | 0.99999785 | -6.3627666 | down |
| KI - WT, cell_origin: iPSC | P30520 | ADSS2 | 4 | Homo sapien | -0.0717528 | -0.7847494 | 0.64124391 | -0.4644648 | -0.2143339 | 0.83314783 | 0.99999785 | -6.3628729 | down |
| KI - WT, cell_origin: iPSC | P53582 | METAP1 | 6 | Homo sapien | 0.11470223 | -1.0276101 | 1.25701457 | -1.3963572 | 0.21398798 | 0.83343256 | 0.99999785 | -6.3629394 | up |
| KI - WT, cell_origin: iPSC | O94964 | SOGA1 | 16 | Homo sapien | -0.1156813 | -1.2682251 | 1.03686255 | -1.4185858 | -0.2138986 | 0.83350098 | 0.99999785 | -6.3629591 | down |
| KI - WT, cell_origin: iPSC | P42574 | CASP3 | 3 | Homo sapien | 0.09029691 | -0.8100738 | 0.99066764 | 0.22413205 | 0.21368972 | 0.83365569 | 0.99999785 | -6.363008 | up |
| KI - WT, cell_origin: iPSC | O43324 | EEF1E1 | 7 | Homo sapien | 0.11032145 | -0.9901665 | 1.21080937 | 0.82912602 | 0.21363728 | 0.83370115 | 0.99999785 | -6.3630169 | up |
| KI - WT, cell_origin: iPSC | P78316 | NOP14 | 5 | Homo sapien | 0.09037335 | -0.8108382 | 0.99158492 | -1.688926 | 0.21357641 | 0.83372799 | 0.99999785 | -6.3630403 | up |
| KI - WT, cell_origin: iPSC | Q9BQ67 | GRWD1 | 8 | Homo sapien | 0.0487855 | -0.438029 | 0.53560004 | 1.19229079 | 0.21343564 | 0.83383582 | 0.99999785 | -6.3630714 | up |
| KI - WT, cell_origin: iPSC | P16455 | MGMT | 7 | Homo sapien | 0.18796181 | -1.689937 | 2.06586067 | 0.54188883 | 0.21330439 | 0.83395612 | 0.99999785 | -6.3630903 | up |
| KI - WT, cell_origin: iPSC | Q8WU90 | ZC3H15 | 15 | Homo sapien | -0.1414806 | -1.5584765 | 1.27551534 | 1.60526889 | -0.2127799 | 0.83435789 | 0.99999785 | -6.3632059 | down |
| KI - WT, cell_origin: iPSC | C9JLW8 | MCRIP1 | 5 | Homo sapien | -0.0655425 | -0.7216659 | 0.59058084 | -0.1778375 | -0.2127538 | 0.83435819 | 0.99999785 | -6.3632215 | down |
| KI - WT, cell_origin: iPSC | Q6DCA0 | AMMECR1L | 4 | Homo sapien | -0.1518348 | -1.6746407 | 1.37097099 | -2.2942955 | -0.2124855 | 0.83458341 | 0.99999785 | -6.3632706 | down |
| KI - WT, cell_origin: iPSC | O60678 | PRMT3 | 7 | Homo sapien | -0.085853 | -0.9481006 | 0.77639468 | -0.786383 | -0.2121003 | 0.83486474 | 0.99999785 | -6.363362 | down |
| KI - WT, cell_origin: iPSC | P46109 | CRKL | 19 | Homo sapien | 0.07898596 | -0.7151803 | 0.87315219 | 4.091438 | 0.21195362 | 0.83499089 | 0.99999785 | -6.3633872 | up |
| KI - WT, cell_origin: iPSC | P26373 | RPL13 | 12 | Homo sapien | 0.04292313 | -0.3888743 | 0.47472054 | 4.76092283 | 0.21171472 | 0.83515434 | 0.99999785 | -6.3634494 | up |
| KI - WT, cell_origin: iPSC | O95235 | KIF20A | 20 | Homo sapien | 0.12731559 | -1.1575577 | 1.41218888 | -1.2168796 | 0.21116583 | 0.83559454 | 0.99999785 | -6.3635595 | up |
| KI - WT, cell_origin: iPSC | O00488 | ZNF593 | 2 | Homo sapien | -0.1346538 | -1.4941635 | 1.22485603 | -0.3808045 | -0.2110758 | 0.83566351 | 0.99999785 | -6.3635792 | down |
| KI - WT, cell_origin: iPSC | P81605 | DCD | 4 | Homo sapien | 0.13020326 | -1.1847356 | 1.44514208 | 1.81300489 | 0.21101761 | 0.83570813 | 0.99999785 | -6.3635919 | up |
| KI - WT, cell_origin: iPSC | P19387 | POLR2C | 5 | Homo sapien | 0.07548856 | -0.6872561 | 0.83823319 | 0.3928052 | 0.21081313 | 0.83584948 | 0.99999785 | -6.3636441 | up |
| KI - WT, cell_origin: iPSC | Q5TH69 | ARFGEF3 | 5 | Homo sapien | -0.094106 | -1.0479793 | 0.85976742 | -2.1770532 | -0.2101194 | 0.8363771 | 0.99999785 | -6.3637971 | down |
| KI - WT, cell_origin: iPSC | P28799 | GRN | 4 | Homo sapien | -0.125165 | -1.3951349 | 1.14480482 | -1.493528 | -0.2100351 | 0.83646112 | 0.99999785 | -6.3638057 | down |
| KI - WT, cell_origin: iPSC | Q9Y580 | RBM7 | 5 | Homo sapien | 0.08766539 | -0.8016191 | 0.97694987 | -0.8627908 | 0.20999325 | 0.83647957 | 0.99999785 | -6.3638216 | up |
| KI - WT, cell_origin: iPSC | P61160 | ACTR2 | 7 | Homo sapien | 0.05957909 | -0.5469138 | 0.66607202 | 1.0521433 | 0.20922224 | 0.83706491 | 0.99999785 | -6.3639915 | up |
| KI - WT, cell_origin: iPSC | P62249 | RPS16 | 13 | Homo sapien | -0.0387376 | -0.4331093 | 0.355634 | 4.86086573 | -0.2092027 | 0.8370799 | 0.99999785 | -6.3639957 | down |
| KI - WT, cell_origin: iPSC | P83881 | RPL36A | 7 | Homo sapien | -0.074487 | -0.8337073 | 0.68473333 | 2.90806144 | -0.2089923 | 0.83724695 | 0.99999785 | -6.3640383 | down |
| KI - WT, cell_origin: iPSC | O94903 | PLPBP | 9 | Homo sapien | 0.09442108 | -0.8686252 | 1.05746737 | 0.97355772 | 0.20894124 | 0.8372997 | 0.99999785 | -6.3640426 | up |
| KI - WT, cell_origin: iPSC | Q71RC2 | LARP4 | 7 | Homo sapien | -0.0990819 | -1.1098931 | 0.91172927 | -0.5707644 | -0.2088944 | 0.83733562 | 0.99999785 | -6.3640527 | down |
| KI - WT, cell_origin: iPSC | Q9Y2J4 | AMOTL2 | 3 | Homo sapien | 0.35071551 | -3.2315574 | 3.93298838 | -2.5602426 | 0.20864053 | 0.83753027 | 0.99999785 | -6.3641075 | up |
| KI - WT, cell_origin: iPSC | Q12874 | SF3A3 | 13 | Homo sapien | -0.111312 | -1.2485701 | 1.02594617 | 1.74269687 | -0.208586 | 0.83757207 | 0.99999785 | -6.3641193 | down |
| KI - WT, cell_origin: iPSC | Q9Y333 | LSM2 | 5 | Homo sapien | 0.05151559 | -0.4764511 | 0.57948231 | 1.13013011 | 0.20781256 | 0.83814595 | 0.99999785 | -6.3642952 | up |
| KI - WT, cell_origin: iPSC | Q7Z6J2 | TAMALIN | 8 | Homo sapien | -0.1484962 | -1.674484 | 1.37749156 | -0.0588158 | -0.20738 | 0.83849695 | 0.99999785 | -6.3643786 | down |
| KI - WT, cell_origin: iPSC | Q9Y4Z0 | LSM4 | 5 | Homo sapien | -0.0509203 | -0.573986 | 0.47214537 | 0.12297781 | -0.207336 | 0.8385115 | 0.99999785 | -6.3643975 | down |
| KI - WT, cell_origin: iPSC | P19525 | EIF2AK2 | 9 | Homo sapien | 0.04691556 | -0.4351837 | 0.52901479 | 0.25924995 | 0.20726222 | 0.83856807 | 0.99999785 | -6.3644133 | up |

| | | | | | | | | | | | | | |
|---|---|---|---|---|---|---|---|---|---|---|---|---|---|
| KI - WT, cell_origin: iPSC | Q14566 | MCM6 | 8 | Homo sapien | 0.11921792 | -1.1086001 | 1.34703595 | -2.0311118 | 0.20686874 | 0.83888067 | 0.99999785 | -6.3644922 | up |
| KI - WT, cell_origin: iPSC | Q9P2D0 | IBTK | 11 | Homo sapien | -0.0818501 | -0.9248292 | 0.7611291 | -0.0403096 | -0.2068331 | 0.83890294 | 0.99999785 | -6.3645023 | down |
| KI - WT, cell_origin: iPSC | Q07955 | SRSF1 | 13 | Homo sapien | -0.1313534 | -1.4876105 | 1.22490363 | 3.50038576 | -0.2063962 | 0.83925151 | 0.99999785 | -6.364589 | down |
| KI - WT, cell_origin: iPSC | Q8WVC0 | LEO1 | 5 | Homo sapien | 0.11740809 | -1.1002618 | 1.33507794 | -1.7886997 | 0.20546553 | 0.83996325 | 0.99999785 | -6.3647883 | up |
| KI - WT, cell_origin: iPSC | Q9Y570 | PPME1 | 6 | Homo sapien | -0.1101129 | -1.2546667 | 1.03444095 | -0.8405429 | -0.2050238 | 0.84030453 | 0.99999785 | -6.3648809 | down |
| KI - WT, cell_origin: iPSC | E9PRG8 | C11orf98 | 3 | Homo sapien | -0.0524201 | -0.5973795 | 0.49253941 | 1.52649933 | -0.2048674 | 0.84040555 | 0.99999785 | -6.3649233 | down |
| KI - WT, cell_origin: iPSC | P31350 | RRM2 | 5 | Homo sapien | 0.07894712 | -0.7421122 | 0.90000641 | -0.3269982 | 0.20482141 | 0.8404462 | 0.99999785 | -6.3649304 | up |
| KI - WT, cell_origin: iPSC | Q9ULE6 | PALD1 | 6 | Homo sapien | -0.1206137 | -1.376456 | 1.13522864 | -1.2577033 | -0.2046745 | 0.84057254 | 0.99999785 | -6.3649549 | down |
| KI - WT, cell_origin: iPSC | O75420 | GIGYF1 | 10 | Homo sapien | -0.1141711 | -1.3047373 | 1.07639524 | -1.2049832 | -0.2043642 | 0.8408107 | 0.99999785 | -6.3650205 | down |
| KI - WT, cell_origin: iPSC | Q13616 | CUL1 | 5 | Homo sapien | 0.09484462 | -0.893925 | 1.08361424 | -2.395351 | 0.20429478 | 0.84084508 | 0.99999785 | -6.3650443 | up |
| KI - WT, cell_origin: iPSC | Q9BZ23 | PANK2 | 4 | Homo sapien | 0.07805193 | -0.7358503 | 0.89195421 | -0.6440107 | 0.2042764 | 0.84086402 | 0.99999785 | -6.3650459 | up |
| KI - WT, cell_origin: iPSC | Q9Y282 | ERGIC3 | 7 | Homo sapien | -0.1654921 | -1.8937107 | 1.56272649 | -0.1135923 | -0.2040708 | 0.84103584 | 0.99999785 | -6.3650824 | down |
| KI - WT, cell_origin: iPSC | Q9BRZ2 | TRIM56 | 12 | Homo sapien | 0.06436994 | -0.6078082 | 0.73654809 | 0.60596243 | 0.20395688 | 0.84110445 | 0.99999785 | -6.3651156 | up |
| KI - WT, cell_origin: iPSC | Q9BZX2 | UCK2 | 7 | Homo sapien | 0.11910719 | -1.1285218 | 1.36673613 | -0.6832417 | 0.20344869 | 0.84151339 | 0.99999785 | -6.3652135 | up |
| KI - WT, cell_origin: iPSC | Q13554 | CAMK2B | 9 | Homo sapien | 0.04729004 | -0.4477937 | 0.54237381 | 1.35727357 | 0.20343736 | 0.84150327 | 0.99999785 | -6.365225 | up |
| KI - WT, cell_origin: iPSC | Q9NRY2 | INIP | 2 | Homo sapien | 0.11307631 | -1.0741229 | 1.30027554 | -2.1573644 | 0.20289248 | 0.84192721 | 0.99999785 | -6.3653367 | up |
| KI - WT, cell_origin: iPSC | O43293 | DAPK3 | 6 | Homo sapien | -0.1356762 | -1.5609116 | 1.28955923 | -0.9339625 | -0.2028707 | 0.84195708 | 0.99999785 | -6.3653349 | down |
| KI - WT, cell_origin: iPSC | Q93052 | LPP | 5 | Homo sapien | 0.08242228 | -0.7845771 | 0.94942168 | -0.5233127 | 0.20250871 | 0.84222187 | 0.99999785 | -6.3654171 | up |
| KI - WT, cell_origin: iPSC | P0DPB5 | POLR1D | 3 | Homo sapien | -0.2003144 | -2.3107011 | 1.91007235 | 0.94189664 | -0.2022798 | 0.84241079 | 0.99999785 | -6.3654587 | down |
| KI - WT, cell_origin: iPSC | Q8N6T7 | SIRT6 | 3 | Homo sapien | -0.1275585 | -1.4708793 | 1.21576236 | -3.3089768 | -0.202241 | 0.84242187 | 0.99999785 | -6.3654758 | down |
| KI - WT, cell_origin: iPSC | Q5TZA2 | CROCC | 6 | Homo sapien | 0.17960682 | -1.7183681 | 2.07758174 | -2.91431 | 0.20166695 | 0.84288136 | 0.99999785 | -6.3655867 | up |
| KI - WT, cell_origin: iPSC | Q9UJK0 | TSR3 | 2 | Homo sapien | 0.10232156 | -0.9785368 | 1.18317989 | -3.409696 | 0.20162206 | 0.8428972 | 0.99999785 | -6.365605 | up |
| KI - WT, cell_origin: iPSC | Q9UPN9 | TRIM33 | 13 | Homo sapien | -0.1346602 | -1.5635328 | 1.29421243 | -1.8857546 | -0.200839 | 0.84351723 | 0.99999785 | -6.365759 | down |
| KI - WT, cell_origin: iPSC | P34932 | HSPA4 | 21 | Homo sapien | -0.1067181 | -1.2392356 | 1.02579941 | -0.7640362 | -0.2008147 | 0.84353587 | 0.99999785 | -6.365764 | down |
| KI - WT, cell_origin: iPSC | Q9NPD3 | EXOSC4 | 7 | Homo sapien | 0.10180821 | -0.9790412 | 1.18265758 | -0.89167 | 0.20073356 | 0.8435982 | 0.99999785 | -6.3657809 | up |
| KI - WT, cell_origin: iPSC | O15021 | MAST4 | 11 | Homo sapien | -0.1359748 | -1.5829747 | 1.31102507 | 2.15269828 | -0.2002592 | 0.84396258 | 0.99999785 | -6.3658792 | down |
| KI - WT, cell_origin: iPSC | P04406 | GAPDH | 14 | Homo sapien | 0.12790365 | -1.2354279 | 1.49123514 | 2.57705828 | 0.1999327 | 0.84421337 | 0.99999785 | -6.3659468 | up |
| KI - WT, cell_origin: iPSC | Q9H9T3 | ELP3 | 8 | Homo sapien | 0.08247254 | -0.7975794 | 0.96252449 | -0.8862482 | 0.19962685 | 0.84443538 | 0.99999785 | -6.3660161 | up |
| KI - WT, cell_origin: iPSC | P62487 | POLR2G | 3 | Homo sapien | 0.10252126 | -0.994058 | 1.19910056 | -0.8552697 | 0.19923988 | 0.84474563 | 0.99999785 | -6.3660898 | up |
| KI - WT, cell_origin: iPSC | P28074 | PSMB5 | 4 | Homo sapien | -0.0922322 | -1.0791545 | 0.89469012 | 0.13620968 | -0.1991599 | 0.84480709 | 0.99999785 | -6.3661062 | down |
| KI - WT, cell_origin: iPSC | Q8NCM8 | DYNC2H1 | 7 | Homo sapien | -0.0682174 | -0.799086 | 0.66265125 | -2.5995047 | -0.1987904 | 0.84507266 | 0.99999785 | -6.366191 | down |
| KI - WT, cell_origin: iPSC | Q5VZE5 | NAA35 | 14 | Homo sapien | 0.11805164 | -1.1479653 | 1.38406854 | 0.11647465 | 0.19871694 | 0.84514744 | 0.99999785 | -6.3661974 | up |
| KI - WT, cell_origin: iPSC | Q9ULL5 | PRR12 | 2 | Homo sapien | -0.0853336 | -1.0005846 | 0.82991746 | -0.8853475 | -0.1986222 | 0.84520945 | 0.99999785 | -6.3662219 | down |
| KI - WT, cell_origin: iPSC | Q9H8Y1 | VRTN | 10 | Homo sapien | -0.0449766 | -0.5273418 | 0.43738858 | 0.40938905 | -0.1985869 | 0.84522902 | 0.99999785 | -6.3662328 | down |
| KI - WT, cell_origin: iPSC | Q16181 | SEPTIN7 | 12 | Homo sapien | 0.06761128 | -0.6580172 | 0.7932398 | -0.3718596 | 0.198447 | 0.84533654 | 0.99999785 | -6.3662615 | up |
| KI - WT, cell_origin: iPSC | P36405 | ARL3 | 3 | Homo sapien | 0.09643853 | -0.940857 | 1.13373405 | -0.7043395 | 0.1981301 | 0.84559839 | 0.99999785 | -6.3663178 | up |
| KI - WT, cell_origin: iPSC | Q05397 | PTK2 | 20 | Homo sapien | -0.0534024 | -0.6281887 | 0.5213839 | 0.65865765 | -0.1978764 | 0.84577513 | 0.99999785 | -6.3663784 | down |
| KI - WT, cell_origin: iPSC | Q9HD26 | GOPC | 13 | Homo sapien | 0.12492561 | -1.2214334 | 1.47128459 | -1.0795148 | 0.19773929 | 0.84589874 | 0.99999785 | -6.3663978 | up |
| KI - WT, cell_origin: iPSC | Q96PE3 | INPP4A | 4 | Homo sapien | -0.0772401 | -0.91077 | 0.75628974 | 1.25554103 | -0.1974103 | 0.84614087 | 0.99999785 | -6.36647 | down |
| KI - WT, cell_origin: iPSC | Q8TE02 | ELP5 | 2 | Homo sapien | 0.26953078 | -2.6404434 | 3.17950495 | -0.5008292 | 0.19738839 | 0.84616844 | 0.99999785 | -6.3664695 | up |
| KI - WT, cell_origin: iPSC | Q9HA64 | FN3KRP | 12 | Homo sapien | -0.0807593 | -0.9538373 | 0.79231874 | 3.21189271 | -0.197125 | 0.8463709 | 0.99999785 | -6.3665232 | down |
| KI - WT, cell_origin: iPSC | Q02878 | RPL6 | 17 | Homo sapien | -0.0469339 | -0.5540437 | 0.46017588 | 3.39165671 | -0.1971171 | 0.84635879 | 0.99999785 | -6.3665334 | down |
| KI - WT, cell_origin: iPSC | O15145 | ARPC3 | 9 | Homo sapien | 0.04620325 | -0.4535697 | 0.54597622 | 0.51006637 | 0.19689714 | 0.84652785 | 0.99999785 | -6.3665782 | up |
| KI - WT, cell_origin: iPSC | Q7LG56 | RRM2B | 7 | Homo sapien | 0.07254429 | -0.712669 | 0.85775756 | -0.5919174 | 0.19678832 | 0.84661456 | 0.99999785 | -6.3665989 | up |
| KI - WT, cell_origin: iPSC | Q92796 | DLG3 | 6 | Homo sapien | 0.09314611 | -0.9163606 | 1.10265286 | -1.4180252 | 0.19655017 | 0.84680002 | 0.99999785 | -6.3666462 | up |
| KI - WT, cell_origin: iPSC | Q9BZE4 | GTPBP4 | 24 | Homo sapien | 0.06291119 | -0.6189225 | 0.74474485 | -0.6485657 | 0.19651203 | 0.84682394 | 0.99999785 | -6.3666565 | up |

| | | | | | | | | | | | | | |
|---|---|---|---|---|---|---|---|---|---|---|---|---|---|
| KI - WT, cell_origin: iPSC | Q9Y6E0 | STK24 | 6 | Homo sapien | 0.06836306 | -0.6727185 | 0.80944461 | -0.7869306 | 0.19646951 | 0.84685663 | 0.99999785 | -6.3666651 | up |
| KI - WT, cell_origin: iPSC | P62136 | PPP1CA | 14 | Homo sapien | 0.06449025 | -0.6349495 | 0.76393003 | 1.33731659 | 0.19637375 | 0.84693025 | 0.99999785 | -6.3666845 | up |
| KI - WT, cell_origin: iPSC | Q13185 | CBX3 | 6 | Homo sapien | -0.1196085 | -1.4182144 | 1.17899738 | -0.3915498 | -0.196285 | 0.84701663 | 0.99999785 | -6.3666941 | down |
| KI - WT, cell_origin: iPSC | Q14008 | CKAP5 | 46 | Homo sapien | 0.06745575 | -0.6645513 | 0.79946278 | 0.24433606 | 0.19627482 | 0.84700779 | 0.99999785 | -6.3667039 | up |
| KI - WT, cell_origin: iPSC | Q14562 | DHX8 | 16 | Homo sapien | -0.1221493 | -1.4503476 | 1.20604895 | -1.1469753 | -0.1959885 | 0.84724461 | 0.99999785 | -6.3667542 | down |
| KI - WT, cell_origin: iPSC | Q9UHD8 | SEPTIN9 | 11 | Homo sapien | 0.08985086 | -0.8888852 | 1.06858696 | -0.4259767 | 0.19564061 | 0.84751205 | 0.99999785 | -6.3668247 | up |
| KI - WT, cell_origin: iPSC | P61923 | COPZ1 | 5 | Homo sapien | 0.11314938 | -1.1196087 | 1.34590743 | 1.16160593 | 0.19560353 | 0.84754056 | 0.99999785 | -6.3668322 | up |
| KI - WT, cell_origin: iPSC | P0DPB6 | POLR1D | 3 | Homo sapien | 0.09395769 | -0.929759 | 1.11767435 | -1.7668478 | 0.19550568 | 0.84760237 | 0.99999785 | -6.3668582 | up |
| KI - WT, cell_origin: iPSC | Q6PJG6 | BRAT1 | 14 | Homo sapien | -0.0841267 | -1.0023917 | 0.83413835 | -1.1825153 | -0.1951567 | 0.8478715 | 0.99999785 | -6.3669284 | down |
| KI - WT, cell_origin: iPSC | Q7L2H7 | EIF3M | 18 | Homo sapien | -0.0419306 | -0.499807 | 0.41594579 | 2.58954565 | -0.1950396 | 0.84795621 | 0.99999785 | -6.3669545 | down |
| KI - WT, cell_origin: iPSC | Q9P2J5 | LARS1 | 20 | Homo sapien | 0.06401086 | -0.636088 | 0.76410969 | 0.60041337 | 0.1947305 | 0.84819393 | 0.99999785 | -6.3670167 | up |
| KI - WT, cell_origin: iPSC | P35611 | ADD1 | 12 | Homo sapien | -0.1696934 | -2.027123 | 1.68773616 | -0.6854835 | -0.1946951 | 0.84823913 | 0.99999785 | -6.3670155 | down |
| KI - WT, cell_origin: iPSC | P63244 | RACK1 | 18 | Homo sapien | 0.06217327 | -0.6197115 | 0.74405803 | 3.7031683 | 0.19422249 | 0.84858926 | 0.99999785 | -6.3671168 | up |
| KI - WT, cell_origin: iPSC | Q8IY67 | RAVER1 | 8 | Homo sapien | -0.1083766 | -1.2996658 | 1.08291271 | -0.4497442 | -0.1938744 | 0.84887032 | 0.99999785 | -6.3671804 | down |
| KI - WT, cell_origin: iPSC | P42345 | MTOR | 48 | Homo sapien | 0.05071264 | -0.507355 | 0.6087803 | 1.24802804 | 0.19353926 | 0.84911027 | 0.99999785 | -6.3672558 | up |
| KI - WT, cell_origin: iPSC | P05388 | RPLP0 | 14 | Homo sapien | -0.0397303 | -0.4775976 | 0.39813693 | 5.32971668 | -0.1932499 | 0.84933286 | 0.99999785 | -6.3673137 | down |
| KI - WT, cell_origin: iPSC | P52597 | HNRNPF | 10 | Homo sapien | -0.0779237 | -0.9381222 | 0.78227482 | 2.32804395 | -0.1930136 | 0.84952675 | 0.99999785 | -6.3673553 | down |
| KI - WT, cell_origin: iPSC | P36404 | ARL2 | 2 | Homo sapien | 0.09816956 | -0.9865086 | 1.1828477 | -2.4777977 | 0.19275943 | 0.84971026 | 0.99999785 | -6.3674115 | up |
| KI - WT, cell_origin: iPSC | Q9H4H8 | FAM83D | 4 | Homo sapien | 0.18936276 | -1.913265 | 2.29199056 | -2.200672 | 0.19192632 | 0.85036905 | 0.99999785 | -6.3675691 | up |
| KI - WT, cell_origin: iPSC | P07384 | CAPN1 | 28 | Homo sapien | 0.18296904 | -1.8512 | 2.21713807 | 1.62790124 | 0.19168712 | 0.85055312 | 0.99999785 | -6.3676165 | up |
| KI - WT, cell_origin: iPSC | Q9NX55 | HYPK | 3 | Homo sapien | 0.05286136 | -0.5351758 | 0.64089855 | -0.5415388 | 0.19145785 | 0.85071189 | 0.99999785 | -6.36767 | up |
| KI - WT, cell_origin: iPSC | P18124 | RPL7 | 16 | Homo sapien | -0.0387683 | -0.4703089 | 0.39277238 | 4.41223223 | -0.1913349 | 0.8508065 | 0.99999785 | -6.3676944 | down |
| KI - WT, cell_origin: iPSC | Q9UK41 | VPS28 | 13 | Homo sapien | 0.05880894 | -0.5965768 | 0.71419468 | 1.4043565 | 0.19111116 | 0.85097873 | 0.99999785 | -6.3677386 | up |
| KI - WT, cell_origin: iPSC | Q15014 | MORF4L2 | 5 | Homo sapien | -0.1111367 | -1.3526623 | 1.13038877 | -0.9680507 | -0.1907675 | 0.85126086 | 0.99999785 | -6.3677984 | down |
| KI - WT, cell_origin: iPSC | Q9H0S4 | DDX47 | 7 | Homo sapien | -0.1513704 | -1.8449751 | 1.54223424 | -0.9066902 | -0.1904721 | 0.85148825 | 0.99999785 | -6.3678567 | down |
| KI - WT, cell_origin: iPSC | Q15149 | PLEC | 264 | Homo sapien | -0.0713991 | -0.8708374 | 0.7280393 | 2.68004992 | -0.1902504 | 0.85164658 | 0.99999785 | -6.3679059 | down |
| KI - WT, cell_origin: iPSC | O95070 | YIF1A | 2 | Homo sapien | -0.2401297 | -2.9311029 | 2.45084351 | -0.3504986 | -0.1901686 | 0.85172183 | 0.99999785 | -6.3679164 | down |
| KI - WT, cell_origin: iPSC | Q8IXT5 | RBM12B | 21 | Homo sapien | 0.06364054 | -0.6502653 | 0.7775464 | 0.26275577 | 0.1898596 | 0.85194219 | 0.99999785 | -6.3679851 | up |
| KI - WT, cell_origin: iPSC | O76094 | SRP72 | 18 | Homo sapien | 0.08672208 | -0.8880932 | 1.06153734 | 1.0612843 | 0.18958752 | 0.85216916 | 0.99999785 | -6.3680305 | up |
| KI - WT, cell_origin: iPSC | O94966 | USP19 | 5 | Homo sapien | -0.0644815 | -0.7900846 | 0.66112162 | -1.4536208 | -0.1892673 | 0.85239823 | 0.99999785 | -6.3681012 | down |
| KI - WT, cell_origin: iPSC | P13639 | EEF2 | 35 | Homo sapien | -0.0650682 | -0.7973759 | 0.66723956 | 2.01114047 | -0.1892559 | 0.8524093 | 0.99999785 | -6.3681024 | down |
| KI - WT, cell_origin: iPSC | Q6PGP7 | TTC37 | 21 | Homo sapien | 0.08308969 | -0.8541734 | 1.02035274 | 0.01697668 | 0.18891051 | 0.85268827 | 0.99999785 | -6.368164 | up |
| KI - WT, cell_origin: iPSC | Q96E39 | RBMXL1 | 3 | Homo sapien | -0.0696817 | -0.8556283 | 0.71626478 | -0.1471664 | -0.188852 | 0.85272172 | 0.99999785 | -6.3681807 | down |
| KI - WT, cell_origin: iPSC | Q9P2N5 | RBM27 | 16 | Homo sapien | -0.1150534 | -1.4154919 | 1.18538506 | -0.1650461 | -0.1885437 | 0.8529728 | 0.99999785 | -6.3682347 | down |
| KI - WT, cell_origin: iPSC | Q96RU2 | USP28 | 6 | Homo sapien | 0.12470683 | -1.2849631 | 1.53437671 | -1.838485 | 0.18852771 | 0.85298514 | 0.99999785 | -6.3682378 | up |
| KI - WT, cell_origin: iPSC | Q9Y388 | RBMX2 | 2 | Homo sapien | -0.1780641 | -2.1920985 | 1.83597022 | 0.72011098 | -0.1884135 | 0.8530731 | 0.99999785 | -6.3682601 | down |
| KI - WT, cell_origin: iPSC | P61077 | UBE2D3 | 4 | Homo sapien | -0.1646678 | -2.0305921 | 1.70125664 | 1.88006613 | -0.1880689 | 0.85333848 | 0.99999785 | -6.3683272 | down |
| KI - WT, cell_origin: iPSC | P07237 | P4HB | 13 | Homo sapien | 0.20166925 | -2.0845797 | 2.48791815 | -1.9394857 | 0.18798298 | 0.85340461 | 0.99999785 | -6.3683439 | up |
| KI - WT, cell_origin: iPSC | Q96JC1 | VPS39 | 9 | Homo sapien | 0.05557368 | -0.5743367 | 0.6854841 | -0.2092545 | 0.1879014 | 0.85345011 | 0.99999785 | -6.3683675 | up |
| KI - WT, cell_origin: iPSC | O43290 | SART1 | 24 | Homo sapien | 0.06747536 | -0.6984082 | 0.83335895 | 0.8878391 | 0.18766757 | 0.85363464 | 0.99999785 | -6.3684109 | up |
| KI - WT, cell_origin: iPSC | Q96SB8 | SMC6 | 5 | Homo sapien | 0.1157902 | -1.2013145 | 1.43289489 | -2.1497034 | 0.18732274 | 0.85390893 | 0.99999785 | -6.3684739 | up |
| KI - WT, cell_origin: iPSC | O43175 | PHGDH | 23 | Homo sapien | -0.0821716 | -1.0171544 | 0.85281127 | 1.78067414 | -0.1872925 | 0.85393638 | 0.99999785 | -6.3684779 | down |
| KI - WT, cell_origin: iPSC | O14579 | COPE | 18 | Homo sapien | 0.0766662 | -0.796554 | 0.94988635 | 0.87048399 | 0.1870735 | 0.85410046 | 0.99999785 | -6.3685224 | up |
| KI - WT, cell_origin: iPSC | P02647 | APOA1 | 4 | Homo sapien | 0.19160315 | -1.9911159 | 2.37432217 | -3.0246562 | 0.1870713 | 0.85410676 | 0.99999785 | -6.3685207 | up |
| KI - WT, cell_origin: iPSC | O95453 | PARN | 12 | Homo sapien | 0.14844656 | -1.5446497 | 1.8415428 | -1.5938789 | 0.18684904 | 0.85427796 | 0.99999785 | -6.3685637 | up |
| KI - WT, cell_origin: iPSC | P62899 | RPL31 | 6 | Homo sapien | 0.04480151 | -0.466759 | 0.556362 | 5.57655629 | 0.18652425 | 0.85451095 | 0.99999785 | -6.3686341 | up |

| | | | | | | | | | | | | | |
|---|---|---|---|---|---|---|---|---|---|---|---|---|---|
| KI - WT, cell_origin: iPSC | O43447 | PPIH | 4 | Homo sapien | 0.26070979 | -2.7192316 | 3.24065116 | 0.2123097 | 0.18644554 | 0.85458878 | 0.99999785 | -6.3686416 | up |
| KI - WT, cell_origin: iPSC | Q96QC0 | PPP1R10 | 15 | Homo sapien | 0.07634594 | -0.7984009 | 0.9510928 | 0.09292843 | 0.18594565 | 0.85496606 | 0.99999785 | -6.3687414 | up |
| KI - WT, cell_origin: iPSC | Q68E01 | INTS3 | 10 | Homo sapien | 0.05510696 | -0.5763915 | 0.68660543 | -1.3462286 | 0.18585481 | 0.85502673 | 0.99999785 | -6.368763 | up |
| KI - WT, cell_origin: iPSC | Q9UNP9 | PPIE | 4 | Homo sapien | 0.06382482 | -0.6681374 | 0.795787 | -1.2265537 | 0.18571228 | 0.85513656 | 0.99999785 | -6.3687904 | up |
| KI - WT, cell_origin: iPSC | Q9H4B6 | SAV1 | 4 | Homo sapien | 0.11650986 | -1.2208044 | 1.45382416 | -1.6330265 | 0.18566565 | 0.8551896 | 0.99999785 | -6.3687917 | up |
| KI - WT, cell_origin: iPSC | P59998 | ARPC4 | 5 | Homo sapien | -0.0630162 | -0.7874569 | 0.66142441 | 0.51422202 | -0.1852751 | 0.85547524 | 0.99999785 | -6.3688734 | down |
| KI - WT, cell_origin: iPSC | Q8WZA9 | IRGQ | 3 | Homo sapien | 0.18180356 | -1.9115903 | 2.27519742 | -1.4665998 | 0.18507758 | 0.85564271 | 0.99999785 | -6.3689045 | up |
| KI - WT, cell_origin: iPSC | Q99880 | H2BC13 | 7 | Homo sapien | -0.086162 | -1.0784071 | 0.90608319 | 2.47544709 | -0.1850542 | 0.85566073 | 0.99999785 | -6.368909 | down |
| KI - WT, cell_origin: iPSC | O94808 | GFPT2 | 20 | Homo sapien | -0.1474891 | -1.8461466 | 1.55116842 | 0.45444647 | -0.1850361 | 0.85567469 | 0.99999785 | -6.3689124 | down |
| KI - WT, cell_origin: iPSC | Q13153 | PAK1 | 15 | Homo sapien | 0.1571086 | -1.659735 | 1.97395217 | 0.37213222 | 0.1842828 | 0.85625517 | 0.99999785 | -6.3690563 | up |
| KI - WT, cell_origin: iPSC | Q15019 | SEPTIN2 | 12 | Homo sapien | 0.0551898 | -0.583012 | 0.69339163 | 0.38114576 | 0.18417914 | 0.85631809 | 0.99999785 | -6.3690836 | up |
| KI - WT, cell_origin: iPSC | Q9Y5Q9 | GTF3C3 | 9 | Homo sapien | 0.14012587 | -1.4831081 | 1.76335989 | -2.0224029 | 0.18396684 | 0.85649867 | 0.99999785 | -6.3691165 | up |
| KI - WT, cell_origin: iPSC | Q13242 | SRSF9 | 13 | Homo sapien | -0.146803 | -1.8477073 | 1.55410134 | 1.96250471 | -0.183932 | 0.8565255 | 0.99999785 | -6.3691231 | down |
| KI - WT, cell_origin: iPSC | Q96KC2 | ARL5B | 2 | Homo sapien | 0.065333 | -0.6915055 | 0.82217149 | 0.57054326 | 0.18387602 | 0.85655534 | 0.99999785 | -6.3691397 | up |
| KI - WT, cell_origin: iPSC | O95218 | ZRANB2 | 13 | Homo sapien | 0.13417974 | -1.4297109 | 1.69807033 | 0.46179813 | 0.18284494 | 0.85736343 | 0.99999785 | -6.3693294 | up |
| KI - WT, cell_origin: iPSC | O60488 | ACSL4 | 20 | Homo sapien | -0.1536121 | -1.9448346 | 1.63761048 | 1.596559 | -0.1827587 | 0.85742989 | 0.99999785 | -6.3693457 | down |
| KI - WT, cell_origin: iPSC | Q2M389 | WASHC4 | 7 | Homo sapien | 0.06661133 | -0.7099549 | 0.84317758 | -1.2212848 | 0.1826877 | 0.85746782 | 0.99999785 | -6.3693665 | up |
| KI - WT, cell_origin: iPSC | Q2VPK5 | CTU2 | 6 | Homo sapien | 0.0620668 | -0.6622712 | 0.7864048 | -0.2397833 | 0.18249784 | 0.8576142 | 0.99999785 | -6.3694023 | up |
| KI - WT, cell_origin: iPSC | Q9NVR2 | INTS10 | 3 | Homo sapien | 0.09396376 | -1.0078695 | 1.19579704 | -1.0636712 | 0.18173846 | 0.85821648 | 0.99999785 | -6.369538 | up |
| KI - WT, cell_origin: iPSC | Q15185 | PTGES3 | 4 | Homo sapien | -0.0974666 | -1.2406977 | 1.04576458 | 0.35400962 | -0.181687 | 0.85825614 | 0.99999785 | -6.3695477 | down |
| KI - WT, cell_origin: iPSC | P68366 | TUBA4A | 5 | Homo sapien | 0.08022299 | -0.8643645 | 1.02481049 | 5.06152243 | 0.18099185 | 0.8587922 | 0.99999785 | -6.3696781 | up |
| KI - WT, cell_origin: iPSC | Q9UPP1 | PHF8 | 3 | Homo sapien | 0.09182081 | -0.9889307 | 1.1725723 | -2.2939409 | 0.18094848 | 0.85880897 | 0.99999785 | -6.3696935 | up |
| KI - WT, cell_origin: iPSC | Q9BTA9 | WAC | 5 | Homo sapien | -0.10522 | -1.3465521 | 1.13611214 | -2.5324988 | -0.1805552 | 0.85911609 | 0.99999785 | -6.3697653 | down |
| KI - WT, cell_origin: iPSC | Q01844 | EWSR1 | 6 | Homo sapien | -0.0659549 | -0.8446867 | 0.71277681 | 5.16882516 | -0.1804936 | 0.85917644 | 0.99999785 | -6.3697713 | down |
| KI - WT, cell_origin: iPSC | P63167 | DYNLL1 | 4 | Homo sapien | -0.0443653 | -0.5684482 | 0.47971754 | 2.76394658 | -0.1802948 | 0.85931316 | 0.99999785 | -6.3698156 | down |
| KI - WT, cell_origin: iPSC | O15084 | ANKRD28 | 2 | Homo sapien | 0.08999059 | -0.9745413 | 1.15452244 | -3.3036647 | 0.18004379 | 0.85950679 | 0.99999785 | -6.3698624 | up |
| KI - WT, cell_origin: iPSC | P30876 | POLR2B | 26 | Homo sapien | 0.05524468 | -0.6000374 | 0.71052679 | 0.47703633 | 0.17955681 | 0.85988246 | 0.99999785 | -6.3699529 | up |
| KI - WT, cell_origin: iPSC | P51825 | AFF1 | 6 | Homo sapien | 0.12605602 | -1.3711341 | 1.62324613 | -1.7652026 | 0.17942751 | 0.85999873 | 0.99999785 | -6.3699698 | up |
| KI - WT, cell_origin: iPSC | Q06455 | RUNX1T1 | 2 | Homo sapien | -0.1750721 | -2.2549601 | 1.90481591 | -4.4372714 | -0.1793014 | 0.86008369 | 0.99999785 | -6.3699985 | down |
| KI - WT, cell_origin: iPSC | P17987 | TCP1 | 16 | Homo sapien | 0.06696036 | -0.7287453 | 0.86266603 | 0.80835147 | 0.17925994 | 0.8601164 | 0.99999785 | -6.3700059 | up |
| KI - WT, cell_origin: iPSC | Q9NXC5 | MIOS | 35 | Homo sapien | -0.0426962 | -0.5509164 | 0.4655241 | 1.22128505 | -0.1789272 | 0.86036819 | 0.99999785 | -6.3700696 | down |
| KI - WT, cell_origin: iPSC | Q00534 | CDK6 | 9 | Homo sapien | -0.0731372 | -0.9471099 | 0.80083548 | -1.0929509 | -0.1782573 | 0.86088932 | 0.99999785 | -6.3701916 | down |
| KI - WT, cell_origin: iPSC | Q6PCB5 | RSBN1L | 5 | Homo sapien | -0.1169571 | -1.5159734 | 1.28205918 | -1.1648839 | -0.1781583 | 0.86097787 | 0.99999785 | -6.3702046 | down |
| KI - WT, cell_origin: iPSC | Q9HCS7 | XAB2 | 5 | Homo sapien | 0.09545781 | -1.0478219 | 1.23873755 | -2.1126088 | 0.17785765 | 0.86119812 | 0.99999785 | -6.370265 | up |
| KI - WT, cell_origin: iPSC | O14787 | TNPO2 | 9 | Homo sapien | -0.044621 | -0.5802012 | 0.49095918 | 0.54790125 | -0.1774412 | 0.86151491 | 0.99999785 | -6.3703435 | down |
| KI - WT, cell_origin: iPSC | P36954 | POLR2I | 3 | Homo sapien | 0.05678841 | -0.6248567 | 0.73843348 | -0.8851421 | 0.17743573 | 0.86151912 | 0.99999785 | -6.3703445 | up |
| KI - WT, cell_origin: iPSC | Q6ZWJ1 | STXBP4 | 3 | Homo sapien | -0.2271552 | -2.9604303 | 2.50611991 | -2.9621921 | -0.1771094 | 0.86178725 | 0.99999785 | -6.3703973 | down |
| KI - WT, cell_origin: iPSC | P40938 | RFC3 | 11 | Homo sapien | -0.1100493 | -1.4359221 | 1.21582353 | -0.0256507 | -0.1768837 | 0.86196148 | 0.99999785 | -6.3704387 | down |
| KI - WT, cell_origin: iPSC | Q9C0B1 | FTO | 10 | Homo sapien | -0.1113894 | -1.4534341 | 1.23065518 | -1.3988902 | -0.1768803 | 0.86196409 | 0.99999785 | -6.3704393 | down |
| KI - WT, cell_origin: iPSC | P49368 | CCT3 | 22 | Homo sapien | 0.0578784 | -0.6407475 | 0.75650426 | 1.65565904 | 0.17644589 | 0.86228313 | 0.99999785 | -6.3705256 | up |
| KI - WT, cell_origin: iPSC | Q7L190 | DPPA4 | 9 | Homo sapien | -0.0881758 | -1.1547706 | 0.97841909 | 1.62368702 | -0.1761782 | 0.86250599 | 0.99999785 | -6.3705676 | down |
| KI - WT, cell_origin: iPSC | P62826 | RAN | 6 | Homo sapien | -0.0803331 | -1.0523349 | 0.89166876 | 0.70415047 | -0.1761285 | 0.86254436 | 0.99999785 | -6.3705766 | down |
| KI - WT, cell_origin: iPSC | Q14690 | PDCD11 | 17 | Homo sapien | 0.03911631 | -0.433915 | 0.51214758 | -0.8017363 | 0.17611962 | 0.86253499 | 0.99999785 | -6.3705851 | up |
| KI - WT, cell_origin: iPSC | Q8N1F7 | NUP93 | 11 | Homo sapien | -0.0674504 | -0.8844918 | 0.7495911 | -0.55829 | -0.1758519 | 0.86274582 | 0.99999785 | -6.3706321 | down |
| KI - WT, cell_origin: iPSC | O75475 | PSIP1 | 17 | Homo sapien | -0.1167495 | -1.5400336 | 1.30653453 | -0.5893808 | -0.1748099 | 0.86356229 | 0.99999785 | -6.370816 | down |
| KI - WT, cell_origin: iPSC | P02042;P688 | HBD;HBB | 2 | Homo sapien | -0.1586148 | -2.0957474 | 1.77851775 | 0.09957422 | -0.1744965 | 0.86380419 | 0.99999785 | -6.3708727 | down |

| | | | | | | | | | | | | | |
|---|---|---|---|---|---|---|---|---|---|---|---|---|---|
| KI - WT, cell_origin: iPSC | P21796 | VDAC1 | 5 | Homo sapien | -0.2834366 | -3.7470064 | 3.18013322 | -2.8976165 | -0.1743951 | 0.8638825 | 0.99999785 | -6.370891 | down |
| KI - WT, cell_origin: iPSC | P29084 | GTF2E2 | 3 | Homo sapien | -0.1189549 | -1.5730769 | 1.33516714 | -2.6772335 | -0.1742866 | 0.86395901 | 0.99999785 | -6.3709136 | down |
| KI - WT, cell_origin: iPSC | P62304 | SNRPE | 4 | Homo sapien | -0.1204004 | -1.5932421 | 1.3524414 | 3.26510114 | -0.1742104 | 0.86402512 | 0.99999785 | -6.3709243 | down |
| KI - WT, cell_origin: iPSC | Q6ZRY4 | RBPMS2 | 10 | Homo sapien | -0.1649068 | -2.1841328 | 1.85431917 | -0.6322651 | -0.1740428 | 0.86415456 | 0.99999785 | -6.3709545 | down |
| KI - WT, cell_origin: iPSC | Q13155 | AIMP2 | 9 | Homo sapien | 0.05648449 | -0.634953 | 0.747922 | 0.24760947 | 0.17398667 | 0.86418188 | 0.99999785 | -6.3709713 | up |
| KI - WT, cell_origin: iPSC | P20700 | LMNB1 | 44 | Homo sapien | 0.12646136 | -1.4233607 | 1.67628339 | 2.76422018 | 0.17389152 | 0.86427136 | 0.99999785 | -6.3709817 | up |
| KI - WT, cell_origin: iPSC | O95456 | PSMG1 | 4 | Homo sapien | -0.1010424 | -1.3407603 | 1.1386755 | -3.250561 | -0.1735883 | 0.86448955 | 0.99999785 | -6.3710429 | down |
| KI - WT, cell_origin: iPSC | P48730 | CSNK1D | 10 | Homo sapien | -0.0419883 | -0.5583574 | 0.47438083 | 0.49175221 | -0.1731839 | 0.8648019 | 0.99999785 | -6.3711154 | down |
| KI - WT, cell_origin: iPSC | P50395 | GDI2 | 13 | Homo sapien | -0.0693601 | -0.922811 | 0.78409086 | 0.44806489 | -0.1731458 | 0.86483985 | 0.99999785 | -6.3711187 | down |
| KI - WT, cell_origin: iPSC | Q9H583 | HEATR1 | 21 | Homo sapien | 0.13642453 | -1.5500176 | 1.82286665 | -2.0639353 | 0.1723945 | 0.86542751 | 0.99999785 | -6.3712499 | up |
| KI - WT, cell_origin: iPSC | P29279 | CCN2 | 8 | Homo sapien | -0.1313135 | -1.7547514 | 1.49212429 | -0.9783513 | -0.1723758 | 0.86544197 | 0.99999785 | -6.3712533 | down |
| KI - WT, cell_origin: iPSC | Q9NUQ6 | SPATS2L | 17 | Homo sapien | 0.05650929 | -0.6432257 | 0.75624429 | 0.5364512 | 0.171999 | 0.86571719 | 0.99999785 | -6.3713269 | up |
| KI - WT, cell_origin: iPSC | P19388 | POLR2E | 6 | Homo sapien | 0.05720417 | -0.651336 | 0.76574432 | 0.08939886 | 0.17195029 | 0.86575482 | 0.99999785 | -6.3713356 | up |
| KI - WT, cell_origin: iPSC | Q9NS86 | LANCL2 | 8 | Homo sapien | -0.0585203 | -0.7836898 | 0.66664912 | 1.29545471 | -0.1718847 | 0.8658073 | 0.99999785 | -6.3713465 | down |
| KI - WT, cell_origin: iPSC | Q08170 | SRSF4 | 15 | Homo sapien | 0.08885707 | -1.0132389 | 1.190953 | 1.45058409 | 0.1718205 | 0.86587089 | 0.99999785 | -6.3713521 | up |
| KI - WT, cell_origin: iPSC | Q9Y3U8 | RPL36 | 3 | Homo sapien | -0.0512517 | -0.6867039 | 0.58420043 | 3.00274601 | -0.1717771 | 0.86588859 | 0.99999785 | -6.3713664 | down |
| KI - WT, cell_origin: iPSC | Q96SB4 | SRPK1 | 10 | Homo sapien | 0.04129593 | -0.4732087 | 0.55580054 | 0.60827766 | 0.17094548 | 0.86653117 | 0.99999785 | -6.3715138 | up |
| KI - WT, cell_origin: iPSC | Q9BWH6 | RPAP1 | 11 | Homo sapien | 0.07120754 | -0.8171967 | 0.95961177 | -1.2423648 | 0.17073046 | 0.86670066 | 0.99999785 | -6.3715504 | up |
| KI - WT, cell_origin: iPSC | P08670 | VIM | 35 | Homo sapien | -0.1944054 | -2.6248822 | 2.23607131 | 2.08102116 | -0.1704587 | 0.86692296 | 0.99999785 | -6.3715933 | down |
| KI - WT, cell_origin: iPSC | Q56P03 | EAPP | 2 | Homo sapien | -0.0624514 | -0.8458151 | 0.72091234 | -1.0117443 | -0.1697925 | 0.8674222 | 0.99999785 | -6.371717 | down |
| KI - WT, cell_origin: iPSC | Q15032 | R3HDM1 | 9 | Homo sapien | 0.10365965 | -1.1974065 | 1.40472576 | -1.3681103 | 0.16979025 | 0.86743952 | 0.99999785 | -6.371711 | up |
| KI - WT, cell_origin: iPSC | Q96GX9 | APIP | 4 | Homo sapien | 0.1022108 | -1.1809127 | 1.38533429 | -1.1928506 | 0.16975817 | 0.86746431 | 0.99999785 | -6.3717166 | up |
| KI - WT, cell_origin: iPSC | Q13045 | FLII | 9 | Homo sapien | -0.1107901 | -1.5022794 | 1.28069916 | -1.5902196 | -0.1696772 | 0.86752687 | 0.99999785 | -6.3717309 | down |
| KI - WT, cell_origin: iPSC | Q6P1Q9 | METTL2B | 8 | Homo sapien | -0.0498453 | -0.6780774 | 0.57838674 | -0.6023285 | -0.1689834 | 0.86804756 | 0.99999785 | -6.3718588 | down |
| KI - WT, cell_origin: iPSC | O14653 | GOSR2 | 7 | Homo sapien | 0.28084115 | -3.2685481 | 3.83023044 | -1.5105142 | 0.16862015 | 0.86834383 | 0.99999785 | -6.3719159 | up |
| KI - WT, cell_origin: iPSC | Q96QZ7 | MAGI1 | 28 | Homo sapien | 0.07736093 | -0.900501 | 1.05522281 | 0.2004754 | 0.16859571 | 0.86836273 | 0.99999785 | -6.3719202 | up |
| KI - WT, cell_origin: iPSC | P83731 | RPL24 | 8 | Homo sapien | -0.0372921 | -0.5126933 | 0.43810914 | 4.00360383 | -0.167069 | 0.86952759 | 0.99999785 | -6.3721915 | down |
| KI - WT, cell_origin: iPSC | P30154 | PPP2R1B | 6 | Homo sapien | -0.0705332 | -0.9700472 | 0.82898077 | -0.9693616 | -0.1670335 | 0.86955961 | 0.99999785 | -6.3721958 | down |
| KI - WT, cell_origin: iPSC | Q15648 | MED1 | 15 | Homo sapien | -0.0720721 | -0.9919185 | 0.84777426 | -0.3510017 | -0.1669491 | 0.86963153 | 0.99999785 | -6.3722077 | down |
| KI - WT, cell_origin: iPSC | P13637 | ATP1A3 | 6 | Homo sapien | -0.206158 | -2.8469304 | 2.43461446 | -1.2061739 | -0.1663687 | 0.87008442 | 0.99999785 | -6.3723062 | down |
| KI - WT, cell_origin: iPSC | O15344 | MID1 | 7 | Homo sapien | 0.09087082 | -1.0781165 | 1.2598581 | -1.8132997 | 0.16562718 | 0.87065287 | 0.99999785 | -6.3724355 | up |
| KI - WT, cell_origin: iPSC | Q86UK7 | ZNF598 | 23 | Homo sapien | 0.10262137 | -1.2203846 | 1.42562736 | 1.56398521 | 0.16530209 | 0.87090923 | 0.99999785 | -6.3724892 | up |
| KI - WT, cell_origin: iPSC | Q14CB8 | ARHGAP19 | 2 | Homo sapien | 0.22133553 | -2.6380123 | 3.08068335 | 2.37457356 | 0.16496299 | 0.8711715 | 0.99999785 | -6.3725472 | up |
| KI - WT, cell_origin: iPSC | Q9C0C2 | TNKS1BP1 | 53 | Homo sapien | 0.05886346 | -0.704083 | 0.82180994 | 1.24544558 | 0.16434577 | 0.87163767 | 0.99999785 | -6.3726568 | up |
| KI - WT, cell_origin: iPSC | O96019 | ACTL6A | 9 | Homo sapien | 0.05603463 | -0.6714631 | 0.78353237 | -0.2664816 | 0.16404557 | 0.87186605 | 0.99999785 | -6.3727093 | up |
| KI - WT, cell_origin: iPSC | Q9Y2S6 | TMA7 | 2 | Homo sapien | -0.1788218 | -2.5049092 | 2.14726558 | 2.70262442 | -0.1638313 | 0.87204688 | 0.99999785 | -6.3727397 | down |
| KI - WT, cell_origin: iPSC | P01040 | CSTA | 3 | Homo sapien | -0.0819735 | -1.1515305 | 0.98758355 | 0.25199352 | -0.1633322 | 0.87243301 | 0.99999785 | -6.3728242 | down |
| KI - WT, cell_origin: iPSC | P33993 | MCM7 | 20 | Homo sapien | -0.1120844 | -1.574782 | 1.35061324 | -0.9471533 | -0.1633025 | 0.87245594 | 0.99999785 | -6.3728293 | down |
| KI - WT, cell_origin: iPSC | O94913 | PCF11 | 15 | Homo sapien | -0.1718118 | -2.4144238 | 2.07080017 | -1.9405988 | -0.1632681 | 0.87248261 | 0.99999785 | -6.3728351 | down |
| KI - WT, cell_origin: iPSC | Q9NVH2 | INTS7 | 9 | Homo sapien | 0.13368527 | -1.6160928 | 1.88346336 | -0.0704847 | 0.1628182 | 0.87283067 | 0.99999785 | -6.372911 | up |
| KI - WT, cell_origin: iPSC | Q8NCA5 | FAM98A | 12 | Homo sapien | -0.0761056 | -1.0733703 | 0.921159 | 2.51143736 | -0.162633 | 0.87297395 | 0.99999785 | -6.3729422 | down |
| KI - WT, cell_origin: iPSC | Q9NR12 | PDLIM7 | 14 | Homo sapien | -0.0843969 | -1.1907215 | 1.02192772 | 1.31937376 | -0.1625722 | 0.87302104 | 0.99999785 | -6.3729524 | down |
| KI - WT, cell_origin: iPSC | P46087 | NOP2 | 13 | Homo sapien | -0.0527448 | -0.7465345 | 0.64104499 | -0.3290885 | -0.1619165 | 0.8735135 | 0.99999785 | -6.3730683 | down |
| KI - WT, cell_origin: iPSC | P49754 | VPS41 | 11 | Homo sapien | 0.06431447 | -0.7825426 | 0.91117156 | -1.5147568 | 0.16174776 | 0.87364412 | 0.99999785 | -6.3730966 | up |
| KI - WT, cell_origin: iPSC | Q96CS2 | HAUS1 | 3 | Homo sapien | 0.07797411 | -0.9491483 | 1.10509654 | -1.7092872 | 0.16171182 | 0.87367611 | 0.99999785 | -6.373101 | up |
| KI - WT, cell_origin: iPSC | Q13601 | KRR1 | 8 | Homo sapien | 0.0929327 | -1.1331725 | 1.31903795 | -1.2320432 | 0.16152628 | 0.87383037 | 0.99999785 | -6.3731279 | up |

| | | | | | | | | | | | | | |
|---|---|---|---|---|---|---|---|---|---|---|---|---|---|
| KI - WT, cell_origin: iPSC | Q8WWP7 | GIMAP1 | 12 | Homo sapien | 0.30457783 | -3.7159256 | 4.32508122 | -1.4968187 | 0.16144337 | 0.87389453 | 0.99999785 | -6.3731417 | up |
| KI - WT, cell_origin: iPSC | Q9GZS3 | WDR61 | 10 | Homo sapien | 0.06994526 | -0.8547649 | 0.99465542 | -0.1797444 | 0.16117276 | 0.8741004 | 0.99999785 | -6.3731883 | up |
| KI - WT, cell_origin: iPSC | Q99570 | PIK3R4 | 22 | Homo sapien | 0.06949489 | -0.8500277 | 0.98901745 | -0.8423486 | 0.16101694 | 0.87421774 | 0.99999785 | -6.3732155 | up |
| KI - WT, cell_origin: iPSC | Q96MX6 | DNAAF10 | 2 | Homo sapien | 0.07439984 | -0.9111021 | 1.0599018 | -1.5583245 | 0.16081306 | 0.87437137 | 0.99999785 | -6.3732511 | up |
| KI - WT, cell_origin: iPSC | Q2NL82 | TSR1 | 19 | Homo sapien | 0.09511495 | -1.1671652 | 1.35739512 | 0.63684158 | 0.16058146 | 0.87456163 | 0.99999785 | -6.3732854 | up |
| KI - WT, cell_origin: iPSC | Q9NX02 | NLRP2 | 31 | Homo sapien | -0.0780576 | -1.1146892 | 0.95857395 | -1.3914186 | -0.1604249 | 0.87467598 | 0.99999785 | -6.373314 | down |
| KI - WT, cell_origin: iPSC | Q8NHH9 | ATL2 | 3 | Homo sapien | -0.1351445 | -1.9300512 | 1.65976213 | -3.5537909 | -0.1604002 | 0.87469332 | 0.99999785 | -6.3733188 | down |
| KI - WT, cell_origin: iPSC | P55212 | CASP6 | 5 | Homo sapien | -0.0692282 | -0.9890506 | 0.85059432 | -0.7291715 | -0.1603471 | 0.87473628 | 0.99999785 | -6.3733269 | down |
| KI - WT, cell_origin: iPSC | O95758 | PTBP3 | 12 | Homo sapien | -0.0526113 | -0.7519496 | 0.64672694 | 2.56119554 | -0.1602277 | 0.87482108 | 0.99999785 | -6.3733497 | down |
| KI - WT, cell_origin: iPSC | Q8WVV9 | HNRNPLL | 21 | Homo sapien | -0.0597555 | -0.855381 | 0.73587003 | 0.98958774 | -0.1599878 | 0.87501083 | 0.99999785 | -6.3733878 | down |
| KI - WT, cell_origin: iPSC | P78344 | EIF4G2 | 27 | Homo sapien | 0.09892269 | -1.2200957 | 1.41794104 | 0.78213064 | 0.159826 | 0.8751464 | 0.99999785 | -6.3734106 | up |
| KI - WT, cell_origin: iPSC | Q9H4A6 | GOLPH3 | 4 | Homo sapien | 0.141735 | -1.7516242 | 2.03509417 | -1.1719612 | 0.1595315 | 0.87537439 | 0.99999785 | -6.3734593 | up |
| KI - WT, cell_origin: iPSC | Q9BQ75 | CMSS1 | 6 | Homo sapien | 0.05415537 | -0.6716446 | 0.77995538 | -0.6935416 | 0.15891478 | 0.87583727 | 0.99999785 | -6.3735665 | up |
| KI - WT, cell_origin: iPSC | Q8N5C6 | SRBD1 | 20 | Homo sapien | 0.11799863 | -1.4649307 | 1.70092796 | -0.7462764 | 0.15886118 | 0.87589335 | 0.99999785 | -6.3735698 | up |
| KI - WT, cell_origin: iPSC | Q9P2Q2 | FRMD4A | 4 | Homo sapien | -0.0508928 | -0.7334267 | 0.63164111 | -1.183244 | -0.1588078 | 0.8759201 | 0.99999785 | -6.3735841 | down |
| KI - WT, cell_origin: iPSC | Q13464 | ROCK1 | 15 | Homo sapien | 0.06439383 | -0.8008311 | 0.92961873 | -1.0267074 | 0.15853386 | 0.87613596 | 0.99999785 | -6.3736277 | up |
| KI - WT, cell_origin: iPSC | Q5H9R7 | PPP6R3 | 7 | Homo sapien | -0.0753278 | -1.0875287 | 0.9368731 | -1.5895473 | -0.1585282 | 0.87614094 | 0.99999785 | -6.3736284 | down |
| KI - WT, cell_origin: iPSC | Q9H0D6 | XRN2 | 24 | Homo sapien | 0.06511308 | -0.8103883 | 0.94061447 | 1.09886148 | 0.15846871 | 0.87619335 | 0.99999785 | -6.3736357 | up |
| KI - WT, cell_origin: iPSC | Q15061 | WDR43 | 10 | Homo sapien | -0.1472445 | -2.1364336 | 1.84194451 | -1.441633 | -0.1577486 | 0.87675485 | 0.99999785 | -6.3737521 | down |
| KI - WT, cell_origin: iPSC | Q99467 | CD180 | 3 | Homo sapien | 0.3617215 | -4.5268121 | 5.25025508 | -4.4361697 | 0.1576878 | 0.87680194 | 0.99999785 | -6.373762 | up |
| KI - WT, cell_origin: iPSC | Q14644 | RASA3 | 4 | Homo sapien | 0.07747094 | -0.9697547 | 1.12469653 | -1.8973936 | 0.15757901 | 0.87687503 | 0.99999785 | -6.373784 | up |
| KI - WT, cell_origin: iPSC | P41252 | IARS1 | 31 | Homo sapien | 0.05639715 | -0.7072296 | 0.82002393 | 0.84658985 | 0.15731959 | 0.87707634 | 0.99999785 | -6.3738261 | up |
| KI - WT, cell_origin: iPSC | Q5T0N5 | FNBP1L | 23 | Homo sapien | 0.07734847 | -0.9713231 | 1.12602004 | -0.3515166 | 0.15718628 | 0.87719033 | 0.99999785 | -6.3738437 | up |
| KI - WT, cell_origin: iPSC | Q14151 | SAFB2 | 7 | Homo sapien | -0.1393793 | -2.0393982 | 1.76063959 | -1.6195817 | -0.1563302 | 0.87785341 | 0.99999785 | -6.3739827 | down |
| KI - WT, cell_origin: iPSC | P42226 | STAT6 | 11 | Homo sapien | -0.1199968 | -1.7579882 | 1.51799465 | -1.6793632 | -0.1561206 | 0.87801573 | 0.99999785 | -6.3740165 | down |
| KI - WT, cell_origin: iPSC | Q12872 | SFSWAP | 13 | Homo sapien | -0.1167316 | -1.7115384 | 1.47807511 | -0.4906602 | -0.155985 | 0.87812077 | 0.99999785 | -6.3740384 | down |
| KI - WT, cell_origin: iPSC | Q96CW1 | AP2M1 | 19 | Homo sapien | 0.04729657 | -0.6004423 | 0.69503543 | 2.14390454 | 0.15551396 | 0.8784714 | 0.99999785 | -6.3741198 | up |
| KI - WT, cell_origin: iPSC | Q9HBE1 | PATZ1 | 10 | Homo sapien | -0.1289296 | -1.8977721 | 1.63991287 | -1.5119614 | -0.1553338 | 0.87862528 | 0.99999785 | -6.3741434 | down |
| KI - WT, cell_origin: iPSC | O75449 | KATNA1 | 5 | Homo sapien | 0.08352741 | -1.0717033 | 1.23875814 | -2.6304094 | 0.15399263 | 0.87965024 | 0.99999785 | -6.3743634 | up |
| KI - WT, cell_origin: iPSC | P27694 | RPA1 | 9 | Homo sapien | -0.0469124 | -0.697494 | 0.60366911 | -0.002144 | -0.1535769 | 0.87997245 | 0.99999785 | -6.3744295 | down |
| KI - WT, cell_origin: iPSC | O95394 | PGM3 | 3 | Homo sapien | 0.16131067 | -2.0786857 | 2.40130703 | -3.6171433 | 0.15346811 | 0.88007083 | 0.99999785 | -6.3744416 | up |
| KI - WT, cell_origin: iPSC | Q8IV08 | PLD3 | 6 | Homo sapien | -0.1121359 | -1.6708566 | 1.44658484 | 0.4895408 | -0.1533129 | 0.88019114 | 0.99999785 | -6.3744662 | down |
| KI - WT, cell_origin: iPSC | P30041 | PRDX6 | 12 | Homo sapien | -0.0751471 | -1.1197413 | 0.96944716 | 1.10501053 | -0.1533087 | 0.88019436 | 0.99999785 | -6.3744669 | down |
| KI - WT, cell_origin: iPSC | Q14694 | USP10 | 19 | Homo sapien | -0.069641 | -1.0395379 | 0.90025597 | 0.6423059 | -0.1530177 | 0.88041987 | 0.99999785 | -6.374513 | down |
| KI - WT, cell_origin: iPSC | A8MWD9;P6 | SNRPGP15;SI | 4 | Homo sapien | -0.1006735 | -1.5028193 | 1.3014723 | 1.82235686 | -0.1530116 | 0.88042458 | 0.99999785 | -6.374514 | down |
| KI - WT, cell_origin: iPSC | Q14919 | DRAP1 | 3 | Homo sapien | 0.15264824 | -1.9744399 | 2.27973637 | -3.0685397 | 0.15293563 | 0.88048349 | 0.99999785 | -6.374526 | up |
| KI - WT, cell_origin: iPSC | Q92820 | GGH | 5 | Homo sapien | 0.13568246 | -1.7561238 | 2.02748875 | -1.1665269 | 0.15284435 | 0.88055422 | 0.99999785 | -6.3745405 | up |
| KI - WT, cell_origin: iPSC | Q8IYB5 | SMAP1 | 5 | Homo sapien | -0.0782161 | -1.1691694 | 1.01273719 | 1.24223586 | -0.1527891 | 0.88059708 | 0.99999785 | -6.3745492 | down |
| KI - WT, cell_origin: iPSC | Q58A45 | PAN3 | 13 | Homo sapien | -0.0531542 | -0.7952062 | 0.68889787 | 0.99190282 | -0.1525804 | 0.88074782 | 0.99999785 | -6.3745863 | down |
| KI - WT, cell_origin: iPSC | P62166 | NCS1 | 6 | Homo sapien | 0.13211813 | -1.7148876 | 1.97912382 | -1.5027127 | 0.15243916 | 0.88086827 | 0.99999785 | -6.3746045 | up |
| KI - WT, cell_origin: iPSC | Q9NU22 | MDN1 | 119 | Homo sapien | 0.11069726 | -1.4369504 | 1.65834494 | 1.84594967 | 0.15242883 | 0.88087627 | 0.99999785 | -6.3746061 | up |
| KI - WT, cell_origin: iPSC | P55072 | VCP | 27 | Homo sapien | -0.0513353 | -0.7687425 | 0.66607193 | 0.95665046 | -0.1524116 | 0.88087717 | 0.99999785 | -6.3746135 | down |
| KI - WT, cell_origin: iPSC | P15311 | EZR | 15 | Homo sapien | 0.10935535 | -1.4208812 | 1.63959189 | 0.34157838 | 0.15229437 | 0.88098049 | 0.99999785 | -6.3746274 | up |
| KI - WT, cell_origin: iPSC | P55060 | CSE1L | 17 | Homo sapien | 0.07149847 | -0.934583 | 1.07757998 | -0.3587667 | 0.15144884 | 0.8816359 | 0.99999785 | -6.3747603 | up |
| KI - WT, cell_origin: iPSC | O75116 | ROCK2 | 16 | Homo sapien | -0.0437686 | -0.6596342 | 0.57209695 | -0.9664007 | -0.1513619 | 0.88168943 | 0.99999785 | -6.374779 | down |
| KI - WT, cell_origin: iPSC | O15067 | PFAS | 23 | Homo sapien | -0.0849314 | -1.2850654 | 1.11520266 | 0.50614379 | -0.1508137 | 0.88212827 | 0.99999785 | -6.3748597 | down |

| | | | | | | | | | | | | | |
|---|---|---|---|---|---|---|---|---|---|---|---|---|---|
| KI - WT, cell_origin: iPSC | O14920 | IKBKB | 8 | Homo sapien | -0.0774226 | -1.1719676 | 1.01712241 | -1.3895164 | -0.1507238 | 0.88219509 | 0.99999785 | -6.3748747 | down |
| KI - WT, cell_origin: iPSC | Q9H479 | FN3K | 8 | Homo sapien | 0.0686956 | -0.9027314 | 1.04012264 | -0.270898 | 0.15070273 | 0.88221432 | 0.99999785 | -6.374877 | up |
| KI - WT, cell_origin: iPSC | P78332 | RBM6 | 35 | Homo sapien | -0.0555169 | -0.8409344 | 0.72990055 | 0.477553 | -0.15057 | 0.88230731 | 0.99999785 | -6.3749013 | down |
| KI - WT, cell_origin: iPSC | Q9NVU7 | SDAD1 | 14 | Homo sapien | -0.0798779 | -1.2111347 | 1.05137896 | -0.2024522 | -0.1504762 | 0.88238997 | 0.99999785 | -6.3749123 | down |
| KI - WT, cell_origin: iPSC | Q86UQ0 | ZNF589 | 2 | Homo sapien | -0.037511 | -0.5691576 | 0.49413555 | -0.5619182 | -0.1502712 | 0.88253515 | 0.99999785 | -6.3749492 | down |
| KI - WT, cell_origin: iPSC | Q9UJX3 | ANAPC7 | 6 | Homo sapien | 0.04970633 | -0.6562618 | 0.75567441 | -1.2814552 | 0.14995684 | 0.88277889 | 0.99999785 | -6.3749981 | up |
| KI - WT, cell_origin: iPSC | Q9Y4W2 | LAS1L | 3 | Homo sapien | 0.14533421 | -1.9240175 | 2.21468589 | -2.0114405 | 0.1496704 | 0.88301474 | 0.99999785 | -6.3750375 | up |
| KI - WT, cell_origin: iPSC | Q9UPT8 | ZC3H4 | 9 | Homo sapien | -0.0651373 | -0.9930772 | 0.86280252 | 0.689315 | -0.1495934 | 0.88307445 | 0.99999785 | -6.3750494 | down |
| KI - WT, cell_origin: iPSC | Q14558 | PRPSAP1 | 6 | Homo sapien | 0.09557089 | -1.2671463 | 1.4582881 | -1.8651704 | 0.14945906 | 0.88317863 | 0.99999785 | -6.3750702 | up |
| KI - WT, cell_origin: iPSC | Q68DK2 | ZFYVE26 | 13 | Homo sapien | -0.0545446 | -0.8341377 | 0.72504853 | -0.6106272 | -0.1490245 | 0.88350372 | 0.99999785 | -6.3751417 | down |
| KI - WT, cell_origin: iPSC | Q9UFF9 | CNOT8 | 4 | Homo sapien | 0.05210203 | -0.6925248 | 0.79672886 | -1.2786143 | 0.14902382 | 0.88350248 | 0.99999785 | -6.3751424 | up |
| KI - WT, cell_origin: iPSC | Q9NW38 | FANCL | 6 | Homo sapien | -0.0417633 | -0.6404744 | 0.55694777 | -0.0823292 | -0.1485652 | 0.88385818 | 0.99999785 | -6.375213 | down |
| KI - WT, cell_origin: iPSC | A6NHR9 | SMCHD1 | 4 | Homo sapien | 0.10423711 | -1.3913168 | 1.59979106 | -2.5746136 | 0.148522 | 0.88390367 | 0.99999785 | -6.3752154 | up |
| KI - WT, cell_origin: iPSC | Q9Y6Q5 | AP1M2 | 9 | Homo sapien | 0.08599483 | -1.1487006 | 1.32069024 | 1.45420602 | 0.14842767 | 0.88397848 | 0.99999785 | -6.3752293 | up |
| KI - WT, cell_origin: iPSC | Q9NR56 | MBNL1 | 3 | Homo sapien | -0.0699455 | -1.0743094 | 0.93441848 | -0.5515918 | -0.1484127 | 0.88399012 | 0.99999785 | -6.3752316 | down |
| KI - WT, cell_origin: iPSC | P43405 | SYK | 4 | Homo sapien | -0.1758562 | -2.7059564 | 2.35424388 | -1.5601002 | -0.148123 | 0.88421479 | 0.99999785 | -6.3752761 | down |
| KI - WT, cell_origin: iPSC | Q96N67 | DOCK7 | 22 | Homo sapien | -0.0691981 | -1.0676136 | 0.92921734 | -0.6453827 | -0.1477017 | 0.88454157 | 0.99999785 | -6.3753406 | down |
| KI - WT, cell_origin: iPSC | P28340 | POLD1 | 19 | Homo sapien | 0.17747109 | -2.395931 | 2.75087319 | -1.1483634 | 0.14696784 | 0.88511081 | 0.99999785 | -6.3754525 | up |
| KI - WT, cell_origin: iPSC | Q9Y3X0 | CCDC9 | 4 | Homo sapien | 0.21128443 | -2.8525987 | 3.27516759 | -2.1898421 | 0.1469595 | 0.88511728 | 0.99999785 | -6.3754538 | up |
| KI - WT, cell_origin: iPSC | Q9UBB6 | NCDN | 4 | Homo sapien | -0.0513417 | -0.7957272 | 0.69304384 | -2.2639164 | -0.1468966 | 0.88515262 | 0.99999785 | -6.3754682 | down |
| KI - WT, cell_origin: iPSC | Q9UEW8 | STK39 | 5 | Homo sapien | 0.07379854 | -0.9991604 | 1.14675746 | -1.2188891 | 0.14656906 | 0.88541891 | 0.99999785 | -6.3755136 | up |
| KI - WT, cell_origin: iPSC | O76003 | GLRX3 | 12 | Homo sapien | -0.0558555 | -0.8678277 | 0.75611658 | 1.41773221 | -0.1465356 | 0.88543673 | 0.99999785 | -6.3755215 | down |
| KI - WT, cell_origin: iPSC | Q96GM5 | SMARCD1 | 11 | Homo sapien | -0.067769 | -1.0556619 | 0.92012383 | -0.5788339 | -0.1461921 | 0.88571264 | 0.99999785 | -6.3755702 | down |
| KI - WT, cell_origin: iPSC | P78364 | PHC1 | 2 | Homo sapien | 0.0563055 | -0.76441 | 0.87702097 | -0.8268884 | 0.14613488 | 0.88574649 | 0.99999785 | -6.3755826 | up |
| KI - WT, cell_origin: iPSC | Q9Y5V3 | MAGED1 | 11 | Homo sapien | -0.1060222 | -1.6527333 | 1.44068891 | 0.36303463 | -0.1460797 | 0.88579981 | 0.99999785 | -6.3755872 | down |
| KI - WT, cell_origin: iPSC | Q9Y5J1 | UTP18 | 10 | Homo sapien | -0.0893802 | -1.3931747 | 1.2144142 | -2.267257 | -0.1460545 | 0.8858133 | 0.99999785 | -6.3755932 | down |
| KI - WT, cell_origin: iPSC | Q96RN5 | MED15 | 5 | Homo sapien | -0.0582875 | -0.9104639 | 0.79388886 | -0.4879155 | -0.1457015 | 0.88608392 | 0.99999785 | -6.3756477 | down |
| KI - WT, cell_origin: iPSC | P12532 | CKMT1A | 6 | Homo sapien | -0.1414442 | -2.2170926 | 1.93420428 | -3.4632556 | -0.1452224 | 0.88646502 | 0.99999785 | -6.3757165 | down |
| KI - WT, cell_origin: iPSC | Q13177 | PAK2 | 24 | Homo sapien | 0.1333631 | -1.8254936 | 2.09221979 | 2.94587504 | 0.1450893 | 0.88656831 | 0.99999785 | -6.3757365 | up |
| KI - WT, cell_origin: iPSC | Q15020 | SART3 | 11 | Homo sapien | -0.071085 | -1.1154459 | 0.97327597 | -1.125615 | -0.1450433 | 0.88660239 | 0.99999785 | -6.375744 | down |
| KI - WT, cell_origin: iPSC | Q8IU60 | DCP2 | 2 | Homo sapien | 0.08205513 | -1.1255443 | 1.2896546 | -2.3777768 | 0.14474035 | 0.88682922 | 0.99999785 | -6.3757924 | up |
| KI - WT, cell_origin: iPSC | Q13057 | COASY | 5 | Homo sapien | -0.0799639 | -1.2588285 | 1.09890064 | -1.3197891 | -0.1445548 | 0.88698308 | 0.99999785 | -6.3758167 | down |
| KI - WT, cell_origin: iPSC | P68431;Q16( | H3C1;H3-4;H | 3 | Homo sapien | -0.062418 | -0.9830815 | 0.85824555 | 3.10802091 | -0.1444811 | 0.88704026 | 0.99999785 | -6.3758277 | down |
| KI - WT, cell_origin: iPSC | Q9BY42 | RTF2 | 2 | Homo sapien | 0.08956014 | -1.2380952 | 1.41721552 | -1.2086658 | 0.14375791 | 0.88760154 | 0.99999785 | -6.3759356 | up |
| KI - WT, cell_origin: iPSC | Q5JPH6 | EARS2 | 2 | Homo sapien | 0.119489 | -1.6530591 | 1.89203708 | -4.3705646 | 0.14357195 | 0.88773273 | 0.99999785 | -6.3759679 | up |
| KI - WT, cell_origin: iPSC | P61964 | WDR5 | 9 | Homo sapien | -0.0727714 | -1.1549692 | 1.00942647 | -1.0100891 | -0.1433033 | 0.88795438 | 0.99999785 | -6.3760032 | down |
| KI - WT, cell_origin: iPSC | P47756 | CAPZB | 10 | Homo sapien | -0.0506318 | -0.8038289 | 0.70256531 | 2.51739787 | -0.1431929 | 0.88803029 | 0.99999785 | -6.376023 | down |
| KI - WT, cell_origin: iPSC | P49790 | NUP153 | 4 | Homo sapien | 0.1404315 | -1.9575969 | 2.23845991 | -2.4329504 | 0.14264467 | 0.88846564 | 0.99999785 | -6.3761007 | up |
| KI - WT, cell_origin: iPSC | O60347 | TBC1D12 | 3 | Homo sapien | -0.082731 | -1.3183532 | 1.15289115 | -2.1506133 | -0.1426387 | 0.88846292 | 0.99999785 | -6.3761041 | down |
| KI - WT, cell_origin: iPSC | Q9Y294 | ASF1A | 3 | Homo sapien | -0.0525199 | -0.8387392 | 0.73369943 | -0.5804701 | -0.1422865 | 0.88873286 | 0.99999785 | -6.3761573 | down |
| KI - WT, cell_origin: iPSC | P62081 | RPS7 | 10 | Homo sapien | -0.0314581 | -0.5041176 | 0.44120135 | 5.10524565 | -0.1417503 | 0.88914699 | 0.99999785 | -6.3762369 | down |
| KI - WT, cell_origin: iPSC | P08621 | SNRNP70 | 18 | Homo sapien | -0.0703924 | -1.1286875 | 0.98790275 | 1.96170763 | -0.1417494 | 0.88916063 | 0.99999785 | -6.3762325 | down |
| KI - WT, cell_origin: iPSC | Q8N9Q2 | SREK1IP1 | 5 | Homo sapien | 0.08804234 | -1.2388904 | 1.41497504 | 0.10765658 | 0.14139859 | 0.88943302 | 0.99999785 | -6.376284 | up |
| KI - WT, cell_origin: iPSC | O14757 | CHEK1 | 14 | Homo sapien | -0.0980801 | -1.5784814 | 1.38232112 | -0.3910444 | -0.14119 | 0.88959498 | 0.99999785 | -6.3763145 | down |
| KI - WT, cell_origin: iPSC | Q99575 | POP1 | 5 | Homo sapien | -0.0661351 | -1.065452 | 0.93318176 | -0.7192154 | -0.1410365 | 0.88971419 | 0.99999785 | -6.3763369 | down |
| KI - WT, cell_origin: iPSC | P43487 | RANBP1 | 3 | Homo sapien | -0.1278319 | -2.0597561 | 1.80409229 | -0.0238317 | -0.1410105 | 0.88973434 | 0.99999785 | -6.3763407 | down |

| | | | | | | | | | | | | | |
|---|---|---|---|---|---|---|---|---|---|---|---|---|---|
| KI - WT, cell_origin: iPSC | Q9BYJ9 | YTHDF1 | 7 | Homo sapien | -0.0368958 | -0.5947007 | 0.52090903 | 0.52127327 | -0.1408753 | 0.88982644 | 0.99999785 | -6.3763648 | down |
| KI - WT, cell_origin: iPSC | O15260 | SURF4 | 5 | Homo sapien | 0.15889781 | -2.2519978 | 2.56979339 | 0.72303019 | 0.14045651 | 0.89016452 | 0.99999785 | -6.3764214 | up |
| KI - WT, cell_origin: iPSC | Q9BXY0 | MAK16 | 2 | Homo sapien | 0.05501315 | -0.7801754 | 0.89020168 | -1.2350875 | 0.1402903 | 0.89028103 | 0.99999785 | -6.3764499 | up |
| KI - WT, cell_origin: iPSC | O14647 | CHD2 | 3 | Homo sapien | -0.0692862 | -1.1241589 | 0.9855865 | -1.8530511 | -0.1399115 | 0.89057821 | 0.99999785 | -6.3765038 | down |
| KI - WT, cell_origin: iPSC | P49257 | LMAN1 | 16 | Homo sapien | 0.15584468 | -2.2236823 | 2.53537165 | 2.03783958 | 0.13957374 | 0.89085006 | 0.99999785 | -6.3765494 | up |
| KI - WT, cell_origin: iPSC | Q08AD1 | CAMSAP2 | 7 | Homo sapien | -0.0455455 | -0.7410031 | 0.64991205 | -1.1731435 | -0.1394809 | 0.8909094 | 0.99999785 | -6.3765672 | down |
| KI - WT, cell_origin: iPSC | Q5VU43 | PDE4DIP | 2 | Homo sapien | -0.0990334 | -1.6133612 | 1.41529446 | -0.8428545 | -0.1393683 | 0.89100959 | 0.99999785 | -6.3765791 | down |
| KI - WT, cell_origin: iPSC | Q7Z460 | CLASP1 | 31 | Homo sapien | 0.04594057 | -0.6570737 | 0.74895483 | -0.1663693 | 0.13917842 | 0.89114434 | 0.99999785 | -6.3766108 | up |
| KI - WT, cell_origin: iPSC | P15927 | RPA2 | 2 | Homo sapien | 0.05037964 | -0.7217319 | 0.82249115 | -0.7960681 | 0.13896987 | 0.89130663 | 0.99999785 | -6.3766407 | up |
| KI - WT, cell_origin: iPSC | Q12904 | AIMP1 | 12 | Homo sapien | 0.0530238 | -0.7600148 | 0.86606245 | 0.19574279 | 0.1389241 | 0.89134568 | 0.99999785 | -6.3766461 | up |
| KI - WT, cell_origin: iPSC | Q96AG4 | LRRC59 | 8 | Homo sapien | -0.1543341 | -2.523659 | 2.21499085 | -0.2537825 | -0.138816 | 0.89143855 | 0.99999785 | -6.3766586 | down |
| KI - WT, cell_origin: iPSC | Q9NQX3 | GPHN | 9 | Homo sapien | 0.05188324 | -0.7463338 | 0.85010032 | -0.613243 | 0.13845281 | 0.8917107 | 0.99999785 | -6.3767141 | up |
| KI - WT, cell_origin: iPSC | Q9UJU6 | DBNL | 11 | Homo sapien | -0.0546349 | -0.8972937 | 0.78802383 | 1.11268154 | -0.1381337 | 0.89196269 | 0.99999785 | -6.3767584 | down |
| KI - WT, cell_origin: iPSC | Q9H2U1 | DHX36 | 36 | Homo sapien | 0.04006427 | -0.5783183 | 0.65844683 | 1.80684812 | 0.13798746 | 0.89206952 | 0.99999785 | -6.3767816 | up |
| KI - WT, cell_origin: iPSC | Q9Y6X9 | MORC2 | 5 | Homo sapien | -0.0888793 | -1.4618567 | 1.28409801 | -1.189646 | -0.1379557 | 0.89210679 | 0.99999785 | -6.3767819 | down |
| KI - WT, cell_origin: iPSC | Q86UW6 | N4BP2 | 6 | Homo sapien | 0.05809536 | -0.8389083 | 0.95509899 | -2.0454641 | 0.13793897 | 0.8921072 | 0.99999785 | -6.3767886 | up |
| KI - WT, cell_origin: iPSC | Q99543 | DNAJC2 | 15 | Homo sapien | -0.0632898 | -1.043299 | 0.91671936 | -0.6815239 | -0.1376199 | 0.8923665 | 0.99999785 | -6.3768302 | down |
| KI - WT, cell_origin: iPSC | Q6ZSZ5 | ARHGEF18 | 7 | Homo sapien | -0.1617201 | -2.6672908 | 2.3438505 | -1.5726511 | -0.1375498 | 0.89242217 | 0.99999785 | -6.3768398 | down |
| KI - WT, cell_origin: iPSC | Q9BXB5 | OSBPL10 | 15 | Homo sapien | 0.04183576 | -0.6072807 | 0.69095224 | 0.22182472 | 0.13726654 | 0.89262964 | 0.99999785 | -6.3768843 | up |
| KI - WT, cell_origin: iPSC | P48634 | PRRC2A | 30 | Homo sapien | -0.0546693 | -0.9039805 | 0.79464178 | 1.9084083 | -0.1371381 | 0.89273617 | 0.99999785 | -6.3769003 | down |
| KI - WT, cell_origin: iPSC | Q8TDM6 | DLG5 | 3 | Homo sapien | 0.08091209 | -1.1773798 | 1.33920393 | -1.2801369 | 0.13703591 | 0.89282137 | 0.99999785 | -6.3769129 | up |
| KI - WT, cell_origin: iPSC | P55084 | HADHB | 4 | Homo sapien | 0.32359768 | -4.742334 | 5.38952936 | -1.473362 | 0.13612831 | 0.89352655 | 0.99999785 | -6.3770413 | up |
| KI - WT, cell_origin: iPSC | Q9BRR8 | GPATCH1 | 8 | Homo sapien | 0.0693395 | -1.0172716 | 1.1559506 | -1.5194616 | 0.13596064 | 0.8936523 | 0.99999785 | -6.3770664 | up |
| KI - WT, cell_origin: iPSC | Q8IWS0 | PHF6 | 6 | Homo sapien | -0.0635867 | -1.0620557 | 0.93488226 | 0.21415732 | -0.135717 | 0.89384618 | 0.99999785 | -6.3770992 | down |
| KI - WT, cell_origin: iPSC | P63220 | RPS21 | 8 | Homo sapien | -0.0396414 | -0.6621559 | 0.58287302 | 3.60696803 | -0.135625 | 0.89390524 | 0.99999785 | -6.3771162 | down |
| KI - WT, cell_origin: iPSC | Q9UHQ1 | NARF | 5 | Homo sapien | -0.0961212 | -1.6121703 | 1.41992798 | -1.3376466 | -0.1351164 | 0.89431288 | 0.99999785 | -6.3771834 | down |
| KI - WT, cell_origin: iPSC | Q9BRD0 | BUD13 | 7 | Homo sapien | -0.0495399 | -0.8313959 | 0.73231608 | -1.7722561 | -0.1349485 | 0.894431 | 0.99999785 | -6.3772109 | down |
| KI - WT, cell_origin: iPSC | Q71U36 | TUBA1A | 27 | Homo sapien | 0.02848774 | -0.4214659 | 0.47844135 | 7.47382842 | 0.13484349 | 0.89451263 | 0.99999785 | -6.3772256 | up |
| KI - WT, cell_origin: iPSC | P32121 | ARRB2 | 4 | Homo sapien | -0.047981 | -0.8071576 | 0.7111956 | -0.1803653 | -0.1346162 | 0.89469074 | 0.99999785 | -6.3772568 | down |
| KI - WT, cell_origin: iPSC | P53350 | PLK1 | 10 | Homo sapien | 0.04996635 | -0.7408133 | 0.84074602 | -0.4451279 | 0.13459156 | 0.89471108 | 0.99999785 | -6.3772599 | up |
| KI - WT, cell_origin: iPSC | Q8NHV1 | GIMAP7 | 7 | Homo sapien | -0.1336322 | -2.2566292 | 1.98936474 | -1.1601232 | -0.1341418 | 0.89507031 | 0.99999785 | -6.3773193 | down |
| KI - WT, cell_origin: iPSC | Q15334 | LLGL1 | 11 | Homo sapien | -0.0490753 | -0.8287247 | 0.73057416 | -0.7100298 | -0.1340705 | 0.8951149 | 0.99999785 | -6.3773327 | down |
| KI - WT, cell_origin: iPSC | P35249 | RFC4 | 15 | Homo sapien | -0.0651484 | -1.1007181 | 0.97042133 | -1.3047007 | -0.1340389 | 0.89514581 | 0.99999785 | -6.377335 | down |
| KI - WT, cell_origin: iPSC | Q9BVP2 | GNL3 | 18 | Homo sapien | 0.16019454 | -2.3976618 | 2.71805088 | 0.27468511 | 0.13346701 | 0.89559483 | 0.99999785 | -6.3774128 | up |
| KI - WT, cell_origin: iPSC | P67870 | CSNK2B | 5 | Homo sapien | 0.04811162 | -0.7202959 | 0.81651912 | 1.33507164 | 0.13337217 | 0.89565947 | 0.99999785 | -6.3774288 | up |
| KI - WT, cell_origin: iPSC | Q9H0L4 | CSTF2T | 12 | Homo sapien | -0.0812495 | -1.3814545 | 1.21895541 | 0.17753976 | -0.1331716 | 0.89582449 | 0.99999785 | -6.3774536 | down |
| KI - WT, cell_origin: iPSC | Q06265 | EXOSC9 | 4 | Homo sapien | -0.0416337 | -0.7078827 | 0.62461527 | -0.0133146 | -0.1330909 | 0.895875 | 0.99999785 | -6.3774686 | down |
| KI - WT, cell_origin: iPSC | P78345 | RPP38 | 2 | Homo sapien | -0.0939767 | -1.6009122 | 1.41295889 | -1.6074347 | -0.1329008 | 0.89603495 | 0.99999785 | -6.3774909 | down |
| KI - WT, cell_origin: iPSC | Q8N5I9 | NOPCHAP1 | 4 | Homo sapien | 0.11067237 | -1.6666877 | 1.88803243 | -2.1259236 | 0.13269855 | 0.8961922 | 0.99999785 | -6.3775187 | up |
| KI - WT, cell_origin: iPSC | Q2M1K9 | ZNF423 | 9 | Homo sapien | 0.07681742 | -1.15866 | 1.3122948 | -0.7657091 | 0.13250349 | 0.89634384 | 0.99999785 | -6.3775455 | up |
| KI - WT, cell_origin: iPSC | P24534 | EEF1B2 | 5 | Homo sapien | -0.0601923 | -1.0335055 | 0.91312096 | 0.58464231 | -0.1317924 | 0.89689669 | 0.99999785 | -6.3776428 | down |
| KI - WT, cell_origin: iPSC | Q9Y5U2 | TSSC4 | 2 | Homo sapien | -0.040597 | -0.6980967 | 0.61690264 | 0.43628213 | -0.1315039 | 0.89710898 | 0.99999785 | -6.377686 | down |
| KI - WT, cell_origin: iPSC | O15182 | CETN3 | 4 | Homo sapien | -0.0729232 | -1.2546293 | 1.10878288 | -1.8166146 | -0.1314887 | 0.89712962 | 0.99999785 | -6.3776852 | down |
| KI - WT, cell_origin: iPSC | P22061 | PCMT1 | 9 | Homo sapien | 0.03643884 | -0.5546838 | 0.62756146 | 1.78268667 | 0.13128847 | 0.8972765 | 0.99999785 | -6.3777153 | up |
| KI - WT, cell_origin: iPSC | Q5T6F2 | UBAP2 | 11 | Homo sapien | -0.0812723 | -1.4021528 | 1.2396081 | 0.13617871 | -0.1311238 | 0.89741653 | 0.99999785 | -6.3777338 | down |
| KI - WT, cell_origin: iPSC | P61586 | RHOA | 17 | Homo sapien | 0.04698158 | -0.7167589 | 0.81072206 | 7.79235811 | 0.1310944 | 0.89743941 | 0.99999785 | -6.3777378 | up |

| | | | | | | | | | | | | | |
|---|---|---|---|---|---|---|---|---|---|---|---|---|---|
| KI - WT, cell_origin: iPSC | Q8N4J0 | CARNMT1 | 4 | Homo sapien | 0.1331094 | -2.0344065 | 2.30062526 | -2.7098454 | 0.13087263 | 0.89761185 | 0.99999785 | -6.3777679 | up |
| KI - WT, cell_origin: iPSC | P53365 | ARFIP2 | 5 | Homo sapien | 0.05397436 | -0.8261699 | 0.93411858 | -0.6516367 | 0.1306326 | 0.89779012 | 0.99999785 | -6.3778031 | up |
| KI - WT, cell_origin: iPSC | O43913 | ORC5 | 6 | Homo sapien | -0.0654721 | -1.1363029 | 1.00535868 | -1.2397702 | -0.1302885 | 0.89806463 | 0.99999785 | -6.3778474 | down |
| KI - WT, cell_origin: iPSC | Q96FW1 | OTUB1 | 6 | Homo sapien | -0.0510165 | -0.8913584 | 0.78932551 | -1.7056302 | -0.1292986 | 0.89882411 | 0.99999785 | -6.3779837 | down |
| KI - WT, cell_origin: iPSC | Q9BRJ6 | C7orf50 | 5 | Homo sapien | -0.0346732 | -0.6062915 | 0.53694507 | 0.01850312 | -0.1291896 | 0.89890889 | 0.99999785 | -6.3779983 | down |
| KI - WT, cell_origin: iPSC | P40937 | RFC5 | 10 | Homo sapien | 0.06707272 | -1.0432337 | 1.1773791 | -0.3763569 | 0.12873761 | 0.89927229 | 0.99999785 | -6.3780549 | up |
| KI - WT, cell_origin: iPSC | Q92841 | DDX17 | 29 | Homo sapien | -0.0985725 | -1.7304704 | 1.53332546 | 4.34466264 | -0.1287256 | 0.89928162 | 0.99999785 | -6.3780565 | down |
| KI - WT, cell_origin: iPSC | Q9UER7 | DAXX | 3 | Homo sapien | 0.05203254 | -0.8091638 | 0.91322889 | -2.2922594 | 0.12868045 | 0.89930499 | 0.99999785 | -6.3780662 | up |
| KI - WT, cell_origin: iPSC | O95793 | STAU1 | 21 | Homo sapien | 0.0391172 | -0.6085704 | 0.68680485 | 2.1649584 | 0.12862988 | 0.89934433 | 0.99999785 | -6.378073 | up |
| KI - WT, cell_origin: iPSC | Q5VTB9 | RNF220 | 3 | Homo sapien | 0.05048414 | -0.7857051 | 0.88667342 | -1.3496088 | 0.12858493 | 0.8993793 | 0.99999785 | -6.378079 | up |
| KI - WT, cell_origin: iPSC | Q2TAY7 | SMU1 | 10 | Homo sapien | -0.0535831 | -0.9424856 | 0.83531943 | 0.53953816 | -0.1284436 | 0.89949814 | 0.99999785 | -6.378095 | down |
| KI - WT, cell_origin: iPSC | Q16775 | HAGH | 3 | Homo sapien | -0.0571498 | -1.0083358 | 0.89403615 | -2.1130148 | -0.1279644 | 0.89986205 | 0.99999785 | -6.3781614 | down |
| KI - WT, cell_origin: iPSC | Q9NPF5 | DMAP1 | 6 | Homo sapien | 0.04068054 | -0.6398544 | 0.72121547 | -0.6296328 | 0.12731395 | 0.90036813 | 0.99999785 | -6.3782473 | up |
| KI - WT, cell_origin: iPSC | P53396 | ACLY | 33 | Homo sapien | -0.052435 | -0.9306431 | 0.82577308 | 0.70190509 | -0.127222 | 0.90044849 | 0.99999785 | -6.3782567 | down |
| KI - WT, cell_origin: iPSC | O75534 | CSDE1 | 30 | Homo sapien | -0.0298479 | -0.5300732 | 0.47037738 | 1.63627953 | -0.1270831 | 0.90054778 | 0.99999785 | -6.3782777 | down |
| KI - WT, cell_origin: iPSC | Q96KR1 | ZFR | 16 | Homo sapien | -0.1106313 | -1.9665105 | 1.74524786 | -0.4787312 | -0.1270371 | 0.90059513 | 0.99999785 | -6.3782802 | down |
| KI - WT, cell_origin: iPSC | Q8IWZ8 | SUGP1 | 6 | Homo sapien | 0.07255282 | -1.1451307 | 1.29023631 | -1.6559991 | 0.12697617 | 0.90064255 | 0.99999785 | -6.3782882 | up |
| KI - WT, cell_origin: iPSC | Q96BK5 | PINX1 | 5 | Homo sapien | 0.06370913 | -1.0059578 | 1.13337602 | -0.764754 | 0.12692743 | 0.90068047 | 0.99999785 | -6.3782946 | up |
| KI - WT, cell_origin: iPSC | Q9NQZ2 | UTP3 | 2 | Homo sapien | -0.051759 | -0.9211961 | 0.81767812 | 0.06107938 | -0.126832 | 0.90074937 | 0.99999785 | -6.3783088 | down |
| KI - WT, cell_origin: iPSC | Q9NP50 | SINHCAF | 3 | Homo sapien | -0.1396522 | -2.5034487 | 2.22414441 | -5.1139497 | -0.1258442 | 0.90151427 | 0.99999785 | -6.3784392 | down |
| KI - WT, cell_origin: iPSC | O75131 | CPNE3 | 2 | Homo sapien | -0.0497882 | -0.8958225 | 0.79624602 | -0.5001584 | -0.1253594 | 0.90189255 | 0.99999785 | -6.3785019 | down |
| KI - WT, cell_origin: iPSC | Q07666 | KHDRBS1 | 12 | Homo sapien | -0.0755998 | -1.3645585 | 1.21335891 | 1.5055131 | -0.1249925 | 0.90218611 | 0.99999785 | -6.378547 | down |
| KI - WT, cell_origin: iPSC | O15042 | U2SURP | 36 | Homo sapien | -0.0660667 | -1.1944861 | 1.06235279 | 1.45943183 | -0.1247712 | 0.90235834 | 0.99999785 | -6.3785757 | down |
| KI - WT, cell_origin: iPSC | Q8IX01 | SUGP2 | 14 | Homo sapien | -0.0569805 | -1.0323157 | 0.91835478 | -0.2296485 | -0.1245015 | 0.90256824 | 0.99999785 | -6.3786105 | down |
| KI - WT, cell_origin: iPSC | Q9H2P0 | ADNP | 5 | Homo sapien | -0.1014983 | -1.8395634 | 1.63656685 | -1.5803946 | -0.12445 | 0.90260827 | 0.99999785 | -6.3786171 | down |
| KI - WT, cell_origin: iPSC | Q13617 | CUL2 | 6 | Homo sapien | -0.0435059 | -0.7894344 | 0.70242255 | -1.0439358 | -0.1242198 | 0.90277614 | 0.99999785 | -6.3786502 | down |
| KI - WT, cell_origin: iPSC | Q9BZF9 | UACA | 29 | Homo sapien | -0.0466464 | -0.8473943 | 0.75410144 | -1.0145283 | -0.1240739 | 0.90289048 | 0.99999785 | -6.3786687 | down |
| KI - WT, cell_origin: iPSC | Q9NYR9 | NKIRAS2 | 4 | Homo sapien | -0.0608086 | -1.1064762 | 0.98485901 | -1.3764178 | -0.1238951 | 0.90303502 | 0.99999785 | -6.3786901 | down |
| KI - WT, cell_origin: iPSC | O00762 | UBE2C | 4 | Homo sapien | -0.0245564 | -0.4477108 | 0.39859792 | -0.1632819 | -0.1235965 | 0.90326127 | 0.99999785 | -6.3787302 | down |
| KI - WT, cell_origin: iPSC | O95486 | SEC24A | 20 | Homo sapien | 0.05414263 | -0.8803971 | 0.98868233 | 0.94299351 | 0.12346503 | 0.90337493 | 0.99999785 | -6.3787436 | up |
| KI - WT, cell_origin: iPSC | Q9ULA0 | DNPEP | 18 | Homo sapien | -0.0593139 | -1.0849503 | 0.96632254 | 2.4600028 | -0.1232439 | 0.90354706 | 0.99999785 | -6.3787719 | down |
| KI - WT, cell_origin: iPSC | P11766 | ADH5 | 2 | Homo sapien | 0.04858657 | -0.7910615 | 0.88823463 | -1.4640942 | 0.12324196 | 0.9035373 | 0.99999785 | -6.3787755 | up |
| KI - WT, cell_origin: iPSC | P55884 | EIF3B | 31 | Homo sapien | -0.0276761 | -0.5061982 | 0.45084597 | 3.8038762 | -0.1231807 | 0.90358497 | 0.99999785 | -6.3787833 | down |
| KI - WT, cell_origin: iPSC | Q16851 | UGP2 | 13 | Homo sapien | -0.3099273 | -5.6793355 | 5.05948086 | -1.4009089 | -0.1230087 | 0.90373015 | 0.99999785 | -6.3788019 | down |
| KI - WT, cell_origin: iPSC | Q14974 | KPNB1 | 24 | Homo sapien | 0.03284226 | -0.5358622 | 0.60154671 | 1.47506409 | 0.12299461 | 0.90372987 | 0.99999785 | -6.378807 | up |
| KI - WT, cell_origin: iPSC | P22392 | NME2 | 7 | Homo sapien | 0.04415062 | -0.7224531 | 0.81075437 | 1.34664289 | 0.12267969 | 0.90397791 | 0.99999785 | -6.3788462 | up |
| KI - WT, cell_origin: iPSC | Q96GI7 | FAM89A | 4 | Homo sapien | -0.0887601 | -1.6339717 | 1.4564515 | 0.48084962 | -0.1224143 | 0.90419283 | 0.99999785 | -6.3788774 | down |
| KI - WT, cell_origin: iPSC | P40925 | MDH1 | 4 | Homo sapien | -0.0730897 | -1.3461908 | 1.20001145 | 0.08554092 | -0.1223476 | 0.90424478 | 0.99999785 | -6.3788859 | down |
| KI - WT, cell_origin: iPSC | Q9ULT8 | HECTD1 | 8 | Homo sapien | 0.05984953 | -0.9825121 | 1.1022112 | -2.3325868 | 0.12228749 | 0.90428041 | 0.99999785 | -6.3788968 | up |
| KI - WT, cell_origin: iPSC | P78347 | GTF2I | 17 | Homo sapien | 0.102511 | -1.6848677 | 1.88988973 | -0.1279904 | 0.12222394 | 0.90434105 | 0.99999785 | -6.3789015 | up |
| KI - WT, cell_origin: iPSC | Q99961 | SH3GL1 | 11 | Homo sapien | 0.03930144 | -0.6456562 | 0.72425907 | 0.74371445 | 0.12220372 | 0.90434563 | 0.99999785 | -6.3789074 | up |
| KI - WT, cell_origin: iPSC | Q12851 | MAP4K2 | 4 | Homo sapien | 0.05377122 | -0.8841976 | 0.99173999 | -0.9512981 | 0.12213569 | 0.9044046 | 0.99999785 | -6.3789142 | up |
| KI - WT, cell_origin: iPSC | Q14C86 | GAPVD1 | 7 | Homo sapien | 0.03734452 | -0.6151874 | 0.68987648 | -0.831011 | 0.12188907 | 0.90459062 | 0.99999785 | -6.3789472 | up |
| KI - WT, cell_origin: iPSC | P07814 | EPRS1 | 38 | Homo sapien | 0.03935094 | -0.6490773 | 0.72777918 | 0.68895107 | 0.12174079 | 0.90470608 | 0.99999785 | -6.3789659 | up |
| KI - WT, cell_origin: iPSC | O60333 | KIF1B | 9 | Homo sapien | -0.0447436 | -0.8282081 | 0.73872092 | -0.2859182 | -0.1216486 | 0.90478023 | 0.99999785 | -6.3789768 | down |
| KI - WT, cell_origin: iPSC | Q06124 | PTPN11 | 15 | Homo sapien | -0.0492303 | -0.9125787 | 0.81411805 | 0.74456116 | -0.1214964 | 0.90490384 | 0.99999785 | -6.3789944 | down |

| | | | | | | | | | | | | | |
|---|---|---|---|---|---|---|---|---|---|---|---|---|---|
| KI - WT, cell_origin: iPSC | O14910 | LIN7A | 3 | Homo sapien | -0.0651745 | -1.2085808 | 1.07823192 | 0.40170052 | -0.1214728 | 0.90492582 | 0.99999785 | -6.3789963 | down |
| KI - WT, cell_origin: iPSC | Q01433 | AMPD2 | 16 | Homo sapien | -0.039955 | -0.7422196 | 0.66230961 | 0.1392345 | -0.1211741 | 0.90514733 | 0.99999785 | -6.3790371 | down |
| KI - WT, cell_origin: iPSC | P25398 | RPS12 | 8 | Homo sapien | 0.02242692 | -0.3721474 | 0.41700125 | 5.21677471 | 0.12105435 | 0.90524059 | 0.99999785 | -6.3790522 | up |
| KI - WT, cell_origin: iPSC | Q9UNF1 | MAGED2 | 16 | Homo sapien | 0.11119705 | -1.8526824 | 2.07507651 | 0.4592646 | 0.12066486 | 0.90555493 | 0.99999785 | -6.3790977 | up |
| KI - WT, cell_origin: iPSC | Q96BD8 | SKA1 | 4 | Homo sapien | -0.0514486 | -0.9623973 | 0.85950009 | -1.927627 | -0.1202871 | 0.90583807 | 0.99999785 | -6.379148 | down |
| KI - WT, cell_origin: iPSC | Q5TA45 | INTS11 | 6 | Homo sapien | 0.04697469 | -0.7871549 | 0.88110431 | -1.2289766 | 0.11994332 | 0.90610609 | 0.99999785 | -6.3791907 | up |
| KI - WT, cell_origin: iPSC | P62857 | RPS28 | 6 | Homo sapien | 0.03481395 | -0.5843329 | 0.65396076 | 6.42714308 | 0.11981156 | 0.90621676 | 0.99999785 | -6.3792047 | up |
| KI - WT, cell_origin: iPSC | Q9Y5Q8 | GTF3C5 | 7 | Homo sapien | 0.06568698 | -1.1043503 | 1.23572427 | -2.239186 | 0.11958766 | 0.90638563 | 0.99999785 | -6.3792341 | up |
| KI - WT, cell_origin: iPSC | P06748 | NPM1 | 13 | Homo sapien | -0.0477595 | -0.8990495 | 0.80353054 | 3.2778393 | -0.1195596 | 0.90641564 | 0.99999785 | -6.3792352 | down |
| KI - WT, cell_origin: iPSC | Q9NSY1 | BMP2K | 11 | Homo sapien | 0.08964334 | -1.5120643 | 1.69135101 | -1.180068 | 0.11927162 | 0.90663989 | 0.99999785 | -6.3792708 | up |
| KI - WT, cell_origin: iPSC | O15164 | TRIM24 | 6 | Homo sapien | -0.0968896 | -1.8283217 | 1.63454239 | -2.7657606 | -0.1192544 | 0.90665333 | 0.99999785 | -6.3792729 | down |
| KI - WT, cell_origin: iPSC | P22234 | PAICS | 17 | Homo sapien | 0.05216463 | -0.8822712 | 0.98660045 | 0.54119851 | 0.11896769 | 0.90687659 | 0.99999785 | -6.3793083 | up |
| KI - WT, cell_origin: iPSC | O00534 | VWA5A | 5 | Homo sapien | -0.0558491 | -1.0575086 | 0.94581041 | -0.1688818 | -0.1188224 | 0.90698973 | 0.99999785 | -6.3793262 | down |
| KI - WT, cell_origin: iPSC | Q14192 | FHL2 | 4 | Homo sapien | -0.0465575 | -0.8840909 | 0.79097586 | -0.2835974 | -0.1184147 | 0.90729969 | 0.99999785 | -6.3793785 | down |
| KI - WT, cell_origin: iPSC | P53621 | COPA | 44 | Homo sapien | 0.04721598 | -0.8059049 | 0.90033683 | 2.07132368 | 0.11791278 | 0.90769334 | 0.99999785 | -6.3794391 | up |
| KI - WT, cell_origin: iPSC | P17858 | PFKL | 16 | Homo sapien | 0.07419906 | -1.2687473 | 1.4171454 | -0.1013465 | 0.1177449 | 0.90782902 | 0.99999785 | -6.3794582 | up |
| KI - WT, cell_origin: iPSC | Q9UJX2 | CDC23 | 5 | Homo sapien | -0.0409274 | -0.7836569 | 0.70180212 | -1.4301781 | -0.1173608 | 0.90811752 | 0.99999785 | -6.3795081 | down |
| KI - WT, cell_origin: iPSC | Q9H7C9 | AAMDC | 4 | Homo sapien | -0.0412364 | -0.7915552 | 0.70908249 | 0.21330612 | -0.1170601 | 0.90835317 | 0.99999785 | -6.3795442 | down |
| KI - WT, cell_origin: iPSC | Q9Y5A9 | YTHDF2 | 12 | Homo sapien | 0.03511091 | -0.6038645 | 0.67408628 | 1.54659787 | 0.11703013 | 0.90837513 | 0.99999785 | -6.3795482 | up |
| KI - WT, cell_origin: iPSC | Q9ULR0 | ISY1 | 4 | Homo sapien | -0.0419485 | -0.8077821 | 0.7238851 | -0.7704462 | -0.1166615 | 0.90866259 | 0.99999785 | -6.3795927 | down |
| KI - WT, cell_origin: iPSC | Q96HQ2 | CDKN2AIPNL | 4 | Homo sapien | 0.04763933 | -0.8254569 | 0.92073561 | -0.2111714 | 0.11624074 | 0.90899484 | 0.99999785 | -6.3796422 | up |
| KI - WT, cell_origin: iPSC | O94979 | SEC31A | 25 | Homo sapien | -0.0404525 | -0.7822916 | 0.70138669 | 0.25387039 | -0.1161463 | 0.90906502 | 0.99999785 | -6.3796546 | down |
| KI - WT, cell_origin: iPSC | O00410 | IPO5 | 35 | Homo sapien | 0.04610551 | -0.7997133 | 0.8919243 | 1.41930146 | 0.11613326 | 0.90907965 | 0.99999785 | -6.3796548 | up |
| KI - WT, cell_origin: iPSC | Q8WUM4 | PDCD6IP | 19 | Homo sapien | 0.03307316 | -0.5749495 | 0.64109583 | -0.2066849 | 0.11584988 | 0.90929475 | 0.99999785 | -6.3796905 | up |
| KI - WT, cell_origin: iPSC | P32929 | CTH | 4 | Homo sapien | -0.0610431 | -1.1873437 | 1.06525749 | -1.5085913 | -0.1154863 | 0.90958646 | 0.99999785 | -6.3797317 | down |
| KI - WT, cell_origin: iPSC | Q86UU0 | BCL9L | 2 | Homo sapien | 0.4626897 | -8.0946488 | 9.02002816 | -6.0240304 | 0.1152268 | 0.90979081 | 0.99999785 | -6.3797621 | up |
| KI - WT, cell_origin: iPSC | O75953 | DNAJB5 | 2 | Homo sapien | 0.06084083 | -1.0642151 | 1.18589679 | -2.9039024 | 0.11517565 | 0.90982015 | 0.99999785 | -6.3797711 | up |
| KI - WT, cell_origin: iPSC | O95239 | KIF4A | 5 | Homo sapien | 0.06222133 | -1.0887655 | 1.21320815 | -2.1918053 | 0.11515312 | 0.90984039 | 0.99999785 | -6.3797731 | up |
| KI - WT, cell_origin: iPSC | Q9H3P2 | NELFA | 5 | Homo sapien | 0.07706751 | -1.3509984 | 1.50513346 | -2.0516784 | 0.11500731 | 0.90996184 | 0.99999785 | -6.3797883 | up |
| KI - WT, cell_origin: iPSC | Q8TD55 | PLEKHO2 | 10 | Homo sapien | -0.047296 | -0.9236747 | 0.82908258 | -0.0163712 | -0.1149778 | 0.90997996 | 0.99999785 | -6.3797931 | down |
| KI - WT, cell_origin: iPSC | P61254 | RPL26 | 3 | Homo sapien | 0.04664052 | -0.8185902 | 0.91187122 | 2.11153611 | 0.11485461 | 0.91007742 | 0.99999785 | -6.3798074 | up |
| KI - WT, cell_origin: iPSC | O95819 | MAP4K4 | 17 | Homo sapien | 0.04898206 | -0.8601418 | 0.95810596 | 0.58614583 | 0.11481127 | 0.91011333 | 0.99999785 | -6.379812 | up |
| KI - WT, cell_origin: iPSC | Q99471 | PFDN5 | 4 | Homo sapien | 0.05778776 | -1.015939 | 1.13151446 | -0.6942644 | 0.11469499 | 0.91020521 | 0.99999785 | -6.3798254 | up |
| KI - WT, cell_origin: iPSC | O43143 | DHX15 | 39 | Homo sapien | -0.0585881 | -1.1512518 | 1.03407562 | 2.02529687 | -0.1142681 | 0.91053783 | 0.99999785 | -6.379876 | down |
| KI - WT, cell_origin: iPSC | P55145 | MANF | 8 | Homo sapien | -0.198745 | -3.9133951 | 3.51590504 | -3.4190621 | -0.1140199 | 0.91073128 | 0.99999785 | -6.3799054 | down |
| KI - WT, cell_origin: iPSC | P36969 | GPX4 | 11 | Homo sapien | 0.03433141 | -0.6081756 | 0.67683845 | 1.55464619 | 0.11380291 | 0.91089001 | 0.99999785 | -6.3799339 | up |
| KI - WT, cell_origin: iPSC | O43312 | MTSS1 | 2 | Homo sapien | -0.0779418 | -1.5381014 | 1.38221777 | -3.2038172 | -0.1137014 | 0.91097131 | 0.99999785 | -6.3799452 | down |
| KI - WT, cell_origin: iPSC | Q6ZS17 | RIPOR1 | 37 | Homo sapien | 0.08937275 | -1.5883534 | 1.76709893 | 0.33065922 | 0.11352367 | 0.91111802 | 0.99999785 | -6.3799639 | up |
| KI - WT, cell_origin: iPSC | O75122 | CLASP2 | 17 | Homo sapien | 0.05976561 | -1.0649819 | 1.18451312 | -0.0618623 | 0.11323969 | 0.91133935 | 0.99999785 | -6.3799972 | up |
| KI - WT, cell_origin: iPSC | Q9BY77 | POLDIP3 | 8 | Homo sapien | -0.0598489 | -1.1867313 | 1.0670334 | -0.4652205 | -0.1131828 | 0.91138371 | 0.99999785 | -6.3800039 | down |
| KI - WT, cell_origin: iPSC | Q13126 | MTAP | 6 | Homo sapien | -0.0620202 | -1.2307287 | 1.10668829 | -0.5079816 | -0.1130913 | 0.91145499 | 0.99999785 | -6.3800146 | down |
| KI - WT, cell_origin: iPSC | Q86TG7 | PEG10 | 2 | Homo sapien | 0.08956579 | -1.6004261 | 1.77955769 | -2.261428 | 0.11294316 | 0.91157046 | 0.99999785 | -6.380032 | up |
| KI - WT, cell_origin: iPSC | P17844 | DDX5 | 41 | Homo sapien | -0.0876166 | -1.7414849 | 1.56625171 | 6.4901006 | -0.1128984 | 0.91160536 | 0.99999785 | -6.3800372 | down |
| KI - WT, cell_origin: iPSC | Q9UJY4 | GGA2 | 2 | Homo sapien | -0.043899 | -0.8726797 | 0.78488176 | -1.0283974 | -0.1128232 | 0.91165539 | 0.99999785 | -6.3800484 | down |
| KI - WT, cell_origin: iPSC | Q7Z2T5 | TRMT1L | 11 | Homo sapien | -0.1119142 | -2.2283204 | 2.0044921 | -1.8121378 | -0.1126908 | 0.91176718 | 0.99999785 | -6.3800615 | down |
| KI - WT, cell_origin: iPSC | Q8WUQ7 | CACTIN | 7 | Homo sapien | -0.0628906 | -1.2569282 | 1.13114699 | -0.6963564 | -0.1122458 | 0.91211403 | 0.99999785 | -6.3801133 | down |

| | | | | | | | | | | | | | |
|---|---|---|---|---|---|---|---|---|---|---|---|---|---|
| KI - WT, cell_origin: iPSC | Q7Z5K2 | WAPL | 5 | Homo sapien | 0.07337914 | -1.3235709 | 1.47032915 | -1.5411604 | 0.11194228 | 0.91235061 | 0.99999785 | -6.3801486 | up |
| KI - WT, cell_origin: iPSC | Q13523 | PRPF4B | 5 | Homo sapien | -0.0839295 | -1.6822432 | 1.51438411 | -2.1680275 | -0.1119065 | 0.91237852 | 0.99999785 | -6.3801527 | down |
| KI - WT, cell_origin: iPSC | Q96ST8 | CEP89 | 2 | Homo sapien | -0.0859346 | -1.7226964 | 1.55082721 | -1.7046856 | -0.1118884 | 0.91239262 | 0.99999785 | -6.3801548 | down |
| KI - WT, cell_origin: iPSC | Q8NAF0 | ZNF579 | 2 | Homo sapien | 0.06950844 | -1.2561306 | 1.39514753 | -2.2696189 | 0.11171956 | 0.91252091 | 0.99999785 | -6.3801753 | up |
| KI - WT, cell_origin: iPSC | P06703 | S100A6 | 2 | Homo sapien | -0.06365 | -1.281108 | 1.15380807 | 0.88467732 | -0.1114157 | 0.91276106 | 0.99999785 | -6.3802095 | down |
| KI - WT, cell_origin: iPSC | Q06210 | GFPT1 | 27 | Homo sapien | -0.0990409 | -1.9987769 | 1.80069508 | 0.82518765 | -0.1111025 | 0.91300529 | 0.99999785 | -6.3802456 | down |
| KI - WT, cell_origin: iPSC | Q92925 | SMARCD2 | 4 | Homo sapien | -0.0600207 | -1.2127395 | 1.09269805 | -1.3843284 | -0.1109636 | 0.91311358 | 0.99999785 | -6.3802616 | down |
| KI - WT, cell_origin: iPSC | O43776 | NARS1 | 7 | Homo sapien | -0.0532211 | -1.0762119 | 0.96976967 | -0.7629959 | -0.1108701 | 0.91318642 | 0.99999785 | -6.3802723 | down |
| KI - WT, cell_origin: iPSC | Q13315 | ATM | 58 | Homo sapien | -0.0574197 | -1.1632807 | 1.04844141 | 0.30686127 | -0.1106528 | 0.91335583 | 0.99999785 | -6.3802973 | down |
| KI - WT, cell_origin: iPSC | Q16831 | UPP1 | 13 | Homo sapien | -0.0598169 | -1.2128302 | 1.09319632 | -0.2237941 | -0.1105585 | 0.91342934 | 0.99999785 | -6.3803081 | down |
| KI - WT, cell_origin: iPSC | Q96DI7 | SNRNP40 | 11 | Homo sapien | -0.0457408 | -0.9278966 | 0.83641499 | 0.89603692 | -0.1104835 | 0.91348538 | 0.99999785 | -6.3803173 | down |
| KI - WT, cell_origin: iPSC | P17480 | UBTF | 11 | Homo sapien | -0.0382795 | -0.7766185 | 0.7000594 | 0.65190175 | -0.1104319 | 0.91351969 | 0.99999785 | -6.3803248 | down |
| KI - WT, cell_origin: iPSC | Q8IXM2 | BAP18 | 2 | Homo sapien | 0.03505792 | -0.6411396 | 0.71125541 | -0.1056709 | 0.11042114 | 0.91352638 | 0.99999785 | -6.3803265 | up |
| KI - WT, cell_origin: iPSC | Q96GD4 | AURKB | 10 | Homo sapien | -0.0268392 | -0.5461538 | 0.49247539 | -0.2422551 | -0.1100724 | 0.9137983 | 0.99999785 | -6.3803663 | down |
| KI - WT, cell_origin: iPSC | P41743 | PRKCI | 5 | Homo sapien | -0.0666073 | -1.3577269 | 1.22451232 | -1.8934262 | -0.1099405 | 0.91391121 | 0.99999785 | -6.3803787 | down |
| KI - WT, cell_origin: iPSC | P62910 | RPL32 | 8 | Homo sapien | -0.0280377 | -0.5718575 | 0.51578215 | 2.78471307 | -0.1098062 | 0.91400592 | 0.99999785 | -6.3803966 | down |
| KI - WT, cell_origin: iPSC | P31689 | DNAJA1 | 16 | Homo sapien | -0.0332387 | -0.6828816 | 0.61640429 | 1.61283681 | -0.1089704 | 0.91465767 | 0.99999785 | -6.3804913 | down |
| KI - WT, cell_origin: iPSC | P28702 | RXRB | 3 | Homo sapien | -0.0545327 | -1.1263581 | 1.01729265 | -2.0889593 | -0.108372 | 0.91512605 | 0.99999785 | -6.3805582 | down |
| KI - WT, cell_origin: iPSC | P30260 | CDC27 | 3 | Homo sapien | 0.04282748 | -0.7997675 | 0.88542241 | -1.617933 | 0.10825384 | 0.91521653 | 0.99999785 | -6.3805719 | up |
| KI - WT, cell_origin: iPSC | Q86U42 | PABPN1 | 5 | Homo sapien | -0.0399651 | -0.8272062 | 0.74727603 | -0.4064245 | -0.1081356 | 0.91531084 | 0.99999785 | -6.3805846 | down |
| KI - WT, cell_origin: iPSC | P12236 | SLC25A6 | 6 | Homo sapien | 0.1538358 | -2.8798317 | 3.18750329 | -1.751478 | 0.10806668 | 0.91537236 | 0.99999785 | -6.3805903 | up |
| KI - WT, cell_origin: iPSC | P56545 | CTBP2 | 14 | Homo sapien | -0.0729202 | -1.5115064 | 1.36566592 | 0.87549726 | -0.1080226 | 0.91540675 | 0.99999785 | -6.3805952 | down |
| KI - WT, cell_origin: iPSC | P62805 | H4C1 | 6 | Homo sapien | 0.04948497 | -0.9316091 | 1.03057902 | 4.06291398 | 0.10748924 | 0.91582269 | 0.99999785 | -6.3806548 | up |
| KI - WT, cell_origin: iPSC | Q9H269 | VPS16 | 15 | Homo sapien | -0.0307725 | -0.6431962 | 0.58165126 | 0.01693824 | -0.1070163 | 0.91618181 | 0.99999785 | -6.3807099 | down |
| KI - WT, cell_origin: iPSC | Q9Y5B6 | PAXBP1 | 9 | Homo sapien | 0.06802741 | -1.2896077 | 1.42566248 | -1.0729736 | 0.10678328 | 0.9163733 | 0.99999785 | -6.3807331 | up |
| KI - WT, cell_origin: iPSC | Q96KN1 | LRATD2 | 4 | Homo sapien | -0.0988441 | -2.0785414 | 1.88085316 | -1.1225643 | -0.1064031 | 0.91666983 | 0.99999785 | -6.3807751 | down |
| KI - WT, cell_origin: iPSC | Q9NV31 | IMP3 | 5 | Homo sapien | 0.03374352 | -0.6422138 | 0.70970083 | -0.8932756 | 0.10631898 | 0.91672576 | 0.99999785 | -6.3807869 | up |
| KI - WT, cell_origin: iPSC | O14617 | AP3D1 | 21 | Homo sapien | 0.04279448 | -0.8156128 | 0.90120179 | -0.3830415 | 0.106197 | 0.91682381 | 0.99999785 | -6.3807996 | up |
| KI - WT, cell_origin: iPSC | Q7LBC6 | KDM3B | 13 | Homo sapien | 0.05718025 | -1.0912743 | 1.20563485 | -1.5067942 | 0.10609865 | 0.91690641 | 0.99999785 | -6.3808089 | up |
| KI - WT, cell_origin: iPSC | O94762 | RECQL5 | 6 | Homo sapien | 0.07228832 | -1.3805953 | 1.52517191 | 1.14997755 | 0.10603266 | 0.9169588 | 0.99999785 | -6.3808159 | up |
| KI - WT, cell_origin: iPSC | Q9UN37 | VPS4A | 4 | Homo sapien | -0.0464799 | -0.9820592 | 0.88909941 | -0.5836078 | -0.1058563 | 0.91709376 | 0.99999785 | -6.3808359 | down |
| KI - WT, cell_origin: iPSC | P0DOX2 | | 3 | Homo sapien | -0.0506097 | -1.0706485 | 0.96942903 | 6.9121621 | -0.1057352 | 0.91719081 | 0.99999785 | -6.3808485 | down |
| KI - WT, cell_origin: iPSC | Q5HYW2 | NHSL2 | 8 | Homo sapien | 0.0276162 | -0.5309307 | 0.58616308 | -0.1961575 | 0.10530377 | 0.91751778 | 0.99999785 | -6.3808982 | up |
| KI - WT, cell_origin: iPSC | Q92613 | JADE3 | 2 | Homo sapien | 0.07689454 | -1.4813205 | 1.63510953 | -1.3675007 | 0.10516482 | 0.91763578 | 0.99999785 | -6.3809108 | up |
| KI - WT, cell_origin: iPSC | Q12834 | CDC20 | 10 | Homo sapien | -0.0737525 | -1.5704603 | 1.42295521 | -2.2001648 | -0.1050128 | 0.91775437 | 0.99999785 | -6.3809274 | down |
| KI - WT, cell_origin: iPSC | Q7Z333 | SETX | 3 | Homo sapien | -0.0720852 | -1.5360488 | 1.39187844 | -2.7619367 | -0.1049051 | 0.91783395 | 0.99999785 | -6.3809402 | down |
| KI - WT, cell_origin: iPSC | O75175 | CNOT3 | 11 | Homo sapien | -0.0333669 | -0.7109499 | 0.64421613 | 0.5212715 | -0.10488 | 0.91784839 | 0.99999785 | -6.3809443 | down |
| KI - WT, cell_origin: iPSC | Q9Y696 | CLIC4 | 3 | Homo sapien | -0.0616303 | -1.3147471 | 1.19148654 | -2.0049181 | -0.1047952 | 0.91792181 | 0.99999785 | -6.3809516 | down |
| KI - WT, cell_origin: iPSC | Q9NUW8 | TDP1 | 5 | Homo sapien | -0.0637256 | -1.3604193 | 1.23296801 | -1.8654541 | -0.1047319 | 0.91797349 | 0.99999785 | -6.3809579 | down |
| KI - WT, cell_origin: iPSC | Q8NCW5 | NAXE | 14 | Homo sapien | -0.0654306 | -1.3972887 | 1.26642757 | 1.78968508 | -0.1046948 | 0.91800245 | 0.99999785 | -6.3809619 | down |
| KI - WT, cell_origin: iPSC | Q9NXV6 | CDKN2AIP | 7 | Homo sapien | -0.0687532 | -1.4692817 | 1.33177537 | -1.3195532 | -0.1046172 | 0.91806301 | 0.99999785 | -6.3809703 | down |
| KI - WT, cell_origin: iPSC | Q9NTZ6 | RBM12 | 9 | Homo sapien | -0.034846 | -0.7453461 | 0.67565414 | -0.664192 | -0.1044547 | 0.91818024 | 0.99999785 | -6.3809904 | down |
| KI - WT, cell_origin: iPSC | Q96AC1 | FERMT2 | 2 | Homo sapien | 0.08437975 | -1.6392412 | 1.80800068 | -2.3163628 | 0.10432752 | 0.91828901 | 0.99999785 | -6.3810017 | up |
| KI - WT, cell_origin: iPSC | Q9HBI1 | PARVB | 4 | Homo sapien | 0.07690921 | -1.4970549 | 1.65087337 | -1.8030327 | 0.1041324 | 0.91844124 | 0.99999785 | -6.3810228 | up |
| KI - WT, cell_origin: iPSC | P08708 | RPS17 | 9 | Homo sapien | -0.0195535 | -0.419878 | 0.38077102 | 4.79651237 | -0.1040283 | 0.91851298 | 0.99999785 | -6.3810364 | down |
| KI - WT, cell_origin: iPSC | Q86XI2 | NCAPG2 | 3 | Homo sapien | 0.03927632 | -0.7671296 | 0.84568221 | -1.3628244 | 0.10373297 | 0.91874343 | 0.99999785 | -6.3810682 | up |

| | | | | | | | | | | | | | |
|---|---|---|---|---|---|---|---|---|---|---|---|---|---|
| KI - WT, cell_origin: iPSC | Q9NZL9 | MAT2B | 7 | Homo sapien | -0.0502168 | -1.08675 | 0.98631634 | 0.09679984 | -0.1032449 | 0.91913372 | 0.99999785 | -6.3811181 | down |
| KI - WT, cell_origin: iPSC | Q9P1Y5 | CAMSAP3 | 8 | Homo sapien | 0.02146228 | -0.4240045 | 0.46692904 | -0.319449 | 0.10261249 | 0.91961783 | 0.99999785 | -6.3811879 | up |
| KI - WT, cell_origin: iPSC | O95810 | CAVIN2 | 3 | Homo sapien | 0.06746772 | -1.3406494 | 1.47558482 | -1.8995668 | 0.10210797 | 0.92002089 | 0.99999785 | -6.3812391 | up |
| KI - WT, cell_origin: iPSC | P23396 | RPS3 | 24 | Homo sapien | 0.0170492 | -0.3393784 | 0.37347676 | 5.99293732 | 0.10187615 | 0.92019252 | 0.99999785 | -6.3812659 | up |
| KI - WT, cell_origin: iPSC | Q5UIP0 | RIF1 | 17 | Homo sapien | 0.07379449 | -1.4724082 | 1.61999722 | -2.2412922 | 0.10170911 | 0.92033216 | 0.99999785 | -6.3812812 | up |
| KI - WT, cell_origin: iPSC | Q3ZCX4 | ZNF568 | 2 | Homo sapien | 0.13641301 | -2.7313743 | 3.00420029 | -3.9212844 | 0.10137042 | 0.92059649 | 0.99999785 | -6.3813168 | up |
| KI - WT, cell_origin: iPSC | P78371 | CCT2 | 24 | Homo sapien | -0.0575087 | -1.268212 | 1.15319467 | 1.13544522 | -0.1012274 | 0.92070811 | 0.99999785 | -6.3813318 | down |
| KI - WT, cell_origin: iPSC | O15258 | RER1 | 7 | Homo sapien | 0.09808151 | -1.9712854 | 2.16744841 | 0.61449674 | 0.10100713 | 0.92088002 | 0.99999785 | -6.3813549 | up |
| KI - WT, cell_origin: iPSC | Q96BW1 | UPRT | 2 | Homo sapien | 0.04538118 | -0.9129984 | 1.00376076 | -0.8421103 | 0.10089516 | 0.92096495 | 0.99999785 | -6.3813672 | up |
| KI - WT, cell_origin: iPSC | Q86WP2 | GPBP1 | 4 | Homo sapien | -0.0870494 | -1.9279246 | 1.75382583 | -2.9630953 | -0.1007729 | 0.92106285 | 0.99999785 | -6.3813794 | down |
| KI - WT, cell_origin: iPSC | Q9BSJ6 | PIMREG | 4 | Homo sapien | 0.09213618 | -1.8572058 | 2.04147813 | -2.9088608 | 0.10072668 | 0.9210989 | 0.99999785 | -6.3813842 | up |
| KI - WT, cell_origin: iPSC | Q96SI9 | STRBP | 9 | Homo sapien | -0.1181768 | -2.6242108 | 2.38785711 | -2.2221799 | -0.1004958 | 0.92127913 | 0.99999785 | -6.3814083 | down |
| KI - WT, cell_origin: iPSC | Q13085 | ACACA | 63 | Homo sapien | 0.05864721 | -1.1861914 | 1.30348585 | 0.66293677 | 0.10040069 | 0.92135335 | 0.99999785 | -6.3814182 | up |
| KI - WT, cell_origin: iPSC | Q9H6R7 | WDCP | 10 | Homo sapien | -0.0426314 | -0.9471973 | 0.86193455 | -1.0754244 | -0.1003939 | 0.92135224 | 0.99999785 | -6.3814205 | down |
| KI - WT, cell_origin: iPSC | P25788 | PSMA3 | 5 | Homo sapien | 0.06220162 | -1.2582582 | 1.38266144 | -0.5897495 | 0.10038732 | 0.92136378 | 0.99999785 | -6.3814196 | up |
| KI - WT, cell_origin: iPSC | Q9BTD8 | RBM42 | 5 | Homo sapien | -0.0304451 | -0.6773518 | 0.61646156 | -1.0331201 | -0.1002341 | 0.9214742 | 0.99999785 | -6.3814378 | down |
| KI - WT, cell_origin: iPSC | Q9Y4K4 | MAP4K5 | 3 | Homo sapien | 0.06616585 | -1.3497943 | 1.48212603 | -1.8640328 | 0.099583 | 0.9219916 | 0.99999785 | -6.3815029 | up |
| KI - WT, cell_origin: iPSC | Q8TDR0 | TRAF3IP1 | 6 | Homo sapien | 0.03628093 | -0.7398851 | 0.81244692 | -1.0652867 | 0.09955508 | 0.92200432 | 0.99999785 | -6.381508 | up |
| KI - WT, cell_origin: iPSC | Q8IX04 | UEVLD | 10 | Homo sapien | -0.0394174 | -0.8837919 | 0.80495709 | -0.8887528 | -0.0994397 | 0.92209675 | 0.99999785 | -6.3815193 | down |
| KI - WT, cell_origin: iPSC | P26358 | DNMT1 | 24 | Homo sapien | 0.08691398 | -1.778375 | 1.95220292 | -0.7668504 | 0.09929923 | 0.92221311 | 0.99999785 | -6.3815322 | up |
| KI - WT, cell_origin: iPSC | Q14194 | CRMP1 | 10 | Homo sapien | 0.03587866 | -0.7338653 | 0.80562263 | -0.463609 | 0.09927978 | 0.92222032 | 0.99999785 | -6.3815361 | up |
| KI - WT, cell_origin: iPSC | Q09666 | AHNAK | 235 | Homo sapien | 0.0967808 | -1.9811656 | 2.17472716 | 1.50861214 | 0.09925611 | 0.92224678 | 0.99999785 | -6.3815366 | up |
| KI - WT, cell_origin: iPSC | Q9H7E9 | C8orf33 | 2 | Homo sapien | -0.0420335 | -0.9446353 | 0.86056833 | -0.5665862 | -0.099216 | 0.92227395 | 0.99999785 | -6.3815417 | down |
| KI - WT, cell_origin: iPSC | Q96FZ7 | CHMP6 | 8 | Homo sapien | 0.05019521 | -1.0340451 | 1.13443555 | 2.88863852 | 0.09865955 | 0.92271248 | 0.99999785 | -6.3815978 | up |
| KI - WT, cell_origin: iPSC | P68371 | TUBB4B | 23 | Homo sapien | -0.020253 | -0.4576586 | 0.41715257 | 6.24089051 | -0.0986155 | 0.92273791 | 0.99999785 | -6.3816045 | down |
| KI - WT, cell_origin: iPSC | Q8WWY3 | PRPF31 | 11 | Homo sapien | -0.0575743 | -1.3022759 | 1.18712727 | -1.3511404 | -0.0985748 | 0.92277866 | 0.99999785 | -6.3816064 | down |
| KI - WT, cell_origin: iPSC | P35250 | RFC2 | 13 | Homo sapien | -0.0488418 | -1.1055227 | 1.00783911 | 0.27192491 | -0.0985032 | 0.92283455 | 0.99999785 | -6.3816137 | down |
| KI - WT, cell_origin: iPSC | Q92769 | HDAC2 | 5 | Homo sapien | 0.05728406 | -1.1840233 | 1.29859144 | -1.3740941 | 0.09834603 | 0.92295723 | 0.99999785 | -6.3816298 | up |
| KI - WT, cell_origin: iPSC | O60264 | SMARCA5 | 27 | Homo sapien | -0.0729384 | -1.6560068 | 1.51012999 | -1.1657463 | -0.0981881 | 0.92308051 | 0.99999785 | -6.3816458 | down |
| KI - WT, cell_origin: iPSC | P63104 | YWHAZ | 10 | Homo sapien | -0.0247249 | -0.5619616 | 0.51251187 | 3.46221558 | -0.0980185 | 0.92320398 | 0.99999785 | -6.3816652 | down |
| KI - WT, cell_origin: iPSC | Q9ULM3 | YEATS2 | 2 | Homo sapien | 0.18193496 | -3.7783918 | 4.14226169 | -2.5292385 | 0.09790109 | 0.92330461 | 0.99999785 | -6.381675 | up |
| KI - WT, cell_origin: iPSC | O60256 | PRPSAP2 | 10 | Homo sapien | 0.04633825 | -0.9637038 | 1.05638033 | 0.52275665 | 0.09776931 | 0.92340749 | 0.99999785 | -6.3816884 | up |
| KI - WT, cell_origin: iPSC | P50914 | RPL14 | 6 | Homo sapien | -0.0226605 | -0.5167045 | 0.47138344 | 3.94548495 | -0.0976887 | 0.92346157 | 0.99999785 | -6.3816987 | down |
| KI - WT, cell_origin: iPSC | Q86U90 | YRDC | 4 | Homo sapien | -0.0485383 | -1.1107465 | 1.01366996 | -0.5756546 | -0.0973816 | 0.92371018 | 0.99999785 | -6.3817276 | down |
| KI - WT, cell_origin: iPSC | Q12923 | PTPN13 | 27 | Homo sapien | 0.08318008 | -1.7372993 | 1.90365945 | -2.1677294 | 0.09737241 | 0.92371737 | 0.99999785 | -6.3817285 | up |
| KI - WT, cell_origin: iPSC | Q86YS7 | C2CD5 | 12 | Homo sapien | 0.03869777 | -0.8097488 | 0.88714435 | -0.7748078 | 0.09715818 | 0.92387843 | 0.99999785 | -6.3817516 | up |
| KI - WT, cell_origin: iPSC | P28370 | SMARCA1 | 7 | Homo sapien | 0.05658322 | -1.187141 | 1.30030746 | -0.8173209 | 0.09695405 | 0.92404403 | 0.99999785 | -6.3817706 | up |
| KI - WT, cell_origin: iPSC | P20073 | ANXA7 | 5 | Homo sapien | -0.0447137 | -1.03014 | 0.94071248 | -1.5520562 | -0.0966549 | 0.92427108 | 0.99999785 | -6.3818022 | down |
| KI - WT, cell_origin: iPSC | Q9NVE7 | PANK4 | 3 | Homo sapien | -0.0806367 | -1.8683015 | 1.70702808 | -3.3333544 | -0.0961138 | 0.92469801 | 0.99999785 | -6.3818552 | down |
| KI - WT, cell_origin: iPSC | Q6UXN9 | WDR82 | 8 | Homo sapien | 0.05615267 | -1.1897008 | 1.30200618 | 0.2495606 | 0.09605186 | 0.9247485 | 0.99999785 | -6.3818608 | up |
| KI - WT, cell_origin: iPSC | Q13888;Q6P | GTF2H2;GTF2 | 3 | Homo sapien | 0.06104393 | -1.3006702 | 1.4227581 | -2.1636534 | 0.0955342 | 0.92515274 | 0.99999785 | -6.3819122 | up |
| KI - WT, cell_origin: iPSC | Q99459 | CDC5L | 22 | Homo sapien | -0.0401589 | -0.9382783 | 0.85796055 | 0.9772594 | -0.0952835 | 0.92534744 | 0.99999785 | -6.3819373 | down |
| KI - WT, cell_origin: iPSC | Q9BYG5 | PARD6B | 5 | Homo sapien | -0.0943866 | -2.2097378 | 2.02096468 | -0.1843636 | -0.0950889 | 0.92550045 | 0.99999785 | -6.3819562 | down |
| KI - WT, cell_origin: iPSC | Q9Y265 | RUVBL1 | 17 | Homo sapien | -0.0298715 | -0.699013 | 0.63927001 | 1.66856478 | -0.0950776 | 0.92550063 | 0.99999785 | -6.3819593 | down |
| KI - WT, cell_origin: iPSC | Q12933 | TRAF2 | 21 | Homo sapien | 0.03737334 | -0.8005539 | 0.87530058 | 0.30685307 | 0.09501091 | 0.92555532 | 0.99999785 | -6.3819653 | up |
| KI - WT, cell_origin: iPSC | Q9H3U1 | UNC45A | 10 | Homo sapien | 0.02470995 | -0.5329625 | 0.58238237 | -0.7129859 | 0.09436968 | 0.9260536 | 0.99999785 | -6.3820288 | up |

| | | | | | | | | | | | | | |
|---|---|---|---|---|---|---|---|---|---|---|---|---|---|
| KI - WT, cell_origin: iPSC | Q9UII4 | HERC5 | 10 | Homo sapien | -0.0480474 | -1.1325139 | 1.03641921 | -2.5332387 | -0.0943611 | 0.92606028 | 0.99999785 | -6.3820296 | down |
| KI - WT, cell_origin: iPSC | P31946 | YWHAB | 12 | Homo sapien | -0.0219955 | -0.5189491 | 0.47495816 | 4.08423249 | -0.0942664 | 0.92613427 | 0.99999785 | -6.3820389 | down |
| KI - WT, cell_origin: iPSC | P35580 | MYH10 | 30 | Homo sapien | -0.0296505 | -0.700695 | 0.64139399 | 0.20916255 | -0.0941067 | 0.92625903 | 0.99999785 | -6.3820545 | down |
| KI - WT, cell_origin: iPSC | P19338 | NCL | 30 | Homo sapien | -0.0274941 | -0.6530472 | 0.59805909 | 4.23029292 | -0.0936204 | 0.92664073 | 0.99999785 | -6.3821013 | down |
| KI - WT, cell_origin: iPSC | Q9NVX7 | KBTBD4 | 8 | Homo sapien | 0.07066599 | -1.553667 | 1.69499902 | -0.8877382 | 0.09271238 | 0.92735668 | 0.99999785 | -6.3821875 | up |
| KI - WT, cell_origin: iPSC | Q69YN4 | VIRMA | 7 | Homo sapien | 0.06521251 | -1.4342876 | 1.56471262 | -1.7830052 | 0.09268016 | 0.92738185 | 0.99999785 | -6.3821906 | up |
| KI - WT, cell_origin: iPSC | P62851 | RPS25 | 6 | Homo sapien | 0.0325552 | -0.7191337 | 0.78424405 | 6.40127736 | 0.09229637 | 0.92768165 | 0.99999785 | -6.3822273 | up |
| KI - WT, cell_origin: iPSC | P20042 | EIF2S2 | 14 | Homo sapien | 0.02367901 | -0.5242791 | 0.5716371 | 1.72636126 | 0.09203562 | 0.92787696 | 0.99999785 | -6.3822541 | up |
| KI - WT, cell_origin: iPSC | Q96T58 | SPEN | 8 | Homo sapien | 0.05682378 | -1.2588391 | 1.37248663 | -2.1447104 | 0.09202896 | 0.92788852 | 0.99999785 | -6.3822533 | up |
| KI - WT, cell_origin: iPSC | O43660 | PLRG1 | 2 | Homo sapien | 0.03313316 | -0.7339123 | 0.80017858 | -0.5105728 | 0.09200506 | 0.9279018 | 0.99999785 | -6.3822568 | up |
| KI - WT, cell_origin: iPSC | O95400 | CD2BP2 | 7 | Homo sapien | -0.0404929 | -0.9802587 | 0.89927288 | -0.773028 | -0.0918071 | 0.92806115 | 0.99999785 | -6.3822746 | down |
| KI - WT, cell_origin: iPSC | P53602 | MVD | 5 | Homo sapien | -0.0639345 | -1.5483489 | 1.42047983 | -0.918566 | -0.0917873 | 0.92807931 | 0.99999785 | -6.3822759 | down |
| KI - WT, cell_origin: iPSC | O75525 | KHDRBS3 | 2 | Homo sapien | -0.0406041 | -0.9842901 | 0.90308191 | -1.5906031 | -0.0916487 | 0.92818068 | 0.99999785 | -6.3822906 | down |
| KI - WT, cell_origin: iPSC | Q9Y2R4 | DDX52 | 9 | Homo sapien | 0.05679821 | -1.2706055 | 1.3842019 | -2.8183813 | 0.09114384 | 0.92857549 | 0.99999785 | -6.3823383 | up |
| KI - WT, cell_origin: iPSC | Q9Y2Z0 | SUGT1 | 6 | Homo sapien | 0.0360154 | -0.8059497 | 0.87798054 | -0.5623969 | 0.09111964 | 0.92859511 | 0.99999785 | -6.3823405 | up |
| KI - WT, cell_origin: iPSC | Q15650 | TRIP4 | 14 | Homo sapien | -0.0276802 | -0.6748733 | 0.61951282 | 1.09421752 | -0.091091 | 0.928615 | 0.99999785 | -6.3823437 | down |
| KI - WT, cell_origin: iPSC | P42285 | MTREX | 19 | Homo sapien | -0.0476516 | -1.162769 | 1.06746579 | -1.0487651 | -0.0910667 | 0.92864231 | 0.99999785 | -6.3823442 | down |
| KI - WT, cell_origin: iPSC | Q13409 | DYNC1I2 | 6 | Homo sapien | 0.04452335 | -0.9975967 | 1.08664338 | -0.5190308 | 0.09104843 | 0.92865657 | 0.99999785 | -6.3823459 | up |
| KI - WT, cell_origin: iPSC | P62318 | SNRPD3 | 6 | Homo sapien | -0.0676407 | -1.6509293 | 1.51564788 | 3.08338746 | -0.0910438 | 0.92866018 | 0.99999785 | -6.3823463 | down |
| KI - WT, cell_origin: iPSC | P16401 | H1-5 | 4 | Homo sapien | 0.06146198 | -1.3828808 | 1.50580478 | 3.1287897 | 0.09068566 | 0.92893999 | 0.99999785 | -6.38238 | up |
| KI - WT, cell_origin: iPSC | Q8N3P4 | VPS8 | 5 | Homo sapien | 0.03387989 | -0.7666191 | 0.83437892 | 0.00165687 | 0.09015462 | 0.92934879 | 0.99999785 | -6.3824312 | up |
| KI - WT, cell_origin: iPSC | Q5JTH9 | RRP12 | 30 | Homo sapien | -0.0304689 | -0.7526308 | 0.69169289 | 0.54434373 | -0.0898642 | 0.92957439 | 0.99999785 | -6.3824585 | down |
| KI - WT, cell_origin: iPSC | Q9UPQ9 | TNRC6B | 28 | Homo sapien | -0.0408533 | -1.0108226 | 0.92911602 | 0.64865513 | -0.0897577 | 0.92966507 | 0.99999785 | -6.3824668 | down |
| KI - WT, cell_origin: iPSC | P51911 | CNN1 | 2 | Homo sapien | -0.092691 | -2.298907 | 2.11352497 | -4.1551791 | -0.0895203 | 0.92984834 | 0.99999785 | -6.3824893 | down |
| KI - WT, cell_origin: iPSC | P49840 | GSK3A | 3 | Homo sapien | 0.03026316 | -0.6919849 | 0.75251125 | -0.6111758 | 0.08924165 | 0.93006015 | 0.99999785 | -6.3825165 | up |
| KI - WT, cell_origin: iPSC | P16989 | YBX3 | 9 | Homo sapien | -0.0298067 | -0.7434928 | 0.68387939 | 4.0049191 | -0.0889743 | 0.93027274 | 0.99999785 | -6.3825403 | down |
| KI - WT, cell_origin: iPSC | Q99590 | SCAF11 | 7 | Homo sapien | 0.06944067 | -1.5966427 | 1.73552407 | -2.0144681 | 0.08882179 | 0.93039636 | 0.99999785 | -6.3825534 | up |
| KI - WT, cell_origin: iPSC | Q96IZ0 | PAWR | 5 | Homo sapien | 0.04446324 | -1.0226553 | 1.1115818 | -1.1603393 | 0.08879546 | 0.93041694 | 0.99999785 | -6.3825558 | up |
| KI - WT, cell_origin: iPSC | O15049 | N4BP3 | 11 | Homo sapien | -0.0364766 | -0.9150974 | 0.84214422 | 3.93963037 | -0.0884739 | 0.93066824 | 0.99999785 | -6.3825854 | down |
| KI - WT, cell_origin: iPSC | Q9Y266 | NUDC | 14 | Homo sapien | -0.0401146 | -1.006411 | 0.92618188 | 0.301246 | -0.0884696 | 0.93067157 | 0.99999785 | -6.3825858 | down |
| KI - WT, cell_origin: iPSC | Q6IN85 | PPP4R3A | 8 | Homo sapien | 0.0569411 | -1.3208935 | 1.43477568 | -0.3864124 | 0.08807063 | 0.93098337 | 0.99999785 | -6.3826223 | up |
| KI - WT, cell_origin: iPSC | Q96RT7 | TUBGCP6 | 7 | Homo sapien | -0.0294241 | -0.7463339 | 0.68748576 | -0.8540857 | -0.0874134 | 0.93148901 | 0.99999785 | -6.3826837 | down |
| KI - WT, cell_origin: iPSC | Q9BRP8 | PYM1 | 4 | Homo sapien | 0.02232384 | -0.5233354 | 0.5679831 | 1.12284202 | 0.08713386 | 0.93170755 | 0.99999785 | -6.382709 | up |
| KI - WT, cell_origin: iPSC | Q13395 | TARBP1 | 19 | Homo sapien | 0.03234699 | -0.7588628 | 0.82355678 | 0.5524992 | 0.08708836 | 0.93174547 | 0.99999785 | -6.3827126 | up |
| KI - WT, cell_origin: iPSC | O43592 | XPOT | 17 | Homo sapien | -0.0421591 | -1.0759109 | 0.99159284 | 0.5712504 | -0.0869114 | 0.93188932 | 0.99999785 | -6.3827274 | down |
| KI - WT, cell_origin: iPSC | Q15427 | SF3B4 | 6 | Homo sapien | 0.04404846 | -1.0363269 | 1.12442383 | -0.8470919 | 0.08688772 | 0.93190786 | 0.99999785 | -6.3827295 | up |
| KI - WT, cell_origin: iPSC | Q01813 | PFKP | 24 | Homo sapien | -0.0714163 | -1.8289019 | 1.68606933 | 0.05389956 | -0.086598 | 0.93213429 | 0.99999785 | -6.3827556 | down |
| KI - WT, cell_origin: iPSC | O00170 | AIP | 9 | Homo sapien | 0.03671635 | -0.8670574 | 0.94049008 | 0.30539431 | 0.08656429 | 0.93215875 | 0.99999785 | -6.382759 | up |
| KI - WT, cell_origin: iPSC | P35625 | TIMP3 | 4 | Homo sapien | 0.04670284 | -1.1069391 | 1.20034482 | -1.0123137 | 0.08627293 | 0.93238837 | 0.99999785 | -6.3827847 | up |
| KI - WT, cell_origin: iPSC | Q99683 | MAP3K5 | 21 | Homo sapien | 0.02374304 | -0.5638317 | 0.61131779 | -0.4284478 | 0.08606229 | 0.93254519 | 0.99999785 | -6.3828051 | up |
| KI - WT, cell_origin: iPSC | Q9BWJ5 | SF3B5 | 4 | Homo sapien | -0.101445 | -2.6177287 | 2.41483874 | -0.4177532 | -0.0859158 | 0.93266748 | 0.99999785 | -6.3828165 | down |
| KI - WT, cell_origin: iPSC | Q9H5Z1 | DHX35 | 5 | Homo sapien | 0.04679709 | -1.115826 | 1.20942019 | -1.4037271 | 0.08577925 | 0.93277426 | 0.99999785 | -6.3828287 | up |
| KI - WT, cell_origin: iPSC | Q9Y295 | DRG1 | 16 | Homo sapien | 0.05844544 | -1.3944622 | 1.51135305 | 1.60371029 | 0.08572648 | 0.9328155 | 0.99999785 | -6.3828334 | up |
| KI - WT, cell_origin: iPSC | Q6P2H3 | CEP85 | 7 | Homo sapien | -0.0387079 | -1.0013289 | 0.9239132 | -1.3405214 | -0.0856568 | 0.9328645 | 0.99999785 | -6.3828407 | down |
| KI - WT, cell_origin: iPSC | Q9Y520 | PRRC2C | 45 | Homo sapien | -0.0370856 | -0.9601707 | 0.8859994 | 1.93771151 | -0.0856183 | 0.9329001 | 0.99999785 | -6.382843 | down |
| KI - WT, cell_origin: iPSC | O15355 | PPM1G | 7 | Homo sapien | -0.0318677 | -0.8247646 | 0.76102928 | -0.9759012 | -0.0856011 | 0.93290589 | 0.99999785 | -6.3828461 | down |

| | | | | | | | | | | | | |
|---|---|---|---|---|---|---|---|---|---|---|---|---|
| KI - WT, cell_origin: iPSC | O94927 | HAUS5 | 3 | Homo sapien | 0.08431254 | -2.0186595 | 2.18728457 | -2.820219 | 0.08543996 | 0.93303946 | 0.99999785 | -6.3828588 up |
| KI - WT, cell_origin: iPSC | P53990 | IST1 | 2 | Homo sapien | 0.06592548 | -1.5802897 | 1.71214065 | -0.9244131 | 0.08534322 | 0.93311509 | 0.99999785 | -6.3828674 up |
| KI - WT, cell_origin: iPSC | P13645 | KRT10 | 27 | Homo sapien | 0.05764223 | -1.3827296 | 1.49801405 | 3.90033414 | 0.08528419 | 0.93316123 | 0.99999785 | -6.3828726 up |
| KI - WT, cell_origin: iPSC | O15511 | ARPC5 | 3 | Homo sapien | 0.03931532 | -0.9439943 | 1.02262491 | 0.26398406 | 0.08520674 | 0.93322177 | 0.99999785 | -6.3828794 up |
| KI - WT, cell_origin: iPSC | O75448 | MED24 | 7 | Homo sapien | 0.04112035 | -0.987565 | 1.0698057 | -1.8722155 | 0.08514711 | 0.93326229 | 0.99999785 | -6.382886 up |
| KI - WT, cell_origin: iPSC | P42766 | RPL35 | 3 | Homo sapien | -0.0233434 | -0.6104099 | 0.56372319 | 5.31074118 | -0.0846999 | 0.93361221 | 0.99999785 | -6.3829252 down |
| KI - WT, cell_origin: iPSC | Q86TN4 | TRPT1 | 2 | Homo sapien | 0.04402798 | -1.0663646 | 1.15442059 | -0.2199921 | 0.08449959 | 0.93377457 | 0.99999785 | -6.3829416 up |
| KI - WT, cell_origin: iPSC | Q9Y314 | NOSIP | 8 | Homo sapien | 0.07027899 | -1.7053981 | 1.84595611 | 0.39856839 | 0.08434588 | 0.93389473 | 0.99999785 | -6.3829551 up |
| KI - WT, cell_origin: iPSC | P26641 | EEF1G | 10 | Homo sapien | -0.0392529 | -1.03133 | 0.95282414 | 0.1136351 | -0.0843196 | 0.93391524 | 0.99999785 | -6.3829573 down |
| KI - WT, cell_origin: iPSC | P62841 | RPS15 | 8 | Homo sapien | -0.0205913 | -0.5415962 | 0.50041356 | 4.22228722 | -0.0841749 | 0.93402075 | 0.99999785 | -6.3829716 down |
| KI - WT, cell_origin: iPSC | P62993 | GRB2 | 8 | Homo sapien | 0.02386671 | -0.5805021 | 0.62823557 | 0.70503676 | 0.08410663 | 0.93407412 | 0.99999785 | -6.3829775 up |
| KI - WT, cell_origin: iPSC | Q05048 | CSTF1 | 11 | Homo sapien | -0.0288504 | -0.7598543 | 0.7021535 | 0.04358192 | -0.0840578 | 0.93411249 | 0.99999785 | -6.3829817 down |
| KI - WT, cell_origin: iPSC | Q7L2E3 | DHX30 | 29 | Homo sapien | -0.0250438 | -0.662984 | 0.61289642 | 1.00538604 | -0.0836103 | 0.93446218 | 0.99999785 | -6.3830206 down |
| KI - WT, cell_origin: iPSC | Q96CP6 | GRAMD1A | 4 | Homo sapien | 0.16571074 | -4.0718826 | 4.40330407 | -2.1753235 | 0.08333621 | 0.93468409 | 0.99999785 | -6.3830428 up |
| KI - WT, cell_origin: iPSC | O43847 | NRDC | 5 | Homo sapien | 0.05330827 | -1.3096897 | 1.41630626 | -2.3249696 | 0.08333577 | 0.93468241 | 0.99999785 | -6.3830432 up |
| KI - WT, cell_origin: iPSC | Q6PJI9 | WDR59 | 19 | Homo sapien | 0.02372994 | -0.5860936 | 0.63355346 | 0.51071626 | 0.08287665 | 0.93503586 | 0.99999785 | -6.3830839 up |
| KI - WT, cell_origin: iPSC | P46781 | RPS9 | 14 | Homo sapien | 0.01575473 | -0.3897827 | 0.4212922 | 5.79791903 | 0.08274077 | 0.93514211 | 0.99999785 | -6.3830956 up |
| KI - WT, cell_origin: iPSC | Q9H0G5 | NSRP1 | 10 | Homo sapien | 0.05920467 | -1.4679822 | 1.58639152 | -0.3051689 | 0.08261638 | 0.93524689 | 0.99999785 | -6.3831047 up |
| KI - WT, cell_origin: iPSC | Q96T21 | SECISBP2 | 2 | Homo sapien | -0.0322149 | -0.8667335 | 0.80230379 | -1.3273684 | -0.0822167 | 0.93555196 | 0.99999785 | -6.3831403 down |
| KI - WT, cell_origin: iPSC | Q13303 | KCNAB2 | 3 | Homo sapien | -0.0359089 | -0.9675548 | 0.89573706 | -1.6907145 | -0.0820911 | 0.93565028 | 0.99999785 | -6.383151 down |
| KI - WT, cell_origin: iPSC | P68036 | UBE2L3 | 4 | Homo sapien | -0.0445603 | -1.2038562 | 1.11473563 | -0.4280578 | -0.0819136 | 0.93579637 | 0.99999785 | -6.3831646 down |
| KI - WT, cell_origin: iPSC | Q9NRW3 | APOBEC3C | 4 | Homo sapien | -0.0603081 | -1.6331388 | 1.51252262 | 1.1025945 | -0.0817139 | 0.93595253 | 0.99999785 | -6.3831815 down |
| KI - WT, cell_origin: iPSC | P52907 | CAPZA1 | 11 | Homo sapien | 0.02348265 | -0.5893643 | 0.63632962 | 3.02832544 | 0.08160837 | 0.93602765 | 0.99999785 | -6.383192 up |
| KI - WT, cell_origin: iPSC | Q9H9A7 | RMI1 | 3 | Homo sapien | -0.0351605 | -0.955766 | 0.88544494 | -0.9464223 | -0.0813636 | 0.93622211 | 0.99999785 | -6.383212 down |
| KI - WT, cell_origin: iPSC | O60216 | RAD21 | 8 | Homo sapien | -0.0392179 | -1.0683296 | 0.98989374 | -1.7114321 | -0.0811775 | 0.93636672 | 0.99999785 | -6.3832279 down |
| KI - WT, cell_origin: iPSC | Q68DQ2 | CRYBG3 | 27 | Homo sapien | -0.0390256 | -1.063868 | 0.98581694 | -0.1663641 | -0.0811511 | 0.93639266 | 0.99999785 | -6.383229 down |
| KI - WT, cell_origin: iPSC | P61313 | RPL15 | 11 | Homo sapien | 0.02249926 | -0.5680346 | 0.61303315 | 3.56616891 | 0.08114526 | 0.93638982 | 0.99999785 | -6.383231 up |
| KI - WT, cell_origin: iPSC | Q9UPT5 | EXOC7 | 23 | Homo sapien | 0.07918147 | -2.00488 | 2.16324295 | -0.5821933 | 0.08096837 | 0.93653554 | 0.99999785 | -6.3832444 up |
| KI - WT, cell_origin: iPSC | Q9UKN8 | GTF3C4 | 10 | Homo sapien | 0.14642232 | -3.7374301 | 4.03027471 | 0.39309041 | 0.08034279 | 0.93702474 | 0.99999785 | -6.3832967 up |
| KI - WT, cell_origin: iPSC | Q08945 | SSRP1 | 18 | Homo sapien | -0.04392 | -1.2111761 | 1.12333607 | 0.88956417 | -0.080186 | 0.93714735 | 0.99999785 | -6.3833097 down |
| KI - WT, cell_origin: iPSC | P62269 | RPS18 | 14 | Homo sapien | -0.0138773 | -0.384692 | 0.35693748 | 5.36289228 | -0.0797052 | 0.93751612 | 0.99999785 | -6.383351 down |
| KI - WT, cell_origin: iPSC | Q9H0F7 | ARL6 | 4 | Homo sapien | -0.0617226 | -1.7126917 | 1.58924646 | -2.5218829 | -0.0796724 | 0.93754906 | 0.99999785 | -6.3833523 down |
| KI - WT, cell_origin: iPSC | Q16763 | UBE2S | 4 | Homo sapien | -0.0511153 | -1.421941 | 1.3197103 | -0.8985658 | -0.0794642 | 0.93771188 | 0.99999785 | -6.3833695 down |
| KI - WT, cell_origin: iPSC | Q9BTL3 | RAMAC | 4 | Homo sapien | 0.03322381 | -0.8576327 | 0.92408034 | 0.00126294 | 0.07945984 | 0.93771262 | 0.99999785 | -6.3833703 up |
| KI - WT, cell_origin: iPSC | Q14956 | GPNMB | 2 | Homo sapien | 0.03702379 | -0.9614226 | 1.03547015 | 2.4564549 | 0.07902391 | 0.93805622 | 0.99999785 | -6.3834056 up |
| KI - WT, cell_origin: iPSC | P49642 | PRIM1 | 4 | Homo sapien | -0.1037897 | -2.9083476 | 2.70076821 | -1.8297925 | -0.0788664 | 0.93817939 | 0.99999785 | -6.3834185 down |
| KI - WT, cell_origin: iPSC | Q86UD0 | SAPCD2 | 2 | Homo sapien | -0.0420702 | -1.1790984 | 1.09495798 | -1.9714447 | -0.0788173 | 0.93821278 | 0.99999785 | -6.3834235 down |
| KI - WT, cell_origin: iPSC | P08237 | PFKM | 7 | Homo sapien | 0.05472724 | -1.4348274 | 1.54428184 | 1.21773855 | 0.07829778 | 0.93862415 | 0.99999785 | -6.3834648 up |
| KI - WT, cell_origin: iPSC | P46782 | RPS5 | 10 | Homo sapien | 0.01380993 | -0.3630781 | 0.39069799 | 5.56998528 | 0.07804025 | 0.93881849 | 0.99999785 | -6.383487 up |
| KI - WT, cell_origin: iPSC | P40227 | CCT6A | 15 | Homo sapien | 0.02483463 | -0.6545815 | 0.70425074 | 1.10502885 | 0.07785052 | 0.93896692 | 0.99999785 | -6.3835024 up |
| KI - WT, cell_origin: iPSC | Q96EK7 | FAM120B | 16 | Homo sapien | 0.02031456 | -0.5383504 | 0.57897956 | 0.38554155 | 0.07744539 | 0.93928385 | 0.99999785 | -6.383535 up |
| KI - WT, cell_origin: iPSC | O15027 | SEC16A | 56 | Homo sapien | 0.04017075 | -1.0653461 | 1.14568757 | 3.0728245 | 0.07743672 | 0.93929767 | 0.99999785 | -6.3835343 up |
| KI - WT, cell_origin: iPSC | Q14966 | ZNF638 | 14 | Homo sapien | -0.0383173 | -1.0938439 | 1.01720928 | -1.345422 | -0.0773496 | 0.93936396 | 0.99999785 | -6.3835417 down |
| KI - WT, cell_origin: iPSC | Q9HB71 | CACYBP | 7 | Homo sapien | -0.0402609 | -1.1515597 | 1.07103801 | 1.12453945 | -0.0772066 | 0.93947768 | 0.99999785 | -6.3835527 down |
| KI - WT, cell_origin: iPSC | Q9BRF8 | CPPED1 | 3 | Homo sapien | -0.0295103 | -0.843964 | 0.78494342 | -0.6390777 | -0.0771816 | 0.93949203 | 0.99999785 | -6.3835557 down |
| KI - WT, cell_origin: iPSC | Q12955 | ANK3 | 9 | Homo sapien | -0.0388839 | -1.1238828 | 1.04611493 | -2.4447299 | -0.0763274 | 0.94015853 | 0.99999785 | -6.383624 down |

| | | | | | | | | | | | | | |
|---|---|---|---|---|---|---|---|---|---|---|---|---|---|
| KI - WT, cell_origin: iPSC | Q15437 | SEC23B | 19 | Homo sapien | 0.02989734 | -0.8061892 | 0.86598383 | 0.68855423 | 0.07618277 | 0.94027526 | 0.99999785 | -6.3836348 | up |
| KI - WT, cell_origin: iPSC | P50990 | CCT8 | 28 | Homo sapien | 0.023315 | -0.6288754 | 0.67550538 | 1.41838734 | 0.07613788 | 0.9403068 | 0.99999785 | -6.383639 | up |
| KI - WT, cell_origin: iPSC | Q8TEX9 | IPO4 | 7 | Homo sapien | 0.0238732 | -0.6450465 | 0.69279286 | -0.5629663 | 0.07601099 | 0.94040607 | 0.99999785 | -6.383649 | up |
| KI - WT, cell_origin: iPSC | Q6P9B9 | INTS5 | 8 | Homo sapien | 0.03241507 | -0.8798738 | 0.94470396 | -1.1660377 | 0.07568895 | 0.94066009 | 0.99999785 | -6.3836739 | up |
| KI - WT, cell_origin: iPSC | Q9UJ70 | NAGK | 10 | Homo sapien | -0.0264725 | -0.7721743 | 0.71922926 | -0.2851082 | -0.0756094 | 0.94072046 | 0.99999785 | -6.3836805 | down |
| KI - WT, cell_origin: iPSC | P29353 | SHC1 | 8 | Homo sapien | -0.0309131 | -0.9020114 | 0.84018514 | 0.17692509 | -0.0756063 | 0.94072643 | 0.99999785 | -6.3836801 | down |
| KI - WT, cell_origin: iPSC | P13693 | TPT1 | 3 | Homo sapien | -0.0628257 | -1.8362986 | 1.71064724 | -2.1001994 | -0.0754944 | 0.9408171 | 0.99999785 | -6.3836882 | down |
| KI - WT, cell_origin: iPSC | Q9H814 | PHAX | 5 | Homo sapien | 0.03626083 | -0.9909917 | 1.06351335 | -0.8997862 | 0.07522512 | 0.9410278 | 0.99999785 | -6.3837093 | up |
| KI - WT, cell_origin: iPSC | P05387 | RPLP2 | 5 | Homo sapien | -0.0148136 | -0.4343663 | 0.40473915 | 6.12759211 | -0.0751993 | 0.94104118 | 0.99999785 | -6.3837126 | down |
| KI - WT, cell_origin: iPSC | Q9BWF3 | RBM4 | 17 | Homo sapien | -0.0470288 | -1.3798833 | 1.28582567 | 1.06429951 | -0.0751941 | 0.9410521 | 0.99999785 | -6.3837117 | down |
| KI - WT, cell_origin: iPSC | Q9UBU9 | NXF1 | 18 | Homo sapien | 0.02861216 | -0.7822024 | 0.83942677 | -0.3154919 | 0.07516851 | 0.94106701 | 0.99999785 | -6.3837146 | up |
| KI - WT, cell_origin: iPSC | Q9BRX9 | WDR83 | 4 | Homo sapien | 0.0473948 | -1.300942 | 1.39573157 | -2.2356956 | 0.07489818 | 0.94128195 | 0.99999785 | -6.383735 | up |
| KI - WT, cell_origin: iPSC | Q9H6Z4 | RANBP3 | 8 | Homo sapien | -0.0479751 | -1.4140584 | 1.31810815 | -1.3318766 | -0.0748413 | 0.9413281 | 0.99999785 | -6.3837391 | down |
| KI - WT, cell_origin: iPSC | Q9NVP1 | DDX18 | 19 | Homo sapien | 0.06612908 | -1.820757 | 1.95301517 | -0.8058003 | 0.07468771 | 0.94144826 | 0.99999785 | -6.383751 | up |
| KI - WT, cell_origin: iPSC | P30050 | RPL12 | 8 | Homo sapien | 0.01550303 | -0.4274228 | 0.45842882 | 4.74218412 | 0.07454615 | 0.94155224 | 0.99999785 | -6.3837632 | up |
| KI - WT, cell_origin: iPSC | P52292 | KPNA2 | 14 | Homo sapien | 0.07379321 | -2.0378244 | 2.1854108 | 0.63287481 | 0.0744738 | 0.94161563 | 0.99999785 | -6.3837676 | up |
| KI - WT, cell_origin: iPSC | P29692 | EEF1D | 12 | Homo sapien | -0.0315563 | -0.9357447 | 0.87263209 | 0.89921125 | -0.07437 | 0.941696 | 0.99999785 | -6.3837757 | down |
| KI - WT, cell_origin: iPSC | P07305 | H1-0 | 4 | Homo sapien | 0.06628012 | -1.8355605 | 1.96812078 | -1.2447952 | 0.07426968 | 0.94177534 | 0.99999785 | -6.3837833 | up |
| KI - WT, cell_origin: iPSC | O75153 | CLUH | 8 | Homo sapien | 0.0589561 | -1.6366417 | 1.7545539 | -1.4632077 | 0.07409831 | 0.94190942 | 0.99999785 | -6.3837965 | up |
| KI - WT, cell_origin: iPSC | Q92989 | CLP1 | 4 | Homo sapien | -0.0288023 | -0.8585239 | 0.80091929 | 0.12875442 | -0.0739456 | 0.94202418 | 0.99999785 | -6.3838091 | down |
| KI - WT, cell_origin: iPSC | P10619 | CTSA | 2 | Homo sapien | 0.06142801 | -1.7107674 | 1.8336234 | -1.8751386 | 0.07386815 | 0.94208951 | 0.99999785 | -6.3838141 | up |
| KI - WT, cell_origin: iPSC | Q96S21 | RAB40C | 2 | Homo sapien | 0.02132151 | -0.5936509 | 0.63629391 | -1.2337226 | 0.07384174 | 0.94210346 | 0.99999785 | -6.3838174 | up |
| KI - WT, cell_origin: iPSC | P61201 | COPS2 | 4 | Homo sapien | 0.03955574 | -1.1033447 | 1.18245614 | -2.2814839 | 0.07372195 | 0.94219863 | 0.99999785 | -6.3838263 | up |
| KI - WT, cell_origin: iPSC | P53618 | COPB1 | 32 | Homo sapien | 0.03318321 | -0.9269531 | 0.99331951 | 0.6400752 | 0.07365257 | 0.94225819 | 0.99999785 | -6.3838306 | up |
| KI - WT, cell_origin: iPSC | Q9NX24 | NHP2 | 3 | Homo sapien | 0.03280118 | -0.9173549 | 0.98295725 | -0.1449462 | 0.07356395 | 0.94232672 | 0.99999785 | -6.3838375 | up |
| KI - WT, cell_origin: iPSC | Q9BV57 | ADI1 | 3 | Homo sapien | 0.05074407 | -1.442862 | 1.54435017 | -2.7764484 | 0.07238223 | 0.94324925 | 0.99999785 | -6.3839273 | up |
| KI - WT, cell_origin: iPSC | Q9C035 | TRIM5 | 24 | Homo sapien | -0.0178899 | -0.5447302 | 0.50895031 | 1.78913099 | -0.0723219 | 0.94329288 | 0.99999785 | -6.3839325 | down |
| KI - WT, cell_origin: iPSC | Q9UKD2 | MRTO4 | 10 | Homo sapien | 0.03321843 | -0.9511249 | 1.01756173 | 0.12601616 | 0.07191756 | 0.94361587 | 0.99999785 | -6.3839615 | up |
| KI - WT, cell_origin: iPSC | Q9C0J8 | WDR33 | 11 | Homo sapien | 0.02913706 | -0.8351068 | 0.89338096 | -0.4929038 | 0.071817 | 0.94368998 | 0.99999785 | -6.3839698 | up |
| KI - WT, cell_origin: iPSC | P50991 | CCT4 | 20 | Homo sapien | 0.02413064 | -0.6925375 | 0.74079879 | 1.1759224 | 0.07171539 | 0.94376811 | 0.99999785 | -6.3839776 | up |
| KI - WT, cell_origin: iPSC | Q93100 | PHKB | 17 | Homo sapien | 0.0198045 | -0.5714212 | 0.61103017 | 1.75890244 | 0.07134283 | 0.94405916 | 0.99999785 | -6.3840053 | up |
| KI - WT, cell_origin: iPSC | O15372 | EIF3H | 17 | Homo sapien | -0.0184562 | -0.5773803 | 0.54046788 | 2.58418799 | -0.0703281 | 0.94485343 | 0.99999785 | -6.3840798 | down |
| KI - WT, cell_origin: iPSC | P48444 | ARCN1 | 20 | Homo sapien | 0.03102276 | -0.9096451 | 0.97169063 | 0.87026066 | 0.07028238 | 0.94489559 | 0.99999785 | -6.384082 | up |
| KI - WT, cell_origin: iPSC | P62314 | SNRPD1 | 5 | Homo sapien | -0.0402836 | -1.2699431 | 1.18937591 | 0.97877272 | -0.0698145 | 0.94526176 | 0.99999785 | -6.384116 | down |
| KI - WT, cell_origin: iPSC | P41091 | EIF2S3 | 16 | Homo sapien | 0.02003349 | -0.5925221 | 0.63258914 | 1.62084037 | 0.06965477 | 0.94538049 | 0.99999785 | -6.3841286 | up |
| KI - WT, cell_origin: iPSC | Q8N7H5 | PAF1 | 7 | Homo sapien | -0.0281564 | -0.8899804 | 0.8336675 | -0.6102041 | -0.0695948 | 0.94542929 | 0.99999785 | -6.3841326 | down |
| KI - WT, cell_origin: iPSC | Q9H7B2 | RPF2 | 6 | Homo sapien | -0.0274282 | -0.8694888 | 0.81463229 | -0.6813536 | -0.0693861 | 0.94559268 | 0.99999785 | -6.3841477 | down |
| KI - WT, cell_origin: iPSC | O15075 | DCLK1 | 5 | Homo sapien | -0.0755305 | -2.3968763 | 2.24581531 | -2.809187 | -0.0693402 | 0.94563307 | 0.99999785 | -6.3841502 | down |
| KI - WT, cell_origin: iPSC | Q9H307 | PNN | 19 | Homo sapien | -0.0355806 | -1.1293552 | 1.05819399 | 0.30519855 | -0.0693247 | 0.94564517 | 0.99999785 | -6.3841513 | down |
| KI - WT, cell_origin: iPSC | Q13123 | IK | 12 | Homo sapien | -0.0395827 | -1.2636575 | 1.18449205 | -0.4232335 | -0.0689128 | 0.94596757 | 0.99999785 | -6.3841808 | down |
| KI - WT, cell_origin: iPSC | Q15393 | SF3B3 | 36 | Homo sapien | -0.0237588 | -0.7600866 | 0.71256892 | 1.77566334 | -0.0687271 | 0.9461075 | 0.99999785 | -6.384195 | down |
| KI - WT, cell_origin: iPSC | Q9NWZ5 | UCKL1 | 5 | Homo sapien | -0.0333266 | -1.0681918 | 1.00153863 | -1.606645 | -0.0686001 | 0.94620796 | 0.99999785 | -6.3842038 | down |
| KI - WT, cell_origin: iPSC | P50995 | ANXA11 | 15 | Homo sapien | -0.0423895 | -1.3664236 | 1.28164461 | -0.3243652 | -0.0682278 | 0.94650378 | 0.99999785 | -6.3842295 | down |
| KI - WT, cell_origin: iPSC | Q9NVI1 | FANCI | 5 | Homo sapien | 0.05670613 | -1.7192442 | 1.83265641 | -3.2147367 | 0.06804084 | 0.94664937 | 0.99999785 | -6.3842428 | up |
| KI - WT, cell_origin: iPSC | Q8IY81 | FTSJ3 | 17 | Homo sapien | 0.08195936 | -2.488196 | 2.65211476 | -1.7208222 | 0.06795815 | 0.94671485 | 0.99999785 | -6.3842485 | up |
| KI - WT, cell_origin: iPSC | Q9BXF3 | CECR2 | 16 | Homo sapien | -0.033353 | -1.0802732 | 1.01356726 | -1.2427682 | -0.0678829 | 0.9467723 | 0.99999785 | -6.3842541 | down |

| | | | | | | | | | | | | | |
|---|---|---|---|---|---|---|---|---|---|---|---|---|---|
| KI - WT, cell_origin: iPSC | Q13283 | G3BP1 | 16 | Homo sapien | -0.0393876 | -1.2796237 | 1.20084845 | 4.0928991 | -0.0676796 | 0.9469329 | 0.99999785 | -6.3842681 | down |
| KI - WT, cell_origin: iPSC | Q15046 | KARS1 | 16 | Homo sapien | -0.0204404 | -0.6666444 | 0.62576358 | 0.60158346 | -0.0673689 | 0.94716999 | 0.99999785 | -6.3842909 | down |
| KI - WT, cell_origin: iPSC | P30414 | NKTR | 4 | Homo sapien | -0.0619367 | -2.023239 | 1.89936552 | -3.6320913 | -0.0672888 | 0.94723735 | 0.99999785 | -6.3842957 | down |
| KI - WT, cell_origin: iPSC | Q9UBN7 | HDAC6 | 15 | Homo sapien | 0.02272365 | -0.6971533 | 0.74260059 | 2.6610577 | 0.06723625 | 0.94727488 | 0.99999785 | -6.3843 | up |
| KI - WT, cell_origin: iPSC | Q9Y2K7 | KDM2A | 4 | Homo sapien | 0.02375891 | -0.7321384 | 0.77965622 | -1.3800207 | 0.06694273 | 0.94750367 | 0.99999785 | -6.3843205 | up |
| KI - WT, cell_origin: iPSC | Q9NQG6 | MIEF1 | 2 | Homo sapien | 0.04501529 | -1.388881 | 1.47891153 | -2.5098141 | 0.06689237 | 0.9475476 | 0.99999785 | -6.3843233 | up |
| KI - WT, cell_origin: iPSC | P49458 | SRP9 | 6 | Homo sapien | -0.0212842 | -0.7068396 | 0.66427125 | 2.95574661 | -0.0661275 | 0.94814261 | 0.99999785 | -6.3843766 | down |
| KI - WT, cell_origin: iPSC | Q9NZT2 | OGFR | 12 | Homo sapien | -0.0298159 | -0.9914402 | 0.93180844 | 1.03114704 | -0.0660761 | 0.94818824 | 0.99999785 | -6.3843793 | down |
| KI - WT, cell_origin: iPSC | O75347 | TBCA | 2 | Homo sapien | 0.04453733 | -1.3927839 | 1.48185854 | -1.7828493 | 0.06603477 | 0.94822058 | 0.99999785 | -6.3843821 | up |
| KI - WT, cell_origin: iPSC | Q5JVS0 | HABP4 | 5 | Homo sapien | -0.0319811 | -1.0696861 | 1.00572395 | -3.0703557 | -0.0656385 | 0.94852483 | 0.99999785 | -6.3844101 | down |
| KI - WT, cell_origin: iPSC | P63172 | DYNLT1 | 2 | Homo sapien | 0.02485888 | -0.7833776 | 0.83309533 | -0.845359 | 0.06551291 | 0.9486242 | 0.99999785 | -6.3844185 | up |
| KI - WT, cell_origin: iPSC | Q9NR50 | EIF2B3 | 7 | Homo sapien | 0.01790267 | -0.5650596 | 0.60086493 | -0.5960051 | 0.06540593 | 0.94870697 | 0.99999785 | -6.3844259 | up |
| KI - WT, cell_origin: iPSC | Q8IYD1 | GSPT2 | 9 | Homo sapien | 0.02544796 | -0.8064431 | 0.85733906 | -0.6874869 | 0.06516183 | 0.94889962 | 0.99999785 | -6.3844422 | up |
| KI - WT, cell_origin: iPSC | Q08J23 | NSUN2 | 11 | Homo sapien | 0.04590746 | -1.4564515 | 1.54826645 | 0.07272254 | 0.06511963 | 0.94893707 | 0.99999785 | -6.3844443 | up |
| KI - WT, cell_origin: iPSC | Q9BQ52 | ELAC2 | 2 | Homo sapien | -0.0449204 | -1.5144306 | 1.42458985 | -3.7745547 | -0.0651044 | 0.94894307 | 0.99999785 | -6.3844463 | down |
| KI - WT, cell_origin: iPSC | O14933 | UBE2L6 | 3 | Homo sapien | 0.03669358 | -1.1681848 | 1.241572 | 0.9068815 | 0.06490068 | 0.9491085 | 0.99999785 | -6.3844591 | up |
| KI - WT, cell_origin: iPSC | O75643 | SNRNP200 | 79 | Homo sapien | -0.0480585 | -1.6267101 | 1.53059306 | 0.82394735 | -0.0648764 | 0.94912754 | 0.99999785 | -6.3844607 | down |
| KI - WT, cell_origin: iPSC | P29590 | PML | 9 | Homo sapien | 0.03238245 | -1.0315365 | 1.09630139 | -0.7318883 | 0.06486397 | 0.94913725 | 0.99999785 | -6.3844615 | up |
| KI - WT, cell_origin: iPSC | Q7L9L4;Q9H8 | MOB1B;MOE | 2 | Homo sapien | 0.01740157 | -0.5544303 | 0.58923349 | 1.31715607 | 0.06481265 | 0.94917154 | 0.99999785 | -6.3844659 | up |
| KI - WT, cell_origin: iPSC | Q9BRP1 | PDCD2L | 10 | Homo sapien | -0.036389 | -1.2331575 | 1.16037941 | 1.38623633 | -0.0647982 | 0.94918876 | 0.99999785 | -6.384466 | down |
| KI - WT, cell_origin: iPSC | Q53EL6 | PDCD4 | 21 | Homo sapien | -0.0366574 | -1.2504611 | 1.17714634 | 2.73198131 | -0.0643599 | 0.94953191 | 0.99999785 | -6.3844953 | down |
| KI - WT, cell_origin: iPSC | A0FGR8 | ESYT2 | 9 | Homo sapien | -0.0757008 | -2.5961141 | 2.44471256 | -2.3186375 | -0.0640075 | 0.94980785 | 0.99999785 | -6.3845188 | down |
| KI - WT, cell_origin: iPSC | O43818 | RRP9 | 2 | Homo sapien | 0.03326076 | -1.0804388 | 1.14696036 | -1.9216933 | 0.06361829 | 0.95010856 | 0.99999785 | -6.3845451 | up |
| KI - WT, cell_origin: iPSC | P54132 | BLM | 14 | Homo sapien | -0.0806354 | -2.7857567 | 2.62448589 | -0.1970735 | -0.0635245 | 0.95018604 | 0.99999785 | -6.3845507 | down |
| KI - WT, cell_origin: iPSC | P52434 | POLR2H | 4 | Homo sapien | 0.02496712 | -0.8132244 | 0.86315867 | -0.3879591 | 0.06345166 | 0.95023905 | 0.99999785 | -6.3845561 | up |
| KI - WT, cell_origin: iPSC | Q9UHB6 | LIMA1 | 3 | Homo sapien | 0.04884568 | -1.6008874 | 1.69857879 | -2.1242368 | 0.0630979 | 0.95052012 | 0.99999785 | -6.3845787 | up |
| KI - WT, cell_origin: iPSC | Q06330 | RBPJ | 7 | Homo sapien | 0.05947934 | -1.9504268 | 2.0693855 | -1.5104392 | 0.06306563 | 0.95054538 | 0.99999785 | -6.3845808 | up |
| KI - WT, cell_origin: iPSC | P55769 | SNU13 | 5 | Homo sapien | 0.02614845 | -0.8652157 | 0.91751258 | 1.30269808 | 0.06250716 | 0.95098134 | 0.99999785 | -6.3846173 | up |
| KI - WT, cell_origin: iPSC | O95347 | SMC2 | 18 | Homo sapien | 0.02228227 | -0.7374726 | 0.78203712 | -0.4991889 | 0.0624669 | 0.95100909 | 0.99999785 | -6.3846205 | up |
| KI - WT, cell_origin: iPSC | Q9BVG4 | PBDC1 | 9 | Homo sapien | -0.0219423 | -0.779214 | 0.73532929 | -0.2450348 | -0.0617165 | 0.95159694 | 0.99999785 | -6.3846688 | down |
| KI - WT, cell_origin: iPSC | P54687 | BCAT1 | 5 | Homo sapien | 0.02795474 | -0.9412807 | 0.9971902 | -0.7158185 | 0.06146058 | 0.95180164 | 0.99999785 | -6.3846845 | up |
| KI - WT, cell_origin: iPSC | P04040 | CAT | 4 | Homo sapien | 0.02338136 | -0.7878186 | 0.83458135 | -0.9573758 | 0.06139258 | 0.95185072 | 0.99999785 | -6.3846894 | up |
| KI - WT, cell_origin: iPSC | Q9UDY2 | TJP2 | 50 | Homo sapien | -0.0373868 | -1.3367586 | 1.26198489 | 2.17689912 | -0.061318 | 0.95191402 | 0.99999785 | -6.3846935 | down |
| KI - WT, cell_origin: iPSC | Q96I25 | RBM17 | 14 | Homo sapien | -0.0184933 | -0.6624374 | 0.62545079 | 1.20616549 | -0.0611655 | 0.95202787 | 0.99999785 | -6.384704 | down |
| KI - WT, cell_origin: iPSC | O15294 | OGT | 13 | Homo sapien | 0.01818032 | -0.6162292 | 0.65258985 | 0.37438896 | 0.06103394 | 0.9521309 | 0.99999785 | -6.3847123 | up |
| KI - WT, cell_origin: iPSC | O43172 | PRPF4 | 14 | Homo sapien | 0.02443865 | -0.8291358 | 0.87801312 | -0.6751192 | 0.06098931 | 0.95216751 | 0.99999785 | -6.3847149 | up |
| KI - WT, cell_origin: iPSC | Q9H5Z6 | FAM124B | 12 | Homo sapien | -0.0351643 | -1.2642594 | 1.19393071 | 1.05667992 | -0.0609705 | 0.95218616 | 0.99999785 | -6.3847155 | down |
| KI - WT, cell_origin: iPSC | Q13042 | CDC16 | 3 | Homo sapien | 0.0213058 | -0.723613 | 0.76622457 | -1.6406714 | 0.0609156 | 0.95222359 | 0.99999785 | -6.3847198 | up |
| KI - WT, cell_origin: iPSC | Q9BWW4 | SSBP3 | 3 | Homo sapien | -0.0350348 | -1.2664049 | 1.19633522 | 1.12373045 | -0.0606337 | 0.95244994 | 0.99999785 | -6.3847367 | down |
| KI - WT, cell_origin: iPSC | P35251 | RFC1 | 24 | Homo sapien | 0.02444794 | -0.8374466 | 0.88634248 | -0.6171879 | 0.06042353 | 0.95261065 | 0.99999785 | -6.3847505 | up |
| KI - WT, cell_origin: iPSC | P63272 | SUPT4H1 | 3 | Homo sapien | -0.0179618 | -0.6518035 | 0.6158798 | 0.00867663 | -0.0603545 | 0.95266312 | 0.99999785 | -6.384755 | down |
| KI - WT, cell_origin: iPSC | Q9UHR6 | ZNHIT2 | 2 | Homo sapien | -0.0379641 | -1.3825929 | 1.30666466 | -2.562219 | -0.0601435 | 0.95282997 | 0.99999785 | -6.384768 | down |
| KI - WT, cell_origin: iPSC | P52701 | MSH6 | 16 | Homo sapien | -0.0734381 | -2.6784839 | 2.53160763 | -2.451727 | -0.0600771 | 0.95288588 | 0.99999785 | -6.3847715 | down |
| KI - WT, cell_origin: iPSC | Q9C0H6 | KLHL4 | 18 | Homo sapien | 0.04521468 | -1.5717634 | 1.66219281 | -0.4713799 | 0.0595906 | 0.95326689 | 0.99999785 | -6.3848017 | up |
| KI - WT, cell_origin: iPSC | Q8WXD5 | GEMIN6 | 4 | Homo sapien | -0.0356557 | -1.3139994 | 1.24268806 | -1.6722641 | -0.0594406 | 0.95338434 | 0.99999785 | -6.384811 | down |
| KI - WT, cell_origin: iPSC | O75694 | NUP155 | 19 | Homo sapien | 0.03807572 | -1.3290549 | 1.40520637 | -0.8321746 | 0.0593527 | 0.95345322 | 0.99999785 | -6.3848164 | up |

| | | | | | | | | | | | | | |
|---|---|---|---|---|---|---|---|---|---|---|---|---|---|
| KI - WT, cell_origin: iPSC | Q9UHD1 | CHORDC1 | 5 | Homo sapien | -0.0270094 | -0.9969275 | 0.9429087 | -1.9802748 | -0.0593089 | 0.95348215 | 0.99999785 | -6.3848199 | down |
| KI - WT, cell_origin: iPSC | P17252 | PRKCA | 2 | Homo sapien | 0.06599095 | -2.3118065 | 2.44378836 | -2.4266218 | 0.05914416 | 0.95361656 | 0.99999785 | -6.3848292 | up |
| KI - WT, cell_origin: iPSC | O15013 | ARHGEF10 | 3 | Homo sapien | 0.04428952 | -1.5546876 | 1.64326664 | -3.2083962 | 0.05900335 | 0.95372309 | 0.99999785 | -6.3848384 | up |
| KI - WT, cell_origin: iPSC | Q15459 | SF3A1 | 26 | Homo sapien | -0.0326942 | -1.224066 | 1.15867758 | -0.8161221 | -0.0584825 | 0.95413482 | 0.99999785 | -6.3848695 | down |
| KI - WT, cell_origin: iPSC | Q9Y3T9 | NOC2L | 10 | Homo sapien | 0.02208931 | -0.7831788 | 0.82735746 | -1.1308754 | 0.05842265 | 0.9541764 | 0.99999785 | -6.3848739 | up |
| KI - WT, cell_origin: iPSC | Q29RF7 | PDS5A | 14 | Homo sapien | 0.03557899 | -1.266946 | 1.33810394 | -1.3157744 | 0.05821165 | 0.95434699 | 0.99999785 | -6.3848859 | up |
| KI - WT, cell_origin: iPSC | O75909 | CCNK | 7 | Homo sapien | 0.04991189 | -1.7808499 | 1.88067373 | -0.768007 | 0.05809979 | 0.95443461 | 0.99999785 | -6.3848927 | up |
| KI - WT, cell_origin: iPSC | Q9NXX6 | NSMCE4A | 4 | Homo sapien | 0.02330848 | -0.83355 | 0.88016693 | -1.4448055 | 0.05793548 | 0.95455805 | 0.99999785 | -6.3849033 | up |
| KI - WT, cell_origin: iPSC | Q9NQT4 | EXOSC5 | 4 | Homo sapien | 0.01723273 | -0.6233137 | 0.65777914 | -1.1955321 | 0.05729849 | 0.9550571 | 0.99999785 | -6.3849413 | up |
| KI - WT, cell_origin: iPSC | Q92615 | LARP4B | 11 | Homo sapien | -0.0184091 | -0.7090648 | 0.67224652 | 0.51175821 | -0.056769 | 0.9554719 | 0.99999785 | -6.3849726 | down |
| KI - WT, cell_origin: iPSC | Q9UHB9 | SRP68 | 21 | Homo sapien | 0.03075582 | -1.1309459 | 1.19245748 | 0.98635383 | 0.05642025 | 0.9557503 | 0.99999785 | -6.3849923 | up |
| KI - WT, cell_origin: iPSC | O43314 | PPIP5K2 | 4 | Homo sapien | -0.0858633 | -3.3321483 | 3.16042163 | -3.0096403 | -0.0563668 | 0.95579217 | 0.99999785 | -6.3849955 | down |
| KI - WT, cell_origin: iPSC | P63279 | UBE2I | 3 | Homo sapien | -0.0193375 | -0.7517007 | 0.71302561 | -0.9686171 | -0.056236 | 0.95588955 | 0.99999785 | -6.3850038 | down |
| KI - WT, cell_origin: iPSC | Q8TDN6 | BRIX1 | 8 | Homo sapien | -0.0537188 | -2.0916155 | 1.98417793 | -1.4719354 | -0.0561754 | 0.9559421 | 0.99999785 | -6.3850066 | down |
| KI - WT, cell_origin: iPSC | Q9Y3C1 | NOP16 | 5 | Homo sapien | -0.0253547 | -0.9928797 | 0.94217024 | 0.09429967 | -0.0558469 | 0.95619944 | 0.99999785 | -6.3850257 | down |
| KI - WT, cell_origin: iPSC | Q8TF05 | PPP4R1 | 7 | Homo sapien | 0.03357872 | -1.2607018 | 1.32785921 | -1.6607103 | 0.05528892 | 0.95663661 | 0.99999785 | -6.3850579 | up |
| KI - WT, cell_origin: iPSC | O15514 | POLR2D | 3 | Homo sapien | -0.0198057 | -0.7867397 | 0.7471283 | -1.0924439 | -0.0550011 | 0.95685709 | 0.99999785 | -6.385075 | down |
| KI - WT, cell_origin: iPSC | Q5JSZ5 | PRRC2B | 27 | Homo sapien | 0.02193096 | -0.8301942 | 0.87405616 | 0.26493202 | 0.05482801 | 0.9569948 | 0.99999785 | -6.3850846 | up |
| KI - WT, cell_origin: iPSC | Q96A19 | CCDC102A | 2 | Homo sapien | -0.0245495 | -0.98257 | 0.93347097 | -3.0381111 | -0.0545768 | 0.95718959 | 0.99999785 | -6.3850991 | down |
| KI - WT, cell_origin: iPSC | P58107 | EPPK1 | 65 | Homo sapien | -0.0498836 | -2.0029239 | 1.9031567 | 0.05293135 | -0.0544313 | 0.95730852 | 0.99999785 | -6.3851066 | down |
| KI - WT, cell_origin: iPSC | O60870 | KIN | 10 | Homo sapien | 0.02594303 | -0.9917678 | 1.04365386 | -0.8559622 | 0.05432486 | 0.95739193 | 0.99999785 | -6.3851126 | up |
| KI - WT, cell_origin: iPSC | P04843 | RPN1 | 10 | Homo sapien | 0.05054144 | -1.9374989 | 2.03858179 | -1.6424627 | 0.05417823 | 0.95750682 | 0.99999785 | -6.3851209 | up |
| KI - WT, cell_origin: iPSC | Q9UI95 | MAD2L2 | 2 | Homo sapien | -0.0116608 | -0.4716076 | 0.44828598 | -0.5083228 | -0.053996 | 0.9576447 | 0.99999785 | -6.3851318 | down |
| KI - WT, cell_origin: iPSC | Q9Y2V2 | CARHSP1 | 6 | Homo sapien | 0.03703941 | -1.4295345 | 1.50361332 | 1.24757464 | 0.05382233 | 0.95778566 | 0.99999785 | -6.3851408 | up |
| KI - WT, cell_origin: iPSC | Q9UPU7 | TBC1D2B | 8 | Homo sapien | -0.084227 | -3.439229 | 3.27077489 | -0.9751838 | -0.0535009 | 0.9580375 | 0.99999785 | -6.3851587 | down |
| KI - WT, cell_origin: iPSC | P11387 | TOP1 | 23 | Homo sapien | 0.03828816 | -1.5032484 | 1.57982469 | 2.4391181 | 0.05293136 | 0.95848377 | 0.99999785 | -6.3851901 | up |
| KI - WT, cell_origin: iPSC | Q9H6S0 | YTHDC2 | 24 | Homo sapien | 0.01368449 | -0.5370384 | 0.56440734 | -0.0444698 | 0.05292188 | 0.95848639 | 0.99999785 | -6.3851913 | up |
| KI - WT, cell_origin: iPSC | Q9UGU0 | TCF20 | 5 | Homo sapien | -0.0213901 | -0.8846516 | 0.84187143 | -1.4346588 | -0.0527728 | 0.95860326 | 0.99999785 | -6.3851994 | down |
| KI - WT, cell_origin: iPSC | Q13228 | SELENBP1 | 2 | Homo sapien | -0.0893479 | -3.7074625 | 3.52876677 | -3.3147992 | -0.0526265 | 0.95872267 | 0.99999785 | -6.3852068 | down |
| KI - WT, cell_origin: iPSC | P40926 | MDH2 | 8 | Homo sapien | 0.06609704 | -2.6233847 | 2.75557882 | -1.560233 | 0.05237401 | 0.9589205 | 0.99999785 | -6.3852205 | up |
| KI - WT, cell_origin: iPSC | Q93062 | RBPMS | 7 | Homo sapien | 0.01544881 | -0.6146983 | 0.64559587 | 0.67756071 | 0.05221469 | 0.9590406 | 0.99999785 | -6.3852298 | up |
| KI - WT, cell_origin: iPSC | P62316 | SNRPD2 | 8 | Homo sapien | -0.0430203 | -1.798942 | 1.71290146 | 2.79054818 | -0.052212 | 0.95904744 | 0.99999785 | -6.3852293 | down |
| KI - WT, cell_origin: iPSC | Q16658 | FSCN1 | 11 | Homo sapien | -0.0186072 | -0.7803868 | 0.74317234 | -0.0286134 | -0.0520279 | 0.9591878 | 0.99999785 | -6.3852398 | down |
| KI - WT, cell_origin: iPSC | Q13049 | TRIM32 | 36 | Homo sapien | -0.0384688 | -1.6182888 | 1.54135121 | 5.11821831 | -0.0518924 | 0.9592979 | 0.99999785 | -6.3852466 | down |
| KI - WT, cell_origin: iPSC | Q7L5N1 | COPS6 | 3 | Homo sapien | 0.02050612 | -0.822678 | 0.86369029 | -1.997077 | 0.05179653 | 0.9593683 | 0.99999785 | -6.3852523 | up |
| KI - WT, cell_origin: iPSC | O76024 | WFS1 | 2 | Homo sapien | 0.1122687 | -4.5629333 | 4.78747067 | -4.1686906 | 0.05117533 | 0.9598598 | 0.99999785 | -6.3852849 | up |
| KI - WT, cell_origin: iPSC | A1X283 | SH3PXD2B | 15 | Homo sapien | -0.0256348 | -1.0983505 | 1.047081 | -0.2513517 | -0.0509269 | 0.96005451 | 0.99999785 | -6.385298 | down |
| KI - WT, cell_origin: iPSC | P78406 | RAE1 | 11 | Homo sapien | 0.01817511 | -0.7564362 | 0.79278646 | 0.22246692 | 0.04997958 | 0.96079336 | 0.99999785 | -6.385348 | up |
| KI - WT, cell_origin: iPSC | Q03111 | MLLT1 | 5 | Homo sapien | 0.03523999 | -1.468279 | 1.53875898 | -0.2006753 | 0.04994928 | 0.9608206 | 0.99999785 | -6.3853491 | up |
| KI - WT, cell_origin: iPSC | Q9P265 | DIP2B | 10 | Homo sapien | 0.01981283 | -0.8302557 | 0.86988133 | -0.3487751 | 0.049649 | 0.96105277 | 0.99999785 | -6.385365 | up |
| KI - WT, cell_origin: iPSC | Q99638 | RAD9A | 2 | Homo sapien | -0.0412455 | -1.8231648 | 1.74067384 | -4.3618676 | -0.0492978 | 0.96132667 | 0.99999785 | -6.3853832 | down |
| KI - WT, cell_origin: iPSC | Q9NRY4 | ARHGAP35 | 45 | Homo sapien | 0.01751386 | -0.7402784 | 0.77530613 | 1.36877667 | 0.04922972 | 0.96138101 | 0.99999785 | -6.3853866 | up |
| KI - WT, cell_origin: iPSC | Q96SU4 | OSBPL9 | 10 | Homo sapien | 0.01456409 | -0.6176266 | 0.64675478 | -0.7266407 | 0.04906536 | 0.96150888 | 0.99999785 | -6.385395 | up |
| KI - WT, cell_origin: iPSC | P42224 | STAT1 | 11 | Homo sapien | -0.0322014 | -1.4314644 | 1.36706154 | -0.7806222 | -0.0490432 | 0.96153074 | 0.99999785 | -6.3853956 | down |
| KI - WT, cell_origin: iPSC | Q5VTR2 | RNF20 | 13 | Homo sapien | -0.0336952 | -1.5003996 | 1.43300923 | -1.7096634 | -0.0489585 | 0.96159711 | 0.99999785 | -6.3853999 | down |
| KI - WT, cell_origin: iPSC | Q13188 | STK3 | 5 | Homo sapien | 0.01789717 | -0.7699963 | 0.80579066 | -1.2091283 | 0.04837895 | 0.96204692 | 0.99999785 | -6.3854297 | up |

| | | | | | | | | | | | | | |
|---|---|---|---|---|---|---|---|---|---|---|---|---|---|
| KI - WT, cell_origin: iPSC | Q8NC69 | KCTD6 | 7 | Homo sapien | -0.0191159 | -0.8653596 | 0.82712772 | -0.0533511 | -0.0481192 | 0.96225185 | 0.99999785 | -6.3854425 | down |
| KI - WT, cell_origin: iPSC | O15347 | HMGB3 | 4 | Homo sapien | -0.0556653 | -2.5217447 | 2.41041417 | 4.07449732 | -0.0481038 | 0.96226695 | 0.99999785 | -6.3854429 | down |
| KI - WT, cell_origin: iPSC | Q92804 | TAF15 | 11 | Homo sapien | 0.02639111 | -1.1432416 | 1.19602384 | 2.1599148 | 0.0480851 | 0.96228162 | 0.99999785 | -6.3854439 | up |
| KI - WT, cell_origin: iPSC | Q9NRX1 | PNO1 | 6 | Homo sapien | 0.03498913 | -1.5322517 | 1.60222992 | -1.9476973 | 0.04757731 | 0.96267961 | 0.99999785 | -6.385469 | up |
| KI - WT, cell_origin: iPSC | P51784 | USP11 | 7 | Homo sapien | 0.03106083 | -1.3632621 | 1.42538374 | -1.8486755 | 0.04747359 | 0.9627609 | 0.99999785 | -6.3854742 | up |
| KI - WT, cell_origin: iPSC | Q9UBQ5 | EIF3K | 9 | Homo sapien | -0.0110903 | -0.5106093 | 0.48842866 | 3.09162996 | -0.0472859 | 0.96290374 | 0.99999785 | -6.3854839 | down |
| KI - WT, cell_origin: iPSC | Q15021 | NCAPD2 | 14 | Homo sapien | 0.01803855 | -0.7987077 | 0.83478481 | -1.0934063 | 0.04704122 | 0.96309592 | 0.99999785 | -6.3854958 | up |
| KI - WT, cell_origin: iPSC | Q9Y4C1 | KDM3A | 13 | Homo sapien | 0.02294122 | -1.0179933 | 1.06387576 | -1.0133199 | 0.04696729 | 0.96315774 | 0.99999785 | -6.3854989 | up |
| KI - WT, cell_origin: iPSC | Q12824 | SMARCB1 | 2 | Homo sapien | -0.0273815 | -1.272275 | 1.21751197 | -0.4550545 | -0.0468735 | 0.96323123 | 0.99999785 | -6.3855035 | down |
| KI - WT, cell_origin: iPSC | Q86VP6 | CAND1 | 16 | Homo sapien | 0.01480268 | -0.6610691 | 0.69067448 | -0.4916 | 0.04664613 | 0.96340524 | 0.99999785 | -6.385515 | up |
| KI - WT, cell_origin: iPSC | P63261 | ACTG1 | 21 | Homo sapien | -0.0160411 | -0.7506865 | 0.71860425 | 5.86229006 | -0.0465328 | 0.96349831 | 0.99999785 | -6.38552 | down |
| KI - WT, cell_origin: iPSC | O00566 | MPHOSPH10 | 2 | Homo sapien | -0.0133181 | -0.6234322 | 0.59679601 | -1.2458329 | -0.0464912 | 0.96352667 | 0.99999785 | -6.3855225 | down |
| KI - WT, cell_origin: iPSC | P02788 | LTF | 3 | Homo sapien | 0.02778065 | -1.2533052 | 1.30886655 | 0.80658935 | 0.04621326 | 0.96374877 | 0.99999785 | -6.3855354 | up |
| KI - WT, cell_origin: iPSC | P18206 | VCL | 10 | Homo sapien | -0.0244066 | -1.1555829 | 1.10676964 | -1.2295886 | -0.0459811 | 0.96393073 | 0.99999785 | -6.3855465 | down |
| KI - WT, cell_origin: iPSC | P31153 | MAT2A | 13 | Homo sapien | 0.02164354 | -0.9822657 | 1.02555281 | 1.46440014 | 0.04594478 | 0.96395921 | 0.99999785 | -6.3855482 | up |
| KI - WT, cell_origin: iPSC | Q9UPW5 | AGTPBP1 | 5 | Homo sapien | -0.0190633 | -0.9066976 | 0.86857094 | -1.9679023 | -0.0457409 | 0.96411487 | 0.99999785 | -6.3855584 | down |
| KI - WT, cell_origin: iPSC | Q92922 | SMARCC1 | 22 | Homo sapien | -0.0572785 | -2.7298356 | 2.61527874 | -0.5572948 | -0.0456737 | 0.96417166 | 0.99999785 | -6.3855611 | down |
| KI - WT, cell_origin: iPSC | Q9UK99 | FBXO3 | 2 | Homo sapien | -0.0156539 | -0.7462957 | 0.71498791 | -0.703136 | -0.0456308 | 0.96420122 | 0.99999785 | -6.3855636 | down |
| KI - WT, cell_origin: iPSC | Q86Y13 | DZIP3 | 8 | Homo sapien | 0.03594566 | -1.6443272 | 1.71621857 | -2.4405366 | 0.04558994 | 0.96423735 | 0.99999785 | -6.385565 | up |
| KI - WT, cell_origin: iPSC | Q9NXA8 | SIRT5 | 10 | Homo sapien | -0.0122053 | -0.5828873 | 0.55847674 | 1.45877408 | -0.0455505 | 0.96426412 | 0.99999785 | -6.3855674 | down |
| KI - WT, cell_origin: iPSC | O00233 | PSMD9 | 8 | Homo sapien | 0.02183897 | -1.0000551 | 1.043733 | -0.4136464 | 0.04554373 | 0.96427358 | 0.99999785 | -6.3855672 | up |
| KI - WT, cell_origin: iPSC | P14324 | FDPS | 3 | Homo sapien | 0.01523535 | -0.6980559 | 0.72852664 | -0.7020947 | 0.04549097 | 0.96431081 | 0.99999785 | -6.3855702 | up |
| KI - WT, cell_origin: iPSC | Q9P287 | BCCIP | 4 | Homo sapien | -0.0274647 | -1.3141873 | 1.25925777 | -1.6157241 | -0.0454876 | 0.96431757 | 0.99999785 | -6.3855699 | down |
| KI - WT, cell_origin: iPSC | Q6DKI1 | RPL7L1 | 4 | Homo sapien | 0.01772483 | -0.8193524 | 0.8548021 | -0.8134146 | 0.04510486 | 0.96461455 | 0.99999785 | -6.3855882 | up |
| KI - WT, cell_origin: iPSC | Q49A26 | GLYR1 | 11 | Homo sapien | 0.02556243 | -1.1856787 | 1.23680357 | -0.6264802 | 0.04497528 | 0.96471917 | 0.99999785 | -6.3855939 | up |
| KI - WT, cell_origin: iPSC | O95251 | KAT7 | 10 | Homo sapien | -0.0264877 | -1.2843333 | 1.23135793 | -2.229355 | -0.0448606 | 0.96480669 | 0.99999785 | -6.3855995 | down |
| KI - WT, cell_origin: iPSC | Q9NWT1 | PAK1IP1 | 4 | Homo sapien | 0.03298233 | -1.5400232 | 1.60598789 | -2.2952403 | 0.04468415 | 0.96494739 | 0.99999785 | -6.3856074 | up |
| KI - WT, cell_origin: iPSC | Q6VN20 | RANBP10 | 5 | Homo sapien | -0.0136771 | -0.6675452 | 0.64019111 | -0.9547767 | -0.0445495 | 0.96504893 | 0.99999785 | -6.3856141 | down |
| KI - WT, cell_origin: iPSC | Q14680 | MELK | 8 | Homo sapien | 0.02403481 | -1.1263548 | 1.17442442 | -1.0064816 | 0.04452441 | 0.96507261 | 0.99999785 | -6.3856148 | up |
| KI - WT, cell_origin: iPSC | P14625 | HSP90B1 | 11 | Homo sapien | 0.02907371 | -1.3662131 | 1.42436047 | -0.7359227 | 0.04440576 | 0.96516561 | 0.99999785 | -6.3856203 | up |
| KI - WT, cell_origin: iPSC | Q9BTT4 | MED10 | 4 | Homo sapien | -0.0167812 | -0.8221248 | 0.78856244 | -1.0831575 | -0.04438 | 0.96518191 | 0.99999785 | -6.3856219 | down |
| KI - WT, cell_origin: iPSC | P49327 | FASN | 93 | Homo sapien | -0.0247307 | -1.237257 | 1.1877956 | 2.26429466 | -0.0434658 | 0.96590249 | 0.99999785 | -6.3856631 | down |
| KI - WT, cell_origin: iPSC | Q96PU5 | NEDD4L | 9 | Homo sapien | 0.01580979 | -0.7590938 | 0.79071344 | -0.5507278 | 0.04345597 | 0.96590671 | 0.99999785 | -6.3856639 | up |
| KI - WT, cell_origin: iPSC | Q01970 | PLCB3 | 4 | Homo sapien | -0.0655773 | -3.2935513 | 3.16239667 | -2.0273649 | -0.0432938 | 0.96603729 | 0.99999785 | -6.3856708 | down |
| KI - WT, cell_origin: iPSC | Q01860 | POU5F1 | 9 | Homo sapien | 0.06536991 | -3.1824114 | 3.31315121 | -0.0692774 | 0.0428937 | 0.96635097 | 0.99999785 | -6.3856887 | up |
| KI - WT, cell_origin: iPSC | Q53GS9 | USP39 | 20 | Homo sapien | -0.0364709 | -1.8574911 | 1.78454925 | 0.21530347 | -0.042681 | 0.96651775 | 0.99999785 | -6.3856981 | down |
| KI - WT, cell_origin: iPSC | Q15054 | POLD3 | 3 | Homo sapien | -0.0377695 | -1.9350097 | 1.85947067 | -1.8244745 | -0.042425 | 0.96671843 | 0.99999785 | -6.3857094 | down |
| KI - WT, cell_origin: iPSC | Q9NY27 | PPP4R2 | 5 | Homo sapien | -0.0168663 | -0.8657131 | 0.83198051 | -1.1291361 | -0.0423227 | 0.96679539 | 0.99999785 | -6.3857143 | down |
| KI - WT, cell_origin: iPSC | Q14683 | SMC1A | 32 | Homo sapien | -0.0210628 | -1.0867534 | 1.04462789 | -0.6982124 | -0.0421198 | 0.96695766 | 0.99999785 | -6.3857228 | down |
| KI - WT, cell_origin: iPSC | P35606 | COPB2 | 32 | Homo sapien | 0.01625192 | -0.8085702 | 0.84107408 | 2.0979131 | 0.04197447 | 0.96706928 | 0.99999785 | -6.3857294 | up |
| KI - WT, cell_origin: iPSC | P55795 | HNRNPH2 | 7 | Homo sapien | -0.0306752 | -1.5949707 | 1.53362031 | 1.73272946 | -0.0417898 | 0.96721638 | 0.99999785 | -6.3857372 | down |
| KI - WT, cell_origin: iPSC | O14776 | TCERG1 | 10 | Homo sapien | -0.0357843 | -1.8764648 | 1.80489629 | -1.3026562 | -0.0414301 | 0.9674984 | 0.99999785 | -6.3857527 | down |
| KI - WT, cell_origin: iPSC | Q13217 | DNAJC3 | 11 | Homo sapien | 0.0333777 | -1.694116 | 1.76087135 | -0.0020898 | 0.04117582 | 0.96769777 | 0.99999785 | -6.3857636 | up |
| KI - WT, cell_origin: iPSC | Q86VS8 | HOOK3 | 8 | Homo sapien | -0.0142253 | -0.7518985 | 0.72344799 | -0.6161564 | -0.0410711 | 0.96777618 | 0.99999785 | -6.3857684 | down |
| KI - WT, cell_origin: iPSC | Q8NBS9 | TXNDC5 | 19 | Homo sapien | 0.05999258 | -3.0535928 | 3.17357794 | 1.64114848 | 0.04106191 | 0.96778709 | 0.99999785 | -6.3857684 | up |
| KI - WT, cell_origin: iPSC | Q12789 | GTF3C1 | 18 | Homo sapien | 0.02061409 | -1.0547066 | 1.09593476 | -1.1244004 | 0.04085343 | 0.96795054 | 0.99999785 | -6.3857773 | up |

| | | | | | | | | | | | | | |
|---|---|---|---|---|---|---|---|---|---|---|---|---|---|
| KI - WT, cell_origin: iPSC | Q9UBS4 | DNAJB11 | 2 | Homo sapien | 0.03484002 | -1.8010271 | 1.8707072 | -1.2763992 | 0.04044265 | 0.96827261 | 0.99999785 | -6.3857946 | up |
| KI - WT, cell_origin: iPSC | Q684P5 | RAP1GAP2 | 17 | Homo sapien | -0.0370966 | -1.9950412 | 1.920848 | -1.7288912 | -0.0403772 | 0.96832395 | 0.99999785 | -6.3857973 | down |
| KI - WT, cell_origin: iPSC | Q9NQP4 | PFDN4 | 2 | Homo sapien | 0.01671349 | -0.8668808 | 0.90030776 | -0.2704168 | 0.0402991 | 0.96838347 | 0.99999785 | -6.3858008 | up |
| KI - WT, cell_origin: iPSC | O76021 | RSL1D1 | 17 | Homo sapien | -0.0888624 | -4.811279 | 4.63355424 | -0.5286303 | -0.0401011 | 0.96854041 | 0.99999785 | -6.3858088 | down |
| KI - WT, cell_origin: iPSC | O14646 | CHD1 | 30 | Homo sapien | -0.0271522 | -1.4709714 | 1.41666711 | -0.7791833 | -0.0400769 | 0.96855939 | 0.99999785 | -6.3858099 | down |
| KI - WT, cell_origin: iPSC | P28072 | PSMB6 | 4 | Homo sapien | 0.01887465 | -0.9850352 | 1.02278452 | 0.37526047 | 0.04006698 | 0.96856715 | 0.99999785 | -6.3858103 | up |
| KI - WT, cell_origin: iPSC | Q08380 | LGALS3BP | 3 | Homo sapien | -0.0239174 | -1.2992999 | 1.25146512 | -1.3770022 | -0.0399646 | 0.96864743 | 0.99999785 | -6.3858145 | down |
| KI - WT, cell_origin: iPSC | Q9GZS1 | POLR1E | 4 | Homo sapien | 0.02966193 | -1.5542481 | 1.61357201 | -1.1311017 | 0.03990904 | 0.96869099 | 0.99999785 | -6.3858168 | up |
| KI - WT, cell_origin: iPSC | Q9UI30 | TRMT112 | 4 | Homo sapien | -0.0128062 | -0.6984407 | 0.67282841 | 0.9067228 | -0.0397801 | 0.96878849 | 0.99999785 | -6.3858225 | down |
| KI - WT, cell_origin: iPSC | P09543 | CNP | 31 | Homo sapien | 0.01745843 | -0.9184716 | 0.95338849 | 5.7017208 | 0.03975246 | 0.96881375 | 0.99999785 | -6.3858233 | up |
| KI - WT, cell_origin: iPSC | Q9Y2D5 | AKAP2 | 11 | Homo sapien | 0.02993663 | -1.5806289 | 1.64050212 | -2.2991547 | 0.03961201 | 0.96892388 | 0.99999785 | -6.3858291 | up |
| KI - WT, cell_origin: iPSC | P27348 | YWHAQ | 9 | Homo sapien | -0.0147208 | -0.8114736 | 0.782032 | 2.98796194 | -0.0393627 | 0.96911769 | 0.99999785 | -6.3858394 | down |
| KI - WT, cell_origin: iPSC | Q86X76 | NIT1 | 3 | Homo sapien | -0.0331916 | -1.8357637 | 1.76938047 | -1.6666643 | -0.0392408 | 0.96921492 | 0.99999785 | -6.3858442 | down |
| KI - WT, cell_origin: iPSC | O75382 | TRIM3 | 7 | Homo sapien | 0.01684787 | -0.9062614 | 0.93995712 | -0.7405639 | 0.03888431 | 0.96949285 | 0.99999785 | -6.3858588 | up |
| KI - WT, cell_origin: iPSC | Q9HAN9 | NMNAT1 | 10 | Homo sapien | -0.0233576 | -1.3132889 | 1.26657377 | -0.5264302 | -0.038589 | 0.96972603 | 0.99999785 | -6.3858705 | down |
| KI - WT, cell_origin: iPSC | P24928 | POLR2A | 36 | Homo sapien | -0.0108438 | -0.6160149 | 0.59432724 | 0.31509321 | -0.0381631 | 0.97005647 | 0.99999785 | -6.3858878 | down |
| KI - WT, cell_origin: iPSC | Q96EV2 | RBM33 | 2 | Homo sapien | -0.0202665 | -1.1578117 | 1.11727874 | -0.9263617 | -0.0379675 | 0.97021332 | 0.99999785 | -6.3858952 | down |
| KI - WT, cell_origin: iPSC | Q99856 | ARID3A | 7 | Homo sapien | -0.014631 | -0.8499305 | 0.82066846 | -0.8758189 | -0.0373112 | 0.97072547 | 0.99999785 | -6.385921 | down |
| KI - WT, cell_origin: iPSC | Q8WTT2 | NOC3L | 3 | Homo sapien | -0.0306251 | -1.7844795 | 1.72322921 | -2.7374954 | -0.0372123 | 0.97080549 | 0.99999785 | -6.3859246 | down |
| KI - WT, cell_origin: iPSC | Q86US8 | SMG6 | 16 | Homo sapien | -0.015865 | -0.9273695 | 0.8956396 | -0.6750441 | -0.0370819 | 0.97090622 | 0.99999785 | -6.3859298 | down |
| KI - WT, cell_origin: iPSC | Q99615 | DNAJC7 | 11 | Homo sapien | -0.0419161 | -2.463293 | 2.37946077 | -1.1053417 | -0.036891 | 0.97105744 | 0.99999785 | -6.385937 | down |
| KI - WT, cell_origin: iPSC | P49915 | GMPS | 28 | Homo sapien | -0.0179888 | -1.0603157 | 1.02433799 | 1.28574855 | -0.0367791 | 0.97114519 | 0.99999785 | -6.3859412 | down |
| KI - WT, cell_origin: iPSC | Q13191 | CBLB | 4 | Homo sapien | 0.02233802 | -1.2774415 | 1.32211752 | -1.9570577 | 0.03662497 | 0.97126609 | 0.99999785 | -6.3859471 | up |
| KI - WT, cell_origin: iPSC | Q6R327 | RICTOR | 32 | Homo sapien | 0.01862225 | -1.0702094 | 1.10745386 | 0.1173687 | 0.03644801 | 0.97140485 | 0.99999785 | -6.3859538 | up |
| KI - WT, cell_origin: iPSC | Q96HW7 | INTS4 | 13 | Homo sapien | -0.0356328 | -2.1214265 | 2.05016098 | -1.093631 | -0.0364066 | 0.97143731 | 0.99999785 | -6.3859554 | down |
| KI - WT, cell_origin: iPSC | Q86W56 | PARG | 6 | Homo sapien | 0.01260205 | -0.7385328 | 0.76373686 | -1.2125918 | 0.03573245 | 0.97196274 | 0.99999785 | -6.3859809 | up |
| KI - WT, cell_origin: iPSC | P18615 | NELFE | 8 | Homo sapien | 0.01411318 | -0.8480479 | 0.87627422 | 0.16699934 | 0.03487491 | 0.97263696 | 0.99999785 | -6.3860121 | up |
| KI - WT, cell_origin: iPSC | Q92878 | RAD50 | 24 | Homo sapien | 0.02532652 | -1.5561685 | 1.6068215 | -0.8964256 | 0.03412794 | 0.97322428 | 0.99999785 | -6.3860387 | up |
| KI - WT, cell_origin: iPSC | A6NJG2 | SOWAHD | 7 | Homo sapien | 0.03517188 | -2.1642105 | 2.23455422 | -1.6659648 | 0.03407982 | 0.97326202 | 0.99999785 | -6.3860404 | up |
| KI - WT, cell_origin: iPSC | P62701 | RPS4X | 24 | Homo sapien | 0.00705004 | -0.4352549 | 0.44935493 | 5.59013169 | 0.03394762 | 0.97336261 | 0.99999785 | -6.3860453 | up |
| KI - WT, cell_origin: iPSC | Q9BUF5 | TUBB6 | 15 | Homo sapien | 0.01167165 | -0.722843 | 0.74618633 | 5.07621051 | 0.03386369 | 0.97343151 | 0.99999785 | -6.386048 | up |
| KI - WT, cell_origin: iPSC | Q08211 | DHX9 | 51 | Homo sapien | 0.02007392 | -1.2514573 | 1.29160518 | 3.26548324 | 0.03364399 | 0.97360382 | 0.99999785 | -6.3860557 | up |
| KI - WT, cell_origin: iPSC | Q9Y6K1 | DNMT3A | 8 | Homo sapien | 0.0372434 | -2.3567979 | 2.43128469 | -1.2717743 | 0.03315279 | 0.97398905 | 0.99999785 | -6.3860727 | up |
| KI - WT, cell_origin: iPSC | P09493 | TPM1 | 5 | Homo sapien | 0.01463723 | -0.9327663 | 0.96204072 | -0.1875782 | 0.03292263 | 0.97416919 | 0.99999785 | -6.3860806 | up |
| KI - WT, cell_origin: iPSC | Q13535 | ATR | 46 | Homo sapien | 0.01378576 | -0.9002268 | 0.9277983 | -0.0969615 | 0.03213739 | 0.97478463 | 0.99999785 | -6.3861071 | up |
| KI - WT, cell_origin: iPSC | Q9BW83 | IFT27 | 6 | Homo sapien | 0.009257 | -0.6101981 | 0.62871212 | -1.3802028 | 0.03182734 | 0.9750257 | 0.99999785 | -6.3861176 | up |
| KI - WT, cell_origin: iPSC | Q8NI22 | MCFD2 | 2 | Homo sapien | -0.0513192 | -3.5140533 | 3.41141497 | 0.38182037 | -0.0315837 | 0.97521968 | 0.99999785 | -6.3861254 | down |
| KI - WT, cell_origin: iPSC | P78356 | PIP4K2B | 3 | Homo sapien | -0.0152784 | -1.0710838 | 1.0405269 | -0.7949344 | -0.0308388 | 0.97580391 | 0.99999785 | -6.3861495 | down |
| KI - WT, cell_origin: iPSC | Q14674 | ESPL1 | 8 | Homo sapien | -0.0172542 | -1.210962 | 1.17645369 | -2.5110791 | -0.0307872 | 0.97584196 | 0.99999785 | -6.3861513 | down |
| KI - WT, cell_origin: iPSC | Q9H3G5 | CPVL | 8 | Homo sapien | 0.01511143 | -1.0328861 | 1.06310892 | -0.3183391 | 0.03072895 | 0.97589007 | 0.99999785 | -6.386153 | up |
| KI - WT, cell_origin: iPSC | Q96ST2 | IWS1 | 7 | Homo sapien | -0.0182092 | -1.2844082 | 1.24798974 | -0.3768654 | -0.0306473 | 0.97595413 | 0.99999785 | -6.3861556 | down |
| KI - WT, cell_origin: iPSC | Q6P158 | DHX57 | 33 | Homo sapien | -0.0101135 | -0.7144911 | 0.69426405 | 1.08260614 | -0.03058 | 0.97600415 | 0.99999785 | -6.3861579 | down |
| KI - WT, cell_origin: iPSC | P54136 | RARS1 | 23 | Homo sapien | 0.01002968 | -0.6921139 | 0.71217323 | 1.49506097 | 0.03042291 | 0.97612736 | 0.99999785 | -6.3861629 | up |
| KI - WT, cell_origin: iPSC | Q96EI5 | TCEAL4 | 7 | Homo sapien | -0.0137813 | -0.9845233 | 0.95696072 | -1.2659209 | -0.0302415 | 0.97627046 | 0.99999785 | -6.3861685 | down |
| KI - WT, cell_origin: iPSC | Q9H8M2 | BRD9 | 4 | Homo sapien | 0.01686183 | -1.1741682 | 1.20789185 | -1.8107945 | 0.03016623 | 0.97633078 | 0.99999785 | -6.3861708 | up |
| KI - WT, cell_origin: iPSC | O00303 | EIF3F | 12 | Homo sapien | 0.00682075 | -0.4897485 | 0.50338996 | 3.2824598 | 0.02925444 | 0.97704396 | 0.99999785 | -6.386199 | up |

| | | | | | | | | | | | | |
|---|---|---|---|---|---|---|---|---|---|---|---|---|
| KI - WT, cell_origin: iPSC | P19784 | CSNK2A2 | 14 | Homo sapien | 0.00971712 | -0.6985358 | 0.71797002 | 0.67279882 | 0.02922098 | 0.97707027 | 0.99999785 | -6.3862 up |
| KI - WT, cell_origin: iPSC | Q8WXE0 | CASKIN2 | 11 | Homo sapien | 0.02604119 | -1.8748003 | 1.92688267 | -1.3178757 | 0.02919559 | 0.97709278 | 0.99999785 | -6.3862006 up |
| KI - WT, cell_origin: iPSC | P23526 | AHCY | 7 | Homo sapien | 0.01074891 | -0.7762499 | 0.79774772 | 0.19145301 | 0.02909358 | 0.97717082 | 0.99999785 | -6.3862038 up |
| KI - WT, cell_origin: iPSC | P62913 | RPL11 | 9 | Homo sapien | 0.00648888 | -0.4686251 | 0.48160291 | 5.0676469 | 0.02908785 | 0.97717465 | 0.99999785 | -6.3862041 up |
| KI - WT, cell_origin: iPSC | P07355 | ANXA2 | 28 | Homo sapien | 0.01717122 | -1.2507464 | 1.28508887 | 5.10732991 | 0.02886107 | 0.97735517 | 0.99999785 | -6.3862107 up |
| KI - WT, cell_origin: iPSC | Q9UFW8 | CGGBP1 | 8 | Homo sapien | 0.01745867 | -1.2743621 | 1.3092794 | -0.6577824 | 0.02880125 | 0.97740209 | 0.99999785 | -6.3862125 up |
| KI - WT, cell_origin: iPSC | P62750 | RPL23A | 11 | Homo sapien | -0.0066548 | -0.5033308 | 0.49002124 | 5.47750229 | -0.0285366 | 0.97760712 | 0.99999785 | -6.3862205 down |
| KI - WT, cell_origin: iPSC | O75506 | HSBP1 | 3 | Homo sapien | -0.0188649 | -1.4301095 | 1.39237969 | 0.02086583 | -0.0284875 | 0.97764818 | 0.99999785 | -6.3862218 down |
| KI - WT, cell_origin: iPSC | Q9Y6Y0 | IVNS1ABP | 16 | Homo sapien | -0.0322946 | -2.4769213 | 2.41233203 | 0.6147774 | -0.0281527 | 0.97791079 | 0.99999785 | -6.3862316 down |
| KI - WT, cell_origin: iPSC | Q66GS9 | CEP135 | 5 | Homo sapien | 0.01281223 | -0.9645249 | 0.99014935 | -2.3690526 | 0.0279203 | 0.97809057 | 0.99999785 | -6.3862386 up |
| KI - WT, cell_origin: iPSC | O43264 | ZW10 | 8 | Homo sapien | -0.0120766 | -0.9468446 | 0.92269127 | -0.4874778 | -0.0275284 | 0.97839989 | 0.99999785 | -6.3862497 down |
| KI - WT, cell_origin: iPSC | P02765 | AHSG | 4 | Homo sapien | 0.04411752 | -3.3795581 | 3.46779311 | -2.7159914 | 0.02746128 | 0.97845316 | 0.99999785 | -6.3862516 up |
| KI - WT, cell_origin: iPSC | Q8N3C0 | ASCC3 | 54 | Homo sapien | 0.00819627 | -0.6330737 | 0.64946629 | 1.17016945 | 0.0272217 | 0.97863862 | 0.99999785 | -6.3862585 up |
| KI - WT, cell_origin: iPSC | Q9NVM6 | DNAJC17 | 6 | Homo sapien | 0.01966622 | -1.5273365 | 1.56666893 | -3.5500124 | 0.02707658 | 0.9787527 | 0.99999785 | -6.3862626 up |
| KI - WT, cell_origin: iPSC | P27816 | MAP4 | 21 | Homo sapien | 0.00736555 | -0.5777671 | 0.59249819 | 1.42376633 | 0.02680962 | 0.97896191 | 0.99999785 | -6.3862701 up |
| KI - WT, cell_origin: iPSC | Q7Z4Q2 | HEATR3 | 4 | Homo sapien | 0.01452797 | -1.1495078 | 1.17856375 | -1.574304 | 0.02659569 | 0.97913188 | 0.99999785 | -6.3862758 up |
| KI - WT, cell_origin: iPSC | Q9Y2X9 | ZNF281 | 9 | Homo sapien | 0.01292174 | -1.0266113 | 1.05245481 | -0.7730878 | 0.0264902 | 0.9792149 | 0.99999785 | -6.3862787 up |
| KI - WT, cell_origin: iPSC | P26640 | VARS1 | 8 | Homo sapien | -0.0176293 | -1.4360353 | 1.40077662 | -1.4990646 | -0.0264873 | 0.97921719 | 0.99999785 | -6.3862788 down |
| KI - WT, cell_origin: iPSC | Q9BSJ8 | ESYT1 | 22 | Homo sapien | 0.0251715 | -2.0353376 | 2.08568061 | -0.4526887 | 0.02603377 | 0.97957295 | 0.99999785 | -6.3862911 up |
| KI - WT, cell_origin: iPSC | Q8NFG4 | FLCN | 12 | Homo sapien | -0.0096332 | -0.8127654 | 0.79349907 | 0.72505235 | -0.0255506 | 0.97995036 | 0.99999785 | -6.3863042 down |
| KI - WT, cell_origin: iPSC | P39060 | COL18A1 | 7 | Homo sapien | 0.02581344 | -2.1300346 | 2.1816615 | -0.889556 | 0.02551703 | 0.9799783 | 0.99999785 | -6.386305 up |
| KI - WT, cell_origin: iPSC | P04083 | ANXA1 | 14 | Homo sapien | -0.0149055 | -1.2730178 | 1.24320672 | 0.64051527 | -0.0252482 | 0.9801892 | 0.99999785 | -6.386312 down |
| KI - WT, cell_origin: iPSC | Q15428 | SF3A2 | 8 | Homo sapien | -0.0156317 | -1.345557 | 1.31429352 | 0.09308866 | -0.0250486 | 0.9803458 | 0.99999785 | -6.3863172 down |
| KI - WT, cell_origin: iPSC | O14807 | MRAS | 8 | Homo sapien | 0.01784559 | -1.5020631 | 1.53775433 | -1.0847579 | 0.02502164 | 0.98036692 | 0.99999785 | -6.3863179 up |
| KI - WT, cell_origin: iPSC | Q8IVL1 | NAV2 | 6 | Homo sapien | -0.0205152 | -1.7691533 | 1.72812293 | -3.0290315 | -0.0250022 | 0.9803822 | 0.99999785 | -6.3863184 down |
| KI - WT, cell_origin: iPSC | Q9NSV4 | DIAPH3 | 11 | Homo sapien | -0.014268 | -1.2396333 | 1.21109726 | -1.4503236 | -0.0248142 | 0.98052966 | 0.99999785 | -6.3863233 down |
| KI - WT, cell_origin: iPSC | Q92620 | DHX38 | 7 | Homo sapien | 0.02197693 | -1.881736 | 1.92568991 | -2.3973571 | 0.02460186 | 0.98069622 | 0.99999785 | -6.3863287 up |
| KI - WT, cell_origin: iPSC | Q6P2Q9 | PRPF8 | 99 | Homo sapien | -0.0199438 | -1.7712696 | 1.731382 | 0.87786069 | -0.0242685 | 0.98095772 | 0.99999785 | -6.3863372 down |
| KI - WT, cell_origin: iPSC | P11172 | UMPS | 3 | Homo sapien | -0.0200048 | -1.7771204 | 1.73711076 | -2.9739477 | -0.0242625 | 0.98096241 | 0.99999785 | -6.3863373 down |
| KI - WT, cell_origin: iPSC | Q9Y657 | SPIN1 | 3 | Homo sapien | -0.0090072 | -0.8040872 | 0.78607284 | -0.7386808 | -0.0241303 | 0.98106434 | 0.99999785 | -6.3863408 down |
| KI - WT, cell_origin: iPSC | P45984 | MAPK9 | 4 | Homo sapien | 0.00993441 | -0.8827567 | 0.90262547 | -0.7539512 | 0.02370728 | 0.98139669 | 0.99999785 | -6.3863512 up |
| KI - WT, cell_origin: iPSC | P15531 | NME1 | 4 | Homo sapien | 0.00956364 | -0.8573053 | 0.87643262 | -0.6862828 | 0.02350108 | 0.98155828 | 0.99999785 | -6.3863563 up |
| KI - WT, cell_origin: iPSC | Q9ULJ6 | ZMIZ1 | 3 | Homo sapien | -0.0217956 | -1.9989291 | 1.9553379 | -2.7371164 | -0.0234928 | 0.98156624 | 0.99999785 | -6.3863564 down |
| KI - WT, cell_origin: iPSC | P80723 | BASP1 | 15 | Homo sapien | -0.0205191 | -1.8843856 | 1.84334743 | 2.66259919 | -0.023461 | 0.98159125 | 0.99999785 | -6.3863572 down |
| KI - WT, cell_origin: iPSC | Q9NUQ8 | ABCF3 | 5 | Homo sapien | -0.0128681 | -1.1914236 | 1.16568734 | -1.9009291 | -0.023262 | 0.98174636 | 0.99999785 | -6.386362 down |
| KI - WT, cell_origin: iPSC | Q15773 | MLF2 | 2 | Homo sapien | -0.0116938 | -1.0990173 | 1.07562963 | -1.385732 | -0.0229159 | 0.98201834 | 0.99999785 | -6.3863703 down |
| KI - WT, cell_origin: iPSC | P06493 | CDK1 | 11 | Homo sapien | 0.02898552 | -2.6738277 | 2.73179871 | -0.0758087 | 0.02285427 | 0.9820672 | 0.99999785 | -6.3863717 up |
| KI - WT, cell_origin: iPSC | Q9UMS4 | PRPF19 | 11 | Homo sapien | -0.0135658 | -1.2818496 | 1.25471812 | -0.2067626 | -0.0227945 | 0.9821141 | 0.99999785 | -6.3863731 down |
| KI - WT, cell_origin: iPSC | P22087 | FBL | 14 | Homo sapien | -0.0103743 | -0.9825608 | 0.96181217 | 0.26656547 | -0.0227412 | 0.98215594 | 0.99999785 | -6.3863744 down |
| KI - WT, cell_origin: iPSC | P53992 | SEC24C | 21 | Homo sapien | 0.00727367 | -0.6889154 | 0.70346274 | 1.59673775 | 0.02225186 | 0.98253778 | 0.99999785 | -6.3863859 up |
| KI - WT, cell_origin: iPSC | Q9NPQ8 | RIC8A | 6 | Homo sapien | -0.009309 | -0.9007895 | 0.88217162 | 0.59460698 | -0.0222499 | 0.98254085 | 0.99999785 | -6.3863859 down |
| KI - WT, cell_origin: iPSC | P14868 | DARS1 | 23 | Homo sapien | 0.00667995 | -0.639013 | 0.65237288 | 1.46370642 | 0.02203368 | 0.98270897 | 0.99999785 | -6.3863909 up |
| KI - WT, cell_origin: iPSC | Q9Y5B9 | SUPT16H | 26 | Homo sapien | -0.0133804 | -1.3150712 | 1.2883103 | 0.6267984 | -0.0219061 | 0.98281107 | 0.99999785 | -6.3863937 down |
| KI - WT, cell_origin: iPSC | Q9NR30 | DDX21 | 30 | Homo sapien | 0.02759283 | -2.6765719 | 2.73175758 | -1.3520089 | 0.0217453 | 0.98293721 | 0.99999785 | -6.3863974 up |
| KI - WT, cell_origin: iPSC | Q86W92 | PPFIBP1 | 26 | Homo sapien | -0.0126648 | -1.2621639 | 1.23683419 | -0.0658628 | -0.0216006 | 0.98305072 | 0.99999785 | -6.3864006 down |
| KI - WT, cell_origin: iPSC | Q9NPI6 | DCP1A | 3 | Homo sapien | 0.0074013 | -0.7317367 | 0.74653931 | -1.1954971 | 0.02132663 | 0.98326374 | 0.99999785 | -6.3864068 up |

| | | | | | | | | | | | | | |
|---|---|---|---|---|---|---|---|---|---|---|---|---|---|
| KI - WT, cell_origin: iPSC | Q5XPI4 | RNF123 | 3 | Homo sapien | -0.0086186 | -0.8695411 | 0.85230398 | -0.9541856 | -0.0213244 | 0.98326595 | 0.99999785 | -6.3864068 | down |
| KI - WT, cell_origin: iPSC | Q92785 | DPF2 | 6 | Homo sapien | 0.00898552 | -0.8939224 | 0.91189345 | -0.9938674 | 0.02119911 | 0.98336437 | 0.99999785 | -6.3864096 | up |
| KI - WT, cell_origin: iPSC | Q01518 | CAP1 | 7 | Homo sapien | 0.01014744 | -1.0176973 | 1.03799214 | 0.57807705 | 0.0210393 | 0.9834911 | 0.99999785 | -6.386413 | up |
| KI - WT, cell_origin: iPSC | O15063 | GARRE1 | 7 | Homo sapien | -0.0085796 | -0.8819019 | 0.8647427 | -0.3842993 | -0.0209286 | 0.98357682 | 0.99999785 | -6.3864155 | down |
| KI - WT, cell_origin: iPSC | Q5TC82 | RC3H1 | 9 | Homo sapien | 0.00969511 | -0.9953753 | 1.01476553 | -1.1520704 | 0.02055362 | 0.98387164 | 0.99999785 | -6.3864235 | up |
| KI - WT, cell_origin: iPSC | Q5T4S7 | UBR4 | 52 | Homo sapien | 0.00852684 | -0.9402154 | 0.95726909 | -0.2583538 | 0.01915182 | 0.98497173 | 0.99999785 | -6.3864524 | up |
| KI - WT, cell_origin: iPSC | Q8ND24 | RNF214 | 5 | Homo sapien | -0.0060116 | -0.6764368 | 0.66441363 | -1.0519629 | -0.0190975 | 0.98501281 | 0.99999785 | -6.3864535 | down |
| KI - WT, cell_origin: iPSC | Q8WUM0 | NUP133 | 5 | Homo sapien | 0.0095294 | -1.0599289 | 1.07898768 | -2.0080997 | 0.01898004 | 0.98510536 | 0.99999785 | -6.3864558 | up |
| KI - WT, cell_origin: iPSC | Q9BYB4 | GNB1L | 6 | Homo sapien | -0.0109903 | -1.2461454 | 1.22416478 | -0.4575233 | -0.0189623 | 0.98512063 | 0.99999785 | -6.3864561 | down |
| KI - WT, cell_origin: iPSC | O60566 | BUB1B | 23 | Homo sapien | -0.0087926 | -1.0087291 | 0.99114385 | -2.116296 | -0.0187277 | 0.98530298 | 0.99999785 | -6.3864608 | down |
| KI - WT, cell_origin: iPSC | Q92576 | PHF3 | 13 | Homo sapien | 0.01844501 | -2.1009574 | 2.13784745 | -0.461508 | 0.01854675 | 0.98544668 | 0.99999785 | -6.3864642 | up |
| KI - WT, cell_origin: iPSC | Q14980 | NUMA1 | 11 | Homo sapien | -0.014565 | -1.7196462 | 1.69051623 | -2.787196 | -0.018204 | 0.98571556 | 0.99999785 | -6.3864707 | down |
| KI - WT, cell_origin: iPSC | O60306 | AQR | 14 | Homo sapien | -0.0150821 | -1.8251572 | 1.79499297 | -0.5398282 | -0.0177569 | 0.98606636 | 0.99999785 | -6.3864791 | down |
| KI - WT, cell_origin: iPSC | Q03701 | CEBPZ | 19 | Homo sapien | -0.0131496 | -1.6221713 | 1.59587211 | -2.3976927 | -0.0174161 | 0.98633374 | 0.99999785 | -6.3864853 | down |
| KI - WT, cell_origin: iPSC | Q96CN7 | ISOC1 | 8 | Homo sapien | -0.009587 | -1.1864091 | 1.16723501 | 1.65580684 | -0.017361 | 0.98637699 | 0.99999785 | -6.3864863 | down |
| KI - WT, cell_origin: iPSC | P60660 | MYL6 | 7 | Homo sapien | -0.0049792 | -0.6189487 | 0.60899031 | 1.96900014 | -0.0172724 | 0.98644498 | 0.99999785 | -6.3864879 | down |
| KI - WT, cell_origin: iPSC | Q9HBI0 | PARVG | 9 | Homo sapien | 0.02442839 | -3.069712 | 3.11856882 | -2.4485154 | 0.01682508 | 0.98679749 | 0.99999785 | -6.3864958 | up |
| KI - WT, cell_origin: iPSC | P61758 | VBP1 | 7 | Homo sapien | -0.0085739 | -1.1181892 | 1.10104148 | -0.17042 | -0.0164667 | 0.98707869 | 0.99999785 | -6.3865019 | down |
| KI - WT, cell_origin: iPSC | O43395 | PRPF3 | 14 | Homo sapien | -0.0074197 | -0.9785099 | 0.96367046 | -0.2127143 | -0.0162829 | 0.98722291 | 0.99999785 | -6.3865051 | down |
| KI - WT, cell_origin: iPSC | Q6NZ67 | MZT2B | 5 | Homo sapien | 0.00314554 | -0.4311056 | 0.43739667 | 0.99656171 | 0.01542743 | 0.98789272 | 0.99999785 | -6.3865192 | up |
| KI - WT, cell_origin: iPSC | Q8NFF5 | FLAD1 | 7 | Homo sapien | -0.0123086 | -1.7139284 | 1.68931113 | -1.3796538 | -0.0154152 | 0.98790372 | 0.99999785 | -6.3865193 | down |
| KI - WT, cell_origin: iPSC | P09012 | SNRPA | 7 | Homo sapien | -0.0127087 | -1.772139 | 1.74672149 | 1.60806076 | -0.0153933 | 0.98792087 | 0.99999785 | -6.3865197 | down |
| KI - WT, cell_origin: iPSC | P05787 | KRT8 | 23 | Homo sapien | 0.01391356 | -1.9264137 | 1.95424078 | 0.54139411 | 0.01528149 | 0.98800863 | 0.99999785 | -6.3865215 | up |
| KI - WT, cell_origin: iPSC | Q00535 | CDK5 | 3 | Homo sapien | -0.0067109 | -0.946718 | 0.93329622 | -0.2631961 | -0.0152124 | 0.98806256 | 0.99999785 | -6.3865226 | down |
| KI - WT, cell_origin: iPSC | Q69YN2 | CWF19L1 | 4 | Homo sapien | 0.00662812 | -0.9250397 | 0.93829594 | -1.4775152 | 0.01515393 | 0.98810764 | 0.99999785 | -6.3865235 | up |
| KI - WT, cell_origin: iPSC | Q14244 | MAP7 | 8 | Homo sapien | -0.0109533 | -1.5654483 | 1.54354161 | -0.0126922 | -0.0150162 | 0.98821681 | 0.99999785 | -6.3865256 | down |
| KI - WT, cell_origin: iPSC | Q99417 | MYCBP | 6 | Homo sapien | -0.0076423 | -1.1153766 | 1.10009196 | 0.42519326 | -0.0147025 | 0.98846291 | 0.99999785 | -6.3865305 | down |
| KI - WT, cell_origin: iPSC | Q9Y4E5 | ZNF451 | 3 | Homo sapien | 0.01771286 | -2.6034624 | 2.63888812 | -3.0525268 | 0.01440107 | 0.98869944 | 0.99999785 | -6.386535 | up |
| KI - WT, cell_origin: iPSC | Q6P5Z2 | PKN3 | 15 | Homo sapien | -0.0056087 | -0.8366808 | 0.82546331 | 1.27863509 | -0.0143772 | 0.98871738 | 0.99999785 | -6.3865354 | down |
| KI - WT, cell_origin: iPSC | Q86V81 | ALYREF | 9 | Homo sapien | -0.0070984 | -1.0593486 | 1.04515181 | 0.9641052 | -0.0143762 | 0.98871895 | 0.99999785 | -6.3865354 | down |
| KI - WT, cell_origin: iPSC | Q96EP0 | RNF31 | 9 | Homo sapien | 0.00446152 | -0.6573782 | 0.66630125 | -0.9466303 | 0.01435719 | 0.98873257 | 0.99999785 | -6.3865357 | up |
| KI - WT, cell_origin: iPSC | Q9H900 | ZWILCH | 6 | Homo sapien | -0.0039525 | -0.5990829 | 0.59117794 | -1.3210513 | -0.0141448 | 0.98889923 | 0.99999785 | -6.3865388 | down |
| KI - WT, cell_origin: iPSC | Q99439 | CNN2 | 12 | Homo sapien | 0.00583687 | -0.9001199 | 0.91179367 | 1.40005626 | 0.01372912 | 0.98922654 | 0.99999785 | -6.3865448 | up |
| KI - WT, cell_origin: iPSC | P51610 | HCFC1 | 9 | Homo sapien | 0.00870978 | -1.3470999 | 1.36451946 | -1.0794942 | 0.01369023 | 0.9892572 | 0.99999785 | -6.3865454 | up |
| KI - WT, cell_origin: iPSC | P43243 | MATR3 | 35 | Homo sapien | -0.0099797 | -1.5700886 | 1.55012916 | 1.5912731 | -0.0136322 | 0.98930276 | 0.99999785 | -6.3865462 | down |
| KI - WT, cell_origin: iPSC | O95372 | LYPLA2 | 8 | Homo sapien | 0.00365964 | -0.5758658 | 0.58318508 | 2.63999424 | 0.01344947 | 0.9894449 | 0.99999785 | -6.3865488 | up |
| KI - WT, cell_origin: iPSC | Q96B36 | AKT1S1 | 3 | Homo sapien | 0.00605261 | -0.9581619 | 0.97026714 | -1.4337064 | 0.01337157 | 0.98950637 | 0.99999785 | -6.3865499 | up |
| KI - WT, cell_origin: iPSC | Q99622 | C12orf57 | 3 | Homo sapien | -0.0070125 | -1.1439981 | 1.12997306 | -1.201595 | -0.0131439 | 0.98968592 | 0.99999785 | -6.386553 | down |
| KI - WT, cell_origin: iPSC | O75691 | UTP20 | 7 | Homo sapien | -0.0067644 | -1.1131128 | 1.09958398 | -1.8760691 | -0.0130243 | 0.9897789 | 0.99999785 | -6.3865546 | down |
| KI - WT, cell_origin: iPSC | Q9NRF8 | CTPS2 | 15 | Homo sapien | -0.0064862 | -1.069035 | 1.0560626 | 0.99824783 | -0.013009 | 0.98979172 | 0.99999785 | -6.3865548 | down |
| KI - WT, cell_origin: iPSC | Q14781 | CBX2 | 9 | Homo sapien | 0.00530949 | -0.8670057 | 0.87762467 | -0.9820022 | 0.01296573 | 0.98982486 | 0.99999785 | -6.3865554 | up |
| KI - WT, cell_origin: iPSC | P41240 | CSK | 17 | Homo sapien | 0.00549742 | -0.9113708 | 0.92236566 | 1.74379533 | 0.01277682 | 0.98997378 | 0.99999785 | -6.3865579 | up |
| KI - WT, cell_origin: iPSC | Q8IX12 | CCAR1 | 19 | Homo sapien | -0.0118276 | -2.0791737 | 2.0555186 | -0.1923759 | -0.0121923 | 0.99043259 | 0.99999785 | -6.3865655 | down |
| KI - WT, cell_origin: iPSC | Q05086 | UBE3A | 19 | Homo sapien | -0.0040691 | -0.7912581 | 0.78311987 | 1.03736341 | -0.0110113 | 0.99135854 | 0.99999785 | -6.3865797 | down |
| KI - WT, cell_origin: iPSC | Q86UE4 | MTDH | 4 | Homo sapien | 0.01126435 | -2.171312 | 2.19384068 | -0.3343065 | 0.01099864 | 0.99136921 | 0.99999785 | -6.3865798 | up |
| KI - WT, cell_origin: iPSC | Q9NY93 | DDX56 | 4 | Homo sapien | 0.00398278 | -0.7702521 | 0.77821766 | -1.1046915 | 0.01095604 | 0.99140164 | 0.99999785 | -6.3865803 | up |

| | | | | | | | | | | | | | |
|---|---|---|---|---|---|---|---|---|---|---|---|---|---|
| KI - WT, cell_origin: iPSC | Q5C9Z4 | NOM1 | 5 | Homo sapien | 0.0063298 | -1.2397365 | 1.25239614 | -1.930302 | 0.01082423 | 0.99150586 | 0.99999785 | -6.3865818 | up |
| KI - WT, cell_origin: iPSC | Q9UG63 | ABCF2 | 7 | Homo sapien | 0.00599662 | -1.1850536 | 1.19704688 | -0.7637214 | 0.01072948 | 0.99158041 | 0.99999785 | -6.3865828 | up |
| KI - WT, cell_origin: iPSC | P10599 | TXN | 5 | Homo sapien | -0.0029808 | -0.602563 | 0.59660145 | 4.50060466 | -0.010589 | 0.99168978 | 0.99999785 | -6.3865844 | down |
| KI - WT, cell_origin: iPSC | Q13772 | NCOA4 | 2 | Homo sapien | 0.0065375 | -1.3226749 | 1.33574987 | -2.6574517 | 0.01047696 | 0.99177789 | 0.99999785 | -6.3865856 | up |
| KI - WT, cell_origin: iPSC | Q96JM3 | CHAMP1 | 9 | Homo sapien | -0.0060288 | -1.2452758 | 1.23321826 | -1.4789892 | -0.0103675 | 0.99186446 | 0.99999785 | -6.3865868 | down |
| KI - WT, cell_origin: iPSC | P48681 | NES | 19 | Homo sapien | 0.00382265 | -0.7852262 | 0.79287149 | -0.8247723 | 0.01031883 | 0.99190182 | 0.99999785 | -6.3865873 | up |
| KI - WT, cell_origin: iPSC | O43252 | PAPSS1 | 14 | Homo sapien | -0.004904 | -1.0471281 | 1.03732003 | 0.24129723 | -0.0100275 | 0.99213121 | 0.99999785 | -6.3865904 | down |
| KI - WT, cell_origin: iPSC | Q9BQE3 | TUBA1C | 3 | Homo sapien | -0.0029011 | -0.6675249 | 0.66172272 | 1.79601219 | -0.0092966 | 0.99270393 | 0.99999785 | -6.3865977 | down |
| KI - WT, cell_origin: iPSC | Q8WVS4 | DYNC2I1 | 4 | Homo sapien | 0.00691726 | -1.5895401 | 1.6033746 | -3.1138112 | 0.00923069 | 0.99275603 | 0.99999785 | -6.3865984 | up |
| KI - WT, cell_origin: iPSC | P33316 | DUT | 8 | Homo sapien | -0.0077482 | -1.8023391 | 1.78684266 | 0.05371764 | -0.0092011 | 0.99277973 | 0.99999785 | -6.3865986 | down |
| KI - WT, cell_origin: iPSC | Q9UBB4 | ATXN10 | 19 | Homo sapien | -0.0103815 | -2.4255689 | 2.40480587 | 1.42772763 | -0.0091604 | 0.99281168 | 0.99999785 | -6.386599 | down |
| KI - WT, cell_origin: iPSC | Q9HC35 | EML4 | 6 | Homo sapien | -0.0035542 | -0.8725513 | 0.86544288 | -0.5522475 | -0.0087125 | 0.99316257 | 0.99999785 | -6.3866032 | down |
| KI - WT, cell_origin: iPSC | P62829 | RPL23 | 6 | Homo sapien | -0.0022893 | -0.5795276 | 0.57494891 | 5.01595096 | -0.0084476 | 0.99337032 | 0.99999785 | -6.3866055 | down |
| KI - WT, cell_origin: iPSC | Q8WW12 | PCNP | 3 | Homo sapien | -0.0040298 | -1.0754895 | 1.06742983 | -1.5096413 | -0.008013 | 0.99371171 | 0.99999785 | -6.3866092 | down |
| KI - WT, cell_origin: iPSC | Q96T88 | UHRF1 | 4 | Homo sapien | -0.0043923 | -1.25074 | 1.24195537 | -2.614809 | -0.0075067 | 0.99410878 | 0.99999785 | -6.3866133 | down |
| KI - WT, cell_origin: iPSC | Q12768 | WASHC5 | 11 | Homo sapien | 0.00397648 | -1.1269759 | 1.13492886 | -0.4795491 | 0.00749303 | 0.99412004 | 0.99999785 | -6.3866134 | up |
| KI - WT, cell_origin: iPSC | P14678;P631 | SNRPB;SNRPI | 8 | Homo sapien | -0.0042257 | -1.2285147 | 1.22006323 | 2.16033148 | -0.0073557 | 0.99422784 | 0.99999785 | -6.3866145 | down |
| KI - WT, cell_origin: iPSC | O75400 | PRPF40A | 11 | Homo sapien | 0.00484722 | -1.4131775 | 1.42287195 | -0.4973508 | 0.00728469 | 0.99428353 | 0.99999785 | -6.386615 | up |
| KI - WT, cell_origin: iPSC | Q96F86 | EDC3 | 11 | Homo sapien | -0.0037559 | -1.1693946 | 1.16188282 | -1.4346167 | -0.0068667 | 0.99461152 | 0.99999785 | -6.3866181 | down |
| KI - WT, cell_origin: iPSC | Q9H7N4 | SCAF1 | 4 | Homo sapien | 0.00574057 | -1.8664993 | 1.87798046 | -1.688959 | 0.00653425 | 0.99487241 | 0.99999785 | -6.3866204 | up |
| KI - WT, cell_origin: iPSC | Q02880 | TOP2B | 43 | Homo sapien | -0.0059565 | -1.9579937 | 1.94608065 | -0.4735527 | -0.0065029 | 0.99489701 | 0.99999785 | -6.3866206 | down |
| KI - WT, cell_origin: iPSC | P60510 | PPP4C | 3 | Homo sapien | -0.0031929 | -1.0578241 | 1.05143826 | -0.9793831 | -0.0064519 | 0.99493702 | 0.99999785 | -6.3866209 | down |
| KI - WT, cell_origin: iPSC | Q96EY5 | MVB12A | 5 | Homo sapien | 0.00179952 | -0.6260568 | 0.62965581 | 1.65820467 | 0.00610429 | 0.99520925 | 0.99999785 | -6.3866232 | up |
| KI - WT, cell_origin: iPSC | Q99661 | KIF2C | 18 | Homo sapien | -0.0042012 | -1.4906382 | 1.48223583 | -0.4449584 | -0.0060232 | 0.99527344 | 0.99999785 | -6.3866237 | down |
| KI - WT, cell_origin: iPSC | Q8NI36 | WDR36 | 8 | Homo sapien | 0.00560566 | -2.0877159 | 2.09892723 | -1.1328169 | 0.00570681 | 0.99552171 | 0.99999785 | -6.3866256 | up |
| KI - WT, cell_origin: iPSC | Q9UNQ2 | DIMT1 | 14 | Homo sapien | 0.00236227 | -0.8896974 | 0.89442197 | -0.0610686 | 0.00564202 | 0.99557235 | 0.99999785 | -6.386626 | up |
| KI - WT, cell_origin: iPSC | Q9NQ55 | PPAN | 2 | Homo sapien | 0.0032194 | -1.2212284 | 1.22766724 | -3.2749769 | 0.00559982 | 0.99560516 | 0.99999785 | -6.3866263 | up |
| KI - WT, cell_origin: iPSC | Q92833 | JARID2 | 14 | Homo sapien | -0.0025635 | -0.996266 | 0.99113901 | -0.9956085 | -0.0054969 | 0.99568635 | 0.99999785 | -6.3866269 | down |
| KI - WT, cell_origin: iPSC | Q9Y6K9 | IKBKG | 8 | Homo sapien | 0.00225282 | -0.9151244 | 0.91963 | 0.08948735 | 0.00523259 | 0.99589373 | 0.99999785 | -6.3866283 | up |
| KI - WT, cell_origin: iPSC | Q96AX9 | MIB2 | 7 | Homo sapien | 0.0029257 | -1.2045156 | 1.21036702 | -1.2852597 | 0.00516376 | 0.99594785 | 0.99999785 | -6.3866287 | up |
| KI - WT, cell_origin: iPSC | Q9Y4C8 | RBM19 | 13 | Homo sapien | 0.00154831 | -0.7322759 | 0.73537248 | -0.9861783 | 0.00449372 | 0.99647324 | 0.99999785 | -6.3866321 | up |
| KI - WT, cell_origin: iPSC | Q10570 | CPSF1 | 22 | Homo sapien | 0.0025407 | -1.2603302 | 1.26541155 | -1.4775603 | 0.00428742 | 0.99663553 | 0.99999785 | -6.386633 | up |
| KI - WT, cell_origin: iPSC | P43490 | NAMPT | 21 | Homo sapien | -0.003573 | -1.8086033 | 1.80145738 | 0.12168062 | -0.0042184 | 0.99668972 | 0.99999785 | -6.3866333 | down |
| KI - WT, cell_origin: iPSC | Q14692 | BMS1 | 10 | Homo sapien | -0.0015766 | -0.8062293 | 0.8030761 | -0.9227603 | -0.0041733 | 0.99672475 | 0.99999785 | -6.3866335 | down |
| KI - WT, cell_origin: iPSC | Q9NR28 | DIABLO | 5 | Homo sapien | -0.0032486 | -1.6853385 | 1.67884125 | -1.4158153 | -0.0041158 | 0.99677021 | 0.99999785 | -6.3866337 | down |
| KI - WT, cell_origin: iPSC | Q8IY33 | MICALL2 | 7 | Homo sapien | 0.00428198 | -2.3426283 | 2.35119223 | -2.7985205 | 0.00388821 | 0.9969488 | 0.99999785 | -6.3866347 | up |
| KI - WT, cell_origin: iPSC | Q8IY18 | SMC5 | 5 | Homo sapien | -0.0017646 | -0.9753759 | 0.97184664 | -1.934013 | -0.0038602 | 0.99697043 | 0.99999785 | -6.3866348 | down |
| KI - WT, cell_origin: iPSC | Q9NUS5 | AP5S1 | 4 | Homo sapien | -0.0012925 | -0.7274256 | 0.72484053 | -0.3761554 | -0.0037911 | 0.99702469 | 0.99999785 | -6.3866351 | down |
| KI - WT, cell_origin: iPSC | Q53EZ4 | CEP55 | 7 | Homo sapien | 0.00281637 | -1.6377221 | 1.64335485 | -1.3404298 | 0.00365853 | 0.99712905 | 0.99999785 | -6.3866356 | up |
| KI - WT, cell_origin: iPSC | Q15691 | MAPRE1 | 6 | Homo sapien | -0.0018652 | -1.1864631 | 1.18273267 | -1.5224832 | -0.0033555 | 0.99736683 | 0.99999785 | -6.3866367 | down |
| KI - WT, cell_origin: iPSC | Q86U38 | NOP9 | 17 | Homo sapien | -0.001526 | -1.1277409 | 1.12468884 | 0.21466728 | -0.0028877 | 0.99773395 | 0.99999785 | -6.3866382 | down |
| KI - WT, cell_origin: iPSC | Q9Y6R0 | NUMBL | 4 | Homo sapien | 0.00132323 | -0.9962049 | 0.99885131 | 0.34974501 | 0.00282691 | 0.99778164 | 0.99999785 | -6.3866384 | up |
| KI - WT, cell_origin: iPSC | Q8NC51 | SERBP1 | 21 | Homo sapien | -0.0009399 | -0.7242667 | 0.72238689 | 4.26637116 | -0.0027686 | 0.99782731 | 0.99999785 | -6.3866385 | down |
| KI - WT, cell_origin: iPSC | P49207 | RPL34 | 6 | Homo sapien | -0.0010259 | -0.8142264 | 0.81217451 | 2.4116341 | -0.0026875 | 0.99789088 | 0.99999785 | -6.3866388 | down |
| KI - WT, cell_origin: iPSC | Q12905 | ILF2 | 17 | Homo sapien | 0.00102241 | -0.8103945 | 0.81243932 | 0.7693865 | 0.00268411 | 0.99789353 | 0.99999785 | -6.3866388 | up |
| KI - WT, cell_origin: iPSC | P14635 | CCNB1 | 9 | Homo sapien | 0.00322361 | -2.8149092 | 2.82135642 | -1.8346786 | 0.00243771 | 0.99808705 | 0.99999785 | -6.3866394 | up |

| | | | | | | | | | | | | | |
|---|---|---|---|---|---|---|---|---|---|---|---|---|---|
| KI - WT, cell_origin: iPSC | O60583 | CCNT2 | 4 | Homo sapien | -0.0011143 | -1.2068028 | 1.2045742 | -1.2804447 | -0.0019695 | 0.99845446 | 0.99999785 | -6.3866405 | down |
| KI - WT, cell_origin: iPSC | Q9P2E9 | RRBP1 | 24 | Homo sapien | -0.0011837 | -1.3361187 | 1.33375133 | -0.5635122 | -0.0018897 | 0.99851713 | 0.99999785 | -6.3866407 | down |
| KI - WT, cell_origin: iPSC | Q96P11 | NSUN5 | 11 | Homo sapien | 0.00117039 | -2.177442 | 2.17978282 | -1.4378326 | 0.00114486 | 0.99910159 | 0.99999785 | -6.3866418 | up |
| KI - WT, cell_origin: iPSC | O14965 | AURKA | 9 | Homo sapien | 0.00075128 | -1.4003734 | 1.40187592 | -0.7128163 | 0.00114269 | 0.9991033 | 0.99999785 | -6.3866418 | up |
| KI - WT, cell_origin: iPSC | O00471 | EXOC5 | 17 | Homo sapien | 0.00066898 | -2.1745576 | 2.17589559 | -0.5097933 | 0.0006554 | 0.99948569 | 0.99999785 | -6.3866423 | up |
| KI - WT, cell_origin: iPSC | Q9NVN8 | GNL3L | 10 | Homo sapien | 0.00014084 | -1.7495204 | 1.74980206 | -1.20394 | 0.00017155 | 0.99986538 | 0.99999785 | -6.3866425 | up |
| KI - WT, cell_origin: iPSC | Q9UPN6 | SCAF8 | 9 | Homo sapien | 5.0255E-05 | -0.9285794 | 0.92867994 | -0.7039572 | 0.00011531 | 0.99990951 | 0.99999785 | -6.3866425 | up |
| KI - WT, cell_origin: iPSC | P18621 | RPL17 | 11 | Homo sapien | -1.986E-05 | -0.457871 | 0.45783128 | 5.22461043 | -9.24E-05 | 0.99992748 | 0.99999785 | -6.3866425 | down |
| KI - WT, cell_origin: iPSC | Q5T1M5 | FKBP15 | 2 | Homo sapien | 2.4393E-05 | -1.5260939 | 1.52614269 | -3.3281307 | 3.4046E-05 | 0.99997328 | 0.99999785 | -6.3866425 | up |
| KI - WT, cell_origin: iPSC | Q9BUQ8 | DDX23 | 24 | Homo sapien | -1.838E-06 | -1.4326096 | 1.4326059 | 1.20932687 | -2.735E-06 | 0.99999785 | 0.99999785 | -6.3866425 | down |
